# Sparse sampling and rare-variant depletion distort PCA visualizations of population structure: recovery with objective-guided manifold learning

**DOI:** 10.64898/2026.08.11.744230

**Authors:** Jan Kočí, Olga Flegontova, Piya Changmai, Leonid A. Vyazov, Leo R. Cooper, Hossein Ashrafi, Zeynep Şencan, Pavel Flegontov

## Abstract

Principal component analysis (PCA) is routinely used to visualize population structure, yet how sparse sampling and rare-variant depletion affect low-dimensional plots remains poorly understood. Using spatial simulations, we show that these factors interact to distort visualization of genetic landscapes, producing triangular and three-ray patterns, artificial outliers and misleading clines. We develop an objective-guided manifold-learning framework that searches across genotype normalization, PCA representation and dimensionality, distance metrics, and UMAP, densMAP and PHATE parameters. High-dimensional classic PC scores consistently outperform the eigenvectors used in population genetics, but other optimal parameters and ranking objectives depend on data quality, sampling and SNP ascertainment. Across six human and animal datasets, optimized embeddings recover fine-scale structure obscured by PCA and supported by independent genetic evidence. In ancient Eurasia, optimized PHATE resolves Slavic-associated structure corroborated by haplotype-sharing communities, *qpAdm*, and Y-chromosome lineages. These results call for caution in interpreting PCA plots and establish optimized manifold learning as a hypothesis-generating approach.

## Introduction

Principal component analysis (PCA) (see Abdi & Williams 2010 and Greenacre et al. 2022 for general reviews) is widely used as a first-line method in population-genetic studies to provide an overview of genetic structure. Classic multidimensional scaling (MDS) based on genetic distance metrics such as outgroup *f_3_*-statistics is less commonly used; examples include Moreno-Mayar et al. (2018) and Maravall-López et al. (2025). Distribution of individual genomes is routinely inspected in a space formed by just first two PCs and often interpreted in terms of one-dimensional (1D) clines that emerge when initially isolated populations begin to mix (e.g., Lazaridis et al. 2025; Zeng et al. 2025). However, PCA coordinates based on population genetic data do not have unambiguous interpretations (McVean 2009). Imbalanced sampling of isolation-by-distance (IBD) landscapes or isolated populations affects PCA coordinates (Novembre & Stephens 2008; McVean 2009; House & Hahn 2017), which is a manifestation of a general problem: sensitivity of PCA to outliers in the data (Nguyen & Holmes 2019, Armstrong et al. 2022). We note that although numerous PCA variants have been proposed to improve robustness to outliers (see Abegaz et al. 2018 for a review of these “robust PCA” algorithms), in this study we focus on the PCA implementation most widely used in archaeogenetics, population and statistical genetics, namely *smartPCA* (Patterson et al. 2006; Herrando-Pérez et al. 2021).

Another fundamental problem of PCA as a visualization tool is the fact that the signal is spread over *s – 1* or *v – 1* dimensions, whichever is smaller (where *s* and *v* are counts of samples and variables, respectively), but only 2D or, rarely, 3D PC spaces are visualized and interpreted. In other words, classic and well-understood dimensionality reduction (DR) methods such as PCA and MDS are not guaranteed to generate distortion-free embeddings of manifolds (Lever et al. 2017; Moon et al. 2019; Ubbens et al. 2022; Meilă & Zhang 2024). For example, if highly drifted groups are combined with structured populations of much larger effective size (less genetically drifted), first two or three PCs are defined by the drifted groups, and the structure in the rest of the dataset is hidden in higher PCs (see an illustration in the Southeast Asian case study below). Thus, PCA does not necessarily produce an optimal low-dimensional embedding (LDE) for an arbitrary set of individuals, and the satisfactory performance of modern reference panels widely used in human archaeogenetics [e.g., West Eurasian (Haak et al. 2015) and North Eurasian (Jeong et al. 2019) panels] – onto which ancient individuals are projected to account for non-uniform sequencing technologies and missing data (Liu et al. 2017) – is the result of laborious manual optimization of dataset composition.

Spatially and temporally structured population-genetic data may contain low-dimensional components generated by 2D geography and 1D time, although population divergence, admixture, migration barriers, selection, and sampling can generate more complex geometries. This makes manifold learning (ML) an intuitively attractive approach for recovering such low-dimensional spatiotemporal structure. Manifold learning encompasses a broad class of methods, including dozens of nonlinear dimensionality-reduction approaches (Meilă & Zhang 2024), that embed samples into spaces of low, explicitly specified dimensionality. t-stochastic neighbor embedding (t-SNE; van der Maaten & Hinton 2008) and uniform manifold approximation and projection (UMAP; McInnes et al. 2018) are currently the most widely used manifold-learning methods in biology (Cashman et al. 2025). Another manifold-learning method, PHATE (“potential of heat-diffusion for affinity-based transition embedding”), has been shown to recover both global and local structure better than PCA, UMAP, and t-SNE on single-cell transcriptomic and microbiome datasets and to produce stronger correlation with geography than PCA on a set of present-day human populations (Moon et al. 2019). But it is well-known that all manifold-learning methods under their default settings demonstrate poorly reproducible results and sub-par performance on population-genetic and other data, mainly due to over-clustering and poor preservation of clines (Diaz-Papkovich et al. 2019, 2021; Moon et al. 2019; Battey et al. 2021; Chari & Pachter 2023; Meilă & Zhang 2024), while PCA is less sensitive to local structure but reproduces global structure more faithfully (Nguyen and Holmes 2019; Moon et al. 2019). This limitation has hindered the widespread adoption of manifold-learning methods in population genetics.

Since common distance metrics are unsuitable for spaces of extremely high dimensionality (Altman & Krzywinski 2018), and for reasons of computational efficiency and de-noising, manifold-learning algorithms are usually applied in practice to PC coordinates instead of raw data (Diaz-Papkovich et al. 2019; Moon et al. 2019; Heiser & Lau 2020; Battey et al. 2021; Kobak & Linderman 2021; Chari & Pachter 2023). Even in the much more extensive single-cell transcriptomics literature on DR techniques, large parameter spaces for manifold-learning algorithms – such as the number of input PCs and the number of nearest neighbors – have not been explored systematically (Moon et al. 2019; Heiser & Lau 2020; Chari & Pachter 2023). At the same time, limited parameter optimization is often performed in manifold-learning workflows (Moon et al. 2019; Battey et al. 2021; Kohli et al. 2021; Chari & Pachter 2023). We take a different approach. Rather than seeking a single optimal DR protocol suitable for most datasets (see Moon et al. 2019 for an example of such an approach), we step back and, for each dataset – that is, each set of individuals and SNPs – explore a very broad parameter space encompassing PCA and data-normalization algorithms, distance metrics in PC space, and several manifold-learning algorithms together with their parameters (**Fig. 1**).

**Figure 1.**
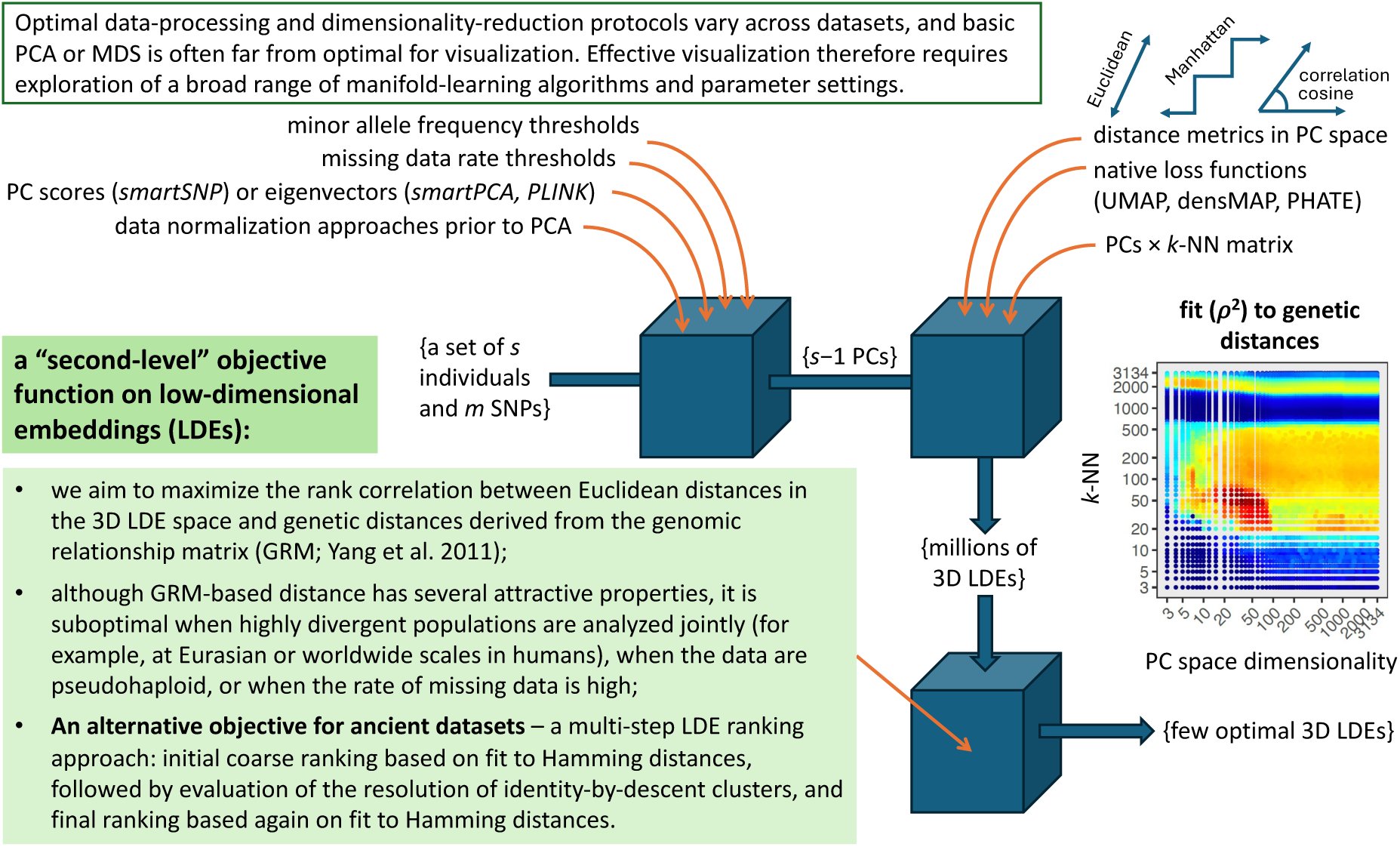
A schematic illustration of the novel manifold-learning optimization protocol for visualizing population structure.

We show that this embedding optimization approach (**Fig. 1**) is highly fruitful when guided by a second-level objective function (for related work in other fields see Xia et al. 2024; Liu et al. 2025; Gildenblat and Pahnke 2026) and examine its performance systematically: on simulated population-genetic data in the form of “stepping-stone landscapes” (SSL; **Box 1**) subsampled in various ways, and across six diverse empirical-data case studies involving both present-day and ancient genome-wide SNP data from humans and other species. Although objective-guided embedding optimization is computationally expensive, it is unsupervised and avoids laborious and subjective optimization of reference individual sets that is often necessary for visualizing population structure with traditional PCA in archaeogenetics (e.g., Gretzinger et al. 2025).

### Box 1. Glossary of terms used in this study.

| Term | Definition/explanation |
| --- | --- |
| Feature (ambient, data) space | A high-dimensional space of variables. |
| Low-dimensional embedding (LDE) | A low-dimensional representation of a feature space generated by dimensionality reduction (DR) methods. |
| Intrinsic dimensionality | The dimensionality of a manifold embedded in a higher-dimensional ambient space. For example, the intrinsic dimensionality of a sphere’s surface is 2, as is that of a Klein bottle in four dimensions and of a rolled-up sheet (Swiss roll) in three dimensions. |
| Local intrinsic dimensionality | The intrinsic dimensionality of a local neighborhood on a manifold. For example, the local intrinsic dimensionality of a tree branch is 1, but at branching points it is 2. |
| Isolation-by-distance (IBD) landscape | A two-dimensional landscape in which migration rates are equal among all points (demes or habitable locations) separated by the same distance, and decline as distance increases. |
| Isolation-by-resistance (IBR) landscape | A two-dimensional landscape with barriers to migration; that is, a landscape in which migration rates vary across space (McRae 2006). |
| Stepping-stone landscape (SSL) | A particular implementation of simulated IBD or IBR landscapes on triangular deme grids introduced by Flegontova et al. (2025). Bidirectional gene flows (“migration edges”) connect nearest neighbors only. |
| Isolation-by-resistance landscape with “flyover” long-distance migrations (IBR-LDM) | An SSL with additional migration edges connecting non-neighboring demes. These rapid long-distance migration events persist for only a limited number of generations and represent a minority of migration edges at any given time (see **Methods** for details of our implementation). |
| Genetic landscape | An IBD, IBR, IBR-LDM, or any similar landscape. |
| Admixture-graph-shaped (AGS) history | A population history composed of isolated panmictic populations that diverge and later admix through pulse-like gene-flow events. In simulations, both divergence and admixture are commonly modeled as instantaneous events involving, respectively, two descendant branches or two ancestral branches. |
| Sparsity | This term may refer either to sparse sampling of a given manifold (landscape) or to the sparsity of a feature matrix in the ambient space. In the latter case, a matrix is considered sparse if it contains many zeros representing missing data (for example, missing genotypes or gene expression values). Sparse matrices tend to be noisier, but sparsity and noise are not equivalent concepts. |
| Fit of LDE distances to genetic or geographic distances | Pearson’s or Spearman’s correlation coefficient (or its square) between all pairwise Euclidean distances in an LDE and the corresponding genetic or geographic distances. |
| $\Delta r^2, \Delta \rho^2$ | The difference in squared Pearson’s or Spearman’s correlation coefficients between a manifold-learning LDE and a reference PCA-based LDE, usually a classic drift-normalized 3D or 2D PC space. |

Numerous population-genetic studies have made substantial progress in improving genotyping and sequencing data quality and have greatly advanced our understanding of biases arising from poor sequencing quality and batch effects (Abegaz et al. 2018; Privé et al. 2020; Meisner et al. 2021; Rohland et al. 2022, Fournier et al. 2025). By contrast, the potential effects of sparse sampling of underlying genetic landscapes have been largely overlooked. For example, Meirmans (2015) noted that prioritizing genotyping quality over sampling density is a common mistake in population genetics, and that gaps in sampling can lead researchers to perceive discrete clusters and infer graph-like population histories (Maier et al. 2023) instead of IBD landscapes (Audzijonyte & Vrijenhoek 2010). To address this gap, in the present study we investigate the effects of progressively sparser sampling of simulated genetic landscapes, as well as the effects of rare-variant depletion, on 2D and 3D LDEs generated by PCA and by the objective-guided manifold-learning optimization protocol. The SSLs we simulated were either uniform (IBD) or included local barriers to gene flow (also termed isolation-by-resistance, or IBR, landscapes; McRae 2006); alternatively, the IBR landscapes were supplemented with long-distance “flyover” gene flows (see **Box 1** for definitions).

## Results

### Faithful PCA visualization of genetic landscapes requires dense sampling and rare variants

We simulated hexagon-shaped SSLs based on a triangular lattice and composed of 331 demes of constant effective size (1,000 diploid individuals). The demes arose via multifurcation and then evolved for ca. 2,500 generations (**Extended Data Fig. 1**). Samples were drawn from these landscapes at two time points: 300 generations before the end of the simulation and at the end of the simulation itself, with three individuals sampled per deme at each time point (993 individuals per time point). Both IBD and IBR landscapes (**Box 1**) were simulated on this grid, and the two time points were independently subjected to various levels and forms of subsampling and SNP filtering (see **Methods** for details):

From the 993 individuals available at each time point, either 200 (∼0.2) or 50 (∼0.05) were sampled at random. The former sampling density still provides relatively uniform coverage of the landscape. Alternatively, 200 or 50 individuals were sampled in a clustered manner to mimic the patchy, biased sampling common in population-genetic studies of humans and other species. In the 200-individual scenario, 15 core demes were selected at random, and each cluster consisted of the core deme and its nearest neighbors. In the 50-individual scenario, 5 such clusters were selected. The resulting clusters were allowed to overlap partially. Then 200 or 50 individuals, respectively, were sampled randomly from the resulting sets of demes. In addition, each subsampled dataset was optionally subjected to rare-variant removal at various minor allele frequency (MAF) thresholds (**Fig. 2**) or to linkage-disequilibrium (LD) SNP pruning.

**Figure 2.**
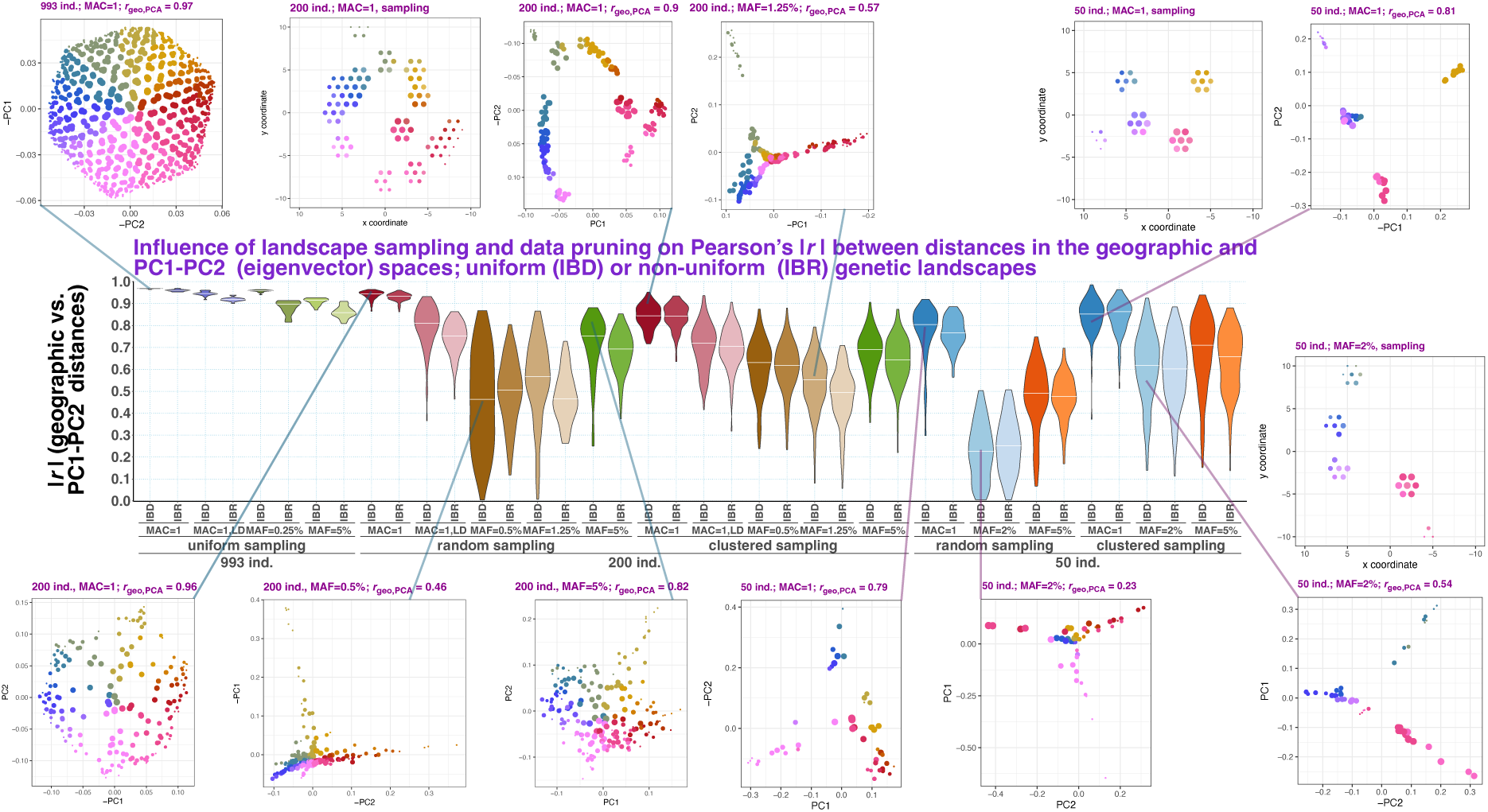
Effects of genetic landscape sampling and data pruning on Pearson’s correlation coefficient (*r*) between all pairwise distances among individuals in the simulated landscape (“geographic”) space and in the PC1–PC2 eigenvector (*U*) space. Violin plots with medians show the distributions of |*r*| across the two sampling time points (300 generations before the “present” and the “present”) × 10 simulation replicates × 5 subsampling replicates. Results are stratified by subsampling intensity, subsampling scheme (random or clustered), SNP-filtering regime (MAF-based or LD-based), and landscape type (IBD or IBR). To illustrate the patterns underlying these correlations, selected PC1–PC2 eigenvector spaces are shown for simulated IBD landscapes, with semi-transparent lines indicating where the corresponding replicates fall within the violin-plot distributions. For clustered sampling, separate panels show sample distributions across the landscape. Point size indicates proximity to the landscape center, point color distinguishes landscape sectors, and point opacity indicates the number of individuals sampled from each deme. The x- and y-axes are scaled identically. Pearson’s correlation coefficient between Euclidean distances in PC1–PC2 space and Euclidean (“geographic”) distances on the landscape is shown above each PC plot.

For 2D IBD landscapes sampled uniformly, densely, and at a single time point, 2D LDEs should ideally correspond almost perfectly to geographic distances. To test this expectation, we examined Pearson’s correlations between all pairwise Euclidean distances in the PC1–PC2 space, based on eigenvectors *U* produced by the *smartPCA*/*PLINK* algorithm (which is by far the most common in archaeogenetics and population genetics; Patterson et al. 2006, Chang et al. 2015), and geographic distances on the simulated IBD and IBR landscapes. Indeed, when all simulated demes were sampled and all polymorphic SNPs were retained [minimum allele count (MAC) = 1; ∼4,960,000–5,419,000 SNPs (**Supplementary Fig. 1**)], Pearson’s correlation coefficients exceeded 0.95 for both IBD and IBR landscapes, and the hexagonal IBD landscapes were visualized almost perfectly in PC1–PC2 space (**Fig. 2**, **Supplementary Fig. 2**). Even under dense, uniform sampling, however, LD-based SNP pruning, which retained ∼526,000–676,000 SNPs (**Supplementary Fig. 1**), led to a significant deterioration in the recovery of geographic distances (for p-values see **Extended Data Fig. 2**). Progressively stricter removal of rare variants then led to a further decline in the geography–PCA correlation (**Fig. 2**). Nevertheless, even at the MAF = 5% threshold, which retained ∼28,000–29,000 SNPs (**Supplementary Fig. 1**), the absolute correlation remained high, exceeding 0.8.

Patchy landscape sampling and depletion of rare variants had a synergistic effect: when combined, they often caused severe degradation of the geographic signal in PCA-based visualizations (**Fig. 2**). For both random and clustered subsampling schemes retaining 200 individuals (6 × 10^-4^ of the combined effective population size), and for both IBD and IBR landscapes, correlations declined in the following order: all variants retained (∼1,442,000– 1,904,000 SNPs retained); all variants retained with LD-based pruning (∼253,000–352,000 SNPs retained); MAF = 0.5% (∼482,000–617,000 SNPs retained); and MAF = 1.25% (∼56,000– 102,000 SNPs retained). However, even more aggressive removal of rare variants (MAF = 5%; (∼28,000–30,000 SNPs retained) led to a significant increase in geography–PCA correlations (**Fig. 2**, **Extended Data Fig. 2**). This effect may underlie the recommendation in the literature to apply MAF-based SNP filtering before PCA visualization and PC-based population stratification (Crosslin et al. 2014; Huckins et al. 2014; Ma & Shi 2020; Privé et al. 2020), as most early studies did not have access to the full allele-frequency spectrum and therefore observed only the transition from mild to more stringent rare-variant removal. Similar patterns were also observed in the sparsest (50-individual) sampling scenario. Interestingly, in this case, clustered sampling yielded significantly higher correlations than random sampling (**Fig. 2**, **Extended Data Fig. 2**). We also note that, because SNP loci are not independent, and because common variants are more informative about global structure whereas rare variants are more informative about local structure, the effects of SNP filtering and pruning cannot be attributed to changing SNP counts alone (compare **Fig. 2** and **Supplementary Fig. 1**). We therefore chose not to equalize SNP counts across filtering and pruning settings, in order to mimic real analytical workflows as closely as possible.

Here, we examine the characteristic patterns produced by patchy sampling and SNP filtering in PC1–PC2 plots of simple IBD landscapes, where geographic and embedding distances correspond almost perfectly under ideal conditions, that is, when all demes are sampled and no SNPs are filtered out (**Fig. 2**, **Supplementary Fig. 2**). First, we examine the effects of SNP filtering and focus on the intermediate subsampling level: 200 individuals, which still provides relatively dense and uniform landscape coverage. Under these conditions, unfiltered SNP datasets allow 2D PCA to faithfully visualize both local microstructure and the hexagonal shape of the landscape, that is, its global structure (**Fig. 2**, **Supplementary Figs. 2a–c & 3a,b**). Progressively more stringent MAF filtering at 0.5% or 1.25% usually causes the hexagonal visualization to collapse: triangular patterns emerge instead, corresponding to tetrahedral patterns in 3D, with some vertices of the hexagonal landscape forming long clines (**Fig. 2**, **Supplementary Fig. 3a,b**). Still more stringent removal of rare variants (MAF = 5%) partially restores the visualization of global structure, and therefore the correlation between LDE and geographic distances, but at the cost of compromised microstructure: neighboring samples are often mixed (**Fig. 2**, **Supplementary Fig. 3a,b**). We note that triangular and tetrahedral patterns are widespread in PCA visualizations across diverse data types and scientific fields, with examples reported in multiple contexts (Hou et al. 2015; Korem et al. 2015; Szekely et al. 2015; Flegontov et al. 2019; Jeong et al. 2019; Sikora et al. 2019; Tao et al. 2023; Allentoft et al. 2024; Salova et al. 2024; Flegontova et al. 2025). Our results indicate that, at least in some cases, such simplicial visualization patterns may signal that the data, or the associated preprocessing and sampling scheme, are suboptimal for faithful low-dimensional visualization, as discussed further below. In many other cases, however, simplicial patterns arise from convex-mixture geometry intrinsic to the data, as in compositional data (Aitchison 1983; Aitchison & Greenacre 2002) or archetypal mixtures (Cutler & Breiman 1994; Ma & Amos 2012).

Importantly, genetic data from AGS or strictly tree-shaped simulations with random topology produce simplicial patterns in PCA visualizations in nearly all cases, irrespective of sampling scheme – whether at the end of the simulation for all branches or at different points through time – and sampling density, including uniform sampling across all leaves as well as random, variable sampling across leaves (**Supplementary Fig. 4** and Flegontova et al. 2025; Vyazov et al., in preparation). Relatively dense sampling of tree or admixture-graph leaves at different points through time generates clinal patterns that closely resemble those observed in IBD landscapes subjected to patchy sampling and intermediate levels of MAF filtering (compare **Supplementary Figs. 3 & 4e,f,m,n**). Thus, in the most general case, PCA visualizations cannot distinguish between landscape-based (Bradburd & Ralph 2019; Flegontova et al. 2025) and phylogenetic models of population history (Patterson et al. 2012; Pickrell & Pritchard 2012). In Vyazov et al. (in preparation), we also showed that simple AGS histories, sampled densely along branches rather than at a single point per branch, produce PC1–PC2 visualizations that can be described as three clines meeting at the center. These visualizations obscure crucial aspects of the simulated history, which are instead recovered by force-directed layouts of IBD-sharing graphs (Vyazov et al., in preparation).

Individuals from marginal demes separated from the rest of the landscape by a sampling gap usually become distant outliers when rare variants that “tether” them to their nearest sampled neighbors are removed (**Supplementary Fig. 3c**). In some cases, very distant outliers and/or horseshoe patterns (Podani & Miklós 2002; Diaconis et al. 2008; Frichot et al. 2012) emerge in visualizations of randomly subsampled hexagonal landscapes subjected to intermediate MAF filtering (**Supplementary Fig. 3d,e**). LD pruning usually has a smaller effect on the visualization than MAF-based filtering (**Supplementary Fig. 3a,c–e**), but it also often leads to “triangularization” (**Supplementary Fig. 3b,f**). We note that variability across simulation and subsampling replicates is high (**Fig. 2**), and that a few random sampling replicates show relatively faithful visualization of global, but not local, structure (e.g., geography–PCA correlation ≳ 0.8) under all SNP-filtering and pruning regimes tested (**Supplementary Fig. 3g**).

Clustered landscape sampling (200 individuals in 15 clusters considered here) is arguably more common in practice than entirely random sampling across demes. This scenario generates highly diverse patterns in PC1–PC2 plots, depending on the shape and symmetry of the sample (**Supplementary Figs. 2d–f & 3h–p**). In general, only unfiltered SNP sets allow even remotely faithful visualization of the underlying sample distribution, with clusters remaining mostly compact and in approximately correct relative positions (**Fig. 2**, **Supplementary Figs. 2d–f & 3h–p**). By contrast, all LD-pruning and SNP-filtering levels tested here produce triangular visualization patterns in the great majority of cases, with some sample clusters remaining compact and others forming long clines (**Fig. 2**, **Supplementary Fig. 3h,i**). The most stringent MAF filtering tested here (5%) does not recover information on the global structure of clustered sample distributions, but instead produces more “puffed-up” clusters (**Supplementary Fig. 3h–j**).

Under SNP pruning and filtering, the smallest and most isolated sample clusters frequently gravitate toward the intersection of clines in PC1–PC2 space (**Supplementary Fig. 3h–j,o**). The explanation for this pattern is as follows. A sample cluster composed of few individuals and separated from the rest of the landscape by a sampling gap – but not by a gene-flow barrier – as in **Supplementary Fig. 3o**, carries specific genetic variants that are rare in the context of the whole landscape. When these variants are removed by a subsample-wide MAF threshold, signals of differential relatedness among, for example, the marginal “magenta” cluster, the more central “magenta” cluster, and the large “blue-purple” and “red-brown” clusters become distorted (**Supplementary Fig. 3o**). As a result, the marginal “magenta” cluster gravitates toward the “blue-purple” cline in both PC1–PC2 (**Supplementary Fig. 3o**) and PC1– PC2–PC3 spaces (not shown). In reality, however, the marginal “magenta” cluster is geographically and genetically equidistant from the “blue-purple” and “red-brown” clusters (**Supplementary Fig. 3o**). In other cases, such isolated clusters gravitate under SNP filtering toward the center of PC1–PC2 (**Supplementary Fig. 3i,j**) and PC1–PC2–PC3 space. They could therefore be misinterpreted as the least drifted individuals, or, depending on relative chronology, as the most admixed. The actual explanation, however, is the loss of genetic variants distinctive of these clusters; but in terms of common variation, these clusters are not strongly divergent from the rest of the landscape. In other words, the situation modeled here can be described as SNP ascertainment bias (Flegontov et al. 2023) caused by highly uneven sampling – that is, subsample-driven SNP ascertainment.

Some clustered sample configurations are clearly more favorable for preserving global structure under SNP pruning and filtering (**Supplementary Fig. 3k**). Sample distributions resembling “ring species” or horseshoes are also recovered only in the absence of SNP filtering, and become “triangularized” under other settings (**Fig. 2**, **Supplementary Fig. 3l–n**). Conversely, some sample distributions subjected to particular SNP-filtering settings yield horseshoe-like PCA patterns suggestive of approximately one-dimensional gradients or “comb-like” trees (Novembre & Stephens 2008; Estavoyer & François 2022; see also **Supplementary Fig. 5**), even though the underlying distribution is more complex (**Supplementary Fig. 3o**). Distant artificial outliers also appear at some SNP-filtering levels (**Supplementary Fig. 3p**). It is well established that PCA visualizations are sensitive to outliers in the data (Nguyen & Holmes 2019; Armstrong et al. 2022). In Vyazov et al. (in preparation), we simulated IBR landscapes of different shapes (“ribbon,” diamond, and hexagon) and “sizes”, with higher gene-flow intensities producing lower overall genetic diversity on the landscape, and sampled temporal transects from every deme. Both small overall genetic diversity, corresponding to a small subcontinental region, and sparse random sampling led to the appearance of distant artificial PC1–PC2 outliers across all landscape shapes tested (Vyazov et al., in preparation). In the present study, subsample-driven SNP ascertainment also sometimes inflates random fluctuations in the data, producing artificial distant outliers, usually at low MAF thresholds (**Supplementary Fig. 3d,e,p**).

Finally, we illustrate the effects of patchy sampling on PCA visualization in the absence of SNP filtering (**Supplementary Fig. 2**). As noted above, under optimal conditions, 2D PCA visualizes our IBD landscapes almost perfectly (**Fig. 2**), owing to their symmetric hexagonal shape (three individuals sampled per deme; ∼5,188,000–5,419,000 SNPs retained). Relatively dense random sampling (200 individuals; ∼1,821,000–1,904,000 SNPs retained) produces noticeable distortions of both global structure, including “triangularization” or “hyperbolization” of the hexagon, and local structure, manifested as exaggerated sampling gaps in the visualizations (**Supplementary Fig. 2a–c**; similar distortions caused by selective oversampling within a small IBD landscape were noted by McVean 2009). Sparse random sampling (50 individuals; ∼562,000–611,000 SNPs retained) produces more severe distortion: the full range of patterns typical of SNP filtering, including collapse of the global structure into triangular or “three-ray” configurations and the appearance of distant outliers (**Supplementary Fig. 2a–c**). As described above, clustered sampling usually introduces substantial distortions into PCA visualizations of global structure: a cluster intermediate between two others may appear equivalent to them, with all three occupying vertices of a triangle; clinal variation may be exaggerated in some clusters but collapsed in others (**Supplementary Fig. 2d–f**). Although 50-individual/5-cluster samples (∼419,000–511,000 SNPs retained) show long tails of rare poor embeddings, their median fits to geographic distances do not differ significantly from those of 200-individual/15-cluster samples at matched MAF thresholds (∼1,442,000–1,797,000 SNPs retained) (**Fig. 2**, **Extended Data Fig. 2**), corresponding to broadly similar visualization quality (**Supplementary Fig. 2d–f**). This suggests that a small number of well-sampled reference points on the landscape may often be sufficient for a relatively faithful, though still imperfect, reconstruction (**Supplementary Figs. 2 & 3**). These reference points must also be well positioned, with some type of symmetry; not all random arrangements of clusters work equally well (**Supplementary Fig. 2**), which explains the very broad distributions of correlation coefficients, especially in the 5-cluster scenario (**Fig. 2**). These observations likely underlie the success of many PCA-based visualizations in population genetics – their good agreement with other genetic analyses, as well as with later studies based on orders-of-magnitude denser sampling – despite the very sparse sampling of genetic landscapes in such work, including pioneering studies in archaeogenetics (e.g., Lazaridis et al. 2025).

**Figure 3.**
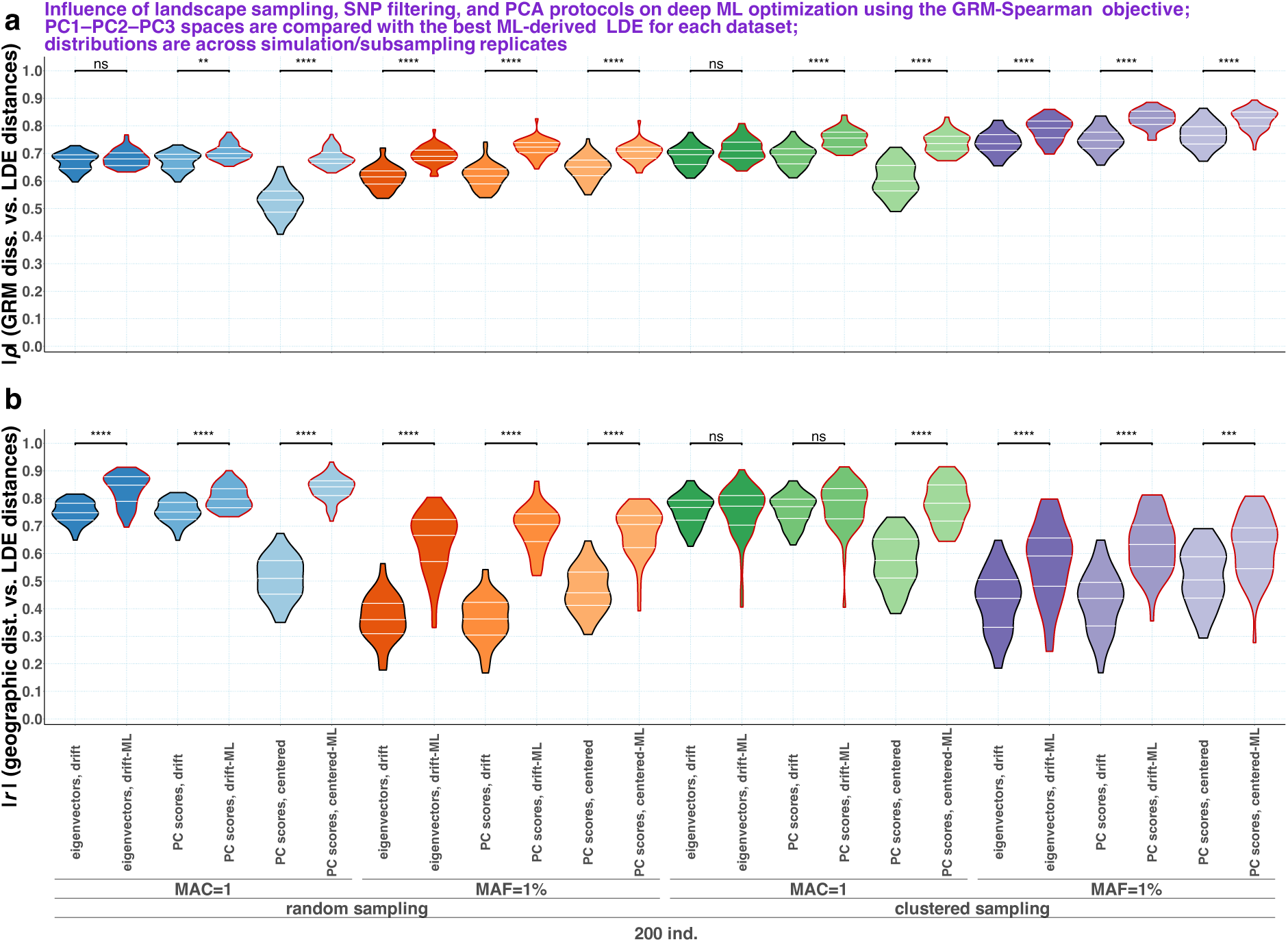
Performance of manifold-learning optimization guided by the best-performing LDE-ranking objective, GRM-Spearman. Violin plots show distributions of correlations between LDE distances and GRM-derived dissimilarities (**a**) or geographic distances (**b**) for the best manifold-learning-derived 3D LDEs, shown as violins with red borders, and for the corresponding 3D PCA LDEs, shown as violins with black borders. Distributions are computed across IBR-LDM simulation and subsampling replicates. Along the x-axes, results are stratified by landscape-sampling approach, SNP-filtering threshold, PC-space type, and LDE type, either PCA or manifold-learning-derived. For each PCA–manifold-learning pair of distributions, Wilcoxon test p-values are shown above the brackets using the following notation: ****, p < 0.0001, highly significant; ***, p < 0.001, highly significant; **, p < 0.01, moderately significant; *, p ≤ 0.05, weakly significant; ns, not significant. P-values from unpaired Wilcoxon tests were adjusted for multiple testing using the Holm method.

The “three-ray” configurations observed in PC1–PC2 spaces under sparse sampling without MAF filtering (**Supplementary Fig. 2**), or under relatively dense sampling with intermediate MAF filtering (**Fig. 2**, **Supplementary Fig. 3a,e,f,h–p**), become overwhelmingly predominant when data purging is intensified further: only 50 individuals are retained by random or clustered sampling, and singleton variants are removed (∼86,000–174,000 SNPs retained; **Fig. 2**, **Supplementary Fig. 6**). Of these, the random sampling regime produces by far the lowest median geography–PCA correlations observed in our study (**Fig. 2**). Stepping-stone IBR landscapes with geometries very different from the 2D hexagonal deme array – namely, 1D strings and rings of demes – yield the same pattern, three rays emanating from a central region in PC1–PC2 space, when subjected to aggressive random subsampling (**Supplementary Fig. 5**).

Most landscape-visualization problems discussed in this section share a common underlying theme: PCA enters a high-dimensional, low-information regime when sparse sampling eliminates the sample configurations needed to resolve local structure, when variables carrying local structure, such as rare variants, are filtered out, or both. These operations destroy the covariance patterns that distinguish nearby points on the manifold. Once the residual sample Gram matrix becomes approximately isotropic, 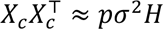 (where *p* is the number of variables, *H* = *I*_*s*_ − **11**^T^/*s*, **1** is the *s*-dimensional column vector of all ones), the observations cease to resemble samples from the original manifold. Instead, they begin to resemble *s* nearly equidistant points, that is, the vertices of a regular (*s – 1*)-simplex, and the original manifold shape becomes largely irrelevant. A line, a 1D ring (**Supplementary Fig. 5**), a branching tree (**Supplementary Fig. 4**), or a 2D triangular lattice of varying shape (**Supplementary Figs. 2**, **3**, **6**) may therefore converge, in the limit, to the same visual artifact once sufficient local information has been removed. The regular-simplex theorem for high-dimension, low-sample-size data provides a mathematical explanation for why the data approach an (*s – 1*)-simplex, where *s* is the number of samples (Hall et al. 2005; Ahn et al. 2007). What still requires a rigorous derivation is why the perturbations that break this near-degeneracy appear to select an *n*-simplex-like configuration in the first *n* PCs, producing three-ray patterns in PC1–PC2 space and four-ray patterns in PC1–PC2–PC3 space. This pattern probably reflects a combination of regular-simplex sample geometry, nearly equal leading eigenvalues (Paul 2007; Jung & Marron 2009; Jung et al. 2012; see the scree plots for 50-individual samples in **Supplementary Fig. 7**), and undersampling-driven eigenvector localization, whereby the leading PCs are driven by a few isolated samples or small sample groups rather than by the original manifold-wide structure.

It is noteworthy that triangular and even three-ray visualization patterns are common in population-genetic studies operating in an extremely sparse sampling regime, for example studies of ancient dogs and wolves, Upper Paleolithic humans, or Native Americans. Here, we do not consider PCA projections onto densely sampled present-day genetic variation, which are very common in archaeogenetics. For example, Kim et al. (2026) present PC1–PC2 eigenvector (*U*) plots based on filtered SNP data (MAF = 5%) for either 158 ancient and modern dogs, wolves, and outgroup species from around the world, or 116 ancient and modern dogs. Both plots show triangular patterns. Botigué et al. (2017) present a PC1–PC2 eigenvector plot based on SNP data derived from outgroup ascertainment, which removes much recent dog-specific rare variation, for ∼75 dogs, mostly modern; this plot shows a triangular pattern. Skoglund et al. (2015) present a PC1–PC2 eigenvector plot based on SNP-array data (∼170,000 SNPs; SNP arrays are typically depleted of rare variants) for hundreds of modern dogs; this plot shows a three-ray pattern. Der Sarkissian et al. (2015) present a PC1– PC2 eigenvector plot based on genotype likelihoods for ∼50 horses, mostly modern; this plot shows a three-ray pattern. Louis et al. (2023) present a PC1–PC2 eigenvector plot based on filtered SNP data (MAF = 5%) for 60 modern dolphins; this plot shows a triangular, possibly three-ray, pattern. Posth et al. (2023) show a typical three-cluster pattern for Upper Paleolithic humans from Eurasia in a 2D MDS based on *f*_3_-statistics of the form *f*_3_(Mbuti; X, Y).

In that analysis, both rare-variant depletion and extremely sparse sampling are present: the analysis is based on the 1240K SNP panel (Rohland et al. 2022; Flegontov et al. 2023), and the MDS plot discussed here includes only 23 individuals sampled across many millennia and the vast territory of Europe. Although that analysis uses MDS rather than PCA, erosion of local-structure information is expected to affect all dimensionality-reduction methods. Very similar methodology and results were reported in an earlier study focused on Upper Paleolithic Eurasia by Fu et al. (2016): 51 individuals were visualized by MDS based on *f*_3_(Mbuti; X, Y), revealing a triangular pattern.

Ribeiro-dos-Santos et al. (2020) present a PC1–PC2 eigenvector plot based on filtered exomic SNP data (MAF = 1%) for 58 modern Native Americans from a small region at the mouth of the Amazon; this plot shows a triangular, four-cluster pattern. Another plot in the same study, based on hundreds of modern Native Americans, mostly from South America, shows a three-ray pattern. Aguilar-Ordoñez et al. (2021) present a PC1–PC2 eigenvector plot based on filtered SNP data (MAF = 5%, LD pruning) for 80 modern Native Americans, mostly from Mexico; this plot shows a triangular pattern. Castro e Silva et al. (2022) present PC1–PC2 eigenvector plots based on LD-pruned SNP data for 87 or fewer modern Native South Americans; these plots show three-ray patterns. Arango-Isaza et al. (2023) present a PC1–PC2 eigenvector plot based on SNP-array data from the Human Origins array for 166 modern Native Americans; this plot shows a triangular pattern. Severson et al. (2022) present a PC2– PC3 plot based on SNP-array data from the Illumina Human610-Quad array for 165 modern and ancient Native Americans, mostly from North America; this plot factors out European admixture on PC1 and shows a three-ray pattern. Castro e Silva et al. (2026) present a PC1– PC2 eigenvector plot based on filtered SNP data (MAF = 5%, LD pruning) for 160 modern Native Americans, mostly from South America; this plot shows a three-ray pattern. We do not explicitly question the conclusions of these earlier studies, because strong drift, isolation, and admixture can generate genuine arms or clusters. Instead, we raise the possibility that the PCA or MDS visualizations presented in them may be strongly distorted because the underlying landscape, or manifold, signal has been eroded in the data.

In summary, the most practically relevant scenario – clustered landscape sampling combined with removal of rare variants – overwhelmingly produces PC1–PC2 plots that preserve little information about global population structure and overemphasize clinal variation in some sample clusters. Similarly, sampling graph-like population histories most often yields PCA score plots that are uninformative about important aspects of the underlying demographic history (**Supplementary Fig. 4**; Flegontova et al. 2025; Vyazov et al., in preparation). The most common problems we identified under sparse sampling, SNP pruning, and subsample-driven SNP ascertainment through MAF filtering are: (1) “triangularization” of complex spatial sample distributions; (2) sampling isolates that appear spuriously close to other clusters or occupy central positions, potentially leading to their misinterpretation as the least drifted or most admixed individuals; and (3) artificial distant outliers. Although unsurprising from the theoretical perspective, these observations call into question the long-standing archaeogenetic practice of interpreting both the global structure and microstructure of PC1– PC2 plots, most often in terms of admixture clines (Haak et al. 2015, Lazaridis et al. 2016, Olalde et al. 2018, Lamnidis et al. 2018, and Lazaridis et al. 2025 provide just a few examples).

### Effects of PCA algorithms and data normalization approaches

Patterson’s *smartPCA* algorithm (Patterson et al. 2006; Price et al. 2006), also implemented in *PLINK* v. 1.9 and v. 2 (Chang et al. 2015), differs in some respects from classic PCA. As illustrated in **Supplementary Fig. 8** and discussed previously by Abraham and Inouye (2014), the properties of eigenvectors (*U*) reported by *smartPCA* and the traditional PC scores (*UΣ*) reported by some other implementations, such as *smartSNP* (Herrando-Pérez et al. 2021), are not the same: the former are bounded between –1 and 1, whereas the latter can take a wide range of positive and negative values. The dispersion of eigenvectors and PC scores along the *n*^th^ axis is related to *n* through complex functions of similar general form, although the exact shape depends on the properties of the dataset (**Supplementary Fig. 8**). In practice, this often leads to different aspect ratios in LDEs based on eigenvectors versus PC scores (see also the empirical-data case studies below).

A snapshot of an IBD or IBR genetic landscape at a given point in time can be viewed as a surface (a 2D manifold) that is folded in complex ways in high-dimensional PC spaces. Consequently, Pearson’s correlation between “geographic” distances measured on our simulated IBR landscapes and Euclidean or Manhattan distances in corresponding *n*-dimensional PC spaces decreases approximately monotonically as *n* increases (up to a value of *n* close to *s − 1*), because the latter are not geodesic distances measured along the manifold (**Supplementary Fig. 9a,b**). We note that geographic–PCA correlations are nearly identical for eigenvectors and PC scores when the PC space dimensionality is 2 (**Extended Data Fig. 3**) or otherwise sufficiently low (**Supplementary Fig. 9a,b**). This is why, for typical visualization applications, the two approaches yield little practical difference. However, in some empirical-data case studies below PC1–PC2–PC3 eigenvector spaces showed lower correlations with genetic distances than did classic PC spaces (**Supplementary Table 1**).

When geographic distance is replaced by Hamming genetic distance between individuals on the simulated IBR landscapes, the *r*-versus-*n* curves peak at *n*>2, and under some conditions the peak correlations are significantly higher in high-dimensional classic PC spaces than in eigenvector spaces (**Supplementary Fig. 9c,d**). This is the case, for example, for aggressively MAF-filtered data (MAF = 5%) from the simulated IBR landscapes. These results suggest that eigenvector space and classic PC space may differ in ways that are relevant for manifold-learning applications, a possibility we explore in the next section.

Patterson et al. (2006) proposed a data pre-processing approach for PCA on genetic data that they showed to outperform traditional centering alone or centering followed by normalization by standard deviation (SD). This approach consists of centering followed by normalization that accounts for the expected dispersion of allele frequencies due to genetic drift, proportional to 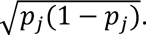 We evaluated all three data-treatment approaches and again examined the linear correlations between pairwise Euclidean distances in PC1–PC2 space and geographic distances on the simulated IBR landscapes (**Extended Data Fig. 3**). Under most conditions we tested, 2D PCA LDEs recover geographic structure most accurately when centering and drift-based normalization are applied, whereas centering followed by SD normalization yields only slightly weaker results. By contrast, centering alone performs substantially worse under dense uniform sampling across all SNP filtering and pruning regimes, as well as in subsampled datasets without rare-variant removal (**Extended Data Fig. 3**). In subsampled datasets subjected to rare-variant removal, however, the pattern changes: correlations are either significantly higher with centering alone at intermediate MAF thresholds or not significantly different among the three data-treatment approaches at MAF = 5%.

Pearson’s correlations between Hamming genetic distances in the IBR simulations and Euclidean distances in the corresponding *n*-dimensional PC spaces reveal a clear pattern: under most conditions (16 of 19), centering alone yields the highest peak correlations among the three data normalization approaches, with these peaks attained at the maximum PC-space dimensionality, *s − 1* (**Supplementary Fig. 9c,d**). At MAF = 5%, however, centering alone performs on par with the other approaches when classic PC scores are used. This emphasis on high-dimensional PC spaces is consistent with earlier landscape-genetic simulations showing that PCA-based genetic distance metrics improved as additional PC axes were included, with a 64-axis PCA metric performing best under some difficult IBR conditions (Shirk et al. 2017). Our result is theoretically expected from the geometry of PCA: unscaled, full-dimensional classic PC scores preserve the Euclidean geometry of the centered genotype matrix (Abraham & Inouye 2014; Greenacre et al. 2022), and centering alone does not alter pairwise genotype differences. These distances should therefore be most closely related to unweighted Hamming (1−IBS) genetic distances. For binary or haploid allele encodings, Hamming distance is exactly proportional to squared Euclidean distance. For diploid allele-count genotypes coded as 0/1/2, as in our simulations, 1−IBS is not identical to squared Euclidean distance because 0-versus-2 genotype contrasts are weighted differently, but the two metrics remain closely related.

By contrast, after SD normalization or Patterson’s drift normalization, full-dimensional PC distances preserve a frequency-weighted genotype distance, often strongly upweighting rarer variants relative to common variants (Patterson et al. 2006; Abraham & Inouye 2014). There is therefore no theoretical reason for these distances to correlate maximally with unweighted Hamming distances; instead, they should be expected to correlate best with a correspondingly weighted genetic distance. The geometry changes again when *smartPCA* eigenvectors are used instead of classic PC scores. In SVD notation, *X* = *U*Σ*V*^T^: classic PC scores are *U*Σ, whereas eigenvectors are essentially *U* (Abraham and Inouye 2014, Greenacre et al. 2022). Distances in *U* divide each PC axis by its singular value, producing a whitened, Mahalanobis-like geometry rather than the original Euclidean genotype geometry. Thus, even in *s − 1* dimensions, eigenvector distances are not expected to preserve raw Hamming distances in the same way as classic PC scores. We return to this issue in the next section, where we examine objective-guided embedding optimization initialized from different types of PC spaces.

### Objective-guided optimized manifold learning outperforms PCA visualization under patchy sampling and SNP filtering

As discussed in the previous section, particular high-dimensional PC representations retain substantial information about inter-sample genetic distances. We therefore asked whether this information could be recovered in LDEs by extensively optimizing nonlinear manifold-learning algorithms (**Fig. 1**). Our central premise was that no single manifold-learning algorithm or hyperparameter setting is expected to be optimal across all target genetic distances and data-generating conditions, including differences in landscape sampling and SNP filtering. Therefore, for each type of PC representation and data normalization, we explored a broad hyperparameter space spanning PC-space dimensionalities, *k*-NN values, input-space distance metrics, and four manifold-learning algorithms: (i) standard UMAP, which constructs a weighted *k*-NN graph and optimizes an LDE to preserve input-space sample neighborhoods, more precisely the corresponding fuzzy topological structure (McInnes et al. 2018); (ii) densMAP, a density-augmented UMAP variant that adds an objective encouraging local densities in the embedding to match those in the input space (Narayan et al. 2021; https://jlmelville.github.io/uwot/); (iii) a density-preserving objective implemented in the *uwot* package, controlled by an intermediate *dens_scale* value of 0.5 and therefore occupying a position between standard UMAP and full densMAP-like density preservation (https://jlmelville.github.io/uwot/); and (iv) PHATE, which builds a diffusion geometry, computes information-geometric distances between diffused affinity profiles, and embeds these distances into a low-dimensional space, with the aim of preserving both local and global nonlinear structure, including continuous progressions and branching patterns (Moon et al. 2019).

Thus, the main components of this manifold-learning optimization framework are: (i) the input representation, namely an *n*-dimensional PC space based on eigenvectors or classic PC scores, with centered and drift-normalized, or centered-only genotype normalization; (ii) the hyperparameter space explored for each manifold-learning algorithm; and (iii) a second-level objective used to rank candidate LDEs and select the best embedding. As second-level objectives, we tested squared Pearson’s and Spearman’s correlation coefficients between all pairwise Euclidean distances in LDE space and five genetic distance matrices: Hamming distance; a Genomic Relationship Matrix (GRM)-derived dissimilarity based on allele-frequency-standardized genotype covariance (Yang et al. 2011; Chang et al. 2015); an unstandardized covariance-derived dissimilarity (Chang et al. 2015); *F_ST_* between individuals; and the closely related *f*_2_-statistic (Patterson et al. 2012) between individuals. GRM-derived and unstandardized covariance-derived dissimilarities are highly correlated (*r*∼0.9), whereas individual-level *F_ST_* and *f*_2_-statistics are nearly identical (**Supplementary Fig. 10**). We therefore focus primarily on three less redundant metrics: Hamming distance, GRM dissimilarity, and *F_ST_*. Because our goal was to improve on PCA-based LDEs, we considered not only raw *r*^2^ and *ρ*^2^, but also Δ*r*^2^ and Δ*ρ*^2^, defined as the differences in squared correlation coefficients between each manifold-learning LDE and a reference PCA-based LDE, namely the classic drift-normalized PC1−PC2−PC3 space.

To test this optimization framework under relatively realistic conditions, we applied it not to simple simulated IBD or IBR landscapes, but to IBR-LDM landscapes: IBR landscapes with “flyover” long-distance gene-flow events placed randomly in space and time (**Box 1** and **Methods**). Because contemporaneous sampling from these landscapes is expected to generate “genetic manifolds” with local intrinsic dimensionality exceeding two in at least some regions, we used 3D rather than 2D LDEs. For simplicity and computational tractability, the parameter space explored above (**Fig. 2**, **Extended Data Fig. 3**) was reduced: only one subsampling density was considered, with 200 individuals sampled from a combined effective population size of 331,000 at each time point; only one epoch, the end of the simulation, was sampled; centering followed by SD normalization was omitted because its behavior was very similar to drift normalization; and only unfiltered SNP data and an intermediate-stringency MAF filter of 1% were tested (**Supplementary Fig. 1**), because the latter condition was previously shown to be the most challenging for genetic landscape visualization (**Fig. 2**, **Supplementary Figs. 3 & 4**).

**Figure 4.**
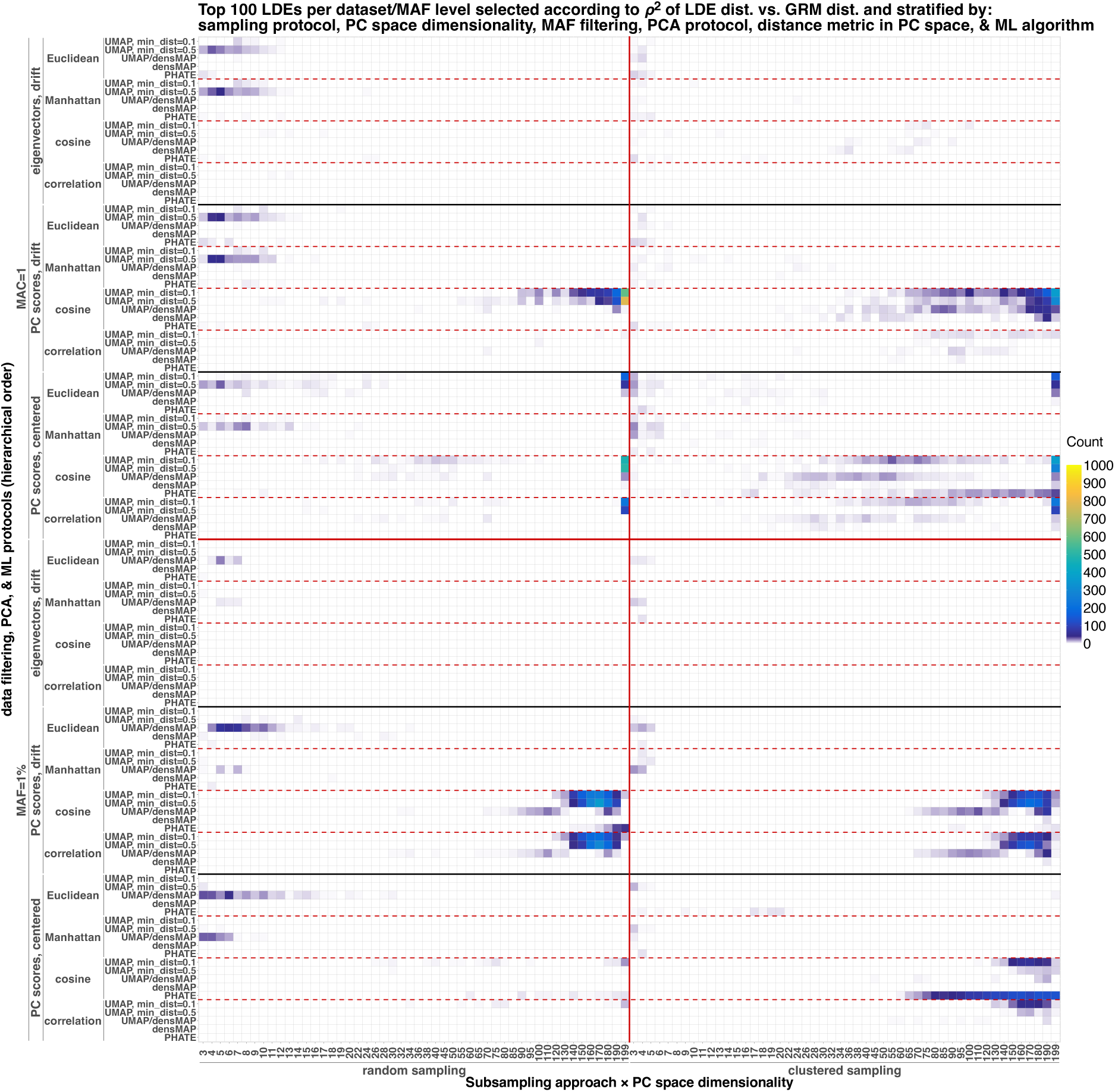
Distribution of top-ranked LDEs in the parameter space of the objective-guided embedding-optimization protocol. Optima in the four regions of parameter space (rectangles delineated by the solid red lines) should be interpreted separately, because each region is derived from a distinct set of subsampling replicates and SNPs. One hundred top-ranked LDEs were generated from each unique IBR-LDM simulated input dataset and selected using the GRM-Spearman objective. An input dataset is defined as a combination of simulation/subsampling replicate and MAF-filtering level, either no MAF filtering or a 1% MAF threshold; all other settings are treated as parameters of the manifold-learning optimization protocol. For clarity, one dimension of the parameter space – the *k*-NN parameter of the manifold-learning algorithms – is collapsed. The density of LDEs at each point in the remaining parameter space is shown on a linear color scale.

Below, we first examine which PC representation, data normalization, input-space distance metric, and PC dimensionality achieve the highest correlations with the selected genetic distances. We then ask whether these high-dimensional-space correlations predict the best fits to genetic distances in manifold-learning-derived LDEs. Finally, we evaluate which of the ten second-level optimization objectives tested – five genetic distances, each combined with either linear or rank correlation – achieve strong fits not only to genetic distance, but also to geographic distance, and whether these fits improve on those obtained from classic drift-normalized PC1–PC2–PC3 spaces.

#### The relationship between genetic distances and PC-space geometry

We return to the *r*-versus-*n* curves for subsampled IBR-LDM landscapes and now explore a wider parameter space: Euclidean, Manhattan, and cosine distances in PC spaces of varying dimensionality, and both linear and rank correlations (**Supplementary Fig. 11**). Nearly perfect correlations with *F_ST_* are achieved by Euclidean distances in centered, unscaled, full-dimensional classic PC space, and this pattern holds across MAF-filtering and landscape-sampling approaches. For data without MAF filtering, both Pearson’s *r* and Spearman’s *ρ* reach median values of approximately 0.95 across simulation and subsampling replicates. These correlations are significantly lower for MAF-filtered data, typically *r*∼0.8–0.9 (**Supplementary Fig. 11a**). Because the 1% MAF threshold was applied separately to each subsampled dataset, the retained SNP set depends on the sampling scheme itself. This can remove variants that are rare in a particular subsample, including variants informative about under-sampled regions or locally differentiated parts of the landscape, thereby altering pairwise genetic distances. Consistent with this interpretation, median *r* is ∼0.8 for landscapes subjected to the most uneven, clustered sampling, but ∼0.9 for landscapes subjected to more uniform, random sampling (see examples of clustered and random sample distributions in **Supplementary Fig. 3**). We note that Euclidean distances in centered, drift-scaled PC or eigenvector spaces represent *F_ST_* substantially less well, both at low and high dimensionalities, especially when rare variants are retained. Manhattan distances show weaker correlations, whereas cosine distances are largely unsuitable for representing *F_ST_* in this setting (**Supplementary Fig. 11a**).

These results are expected because, as discussed above, full-dimensional classic PC scores preserve the Euclidean geometry of the centered genotype matrix (Abraham & Inouye 2014; Greenacre et al. 2022). For allele-frequency vectors, *f*_2_ is geometrically a squared Euclidean distance in allele-frequency space, and Peter (2022) explicitly connects *f*-statistics with PCA geometry. So squared Euclidean distances in full-dimensional, centered, unscaled PC space should be nearly identical to individual-level *f*_2_, apart from constants, missing-data handling, bias correction, and LD effects. Because individual-level *F_ST_* is closely related to *f*_2_ but additionally normalized by heterozygosity/diversity terms (Patterson et al. 2012), it is expected to correlate very strongly with full-dimensional unscaled PC distances when these normalization terms vary little across sample pairs.

Relationships between PC-space distances and Hamming distances broadly resemble those observed for *F_ST_* (**Supplementary Fig. 11b**). However, for MAF-filtered data, centering followed by drift normalization performs nearly as well as, or slightly better than, centering alone. Conversely, for data without MAF filtering, cosine distance in centered classic PC space performs nearly as well as, or slightly better than, Euclidean distance. In all cases, the highest correlations are reached in full-dimensional PC spaces (**Supplementary Fig. 11b**). This pattern is expected because Hamming distance is related to, but not identical to, Euclidean genotype distance. For diploid genotypes, a heterozygote–homozygote difference contributes 1 to both Hamming distance and squared Euclidean distance, whereas an opposite-homozygote difference contributes 2 to Hamming distance but 4 to squared Euclidean distance. Thus, Euclidean distances in full-dimensional, unscaled PC space should correlate strongly with Hamming distance.

The cases in which Euclidean distances in drift-normalized PC spaces perform best likely reflect the effect of locus weighting. Unlike centered, unscaled PCs, drift-normalized PCs do not preserve raw allele-count Euclidean geometry, but instead give relatively greater weight to lower-frequency variants. If, in a given sampling or MAF-filtering setting, Hamming-distance structure is driven disproportionately by rare or locally informative variants, this reweighting can improve the correlation even though the resulting distance is no longer algebraically equivalent to Hamming distance. Similarly, cosine distance in centered classic PC space can outperform Euclidean distance in some settings because it compares the angle between centered genotype-deviation vectors rather than their absolute separation. In practice, this downweights variation in vector length, which may reflect sample-specific heterozygosity, inbreeding, or admixture intensity. If such magnitude differences are less relevant to Hamming distance than the overall direction of allele-sharing deviations, cosine distance can yield tighter correlations.

Finally, we considered the GRM-derived dissimilarity obtained from *PLINK*’s *--make-rel* output. The resulting matrix is a variance-standardized relationship matrix, in which SNP-level covariance is divided by SNP variance. For MAF-filtered data, median Pearson’s *r,* multiplied by –1, exceeds 0.95 for cosine distances in full-dimensional, drift-normalized classic PC spaces, but is substantially lower for other PC-space and distance-metric combinations (**Supplementary Fig. 11c**). For data without MAF filtering, *r* reaches approximately 0.8–0.9 for cosine distances in full-dimensional, centered classic PC spaces, but remains lower in other settings (**Supplementary Fig. 11c**). Because the GRM is a relationship, or similarity, matrix rather than a distance matrix, correlations between PC-space distances and GRM*_ij_* were multiplied by –1; this is equivalent to correlating PC-space distances with 1 – GRM*_ij_*. This quantity should be interpreted as a GRM-derived dissimilarity rather than as the Euclidean distance induced by the GRM kernel.

The observed results are broadly consistent with the geometry of the GRM: *PLINK*’s default GRM is based on allele-frequency-standardized genotype covariance, so full-dimensional drift-normalized classic PC spaces are expected to be closely related to this matrix. The better performance of cosine distance is also expected because 1 – GRM*_ij_* is an inner-product-based dissimilarity, and cosine distance is likewise based on normalized inner products, whereas the closest Euclidean analogue is squared Euclidean distance; here, however, correlations were calculated with unsquared Euclidean distance. Moreover, Euclidean distance additionally depends on differences in vector length, which can arise from differences in heterozygosity, inbreeding, admixture, or rare-variant burden, especially because allele-frequency standardization gives large weights to very rare variants. In unfiltered data, the latter effect makes drift-normalized geometry sensitive to rare-variant burden, local singleton/doubleton structure, and variation in vector norms. In this setting, cosine distances in full-dimensional, centered classic PC space may better capture the direction of shared genotype-deviation profiles while downweighting variation in magnitude, explaining why they can outperform drift-normalized distances when rare variants are retained.

#### High-dimensional-space correlations predict genetic-distance preservation in manifold-learning-derived LDEs

Subsets of 50 *n*-dimensional PC representations derived from the IBR-LDM simulations, with *n* ranging from 3 to 199, were used as input for the manifold-learning optimization pipeline described above (**Fig. 1**). For each representation, correlations with geographic distance and selected genetic distances – Hamming, GRM-derived dissimilarity, and *F_ST_* – are known (**Supplementary Fig. 11**). For each unique input, the best manifold-learning-derived LDE was selected according to either Pearson’s or Spearman’s correlation with each of these four metrics. We then matched the correlations between the original *n*-dimensional input and each distance metric to the corresponding correlations between the optimized 3D LDE and the same metric (**Extended Data Fig. 4**).

When random and clustered sampling schemes are analyzed together, and results for unfiltered SNP sets are pooled with those for MAF-filtered sets (**Extended Data Fig. 4a**), correlation with *F_ST_* in the input PC space is a strong predictor of manifold-learning performance: deviance explained by nonlinear generalized additive models (GAMs) reaches ∼71–77%. The same is true for Pearson’s correlation with GRM-derived dissimilarity, for which the GAM explains 73% of the deviance; however, the relationship plateaus at a PC-space/GRM-derived distance correlation of ∼0.5 (**Extended Data Fig. 4a**). By contrast, correlation with Hamming distance is a weaker predictor, with GAM deviance explained of 58–62%.

When the results are stratified by sampling scheme and MAF-filtering threshold, it becomes clear that the poor performance of Hamming distance as a predictor is driven by a subset of MAF-filtered input spaces (**Extended Data Fig. 4b**). Overall, the strongest predictors are Pearson’s correlation with GRM-derived dissimilarity (deviance explained = 68–86%; the maximum correlation achieved after manifold learning remains modest, at approximately 0.6), Spearman’s correlation with GRM-derived dissimilarity (deviance explained = 37–75%), Spearman’s correlation with *F_ST_* (deviance explained = 69–83%), and Pearson’s correlation with Hamming distance for unfiltered SNP sets (deviance explained = 70–72%). In the next section, we examine whether any of these second-level objectives achieve a good fit to both genetic and geographic distances – that is, whether they reconstruct the “genetic landscape” – and whether they outperform PCA in this respect.

#### Genetic landscape reconstruction with objective-guided optimized manifold learning

Applying any of the ten second-level objectives used to rank outputs of manifold-learning optimization significantly improves the fit to genetic distances relative to the corresponding PC1–PC2–PC3 space across nearly all landscape-sampling and SNP-filtering approaches tested (**Table 1**). In some cases, the improvement in correlation coefficient is substantial – for example, from ∼0.35 to ∼0.75 under random sampling with MAC = 1, centered classic PC spaces, and the *F_ST_*-Spearman objective (**Supplementary Fig. 12**). In other cases, such as the GRM-Spearman objective, the improvement is modest, but it is still statistically significant in most cases (**Fig. 3a**, **Table 1**, **Supplementary Fig. 12**). This result is unsurprising, given that each optimization run explores a very wide parameter space (**Fig. 1**, **Supplementary Dataset 1**).

**Table 1.**
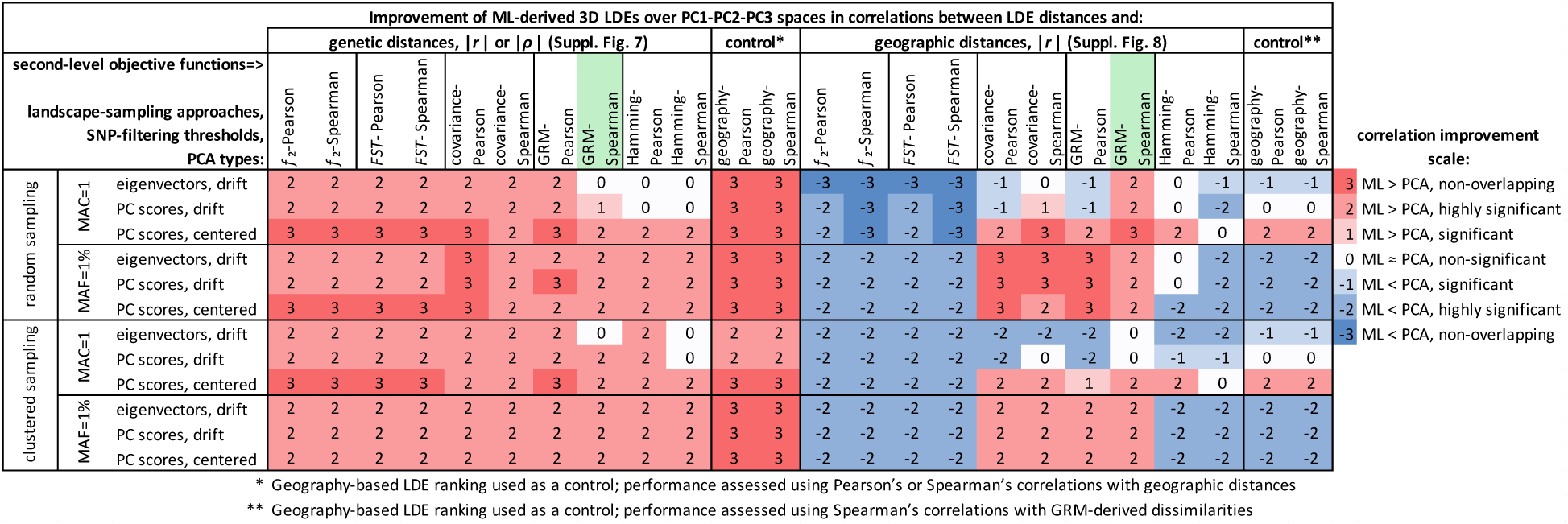
Comparing the performance of second-level objectives used to select the best manifold-learning-derived LDE: summary of distribution comparisons shown in **Supplementary Figs. 12 & 13**. Differences between the fits of PC1–PC2–PC3 spaces and the best manifold-learning-derived 3D LDEs to genetic or geographic distances are graded on a scale from 3, indicating that manifold learning improves over PCA visualizations and that the correlation coefficient distributions do not overlap, to −3, indicating that manifold-learning-derived LDEs perform worse than PCA and that the distributions do not overlap. Adjusted Wilcoxon test p-values < 0.001 are considered highly significant, whereas p > 0.05 is considered non-significant. Ten second-level objective functions are shown in columns, while landscape-sampling approaches, SNP-filtering thresholds, and PC-space types are shown in rows. Performance was assessed using Pearson’s or Spearman’s correlations with genetic distances in the left half of the table, corresponding to **Supplementary Fig. 12**, and using Pearson’s correlations with geographic distances for the same top-ranked LDEs in the right half, corresponding to **Supplementary Fig. 13**. As a control, we reversed the protocol: LDEs were ranked by their fit to geographic distances, and Spearman’s correlations with GRM-derived dissimilarities were then assessed for the top-ranked LDEs.

However, improvement in the fit to both genetic and geographic distances, using PC1–PC2– PC3 spaces as the baseline, is much rarer (**Table 1**). The *F_ST_* and *f*_2_ objectives, which are nearly identical to each other (**Supplementary Fig. 10**), fail across the board with respect to Pearson’s correlation between LDE distances and geographic distances, under all landscape-sampling and SNP-filtering approaches tested (**Table 1**, **Supplementary Fig. 13a–d**). These declines in correlation are often substantial, for example from ∼0.5 to ∼0.09 (**Supplementary Fig. 13b,d**). The only second-level objective that achieves median Pearson’s correlations with geographic distances that are consistently as good as, or significantly better than, those obtained in the corresponding PC1–PC2–PC3 spaces is GRM-Spearman (**Fig. 3b**, **Table 1**). GRM-Pearson performs worse under some conditions, as do covariance-Pearson and covariance-Spearman (**Table 1**, **Supplementary Fig. 13e–g**), and especially Hamming-Pearson and Hamming-Spearman. The latter two objectives are almost as bad in this respect as the *f*_2_- and *F_ST_*-based ones (**Table 1**, **Supplementary Fig. 13i,j**). Thus, we prioritize the GRM-Spearman objective for practical applications, at least for high-quality empirical datasets – diploid and with low missingness – whose levels of genetic diversity are comparable to those of our simulated landscapes, that is, to intracontinental diversity in humans (**Supplementary Fig. 14a**). We further compare different LDE-ranking objectives below in case studies based on real genetic data. Inspecting the results of manifold-learning optimization guided by the GRM-Spearman objective, we find that manifold-learning optimization is most useful – that is, it achieves the greatest improvement over direct PCA visualization in fits to both GRM-derived dissimilarities and geographic distances – for randomly sampled landscapes with MAF-filtered SNP data (**Fig. 3**). This comparison applies to the best-performing PC-space type (**Extended Data Fig. 3**): classic drift-normalized PC spaces.

The finding that a GRM-based LDE-ranking objective recovers landscape patterns – often distorted in PC1–PC2 (**Supplementary Figs. 3 & 4**) or PC1–PC2–PC3 spaces (**Supplementary Dataset 2**) – better than the other objectives we tested is consistent with the properties of GRM-derived dissimilarity (Yang et al. 2011). This measure is sensitive to shared rare variants, which are informative about migration and admixture among close neighbors, and therefore may “anchor” sampled individuals in the simulated genetic landscape more effectively than the other genetic distance measures tested.

We also showed that, although manifold-learning-derived LDEs that fit geographic distances very well can be found within the large parameter spaces we explore (**Supplementary Fig. 12k,l**), these LDEs usually fit GRM-derived dissimilarities worse than the corresponding PC1– PC2–PC3 spaces (**Table 1**, **Supplementary Fig. 13k,l**). This demonstrates that IBR landscapes with long-distance gene flows generate “genetic manifolds” that do not fully conform to geography.

#### Dissecting the manifold-learning optimization process

Above, we treated the manifold-learning optimization algorithm as a black box that produces a single best embedding for each input dataset, selected using a given LDE-ranking objective. Here, we examine which manifold-learning algorithms, or loss functions, and parameter settings produce these best-fitting LDEs. Crucially, by a unique input dataset we mean a specific combination of individuals, defined by the simulation and subsampling replicate, and SNPs, determined by the MAF filter. Other data-processing choices, such as PC-space type, are treated as part of the hyperparameter space. We focus on the GRM-Spearman LDE-ranking objective (**Fig. 4**), which performed best in our IBR-LDM landscape simulations (**Table 1**); results for the other objectives are presented in **Supplementary Figs. 7 & 15**.

To visualize these patterns more smoothly, we map not only the single best LDE, but also the top 100 LDEs for each unique input dataset, in a simplified hyperparameter space that retains all relevant dimensions except variation in the *k*-NN parameter (**Fig. 4**, **Supplementary Fig. 15**). A key observation is that no single manifold-learning algorithm or PC-space distance metric is optimal across the four dataset types we explored: random versus clustered sampling, and all SNPs versus SNPs with MAF > 1%. The most conspicuous differences are between datasets retaining all SNPs and those subjected to MAF filtering. In the former case, two main optima are observed: (1) approximately full-dimensional drift-normalized classic PC spaces analyzed with UMAP using the cosine distance metric; and (2) full-dimensional centered classic PC spaces analyzed with UMAP using the cosine, correlation, or Euclidean distance metric (**Fig. 4**). In the latter case, the main optima are different: (1) high-dimensional, but not full-dimensional, drift-normalized classic PC spaces analyzed with UMAP using the cosine or correlation distance metric; and (2) high-dimensional and full-dimensional centered classic PC spaces analyzed with PHATE using the cosine distance metric (**Fig. 4**). The latter optimum is specific to the clustered landscape-sampling approach and represents the clearest difference between clustered and randomly sampled datasets.

Only a few top-ranked LDEs are based on low-dimensional PC spaces (*n* ≤ 10). These secondary optima are specific to randomly sampled landscapes and correspond to all PCA types and to UMAP, or to a loss function intermediate between UMAP and densMAP, using Euclidean or Manhattan distances (**Fig. 4**). This region of parameter space is closest to the most common manifold-learning approach in population genetics: UMAP applied to a relatively low-dimensional eigenvector (*U*) input and using Euclidean distance (Diaz-Papkovich et al. 2019, 2021). These results demonstrate the value of optimization. By varying PCA type, input-space dimensionality, distance metric, and even the manifold-learning algorithm itself, we move beyond clearly suboptimal default settings and identify distinct optima that depend on the properties of the data. We expect real archaeogenetic datasets to most often resemble the clustered, MAF-filtered scenario, because of major gaps in spatiotemporal coverage (Racimo et al. 2020; Muktupavela et al. 2022), the difficulty of calling rare variants in error-rich ancient DNA data (Hui et al. 2020; Sousa da Mota et al. 2023), and widespread reliance on SNP arrays for targeted enrichment and PCA projection. Notably, PHATE, a relatively uncommon manifold-learning algorithm, often emerges as the top-performing method under these conditions. This is consistent with our human SNP-array case studies, which were prioritized for empirical testing and consistently identified PHATE as either the best-performing or one of the best-performing algorithms (**Table 2**). Another important take-home message is that the eigenvector spaces overwhelmingly used in population genetics are clearly suboptimal as inputs for manifold-learning-based LDE construction.

**Table 2.**
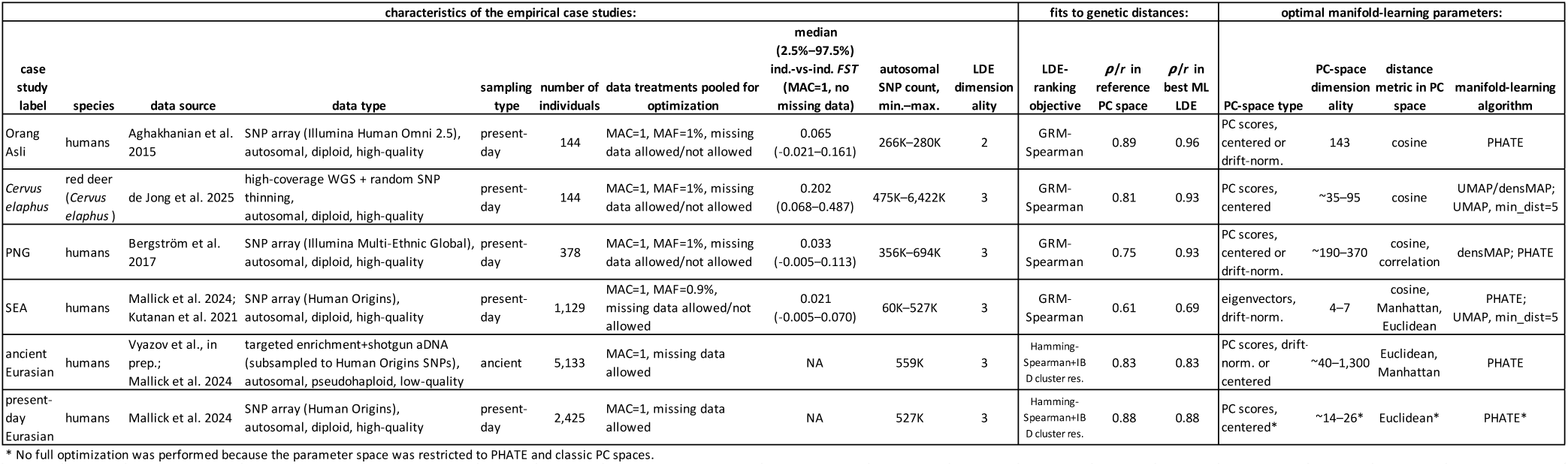
Summary of the six empirical case studies, including dataset metadata and optimal manifold-learning parameters.

| characteristics of the empirical case studies: |  |  |  |  |  |  |  |  |  | fits to genetic distances: |  |  | optimal manifold-learning parameters: |  |  |  |
| --- | --- | --- | --- | --- | --- | --- | --- | --- | --- | --- | --- | --- | --- | --- | --- | --- |
| case study label | species | data source | data type | sampling type | number of individuals | data treatments pooled for optimization | median (2.5%–97.5%) ind.-vs-ind. $F_{ST}$ (MAC=1, no missing data) | autosomal SNP count, min.–max. | LDE dimensionality | LDE-ranking objective | $\rho/r$ in reference PC space | $\rho/r$ in best ML LDE | PC-space type | PC-space dimensionality | distance metric in PC space | manifold-learning algorithm |
| Orang Asli | humans | Aghakhanian et al. 2015 | SNP array (Illumina Human Omni 2.5), autosomal, diploid, high-quality | present-day | 144 | MAC=1, MAF=1%, missing data allowed/not allowed | 0.065 (-0.021–0.161) | 266K–280K | 2 | GRM-Spearman | 0.89 | 0.96 | PC scores, centered or drift-norm. | 143 | cosine | PHATE |
| Cervus elaphus | red deer (Cervus elaphus) | de Jong et al. 2025 | high-coverage WGS + random SNP thinning, autosomal, diploid, high-quality | present-day | 144 | MAC=1, MAF=1%, missing data allowed/not allowed | 0.202 (0.068–0.487) | 475K–6,422K | 3 | GRM-Spearman | 0.81 | 0.93 | PC scores, centered | ~35–95 | cosine | UMAP/densMAP; UMAP, min_dist=5 |
| PNG | humans | Bergström et al. 2017 | SNP array (Illumina Multi-Ethnic Global), autosomal, diploid, high-quality | present-day | 378 | MAC=1, MAF=1%, missing data allowed/not allowed | 0.033 (-0.005–0.113) | 356K–694K | 3 | GRM-Spearman | 0.75 | 0.93 | PC scores, centered or drift-norm. | ~190–370 | cosine, correlation | densMAP; PHATE |
| SEA | humans | Mallick et al. 2024; Kutanan et al. 2021 | SNP array (Human Origins), autosomal, diploid, high-quality | present-day | 1,129 | MAC=1, MAF=0.9%, missing data allowed/not allowed | 0.021 (-0.005–0.070) | 60K–527K | 3 | GRM-Spearman | 0.61 | 0.69 | eigenvectors, drift-norm. | 4–7 | cosine, Manhattan, Euclidean | PHATE; UMAP, min_dist=5 |
| ancient Eurasian | humans | Vyazov et al., in prep.; Mallick et al. 2024 | targeted enrichment+shotgun aDNA (subsampled to Human Origins SNPs), autosomal, pseudohaploid, low-quality | ancient | 5,133 | MAC=1, missing data allowed | NA | 559K | 3 | Hamming-Spearman+IB D cluster res. | 0.83 | 0.83 | PC scores, drift-norm. or centered | ~40–1,300 | Euclidean, Manhattan | PHATE |
| present-day Eurasian | humans | Mallick et al. 2024 | SNP array (Human Origins), autosomal, diploid, high-quality | present-day | 2,425 | MAC=1, missing data allowed | NA | 527K | 3 | Hamming-Spearman+IB D cluster res. | 0.88 | 0.88 | PC scores, centered* | ~14–26* | Euclidean* | PHATE* |

\* No full optimization was performed because the parameter space was restricted to PHATE and classic PC spaces.

Distributions of top-ranked LDEs selected using the other second-level objectives are shown in the same parameter spaces in **Supplementary Fig. 15**. The most complete parameter spaces – sets of 228 PC-space dimensionality versus *k*-NN biplots per objective – are shown in **Supplementary Dataset 1**. These biplots show the median (across simulation and subsampling replicates) Δ*r*^2^ or Δ*ρ*^2^ between manifold-learning-derived LDEs and the corresponding classic drift-normalized PC1–PC2–PC3 spaces. We also provide an interactive application that visualizes all PC1–PC2–PC3 spaces for the IBR-LDM simulations and generates any manifold-learning-derived LDE within our parameter space on the fly (**Supplementary Dataset 2**).

Comparison of the manifold-learning optima identified above with scree plots generated from the same data shows that the conventional Cattell elbow criterion (Cattell 1966; Jackson 1993; Peres-Neto et al. 2005), widely used in statistical-genetic and population-genetic PCA applications (Luu et al. 2017; Abegaz et al. 2018; Tyrmi et al. 2020; Pélissié et al. 2022), is not suitable for manifold-learning optimization. In our data derived from IBR-LDM landscapes, the inflection points of the eigenvalue curves occur at approximately PC 20, or even earlier, across all data-normalization and SNP-filtering strategies. This pattern is especially clear for MAF-filtered data and centered PC spaces (**Supplementary Fig. 7a**). By contrast, the main manifold-learning optima occur either in full-dimensional PC spaces (*n* = 199) or, in the case of clustered sampling with MAF = 1%, at dimensions of at least *n* ≈ 70 (**Fig. 4**).

### Applying manifold-learning optimization to empirical population-genetic data

We tested our objective-guided embedding-optimization approach on six case studies using real genetic data. Key parameters and outcomes of these tests are summarized in **Table 2**. We begin by describing results for scenarios most closely resembling our simulated data: contemporaneous sampling; moderate levels of genetic differentiation, subcontinental in the case of humans; and high-quality diploid data from humans or animals. We then move to more divergent scenarios, including high-quality data from human populations spanning intercontinental levels of diversity, and low-quality ancient human data projected onto high-quality present-day data with high levels of genetic differentiation.

#### Orang Asli – present-day indigenous groups from the Malay Peninsula (Aghakhanian et al. 2015)

We begin with the most geographically and genetically “compact” human dataset analyzed in this study: Illumina Human Omni 2.5 SNP-array data (Aghakhanian et al. 2015) for nine Indigenous Orang Asli present-day groups from the Malay Peninsula (144 individuals and ca. 280,000 autosomal SNPs polymorphic in the dataset; **Table 2**, **Supplementary Table 2**). First, we generated combinations of three PCA protocols – eigenvectors, centered and drift-normalized; classic PC scores, either centered only or centered and drift-normalized – and two levels of rare-variant removal: no removal or MAF ≈ 1% (MAC = 3 in this case). Because the *smartPCA/PLINK* PCA algorithm is more robust to missing data than the *smartSNP* algorithm (**Methods**), we also tested two levels of missing-data filtering before applying *smartPCA/PLINK*: missing data allowed or no missing data allowed at the SNP level. Missing data were always removed before calculating classic PC scores with *smartSNP* (**Supplementary Table 1**). We assessed linear and nonlinear fits, using Pearson’s and Spearman’s correlation coefficients, between the resulting PCA LDEs – PC1–PC2 spaces in this case – and six types of genetic distances or dissimilarities applied at the level of individuals (PCA and genetic distances were always calculated using the same SNP sets). Most of these distances/dissimilarities overlapped with the metrics considered for the simulated data (**Supplementary Fig. 10**); however, we also tested a version of Hamming distance with a “flat” missing-data correction in addition to the default missing-data treatment in *PLINK* v. 1.9 (Chang et al. 2015), and calculated outgroup *f*_3_-statistics of the form *f*_3_(Mbuti; X, Y). Remarkably, across all these metrics, PC1–PC2 spaces based on classic PC scores fit the data slightly better than eigenvector-based spaces (**Supplementary Table 1**). Because all PCA LDEs based on classic PC scores showed nearly equivalent fits, we selected the drift-normalized classic PC-score space as the reference PC space (MAC = 1, no missing data allowed), since drift normalization is much more commonly used in practice than centering only.

Although the selected PC1–PC2 space shows high fits to the genetic distances tested (e.g., *ρ*_GRM_ = 0.90; **Supplementary Table 1**), it nevertheless displays the three-ray pattern discussed above, which is typical of PCA visualizations of similarly “compact” datasets or datasets in which little information about local structure is preserved. The southernmost Indigenous group, Seletar, forms a long separate cline, whereas the other eight groups occupy a 1D cline (**Extended Data Fig. 5**) that shows some correspondence to geography (*r*_geogr_ = 0.76 for all individuals).

Inspection of the fits between *n*-dimensional PC spaces and the four key genetic distance metrics (*F_ST_*, Hamming distance, GRM-derived dissimilarity, and outgroup *f*_3_-statistics; **Supplementary Table 3**) revealed a clear pattern in this dataset: correlations approaching 1 were reached in full-dimensional spaces based on classic PC scores, rather than in eigenvector-based PC spaces or spaces of lower dimensionality, for all four metrics (**Supplementary Fig. 16a–e**). In contrast, application of the Cattell elbow criterion (Cattell 1966; Jackson 1993; Peres-Neto et al. 2005) would have selected a PC-space dimensionality close to 10 (**Supplementary Fig. 16f**), which is suboptimal for manifold learning, as we illustrate shortly.

We then performed objective-guided embedding optimization guided by 12 alternative LDE-ranking objectives, corresponding to the fits to the distance metrics listed in **Supplementary Table 1**. The highest PC-space dimensionality explored was *s − 1* = 143. In contrast to the optimization runs on simulated data, where MAC = 1 and MAF = 1% runs were performed separately, here both missing-data filtering levels and both MAF-filtering levels were included in a single optimization run for simplicity. Correlations among all genetic and geographic distances or dissimilarities are shown in **Supplementary Table 3**. We note that outgroup *f*_3_-statistics, a data type often used in archaeogenetics for PCA or MDS (e.g., Fu et al. 2016; Posth et al. 2023; Maravall-López et al. 2025), correlate very well in this dataset with Hamming distance and with GRM-derived or covariance-derived dissimilarity (*r* > 0.95).

We explored 225,264 2D LDEs during objective-guided embedding optimization and applied the GRM-Spearman LDE-ranking objective, which was identified as the best-performing objective for simulated IBR-LDM landscapes (**Table 1**). These landscapes are also relatively compact in terms of genetic differentiation (compare the *F_ST_* values displayed in **Supplementary Fig. 14** and **Supplementary Table 1**), contemporaneous, and based on high-quality data, making them broadly comparable to the Orang Asli dataset. The upper tail of the average fit score, *ρ*² (see **Box 1** for definition), across the *F_ST_*-Spearman, Hamming-Spearman, GRM-Spearman, and *f*_3_-Spearman objectives is populated mostly by LDEs selected under the Hamming-Spearman, Hamming-Pearson, and GRM-Spearman objectives (**Supplementary Dataset 3**). This shows that these objectives achieve the best compromise fit to multiple types of genetic distances, all of which are highly or moderately correlated in this dataset (*ρ* = 0.65– 0.94; **Supplementary Table 3**).

Thus, use of the plain GRM-Spearman objective 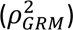 is justified for this case study. This objective selected the following 2D LDE as the global optimum: a dataset with missing data removed and no SNP filtering, a PHATE LDE based on the full-dimensional classic centered PC space, cosine distance in that space, and a moderately low *k*-NN value of 18 (**Supplementary Dataset 3**). This LDE achieved a nearly ideal *ρ*_*GRM*_= 0.96, improving on the reference PC1– PC2 space (no missing data, MAC = 1, classic drift-normalized PC scores), which had *ρ*_*GRM*_ = 0.89 (**Extended Data Fig. 5**). Relative to this reference PC1–PC2 space, the optimized LDE showed no improvement in *ρ*_*Hamming*_(both ∼0.90), a decrease in *ρ*_*F_ST_*_ (from 0.81 to 0.74), an increase in *ρ*_*f*3_ (from 0.80 to 0.90), and an increase in *r*_*geogr*_(from 0.76 to 0.83) (**Extended Data Fig. 5**). We note that the optimum predicted from correlations between GRM-derived dissimilarity and distances in *n*-dimensional PC spaces lies exactly at this position: MAC = 1, classic centered PC scores, full-dimensional PC space, and cosine distance applied to that space (**Supplementary Fig. 16c**).

Inspection of the full parameter space of the optimization run confirms that the optimum of 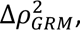 calculated relative to the drift-normalized classic PC space at MAC = 1, is well localized: MAC = 1 or MAF = 1%, input-space dimensionality approaching or equal to *s − 1*, PHATE, cosine distance in classic PC space, and *k*-NN around 20 (**Supplementary Dataset 4**). UMAP and densMAP perform worse, and all manifold-learning methods applied to eigenvector spaces perform substantially worse (**Supplementary Dataset 4**). The poor performance of eigenvector spaces is consistent with both the simulated results and our analysis of *n*-dimensional input PC spaces in **Supplementary Fig. 16**.

Although both the reference PC1–PC2 space and the top-ranked LDE show broadly similar patterns and fit genetic distances well, the better quantitative performance of the latter is reflected in its “unfolding” of approximately 1D clines into 2D gradients. For example, internal structure becomes visible within the southernmost Seletar group, and the groups of the northern cluster – Jehai, Kintaq, and Bateq + Mendriq, which are very close geographic neighbors – become partially resolved (**Extended Data Fig. 5a**; see the map in Aghakhanian et al. 2015). The geographically central Cheq Wong group, together with a few outliers from other groups, also forms a more distinct cluster in the PHATE space. Remarkably, this optimized manifold-learning-derived embedding agrees well with a completely independent analysis: a force-directed layout of the IBD-sharing graph constructed for the same individuals using an IBD segment-length threshold of ≥8 cM (**Extended Data Fig. 5b**). One caveat is that IBD graphs depend on the chosen segment-length cut-off, because different thresholds emphasize relatedness at different time depths. Nevertheless, the positions of outliers within each ethnic group in the PHATE space are more informative about their IBD-connection patterns than their positions in the PC1–PC2 space. For example, Mah Meri individual M11 is the Mah Meri individual closest to the Cheq Wong cluster in the IBD-sharing graph, a pattern visible in the PHATE space but not in the PC1–PC2 embedding (**Extended Data Fig. 5**). Similarly, Temuan individual T5 is attracted toward the “northern” cluster (Bateq, Jehai, Kintaq, Mendriq) in the IBD graph, but this pattern is not clearly resolved in the PC1–PC2 embedding; by contrast, in the PHATE space this individual appears as a distant outlier within the Temuan group (**Extended Data Fig. 5**). The same pattern is observed for the Jakun outlier JKN040.

The top-ranked LDE fits geography well, even when compared with the least realistic, great-circle, geographic distances (**Extended Data Fig. 5a**). Together with the presence of outliers from several groups clustering with other groups, likely representing recent migrants or their admixed descendants, this suggests that Indigenous groups in the Malay Peninsula lie on a genetic landscape approaching an isolation-by-distance scenario.

#### Present-day red deer (Cervus elaphus) (de Jong et al. 2025)

We applied the framework described for the Orang Asli analysis to additional empirical-data case studies. Here, we describe results for a present-day red deer (*Cervus elaphus*) dataset (de Jong et al. 2025) comprising high-coverage shotgun genomes from 144 individuals sampled across Europe, North Africa, Anatolia, and the Caucasus, and including approximately 6,422,000 autosomal SNPs polymorphic in the dataset (**Table 2**, **Supplementary Table 2**). Here and in subsequent case studies, we focused on 3D LDEs instead of the more commonly used 2D LDEs. All tested PC1–PC2–PC3 spaces showed broadly similar fits to the set of genetic distances and dissimilarities used here, with some variability driven by a small number of individuals whose positions changed substantially depending on MAF filtering and missing-data removal (see interactive 3D plots in **Supplementary Dataset 5a–h**). The best fits were obtained for the centered or drift-normalized classic PC spaces **(Supplementary Table 1)**. For consistency with the other case studies, we selected the latter as the reference PC space, using MAC = 1 and allowing no missing data.

Manifold-learning optimization was guided by 14 alternative LDE-ranking objectives (**Supplementary Dataset 3**). We note that, in this dataset, *f*₃-statistics computed using four wapiti (*Cervus canadensis*) individuals as the outgroup correlate less strongly with the other metrics than in the Orang Asli dataset; otherwise, the overall correlation patterns are similar (**Supplementary Table 3**). We explored 225,264 3D LDEs and, because the dataset is broadly similar to our simulated landscapes in genetic differentiation (compare **Supplementary Fig. 14** and **Supplementary Table 1**) and data quality, applied the GRM-Spearman objective. The upper tail of the average fit score, *ρ*², calculated across the *F_ST_*-Spearman, Hamming-Spearman, GRM-Spearman, and *f*₃-Spearman objectives, was again dominated by LDEs selected under the Hamming-Spearman, Hamming-Pearson, and the closely correlated GRM-Spearman and covariance-Spearman objectives (**Supplementary Dataset 3**). The GRM-Spearman objective selected an LDE generated with the algorithm intermediate between standard UMAP and densMAP (*uwot dens_scale* = 0.5) from a high-dimensional, centered classic PC space comprising 60 PCs, using cosine distance in that space and a moderately low *k*-NN value of 22 (**Table 2**, **Supplementary Dataset 3**). This embedding was based on the dataset filtered to remove missing data and retain variants with MAF ≥1%. Embeddings from the same region of parameter space, but generated from the dataset without MAF filtering, achieved only slightly lower fits to GRM-derived dissimilarities (**Supplementary Datasets 3 & 6**). As in the Orang Asli case study, the top-ranked LDE achieved a nearly ideal *ρ*_*GRM*_= 0.93, improving on the reference and input PC1–PC2–PC3 spaces, which both yielded *ρ*_*GRM*_= 0.81 (**Supplementary Table 1**). Relative to this reference PC space, the optimized LDE showed a slight decrease in *ρ*_*Hamming*_, from 0.85 to 0.78; a substantial decrease in *ρ*_*F_ST_*_, from 0.61 to 0.41 (this is consistent with *F_ST_* and *f*₂ being outlier metrics that correlate poorly with the other measures; **Supplementary Table 3**); an increase in *ρ*_*f*3_, from 0.45 to 0.60; and an increase in *r*_*geogr*_, from 0.65 to 0.71. The broader global optimum in the hyperparameter space was located in the following region: MAC = 1 or MAF = 1%; input-space dimensionality of approximately 35–95 PCs; UMAP/densMAP, or UMAP with *min_dist = 0.5*; cosine distance in centered classic PC space; and *k*-NN values of approximately 20–60 (**Table 2**, **Supplementary Datasets 3 & 6**). All manifold-learning methods applied to eigenvector spaces performed substantially worse (**Supplementary Dataset 6**), in line with the results reported above.

Correlations between GRM-derived dissimilarities and distances in *n*-dimensional PC spaces had lower predictive power in the *Cervus elaphus* case study than in the Orang Asli case study. Correlation coefficients approached 1 in full-dimensional classic PC spaces when cosine or correlation distances were applied (**Supplementary Fig. 17c**). This region of hyperparameter space was indeed optimal for manifold learning, with one exception: 35–95-dimensional PC spaces were optimal, rather than the full-dimensional (143-dimensional) PC spaces (**Supplementary Dataset 6**). Application of the Cattell elbow criterion would have selected a PC-space dimensionality of approximately 20–35, depending on the type of PCA (**Supplementary Fig. 17f**), which is suboptimal for manifold learning.

The PC1–PC2 space from the original study (de Jong et al. 2025) and the PC1–PC2–PC3 spaces considered here show very similar simplicial geometries: most European red deer individuals form an approximately 1D cline, whereas individuals from the Italian Peninsula/Sardinia/Tunisia, the Iberian Peninsula, and the Caucasus/Anatolia appear as distant outliers occupying the vertices of a triangle or tetrahedron (**Supplementary Dataset 5a–h**). In contrast, the top-ranked LDE retains the Italian/Sardinian/Tunisian and Iberian populations as outliers, while highlighting population structure within Europe, Anatolia, and the Caucasus, organized along southwest–southeast and northwest–northeast clines (**Supplementary Dataset 5i**). This pattern is consistent with predominant isolation by distance among European red deer populations.

#### Present-day humans from Papua New Guinea (Bergström et al. 2017)

In this section, we analyze SNP-array data generated with the Illumina Infinium Multi-Ethnic Global Array for 378 present-day individuals from Papua New Guinea (Bergström et al. 2017), including approximately 694,000 autosomal SNPs polymorphic in the dataset (**Table 2**, **Supplementary Table 2**). Centered and drift-normalized classic PC1–PC2–PC3 spaces showed better fits to the genetic distances and dissimilarities considered here than eigenvector spaces across all metrics (**Supplementary Table 1**; see interactive 3D plots in **Supplementary Dataset 7a–h**). For consistency with the other case studies, we selected the drift-normalized classic PC space as the reference PC space, using MAC = 1 and allowing no missing data.

Manifold-learning optimization was guided by 14 alternative LDE-ranking objectives (**Supplementary Dataset 3**). We explored 692,512 3D LDEs and applied the GRM-Spearman objective, which selected an LDE generated with densMAP from a nearly full-dimensional, drift-normalized classic PC space comprising 360 PCs, using cosine distance in that space and a high *k*-NN value of 180 (**Table 2**, **Supplementary Dataset 3**). This embedding was based on the dataset filtered to remove missing data and retain all variants (MAC = 1). The top-ranked LDE again achieved a nearly ideal *ρ*_*GRM*_= 0.93, improving on the reference/input PC1–PC2– PC3 space, which yielded *ρ*_*GRM*_ = 0.75 (**Supplementary Table 1**). Consistent with the relatively weak correlations of GRM- and covariance-derived dissimilarities with the other metrics in this dataset (**Supplementary Table 3**), the optimized LDE showed poorer fits to these metrics than the reference PCA LDE: *ρ*_*Hamming*_ decreased from 0.80 to 0.66; *ρ*_*F_ST_*_, from 0.71 to 0.63; *ρ*_*f*3_, from 0.80 to 0.65; and *r*_*geogr*_, from 0.61 to 0.53. The broader global optimum in the hyperparameter space was located in the following region: MAC = 1 or MAF = 1%; input-space dimensionality of approximately 190–370 PCs; densMAP or PHATE; cosine or correlation distance in centered or drift-normalized classic PC space; and *k*-NN values of approximately 100–290 (**Table 2**, **Supplementary Datasets 3 & 8**). All manifold-learning methods applied to eigenvector spaces performed substantially worse (**Supplementary Dataset 8**), in line with the results above.

Correlation coefficients between GRM-derived dissimilarities and distances in *n*-dimensional PC spaces approached 1 in full-dimensional classic PC spaces when cosine or correlation distances were applied (**Supplementary Fig. 18c**). This region of hyperparameter space was indeed optimal for manifold learning, with one exception: PC spaces comprising 190–370 dimensions were optimal, rather than the full 377-dimensional space (**Supplementary Dataset 8**). This pattern closely resembles that observed in the *Cervus elaphus* case study. Application of the Cattell elbow criterion would have selected a PC-space dimensionality of approximately 10 (**Supplementary Fig. 18f**), which is suboptimal for manifold learning.

The original study (Bergström et al. 2017) estimated PC axes separately using Papuan highlanders or lowlanders and projected the other group onto them, precluding direct comparison with our results. Our joint 3D analysis of both groups shows that this potentially artificial *a priori* division is unnecessary. Highlander and lowlander variation are orthogonal in our PC spaces (**Supplementary Dataset 7a–h**): our PC2–PC3 space recapitulates the highlander PC1–PC2 pattern reported in the original study (Fig. 2a and Fig. S10 in that paper), whereas our PC1–PC3 space recapitulates the corresponding lowlander PC1–PC2 pattern (Fig. S11 in that paper). Relative to our PC1–PC2–PC3 spaces (**Supplementary Dataset 7a–h**), the optimized LDE (**Supplementary Dataset 7i**) resolves additional structure within the cluster comprising individuals from the neighboring Southern Highlands and Enga provinces. The lowlander genetic gradients revealed by both PCA and manifold-learning-derived LDEs are largely geographic, forming a semicircle around the central mountain range: East Sepik– Madang–Milne Bay/Central–Gulf–Western provinces. Although this pattern is visible in Fig. S11, it was not discussed in the original study. Thus, the main advantage of our manifold-learning-derived LDE over both previously published and our PC visualizations is that the highlander and lowlander genetic gradients are not analyzed separately and not constrained to be orthogonal, allowing Papuan highlanders to be placed in the broader context of lowlander variation (**Supplementary Dataset 7i**).

#### Present-day humans from Southeast Asia (Mallick et al. 2024 & Kutanan et al. 2021)

We analyzed Human Origins SNP array data for 1,129 present-day individuals from Mainland and Island Southeast Asia (SEA), comprising approximately 527,000 autosomal SNPs polymorphic in the dataset (**Table 2**, **Supplementary Table 2**). The dataset combines a subset of Mallick et al. (2024) with data from Kutanan et al. (2021) and includes all ethnolinguistic groups from the region represented in these studies, except the most genetically divergent Negrito groups, which dominate the first PC when included.

The overall distribution of individuals in PC1–PC2–PC3 space is tetrahedral and dominated by highly drifted groups with low effective population sizes (tribal populations): distinct clines are formed primarily by Hmong-Mien speakers, Austronesian and Austroasiatic speakers, and Tibeto-Burman speakers, whereas large SEA populations such as Thai and Vietnamese gravitate toward the center of the space and form a poorly resolved cloud (see, e.g., **Supplementary Dataset 9b,h**). Notably, this structure fits all tested genetic and geographic distances poorly: the highest observed fits are *ρ*_GRM_ values of only up to 0.63, while the remaining key fits are substantially lower, with a median of 0.48, compared with 0.69–0.85 for the other five case studies (**Supplementary Table 1**). In this respect, the dataset stands out from the other case studies (**Supplementary Table 1**), suggesting that the selected sample composition is particularly suboptimal for PCA-based visualization of genetic structure. It is also distinguished by its low median inter-individual *F_ST_* of approximately 0.02 (**Table 2**, **Supplementary Table 1**). Moreover, some combinations of PCA algorithm, MAF filter, and missing-data removal regime produce distant outliers in PC1–PC2–PC3 space (**Supplementary Dataset 9a,c,g**). Given these limitations of PCA, this dataset provides a particularly informative test of whether manifold-learning optimization can substantially improve the preservation of genetic distances and dissimilarities. The drift-normalized classic PC space (MAC = 1, no missing data) showed the highest or second-highest fits to GRM-derived dissimilarities, Hamming distances, outgroup *f*_3_(Mbuti; X, Y), and geographic distances, and was therefore selected as the reference PC space.

Manifold-learning optimization was guided by 12 alternative LDE-ranking objectives (**Supplementary Dataset 3**). We explored 936,624 3D LDEs and applied the GRM-Spearman objective, which selected an embedding generated with PHATE from a low-dimensional, drift-normalized eigenvector space comprising 7 PCs, using Euclidean distance and a *k*-NN value of 70 (**Table 2**, **Supplementary Dataset 3**). This embedding was based on the dataset retaining missing data and all variants (MAC = 1). The broader global optimum in hyperparameter space occupied the following region: no missing data removed, MAC = 1 or MAF = 1%; input dimensionality of 3–7 PCs; PHATE or UMAP with *min_dist = 0.5*; cosine, Manhattan, or Euclidean distance in drift-normalized eigenvector space; and *k*-NN values of 50–120 for input dimensionalities >3 PCs (**Table 2**, **Supplementary Datasets 3 & 10**). A shallower optimum occurred in the region characteristic of the Orang Asli, *Cervus elaphus*, and PNG datasets: missing data removed, full-dimensional classic PC space with cosine or correlation distance, and UMAP or densMAP (**Supplementary Dataset 10**). This optimization landscape is unusual and may reflect a limitation of our analysis in this case study: because *smartSNP* handles missing data poorly, classic PCA was performed only after complete removal of sites containing missing genotypes, an approach that was probably overly aggressive, reducing the dataset from 436K–527K to only 60K–75K sites. Moreover, because the Human Origins data were assembled from multiple studies with different levels of missingness (median missingness = 7.6% in Mallick et al. 2024 and 1% in Kutanan et al. 2021; **Supplementary Table 2**), batch effects arising from non-random patterns of missing data cannot be excluded. This overly stringent missing-data filter may therefore have removed substantial information from the classic PC spaces, making the full-dimensional PC-score representation inferior to the low-dimensional eigenvector space. We retain this analysis as a cautionary example of how preprocessing choices can strongly affect manifold-learning optimization.

Correlations between GRM-derived dissimilarities and distances in *n*-dimensional PC spaces approached 1 in full-dimensional classic PC spaces when cosine or correlation distance was used (**Supplementary Fig. 19c,g**). As discussed above, however, this region of hyperparameter space represents only a secondary manifold-learning optimum; the primary GRM-Spearman optimum lies in low-dimensional eigenvector spaces derived from datasets without missing-data removal (**Supplementary Dataset 10**). Excluding 3-dimensional eigenvector spaces as inputs for 3D embeddings, since such setups do not extract additional information, this optimum is likely centered around 7 PCs used as input to PHATE (**Supplementary Dataset 10**). The primary optimum is also predicted by the correlation plots: the highest rank correlation with GRM-derived dissimilarities in these datasets (∼0.69 for MAC = 1, matching that of the top-ranked LDE) occurs in 7–9-dimensional eigenvector spaces (**Supplementary Fig. 19g**). By contrast, the Cattell elbow criterion would select approximately 20–50 PCs (**Supplementary Fig. 19f**), corresponding to neither optimum.

The top-ranked LDE achieved *ρ*_*GRM*_= 0.69, improving on the reference PC1–PC2–PC3 space, which yielded *ρ*_*GRM*_ = 0.61 (**Supplementary Table 1**). Consistent with the weak correlations of GRM- and covariance-derived dissimilarities with the other metrics in this dataset (**Supplementary Table 3**), the optimized LDE showed poorer fits to these metrics than the reference PCA LDE: *ρ*_*Hamming*_ decreased from 0.56 to 0.39; *ρ*_*F_ST_*_, from 0.41 to 0.27; *ρ*_*f*3_, from 0.47 to 0.32; and *r*_*geogr*_ increased from 0.24 to 0.30.

Relative to our PC1–PC2–PC3 spaces (e.g., **Supplementary Dataset 9b**), whose first three PCs are dominated by small ethnolinguistic groups, the optimized LDE (**Supplementary Dataset 9i**) resolves a much more prominent cline among Kra-Dai speakers and reveals their nuanced relationships with other groups, including patterns identified independently in our earlier work (Changmai et al. 2023). Most notably, the top-ranked LDE recovers the Indian-admixed SEA populations, including Mon, Burmese, Central Thai, Southern Thai, and Malay (Changmai et al. 2022; Kutanan et al. 2021), and does that in an unsupervised analysis and without South Asian reference groups. These groups are pulled out of the poorly resolved central cloud in PC1–PC2–PC3 space and form a distinct pole in the LDE (**Supplementary Dataset 9i**). Thus, despite the limitations described above, manifold-learning optimization guided by the GRM-Spearman objective appears particularly sensitive to South Asian admixture in SEA – a signal that remained largely undetected for some time in studies relying on conventional PCA, *qpAdm*, and *f*-statistics (compare Lipson et al. 2018 with Changmai et al. 2022).

#### Iron Age–Late Medieval Eurasians projected onto present-day variation: fine-scale European structure and early Slavic dispersal

We assembled both published (Mallick et al. 2024) and novel data (Vyazov et al., in preparation), generated using shotgun sequencing or targeted enrichment, for 5,133 ancient Eurasian individuals dated to ∼125–3,200 years before present. To address high missing rates typical for ancient data and sequencing-technology-driven batch effects (Rohland et al. 2022, Fournier et al. 2025), we projected their genome-wide SNP data onto PCs derived from Human Origins SNP data for 2,425 present-day Eurasians (see the geographic distributions of ancient and modern individuals in **Supplementary Fig. 20a,b** and additional metadata in **Supplementary Table 2**). This projection was performed in parallel using *smartPCA*, which produces eigenvectors, and *smartSNP*, which produces PC scores (see interactive 3D plots in **Supplementary Dataset 11** and fits in **Supplementary Tables 1 & 4**). Notably, the classic PC1– PC2–PC3 spaces fit all genetic distance/dissimilarity measures and great-circle geographic distances substantially better than the eigenvector space (**Supplementary Tables 1 & 4**), likely because of their more favorable aspect ratios (**Supplementary Fig. 21b**, **Supplementary Dataset 11**), and both centered and drift-normalized classic spaces demonstrate very similar fits (**Supplementary Tables 1 & 4**).

Medieval individuals from Slavic cultural contexts display relatively high within-group IBD-sharing, probably due to their low effective population size (*N_e_*): nearly all such individuals fall into a single IBD-sharing community (**Extended Data Fig. 6a**), which we label – based on the spatial (**Supplementary Fig. 20c**) and temporal distribution of its members – as “Central-East Europe 1.8–0.3 kya” (abbreviated as CEE). PC1–PC2 (**Fig. 5a**) and PC1–PC2–PC3 spaces (**Supplementary Dataset 11**) do not clearly distinguish members of the CEE IBD-sharing community from other European groups in the Eurasian context: for example, the “Northwest Europe 2.3–0.2 kya” (NWE) community – with 95.5% of its 246 members buried in Germanic contexts, including Norse (**Supplementary Table 2**) – is only weakly separated from the CEE community in PC1–PC2–PC3 spaces, as reflected by their Mahalanobis distances of ∼1–1.1 standard deviations in all three types of PC spaces (**Supplementary Table 4**). For the present-day IBD communities in PC1–PC2–PC3 spaces, the Mahalanobis distances between the Slavic/Baltic and Celtic/Germanic groups are similarly low, at ∼1.4–1.5 standard deviations (**Supplementary Table 4**).

**Figure 5.**
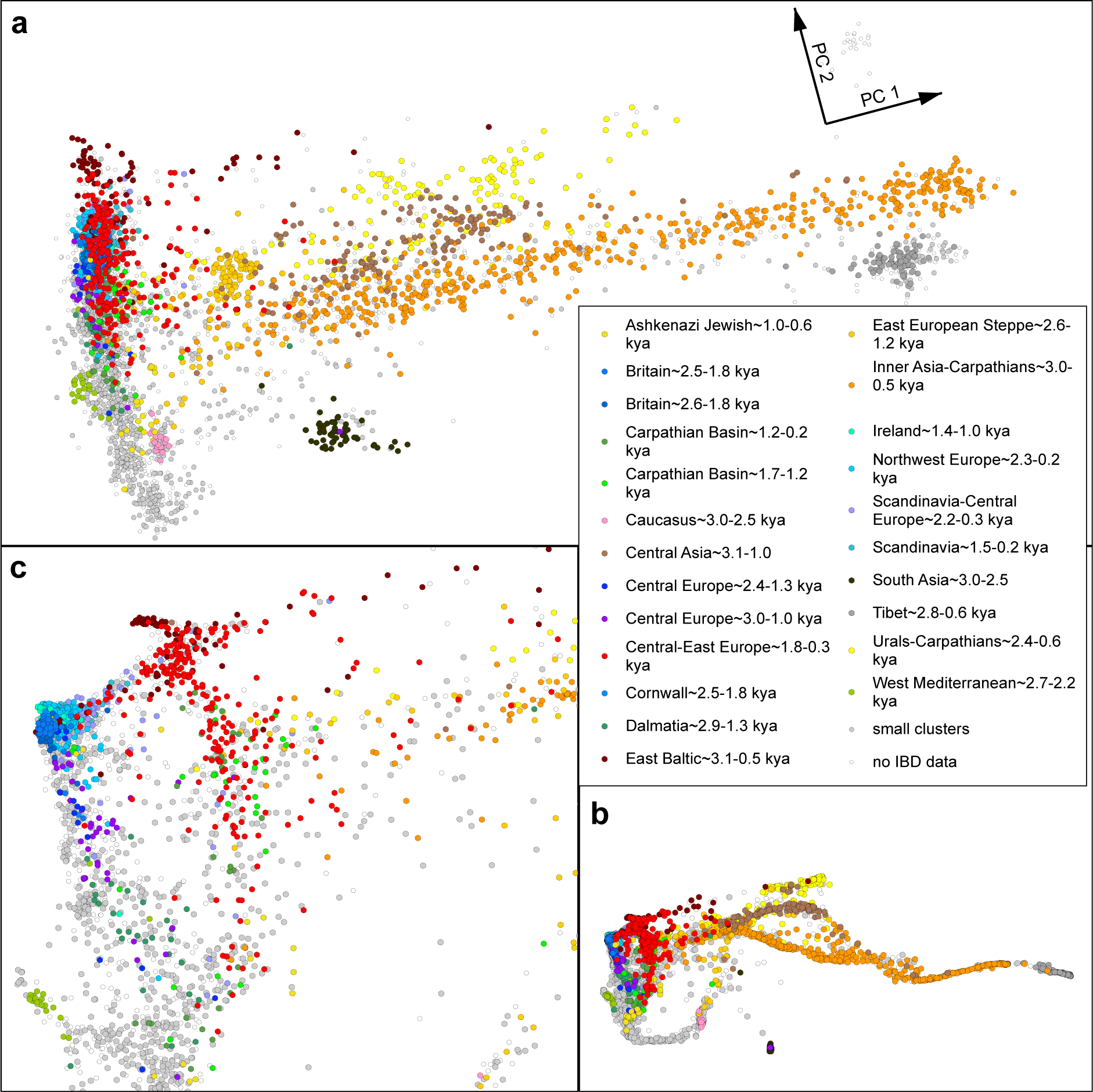
Genetic structure in Eurasia in the Iron Age to Late Medieval periods viewed through the lenses of genome-wide ancestry (embeddings constructed using classic PCA and PHATE) and networks of distant relatives (IBD-sharing communities). (**a**) Ancient individuals projected onto the PC space derived from present-day Eurasians genotyped on the Human Origins SNP panel (data centered and normalized by the allele-frequency dispersion expected under genetic drift; classic PC scores computed with *smartSNP*). Individuals in this and further panels (**b**, **c**) are colored according to the Leiden community-detection results on IBD data. (**b**) An optimized approximately isometric unfolding of the best-fitting PHATE 3D LDE performed with a customized Isomap algorithm and a zoomed-in view on Central and Northern Europe (**c**).

While a European-restricted PCA on a highly curated set of individuals and ca. 400,000 SNPs does separate Slavic and non-Slavic groups (Gretzinger et al. 2025), the approach is not ideal, as it masks signals of important migrations from Asia. To address these well-known limitations of PCA visualizations of human genetic variation across Eurasia (Olalde et al. 2023; Chobanov & Stamov 2025), we applied our objective-driven manifold-learning optimization approach to this dataset, but in contrast to the previous case studies on high-quality data for present-day individuals (**Table 2**), the resulting 3D LDEs were ranked via a multi-step procedure (**Extended Data Fig. 7**); see **Methods** section 5.4 for the rationale behind this approach. A coarse stage based on the GRM-Spearman and Hamming-Spearman objectives identified the preferred manifold-learning and PCA methods (reducing ∼809,000 LDEs to ∼85,000).

We then narrowed the resulting pool of ∼85,000 LDEs to the top 5% based on how well LDEs resolved the 23 largest IBD-sharing communities (**Extended Data Fig. 6a**). We confirmed that the Hamming-Spearman objective is more suitable for our data (pseudo-haploid, with high missing rates and different sequencing/capture technologies included; see **Supplementary Table 2**) than the GRM-based objective (**Extended Data Fig. 7**, **Methods** section 5.4), and we selected a small set of finalists (**Supplementary Dataset 12a–c**, **Supplementary Table 4**) using the former objective. As detailed in the **Methods**, this multi-step LDE ranking approach incorporates IBD information to compensate for the limited ability of the Hamming distance matrix to resolve local genetic structure and avoids the over-clustering often observed with default manifold-learning settings (Diaz-Papkovich et al. 2019, 2021; Moon et al. 2019; Battey et al. 2021; Chari & Pachter 2023; Meilă & Zhang 2024). We further demonstrated that optimizations beginning with the preferred manifold-learning and PCA methods and their ∼85,000 LDEs yield similar qualitative patterns (see interactive 3D LDEs in **Supplementary Dataset 13**) under diverse objectives, including a statistic quantifying preservation of sample neighborhoods in the (*s – 1*)-dimensional PC-space (**Methods** sections 5.5 and 5.8, **Supplementary Table 4**). The persistence of high Mahalanobis distances (>3 standard deviations) between the CEE and NWE IBD-sharing communities across a relatively wide parameter range supports this conclusion (**Supplementary Dataset 14**, **Supplementary Table 4**).

Compared with NNLS (Gretzinger et al. 2025), PANE (de Gennaro et al. 2025), and MOBEST (Schmid & Schiffels 2023) – approaches that infer ancestry or geographic affinity from multidimensional genetic representations – our objective-guided embedding optimization systematically optimizes the dimensionality of the input PC space, whereas these approaches are typically applied at fixed, relatively low dimensionalities (MOBEST: 3 PCs in Gretzinger et al. 2025; NNLS and PANE: 10 PCs). Notably, the PCA-based implementations of these approaches in Gretzinger et al. (2025) and de Gennaro et al. (2025) used *smartPCA* eigenvectors, which our analyses indicate are generally suboptimal inputs for manifold learning and may likewise be suboptimal for admixture-model inference.

A final outcome of the optimization procedure (**Fig. 5b-d**, **Supplementary Dataset 12a**) is a 3D LDE generated by the PHATE algorithm considering 26 nearest neighbors and based on Manhattan distances in a 44-dimensional classic drift-normalized PC space (for a discussion and visualization of the broader parameter optimum, see **Methods** sections 5.3 and 5.4, **Table 2**, **Supplementary Table 4**, and **Supplementary Dataset 14**). Compared with the corresponding PC1–PC2–PC3 space (**Supplementary Dataset 11a**), the best-scoring PHATE LDE yields a nearly identical fit to Hamming genetic distances (*ρ* = 0.83; **Table 2**, **Supplementary Table 4**). In contrast, median Mahalanobis separation across the 23 largest IBD communities (**Extended Data Fig. 6a**) increases markedly – from 3.78 standard deviations for the corresponding 3D PC space to 6.05 for the best-fitting PHATE 3D LDE (**Supplementary Table 4**), moving from the 15.8th to the 97.4th percentile among all PHATE embeddings on classic PC scores (**Supplementary Dataset 14**). This substantial boost in IBD community resolution indicates that the objective-guided manifold-learning optimization and multi-step LDE ranking effectively denoise the PCA-based embedding while maintaining aggregate correlations to genetic distances. For a more straightforward presentation, in **Fig. 5b,c** we show an optimized approximately isometric 2D unfolding of the best 3D LDE (**Methods** section 5.9). This unfolding yields correlations with great-circle geographic, Hamming, and GRM-derived genetic distances that are very close to those of the original 3D LDE (**Supplementary Table 4**).

We validated the best-fitting 3D LDE using an out-of-sample metric: the –log_10_(p-value) from “rotating” *qpAdm* tests (Fernandes et al. 2021; Flegontova et al. 2025) applied to pairs of sample clusters (**Methods** section 5.7). Across subsampling iterations, distances between cluster centroids in the PHATE space exhibited higher correlations with log-p-values of one-source *qpAdm* tests (on PHATE-based clusters) than distances in the drift-normalized classic PC space with one-source *qpAdm* results (on PCA-based clusters). For example, median iteration-wise Spearman correlation coefficients are as follows: *ρ*_PHATE_ ≈ 0.86 vs. *ρ*_PCA_ ≈ 0.82; unpaired Wilcoxon p ≈ 1.8 × 10^-4^.

The best-fitting LDE (shown as an interactive 3D visualization in **Supplementary Dataset 12a** and as the isometric 2D unfolding in **Fig. 5b,c**) reveals a nuanced genetic structure in the Medieval, Roman, and Iron Age Europe and, crucially, detects an ancestral component specific to the putatively Slavic CEE IBD-sharing community. PHATE keeps the major Asian clines, associated with historically recorded migrations of the Iranian, Turkic, and Uralic speakers, separated (**Fig. 5b,c**) and also resolves European populations into two major clines – “West Europe–West Mediterranean” and “East Europe–East Mediterranean” (abbreviated WE–WM and EE–EM, respectively) – a pattern that persists beyond the best-fitting PHATE LDE and is shared by LDEs across much of the parameter space (**Supplementary Dataset 14**, **Supplementary Datasets 12a–c & 13a–h**). We additionally demonstrated that the LDE ranking procedure is robust to the choice of objective (be it a genetic distance, a median inter-IBD-cluster distance, or a measure of how well sample neighborhoods in the (*s – 1*)-dimensional PC-space are preserved), as most European and Asian clines consistently reappear regardless of which objective is used (**Supplementary Table 4**, **Supplementary Datasets 12a–c & 13a– h**).

The earliest members of the CEE IBD community (∼150–550 CE) are found from the Middle Danubian Roman provinces to the Dniester and Don estuaries and the Middle Volga, where they appear as genetic outliers relative to local populations (**Extended Data Fig. 6c**). Despite this wide distribution, they form a compact cluster in the PHATE (but not in PC1–PC2–PC3) space, suggesting limited admixture with surrounding groups and making them the best available proxy for the shared genetic profile of the CEE community—and, by implication, of early Slavic speakers.

#### Present-day humans from Eurasia (Mallick et al. 2024)

We also performed manifold-learning optimization on the reference set of Human Origins SNP data for 2,425 present-day individuals (Mallick et al. 2024) used in the ancient Eurasian case study (**Supplementary Table 2**), exploring alternative optimization objectives or applying the multi-step LDE-ranking procedure (**Supplementary Tables 4 & 5b**). In this run, we explored a narrower parameter space, focusing on the broad optimum identified in the ancient Eurasian case study – restricting the dimensionality-reduction methods to PHATE and classic PCA. As compared to the ancient Eurasian case study, we also broadened the set of optimization objectives: included *ρ*^2^ and *r*^2^ fits to *F_ST_* and outgroup *f*_3_-statistics *f*_3_(Mbuti; X, Y) and omitted Y-chromosome-based cluster-separation metrics (**Supplementary Table 5b**). Visualizations of the main LDE fidelity metrics across the parameter space are provided in **Supplementary Dataset 15**.

Neighborhood preservation in the (*s – 1*)-dimensional classic PC space showed little relationship to *ρ*^2^ fits based on any of the four genetic distance/dissimilarity metrics (Spearman’s *ρ* ranging from 0.12 to –0.40 for *k* = 250 or 50; **Supplementary Table 5b**), contradicting the results on the ancient Eurasian dataset (**Supplementary Table 5a**). However, it did correlate with the median Mahalanobis (or Bhattacharyya) separation among the 41 largest IBD-sharing communities (Spearman’s *ρ* ranging from 0.66 to 0.72 for *k* = 250 or 50; **Supplementary Table 5b**). Consistent with this pattern, when *ρ*^2^ or *r*^2^ fits to genetic distances/dissimilarities were used as the sole optimization targets, the best-scoring LDEs still failed to resolve the major Asian or European clines (**Supplementary Dataset 16a–h**, **Supplementary Table 4**). Conversely, optimizing only the median Mahalanobis separation of IBD clusters produced well-resolved European and Asian structure, but at the cost of pronounced overclustering (**Supplementary Dataset 16i**). Using a neighborhood-preservation measure as the sole optimization objective mitigated this overclustering while retaining clear resolution of both Asian and European clines (**Supplementary Dataset 16j,k**, **Supplementary Table 4**).

Applying our multi-layered ranking scheme (**Extended Data Fig. 7**) – combining IBD-based clustering information with the Hamming-Spearman objective – produced results (**Supplementary Dataset 17**) that closely match those obtained when optimizing for neighborhood preservation (**Supplementary Dataset 16j,k**). The resulting 3D LDEs are broadly elliptical and closely resemble the top-scoring embeddings obtained for the ancient Eurasian dataset (**Supplementary Dataset 12**). The twenty highest-ranked LDEs (including the top five shown in **Supplementary Dataset 17a–e**) occupy a narrow region of parameter space: all but one are based on PC scores without drift-normalization; all rely on Euclidean distances in PC space; PC-space dimensionality ranges from 14 to 26; and *k*-NN ranges from 20 to 40 (**Table 2**, **Supplementary Table 4**). The final output of this optimization run (**Supplementary Dataset 17a**) is a 3D PHATE embedding using 32 nearest neighbors, computed with Euclidean distances in a 22-dimensional classic centered PC space. As in the ancient Eurasian optimization run above, this embedding can be interpreted as a denoised version of the corresponding PC1–PC2–PC3 space (**Supplementary Dataset 11e**). It increases the median Mahalanobis separation among the 41 largest IBD-sharing communities (**Supplementary Table 2**) from 7.45 standard deviations (best PC1–PC2–PC3 space) to 12.03 (best PHATE LDE). The gain is even more pronounced for the two reference IBD communities (Slavic–Baltic vs. Celtic–Germanic), whose Mahalanobis separation rises from 1.53 to 10.02 standard deviations (**Supplementary Table 4**). As in the ancient Eurasian dataset, the classic PC1–PC2– PC3 spaces derived from the present-day dataset fit all genetic distance/dissimilarity measures and great-circle geographic distances substantially better than the eigenvector space (**Supplementary Tables 1 & 4**), likely because of their more favorable aspect ratios (**Supplementary Fig. 21a**, **Supplementary Dataset 11**).

These improvements reflect, among other LDE features, the emergence of two major clines in Europe and West Asia, mirroring those seen in the best-scoring PHATE LDE on ancient Eurasian data (**Supplementary Dataset 12a**): a WE–WM cline and an EE–EM cline. Germanic-, Romance-, and Celtic-speaking populations map primarily to the former, whereas Slavic- and Baltic-speaking populations map to the latter. This represents a substantial improvement over the PC1–PC2–PC3 space (**Supplementary Dataset 11d–f**), where present-day European structure is captured mainly along a south–north axis, with very limited east–west separation.

## Discussion

Our results show that the familiar PC1–PC2 or PC1–PC2–PC3 plot is not a neutral summary of population-genetic data. Under favorable conditions – dense, uniform sampling of a simple isolation-by-distance landscape and retention of all polymorphic variants – PCA reproduced simulated geography almost perfectly. Once sampling became sparse or clustered, however, the same landscapes generated markedly different visualizations, and rare-variant removal or LD pruning generally amplified the distortion. Although the most stringent MAF filter sometimes increased global geography–PCA correlations, it did not restore local structure and often mixed neighboring samples. Complex spatial distributions collapsed into triangles, tetrahedra, or three-ray patterns; genuine gradients were exaggerated in some regions and compressed in others; and marginal samples became artificial outliers or shifted toward intersections of clines. Individuals in central positions could therefore be misinterpreted as minimally drifted or highly admixed even when their placement resulted mainly from sampling gaps and subsample-driven SNP ascertainment. Similar visualization patterns arose from spatial landscapes, trees, and admixture-graph-shaped histories, showing that the apparent topology of a PCA plot does not uniquely identify the demographic process that generated it.

Many of these failures can be understood as consequences of a high-dimensional, low-information regime. Sparse sampling removes the configurations needed to resolve local structure, whereas rare-variant depletion removes variables that connect nearby points on the genetic landscape. As these signals disappear, sample geometry approaches that of nearly equidistant points, and low-dimensional projections increasingly reflect simplex-like geometry and, potentially, eigenvector localization rather than the original landscape. This provides a plausible explanation for recurrent triangular and three-ray patterns in the simulations and in many published datasets. It does not imply that every simplex-like pattern is artifactual: divergence and mixture among genuinely discrete populations can produce related geometry. Rather, the visual pattern alone cannot distinguish these alternatives. A rigorous derivation of the transition to three-ray and four-ray configurations remains an important theoretical problem.

These findings are particularly relevant to archaeogenetics, where sparse and often biased sampling is the norm rather than the exception. Singletons and other rare variants in archaeogenetic studies are often considered particularly error-prone and are therefore removed or subjected to additional filtering before downstream analyses (Lindo et al. 2018; Barlow et al. 2020; Martiniano et al. 2022). Many studies also rely on SNP-enrichment panels (Rohland et al. 2022) dominated by common variants (Flegontov et al. 2023), while PCA is commonly performed on LD-pruned datasets (Crosslin et al. 2014; Privé et al. 2020; Grinde et al. 2024), following established recommendations (Patterson et al. 2006). The effects identified here may therefore distort practical reconstructions of past and present-day genetic landscapes. Restricting PCA to common variants can improve population stratification when the relevant structure consists of discrete populations (Ma & Shi 2020). When variation is spatially continuous, however, maximizing separation among sampled groups can conflict with reconstructing the underlying landscape. Rare variants are particularly informative about local structure and recent shared history (Mathieson & McVean 2012), and filtering should therefore be chosen according to the inferential goal rather than applied automatically.

A second general result concerns the representation used before nonlinear dimensionality reduction. The eigenvectors *U* reported by *smartPCA* and *PLINK* are not geometrically equivalent to conventional PC scores *UΣ*. This distinction is often (but not always) modest in two dimensions but becomes consequential when tens or thousands of PCs are supplied to manifold-learning algorithms. Full-dimensional centered classic PC scores preserve the Euclidean geometry of the centered genotype matrix, whereas eigenvectors rescale successive axes by their singular values and produce a whitened, Mahalanobis-like geometry. Accordingly, high-dimensional classic PC spaces usually represented the tested genetic relationships better and provided substantially better inputs for optimized manifold learning. The optimal input dimensionality for manifold learning was often far higher than suggested by a conventional scree-plot elbow and, in both simulated and empirical datasets, frequently approached the full sample rank. Choosing a small number of PCs because they explain most variance is therefore not generally appropriate when the goal is an informative nonlinear embedding.

The spatial simulation study further demonstrates why no universal manifold-learning protocol should be expected for population-genetic data. As expected, each second-level objective improved preservation of the relationship used to rank the embeddings, but these gains did not necessarily transfer to other relationships. Embeddings selected using *F_ST_*, *f*_2_-statistics, or Hamming distance could fit the target metric well while weakening correspondence with geography. Rank correlation with GRM-derived dissimilarity provided the most consistent compromise between genetic and geographic fidelity for high-quality, diploid datasets with intracontinental-scale genetic differentiation. Even then, optima depended on sampling and filtering: UMAP on approximately full-dimensional classic PC spaces was favored in several unfiltered settings, whereas PHATE frequently emerged under clustered sampling and rare-variant depletion. In the simulations, the common population-genetic workflow – UMAP with Euclidean distance on a small number of *smartPCA* eigenvectors – appeared only among a few secondary optima under random landscape sampling and was generally suboptimal. A closely related pattern – a preference for classic PC-score inputs over low-dimensional *smartPCA* eigenvectors – was observed in five of the six empirical case studies. The Southeast Asian analysis was an informative exception, illustrating how preprocessing constraints may reverse this general pattern. The main conclusion is therefore not that one algorithm replaces PCA visualization of population-genetic data, but that input representation, dimensionality, distance metric, manifold-learning algorithm, neighborhood size, and ranking objective must be optimized jointly.

The empirical-data analyses support this conclusion while revealing its limitations. In the Orang Asli present-day human dataset, PHATE unfolded a largely one-dimensional PCA configuration into geographically interpretable gradients, resolved additional structure, and placed individual outliers in positions concordant with an independent IBD-sharing graph. In present-day red deer, an optimized UMAP–densMAP intermediate resolved genetic gradients within Europe that were compressed in the simplicial PCA representation. In the Papua New Guinea present-day human dataset, optimized densMAP integrated highlander and lowlander variation into one representation and revealed additional structure among neighboring highland provinces. In the Southeast Asian present-day human dataset, where the leading PCs were dominated by highly drifted small ethnolinguistic groups and compressed major populations into a poorly resolved central cloud, optimized PHATE recovered a prominent Kra-Dai cline and finer relationships among regional groups. Most notably, it separated the Indian-admixed Mon, Burmese, Central Thai, Southern Thai, and Malay populations into a distinct pole in a fully unsupervised analysis lacking South Asian reference groups. Yet the red deer, Papua New Guinea, and Southeast Asian embeddings optimized for GRM-derived relationships showed poorer fits to some other genetic metrics than the corresponding PCA embeddings. The unusual Southeast Asian optimum – a PHATE embedding based on seven drift-normalized eigenvectors – may partly reflect the aggressive filtering of missing data before calculating classic PC scores, together with heterogeneous missingness across the source datasets. Manifold-learning optimization can therefore expose structure hidden by the leading PCs, but it cannot eliminate differences among alternative measures of relatedness or make one representation optimal for every biological question.

The ancient Eurasian application provides the clearest demonstration of the historical utility of our approach. Conventional three-dimensional PCA, with ancient individuals projected onto a present-day Eurasian reference panel, weakly separated the Central–East Europe (CEE) IBD-sharing community, which contains nearly all Medieval individuals from Slavic cultural contexts, from the Northwest Europe (NWE) community, whose members were overwhelmingly buried in Germanic contexts, and from other North and Central European IBD communities. A layered search across approximately 809,000 candidate embeddings selected a PHATE representation based on 44 drift-normalized classic PCs, Manhattan distance, and 26 nearest neighbors. It retained essentially the same aggregate fit to Hamming distances as PC1–PC2–PC3 space while increasing median Mahalanobis separation among the 23 largest IBD communities from 3.8 to 6.1 standard deviations and separation of the two focal IBD communities (Central–East European and Northwest European) from approximately 1.1 to 4.1 standard deviations. IBD-community separation was incorporated into the layered ranking procedure, whereas Y-chromosome lineages and one-source *qpAdm* results provided complementary assessments of the selected embedding; similar European and Asian clines also recurred under alternative ranking objectives.

The earliest members of the CEE IBD-sharing community, dated to approximately 150–550 CE and distributed from the Middle Danube to the Dniester and Don estuaries and the Middle Volga, appeared as genetic outliers relative to the local populations in each region. Despite their wide geographic distribution, they formed a compact cluster in the 3D PHATE space but not in PC1–PC2–PC3 space. This generated a historically consequential hypothesis about the early dispersal of ancestry shared by later Slavic speakers and illustrates how optimized embeddings can guide further testing with IBD, *qpAdm*, *f*-statistics, Y-chromosome phylogenies, and archaeology. Manifold-learning optimization of the present-day Eurasian reference set recovered the same West Europe–West Mediterranean and East Europe–East Mediterranean clines and increased the Mahalanobis separation between the Slavic–Baltic and Celtic–Germanic IBD communities from 1.5 to 10.0 standard deviations, further supporting the robustness of the broad population structure revealed in the ancient dataset.

Neither PCA nor manifold learning is a demographic model, and no two- or three-dimensional representation should be interpreted as a unique reconstruction of population history. Global distance preservation, local-neighborhood preservation, density preservation, cluster separation, geographic correspondence, and affinity to independently derived genetic relationships are distinct goals that can favor different embeddings. The strongest results here therefore come not from a single numerical optimum but from structures that persist across broad regions of parameter space and are supported by complementary analyses. For example, in the ancient Eurasian case study we compared fundamentally different LDE-ranking strategies, including agreement with Hamming and GRM-derived dissimilarities, separation of IBD-sharing communities, and preservation of high-dimensional sample neighborhoods. The final PHATE embedding was then assessed from complementary perspectives – including separation of clusters defined by Y-chromosome lineages and agreement with one-source *qpAdm* p-values – while the major European and Asian clines consistently re-emerged under alternative LDE-ranking objectives, supporting the robustness of the result. Importantly, the selected PHATE embedding did not outperform the corresponding PC1–PC2–PC3 space on the tested metrics assessing preservation of full-dimensional PC-space’s geometry; its advantage was concentrated in genetically defined structure. This reinforces that the preferred embedding depends on the evaluation criterion rather than being universally superior.

PHATE, which emerged in our study as a particularly attractive manifold-learning method for empirical population-genetic datasets, has previously been illustrated on human SNP data (Moon et al. 2019) and applied to present-day population-genomic datasets from maize (Wisser et al. 2019), dogs (Dutrow et al. 2022), grapevine (Liu et al. 2024), horses (Li et al. 2025), and sheep (Bionda et al. 2026), but always with default or manually selected input-space dimensionalities and hyperparameters, including elbow-rule-based dimensionality selection in Dutrow et al. (2022). Earlier population-genetic studies have also applied locally linear embedding, t-SNE, UMAP, variational autoencoders, and other nonlinear methods (e.g., Siu et al. 2012; Sakaue et al. 2020; Battey et al. 2021), and limited optimization of input dimensionality and UMAP hyperparameters has been explored (Diaz-Papkovich et al. 2019, 2026). An objective-guided framework has also been developed for comparing dimensionality-reduction algorithms against pedigree and IBD relationships (Ubbens et al. 2022). However, to our knowledge, no previous population-genetic study has jointly optimized genotype preprocessing, PCA representation and dimensionality, input-space distance metric, manifold-learning algorithm, neighborhood scale, and multiple population-genetic ranking objectives.

We also found no optimization effort of comparable scope in the single-cell transcriptomics literature, where manifold-learning optimization and validation are considerably more widespread. The scDEED method (Xia et al. 2024) illustrates some important limitations of current practice. Xia et al. developed a valuable statistical framework for detecting dubious embeddings of single-cell transcriptomic data and optimizing UMAP or t-SNE parameters. The framework is generalizable to other data types and manifold-learning algorithms and evaluates embeddings by comparing sample neighborhoods in low- and high-dimensional spaces. However, optimization is conditional on an input PC space of fixed dimensionality (12 to 50), either adopted from the original empirical studies or selected using a scree-plot elbow; a single fixed dimensionality of 12 was likewise used for simulated data (Xia et al. 2024). The Cattell elbow criterion (Cattell 1966; Peres-Neto et al. 2005) was developed as a heuristic for separating dominant linear factors from the residual eigenvalue tail and has no general theoretical connection to the PC dimensionality that best preserves neighborhoods under nonlinear embedding: we showed that it may underselect, overselect, or fall between distinct optima. Moreover, because scDEED evaluates each embedding against the same fixed PC space used to define the pre-embedding relationships, information discarded by PC truncation cannot be recognized as lost. Joint optimization of input dimensionality, neighborhood scale, and other embedding parameters – preferably evaluated against a common external or held-out target – is therefore necessary for comprehensive manifold-learning optimization.

Optimized manifold learning is best regarded as a hypothesis-generating and integrative framework: it reveals gradients, outliers, and candidate historical connections that are difficult to recognize in PCA and organizes them for testing with haplotype-based, allele-frequency-based, uniparental, geographic, or archaeological evidence. Future work should make this strategy more reliable and efficient. The conditions under which sampling and ascertainment drive a genetic landscape toward simplex-like PCA geometry should be formalized and tested across a broader range of spatial, temporal, and graph-shaped population histories. Exhaustive searches over hundreds of thousands of embeddings could be replaced by adaptive, multi-objective optimization that reports the stability of features across stochastic seeds, genomic resampling, and nearby parameter settings rather than only a single optimum. Ancient-DNA applications require objectives robust to pseudo-haploidy, nonrandom missingness, sequencing and capture heterogeneity, imputation uncertainty, and SNP ascertainment. The present results suggest that integrating complementary information – genotype-based distances, high-dimensional neighborhood preservation, and independently inferred IBD communities – will be more productive than seeking a universal distance measure or algorithm. Broader benchmarks involving simulated continuous-space histories, range expansions, temporal sampling, barriers, and long-distance migrations should clarify which aspects of population history are recoverable in low dimensions and when no trustworthy embedding can be produced.

PCA should remain an important baseline, but its first two or three axes should not be treated as a self-interpreting portrait of population history. Analysts should examine how sampling and SNP ascertainment shape the plot, distinguish eigenvectors from classic PC scores, retain sufficient higher-dimensional information for downstream methods, compare multiple embedding-selection objectives, and validate important structures independently. Our simulations and empirical applications show that biologically and historically meaningful information can remain hidden beyond the leading PCs. Objective-guided manifold learning can recover part of that information and convert it into testable hypotheses, provided that the resulting embedding is evaluated as critically as the PCA visualization it is intended to improve.

## Methods

### 1. Simulated stepping-stone landscapes

The stepping-stone simulations were performed using the *slendr* R package v. 0.8.0 (Petr et al. 2023) with the *msprime* v. 1.2.0 backend. We simulated hexagon-shaped SSLs based on a triangular lattice (similar to those introduced by Flegontova et al. 2025) and composed of 331 panmictic demes of constant effective population size (1,000 diploid individuals). The demes arose via multifurcation (“star radiation”) at generation 470 (counted forward from the past). Beginning at generation 471, three consecutive gene-flow epochs were simulated: pre-LGM, lasting 1,400 generations; LGM, lasting 350 generations; and post-LGM, lasting 770 generations (LGM = Last Glacial Maximum). Each pair of neighboring demes was connected by two gene flows in opposite directions. The following features differed among the IBD, IBR, and IBR-LDM landscapes (**Extended Data Fig. 1**):

- **IBD:** In each epoch – pre-LGM, LGM, and post-LGM – the per-generation gene-flow intensity (identical in both directions and uniform across the landscape) was sampled randomly from a normal distribution centered at 0 with a standard deviation of 0.3. In the LGM epoch, the normal distribution differed slightly: the standard deviation was 0.1 rather than 0.3. The following epoch-specific brackets for gene-flow intensity were applied during random sampling from the normal distributions: pre-LGM, 2.1×10^-5^ to 3.6×10^-3^; LGM, 2.9×10^-6^ to 3.6×10^-3^; post-LGM, 1.9×10^-5^ to 6.5×10^-3^. These values represent the proportion of individuals in a given generation originating from a neighboring deme.
- **IBR:** For each unidirectional gene-flow edge, per-generation gene-flow intensity was sampled independently at random from the normal distributions and brackets described above. This sampling procedure was repeated at the start of each gene-flow epoch – pre-LGM, LGM, and post-LGM (**Extended Data Fig. 1**). Genetic distances (*F_ST_*) among the simulated demes are comparable to intracontinental-scale differentiation in humans, such as that observed within the Americas, Eurasia, and Africa (**Supplementary Fig. 14b**).
- **IBR-LDM:** Long-distance bidirectional gene flows bypassing nearest neighbors were superimposed on the IBR landscape as follows: 300 pairs of non-neighboring demes were selected at random from all possible non-neighbor pairs, and the resulting gene-flow events were distributed randomly across the 2,520 generations of the simulation, each persisting for 10 generations. Per-generation migration rates were assigned independently for the two directions and, with equal probability, were set to 0.05 or 0.1. Genetic distances (*F_ST_*) among the simulated demes are comparable to intracontinental-scale differentiation in humans (**Supplementary Fig. 14a**).

For each of the three landscape types, we generated ten independent simulation replicates, each with a new random resampling of gene-flow intensities. Diploid individuals were sampled from each deme in the post-LGM epoch: three individuals at 300 generations before the end of the simulation and three again at the end of the simulation, yielding 993 individuals per sampling point and 1,986 individuals per simulation replicate. Thus, the densest and most uniform sampling scheme in our study represents 0.003 of the combined effective population size of 331,000 at each time point. Because of the substantial memory requirements of the simulations, genome length was limited to 200 Mbp. The recombination rate was set to 1×10^-8^ per nucleotide per generation, and the mutation rate to 1.25×10^-8^ per nucleotide per generation.

### 2. Subsampling simulated individuals and SNP filtering

The two sampling points (300 generations before the end of the simulation and at the end of the simulation) were independently subjected to subsampling and SNP filtering:

- From the 993 individuals available at each time point, either 200 (∼0.2) or 50 (∼0.05) were sampled at random.
- Alternatively, 200 or 50 individuals were sampled in a clustered manner. In this approach, either 15 or 5 demes were selected at random as “seeds,” all immediate neighbors of the seed demes were then added, and 200 or 50 individuals, respectively, were sampled randomly from the resulting set of demes.

Five subsampling replicates were generated for each simulation replicate and for each sampling regime. Each SNP–individual dataset was filtered to remove non-polymorphic sites (MAC = 1) and multiallelic variants. In addition, each subsampled dataset was optionally subjected to rare-variant removal at various MAF thresholds (**Fig. 2**) or to LD-based SNP pruning, performed in *PLINK* v. 2 with the *--indep-pairwise* option, a 200 kbp window, and an *r*² threshold of 0.5 (or 0.3 for the 993-individual samples, to keep the total number of SNPs at a manageable level). Counts of polymorphic SNP loci in the simulated subsampled datasets are shown in **Supplementary Fig. 1**.

### 3. PCA on simulated and empirical data

For every dataset containing *s* individuals, we computed *s* − 1 PC axes using either *PLINK* v. 2 (Chang et al. 2015), which implements the *smartPCA* algorithm of Patterson et al. (2006) and outputs eigenvectors, or the *smartSNP* R package v. 1.1.0 (default settings: *missing_impute = "mean"* and *program_svd = "RSpectra"*; Herrando-Pérez et al. 2021), which outputs conventional PC scores. *PLINK* v. 2 applies a single data normalization procedure before PCA: centering followed by scaling that accounts for the expected dispersion of allele frequencies due to genetic drift, proportional to 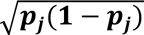 (Patterson et al. 2006). We refer to this procedure as normalization by genetic-drift dispersion, or simply drift normalization. By contrast, *smartSNP* offers several normalization schemes (Herrando-Pérez et al. 2021), which we tested here: centering alone, centering followed by standard-deviation normalization, and centering followed by normalization by genetic-drift dispersion.

### 4. Manifold-learning optimization on simulated and empirical data

Manifold-learning optimization was implemented in R using the *uwot* package v. 0.2.4 for UMAP and densMAP (https://github.com/jlmelville/uwot) and *phateR* v. 1.0.7 for PHATE (https://github.com/KrishnaswamyLab/phateR). For each precomputed PC representation, we varied the number of input PCs, the *k*-nearest-neighbor parameter and the input-space distance metric, while setting the output dimensionality to 3 or 2. UMAP was run with Euclidean, Manhattan, cosine or correlation distance and with *min_dist = 0.1* or *0.5*. We additionally tested two density-aware variants at *min_dist = 0.1*: an intermediate UMAP– densMAP objective (*dens_scale = 0.5*) and densMAP (*dens_scale = 1*), both using *n_trees = 100* and *n_epochs = 1000*; individual embeddings were run with one thread within an outer multicore loop. PHATE was run with Euclidean, cosine or city-block (Manhattan) distances, the SMACOF MDS solver (*mds.solver = "smacof"*), and multithreading. For both *uwot* and *phateR*, the same settings were applied to simulated and empirical data, and other algorithm parameters were left at package defaults.

Each candidate low-dimensional embedding was evaluated from all pairwise Euclidean distances in the output space using squared Pearson and Spearman correlations with the corresponding genetic and geographic pairwise measures. Geographic distances were Euclidean for simulated landscapes and great-circle distances for empirical data. Genetic criteria included Hamming/IBS distance, calculated using *PLINK* v. 1.9; GRM- and covariance-derived dissimilarities, calculated using *PLINK* v. 1.9; individual-level Hudson’s *F_ST_*, calculated using *PLINK* v. 2; individual-level *f*_2_-statistics, calculated using *ADMIXTOOLS 2* (*maxmiss=1*, *adjust_pseudohaploid=FALSE*); and, where available, outgroup *f*_3_-statistics, also calculated using *ADMIXTOOLS 2* (*maxmiss=1*, *adjust_pseudohaploid=FALSE*). Negative *F_ST_*, *f*_2_, and outgroup *f*_3_ estimates were considered artefactual and truncated to zero; *f*-statistic Z-scores were not considered. The same statistics were calculated for the corresponding reference PCA embeddings. When LDEs were ranked by their fit to genetic or geographic distances, LDE or PC spaces were always matched to genetic-distance matrices by both individual set (defined, for example, by the simulation and subsampling replicate), and SNP set, defined by the MAF-filtering and missing-data removal level. Scripts used for objective-guided optimization, visualization, and validation of its outputs are available at https://github.com/flegontovlab/optimized_manifold_learning.

### 5. Manifold-learning optimization and other methods for the Eurasian case studies

#### 5.1. Dataset composition and principal component analysis

To reduce batch effects introduced by jointly analyzing data produced with different aDNA sequencing strategies (primarily shotgun sequencing and the 1240K and Twist enrichment panels; see Rohland et al. 2022) and noise due to low coverage in some samples, autosomal SNP data for 5,133 ancient Eurasian individuals were projected onto PC axes computed from 2,425 present-day Eurasian individuals (**Supplementary Table 2**). Sources of the published ancient and present-day data are listed in **Supplementary Table 2**; the unpublished ancient data will be released with Vyazov et al. (in preparation).

Because our analysis focuses on the Slavic dispersal and on Iron Age and Medieval Europe more broadly, we excluded large portions of Eurasia (the Arabian Peninsula; South, Southeast, and East Asia) from the present-day sample set. Present-day individuals were selected to provide continuous coverage of the remaining regions, with only a few geographic outliers originating from the excluded areas (**Supplementary Fig. 20b**). Ancient individuals were chosen mainly according to the focal time range (125 to ∼3,200 yBP), with the Arabian Peninsula, the Iranian Plateau, South and Southeast Asia largely omitted. This yielded geographic coverage forming a diagonal transect across Eurasia, from Iberia and Iceland to Taiwan and the Japanese Archipelago (**Supplementary Fig. 20a**). For ancient individuals, genotypes were restricted to the Human Origins panel (Patterson et al. 2012), matching the genotyping platform used for the present-day cohort. Across ancient individuals, the number of autosomal SNPs polymorphic in the dataset ranged from 559,141 to 19,434 (559,163 loci in total), with a median of 373,443 SNPs. For present-day individuals, corresponding counts ranged from 526,682 to 490,025 (526,824 loci in total), with a median of 521,822 SNPs.

PC axes for present-day individuals and least-squares projections for ancient individuals were computed using two alternative approaches: *smartPCA* v.16000 (settings: *numoutlieriter: 0, lsqproject: YES, autoshrink: YES*; Patterson et al. 2006; Liu et al. 2017) and the *smartSNP* R package v.1.1.0 (default settings: *missing_impute = "mean"* and *program_svd = "RSpectra"*; Herrando-Pérez et al. 2021). *smartPCA* outputs eigenvectors, whereas *smartSNP* returns conventional PC scores. The alternative data-normalization approaches were identical to those used in the other case studies. For runtime reasons, the *smartPCA* analysis was limited to 200 PCs, whereas *smartSNP* computed the maximum number of components for both present-day and ancient individuals (*s* – 1 = 2,424, where *s* is the number of basis individuals). Eigenvalues and other component-wise statistics are shown in **Supplementary Fig. 21**.

To avoid shrinkage artefacts in PCA – systematic shifts between basis and projected samples that are not fully mitigated by dedicated algorithms available in the *smartPCA* software (Lee et al. 2010; Liu et al. 2017) – and because our primary interest lies in ancient genetic structure – we visualize the PC spaces (PC1–PC2–PC3) for ancient and present-day individuals separately in most cases, and we also keep them separate for manifold-learning applications (see below). Interactive PC1–PC2–PC3 plots generated for the ancient Eurasians are provided in **Supplementary Dataset 11a–c**; the corresponding plots for the present-day basis individuals are shown in **Supplementary Dataset 11d–e**; and the combined plots of present-day and ancient individuals are presented in **Supplementary Dataset 11g–i**.

#### 5.2. IBD inference and community detection for present-day genetic data

Genotype phasing was performed on the Human Origins autosomal dataset using *Beagle* v.5.5, build 27Feb25.75f (Browning et al. 2021). Phasing was conducted separately for each autosome (chromosomes 1–22) without the use of an external reference panel. To ensure accurate modeling of recombination rates during phasing, we utilized the HapMap Phase II genetic map (GRCh37) (The International HapMap Consortium 2007). After phasing, we detected IBD segments using *Hap-IBD* v.1.0 (build 15Jun23.92f) with default settings (Zhou et al. 2020).

We constructed an IBD-sharing graph among 2,425 present-day individuals based on all autosomal IBD segments longer than 2 cM. Because present-day data are of high quality (diploid, with low missing rates; **Supplementary Table 2**) and *N_e_* is expected to be much larger than in the ancient groups, we did not apply the more stringent 8 cM IBD segment length cut-off used for the ancient data (Ringbauer et al. 2024). For each pair of individuals, we first aggregated all segments into a single non-directional, non-redundant pair by summing the segment lengths in cM, yielding a cumulative IBD length per pair. These cumulative lengths were used as edge weights in an undirected graph with individuals as nodes and one edge per pair with non-zero total IBD. Community detection on this graph was performed using the Leiden algorithm (with the modularity objective) as implemented in the *igraph* v.2.1.4 package in R, across a grid of 14 resolution parameters (from 0.2 to 10) spanning coarse to fine partitions. For each resolution, we recorded the community assignment of every individual, the modularity of the corresponding partition, the total number of communities, and the number of communities containing at least 10 individuals.

After examining several alternative partitions, we selected the lowest resolution level that distinguishes three groups: (1) Slavic and Baltic speakers; (2) Celtic and Germanic speakers; and (3) Finnic speakers and northern Russians, who are known to be substantially admixed with indigenous Finnic-speaking populations (Khrunin et al. 2013; Peltola et al. 2023; Agdzhoyan et al. 2024). Although the ancient and present-day datasets differ in sample sizes (5,133 vs. 2,425 individuals), *N_e_* (smaller vs. larger), and IBD segment-length thresholds (8 vs. 2 cM), this choice of resolution allows for meaningful comparison between them. At the selected resolution level, there are 41 IBD-sharing communities composed of 10 or more individuals (**Supplementary Table 2**).

#### 5.3. Manifold-learning optimization in the ancient Eurasian case study

Our testing on simulated stepping-stone landscapes with “flyover” gene flows and on real-world case studies shows that *ρ*^2^ is by far the best objective for ranking LDEs (as compared to objectives based on Hamming distances, *F_ST_*, *f*_2_- and outgroup *f*_3_-statistics), but if deeply diverged populations and/or pseudo-haploid low-coverage data are co-analyzed 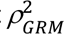 performance becomes worse. Consistent with these findings, Hamming and GRM distances are strongly correlated in our set of 2,425 present-day Eurasian individuals (Pearson *r* = –0.87; Spearman *ρ* = –0.91), whereas the correlations are weaker in the 5,133 ancient Eurasian individuals (*r* = –0.70; *ρ* = –0.75) (**Supplementary Table 3**). Both datasets use Human Origins SNPs, have very similar geographic coverage (**Supplementary Fig. 20**) and similar levels of genetic diversity, but the present-day dataset is technologically homogeneous, diploid, and has a low median missing rate (0.0095; **Supplementary Table 2**). In contrast, the ancient dataset consists of pseudo-haploid data generated using multiple sequencing technologies and exhibits a much higher median missing rate (0.33; **Supplementary Table 2**). These factors collectively reduce the performance of genetic distance metrics. *PLINK* v. 1.9 (Chang et al. 2015) rescales each observed Hamming distance by 1 – (sum of missing variants’ average contribution to distance). If minor-allele frequency is roughly independent of missingness, this adjustment is more accurate than the usual flat scaling by 1 – (missing-call rate).

Given these limitations, for our LDE ranking we prioritized fits to the Hamming distances, computed with *PLINK* v. 1.9, and also incorporated IBD-based clustering information, as detailed in the next **Methods** section 5.4. Another important result informing our decision to de-prioritize the GRM distance comes from a downstream analysis described in **Methods** section 5.5. We evaluated correlations between LDE fidelity metrics indirectly, using their correlations with Euclidean distances in LDE space across a large set of LDEs for the ancient individuals (**Supplementary Table 5a**), and focused on a key measure of how well LDEs preserve the geometry of the (*s – 1*)-dimensional PC space – namely, preservation of *k*-sized sample neighborhoods. This metric (at *k* = 500) correlates strongly with several genetic LDE-fidelity metrics (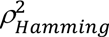 and IBD-based ones; *r* = 0.65–0.79; *ρ* = 0.70–0.89) but shows essentially no correlation with 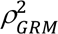 (*r* = 0.29; *ρ* = 0; **Supplementary Table 5a**).

The optimization landscape explored in this study is presented in **Supplementary Dataset 18**, which reports the fits of all 3D LDEs to genetic and geographic distances in the form of differences in squared correlation relative to the baseline PC1–PC2–PC3 space (drift-normalized classic PC space): 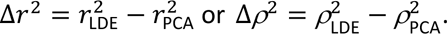 Using either genetic distance metric (Hamming or GRM) and considering both Pearson’s and Spearman’s correlations, PC scores serve as substantially better manifold-learning inputs than eigenvectors (**Supplementary Dataset 18**). Among embedding methods, PHATE (Moon et al. 2019) achieves the strongest fits for both genetic distance metrics, followed by densMAP (Narayan et al. 2021), then a loss function midway between UMAP and densMAP (*dens_scale = 0.5*), and finally standard UMAP (McInnes et al. 2018). Optimizations using Euclidean (*ℓ_2_* norm) and Manhattan (*ℓ_1_* norm) distances in the input PC spaces perform similarly, whereas cosine and correlation affinities are less effective for the ancient Eurasian dataset subjected to the Human Origins ascertainment (**Supplementary Dataset 18**).

The two-dimensional space defined by input PC space dimensionality and *k*-NN shows intricate, highly method-dependent patterns across PCA variants, manifold-learning algorithms, and distance metrics. In particular, PHATE paired with Manhattan or cosine distances yields especially complex, non-smooth fit landscapes (**Supplementary Dataset 18**). This complexity is to be expected. PHATE embeds data by matching low-dimensional distances to diffusion-potential distances derived from a *k*-NN graph and a Markov diffusion process. The associated objective is a nonconvex stress function, and its targets depend nonlinearly on kernel bandwidths, neighborhood size, diffusion time, and a logarithmic transform. Small tweaks to these settings – or to the input representation – can discretely change the graph and transition probabilities, producing a non-smooth loss surface with many local optima. As a result, the optimization landscape over PC space dimensionality, *k*-NN, and distance or affinity metrics is particularly intricate for PHATE. This issue is compounded by our parameter grid (**Supplementary Datasets 18 & 19**): computational limits (and PHATE is by far the slowest algorithm in our panel) prevent us from evaluating every *k*-NN and input PC dimensionality, although for each manifold-learning method we evaluated 151 *k*-NN values (3–5,100 for 5,133 individuals), and 118 PC-space dimensionalities (from 3 up to the maximum of 2,424). Notably, PHATE (and, to a lesser extent, UMAP and densMAP) shows different behavior for PC dimensions <100 versus >100, and at this boundary our step sizes for both *k*-NN and dimensionality increase from 2 to 5 (compare the same optimization landscapes shown on log-scaled axes in **Supplementary Dataset 18** with those on linear axes in **Supplementary Dataset 19**).

#### 5.4. A fine-scale embedding ranking method leveraging IBD-sharing communities

Across both objective functions– 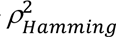 and 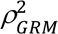 – the most optimal approach is PHATE applied to classic PC scores using either Manhattan or Euclidean (but not cosine) distances, with densMAP applied to the same kind of PC spaces as the consistent runner-up (**Supplementary Datasets 18 & 19**). Since very different sample configurations can yield similar *ρ*^2^ values – and because Hamming and GRM distances each have distinct shortcomings in the ancient Eurasian dataset (see **Methods** section 5.3) – it is essential to evaluate the embeddings carefully. Reviewing LDEs from the global 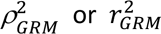 optima (PC space dimensionality >20, *k*-NN between ∼500 and ∼2000; **Supplementary Dataset 18a**) indicates that while many European groups are well resolved – and individuals of predominantly European ancestry make up ∼70% of the dataset – the Asian steppe clines remain poorly resolved in all such embeddings (see an LDEs maximizing 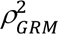 and an LDEs maximizing 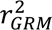 as interactive 3D plots in **Supplementary Dataset 13a,b** and their LDE fidelity metrics in **Supplementary Table 4**). In contrast, LDEs from the global 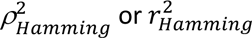 optima (PC space dimensionality close to *s* – 1, *k*-NN from ∼20 to ∼100; see **Supplementary Dataset 18b**) demonstrate good resolution both within the European and Asian parts of the genetic landscape (see an LDEs maximizing 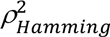 and an LDEs maximizing 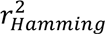 as interactive 3D plots in **Supplementary Dataset 13c,d** and their fidelity metrics in **Supplementary Table 4**).

However, given that Hamming distance is a coarse metric for capturing fine-scale genetic structure, we chose to incorporate IBD-sharing information into the LDE ranking procedure by using IBD-sharing communities (clusters) identified in the IBD graph with the Leiden algorithm (**Extended Data Fig. 6a**). We focused on the 23 largest communities (composed of 20 or more individuals; **Extended Data Fig. 6a**) and, for each PHATE embedding derived from classic PC spaces (the broad optimum in our parameter space), computed median Mahalanobis and Bhattacharyya inter-cluster distances across all community pairs. As a control, we also computed these distances for two benchmark communities that are notoriously difficult to separate with PCA: CEE (primarily Slavic) and NWE (primarily Germanic). Mahalanobis distance was computed between the two cluster centroids using the average covariance Σ_pool_ = (*S*_1_ + *S*_2_)/2, a common, stable choice when covariances differ. Bhattacharyya distance uses Σ = (*S*_1_ + *S*_2_)/2 in the standard closed form:

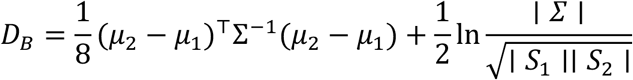

The code (see https://github.com/flegontovlab/optimized_manifold_learning) also adds a tiny ridge (10^-6^) to covariances if needed and falls back to a pseudo-inverse for numerical robustness.

These standard IBD cluster-resolution metrics, as well as 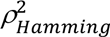 and 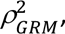 each reach their optimum in different regions of the PHATE parameter space (**Supplementary Dataset 14a,b**, **Supplementary Table 4**). Because ranking solely by median inter-cluster distances can encourage over-clustering (and thus weaken clines) – a common issue with default manifold-learning settings – we selected median Mahalanobis distance (compared with the Bhattacharyya distance, this simpler metric prioritizes centroid separation rather than differences in cluster shape), screened embeddings to form a broad candidate set, and then ranked those candidates by the 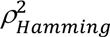 which we prioritized over 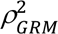 (**Extended Data Fig. 7**). For comparison, see a PHATE LDE maximizing median Mahalanobis distance among the 23 largest IBD-sharing communities as an interactive 3D plot in **Supplementary Dataset 13e**, and its LDE fidelity metrics in **Supplementary Table 4**.

Accordingly, the final embedding was chosen via the following multi-step procedure (**Extended Data Fig. 7**):

- Inspect the full parameter space (809,286 LDEs) and identify the optimal manifold-learning and PCA algorithm choices – by consensus across the two objective functions 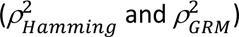 – namely, PHATE and classic PC scores;
- From 85,188 PHATE embeddings on classic PC scores, select the top 5% (4,259 LDEs) by median Mahalanobis distance among the 23 largest IBD-sharing communities;
- From that subset, keep the top 20 by 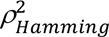 (see **Supplementary Table 4** for details on these top LDEs);
- Within that narrow subset, retain embeddings based on the PCA approach better suited to genetic data (Patterson et al. 2006; data centered and normalized by genetic-drift dispersion) and prefer the larger *k*-NN to better preserve clines, yielding a single best embedding. See three LDEs constructed using the largest *k*-NN values in **Supplementary Dataset 12a–c**.

To reiterate, the selected embedding (visualized as an interactive 3D plot in **Supplementary Dataset 12a**) was constructed from classic PC scores (based on data normalized by genetic-drift dispersion) as input; the default PHATE algorithm (Moon et al. 2019) used Manhattan distance in PC space, 44 PCs, and *k*-NN = 26. This embedding ranks in the top 2.6% of all PHATE embeddings on classic PC scores by median Mahalanobis separation across the large IBD communities. Considering the benchmark pair of IBD-sharing communities (CEE vs. NWE), this LDE is not unusual: 19.4% of PHATE embeddings on classic PC scores attain equal or higher Mahalanobis separation, and 33% achieve equal or higher Bhattacharyya separation of the benchmark communities. Reassuringly, the three LDEs prioritized at the last step of the ranking procedure (**Supplementary Dataset 12a–c**, **Supplementary Table 4**) are qualitatively similar to the LDE that maximizes 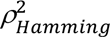 across the entire parameter space (**Supplementary Dataset 13c**, **Supplementary Table 4**) and to the LDE that maximizes median Mahalanobis distance among the 23 largest IBD-sharing communities within the PHATE parameter space (**Supplementary Dataset 13e**, **Supplementary Table 4**). This similarity – evident in both the European and Asian clines – indicates that the more complex procedure does not produce anomalous or erratic results.

The selected 3D PHATE embedding shows only modest Δ*ρ*^2^ relative to the corresponding PC1–PC2–PC3 space (visualized in **Supplementary Dataset 11a,g**): 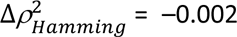 (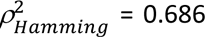 for the 3D PC space) and 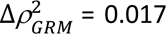 (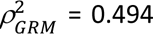 for the 3D PC space). However, median Mahalanobis separation across the 23 largest IBD communities increases markedly – from 3.78 standard deviations for the PC1–PC2–PC3 space to 6.05 for the 3D LDE, moving from the 15.8th to the 97.4th percentile among all PHATE embeddings on classic PC scores. Median Bhattacharyya distance follows the same trend – it increases from 2.48 to 6.07 standard deviations, moving from the 10.3rd to the 77.5th percentile. Notably, the Mahalanobis distance between the CEE and NWE IBD communities also increases – from 1.07 standard deviations for the 3D PC space to 4.11 for the 3D LDE – rising from the 18.3rd to the 80.6th percentile among all PHATE embeddings on classic PC scores (the corresponding Bhattacharyya distance rises from the 19.1st to 67th percentile).

Comparable IBD-sharing communities – consisting predominantly of (1) Slavic and Baltic speakers and (2) Celtic speakers (in their recent past) and Germanic speakers – emerge at an appropriate granularity of the Leiden algorithm in the IBD graph for the present-day basis individuals (**Supplementary Table 2**, **Supplementary Dataset 11d**). A strong signal of shared relatedness between Slavic-speaking and non-Slavic-speaking populations in the Balkans and in Central and Eastern Europe had already been noted in early IBD studies (Hellenthal et al. 2014; Busby et al. 2015). Median Mahalanobis and Bhattacharyya distances among the largest IBD-sharing communities are difficult to compare directly between the ancient and present-day datasets, owing to multiple differences in thresholds and dataset properties (see **Methods** section 5.2), and indeed these distances differ substantially (**Supplementary Table 4**). Nevertheless, it is clear that the separation between the CEE and NWE IBD-sharing communities is also limited in the present-day PC1–PC2–PC3 space, with Mahalanobis distances of 1.35–1.53 and Bhattacharyya distances of 0.41–0.50 standard deviations (**Supplementary Table 4**).

#### 5.5. Quantifying retention of (s – 1)-dimensional PC-space structure

In **Methods** sections 5.3 and 5.4, we evaluated embeddings using correlation to genetic distances and using IBD-based clustering information. As a complementary approach, we assessed how well embeddings preserve the geometry of the ambient input space used by the manifold-learning pipeline. Following Fischer & Ma (2024), we computed four simple metrics that quantify preservation of pairwise distances, *k*-NN sets, local densities, and interpoint angles when mapping from the (*s – 1*)-dimensional PC space to a 3D LDE or PC1– PC2–PC3 space. Let *X* ∈ ℝ^*s*×2424^ denote the high-dimensional representation and *Y* ∈ ℝ^*s*×3^the corresponding 3D embedding.

1. Distance preservation. We assessed global structure by the Spearman rank correlation between all pairwise Euclidean distances in *X* and *Y*. Concretely, we formed the condensed distance vectors d_*X*_ = {∥ *X*_*i*_ − *X*_*j*_ ∥_2_: *i* < *j*} and d_Y_ = {∥ *Y*_*i*_ − *Y*_j_ ∥_2_: *i* < *j*}, and reported *ρ*_*S*_(d_*X*_, d_*Y*_). We approximated this using a uniform random sample of 300,000 unordered pairs.
2. Neighborhood preservation. We quantified local structure with the mean Jaccard index between *k*-NN sets computed in high and low dimensions (using *k* = 500 for the ancient sample set, *k* = 250 for the present-day sample set, which is half the size, and *k* = 50 following Fischer & Ma 2024):

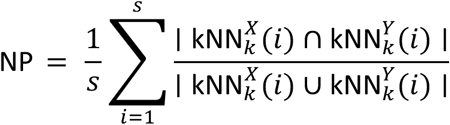 To reduce high-dimensional noise before *k*-NN in *X*, we applied a denoising step different from the ScreeNOT method used by Fischer & Ma (2024) and based on optimal hard SVD thresholding (Gavish–Donoho): after column centering *X*, we kept singular values above the data-driven threshold and reconstructed a denoised matrix, which was then used for *k*-NN calculations.
3. Density preservation. We measured how well relative local densities are preserved. In each space, we first estimated a characteristic radius as the mean distance to the *k*th nearest neighbor across samples (using *k* = 250 for the ancient sample set, *k* = 125 for the present-day sample set, which is half the size, and *k* = 25 following Fischer & Ma 2024), yielding *r*_*X*_ and *r*_F_. For each sample *i*, we counted neighbors within a fixed-radius ball of radius *r*_*X*_ in *X* and *r*_F_ in *Y* (self excluded), producing counts *c*_*X*_(*i*) and *c*_F_(*i*). The density preservation score is the Pearson correlation *r*(*c*_*X*_, *c*_F_) over samples.
4. Angle preservation. To capture preservation of orientations among points, we compared angles subtended at each sample *i* in the two spaces. For computational efficiency, for each *i* we sampled *m* other points uniformly without replacement (*m* = 128) and computed all unordered angles ∠*jik* using normalized dot products in *X* and *Y*. Aggregating over all vertices and sampled pairs, we reported the Pearson correlation between the angle sets in *X* and *Y*. The *R* script performing these four tasks is available at https://github.com/flegontovlab/optimized_manifold_learning.

We assessed Spearman and Pearson correlations of the most important LDE fidelity metrics applied to the PHATE LDEs generated from classic PC scores for the ancient individuals: *ρ*^2^ fits to genetic and/or geographic distances, Mahalanobis and Bhattacharyya distances between IBD-sharing communities, and the four geometry-preservation metrics described in this section (**Supplementary Table 5a**). With *k* = 500 or roughly 1/10 of the sample set (yielding slightly higher correlations as compared to *k* = 50), neighborhood preservation shows strong correlations with median Mahalanobis/Bhattacharyya distances among non-redundant pairs of the 23 IBD clusters (*ρ* = 0.89, *r* = 0.79 for Mahalanobis) and with 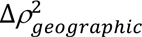 (*ρ* = 0.88, *r* = 0.86). Correlation with 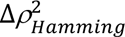 is weaker (*ρ* = 0.70, *r* = 0.65), and correlation with 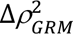 is very poor (*ρ* ≈ 0, *r* = 0.29), suggesting that 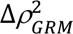 is not an appropriate objective on a technologically heterogeneous, pseudo-haploid, low-coverage dataset. In contrast, the high-dimensional distance-preservation metric is uncorrelated with the genetic/geographic metrics tested (|*ρ*| ≤ 0.25, |*r*| ≤ 0.27); angle preservation is only modestly correlated (|*ρ*| ≤ 0.34, |*r*| ≤ 0.44); and density preservation at *k* = 250 is uncorrelated or even weakly negative (with 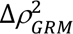: *ρ* = –0.52, *r* = –0.34) (**Supplementary Table 5a**).

The 3D LDE selected in **Methods** section 5.4 does not surpass the drift-normalized classic PC1– PC2–PC3 space on geometry-preservation metrics, though the differences are small for the neighborhood-preservation and distance-preservation metrics: neighborhood (*k* = 500) 0.32→0.30 (78.5th→76.5th percentile among all PHATE embeddings on classic PC scores), distance 0.47→0.45 (46.9th→43.7th), angles 0.44→0.39 (68.2nd→48.1st), and density (*k* = 250) 0.24→0.17 (72.2nd→54.4th). For comparison, we show LDEs that maximize the neighborhood-preservation metric (at *k* = 500 or 50) within the PHATE parameter space (**Supplementary Dataset 13f,g**, **Supplementary Table 4**) and observe that they are qualitatively similar to the former (final) LDE.

#### 5.6. Y-chromosome lineages in the embedding space

To contextualize Y-chromosomal diversity within the embedding space, we first selected a coalescence-time threshold (“coalescence horizon”) on the dated Y-chromosomal tree from the YFull company (see https://github.com/YFullTeam/YTree). Accurate Y-lineage calls were generated manually using our new *callYsnps* tool, as described in Vyazov et al. (in preparation). To limit noise from imprecise lineage calls in low-coverage samples, we excluded individuals below the 25th percentile of SNP counts, retaining 2,197 of 2,930 males from our ancient dataset (**Supplementary Table 2**). We then formed Y-lineage clusters and restricted analyses to clusters containing ≥4 or ≥10 individuals. Clustering quality was evaluated using three standard, complementary indices:

- Silhouette coefficient 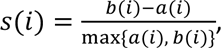 where *a*(*i*) is the mean intra-cluster dissimilarity for point *i* and *b*(*i*) is the minimum mean dissimilarity of *i* to any other cluster; *s*(*i*) ∈ [–1, 1] with higher values indicating more compact and well-separated clusters (Rousseeuw 1987).
- Davies-Bouldin index 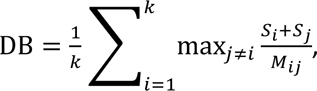 where *S_i_* is the intra-cluster scatter (e.g., average distance to the cluster centroid) and *M_ij_* is the inter-centroid distance; lower values indicate better separation relative to scatter (Davies and Bouldin 1979).
- Caliñski-Harabasz index 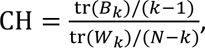 the ratio of between-cluster to within-cluster dispersion; higher values indicate better defined clusters (Caliński and Harabasz 1974).

We evaluated 257 intra-lineage coalescence cut-offs spanning 0–10,000 years ago (ya) (**Supplementary Table 6**). In the highest-scoring 3D LDE space (**Supplementary Dataset 12a**), the Davies-Bouldin index favored 2,600–2,800 ya as the most well-resolved levels, whereas the Caliński-Harabasz index favored 4,600 ya. The Silhouette coefficient identified 0–1,900 ya as optimal (**Supplementary Table 6**). For visualization, we show the 2,800 ya and 4,600 ya cut-offs in the LDE space (on its isometric 2D unfolding; **Supplementary Dataset 20a,b**). Given our focus on Iron Age and later history, we used the 2,800 ya cut-off for primary analyses.

Under the selected settings (2,197 males with high-quality data; ≥10 individuals per Y-lineage; intra-lineage coalescence ∼2,800 ya), we quantified separation of Y-lineage clusters across all PHATE embeddings constructed from classic PC spaces. Following the procedure used for IBD clusters, we computed median Mahalanobis and Bhattacharyya distances for all pairs among these 21 Y-lineages (**Supplementary Dataset 21**, **Supplementary Tables 4 & 5**). The final LDE (**Supplementary Dataset 12a**) ranked above 88.7% of PHATE embeddings on classic PC scores for resolving these Y-chromosomal lineages and outperformed the corresponding drift-normalized PC1–PC2–PC3 space, which ranked at the 36.6th percentile. For comparison, we show a PHATE LDE that maximizes median Mahalanobis distance among the 21 Y-lineages with intra-lineage coalescence around 2,800 ya (**Supplementary Dataset 13h**, **Supplementary Table 4**), which is qualitatively similar to the final LDE.

#### 5.7. External validation of low-dimensional embeddings using qpAdm p-values

To validate the best-scoring 3D LDE in yet another way, we compared inter-cluster Euclidean distances to a proxy of genetic distance derived from one-source “rotating” *qpAdm* tests (Fernandes et al. 2021; Flegontova et al. 2025). In each embedding space – (i) PHATE (3D; **Supplementary Dataset 12a**) and (ii) classic PCA on drift-normalized data (3D; **Supplementary Dataset 11a,g**) – we identified communities via the Leiden algorithm (with the Constant Potts Model objective) on a fully connected, inverse-distance weighted graph, selecting the 30 largest clusters (size ≥15 individuals) per space. For robustness, we generated 10 independent subsamples per space by drawing 10 individuals per cluster without replacement and, for each subsample, computed Euclidean distances between cluster centroids defined on the subsampled individuals. For each space and iteration, we matched unordered cluster pairs to the corresponding one-source *qpAdm* p-values (non-redundant pair set) and quantified association between centroid distance and –log_10_(p) by iteration-wise correlation coefficients (Pearson and Spearman). The *R* script performing these tasks is available at https://github.com/flegontovlab/optimized_manifold_learning.

Across subsampling iterations, distances between cluster centroids in the best-scoring PHATE space exhibited higher correlations with one-source *qpAdm* results (on PHATE-based clusters) than distances between cluster centroids in the drift-normalized classic PC space (on PCA-based clusters). For example, median iteration-wise Spearman correlation coefficients are as follows: *ρ*_PHATE_ = 0.86 vs. *ρ*_PCA_ = 0.82; unpaired Wilcoxon p = 1.8 × 10^-4^.

#### 5.8. Integrative assessment of optimization results for the ancient Eurasian dataset

Because the fidelity of both PCA and PHATE ultimately depends on the present-day basis set (**Supplementary Fig. 20b**, **Supplementary Table 2**), we first applied our panel of LDE fidelity metrics – presented in **Methods** sections 5.3–5.6 and summarized in **Supplementary Table 4** – to PC1–PC2–PC3 spaces for the present-day individuals. Euclidean distances in the basis PC1–PC2–PC3 space (for both centered and centered + drift-normalized data) correlate very strongly with all tested genetic distance metrics – Hamming with different treatments of missing data, GRM, covariance, *F_ST_*, and outgroup *f*_3_(Mbuti; X, Y) – with *ρ* = 0.81–0.88 and *r* = 0.73–0.92 (**Supplementary Table 4**). It is worth noting that these genetic distance metrics are intercorrelated in this dataset (**Supplementary Table 3**). Great-circle geographic distances are also well correlated with LDE distances (*ρ* = 0.78, *r* = 0.80). In contrast, preservation of (*s – 1*)-dimensional sample neighborhoods (range 0–1) is poor at *k* = 250: 0.28 for drift-normalized data and 0.25 for centered data. Distance- and angle-preservation scores relative to the (*s – 1*)-dimensional PC space on centered data are high in the context of the LDEs we examined (0.88 and 0.66, respectively), whereas the same metrics on drift-normalized data perform substantially worse (**Supplementary Table 4**).

Taken together, these LDE fidelity metrics indicate that the first three PCs for the northern Eurasian present-day individuals (**Supplementary Fig. 20b**, **Supplementary Table 2**) capture the underlying “genetic manifolds” well and are not dominated by genetic drift in a few outlying populations. We also see no clear clustering by sequencing technology among ancient individuals and no discernible shift between basis and projected samples (**Supplementary Dataset 11i–h**). While this does not rule out batch effects in higher PCs or guarantee that the chosen basis set is fully optimal, it does not reveal any major issues that would prevent its use as the foundation for our downstream manifold-learning optimization pipeline.

We then return to the optimization results for the manifold-learning methods and reassess them using the same panel of LDE fidelity metrics. Across all evaluated metrics – genetic and geometry preservation – the final LDE outperforms the corresponding PC1–PC2–PC3 space only on genetic metrics: 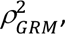 median Mahalanobis and Bhattacharyya distances across the 23 largest IBD-sharing communities and across the 21 Y-lineages, and p-values of one-source *qpAdm* models (**Supplementary Table 4**). Importantly, these genetic metrics are constructed in very different ways, which strengthens confidence in the result’s robustness.

Visual inspection of both the results of the multi-step ranking procedure in **Supplementary Dataset 12a–c** and the LDEs optimized for individual key objectives in **Supplementary Dataset 13a–h** (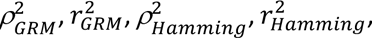 median Mahalanobis distance among the 23 largest IBD-sharing communities, neighborhood-preservation metric at *k* = 500 or *k* = 50, median Mahalanobis distance among the 21 Y-lineages with intra-lineage coalescence around 2,800 ya) further supports that the LDE ranking process is robust to the choice of objective: most of these LDEs display the same clusters and clines across the European and Asian portions of the genetic landscape. For instance, all outcomes produced by the alternative optimization methods (28 PHATE LDEs) show substantially greater separation between the CEE and NWE IBD-sharing communities (Mahalanobis distances ranging from 1.65 to 6.05 standard deviations; median 3.01) compared to the three PC1–PC2–PC3 spaces, which yield distances of only 1.03–1.07 standard deviations (**Supplementary Table 4**). The most divergent LDEs arise when ranking solely by 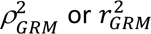 (**Supplementary Dataset 13a,b**), which are weak objectives that are largely uncorrelated with most other genetic and geometric criteria (**Supplementary Table 5a**), and the second most divergent LDE occurs when ranking solely by median Mahalanobis distance among the 21 Y-lineages (**Supplementary Dataset 13h**), an objective that emphasizes genetic information from a single locus at a specific coalescent time horizon and is therefore expected to be biased.

#### 5.9. From a 3D embedding to its approximately isometric 2D unfolding

Many PHATE-derived 3D LDEs exhibited an approximately ellipsoidal geometry (**Supplementary Datasets 12 & 13**), so we developed an isometric (geodesic-distance-preserving) unfolding pipeline based on Isomap (Tenenbaum et al. 2000). We first modeled the surface as a radial field *r*(*θ*,*ϕ*) in spherical coordinates using a tensor-product generalized additive model (from the *mgcv R* package): *r* ∼ te(*θ*, *ϕ*; bs = {“cc”, “tp”}, *k* = {60, 30}) with periodicity enforced for longitude (*θ*) via a cyclic cubic spline, a thin-plate spline for colatitude (*ϕ*), and smoothing selected by the restricted maximum likelihood (REML) algorithm. The fitted surface was evaluated on a longitude-colatitude grid, converted to a triangulated mesh by wrapping the seam and adding smooth polar caps, and all samples were projected radially onto this mesh (with linear extrapolation near the poles), yielding Cartesian coordinates for graph construction. We then built a weighted *k*-NN graph on the projected points; edges were assigned angular weights (central angles between sample directions) and, for each threshold *α* ∈ {5°, 10°, 15°, 20°, 25°, 30°, 40°, 50°} edges with angle >*α* were pruned. Starting from *k* = 5, *k* was increased (up to 2,000 in steps of 20) until the graph was connected, and geodesic distances (shortest paths) were embedded to ℝ^2^ via classic MDS. Among candidate settings, we selected the configuration minimizing Kruskal’s stress–1 while retaining all samples (breaking ties by smaller *k* and smaller *α*); for the dataset analyzed here, this yielded *k* = 820, *α* = 25° and Kruskal’s stress–1 = 0.04. The complete *R* script performing these tasks is available at https://github.com/flegontovlab/optimized_manifold_learning.

## Data Availability Statement

Supplementary datasets for this study (interactive 3D visualizations, sets of large vector figures, and spreadsheets) are available at https://zenodo.org/records/21884011 (doi: 10.5281/zenodo.21884011). Principal software packages used in the study are *slendr* v. 0.7.2.9000 and v. 0.8.0 (Petr et al. 2023, https://github.com/bodkan/slendr), *msprime* v. 1.2.0 (Baumdicker et al. 2022, https://github.com/tskit-dev/msprime), *PLINK* v. 1.9.0-b.8 and v. 2.0.0-a.6.9LM (https://www.cog-genomics.org/plink), *ADMIXTOOLS 2* (Maier et al. 2023, https://github.com/uqrmaie1/admixtools), *uwot* v. 0.2.4 (https://github.com/jlmelville/uwot), *phateR* v. 1.0.7 (https://github.com/KrishnaswamyLab/phateR), and a collection of our own scripts (https://github.com/flegontovlab/optimized_manifold_learning).

## Acknowledgements

J.C., O.F., P.C., and P.F. were supported by the Czech Ministry of Education, Youth and Sports (program ERC CZ, project no. LL2404). Additional funding came from: the Czech Science Foundation (project no. 25-17789S) for O.F. and P.F.; the European Union Operational Programme Just Transition (LERCO project CZ.10.03.01/00/22_003/0000003) for H.A. and P.F.; and a gift from Jean-Francois Clin to P.F. Computational resources were partly funded by the Ministry of Education, Youth and Sports of the Czech Republic through the e-INFRA CZ (ID:90254). We thank David Reich, Menno de Jong, Wibhu Kutanan, and Farhang Aghakhanian for sharing datasets used in their publications.

## Extended Data Figures

**Extended Data Figure 1.**
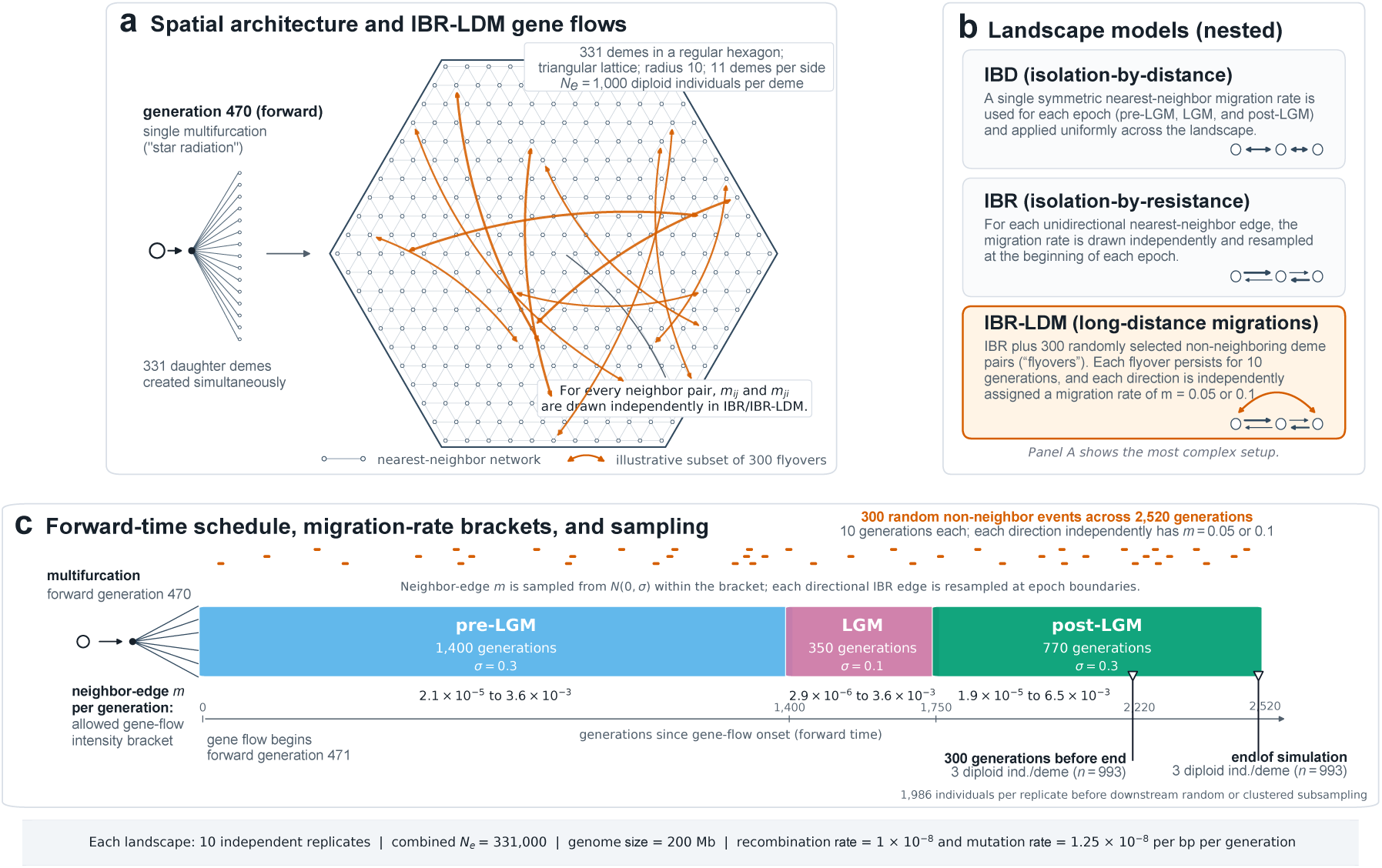
Design of the stepping-stone simulations. (**a**) At forward generation 470, a single multifurcation generated 331 panmictic demes, each with a constant effective size of 1,000 diploid individuals, arranged as a regular hexagonal patch of a triangular lattice. Gray edges indicate nearest-neighbor connections; each connection carries two directed migration edges, one in each direction. Orange arcs show an illustrative subset of the 300 transient long-distance “flyover” connections added in the IBR-LDM simulations. Each flyover connection was bidirectional, with the two directional migration rates assigned independently. (**b**) The three nested types of simulated landscapes. In IBD landscapes, a single symmetric nearest-neighbor migration rate, *m*, was used in each epoch: the same rate applied to all neighboring deme pairs across the landscape, and the two directions of each neighboring pair had identical rates. In IBR landscapes, nearest-neighbor migration was directional: for every ordered nearest-neighbor edge, the migration rate was sampled independently at the beginning of each epoch, so reciprocal rates between the same two demes could differ. In IBR-LDM landscapes, transient bidirectional non-neighbor flyovers were superimposed on the IBR nearest-neighbor landscape. (**c**) Gene-flow epochs, their durations, and migration rate sampling. Gene flow began at forward generation 471 and continued for 2,520 generations, divided into three epochs: pre-LGM, lasting 1,400 generations, with nearest-neighbor *m* = 2.1 × 10^-5^ to 3.6 × 10^-3^; LGM, lasting 350 generations, with nearest-neighbor *m* = 2.9 × 10^-6^ to 3.6 × 10^-3^; and post-LGM, lasting 770 generations, with nearest-neighbor *m* = 1.9 × 10^-5^ to 6.5 × 10^-3^. In IBD landscapes, one symmetric landscape-wide nearest-neighbor migration rate was redrawn at each epoch boundary. In IBR and IBR-LDM landscapes, every directional nearest-neighbor migration rate was independently redrawn at each epoch boundary. The 300 long-distance flyover events in IBR-LDM landscapes were distributed randomly across the simulation, lasted 10 generations each, and consisted of two directed migration edges. For each flyover direction, the migration rate was independently assigned as *m* = 0.05 or 0.1 with equal probability. Three diploid individuals were sampled per deme 300 generations before the end of the simulation and again at the end, yielding 993 individuals per sampling epoch before downstream random or clustered subsampling. Ten independent simulation replicates were generated for each landscape type.

**Extended Data Figure 2.**
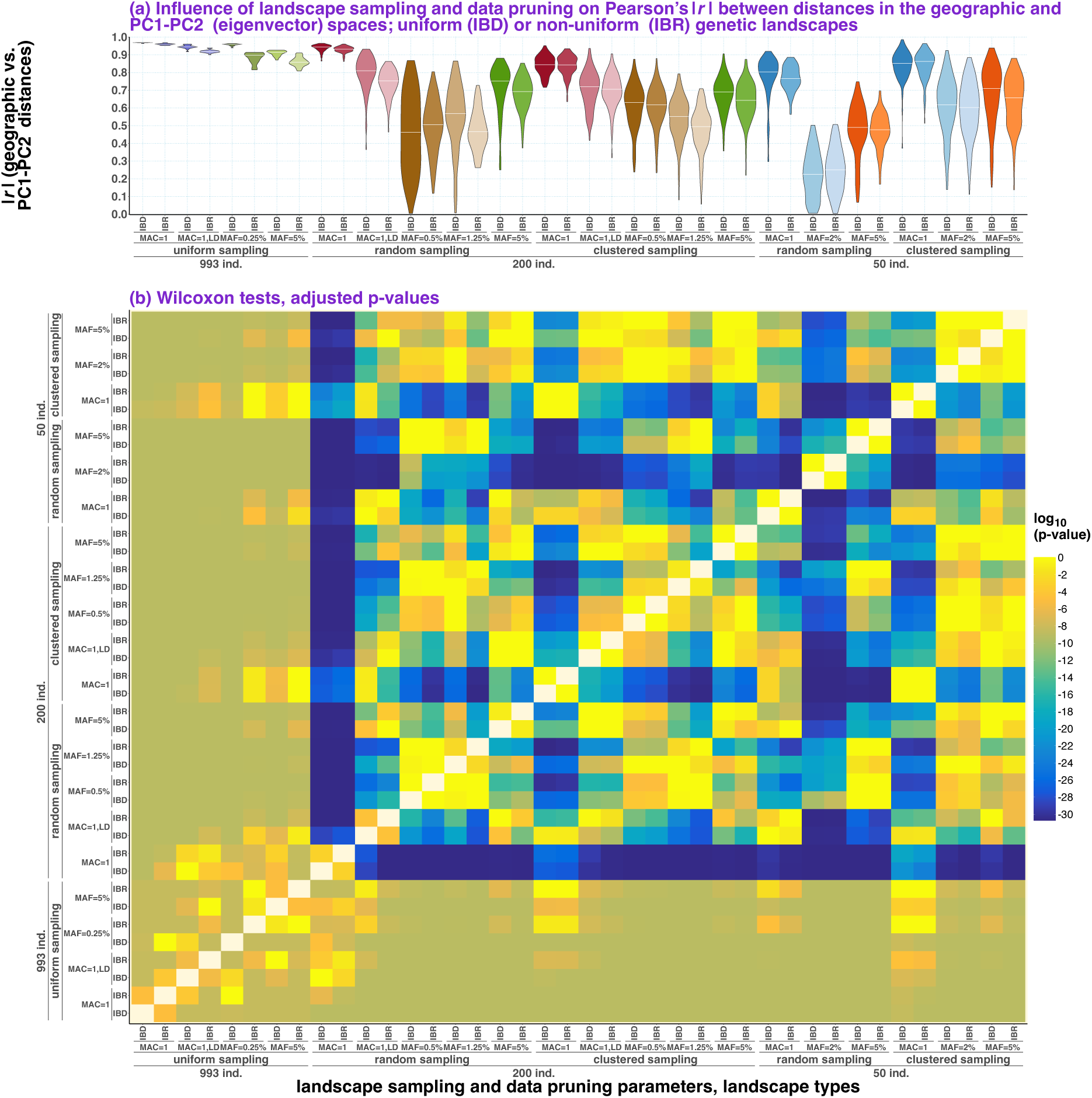
(**a**) Effects of genetic landscape sampling and data pruning on Pearson’s correlation coefficient (*r*) between all pairwise distances among individuals in the simulated landscape (“geographic”) space and in the PC1–PC2 eigenvector space. Violin plots with medians show the distributions of |*r*| across the two sampling time points (300 generations before the “present” and the “present”) × 10 simulation replicates × 5 subsampling replicates. Results are stratified by subsampling intensity, subsampling scheme (random or clustered), SNP-pruning regime (MAF-based or LD-based), and landscape type (IBD or IBR). (**b**) Results of pairwise comparisons among these distributions are shown as a matrix; p-values from unpaired Wilcoxon tests were adjusted for multiple testing using the Holm method.

**Extended Data Figure 3.**
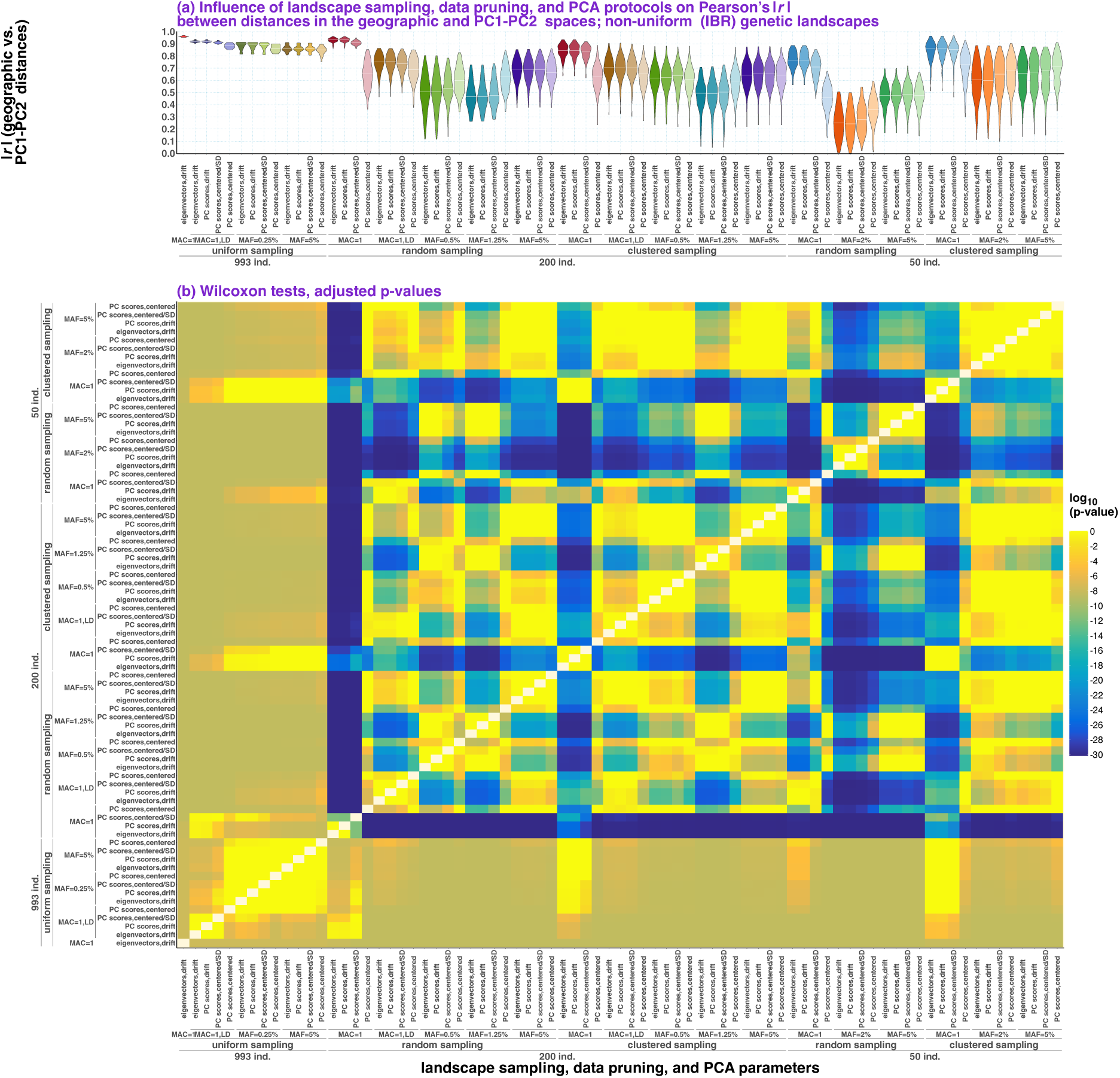
(**a**) Effects of genetic landscape sampling and data pruning on Pearson’s correlation coefficient (*r*) between all pairwise distances among individuals in the simulated landscape (“geographic”) space and in the PC1–PC2 space. Violin plots with medians show the distributions of |*r*| across the two sampling time points (300 generations before the “present” and the “present”) × 10 simulation replicates × 5 subsampling replicates. Results are stratified by subsampling intensity, subsampling scheme (random or clustered), SNP-pruning regime (MAF-based or LD-based), PCA algorithms, and data normalization approaches. (**b**) Results of pairwise comparisons among these distributions are shown as a matrix; p-values from unpaired Wilcoxon tests were adjusted for multiple testing using the Holm method.

**Extended Data Figure 4.**
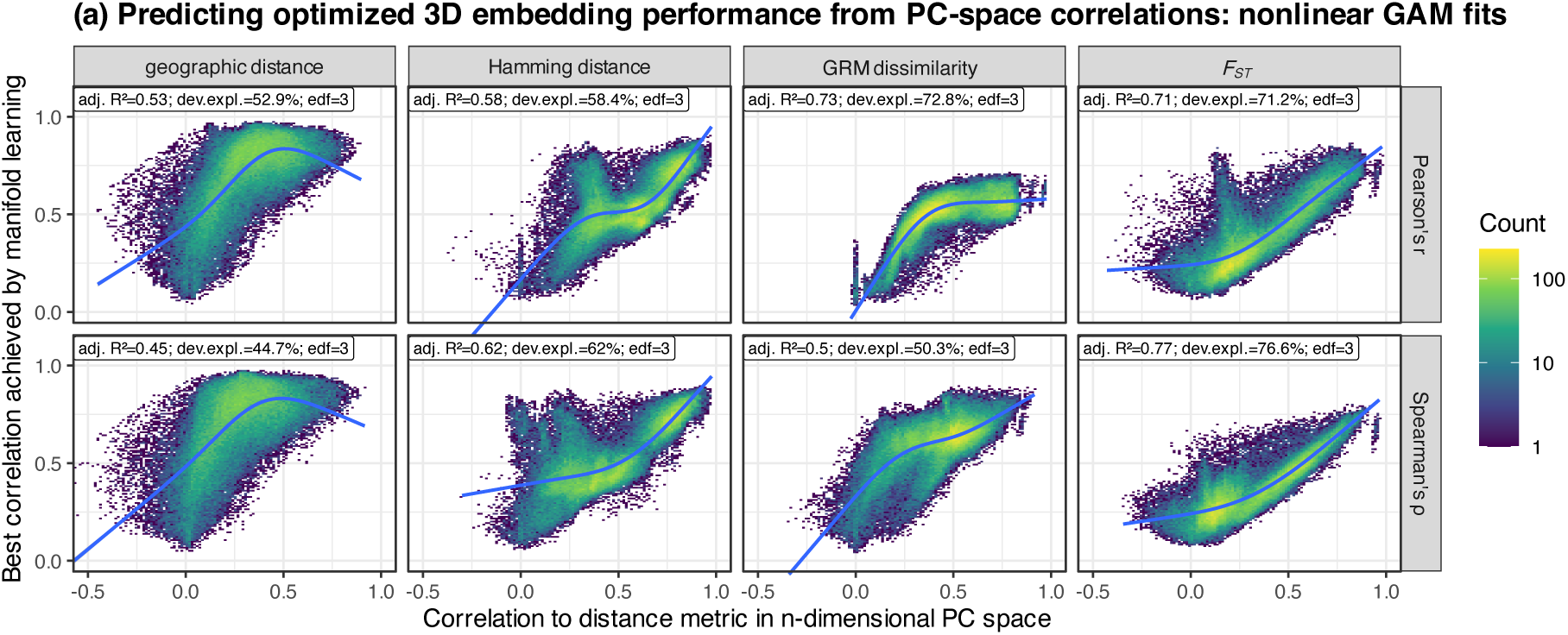
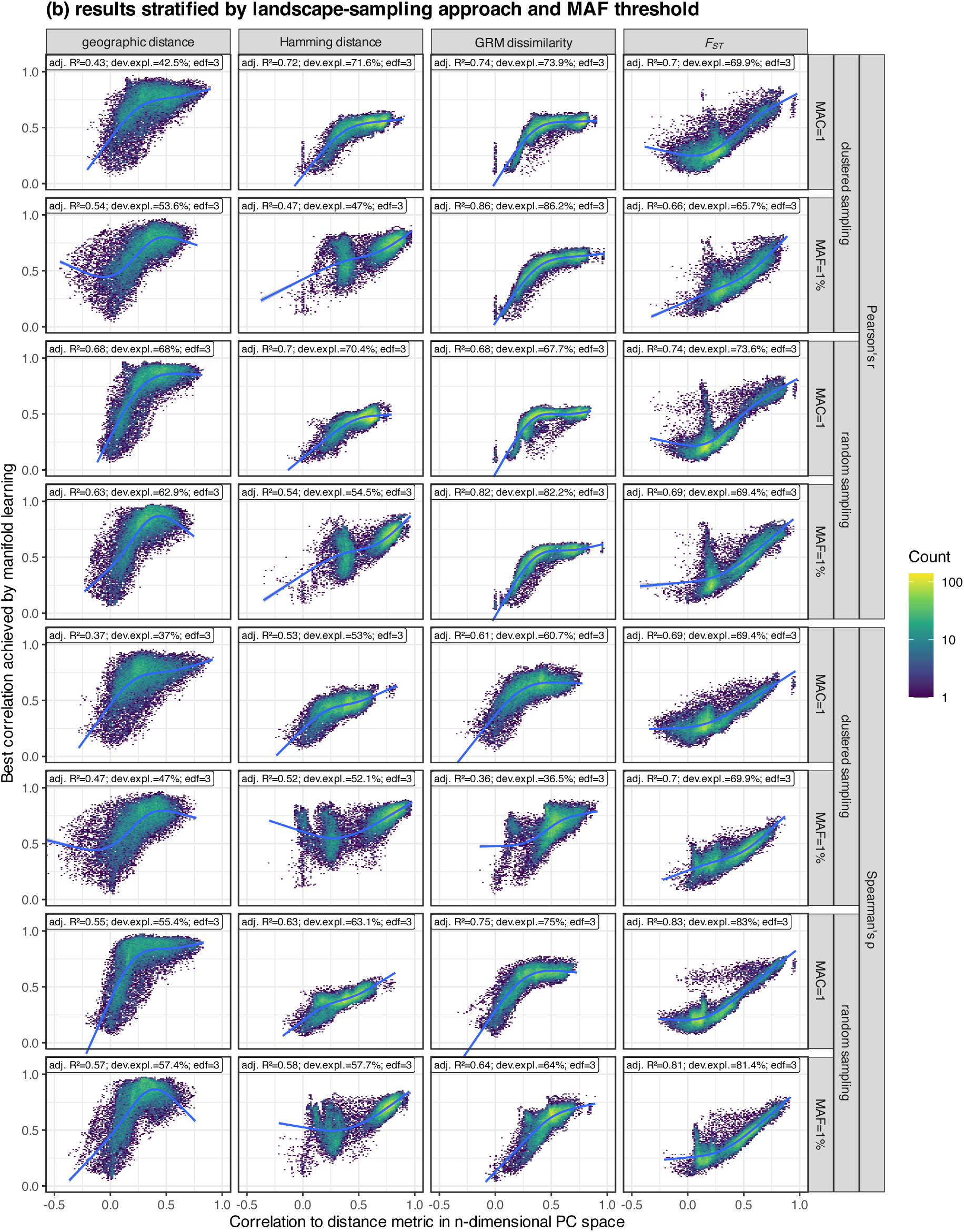
Predictors of manifold-learning optimization performance. X-axes show correlations between geographic or genetic distances – Hamming distance, GRM distance, or more precisely GRM-derived dissimilarity, and *F_ST_* – and distances in *n*-dimensional PC spaces derived from the IBR-LDM simulations. PC-space distances were computed using Euclidean, Manhattan, or cosine metrics. Y-axes show the best manifold-learning optimization result for the corresponding PC space: the correlation between Euclidean distances in a single best optimized 3D LDE and the same geographic or genetic distance. In panel **a**, results are stratified by geographic/genetic distance metric in columns and by correlation coefficient, Pearson’s or Spearman’s, in rows. In panel **b**, results are further stratified by landscape-sampling approach and SNP-filtering threshold in rows. GAMs of the form *y* ∼ *s*(*x*, *k* = *4*) were fitted to each distribution, and key model-fit statistics are shown above each plot: adjusted R², deviance explained, and effective degrees of freedom (edf). Distributions are shown as binned density plots with a log_10_-transformed color scale.

**Extended Data Figure 5.**
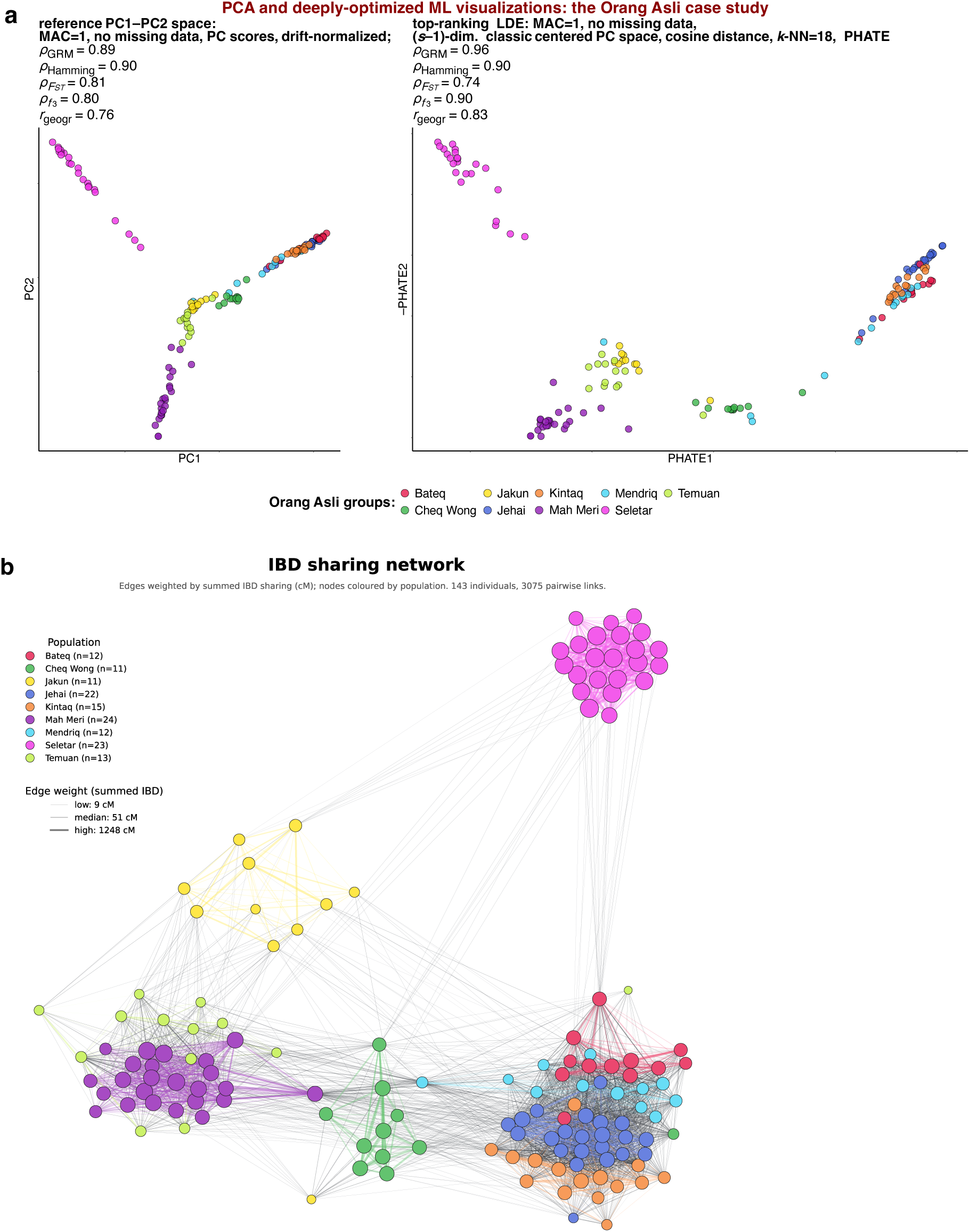
Comparison of PCA and optimized manifold-learning visualization of population structure in the Orang Asli case study. (**a**) Reference PC1–PC2 space and a top-ranked 2D LDE, generated with PHATE from the full-dimensional classic centered PC space, using cosine distance in the input space and *k*-NN = 18. Correlations with genetic distances and great-circle geographic distances are shown for both embeddings. (a) Force-directed layout of the IBD-sharing graph for the same individuals. Autosomal genotypes were statistically phased with *Beagle* v5.5 without a reference panel, using chromosome-specific GRCh37 genetic maps. IBD segments were inferred from the phased haplotypes with *hap-IBD* (Zhou et al. 2020) using the same genetic maps. For the network analysis shown here, only segments ≥8 cM were retained, and edges were weighted by the cumulative length of retained segments shared by each pair of individuals. Individuals with no retained IBD segments ≥8 cM were excluded from the graph. The network was visualized as a weighted Fruchterman–Reingold force-directed graph, with node size proportional to total IBD connectivity and edge width and opacity scaled by cumulative IBD sharing.

**Extended Data Figure 6.**
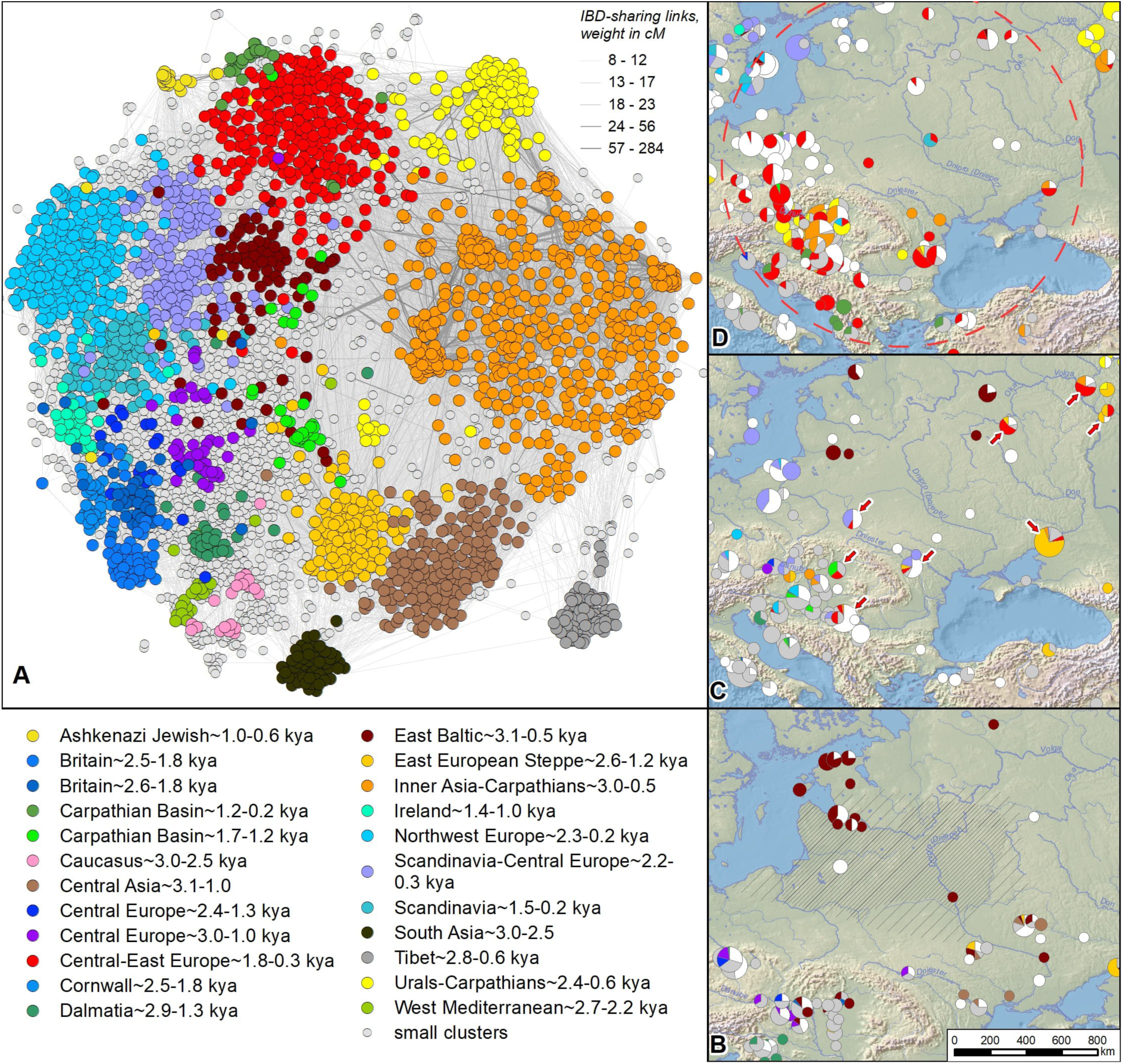
(**a**) A layout (2D LDE) of the IBD-sharing graph for the ancient Eurasian case study generated with the *ForceAtlas2* force-directed algorithm (Jacomy et al. 2014). Highly connected IBD-sharing communities (clusters) identified with the Leiden community-detection algorithm (Traag et al. 2019) are listed in the legend. (**b**) Spatial distribution of the IBD-sharing communities in the periods preceding the Roman and Migration Era. Hatched areas show the geographic distributions of material cultures associated tentatively with Baltic speakers. (**c**) The first appearance of the Slavic-associated Central–East Europe (CEE) IBD-sharing community (shown in red) in the Roman Era and the initial phase of its dispersal in the Migration Period Central– East Europe. (**d**) Expansion limits of the CEE IBD-sharing community in the Medieval period (the red dashed circle). On all the maps, archaeological sites within 100 km are aggregated.

**Extended Data Figure 7.**
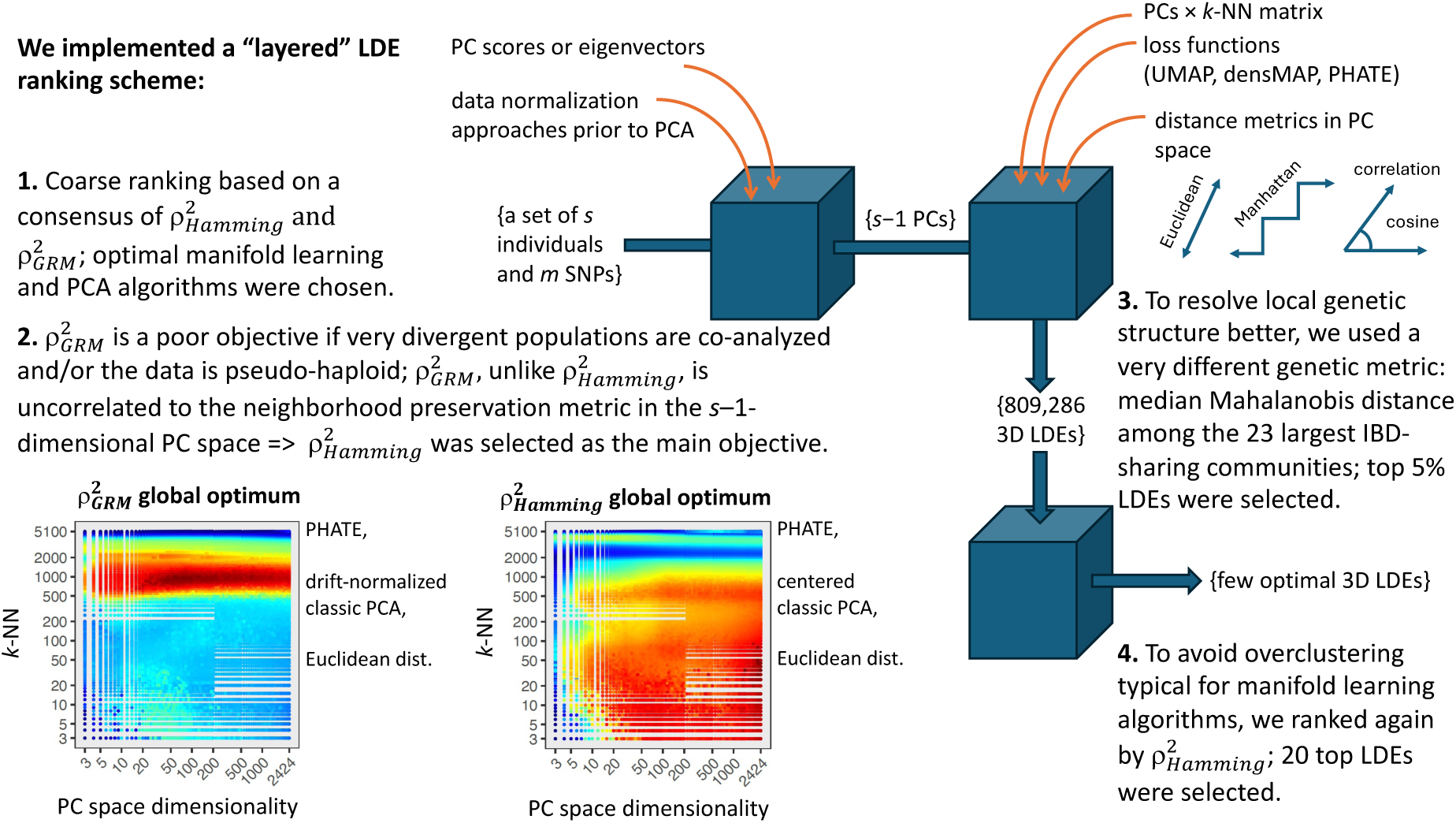
A diagram illustrating the objective-guided manifold-learning optimization protocol used in the ancient and present-day Eurasian case studies.

## Supplementary Information

### Supplementary Figure Legends

**Supplementary Fig. 1**: Counts of polymorphic SNP loci per simulated dataset, shown as violin plots across landscape-sampling strategies, SNP-pruning/filtering settings, and landscape types (IBD, IBR, IBR-LDM). For simplicity, results are shown only for the “present-day” sampling epoch. The y-axis is logarithmic.

**Supplementary Fig. 2**: PCA visualizations of SNP data from simulated IBD landscapes, showing the effect of subsampling independently of SNP filtering; all datasets shown here are unfiltered. Results are shown for three simulation replicates subjected to random landscape sampling (**a–c**) at two densities (200 or 50 individuals), and for three simulation replicates subjected to clustered sampling at the same densities (**d–f**). Results for uniformly and densely sampled landscapes are shown for comparison in each panel, along with results for five subsampling replicates per sampling density level. In each panel, the plots in rows two and four show the sample distributions on the landscape, and the plots below them show the corresponding PC1–PC2 eigenvector spaces. Point size indicates proximity to the landscape center; point colors distinguish landscape sectors; and point opacity indicates the number of individuals sampled from each deme. The x- and y-axes are scaled identically. Pearson’s correlation coefficient between Euclidean distances in PC1–PC2 space and Euclidean (“geographic”) distances on the landscape is shown above each PC plot.

**Supplementary Fig. 3**: PCA visualizations of SNP data from simulated IBD landscapes, with 200 individuals sampled either randomly (**a–g**) or in clusters (**h–p**). The simulation/subsampling replicates shown here were selected to illustrate various common patterns in the data and represent only a fraction of the full simulation dataset. In each panel, the first plot shows the sample distribution on the landscape for a selected simulation and subsampling replicate, and the subsequent plots show PC1–PC2 eigenvector spaces for all SNP-pruning and filtering settings tested. Point size indicates proximity to the landscape center; point colors distinguish landscape sectors; and point opacity indicates the number of individuals sampled from each deme. The x- and y-axes are scaled identically. Pearson’s correlation coefficient between Euclidean distances in PC1–PC2 space and Euclidean (“geographic”) distances on the landscape is shown above each PC plot.

**Supplementary Fig. 4**: PCA visualizations of SNP data from admixture-graph-shaped (AGS) and strictly tree-shaped demographic simulations. In all cases, simulated histories with random topology included 40 leaves and either 15 or zero pulse-like admixture events. Chromosome length was set to 500 Mbp, and *N_e_* was constant, with 1,000 diploid individuals on each branch. Mutation and recombination rates matched those used in the landscape simulations in this study. Maximum tree depth was set to 3,000 generations, and lineage-divergence and admixture dates were assigned quasi-randomly following the algorithm of Flegontova et al. (2025). PC1– PC2 spaces for AGS simulations are shown in panels **a–h**, and those for tree simulations in panels **i–p**. Graph branches were sampled either at the end of the simulation or at different points in time, assigned according to the algorithm of Flegontova et al. (2025): panels **a–d** and **i–l** show “present-day” sampling, whereas panels **e–h** and **m–p** show time-stratified sampling. These two sampling scenarios were applied to independent random topologies, rather than to the same underlying topology. The corresponding simulated graphs or trees are shown below each panel. For each of the four simulation setups – AGS or tree-shaped simulations, with either present-day or time-stratified sampling – 10 simulation replicates were generated; here, we show results for replicates 1–4. In each panel, the first PC1–PC2 plot is based on uniform sampling of all leaves, with 40 leaves and five individuals per leaf. In the remaining plots, either 25% (**a, b, e, f, i, j, m, n**) or 10% (**c, d, g, h, k, l, o, p**) of the original 200-individual sample was randomly retained; these 10 plots show the corresponding subsampling replicates. For time-stratified sampling, point size indicates sampling date, measured in generations before the end of the simulation.

**Supplementary Fig. 5**: PCA visualizations of SNP data from 1D linear (**a–c**) and circular (**d–f**) stepping-stone IBR simulations. Chromosome length was set to 500 Mbp, and *N_e_* was constant, with 1,000 diploid individuals on each branch. Mutation and recombination rates matched those used in the 2D landscape simulations in this study. All other simulation parameters matched those used in the 2D landscape simulations of Flegontova et al. (2025), namely simulations with per-generation gene-flow intensities of ∼10^−4^–10^−3^ in each direction. For each of the two simulation setups – linear and circular – 10 simulation replicates were generated; here, we show results for replicates 1–3. In each panel, the first PC1–PC2 plot is based on uniform sampling of all demes, with 40 demes and five individuals per deme. In the remaining plots, 10% of the original 600-individual sample was randomly retained; these 10 plots show the corresponding subsampling replicates. No MAF-based SNP filtering was performed. Samples were taken at the end of three gene-flow epochs – “pre-LGM,” “LGM,” and “post-LGM” (Flegontova et al. 2025) – with epochs indicated by marker shape. Color indicates angular position on the simulated circle and linear position along the simulated line.

**Supplementary Fig. 6**: PCA visualizations of SNP data from simulated IBD landscapes after extreme data purging: 50 individuals were sampled, with either all variants retained (MAC = 1) or singleton variants removed (MAC = 2; MAF = 2%). Results are shown for three simulation replicates subjected to random landscape-wide sampling (**a–c**) and for the same three simulation replicates subjected to clustered sampling (**d–f**). Results for uniformly and densely sampled landscapes are shown for comparison in each panel, along with results for five subsampling replicates per sampling density level. In each panel, the plots in rows two and four show the sample distributions on the landscape, and the plots below them show the corresponding PC1–PC2 eigenvector spaces. Point size indicates proximity to the landscape center; point colors distinguish landscape sectors; and point opacity indicates the number of individuals sampled from each deme. The x- and y-axes are scaled identically. Pearson’s correlation coefficient between Euclidean distances in PC1–PC2 space and Euclidean (“geographic”) distances on the landscape is shown above each PC plot.

**Supplementary Fig. 7**: Eigenvalues plotted across landscape-sampling, SNP-pruning/filtering, and data-normalization strategies. Each point represents one PC from one simulation replicate, sampling epoch, and subsampling replicate. Results are shown for the “present-day” sampling epoch, classic PC scores calculated using *smartSNP*, and IBR-LDM (panel **a**; 200 or 50 individuals sampled) or IBR landscapes (panel **b**, 200 individuals sampled; panel **c**, 993 or 50 individuals sampled). For clarity, only the first 200 PCs are shown in each case.

**Supplementary Fig. 8**: Component-wise statistics for PCs 1 to *s − 1* (*s* = number of samples): (**a**) eigenvalue and sum of squares; (**b**) variance and spread; and (**c**) mean and median. Panels are stratified by PCA algorithm (eigenvectors reported by *smartPCA* or PC scores reported by *smartSNP*), subsampling intensity, subsampling scheme (random or clustered), and SNP-pruning regime (MAF-based or LD-based). Colors are used for visual distinction only. Y-axis scales vary among panels.

**Supplementary Fig. 9**: Correlations between PC-space distances and geographic or genetic distances in IBR simulations. Pearson’s *r* was calculated between pairwise “geographic” distances on simulated IBR landscapes (**a,b**) or inter-sample Hamming genetic distances (**c, d**) and Euclidean or Manhattan distances in PC spaces of varying dimensionality. Each point represents one combination of simulation replicate, sampling epoch, subsampling replicate, and number of PCs used to calculate distances. Results are stratified by subsampling intensity: datasets of 200 individuals are shown in panels **a** and **c**, whereas datasets of 993 or 50 individuals are shown in panels **b** and **d**. Within each panel, results are further stratified by PCA algorithm and data normalization approach, PC-space distance metric, subsampling scheme (random or clustered), and SNP-pruning regime (MAF-based or LD-based). Colors are used for visual distinction only. Within each row of panels, the setting that attains the highest per-PC median correlation is outlined in orange, and the corresponding median value is shown by a dashed horizontal line.

**Supplementary Fig. 10**: All-vs-all Spearman’s and Pearson’s correlations among distance metrics, summarized across IBR-LDM simulation and subsampling iterations as violin plots. The distances considered are: geographic distance on the simulated landscapes; Hamming distance, or 1 − IBS (Chang et al. 2015); GRM-derived dissimilarity (Yang et al. 2011; Chang et al. 2015); unstandardized covariance-derived dissimilarity (Chang et al. 2015); Euclidean distance between samples in 3D MDS space based on Hamming distances (Chang et al. 2015); *F_ST_*; and *f*_2_-statistics (Patterson et al. 2012). Results are stratified by MAF threshold and landscape-sampling approach.

**Supplementary Fig. 11**: Correlations between PC-space distances and genetic or geographic distances in IBR-LDM simulations. Pearson’s *r* and Spearman’s *ρ* were calculated between pairwise inter-sample genetic distances − *F_ST_* (**a**), Hamming distance (**b**), and GRM-derived dissimilarity (**c**) − or geographic distances on simulated IBR-LDM landscapes (**d**), and Euclidean, Manhattan, or cosine distances in PC spaces of varying dimensionality. Each point represents one combination of simulation replicate, sampling epoch, subsampling replicate, and number of PCs used to calculate distances. Results are stratified by PCA algorithm and data normalization approach, PC-space distance metric, correlation coefficient, subsampling scheme (random or clustered), and SNP-pruning regime (MAF-based or LD-based). Colors are used for visual distinction only. Within each row of panels, the setting that attains the highest per-PC median correlation is outlined in orange, and the corresponding median value is shown by a dashed horizontal line.

**Supplementary Fig. 12**: Comparing the performance of second-level objectives used to select the best manifold-learning-derived LDE, part 1. Second-level objectives are defined as combinations of a genetic distance metric and a correlation coefficient. Violin plots show distributions of correlations, Pearson’s |*r*| or Spearman’s |*ρ*|, between LDE distance and the selected genetic distance for the best manifold-learning-derived 3D LDEs, shown as violins with red borders, and for the corresponding 3D PCA LDEs, shown as violins with black borders. Distributions are computed across IBR-LDM simulation and subsampling replicates. Results are shown for the following objectives: *f*_2_-Pearson and *f*_2_-Spearman (**a**, **b**), *F_ST_*-Pearson and *F_ST_*-Spearman (**c**, **d**), covariance-Pearson and covariance-Spearman (**e**, **f**), GRM-Pearson and GRM-Spearman (**g**, **h**), and Hamming-Pearson and Hamming-Spearman (**i**, **j**). For comparison, we also show results ranked by correlation with geographic distances (**k**, **l**). Along the x-axes, results are stratified by landscape-sampling approach, SNP-filtering threshold, PC-space type, and LDE type, either PCA or manifold-learning-derived. Pairwise comparisons among distributions are shown as matrices below the violin plots; p-values from unpaired Wilcoxon tests were adjusted for multiple testing using the Holm method. For each PCA–manifold-learning pair of distributions, adjusted Wilcoxon test p-values are also shown above the brackets using the following notation: ****, p < 0.0001, highly significant; ***, p < 0.001, highly significant; **, p < 0.01, moderately significant; *, p ≤ 0.05, weakly significant; ns, not significant.

**Supplementary Fig. 13**: Comparing the performance of second-level objectives used to select the best manifold-learning-derived LDE, part 2: recovery of geographic patterns. Violin plots show distributions of Pearson’s correlations between LDE distances and geographic distances for the best manifold-learning-derived 3D LDEs, shown as violins with red borders, and for the corresponding 3D PCA LDEs, shown as violins with black borders. Distributions are computed across IBR-LDM simulation and subsampling replicates. Best manifold-learning-derived LDEs were selected using the following objectives: *f*_2_-Pearson and *f*_2_-Spearman (**a**, **b**), *F_ST_*-Pearson and *F_ST_*-Spearman (**c**, **d**), covariance-Pearson and covariance-Spearman (**e**, **f**), GRM-Pearson and GRM-Spearman (**g**, **h**), and Hamming-Pearson and Hamming-Spearman (**i**, **j**). For comparison, we also show results ranked by correlation with geographic distances (**k**, **l**). Along the x-axes, results are stratified by landscape-sampling approach, SNP-filtering threshold, PC-space type, and LDE type, either PCA or manifold-learning-derived. Pairwise comparisons among distributions are shown as matrices below the violin plots; p-values from unpaired Wilcoxon tests were adjusted for multiple testing using the Holm method. For each PCA–manifold-learning pair of distributions, adjusted Wilcoxon test p-values are also shown above the brackets using the following notation: ****, p < 0.0001, highly significant; ***, p < 0.001, highly significant; **, p < 0.01, moderately significant; *, p ≤ 0.05, weakly significant; ns, not significant.

**Supplementary Fig. 14**: Genetic differentiation measured by Hudson’s *F_ST_* in simulated demes and present-day human groups. The red boxplot shows all-versus-all comparisons among simulated demes from exhaustively sampled IBR-LDM (**a**) or IBR (**b**) landscapes, with three individuals sampled per deme. The remaining boxplots show inter- and intra-continental comparisons among groups in the Simons Genome Diversity Project dataset (Mallick et al. 2016). The simulated-deme distribution was compared with each empirical distribution using an unpaired Wilcoxon test. P-values were adjusted for multiple testing using the FDR method and are indicated as follows: ****, p < 0.0001, highly significant; ***, p < 0.001, highly significant; **, p < 0.01, moderately significant; *, p ≤ 0.05, weakly significant; ns, not significant.

**Supplementary Fig. 15**: Distribution of top-ranked LDEs in the parameter space of the objective-guided embedding-optimization protocol. Optima in the four regions of parameter space (rectangles delineated by the solid red lines) should be interpreted separately, because each region is derived from a distinct set of subsampling replicates and SNPs. One hundred top-ranked LDEs were generated from each unique IBR-LDM simulated input dataset and selected using different objectives: *f*_2_-Pearson and *f*_2_-Spearman (**a**, **b**), *F_ST_*-Pearson and *F_ST_*-Spearman (**c**, **d**), covariance-Pearson and covariance-Spearman (**e**, **f**), GRM-Pearson (**g**), Hamming-Pearson and Hamming-Spearman (**h**, **i**), geography-Pearson and geography-Spearman (**j**, **k**). An input dataset is defined as a combination of simulation/subsampling replicate and MAF-filtering level, either no MAF filtering or a 1% MAF threshold; all other settings are treated as parameters of the manifold-learning-optimization protocol. For clarity, one dimension of the parameter space – the k-NN parameter of the manifold-learning algorithms – is collapsed. The density of LDEs at each point in the remaining parameter space is shown on a linear color scale.

**Supplementary Fig. 16**: Correlations between PC-space distances and genetic or geographic distances in the Orang Asli case study. Pearson’s *r* and Spearman’s *ρ* were calculated between pairwise inter-individual genetic distances − *F_ST_* (**a**), Hamming distance (**b**), GRM-derived dissimilarity (**c**), and outgroup *f*_3_-statistics *f*_3_(Mbuti; X, Y) (**d**) − or great-circle geographic distances (**e**), and Euclidean, Manhattan, cosine, or correlation distances in PC spaces of varying dimensionality. Eigenvalues are also shown in panel **f**. Results are stratified by PCA algorithm and data normalization approach, PC-space distance metric, correlation coefficient, missing-data treatment, and SNP-filtering approach. Colors are used for visual distinction only. Within each row of panels, the setting that attains the highest per-PC correlation is outlined in orange, and the corresponding maximal value is shown by a dashed horizontal line.

**Supplementary Fig. 17**: Correlations between PC-space distances and genetic or geographic distances in the *Cervus elaphus* case study. Pearson’s *r* and Spearman’s *ρ* were calculated between pairwise inter-individual genetic distances − *F_ST_* (**a**), Hamming distance (**b**), GRM-derived dissimilarity (**c**), and outgroup *f*_3_-statistics *f*_3_(wapiti; X, Y) (**d**) − or great-circle geographic distances (**e**), and Euclidean, Manhattan, cosine, or correlation distances in PC spaces of varying dimensionality. Eigenvalues are also shown in panel **f**. Results are stratified by PCA algorithm and data normalization approach, PC-space distance metric, correlation coefficient, missing-data treatment, and SNP-filtering approach. Colors are used for visual distinction only. Within each row of panels, the setting that attains the highest per-PC correlation is outlined in orange, and the corresponding maximal value is shown by a dashed horizontal line.

**Supplementary Fig. 18**: Correlations between PC-space distances and genetic or geographic distances in the PNG case study. Pearson’s *r* and Spearman’s *ρ* were calculated between pairwise inter-individual genetic distances − *F_ST_* (**a**), Hamming distance (**b**), GRM-derived dissimilarity (**c**), and outgroup *f*_3_-statistics *f*_3_(Mbuti; X, Y) (**d**) − or great-circle geographic distances (**e**), and Euclidean, Manhattan, cosine, or correlation distances in PC spaces of varying dimensionality. Eigenvalues are also shown in panel **f**. Results are stratified by PCA algorithm and data normalization approach, PC-space distance metric, correlation coefficient, missing-data treatment, and SNP-filtering approach. Colors are used for visual distinction only. Within each row of panels, the setting that attains the highest per-PC correlation is outlined in orange, and the corresponding maximal value is shown by a dashed horizontal line.

**Supplementary Fig. 19**: Correlations between PC-space distances and genetic or geographic distances in the SEA case study. Pearson’s *r* and Spearman’s *ρ* were calculated between pairwise inter-individual genetic distances − *F_ST_* (**a**), Hamming distance (**b**), GRM-derived dissimilarity (**c,g**), and outgroup *f*_3_-statistics *f*_3_(Mbuti; X, Y) (**d**) − or great-circle geographic distances (**e**), and Euclidean, Manhattan, cosine, or correlation distances in PC spaces of varying dimensionality. Eigenvalues are also shown in panel **f**. To better visualize the primary manifold-learning optimum in eigenvector spaces, panel **g** also shows the results for GRM-derived dissimilarities on logarithmic x-axes. Results are stratified by PCA algorithm and data normalization approach, PC-space distance metric, correlation coefficient, missing-data treatment, and SNP-filtering approach. Colors are used for visual distinction only. Within each row of panels, the setting that attains the highest per-PC correlation is outlined in orange, and the corresponding maximal value is shown by a dashed horizontal line.

**Supplementary Fig. 20**: Locations of the ancient individuals (**a**) and present-day reference individuals (**b, c**) included in the Eurasian case studies are shown on terrain maps of Eurasia. Marker sizes scale non-linearly with the number of individuals at each location. Locations of present-day individuals belonging to the Slavic-Baltic IBD-sharing community are highlighted in panel **c**.

**Supplementary Fig. 21**: Component-wise statistics of PC scores – eigenvalue, sum of squares, variance, range, mean, and median – for PCs 1 to *s* – 1 in the Eurasian case studies. Panels are stratified by PCA algorithm (*smartPCA* eigenvectors or classic PC scores from the *smartSNP* package) and preprocessing (centered with normalization by genetic-drift dispersion vs. centered only), and are shown separately for (**a**) the basis samples (Human Origins; 2,425 present-day individuals) and (**b**) the projected samples (Human Origins SNPs; 5,133 ancient individuals). Colors aid visual distinction only. The x-axis (PC number) is on a logarithmic scale; y-axis scales differ across panels.

### Supplementary Table Legends

**Supplementary Table 1**: Fits of PC1–PC2 or PC1–PC2–PC3 spaces to genetic distances/dissimilarities and geographic distances are shown for the six empirical case studies, together with key metadata for each dataset.

**Supplementary Table 2**: Dataset compositions and meta-data for the six empirical case studies.

**Supplementary Table 3**: Correlations among genetic and geographic distances/dissimilarities across the six empirical case-study datasets.

**Supplementary Table 4**: Assessment of promising LDEs in the ancient and present-day Eurasian case studies. We evaluate a suite of fidelity metrics for: (i) the “ancient” 3D LDEs that maximize different objectives (8 LDEs shown in **Supplementary Dataset 13**); (ii) the outcomes of the multi-step LDE-ranking procedure applied to the ancient Eurasian dataset (20 3D LDEs from step 3, three of which are visualized in **Supplementary Dataset 12**), including the final LDE; (iii) the 2D Isomap unfolding of the final LDE; (iv) three 3D PCAs on ancient Eurasian data; (v) the “present-day” 3D LDEs that maximize different objectives (11 LDEs shown in **Supplementary Dataset 16**); (vi) the outcomes of the multi-step LDE-ranking procedure applied to the present-day Eurasian dataset (20 3D LDEs, five of which are visualized in **Supplementary Dataset 17**), including the final LDE; and (vii) three 3D PCAs on present-day Eurasian data. The following metrics are reported: *ρ*^2^ and *r*^2^ fits to great-circle geographic distances; *ρ*^2^ and *r*^2^ fits to Hamming distances (using either the default or the flat missingness correction in *PLINK* v. 1.9), GRM distances, covariance-based genetic distances; eight cluster-separation metrics derived from IBD or Y-lineage data; and six metrics quantifying preservation of geometry in the (*s – 1*)-dimensional PC space. For most PHATE-based LDEs, *ρ*^2^ and *r*^2^ values are presented as Δ*ρ*^2^ and Δ*r*^2^ relative to the best-fitting PC1–PC2–PC3 space (the classic PCA on centered data normalized by genetic-drift dispersion). For the final PHATE LDE on the ancient Eurasian data, its Isomap unfolding, and all the PCAs, we additionally report *ρ*^2^, *r*^2^, *ρ*, and *r*. For the LDEs on present-day Eurasian data, we also report *ρ* and *r* for two additional genetic distance metrics: *F_ST_* and outgroup *f*_3_(Mbuti; X, Y). The definitions of the IBD cluster-separation metrics and some geometry-preservation metrics differ between the ancient and present-day datasets (see row 47).

**Supplementary Table 5**: (**a**) All-vs-all correlations (Spearman *ρ* or Pearson *r*) among key LDE fidelity metrics in the ancient Eurasian case study: 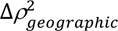 (from great-circle distances), 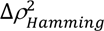 (using the default missingness correction), 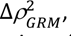 six cluster-separation metrics derived from IBD or Y-lineage data, and six metrics assessing preservation of geometry in the (*s – 1*)-dimensional PC space. Correlation coefficients were computed using all PHATE-based LDEs generated from classic PC scores (85,188 LDEs), which represent the broad global optimum in the parameter space. (**b**) A similar set of correlations computed for the present-day Eurasian case study. The set of LDE fidelity metrics differs from that used in table **a**: 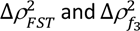 are included, Y-chromosome-based cluster-separation metrics are omitted, and two neighborhood- and density-preservation measures are computed using proportionally smaller *k* values. Correlation coefficients were computed using all PHATE-based LDEs generated from classic PC scores (83,544 LDEs).

**Supplementary Table 6**: Selecting the most informative level of the Y-chromosome phylogeny in the ancient Eurasian case study: evaluation of clustering quality in the PHATE space (final LDE) across a range of inter-lineage coalescence-time thresholds. Three standard indices are reported: Silhouette, Davies–Bouldin, and Caliński–Harabasz.

### Supplementary Datasets

**Supplementary Dataset 1**: Comprehensive summary of manifold-learning-derived LDEs across the full parameter space of the objective-guided embedding-optimization protocol. Each point in the PC-space-dimensionality versus *k*-NN biplots is colored by the median Δ*r*² or Δ*ρ*² between a manifold-learning-derived embedding and the corresponding drift-normalized classic PC1–PC2–PC3 space. Each figure in this dataset consists of a collection of such biplots spanning different PC-space types, PC-space distance metrics, and manifold-learning algorithms. Individual figures are defined by the combination of LDE-ranking objective, landscape-sampling approach, either random or clustered, and MAF-filtering threshold, either no filtering or MAF = 1%. Separate sets of figures show the PC-space-dimensionality versus *k*-NN parameter space using either log_10_ or linear axis scales.

Available at: https://zenodo.org/records/21884011/files/SData_1.zip

**Supplementary Dataset 2**: An interactive R application for visualizing any selected PC1–PC2–PC3 space derived from the IBR-LDM simulations and for generating UMAP, UMAP/densMAP, densMAP, or PHATE 3D LDEs from these data using user-defined parameters. For each embedding, we display Pearson’s correlation with geographic distances and Spearman’s correlation with GRM-derived dissimilarities, the best-performing LDE-ranking objective.

Available at: https://zenodo.org/records/21884011/files/SData_2.zip

**Supplementary Dataset 3**: Parameters and fits of the top 100 LDEs for each objective function are shown for each of the four case studies in which simple, non-layered LDE ranking was used.

Available at: https://zenodo.org/records/21884011/files/SData_3.xlsx

**Supplementary Dataset 4**: Comprehensive summary of manifold-learning-derived LDEs across the full parameter space of the objective-guided embedding-optimization protocol in the Orang Asli case study. Each point in the PC-space-dimensionality versus *k*-NN biplots is colored by the Δ*r*² or Δ*ρ*² between a manifold-learning-derived embedding and the corresponding drift-normalized classic PC1–PC2 space (MAC = 1, no missing data). Each figure in this dataset consists of a collection of such biplots spanning different PC-space types, PC-space distance metrics, and manifold-learning algorithms. Individual figures are defined by the LDE-ranking objective: *F_ST_*, Hamming distance (default), GRM-derived dissimilarity, or outgroup *f*_3_-statistics, each evaluated using either Pearson’s or Spearman’s correlation. Geography-based Pearson and Spearman objectives are shown for comparison. Separate sets of figures show the PC-space-dimensionality versus *k*-NN parameter space using either log_10_ or linear axis scales.

Available at: https://zenodo.org/records/21884011/files/SData_4.zip

**Supplementary Dataset 5**: Alternative PC1–PC2–PC3 spaces (**a–h**) and the top-ranked manifold-learning-derived LDE (**i**) from the *Cervus elaphus* case study, shown as interactive 3D plots. For the PCA algorithm, SNP-filtering scheme, and missing-data removal regime used for each analysis, see the file names and **Supplementary** Table 1. Corresponding goodness-of-fit statistics are provided in **Supplementary** Table 1 and **Supplementary Dataset 3.** Individuals are colored by the geographic regions defined by de Jong et al. (2025).

Available at: https://zenodo.org/records/21884011/files/SData_5.zip

**Supplementary Dataset 6**: Comprehensive summary of manifold-learning-derived LDEs across the full parameter space of the objective-guided embedding-optimization protocol in the *Cervus elaphus* case study. Each point in the PC-space-dimensionality versus *k*-NN biplots is colored by the Δ*r*² or Δ*ρ*² between a manifold-learning-derived embedding and the corresponding drift-normalized classic PC1–PC2–PC3 space (MAC = 1, no missing data). Each figure in this dataset consists of a collection of such biplots spanning different PC-space types, PC-space distance metrics, and manifold-learning algorithms. Individual figures are defined by the LDE-ranking objective: *F_ST_*, Hamming distance (default), GRM-derived dissimilarity, or outgroup *f*_3_-statistics, each evaluated using either Pearson’s or Spearman’s correlation. Geography-based Pearson and Spearman objectives are shown for comparison. Separate sets of figures show the PC-space-dimensionality versus *k*-NN parameter space using either log_10_ or linear axis scales.

Available at: https://zenodo.org/records/21884011/files/SData_6.zip

**Supplementary Dataset 7**: Alternative PC1–PC2–PC3 spaces (**a–h**) and the top-ranked manifold-learning-derived LDE (**i**) from the PNG case study, shown as interactive 3D plots. For the PCA algorithm, SNP-filtering scheme, and missing-data removal regime used for each analysis, see the file names and **Supplementary** Table 1. Corresponding goodness-of-fit statistics are provided in **Supplementary** Table 1 and **Supplementary Dataset 3**. Following Bergström et al. (2017), individuals whose parents were born in the same province are colored according to their province of parental origin, defined by Papua New Guinea provincial boundaries.

Available at: https://zenodo.org/records/21884011/files/SData_7.zip

**Supplementary Dataset 8**: Comprehensive summary of manifold-learning-derived LDEs across the full parameter space of the objective-guided embedding-optimization protocol in the PNG case study. Each point in the PC-space-dimensionality versus *k*-NN biplots is colored by the Δ*r*² or Δ*ρ*² between a manifold-learning-derived embedding and the corresponding drift-normalized classic PC1–PC2–PC3 space (MAC = 1, no missing data). Each figure in this dataset consists of a collection of such biplots spanning different PC-space types, PC-space distance metrics, and manifold-learning algorithms. Individual figures are defined by the LDE-ranking objective: *F_ST_*, Hamming distance (default), GRM-derived dissimilarity, or outgroup *f*_3_-statistics, each evaluated using either Pearson’s or Spearman’s correlation. Geography-based Pearson and Spearman objectives are shown for comparison. Separate sets of figures show the PC-space-dimensionality versus *k*-NN parameter space using either log_10_ or linear axis scales.

Available at: https://zenodo.org/records/21884011/files/SData_8.zip

**Supplementary Dataset 9**: Alternative PC1–PC2–PC3 spaces (**a–h**) and the top-ranked manifold-learning-derived LDE (**i**) from the SEA case study, shown as interactive 3D plots. For the PCA algorithm, SNP-filtering scheme, and missing-data removal regime used for each analysis, see the file names and **Supplementary** Table 1. Corresponding goodness-of-fit statistics are provided in **Supplementary** Table 1 and **Supplementary Dataset 3**. Individuals are colored according to their linguistic affiliation (language family).

Available at: https://zenodo.org/records/21884011/files/SData_9.zip

**Supplementary Dataset 10**: Comprehensive summary of manifold-learning-derived LDEs across the full parameter space of the objective-guided embedding-optimization protocol in the SEA case study. Each point in the PC-space-dimensionality versus *k*-NN biplots is colored by the Δ*r*² or Δ*ρ*² between a manifold-learning-derived embedding and the corresponding drift-normalized classic PC1–PC2–PC3 space (MAC = 1, no missing data). Each figure in this dataset consists of a collection of such biplots spanning different PC-space types, PC-space distance metrics, and manifold-learning algorithms. Individual figures are defined by the LDE-ranking objective: *F_ST_*, Hamming distance (default), GRM-derived dissimilarity, or outgroup *f*_3_-statistics, each evaluated using either Pearson’s or Spearman’s correlation. Geography-based Pearson and Spearman objectives are shown for comparison. Separate sets of figures show the PC-space-dimensionality versus *k*-NN parameter space using either log_10_ or linear axis scales.

Available at: https://zenodo.org/records/21884011/files/SData_10.zip

**Supplementary Dataset 11**: PCA LDEs visualizing the projected (ancient Eurasian; **a–c**) and basis (present-day Eurasian; **d–f**) individuals separately. The PC1–PC2–PC3 spaces are presented as interactive 3D plots. A separate series of interactive plots displays the projected and basis individuals jointly (**g–i**). Shown are results from three PCA approaches: classic PC scores on data that were centered and normalized by genetic-drift dispersion prior to eigendecomposition (**a, d, g**); classic PC scores on centered data (**b, e, h**); and eigenvectors on data centered and normalized by genetic-drift dispersion (**c, f, i**). Ancient Eurasian individuals are colored according to the 23 largest IBD-sharing communities including 20 or more individuals (**Supplementary** Table 2) in all plots; present-day Eurasian individuals are colored according to the 41 largest IBD-sharing communities including 10 or more individuals (**Supplementary** Table 2) in plots **d–f** and shown as grey dots in plots **g–i**. All fidelity metrics for these 3D PC spaces are provided in **Supplementary Table 4**.

Available at: https://zenodo.org/records/21884011/files/SData_11.zip

**Supplementary Dataset 12**: Three prioritized results from the multi-step LDE-ranking procedure applied to 5,133 ancient Eurasian individuals: the final LDE (**a, d**) and two additional LDEs (**b, c**) produced under the highest *k*-NN settings among the top 20 LDEs from step 3. Each LDE is shown as an interactive 3D plot, with individuals colored by the 23 largest IBD-sharing communities (**a–c**) or by 10 geographic regions (**d**). In plot **d**, members of the CEE IBD-sharing community are shown as medium-sized points, while CEE individuals from the Roman and Migration period (eCEE) are emphasized as the largest points. The parameter settings used to generate these LDEs, along with all associated fidelity metrics, are listed in **Supplementary Table 4**.

Available at: https://zenodo.org/records/21884011/files/SData_12.zip

**Supplementary Dataset 13**: Comparison of manifold-learning optimization outcomes (using 5,133 ancient Eurasian individuals) obtained under different objectives. For each of the following key objectives, the best-performing 3D LDE is shown: 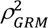 (**a**), 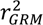 (**b**), 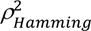 with the default missingness correction (**c**), 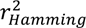 (**d**), median Mahalanobis distance among the 23 largest IBD-sharing communities (**e**), neighborhood-preservation metric at *k* = 500 (**f**) or *k* = 50 (**g**), and median Mahalanobis distance among the 21 Y-lineages with intra-lineage coalescence around 2,800 ya (**h**). The LDEs are displayed as interactive 3D plots, with individuals colored by the 23 largest IBD-sharing communities. Settings used to generate each LDE and all fidelity metrics are provided in **Supplementary Table 4**.

Available at: https://zenodo.org/records/21884011/files/SData_13.zip

**Supplementary Dataset 14**: Illustrating the LDE ranking procedure applied to the ancient Eurasian dataset: Δ*ρ*^2^ fits, Mahalanobis and Bhattacharya distances between IBD-sharing communities, and four metrics assessing how well the geometry of the (*s – 1*)-dimensional PC space (Fischer & Ma 2024) is preserved in an LDE are shown for the broader optimal region of the parameter space: 85,188 PHATE embeddings on classic PC scores. The results are displayed on logarithmic (**a**) or linear (**b**) PC space dimensionality and *k*-NN axes. In each row, the results are grouped by: (i) PCA variant – classic PCA with centering only, and classic PCA with centering plus normalization by genetic-drift dispersion; (ii) distance/affinity in the input PC space – Euclidean (ℓ₂), Manhattan (ℓ₁), and cosine.

Available at: https://zenodo.org/records/21884011/files/SData_14.zip

**Supplementary Dataset 15**: Illustrating the LDE ranking procedure applied to the present-day Eurasian dataset: Δ*ρ*^2^ fits, Mahalanobis and Bhattacharya distances between IBD-sharing communities, and four metrics assessing how well the geometry of the (*s – 1*)-dimensional PC space (Fischer & Ma 2024) is preserved in an LDE are shown for the region of the parameter space we explored: 83,544 PHATE embeddings on classic PC scores. The results are displayed on logarithmic (**a**) or linear (**b**) PC space dimensionality and *k*-NN axes. In each row, the results are grouped by: (i) PCA variant – classic PCA with centering only, and classic PCA with centering plus normalization by genetic-drift dispersion; (ii) distance/affinity in the input PC space – Euclidean (ℓ₂), Manhattan (ℓ₁), and cosine.

Available at: https://zenodo.org/records/21884011/files/SData_15.zip

**Supplementary Dataset 16**: Comparison of manifold-learning optimization outcomes (using 2,425 present-day Eurasian individuals) obtained under different objectives. For each of the following key objectives, the best-performing 3D LDE is shown: 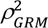 (**a**),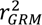 (**b**), 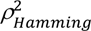 with the default missingness correction (**c**), 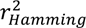 (**d**), 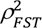 (**e**), 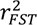 (**f**), 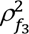(**g**), 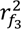 (**h**), median Mahalanobis distance among the 41 largest IBD-sharing communities (**i**), neighborhood-preservation metric at *k* = 250 (**j**) or *k* = 50 (**k**). The LDEs are displayed as interactive 3D plots, with individuals colored by the 41 largest IBD-sharing communities. Settings used to generate each LDE and all fidelity metrics are provided in **Supplementary Table 4**.

Available at: https://zenodo.org/records/21884011/files/SData_16.zip

**Supplementary Dataset 17**: Five prioritized results from the multi-step LDE-ranking procedure applied to 2,425 present-day Eurasian individuals: the final LDE (**a, f**) and the four next highest-scoring LDEs (**b–e**). Each LDE is shown as an interactive 3D plot, with individuals colored by the 41 largest IBD-sharing communities or by groups used for the *HapNe-IBD* analysis (**f**). The parameter settings used to generate these LDEs, along with all associated fidelity metrics, are listed in **Supplementary Table 4**.

Available at: https://zenodo.org/records/21884011/files/SData_17.zip

**Supplementary Dataset 18**: Optimization landscape from objective-guided manifold learning search in the ancient Eurasian case study: fits of 809,286 unique 3D LDE configurations to genetic and geographic distances. Values shown are differences in squared correlation relative to a baseline of classic 3D PC space (with data normalized by genetic-drift dispersion): 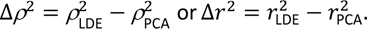. Panels **a–c** use Spearman’s rank correlation, and panels **d–f** use Pearson’s linear correlation. Each panel reports fits to pairwise distances among 5,133 individuals: (**a, d**) Hamming genetic distances; (**b, e**) GRM genetic distances; (**c, f**) great-circle geographic distances. The color scales use a power (warp-top) transform *T*(*x*) = *u*^*k*^ with *k* = 4 to enhance contrast near the best-fitting regions of the landscape. In each panel, the results are grouped by: (i) PCA variant – *smartPCA* eigenvectors, classic PC scores with centering only, and classic PC scores with centering plus normalization by genetic-drift dispersion; (ii) distance/affinity in the input PC space – Euclidean (ℓ₂), Manhattan (ℓ₁), cosine, and correlation; (iii) manifold-learning algorithm and its settings – UMAP (two *min_dist* values), a UMAP–densMAP hybrid (*dens_scale = 0.5*), densMAP (*dens_scale = 1*), and PHATE. Both the PC space dimensionality and *k*-NN axes use a logarithmic scale.

Available at: https://zenodo.org/records/21884011/files/SData_18.zip

**Supplementary Dataset 19**: The optimization landscapes from **Supplementary Dataset 18**, displayed on linear (non-logarithmic) PC space dimensionality and *k*-NN axes.

Available at: https://zenodo.org/records/21884011/files/SData_19.zip

**Supplementary Dataset 20**: Most frequent Y-lineages in the ancient Eurasian dataset at selected intra-lineage coalescence cut-offs, 2,800 ya (**a**) and 4,600 ya (**b**), visualized on the 2D isometric unfolding of the best-fitting 3D LDE. Interactive 2D plots are shown. The sets of 21 (**a**) and 31 (**b**) Y-chromosomal lineages, each represented by at least 10 individuals, were defined using a reduced subset of individuals with SNP counts above the 25th percentile; however, the plots themselves were not filtered by this SNP-count threshold. Males carrying other Y-lineages are shown as grey dots and labeled “zzz”.

Available at: https://zenodo.org/records/21884011/files/SData_20.zip

**Supplementary Dataset 21**: Quantifying Y-lineage separation in PHATE embeddings based on classic PC spaces in the ancient Eurasian case study.

Available at: https://zenodo.org/records/21884011/files/SData_21.zip

## Notes

https://zenodo.org/records/21884011

### Competing Interest Statement

The authors have declared no competing interest.

### Summary of Updates

One author name was corrected, two paragraphs were added to the Discussion, and typos were fixed.

## References

Abdi H and Williams LJ (2010). Principal component analysis. Wiley Interdiscip. Rev. Comput. Stat., 2:433–459. doi: 10.1002/wics.101.

Abegaz F, Chaichoompu K, Génin E, et al. (2018). Principals about principal components in statistical genetics. Brief. Bioinform., 20:2200–2216. doi: 10.1093/bib/bby081.

Abraham G and Inouye M (2014). Fast principal component analysis of large-scale genome-wide data. PLoS One, 9:e93766. doi: 10.1371/journal.pone.0093766.

Agdzhoyan A, Ponomarev G, Pylev V, et al. (2024). The Finnic peoples of Russia: genetic structure inferred from genome-wide and Y-chromosome data. Genes (Basel), 15:1610. doi: 10.3390/genes15121610.

Aghakhanian F, Yunus Y, Naidu R, et al. (2015). Unravelling the genetic history of Negritos and Indigenous populations of Southeast Asia. Genome Biol. Evol., 7:1206–1215. doi: 10.1093/gbe/evv065.

Aguilar-Ordoñez I, Pérez-Villatoro F, García-Ortiz H, et al. (2021). Whole genome variation in 27 Mexican indigenous populations, demographic and biomedical insights. PLoS One, 16:e0249773. doi: 10.1371/journal.pone.0249773.

Ahn J, Marron JS, Muller KM, et al. (2007). The high-dimension, low-sample-size geometric representation holds under mild conditions. Biometrika, 94:760–766. doi: 10.1093/biomet/asm050.

Aitchison J (1983). Principal component analysis of compositional data. Biometrika, 70:57–65. doi: 10.1093/biomet/70.1.57.

Aitchison J and Greenacre M (2002). Biplots of compositional data. J. R. Stat. Soc. Ser. C Appl. Stat., 51:375–392. doi: 10.1111/1467-9876.00275.

Allentoft ME, Sikora M, Refoyo-Martínez A, et al. (2024). Population genomics of post-glacial western Eurasia. Nature, 625:301–311. doi: 10.1038/s41586-023-06865-0.

Altman N and Krzywinski M (2018). The curse(s) of dimensionality. Nat. Methods, 15:399–400. doi: 10.1038/s41592-018-0019-x.

Arango-Isaza E, Capodiferro MR, Aninao MJ, et al. (2023). The genetic history of the Southern Andes from present-day Mapuche ancestry. Curr. Biol., 33:2602–2615.e5. doi: 10.1016/j.cub.2023.05.013.

Armstrong G, Rahman G, Martino C, et al. (2022). Applications and comparison of dimensionality reduction methods for microbiome data. Front. Bioinform., 2:821861. doi: 10.3389/fbinf.2022.821861.

Audzijonyte A and Vrijenhoek RC (2010). When gaps really are gaps: statistical phylogeography of hydrothermal vent invertebrates. Evolution, 64:2369–2384. doi: 10.1111/j.1558-5646.2010.00987.x.

Barlow A, Hartmann S, Gonzalez J, et al. (2020). Consensify: a method for generating pseudohaploid genome sequences from palaeogenomic datasets with reduced error rates. Genes (Basel), 11:50. doi: 10.3390/genes11010050.

Battey CJ, Coffing GC and Kern AD (2021). Visualizing population structure with variational autoencoders. G3 Genes Genomes Genet., 11:jkaa036. doi: 10.1093/g3journal/jkaa036.

Baumdicker F, Bisschop G, Goldstein D, et al. (2022). Efficient ancestry and mutation simulation with msprime 1.0. Genetics, 220:iyab229. doi: 10.1093/genetics/iyab229.

Bergström A, McCarthy SA, Hui R, et al. (2017). A Neolithic expansion, but strong genetic structure, in the independent history of New Guinea. Science, 357:1160–1163. doi: 10.1126/science.aan3842.

Bionda A, Negro A, Floridia V, et al. (2026) Genomic insights into the recent evolution and biodiversity of Italian sheep breeds. Mamm. Genome, 37:5. doi: 10.1007/s00335-025-10170-8.

Botigué LR, Song S, Scheu A, et al. (2017). Ancient European dog genomes reveal continuity since the Early Neolithic. Nat. Commun., 8:16082. doi: 10.1038/ncomms16082.

Bradburd GS and Ralph PL (2019). Spatial population genetics: it’s about time. Annu. Rev. Ecol. Evol. Syst., 50:427–449. doi: 10.1146/annurev-ecolsys-110316-022659.

Browning BL, Tian X, Zhou Y, et al. (2021). Fast two-stage phasing of large-scale sequence data. Am. J. Hum. Genet., 108:1880–1890. doi: 10.1016/j.ajhg.2021.08.005.

Busby GBJ, Hellenthal G, Montinaro F, et al. (2015). The role of recent admixture in forming the contemporary West Eurasian genomic landscape. Curr. Biol., 25:2518–2526. doi: 10.1016/j.cub.2015.08.007.

Caliński T and Harabasz J (1974). A dendrite method for cluster analysis. Commun. Stat., 3:1–27. doi: 10.1080/03610927408827101.

Cashman D, Keller M, Jeon H, et al. (2025). A critical analysis of the usage of dimensionality reduction in four domains. IEEE Trans. Vis. Comput. Graph., 31:9405–9423. doi: 10.1109/TVCG.2025.3567989.

Castro e Silva MA, Ferraz T, Couto-Silva CM, et al. (2022). Population histories and genomic diversity of South American natives. Mol. Biol. Evol., 39:msab339. doi: 10.1093/molbev/msab339.

Castro e Silva MA, Nunes K, Ribeiro MR, et al. (2026). The evolutionary history and unique genetic diversity of Indigenous Americans. Nature, 653:134–145. doi: 10.1038/s41586-026-10406-w.

Cattell RB (1966). The scree test for the number of factors. Multivar. Behav. Res., 1:245–276. doi: 10.1207/s15327906mbr0102_10.

Chang CC, Chow CC, Tellier LC, et al. (2015). Second-generation PLINK: rising to the challenge of larger and richer datasets. GigaScience, 4:7. doi: 10.1186/s13742-015-0047-8.

Changmai P, Jaisamut K, Kampuansai J, et al. (2022). Indian genetic heritage in Southeast Asian populations. PLoS Genet., 18:e1010036. doi: 10.1371/journal.pgen.1010036.

Changmai P, Phongbunchoo Y, Kočí J, Flegontov P. (2023). Reanalyzing the genetic history of Kra-Dai speakers from Thailand and new insights into their genetic interactions beyond Mainland Southeast Asia. Sci. Rep., 13:8371. doi: 10.1038/s41598-023-35507-8.

Chari T and Pachter L (2023). The specious art of single-cell genomics. PLoS Comput. Biol., 19:e1011288. doi: 10.1371/journal.pcbi.1011288.

Chobanov T and Stamov S (2025). Slavs in the closet: computational genomic analysis reveals cryptic Slavic signatures in the Avar Khaganate and their contribution to medieval Croatian population formation. Front. Genet., 16:1610942. doi: 10.3389/fgene.2025.1610942.

Crosslin DR, Tromp G, Burt A, et al. (2014). Controlling for population structure and genotyping platform bias in the eMERGE multi-institutional biobank linked to electronic health records. Front. Genet., 5:352. doi: 10.3389/fgene.2014.00352.

Cutler A and Breiman L (1994). Archetypal analysis. Technometrics, 36:338–347. doi: 10.1080/00401706.1994.10485840.

Davies DL and Bouldin DW (1979). A cluster separation measure. IEEE Trans. Pattern Anal. Mach. Intell., PAMI-1:224–227. doi: 10.1109/TPAMI.1979.4766909.

de Gennaro L, Molinaro L, Raveane A, et al. (2025). PANE: fast and reliable ancestral reconstruction on ancient genotype data with non-negative least square and principal component analysis. Genome Biol., 26:29. doi: 10.1186/s13059-025-03491-z.

de Jong MJ, Anaya G, Niamir A, et al. (2025). Red deer resequencing reveals the importance of sex chromosomes for reconstructing Late Quaternary events. Mol. Biol. Evol., 42:msaf031. doi: 10.1093/molbev/msaf031.

Der Sarkissian C, Ermini L, Schubert M, et al. (2015). Evolutionary genomics and conservation of the endangered Przewalski’s horse. Curr. Biol., 25:2577–2583. doi: 10.1016/j.cub.2015.08.032.

Diaconis P, Goel S and Holmes S (2008). Horseshoes in multidimensional scaling and local kernel methods. Ann. Appl. Stat., 2:777–807. doi: 10.1214/08-AOAS165.

Diaz-Papkovich A, Anderson-Trocmé L, Ben-Eghan C, et al. (2019). UMAP reveals cryptic population structure and phenotype heterogeneity in large genomic cohorts. PLoS Genet., 15:e1008432. doi: 10.1371/journal.pgen.1008432.

Diaz-Papkovich A, Anderson-Trocmé L and Gravel S (2021). A review of UMAP in population genetics. J. Hum. Genet., 66:85–91. doi: 10.1038/s10038-020-00851-4.

Diaz-Papkovich A, Zabad S, Snell H, et al. (2026). Topological stratification of continuous genetic variation in large biobanks. PLoS Genet., 22:e1012068. doi: 10.1371/journal.pgen.1012068.

Dutrow EV, Serpell JA, Ostrander EA. (2022). Domestic dog lineages reveal genetic drivers of behavioral diversification. Cell, 185:4737–4755. doi: 10.1016/j.cell.2022.11.003.

Estavoyer M and François O (2022). Theoretical analysis of principal components in an umbrella model of intraspecific evolution. Theor. Popul. Biol., 148:11–21. doi: 10.1016/j.tpb.2022.08.002.

Fernandes DM, Sirak KA, Ringbauer H, et al. (2021). A genetic history of the pre-contact Caribbean. Nature, 590:103–110. doi: 10.1038/s41586-020-03053-2.

Fischer J and Ma R (2024). Sailing in high-dimensional spaces: low-dimensional embeddings through angle preservation. arXiv:2406.09876. doi: 10.48550/arXiv.2406.09876.

Flegontov P, Altınışık NE, Changmai P, et al. (2019). Palaeo-Eskimo genetic ancestry and the peopling of Chukotka and North America. Nature, 570:236–240. doi: 10.1038/s41586-019-1251-y.

Flegontov P, Işıldak U, Maier R, et al. (2023). Modeling of African population history using *f*-statistics is biased when applying all previously proposed SNP ascertainment schemes. PLoS Genet., 19:e1010931. doi: 10.1371/journal.pgen.1010931.

Flegontova O, Işıldak U, Yüncü E, et al. (2025). Performance of *qpAdm*-based screens for genetic admixture on graph-shaped histories and stepping stone landscapes. Genetics, 230:iyaf047. doi: 10.1093/genetics/iyaf047.

Fournier R, Pearson Fulton A and Reich D. (2025). A SNP panel for co-analysis of capture and shotgun ancient DNA data. bioRxiv:2025.07.30.667733. doi: 10.1101/2025.07.30.667733.

François O and Jay F (2020). Factor analysis of ancient population genomic samples. Nat. Commun., 11:4661. doi: 10.1038/s41467-020-18335-6.

Frichot E, Schoville S, Bouchard G, et al. (2012). Correcting principal component maps for effects of spatial autocorrelation in population genetic data. Front. Genet., 3:254. doi: 10.3389/fgene.2012.00254.

Fu Q, Posth C, Hajdinjak M, et al. (2016). The genetic history of Ice Age Europe. Nature, 534:200–205. doi: 10.1038/nature17993.

Gildenblat J and Pahnke J (2026). Dimensionality reduction with strong global structure preservation. Pattern Anal. Appl., 29:38. doi: 10.1007/s10044-025-01585-9.

Greenacre M, Groenen PJF, Hastie T, et al. (2022). Principal component analysis. Nat. Rev. Methods Primers, 2:100. doi: 10.1038/s43586-022-00184-w.

Gretzinger J, Biermann F, Mager H, et al. (2025). Ancient DNA connects large-scale migration with the spread of Slavs. Nature, 646:384–393. doi: 10.1038/s41586-025-09437-6.

Grinde KE, Browning BL, Reiner AP, et al. (2024). Adjusting for principal components can induce collider bias in genome-wide association studies. PLoS Genet., 20:e1011242. doi: 10.1371/journal.pgen.1011242.

Haak W, Lazaridis I, Patterson N, et al. (2015). Massive migration from the steppe was a source for Indo-European languages in Europe. Nature, 522:207–211. doi: 10.1038/nature14317.

Hall P, Marron JS and Neeman A (2005). Geometric representation of high dimension, low sample size data. J. R. Stat. Soc. Ser. B Stat. Methodol., 67:427–444. doi: 10.1111/j.1467-9868.2005.00510.x.

Heiser CN and Lau KS (2020). A quantitative framework for evaluating single-cell data structure preservation by dimensionality reduction techniques. Cell Rep., 31:107576. doi: 10.1016/j.celrep.2020.107576.

Hellenthal G, Busby GBJ, Band G, et al. (2014). A genetic atlas of human admixture history. Science, 343:747– 751. doi: 10.1126/science.1243518.

Herrando-Pérez S, Tobler R and Huber CD (2021). *smartsnp*, an R package for fast multivariate analyses of big genomic data. Methods Ecol. Evol., 12:2084–2093. doi: 10.1111/2041-210X.13684.

Hou M, Huang Y, Ma L, et al. (2015). Compositional analysis of ternary and binary chemical mixtures by surface-enhanced Raman scattering at trace levels. Nanoscale Res. Lett., 10:437. doi: 10.1186/s11671-015-1142-6.

House GL and Hahn MW (2017). Evaluating methods to visualize patterns of genetic differentiation on a landscape. Mol. Ecol. Resour., 18:448–460. doi: 10.1111/1755-0998.12747.

Huckins LM, Boraska V, Franklin CS, et al. (2014). Using ancestry-informative markers to identify fine structure across 15 populations of European origin. Eur. J. Hum. Genet., 22:1190–1200. doi: 10.1038/ejhg.2014.1.

Hui R, D’Atanasio E, Cassidy LM, et al. (2020). Evaluating genotype imputation pipeline for ultra-low coverage ancient genomes. Sci. Rep., 10:18542. doi: 10.1038/s41598-020-75387-w.

Jackson DA (1993). Stopping rules in principal components analysis: a comparison of heuristical and statistical approaches. Ecology, 74:2204–2214. doi: 10.2307/1939574.

Jacomy M, Venturini T, Heymann S, et al. (2014). *ForceAtlas2*, a continuous graph layout algorithm for handy network visualization designed for the *Gephi* software. PLoS One, 9:e98679. doi: 10.1371/journal.pone.0098679.

Jeong C, Balanovsky O, Lukianova E, et al. (2019). The genetic history of admixture across inner Eurasia. *Nat*. Ecol. Evol., 3:966–976. doi: 10.1038/s41559-019-0878-2.

Jung S and Marron JS (2009). PCA consistency in high dimension, low sample size context. Ann. Stat., 37:4104– 4130. doi: 10.1214/09-AOS709.

Jung S, Sen A and Marron JS (2012). Boundary behavior in high dimension, low sample size asymptotics of PCA. J. Multivar. Anal., 109:190–203. doi: 10.1016/j.jmva.2012.03.005.

Khrunin AV, Khokhrin DV, Filippova IN, et al. (2013). A genome-wide analysis of populations from European Russia reveals a new pole of genetic diversity in northern Europe. PLoS One, 8:e58552. doi: 10.1371/journal.pone.0058552.

Kim H, Kim S, Yu AR, et al. (2026). Ancient East Asian dog lineage is revealed by genome of ancient Korean dogs. PLoS One, 21:e0346864. doi: 10.1371/journal.pone.0346864.

Kobak D and Linderman GC (2021). Initialization is critical for preserving global data structure in both t-SNE and UMAP. Nat. Biotechnol., 39:156–157. doi: 10.1038/s41587-020-00809-z.

Kohli D, Cloninger A and Mishne G (2021). LDLE: low distortion local eigenmaps. J. Mach. Learn. Res., 22:1–64.

Korem Y, Szekely P, Hart Y, et al. (2015). Geometry of the gene expression space of individual cells. PLoS Comput. Biol., 11:e1004224. doi: 10.1371/journal.pcbi.1004224.

Kutanan W, Liu D, Kampuansai J, et al. (2021). Reconstructing the human genetic history of Mainland Southeast Asia: Insights from genome-wide data from Thailand and Laos. Mol. Biol. Evol., 38:3459–3477. doi: 10.1093/molbev/msab124.

Lamnidis TC, Majander K, Jeong C, et al. (2018). Ancient Fennoscandian genomes reveal origin and spread of Siberian ancestry in Europe. Nat. Commun., 9:5018. doi: 10.1038/s41467-018-07483-5.

Lazaridis I, Nadel D, Rollefson G, et al. (2016). Genomic insights into the origin of farming in the ancient Near East. Nature, 536:419–424. doi: 10.1038/nature19310.

Lazaridis I, Patterson N, Anthony D, et al. (2025). The genetic origin of the Indo-Europeans. Nature, 639:132–142. doi: 10.1038/s41586-024-08531-5.

Lee S, Zou F and Wright FA (2010). Convergence and prediction of principal component scores in high-dimensional settings. Ann. Stat., 38:3605–3629. doi: 10.1214/10-AOS821.

Lever J, Krzywinski M and Altman N (2017). Principal component analysis. Nat. Methods, 14:641–642. doi: 10.1038/nmeth.4346.

Li X, Wang Z, Zhu M, et al. (2025). Genomic insights into post-domestication expansion and selection of body size in ponies. Adv. Sci. (Weinh), 12:e2413023. doi: 10.1002/advs.202413023.

Linck E and Battey CJ (2019). Minor allele frequency thresholds strongly affect population structure inference with genomic datasets. Mol. Ecol. Resour., 19:639–647. doi: 10.1111/1755-0998.12995.

Lindo J, Haas R, Hofman C, et al. (2018). The genetic prehistory of the Andean highlands 7000 years BP though European contact. Sci. Adv., 4:eaau4921. doi: 10.1126/sciadv.aau4921.

Lipson M, Cheronet O, Mallick S, et al. (2018). Ancient genomes document multiple waves of migration in Southeast Asian prehistory. Science, 361:92–95. doi: 10.1126/science.aat3188.

Liu LT, Dobriban E and Singer A (2017). ePCA: High Dimensional Exponential Family PCA. arXiv:1611.05550. doi: 10.48550/arXiv.1611.05550.

Liu Z, Wang N, Su Y, et al. (2024). Grapevine pangenome facilitates trait genetics and genomic breeding. Nat. Genet., 56:2804–2814. doi: 10.1038/s41588-024-01967-5.

Liu Z, Ma R and Zhong Y (2025). Assessing and improving reliability of neighbor embedding methods: a map-continuity perspective. Nat. Commun., 16:5037. doi: 10.1038/s41467-025-60434-9.

Louis M, Korlević P, Nykänen M, et al. (2023). Ancient dolphin genomes reveal rapid repeated adaptation to coastal waters. Nat. Commun., 14:4020. doi: 10.1038/s41467-023-39532-z.

Luu K, Bazin E and Blum MGB (2017). *pcadapt*: an R package to perform genome scans for selection based on principal component analysis. Mol. Ecol. Resour., 17:67–77. doi: 10.1111/1755-0998.12592.

Ma J and Amos CI (2012). Principal components analysis of population admixture. PLoS One, 7:e40115. doi: 10.1371/journal.pone.0040115.

Ma S and Shi G (2020). On rare variants in principal component analysis of population stratification. BMC Genet., 21:34. doi: 10.1186/s12863-020-0833-x.

Maier R, Flegontov P, Flegontova O, et al. (2023). On the limits of fitting complex models of population history to *f*-statistics. eLife, 12:e85492. doi: 10.7554/eLife.85492.

Mallick S, Li H, Lipson M, et al. (2016). The Simons Genome Diversity Project: 300 genomes from 142 diverse populations. Nature, 538:201–206. doi: 10.1038/nature18964.

Mallick S, Micco A, Mah M, et al. (2024). The Allen Ancient DNA Resource (AADR): a curated compendium of ancient human genomes. Sci. Data, 11:182. doi: 10.1038/s41597-024-03031-7.

Maravall-López J, Motti JMB, Pastor N, et al. (2025). Eight millennia of continuity of a previously unknown lineage in Argentina. Nature, 649:647–656. doi: 10.1038/s41586-025-09731-3.

Martiniano R, De Sanctis B, Hallast P, et al. (2022). Placing ancient DNA sequences into reference phylogenies. Mol. Biol. Evol., 39:msac017. doi: 10.1093/molbev/msac017.

Mathieson I and McVean G (2012). Differential confounding of rare and common variants in spatially structured populations. Nat. Genet., 44:243–246. doi: 10.1038/ng.1074.

McInnes L, Healy J and Melville J (2018). UMAP: uniform manifold approximation and projection for dimension reduction. arXiv:1802.03426. doi: 10.48550/arXiv.1802.03426.

McRae BH (2006). Isolation by resistance. Evolution, 60:1551–1561. doi: 10.1111/j.0014-3820.2006.tb00500.x.

McVean G (2009). A genealogical interpretation of principal components analysis. PLoS Genet., 5:e1000686. doi: 10.1371/journal.pgen.1000686.

Meilă M and Zhang H (2024). Manifold learning: what, how, and why. Annu. Rev. Stat. Appl., 11:393–417. doi: 10.1146/annurev-statistics-040522-115238.

Meirmans PG (2015). Seven common mistakes in population genetics and how to avoid them. Mol. Ecol., 24:3223–3231. doi: 10.1111/mec.13243.

Meisner J, Liu S, Huang M, et al. (2021). Large-scale inference of population structure in presence of missingness using PCA. Bioinformatics, 37:1868–1875. doi: 10.1093/bioinformatics/btab027.

Moon KR, van Dijk D, Wang Z, et al. (2019). Visualizing structure and transitions in high-dimensional biological data. Nat. Biotechnol., 37:1482–1492. doi: 10.1038/s41587-019-0336-3.

Moreno-Mayar JV, Vinner L, de Barros Damgaard P, et al. (2018). Early human dispersals within the Americas. Science, 362:eaav2621. doi: 10.1126/science.aav2621.

Muktupavela RA, Petr M, Ségurel L, et al. (2022). Modeling the spatiotemporal spread of beneficial alleles using ancient genomes. eLife, 11:e73767. doi: 10.7554/eLife.73767.

Narayan A, Berger B and Cho H (2021). Assessing single-cell transcriptomic variability through density-preserving data visualization. Nat. Biotechnol., 39:765–774. doi: 10.1038/s41587-020-00801-7.

Nguyen LH and Holmes S (2019). Ten quick tips for effective dimensionality reduction. PLoS Comput. Biol., 15:e1006907. doi: 10.1371/journal.pcbi.1006907.

Novembre J and Stephens M (2008). Interpreting principal component analyses of spatial population genetic variation. Nat. Genet., 40:646–649. doi: 10.1038/ng.139.

Olalde I, Brace S, Allentoft ME, et al. (2018). The Beaker phenomenon and the genomic transformation of northwest Europe. Nature, 555:190–196. doi: 10.1038/nature25738.

Olalde I, Carrión P, Mikić I, et al. (2023). A genetic history of the Balkans from Roman frontier to Slavic migrations. Cell, 186:5472–5485. doi: 10.1016/j.cell.2023.10.018.

Patterson N, Price AL and Reich D (2006). Population structure and eigenanalysis. PLoS Genet., 2:e190. doi: 10.1371/journal.pgen.0020190.

Patterson N, Moorjani P, Luo Y, et al. (2012). Ancient admixture in human history. Genetics, 192:1065–1093. doi: 10.1534/genetics.112.145037.

Paul D (2007). Asymptotics of sample eigenstructure for a large dimensional spiked covariance model. Stat. Sin., 17:1617–1642.

Peltola S, Majander K, Makarov N, et al. (2023). Genetic admixture and language shift in the medieval Volga-Oka interfluve. Curr. Biol., 33:174–182.e10. doi: 10.1016/j.cub.2022.11.036.

Peres-Neto PR, Jackson DA and Somers KM (2005). How many principal components? Stopping rules for determining the number of non-trivial axes revisited. Comput. Stat. Data Anal., 49:974–997. doi: 10.1016/j.csda.2004.06.015.

Peter BM (2022). A geometric relationship of F2, F3 and F4-statistics with principal component analysis. Philos. Trans. R. Soc. Lond. B Biol. Sci., 377:20200413. doi: 10.1098/rstb.2020.0413.

Petr M, Haller BC, Ralph PL, et al. (2023). *slendr*: a framework for spatio-temporal population genomic simulations on geographic landscapes. Peer Community J., 3:e121. doi: 10.24072/pcjournal.354.

Pickrell JK and Pritchard JK (2012). Inference of population splits and mixtures from genome-wide allele frequency data. PLoS Genet., 8:e1002967. doi: 10.1371/journal.pgen.1002967.

Podani J and Miklós I (2002). Resemblance coefficients and the horseshoe effect in principal coordinates analysis. Ecology, 83:3331–3343. doi: 10.1890/0012-9658(2002)083[3331:RCATHE]2.0.CO;2.

Posth C, Yu H, Ghalichi A, et al. (2023). Palaeogenomics of Upper Palaeolithic to Neolithic European hunter-gatherers. Nature, 615:117–126. doi: 10.1038/s41586-023-05726-0.

Price AL, Patterson NJ, Plenge RM, et al. (2006). Principal components analysis corrects for stratification in genome-wide association studies. Nat. Genet., 38:904–909. doi: 10.1038/ng1847.

Privé F, Luu K, Blum MGB, et al. (2020). Efficient toolkit implementing best practices for principal component analysis of population genetic data. Bioinformatics, 36:4449–4457. doi: 10.1093/bioinformatics/btaa520.

Pélissié B, Chen YH, Cohen ZP, et al. (2022). Genome resequencing reveals rapid, repeated evolution in the Colorado potato beetle. Mol. Biol. Evol., 39:msac016. doi: 10.1093/molbev/msac016.

Racimo F, Sikora M, Vander Linden M, et al. (2020). The spatiotemporal spread of human migrations during the European Holocene. Proc. Natl. Acad. Sci. U.S.A., 117:8989–9000. doi: 10.1073/pnas.1920051117.

Ribeiro-dos-Santos AM, Vidal AF, Vinasco-Sandoval T, et al. (2020). Exome sequencing of native populations from the Amazon reveals patterns on the peopling of South America. Front. Genet., 11:548507. doi: 10.3389/fgene.2020.548507.

Ringbauer H, Huang Y, Akbari A, et al. (2024). Accurate detection of identity-by-descent segments in human ancient DNA. Nat. Genet., 56:143–151. doi: 10.1038/s41588-023-01582-w.

Rohland N, Mallick S, Mah M, et al. (2022). Three assays for in-solution enrichment of ancient human DNA at more than a million SNPs. Genome Res., 32:2068–2078. doi: 10.1101/gr.276728.122.

Rousseeuw PJ (1987). Silhouettes: a graphical aid to the interpretation and validation of cluster analysis. J. Comput. Appl. Math., 20:53–65. doi: 10.1016/0377-0427(87)90125-7.

Sakaue S, Hirata J, Kanai M, et al. (2020). Dimensionality reduction reveals fine-scale structure in the Japanese population with consequences for polygenic risk prediction. Nat. Commun., 11:1569. doi: 10.1038/s41467-020-15194-z.

Salova J, Vyazov L and Beneš J (2024). When barley and wheat meet millet: cereal cultivation patterns in the forest and forest-steppe of Eastern Europe from the Early Iron Age to the Early Middle Ages. Interdiscip. Archaeol. Nat. Sci. Archaeol., 15:167–181. doi: 10.24916/iansa.2024.2.3.

Schmid C and Schiffels S (2023). Estimating human mobility in Holocene Western Eurasia with large-scale ancient genomic data. Proc. Natl. Acad. Sci. U.S.A., 120:e2218375120. doi: 10.1073/pnas.2218375120.

Severson AL, Byrd BF, Mallott EK, et al. (2022). Ancient and modern genomics of the Ohlone Indigenous population of California. Proc. Natl. Acad. Sci. U.S.A., 119:e2111533119. doi: 10.1073/pnas.2111533119.

Shirk AJ, Landguth EL and Cushman SA (2017). A comparison of individual-based genetic distance metrics for landscape genetics. Mol. Ecol. Resour., 17:1308–1317. doi: 10.1111/1755-0998.12684.

Sikora M, Pitulko VV, Sousa VC, et al. (2019). The population history of northeastern Siberia since the Pleistocene. Nature, 570:182–188. doi: 10.1038/s41586-019-1279-z.

Siu H, Jin L, Xiong M. (2012). Manifold learning for human population structure studies. PLoS One, 7:e29901. doi: 10.1371/journal.pone.0029901.

Skoglund P, Ersmark E, Palkopoulou E, et al. (2015). Ancient wolf genome reveals an early divergence of domestic dog ancestors and admixture into high-latitude breeds. Curr. Biol., 25:1515–1519. doi: 10.1016/j.cub.2015.04.019.

Sousa da Mota B, Rubinacci S, Cruz Dávalos DI, et al. (2023). Imputation of ancient human genomes. Nat. Commun., 14:3660. doi: 10.1038/s41467-023-39202-0.

Szekely P, Korem Y, Moran U, et al. (2015). The mass-longevity triangle: Pareto optimality and the geometry of life-history trait space. PLoS Comput. Biol., 11:e1004524. doi: 10.1371/journal.pcbi.1004524.

Tao L, Yuan H, Zhu K, et al. (2023). Ancient genomes reveal millet farming-related demic diffusion from the Yellow River into southwest China. Curr. Biol., 33:4995–5002. doi: 10.1016/j.cub.2023.09.055.

Tenenbaum JB, de Silva V and Langford JC (2000). A global geometric framework for nonlinear dimensionality reduction. Science, 290:2319–2323. doi: 10.1126/science.290.5500.2319.

The International HapMap Consortium (2007). A second generation human haplotype map of over 3.1 million SNPs. Nature, 449:851–861. doi: 10.1038/nature06258.

Traag VA, Waltman L and van Eck NJ (2019). From Louvain to Leiden: guaranteeing well-connected communities. Sci. Rep., 9:5233. doi: 10.1038/s41598-019-41695-z.

Tyrmi JS, Vuosku J, Acosta JJ, et al. (2020). Genomics of clinal local adaptation in Pinus sylvestris under continuous environmental and spatial genetic setting. G3 Genes Genomes Genet., 10:2683–2696. doi: 10.1534/g3.120.401285.

Ubbens J, Feldmann MJ, Stavness I, et al. (2022). Quantitative evaluation of nonlinear methods for population structure visualization and inference. G3 Genes Genomes Genet., 12:jkac191. doi: 10.1093/g3journal/jkac191.

van der Maaten L and Hinton G (2008). Visualizing data using t-SNE. J. Mach. Learn. Res., 9:2579–2605.

Watson ER, Mora A, Taherian Fard A, et al. (2022). How does the structure of data impact cell-cell similarity? Evaluating how structural properties influence the performance of proximity metrics in single cell RNA-seq data. Brief. Bioinform., 23:bbac387. doi: 10.1093/bib/bbac387.

Wisser RJ, Fang Z, Holland JB, et al. (2019). The Genomic basis for short-term evolution of environmental adaptation in maize. Genetics, 213:1479–1494. doi: 10.1534/genetics.119.302780.

Vyazov LA, Flegontov P, Flegontova O, Cooper LR, Kedem I, Kandell T, Changmai T, Atanasoska-Vrhel N, Kočí J, Kassian AS, Ringbauer H, Birkina N, Đukić K, Hajdu T, Kitov E, Novak M, Petrova D, Šefčáková A, Simalcsik A, Soficaru A, Stashenkov D, Teschler-Nicola M, Valiev R, Voroniatov S, Zagorc B, Batieva EF, Bugarski I, Carić M, Dizdar M, Dugonjić A, Farkaš Z, Jelínek P, Jončić N, Keckarević D, Khokhlov A, Kitova A, Klyukoit A, Kozubová A, Krznar S, Mikašinović V, Musilová M, Nechiporuk A, Rapan Papeša A, Resutík B, Rimpf A, Serykh D, Tresić Pavičić D, Veselinov D, Vulić H, Živković M and David Reich. Slavs before the Slavs: deeply optimized manifold learning and long autosomal haplotypes reveal the origins and early dispersal of the Slavic ancestry. Manuscript in preparation.

Xia L, Lee C and Li JJ (2024). Statistical method scDEED for detecting dubious 2D single-cell embeddings and optimizing t-SNE and UMAP hyperparameters. Nat. Commun., 15:1753. doi: 10.1038/s41467-024-45891-y.

Yang J, Lee SH, Goddard ME, et al. (2011). GCTA: a tool for genome-wide complex trait analysis. Am. J. Hum. Genet., 88:76–82. doi: 10.1016/j.ajhg.2010.11.011.

Zeng TC, Vyazov LA, Kim A, et al. (2025). Ancient DNA reveals the prehistory of the Uralic and Yeniseian peoples. Nature, 644:122–132. doi: 10.1038/s41586-025-09189-3.

Zhou Y, Browning SR and Browning BL (2020). A fast and simple method for detecting identity-by-descent segments in large-scale data. Am. J. Hum. Genet., 106:426–437. doi: 10.1016/j.ajhg.2020.02.010.

