## Supplementary figures and images for "Sparse sampling and rare-variant depletion distort PCA visualizations of population structure: recovery with objective-guided manifold learning"

### Supplementary Figure 1

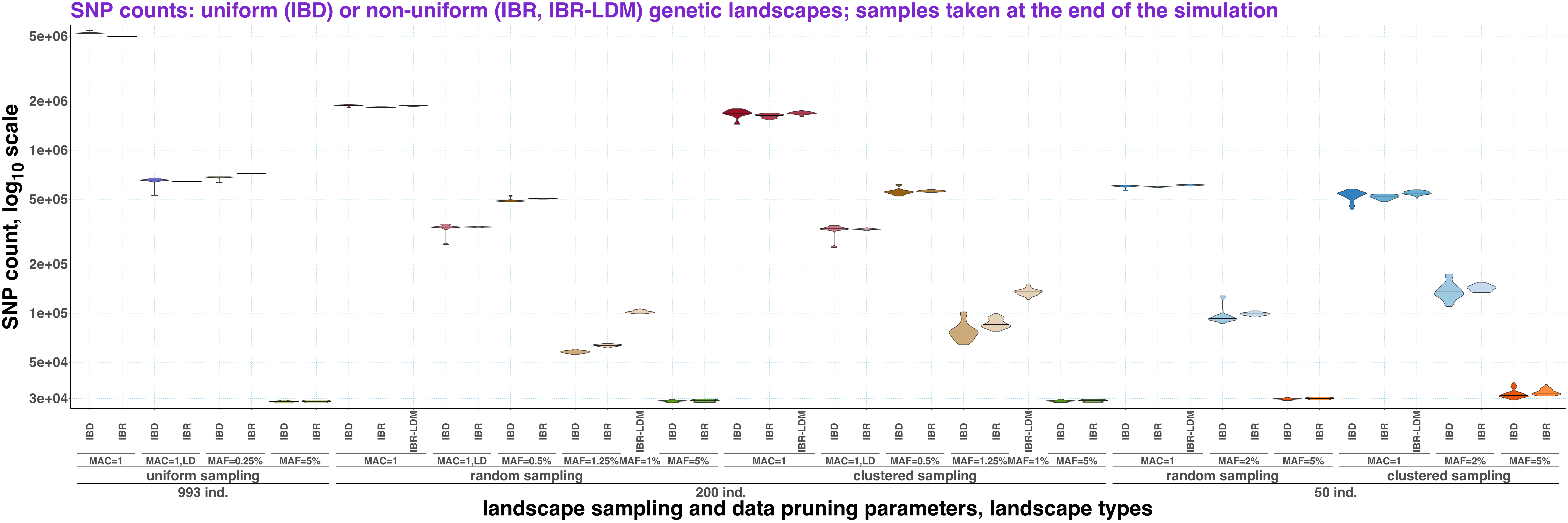

### Supplementary Figure 7a

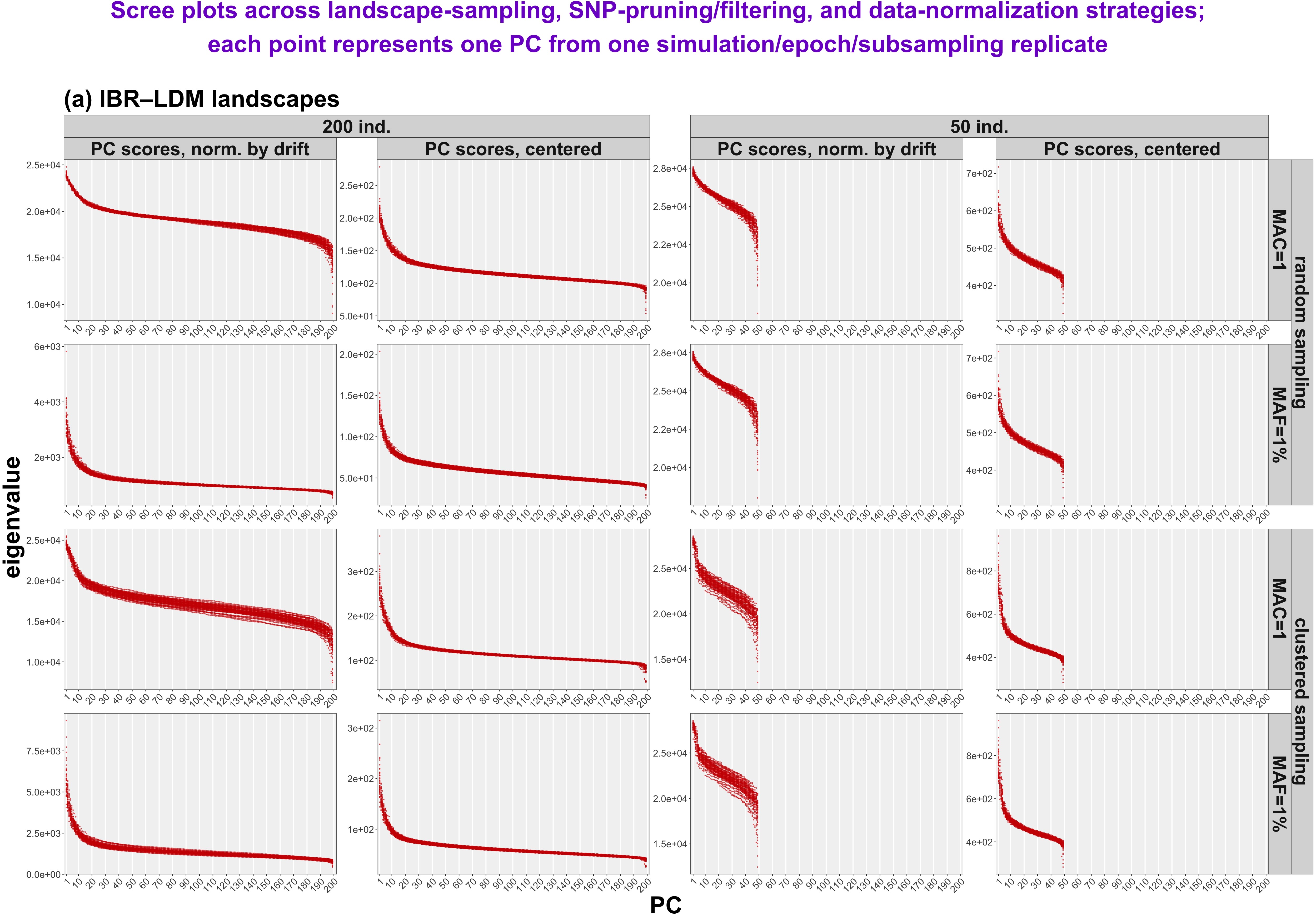

### Supplementary Figure 7b-c

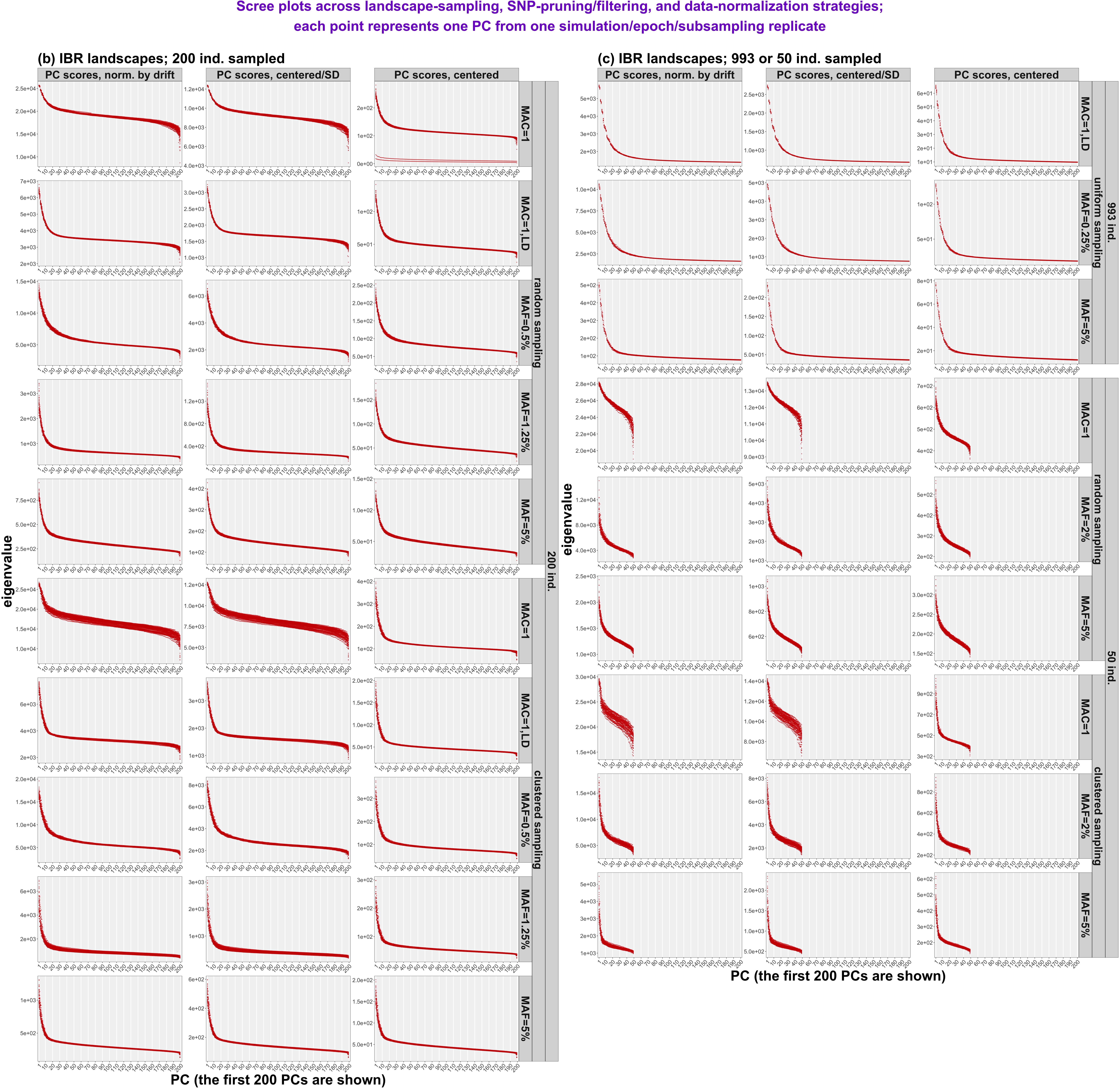

### Supplementary Figure 9a

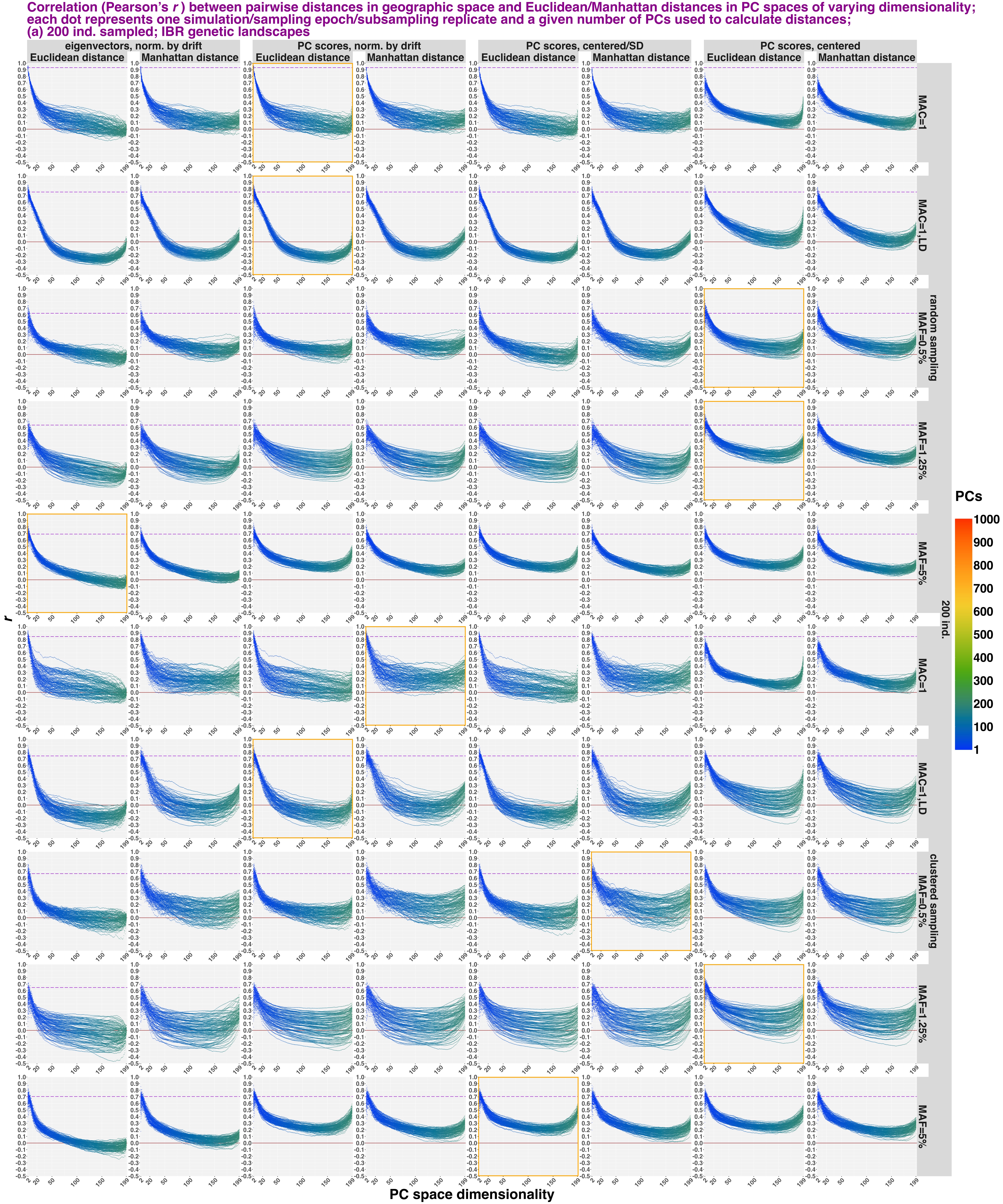

### Supplementary Figure 9b

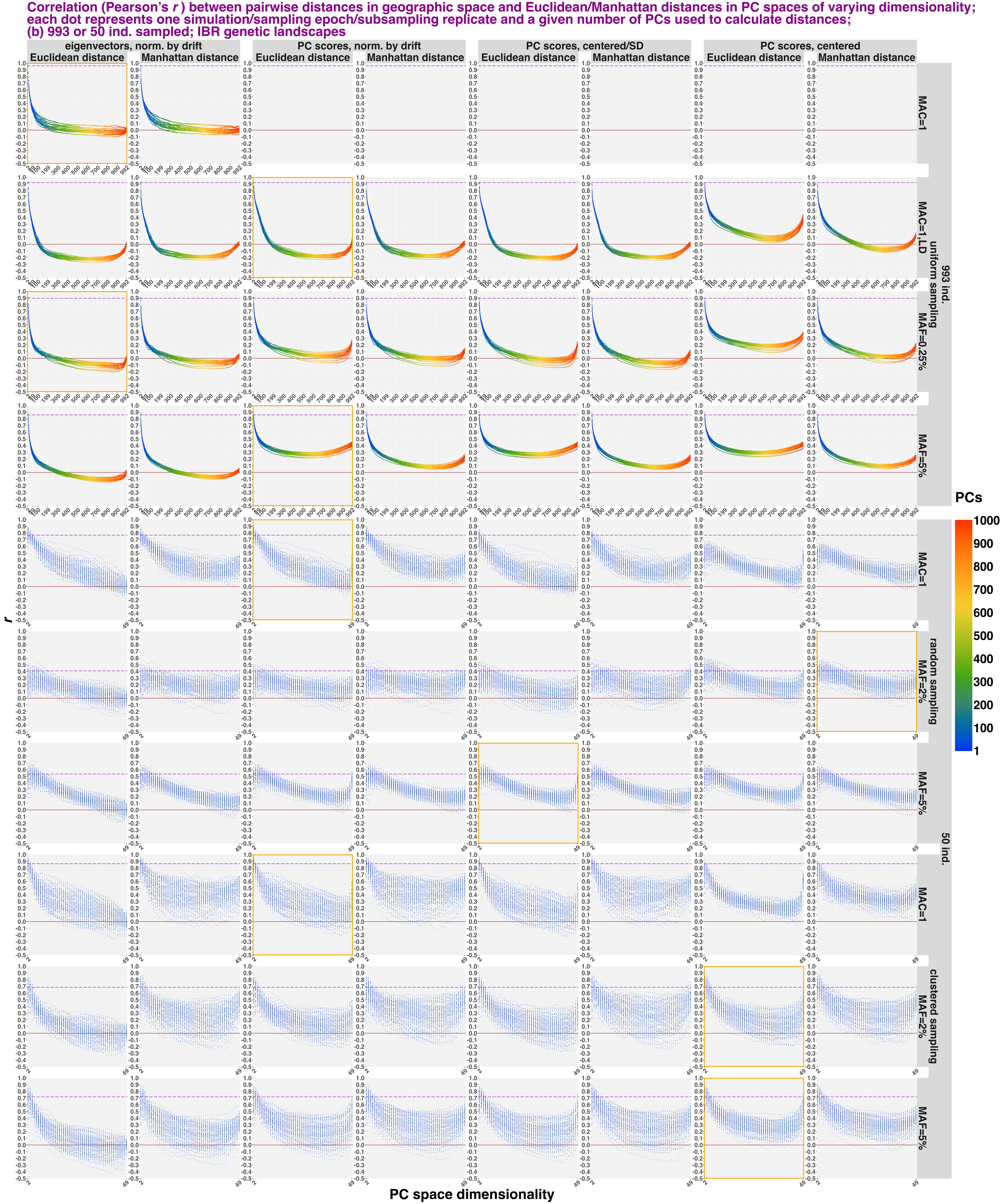

### Supplementary Figure 9c

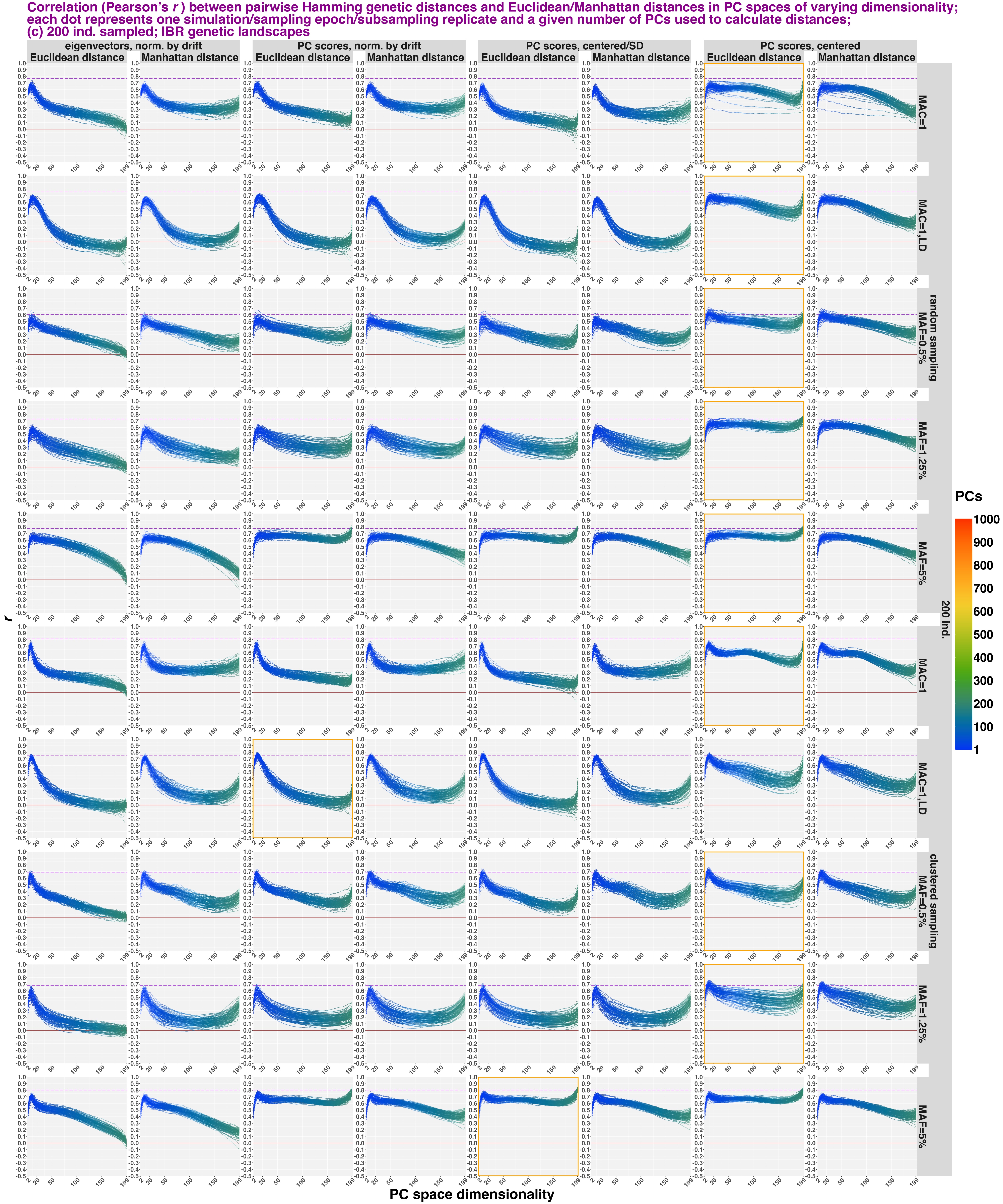

### Supplementary Figure 9d

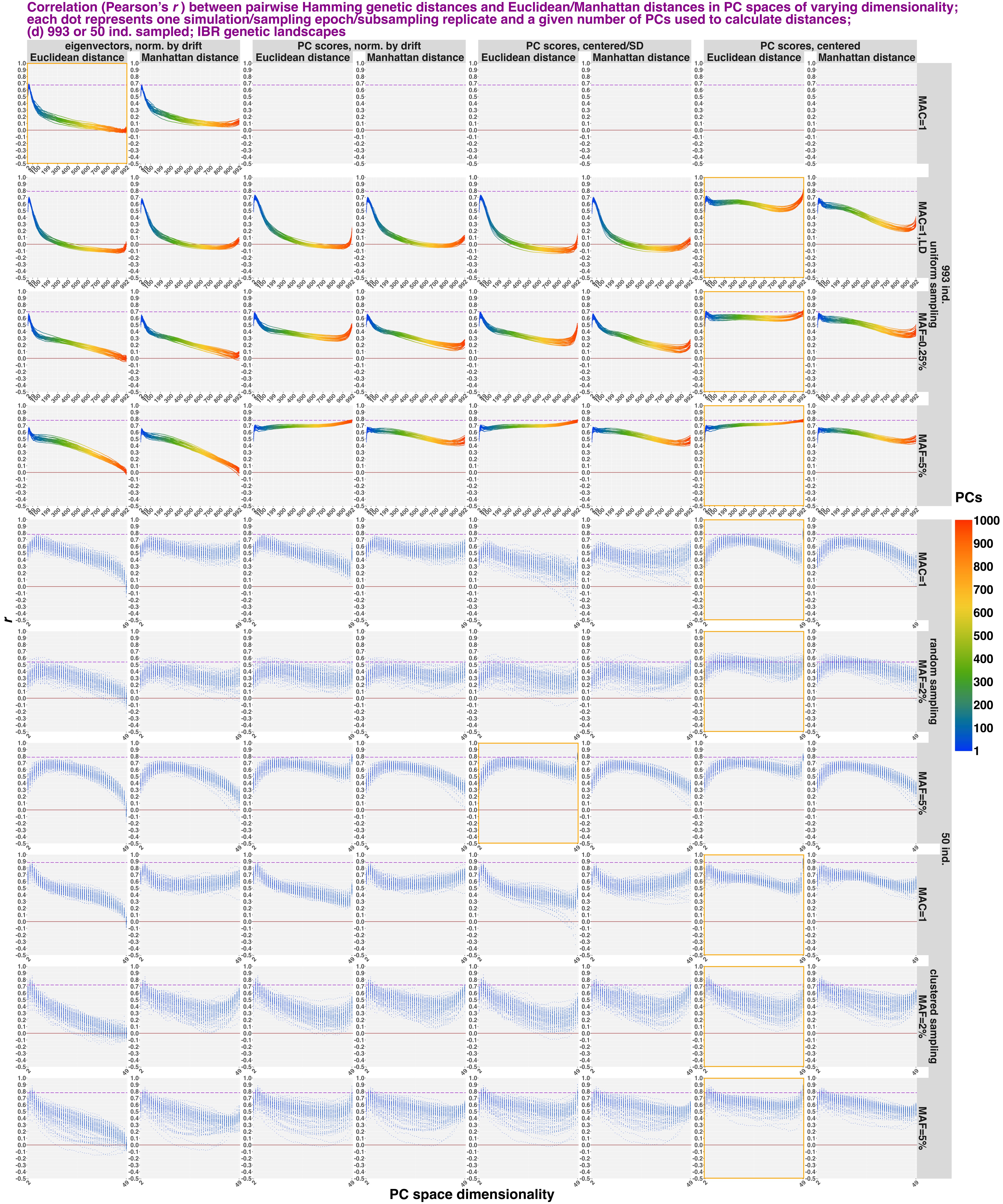

### Supplementary Figure 10

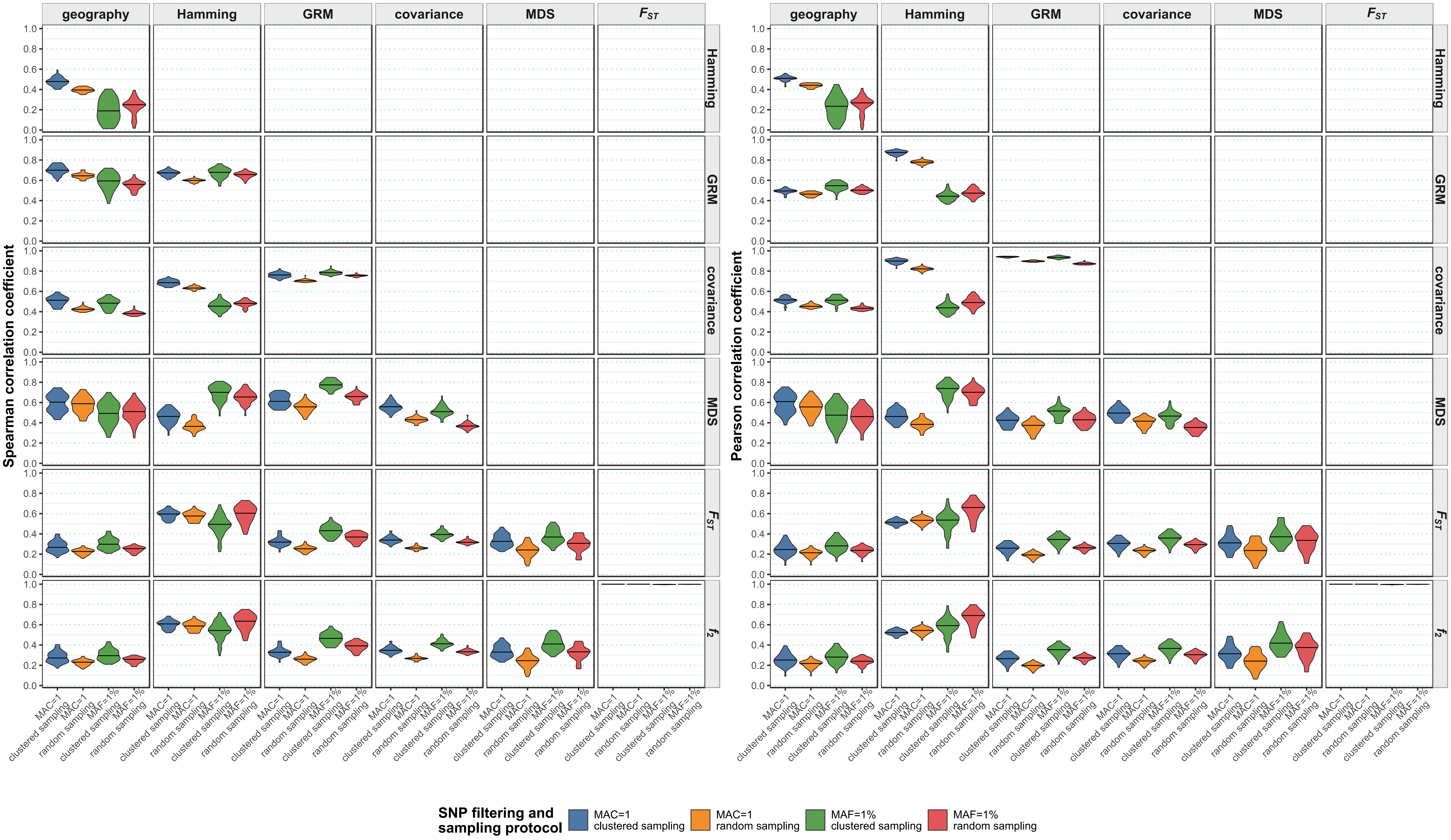

### Supplementary Figure 11a

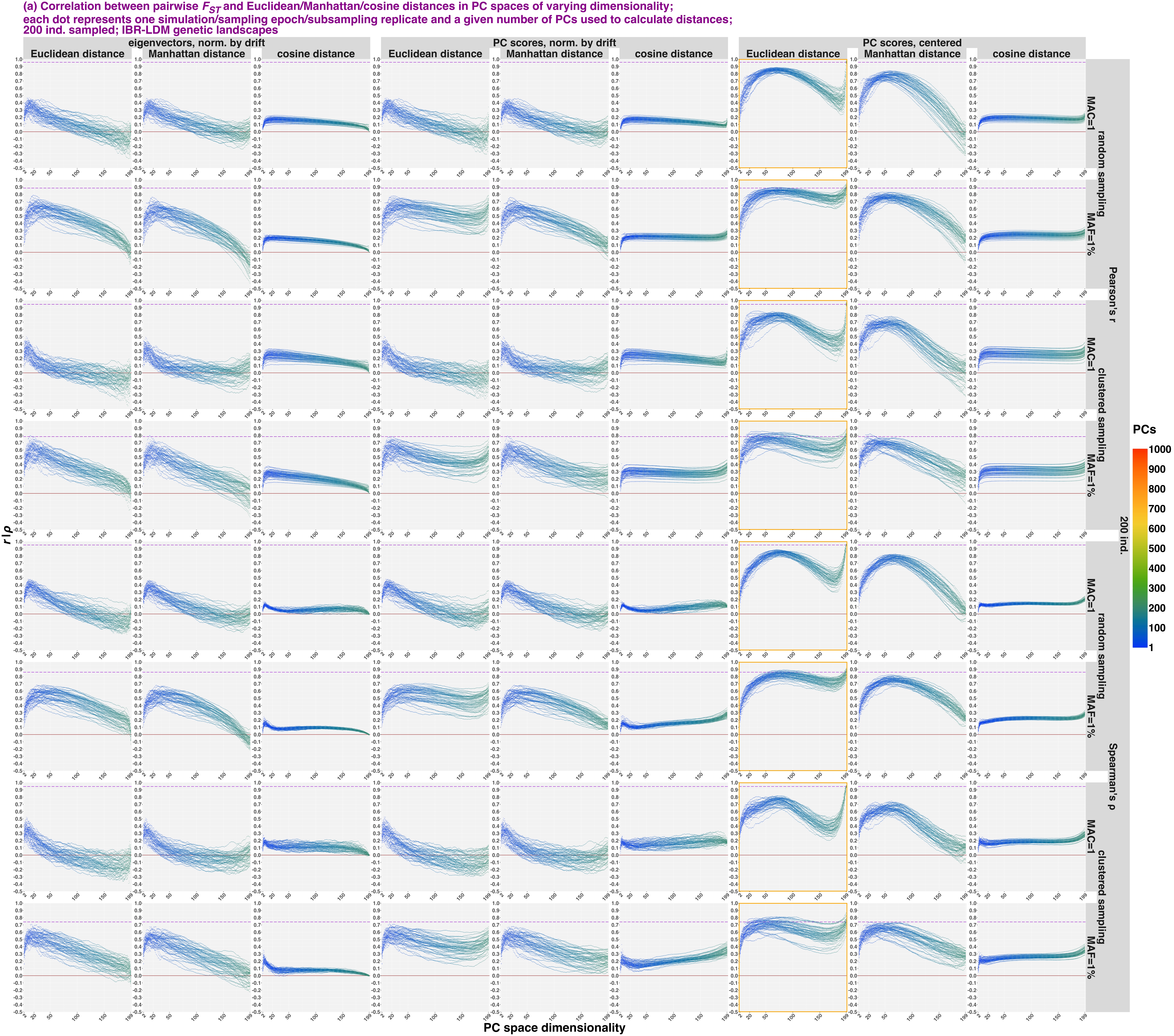

### Supplementary Figure 11b

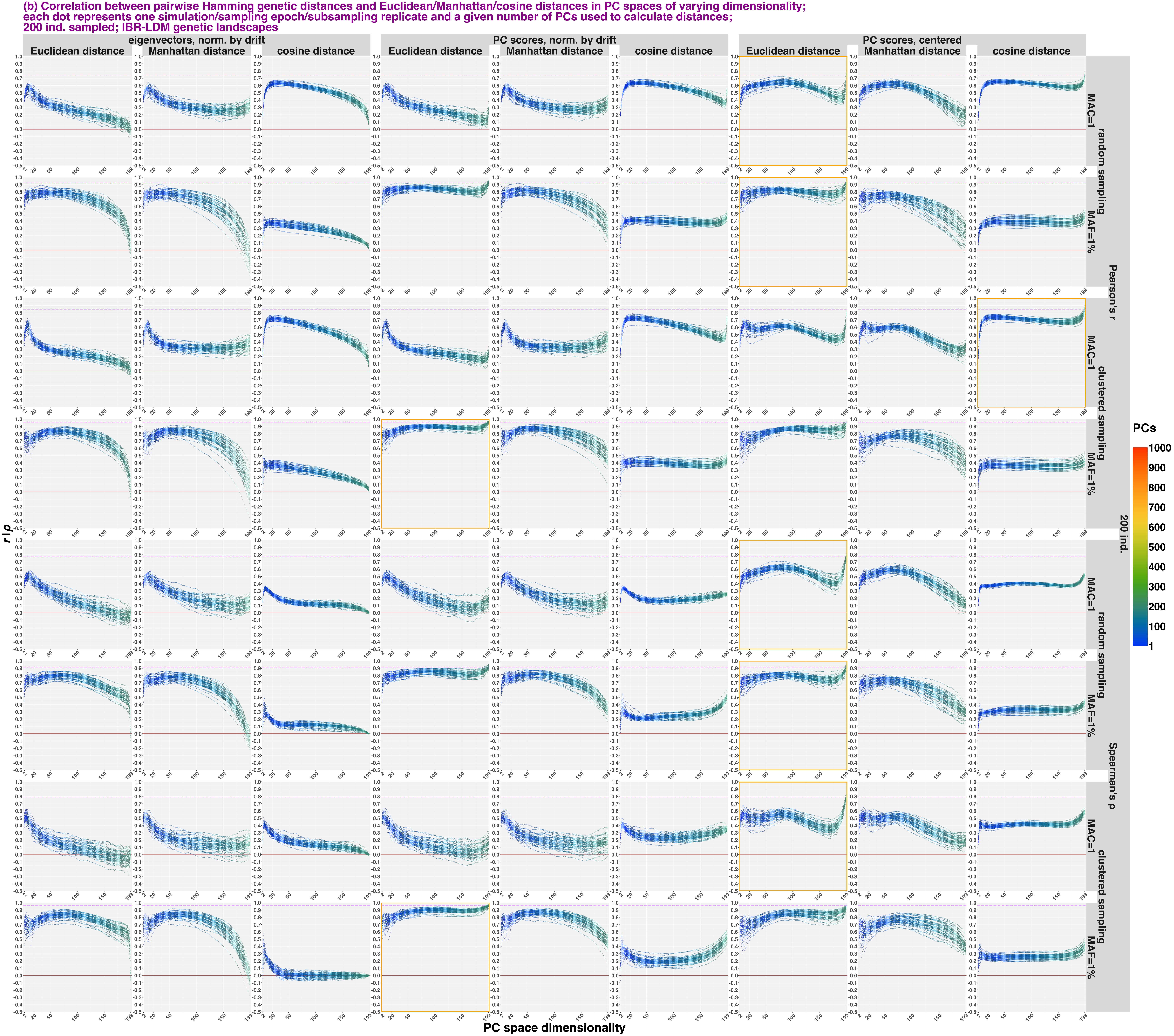

### Supplementary Figure 11c

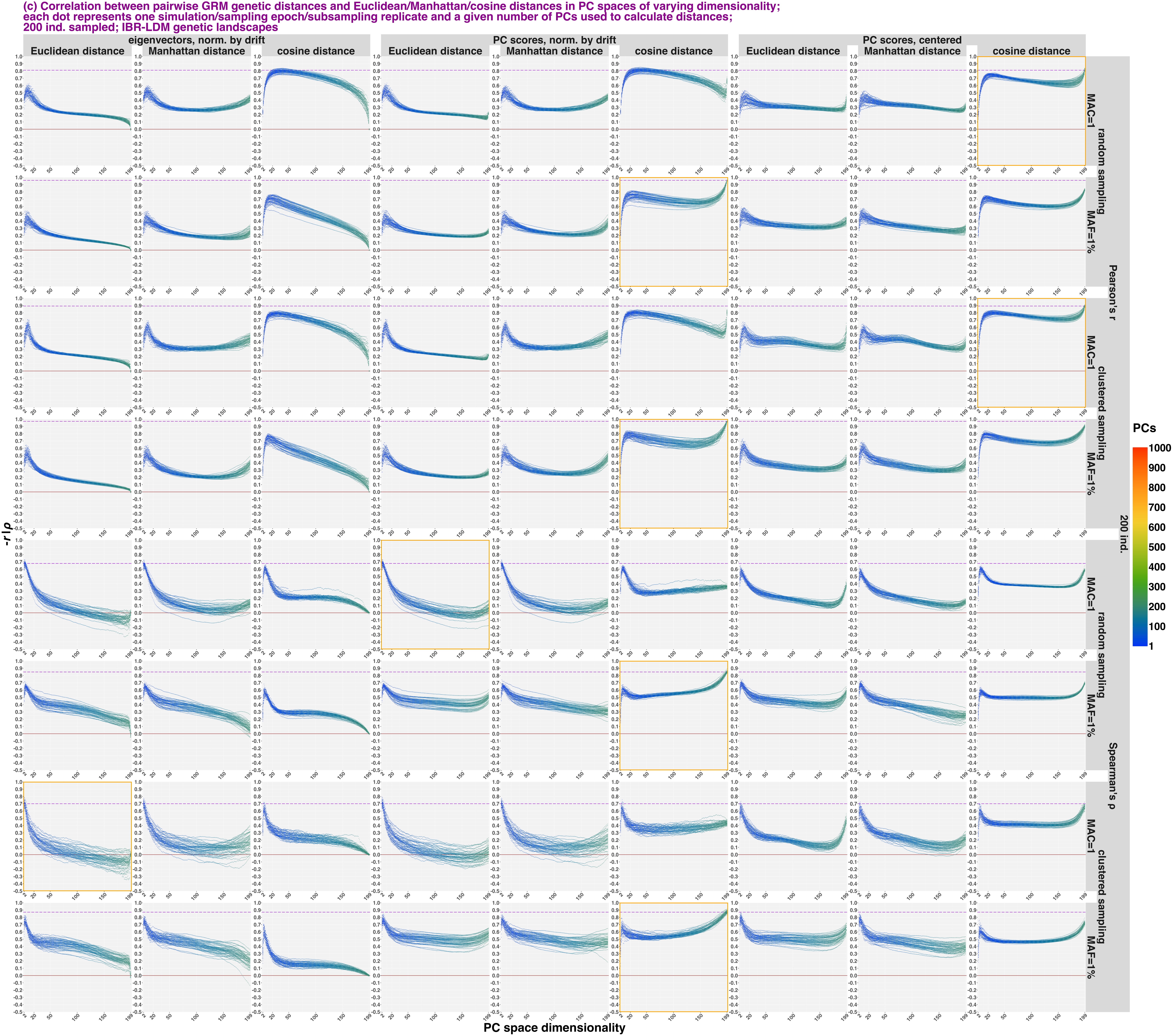

### Supplementary Figure 11d

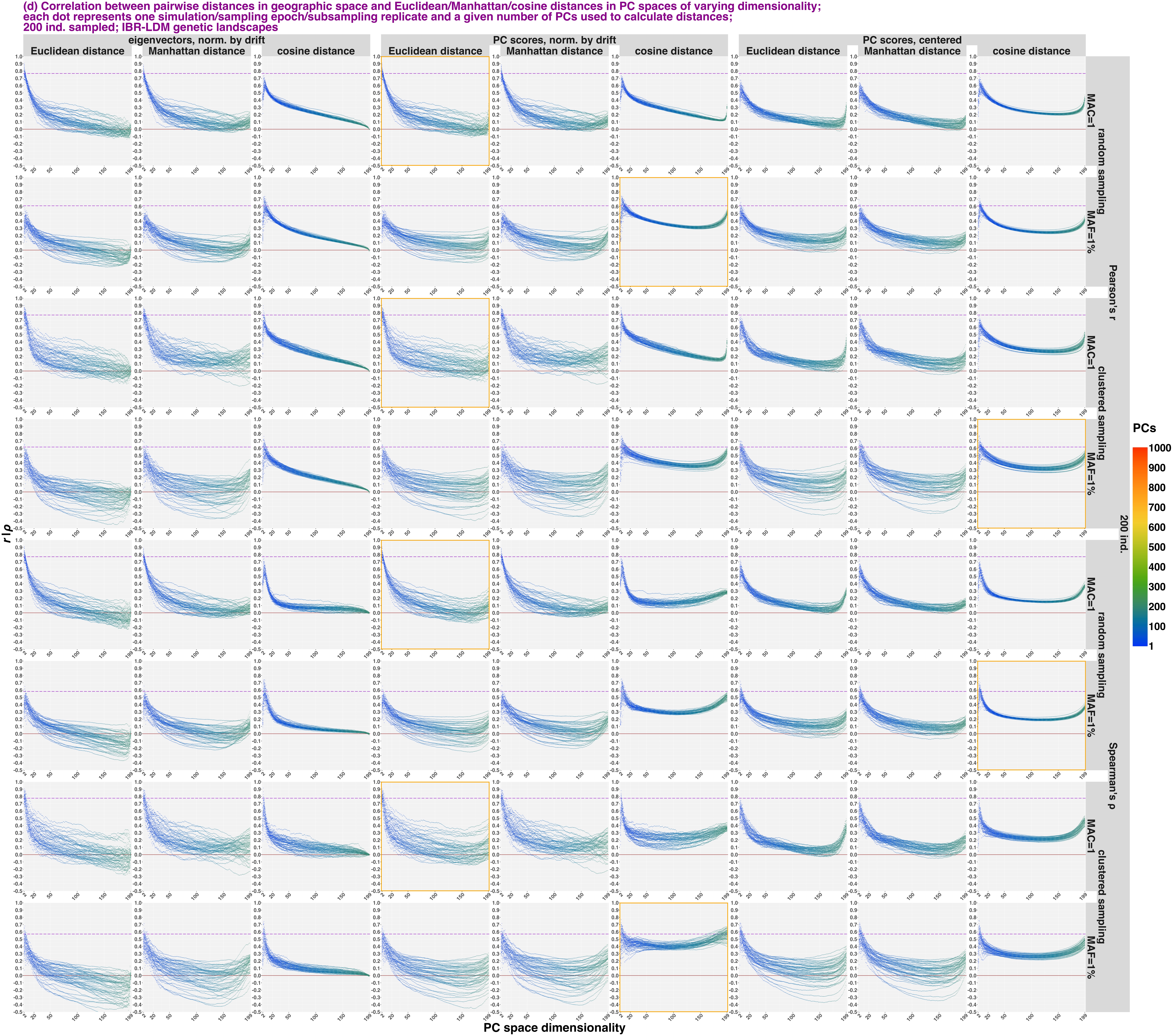

### Supplementary Figure 14

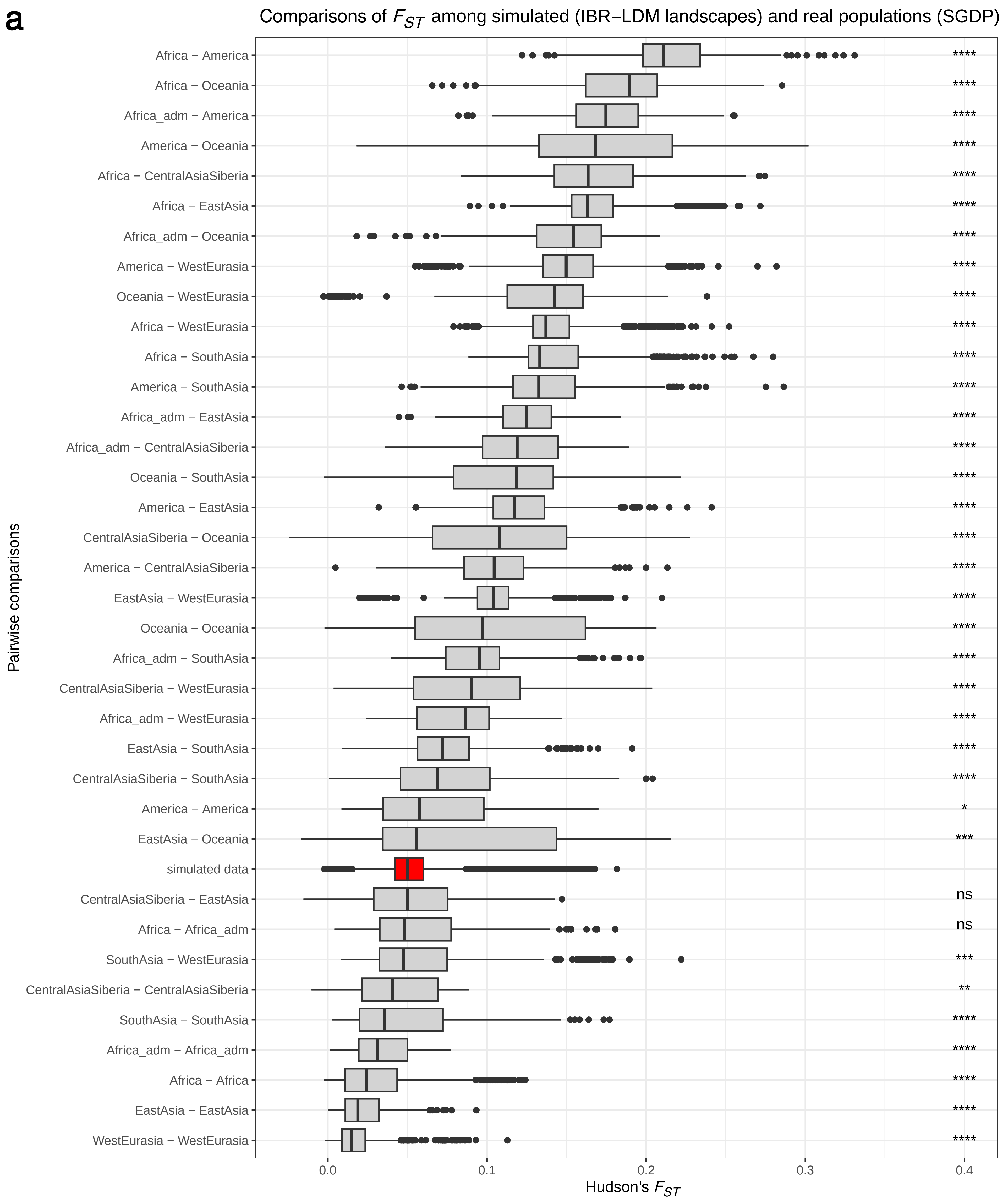

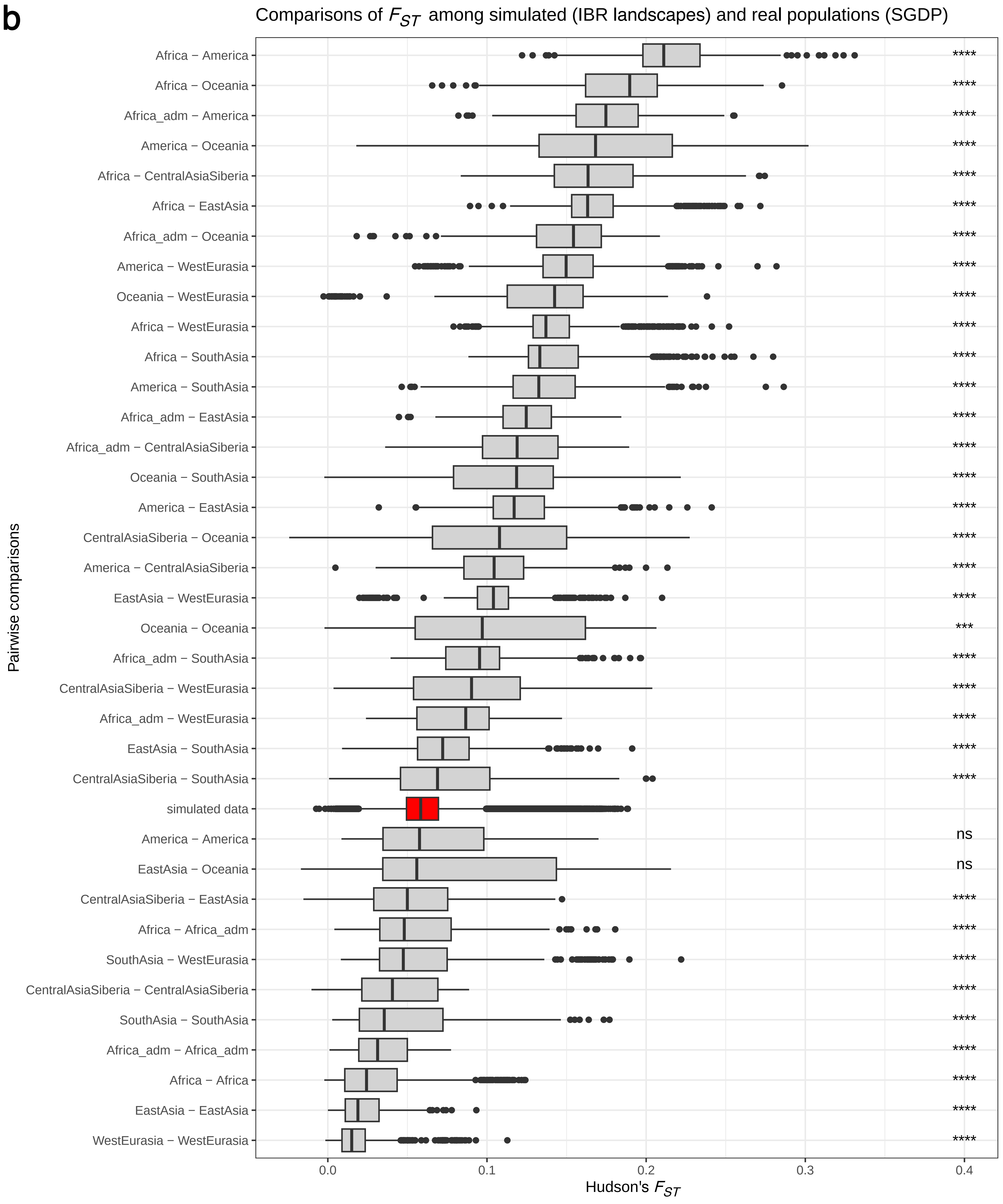

### Supplementary Figure 20a

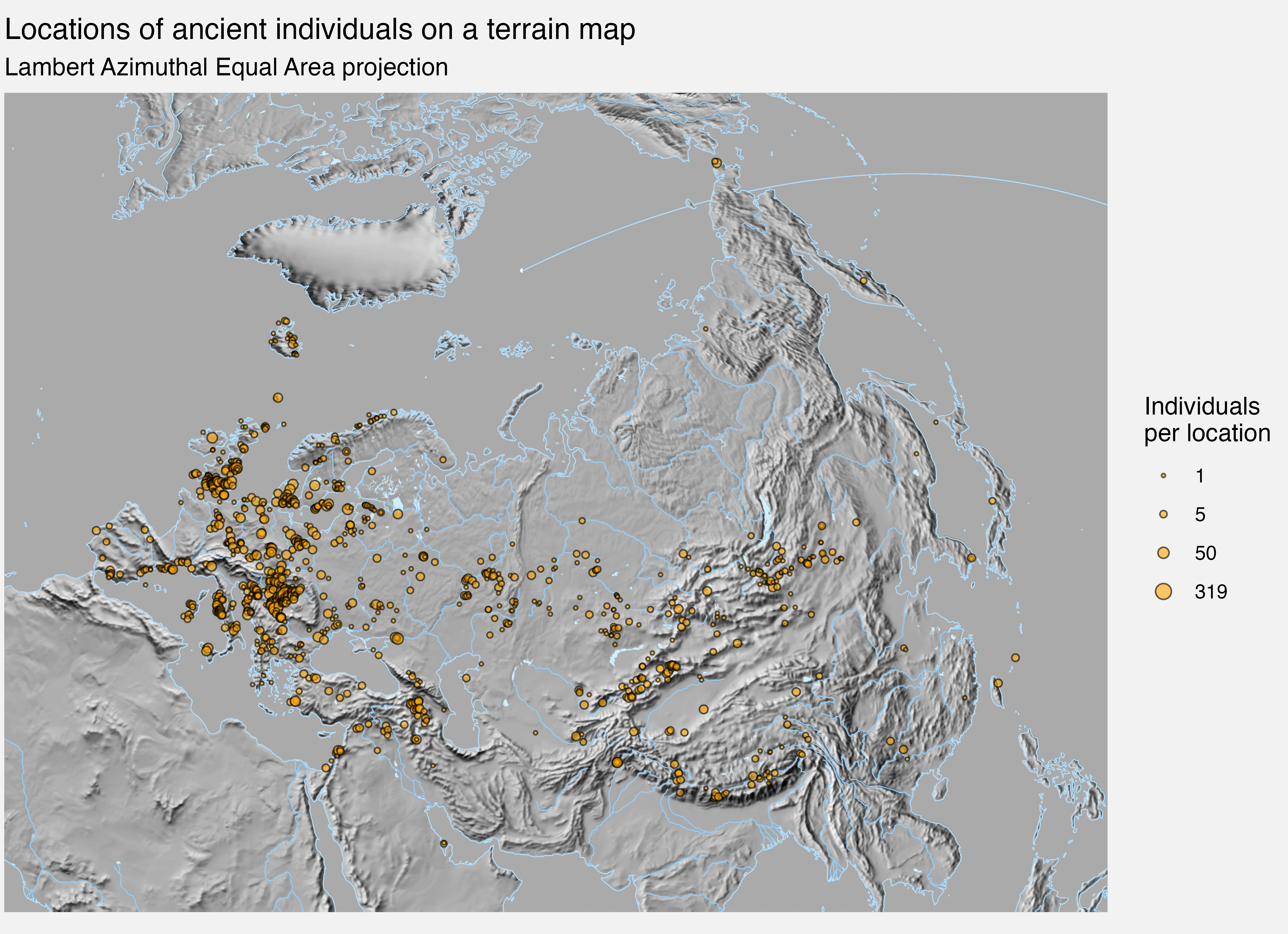

### Supplementary Figure 20b

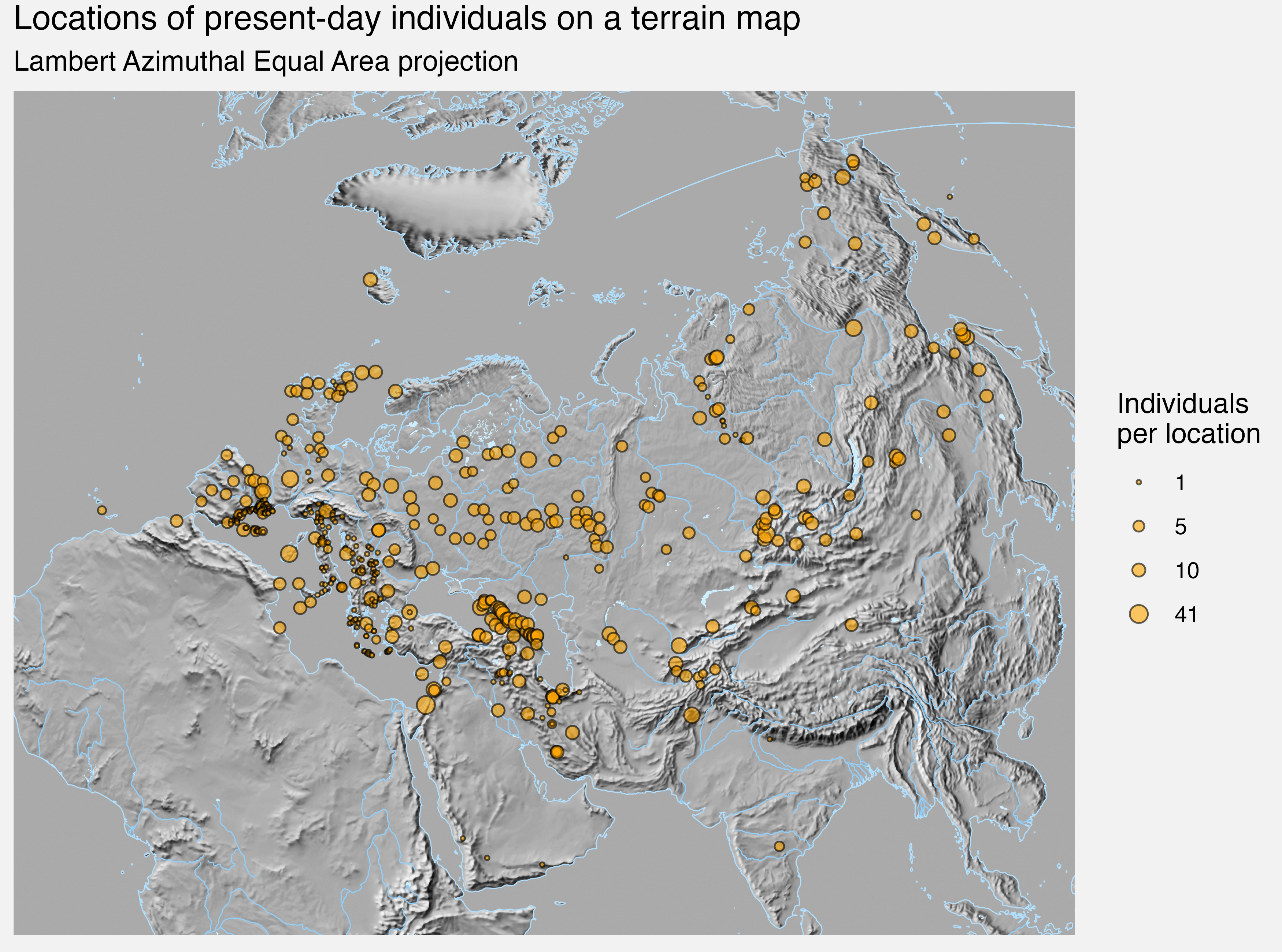

### Supplementary Figure 20c

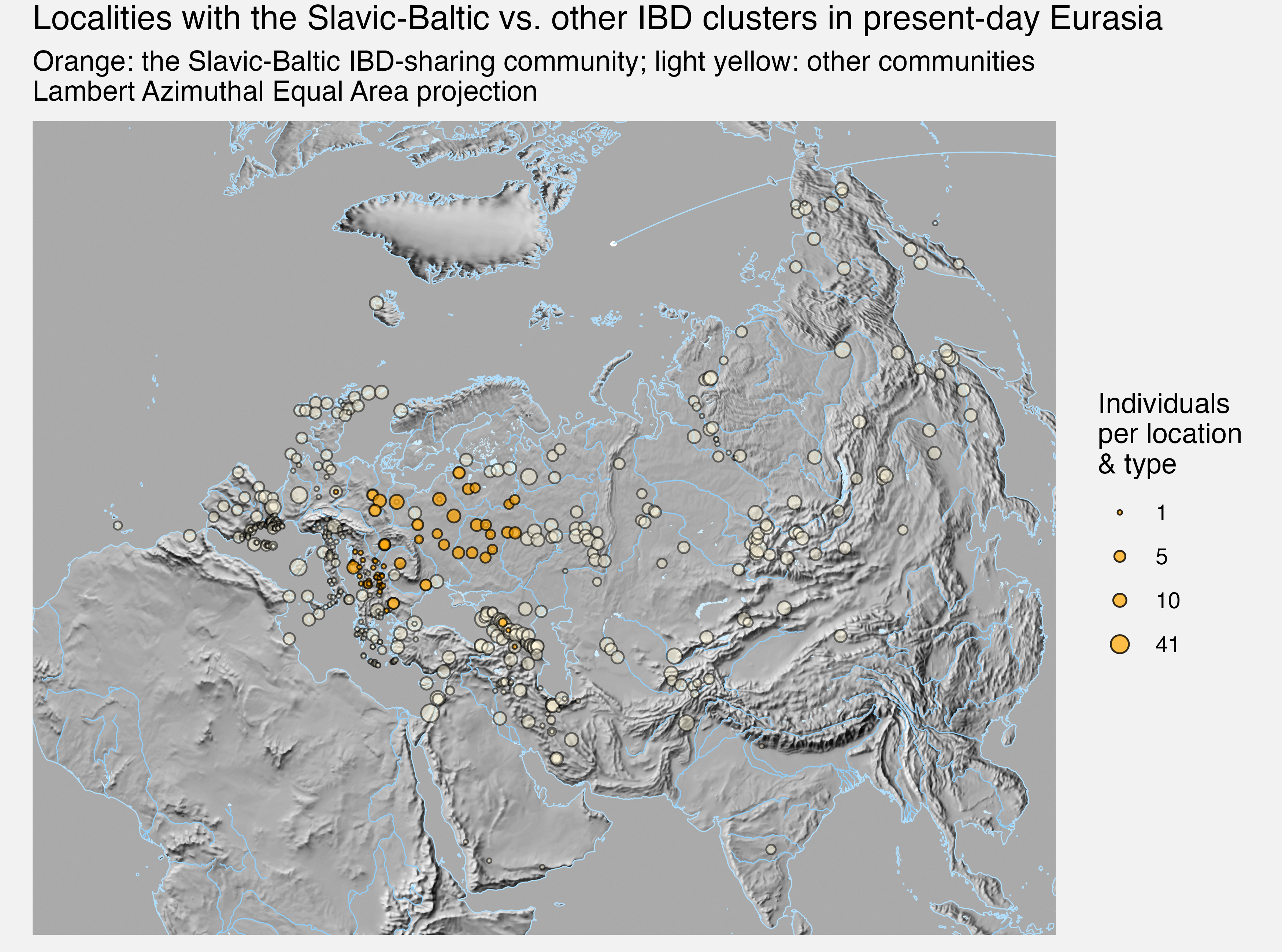
