## Supplementary Figure 2 for "Sparse sampling and rare-variant depletion distort PCA visualizations of population structure: recovery with objective-guided manifold learning"

**a**

**effects of sparse sampling on visualization of IBD landscapes in PC1−PC2 space (eigenvectors, norm. by drift); random sampling, simulation repl. 2, 300 gen. BP, MAC=1**

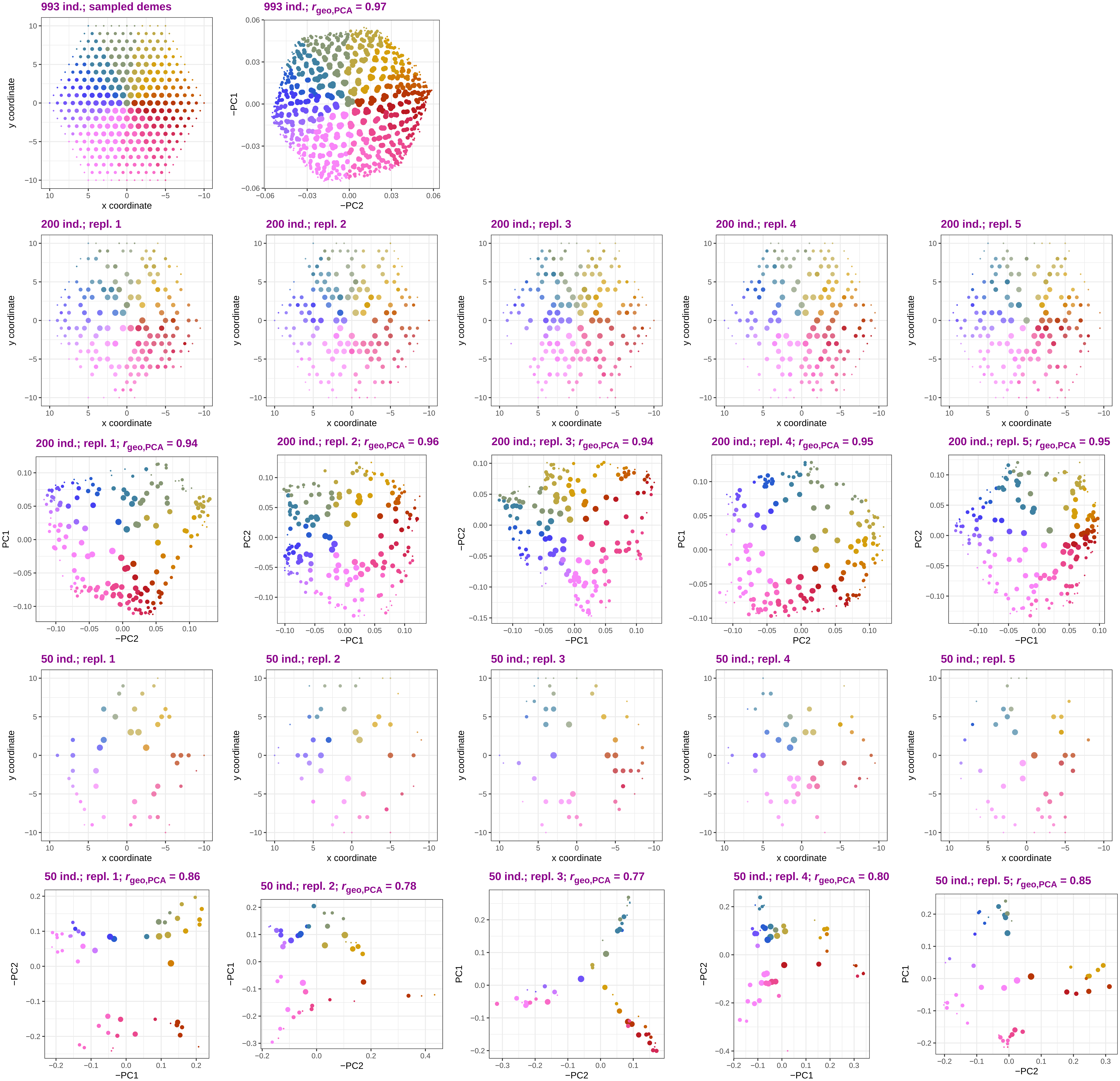

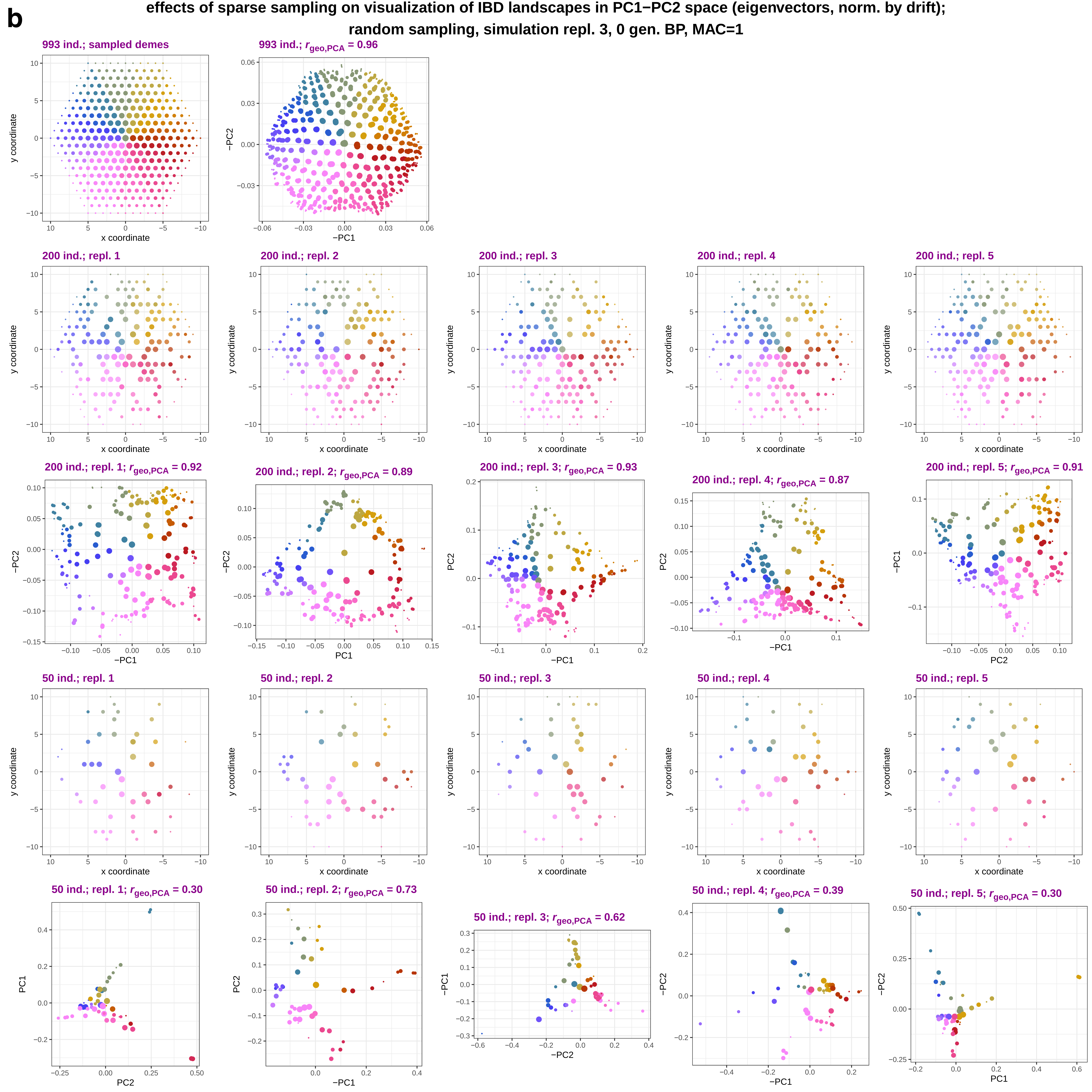

b
effects of sparse sampling on visualization of IBD landscapes in PC1−PC2 space (eigenvectors, norm. by drift);
random sampling, simulation repl. 3, 0 gen. BP, MAC=1
993 ind.; sampled demes
993 ind.; $r_{geo,PCA}$ = 0.96
200 ind.; repl. 1
200 ind.; repl. 2
200 ind.; repl. 3
200 ind.; repl. 4
200 ind.; repl. 5
200 ind.; repl. 1; $r_{geo,PCA}$ = 0.92
200 ind.; repl. 2; $r_{geo,PCA}$ = 0.89
200 ind.; repl. 3; $r_{geo,PCA}$ = 0.93
200 ind.; repl. 4; $r_{geo,PCA}$ = 0.87
200 ind.; repl. 5; $r_{geo,PCA}$ = 0.91
50 ind.; repl. 1
50 ind.; repl. 2
50 ind.; repl. 3
50 ind.; repl. 4
50 ind.; repl. 5
50 ind.; repl. 1; $r_{geo,PCA}$ = 0.30
50 ind.; repl. 2; $r_{geo,PCA}$ = 0.73
50 ind.; repl. 3; $r_{geo,PCA}$ = 0.62
50 ind.; repl. 4; $r_{geo,PCA}$ = 0.39
50 ind.; repl. 5; $r_{geo,PCA}$ = 0.30
x coordinate
y coordinate
PC1
−PC1
PC2
−PC2

**c**

### effects of sparse sampling on visualization of IBD landscapes in PC1−PC2 space (eigenvectors, norm. by drift); random sampling, simulation repl. 8, 300 gen. BP, MAC=1

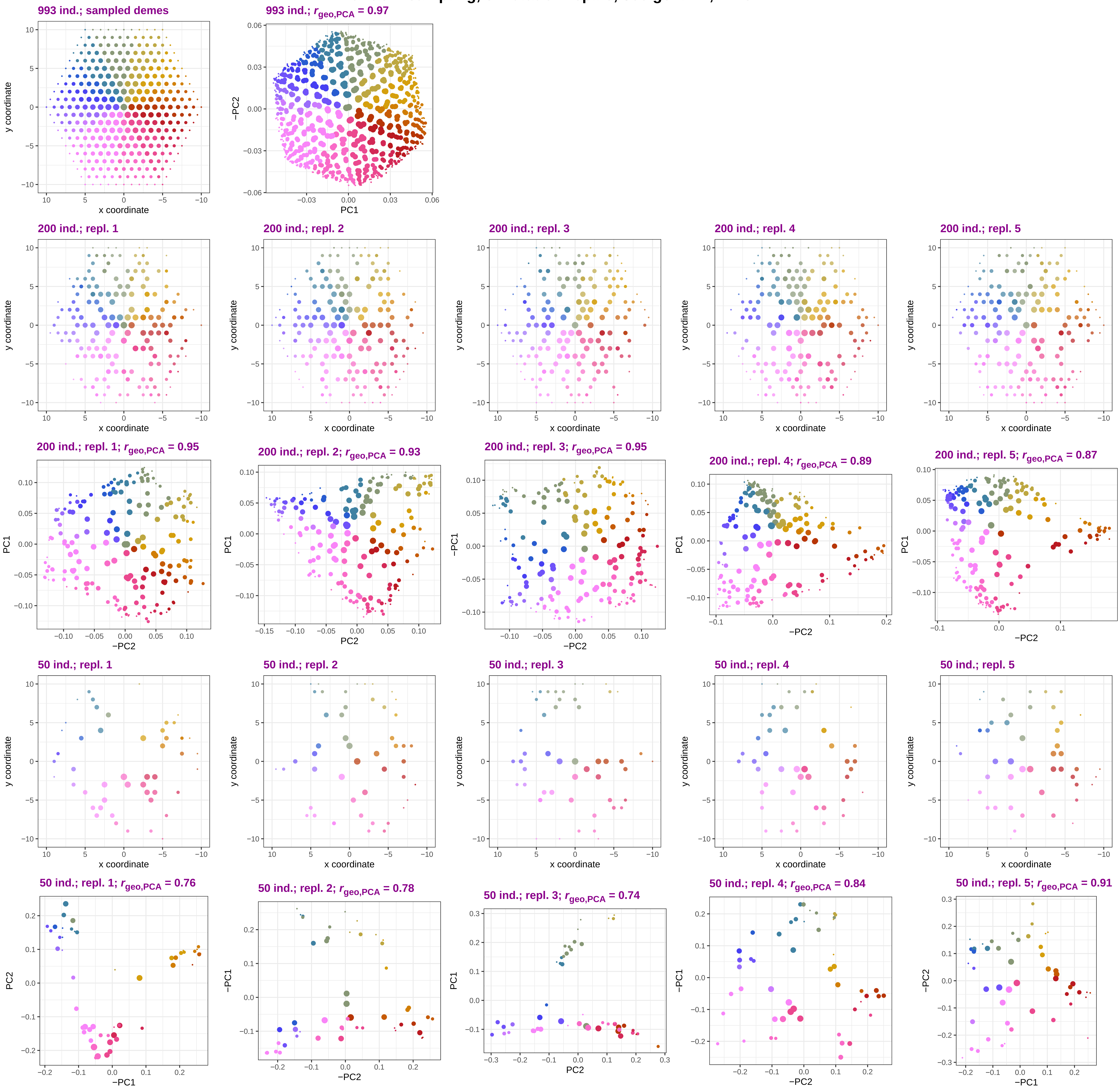

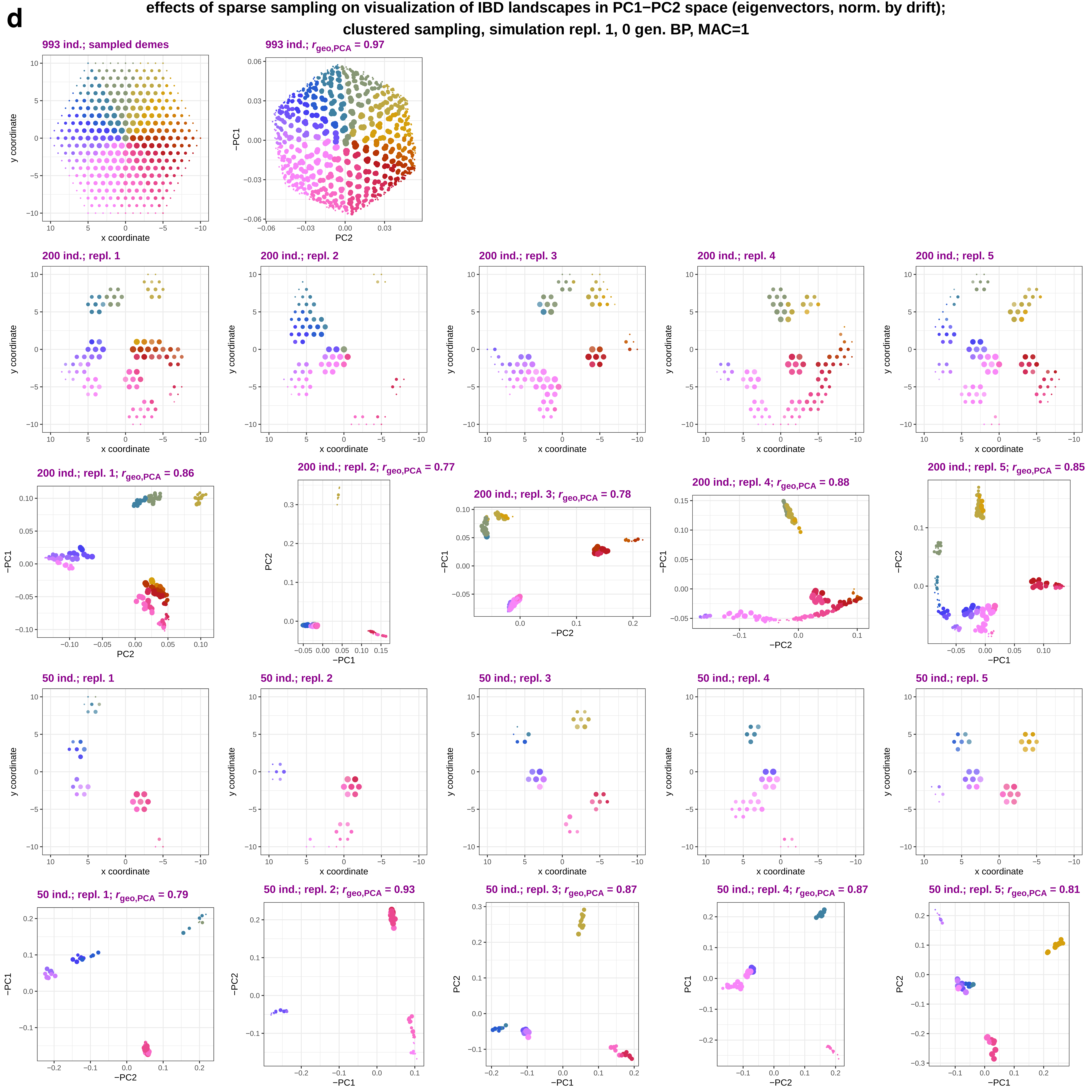
d
effects of sparse sampling on visualization of IBD landscapes in PC1−PC2 space (eigenvectors, norm. by drift);
clustered sampling, simulation repl. 1, 0 gen. BP, MAC=1
993 ind.; sampled demes
993 ind.; $r_{geo,PCA}$ = 0.97
200 ind.; repl. 1
200 ind.; repl. 2
200 ind.; repl. 3
200 ind.; repl. 4
200 ind.; repl. 5
200 ind.; repl. 1; $r_{geo,PCA}$ = 0.86
200 ind.; repl. 2; $r_{geo,PCA}$ = 0.77
200 ind.; repl. 3; $r_{geo,PCA}$ = 0.78
200 ind.; repl. 4; $r_{geo,PCA}$ = 0.88
200 ind.; repl. 5; $r_{geo,PCA}$ = 0.85
50 ind.; repl. 1
50 ind.; repl. 2
50 ind.; repl. 3
50 ind.; repl. 4
50 ind.; repl. 5
50 ind.; repl. 1; $r_{geo,PCA}$ = 0.79
50 ind.; repl. 2; $r_{geo,PCA}$ = 0.93
50 ind.; repl. 3; $r_{geo,PCA}$ = 0.87
50 ind.; repl. 4; $r_{geo,PCA}$ = 0.87
50 ind.; repl. 5; $r_{geo,PCA}$ = 0.81
x coordinate
y coordinate
PC1
−PC1
PC2
−PC2

**e** **effects of sparse sampling on visualization of IBD landscapes in PC1−PC2 space (eigenvectors, norm. by drift); clustered sampling, simulation repl. 4, 0 gen. BP, MAC=1**

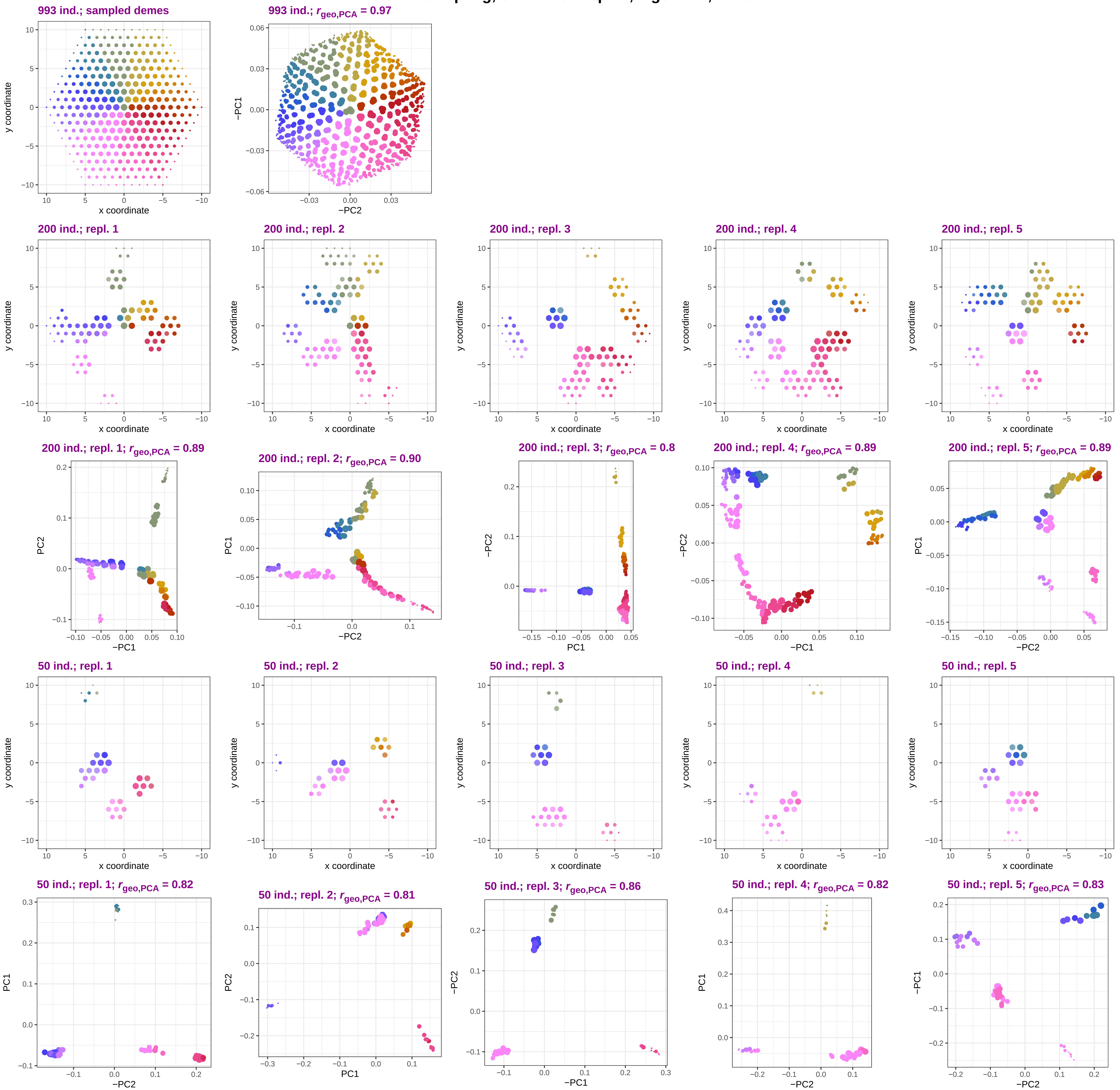

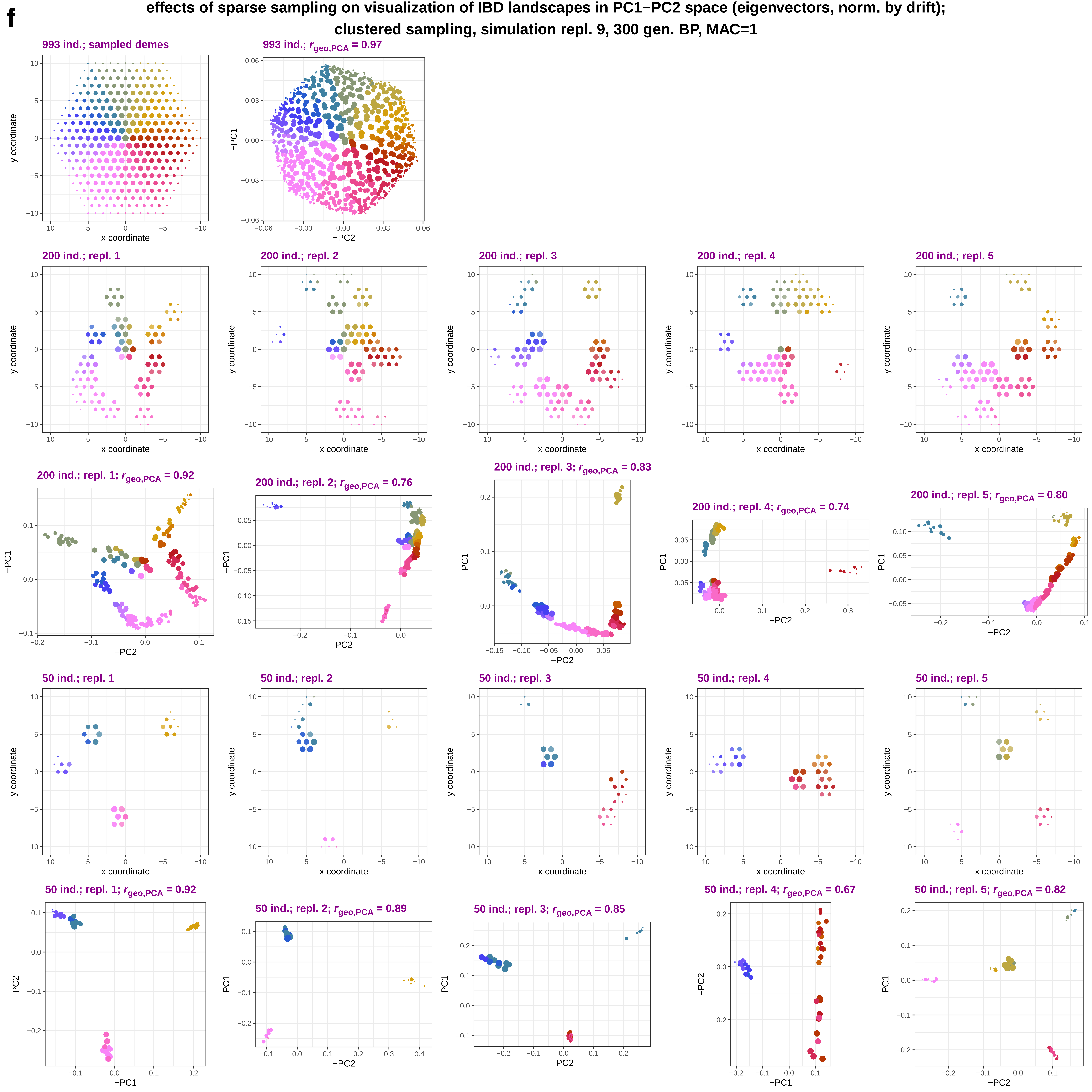
f
effects of sparse sampling on visualization of IBD landscapes in PC1−PC2 space (eigenvectors, norm. by drift);
clustered sampling, simulation repl. 9, 300 gen. BP, MAC=1
993 ind.; sampled demes
y coordinate
x coordinate
993 ind.; $r_{geo,PCA}$ = 0.97
−PC1
−PC2
200 ind.; repl. 1
200 ind.; repl. 2
200 ind.; repl. 3
200 ind.; repl. 4
200 ind.; repl. 5
200 ind.; repl. 1; $r_{geo,PCA}$ = 0.92
−PC1
−PC2
200 ind.; repl. 2; $r_{geo,PCA}$ = 0.76
−PC1
PC2
200 ind.; repl. 3; $r_{geo,PCA}$ = 0.83
PC1
−PC2
200 ind.; repl. 4; $r_{geo,PCA}$ = 0.74
PC1
−PC2
200 ind.; repl. 5; $r_{geo,PCA}$ = 0.80
PC1
−PC2
50 ind.; repl. 1
50 ind.; repl. 2
50 ind.; repl. 3
50 ind.; repl. 4
50 ind.; repl. 5
50 ind.; repl. 1; $r_{geo,PCA}$ = 0.92
PC2
−PC1
50 ind.; repl. 2; $r_{geo,PCA}$ = 0.89
PC1
−PC2
50 ind.; repl. 3; $r_{geo,PCA}$ = 0.85
PC1
−PC2
50 ind.; repl. 4; $r_{geo,PCA}$ = 0.67
−PC2
−PC1
50 ind.; repl. 5; $r_{geo,PCA}$ = 0.82
PC1
−PC2
