## Supplementary Figure 3 for "Sparse sampling and rare-variant depletion distort PCA visualizations of population structure: recovery with objective-guided manifold learning"

a

**effects of SNP filtering and pruning on visualization of IBD landscapes in PC1−PC2 space (eigenvectors, norm. by drift); random sampling (200 ind.), simulation repl. 1, 0 gen. BP, subsampling repl. 1**

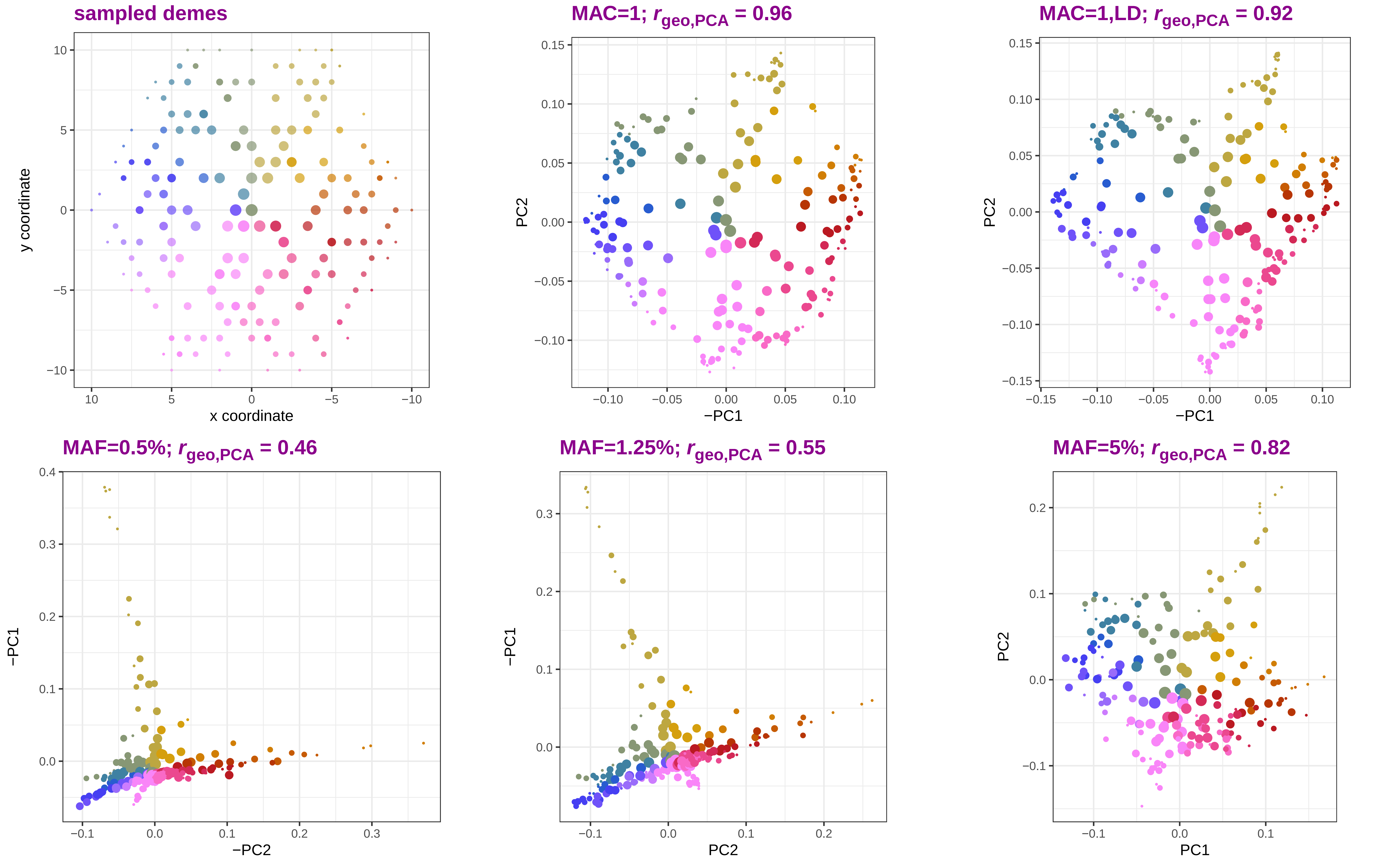

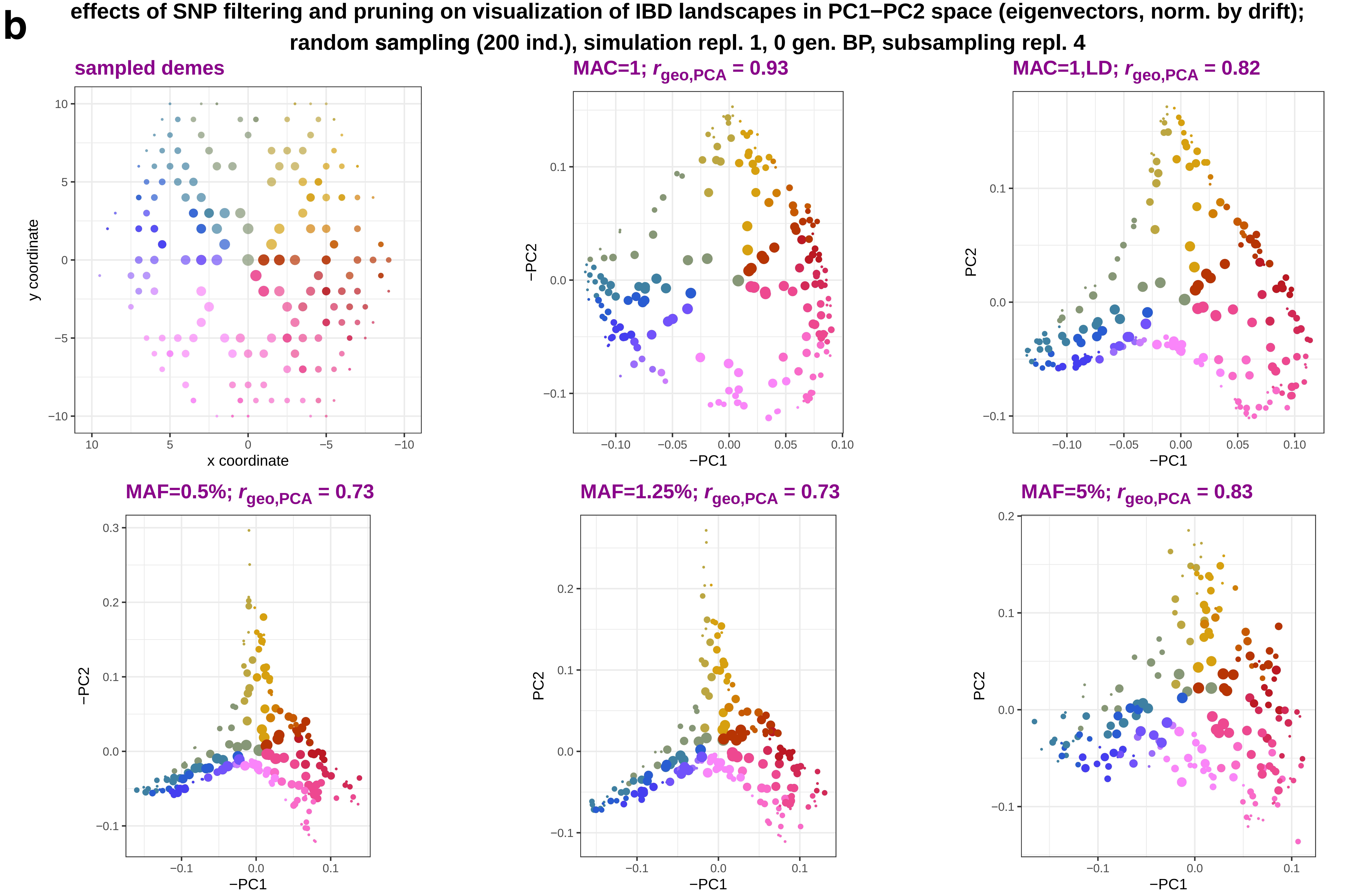
b
effects of SNP filtering and pruning on visualization of IBD landscapes in PC1−PC2 space (eigenvectors, norm. by drift);
random sampling (200 ind.), simulation repl. 1, 0 gen. BP, subsampling repl. 4
sampled demes
x coordinate
y coordinate
MAC=1; r geo,PCA = 0.93
MAC=1,LD; r geo,PCA = 0.82
MAF=0.5%; r geo,PCA = 0.73
MAF=1.25%; r geo,PCA = 0.73
MAF=5%; r geo,PCA = 0.83
−PC1
−PC2
PC2

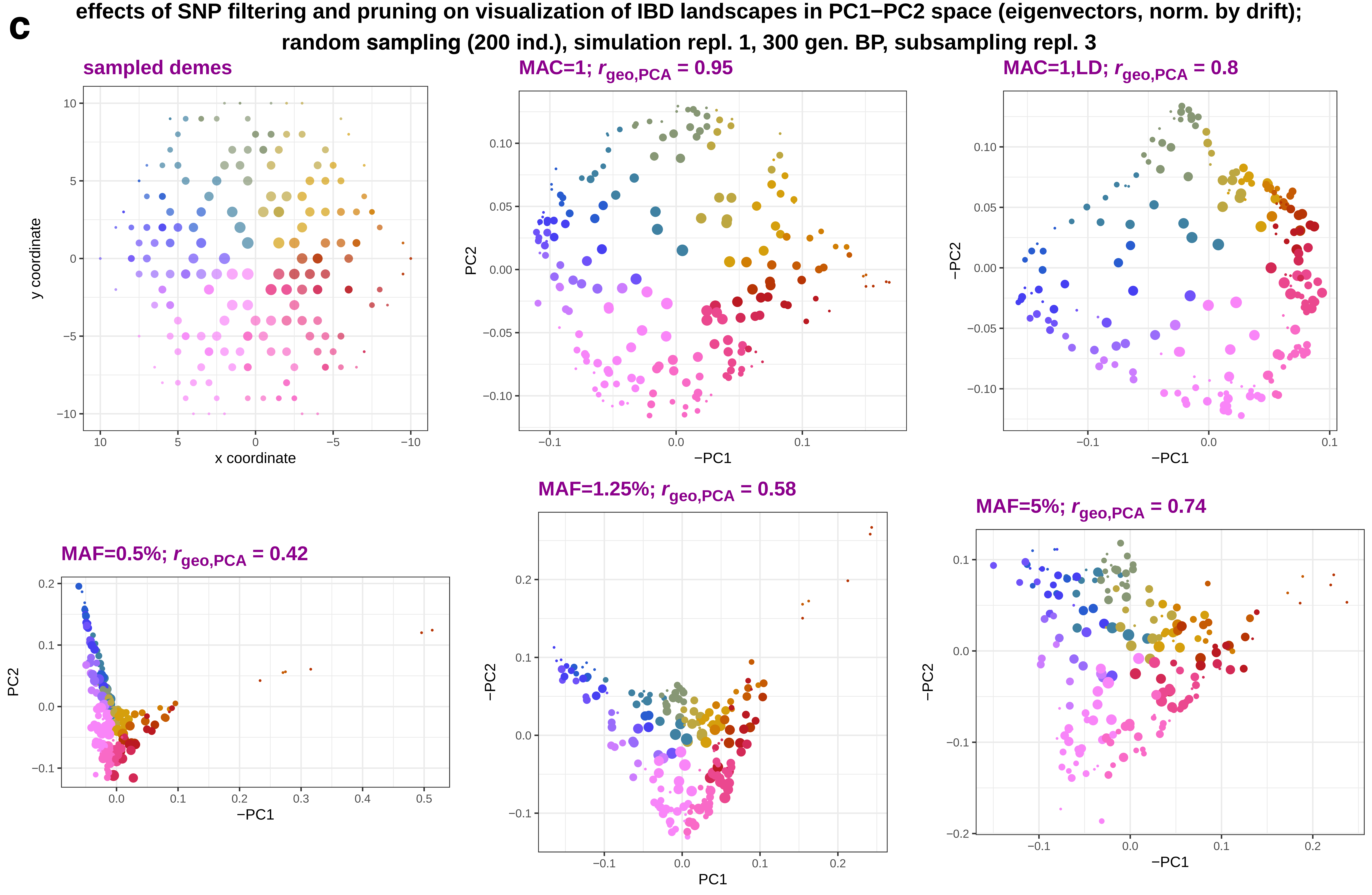
c
effects of SNP filtering and pruning on visualization of IBD landscapes in PC1−PC2 space (eigenvectors, norm. by drift);
random sampling (200 ind.), simulation repl. 1, 300 gen. BP, subsampling repl. 3
sampled demes
y coordinate
x coordinate
MAC=1; $r_{geo,PCA}$ = 0.95
PC2
−PC1
MAC=1,LD; $r_{geo,PCA}$ = 0.8
−PC2
−PC1
MAF=0.5%; $r_{geo,PCA}$ = 0.42
PC2
−PC1
MAF=1.25%; $r_{geo,PCA}$ = 0.58
−PC2
PC1
MAF=5%; $r_{geo,PCA}$ = 0.74
−PC2
−PC1

**d** **effects of SNP filtering and pruning on visualization of IBD landscapes in PC1−PC2 space (eigenvectors, norm. by drift); random sampling (200 ind.), simulation repl. 2, 0 gen. BP, subsampling repl. 4**

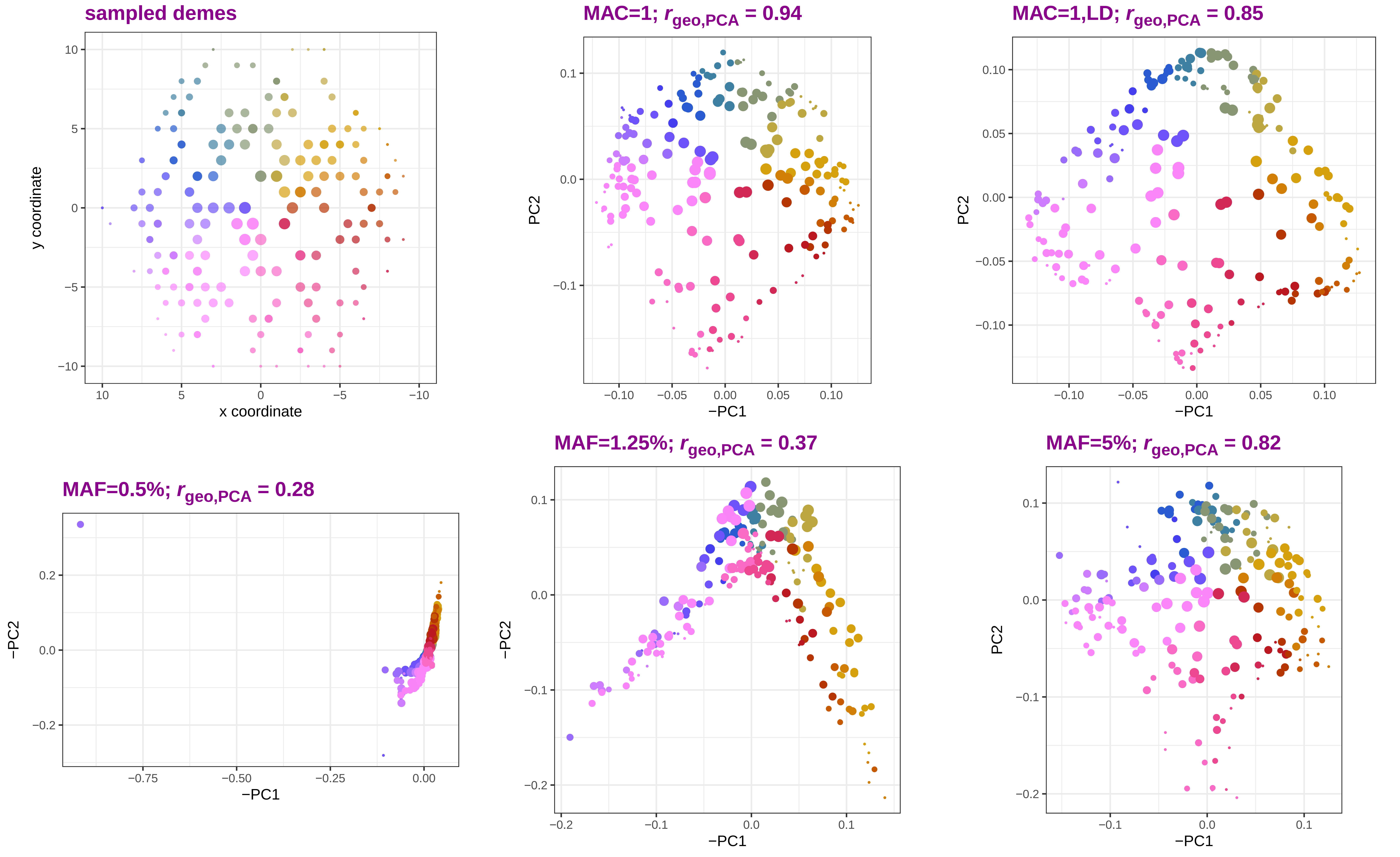

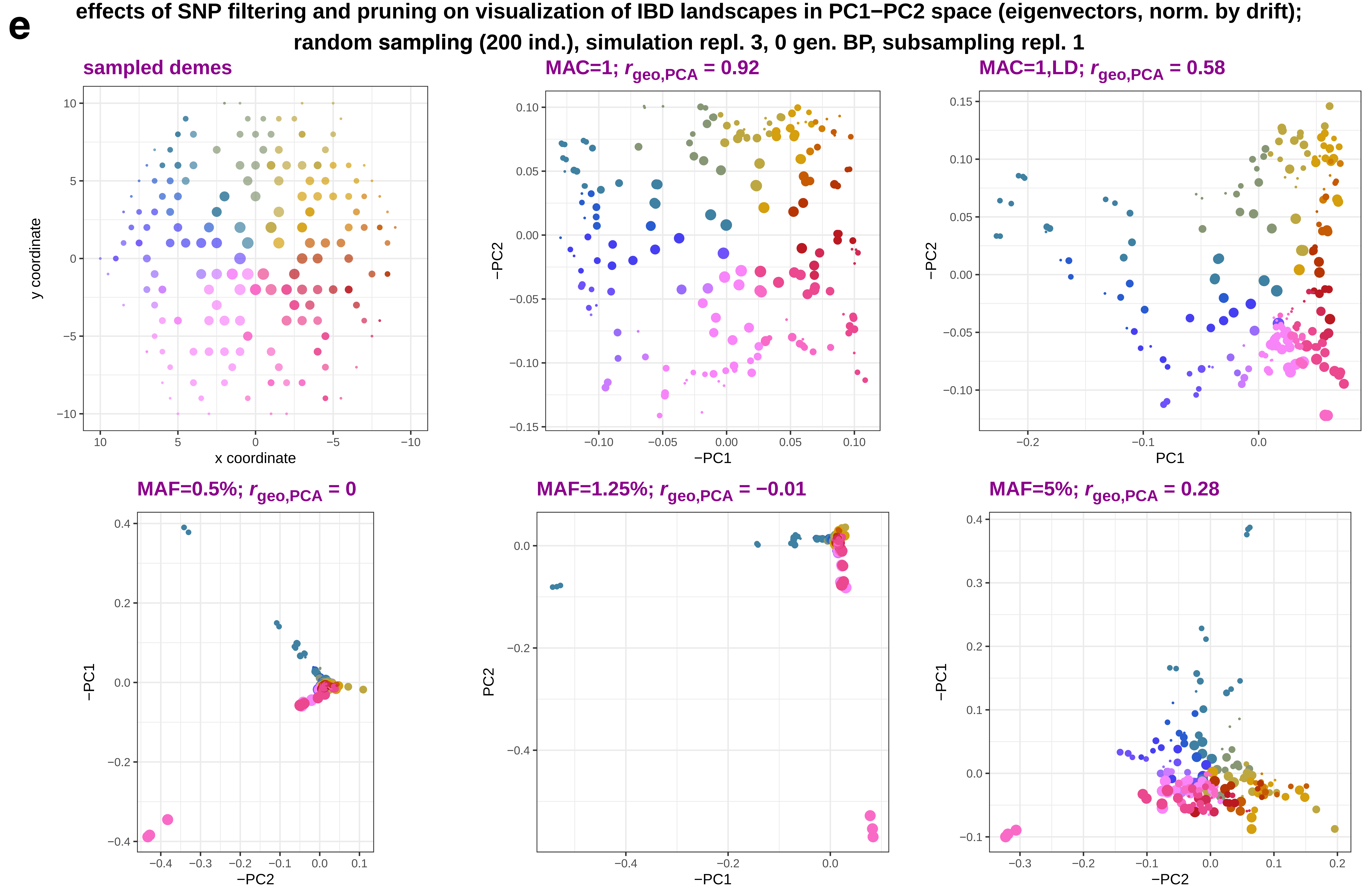
e
effects of SNP filtering and pruning on visualization of IBD landscapes in PC1−PC2 space (eigenvectors, norm. by drift);
random sampling (200 ind.), simulation repl. 3, 0 gen. BP, subsampling repl. 1
sampled demes
MAC=1; $r_{geo,PCA}$ = 0.92
MAC=1,LD; $r_{geo,PCA}$ = 0.58
MAF=0.5%; $r_{geo,PCA}$ = 0
MAF=1.25%; $r_{geo,PCA}$ = −0.01
MAF=5%; $r_{geo,PCA}$ = 0.28
y coordinate
x coordinate
−PC2
−PC1
PC1
PC2
−PC1
−PC2
PC2
−PC1
−PC1
−PC2

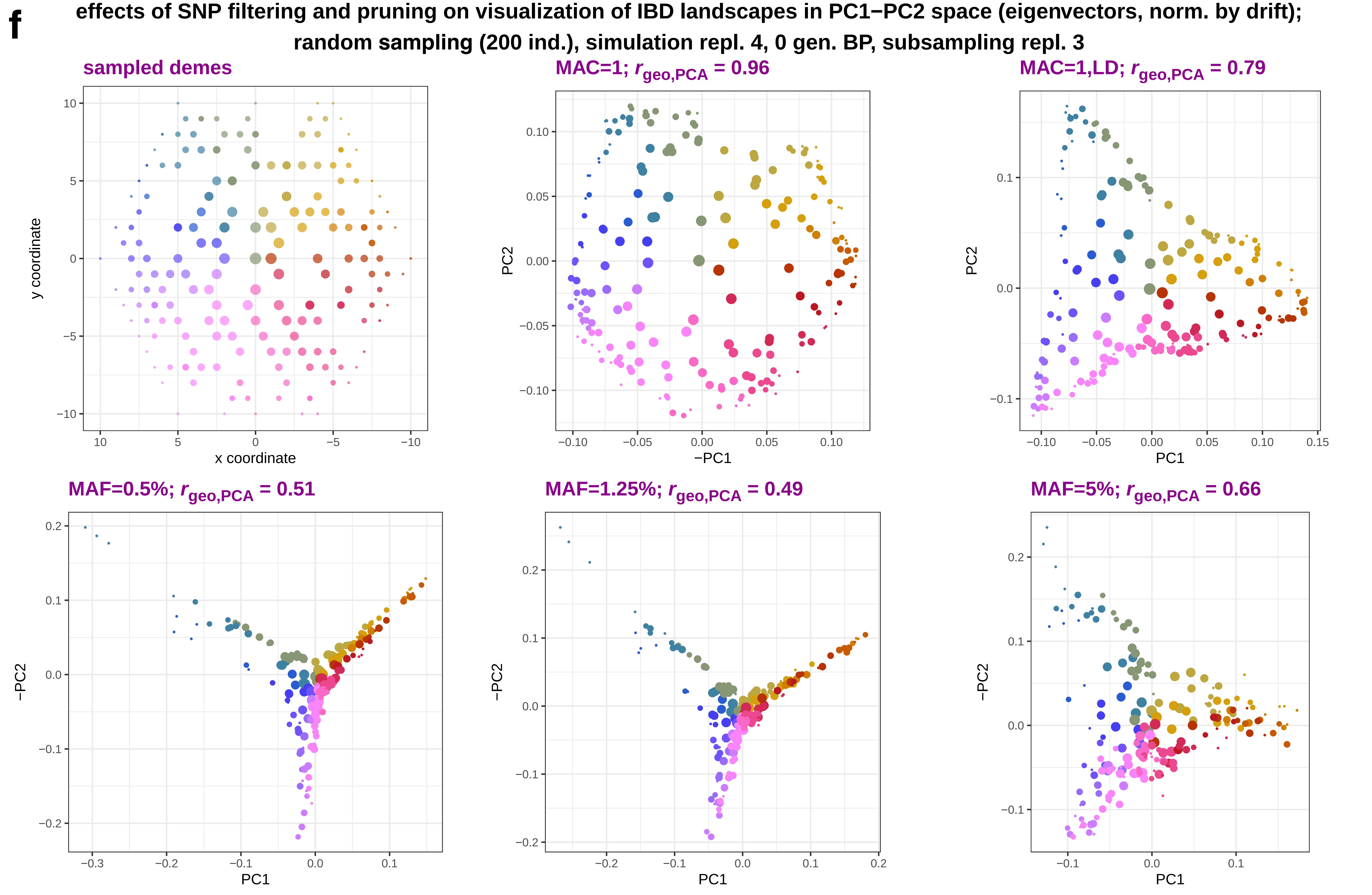
f
effects of SNP filtering and pruning on visualization of IBD landscapes in PC1−PC2 space (eigenvectors, norm. by drift);
random sampling (200 ind.), simulation repl. 4, 0 gen. BP, subsampling repl. 3
sampled demes
MAC=1; $r_{geo,PCA}$ = 0.96
MAC=1,LD; $r_{geo,PCA}$ = 0.79
MAF=0.5%; $r_{geo,PCA}$ = 0.51
MAF=1.25%; $r_{geo,PCA}$ = 0.49
MAF=5%; $r_{geo,PCA}$ = 0.66
x coordinate
y coordinate
−PC1
PC1
PC2
−PC2

**g** **effects of SNP filtering and pruning on visualization of IBD landscapes in PC1−PC2 space (eigenvectors, norm. by drift); random sampling (200 ind.), simulation repl. 10, 0 gen. BP, subsampling repl. 3**

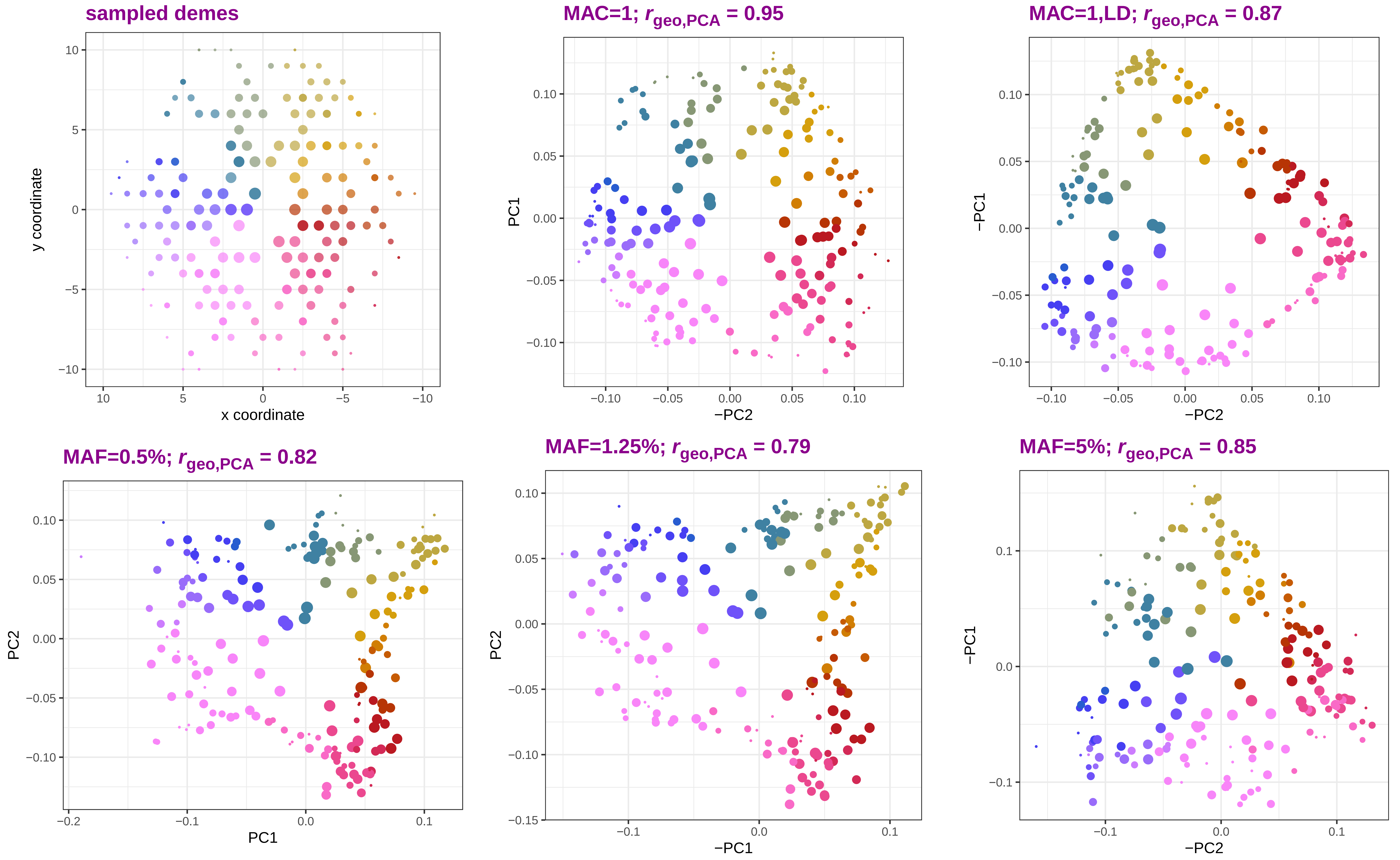

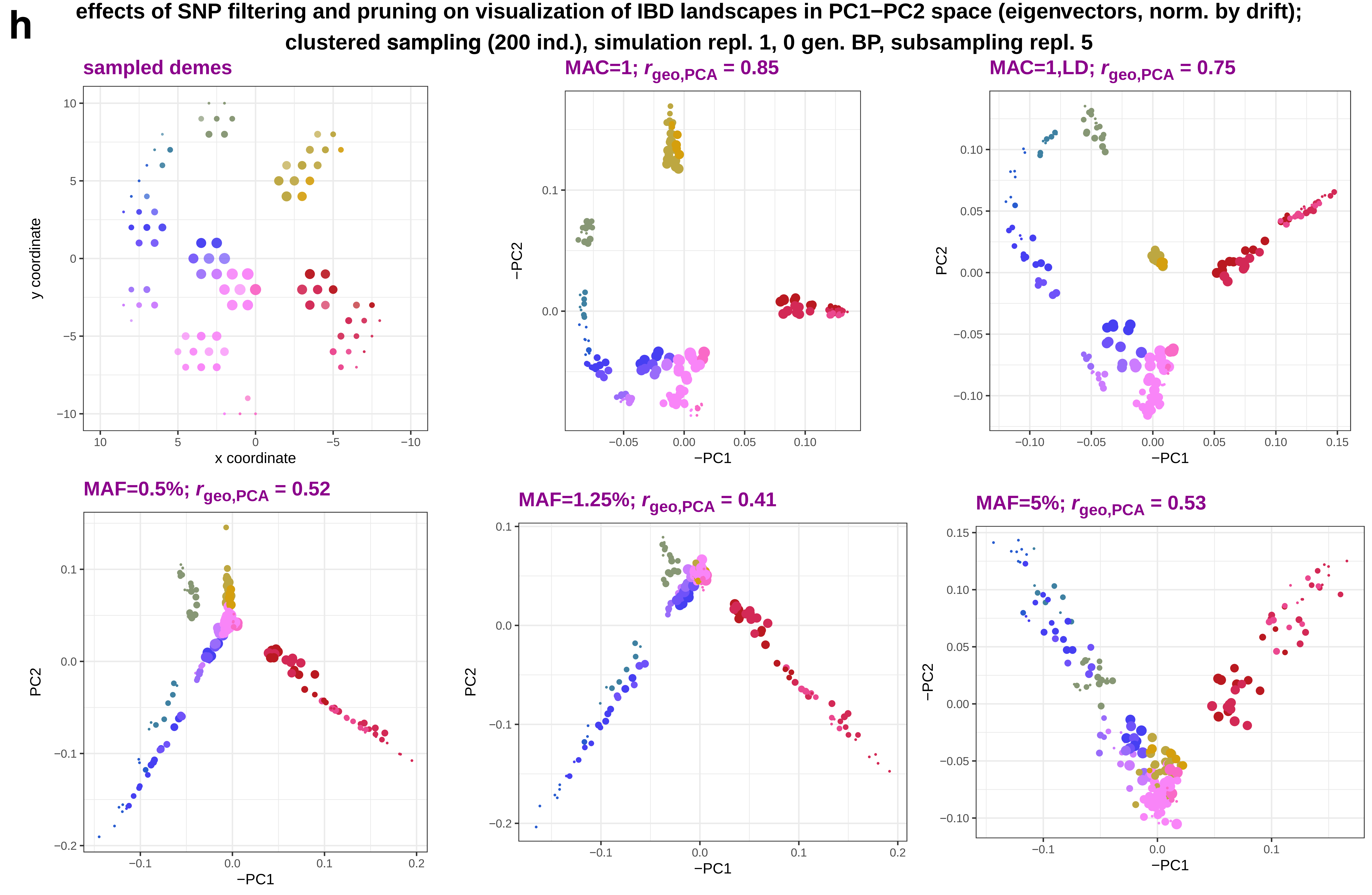
h
effects of SNP filtering and pruning on visualization of IBD landscapes in PC1−PC2 space (eigenvectors, norm. by drift);
clustered sampling (200 ind.), simulation repl. 1, 0 gen. BP, subsampling repl. 5
sampled demes
y coordinate
x coordinate
MAC=1; $r_{geo,PCA}$ = 0.85
−PC2
−PC1
MAC=1,LD; $r_{geo,PCA}$ = 0.75
PC2
−PC1
MAF=0.5%; $r_{geo,PCA}$ = 0.52
PC2
−PC1
MAF=1.25%; $r_{geo,PCA}$ = 0.41
PC2
−PC1
MAF=5%; $r_{geo,PCA}$ = 0.53
−PC2
−PC1

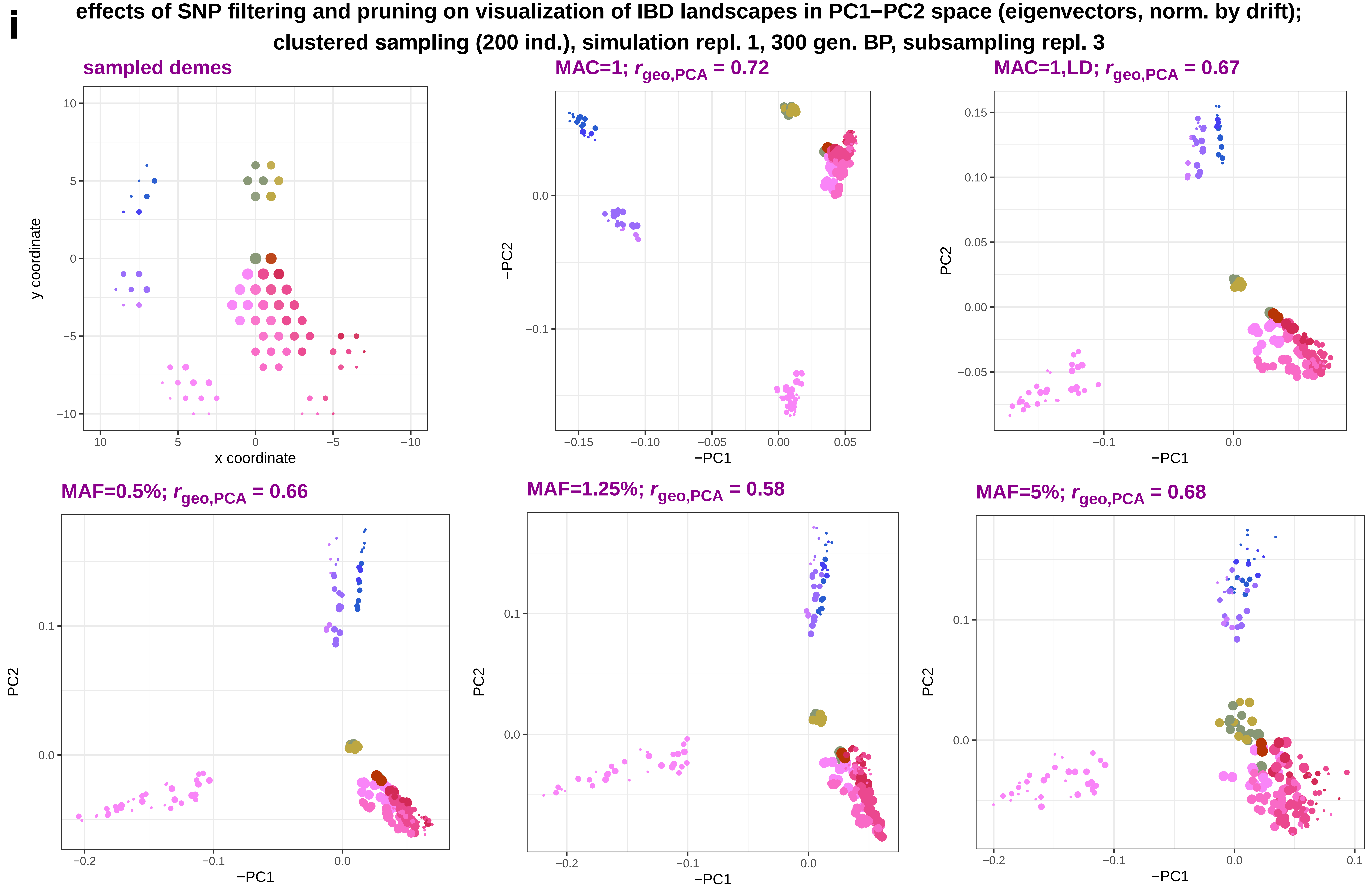
i
effects of SNP filtering and pruning on visualization of IBD landscapes in PC1−PC2 space (eigenvectors, norm. by drift);
clustered sampling (200 ind.), simulation repl. 1, 300 gen. BP, subsampling repl. 3
sampled demes
MAC=1; $r_{geo,PCA}$ = 0.72
MAC=1,LD; $r_{geo,PCA}$ = 0.67
MAF=0.5%; $r_{geo,PCA}$ = 0.66
MAF=1.25%; $r_{geo,PCA}$ = 0.58
MAF=5%; $r_{geo,PCA}$ = 0.68
y coordinate
x coordinate
−PC1
−PC2
PC2

**j** **effects of SNP filtering and pruning on visualization of IBD landscapes in PC1−PC2 space (eigenvectors, norm. by drift); clustered sampling (200 ind.), simulation repl. 1, 0 gen. BP, subsampling repl. 2**

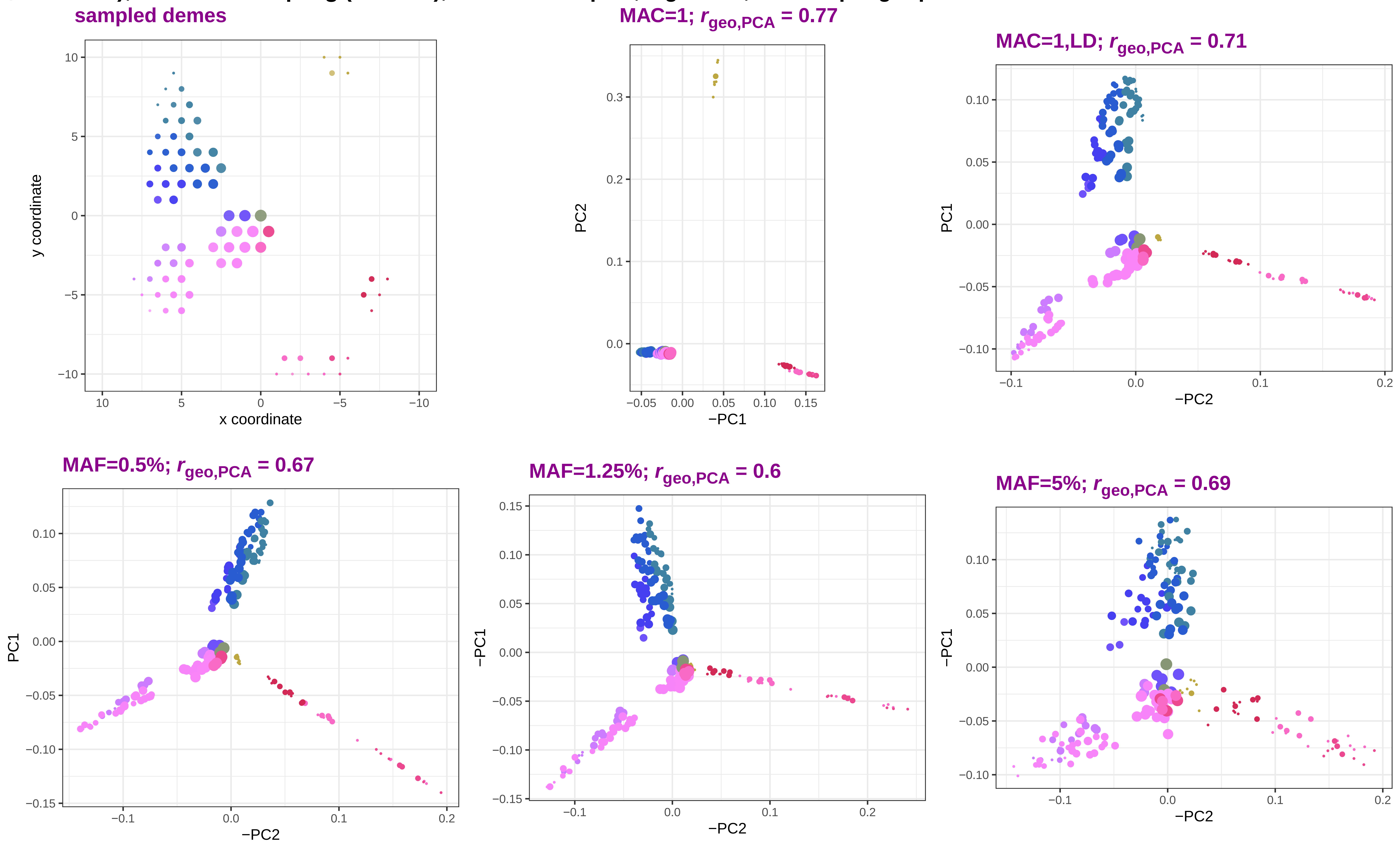

**k** **effects of SNP filtering and pruning on visualization of IBD landscapes in PC1−PC2 space (eigenvectors, norm. by drift); clustered sampling (200 ind.), simulation repl. 2, 0 gen. BP, subsampling repl. 3**

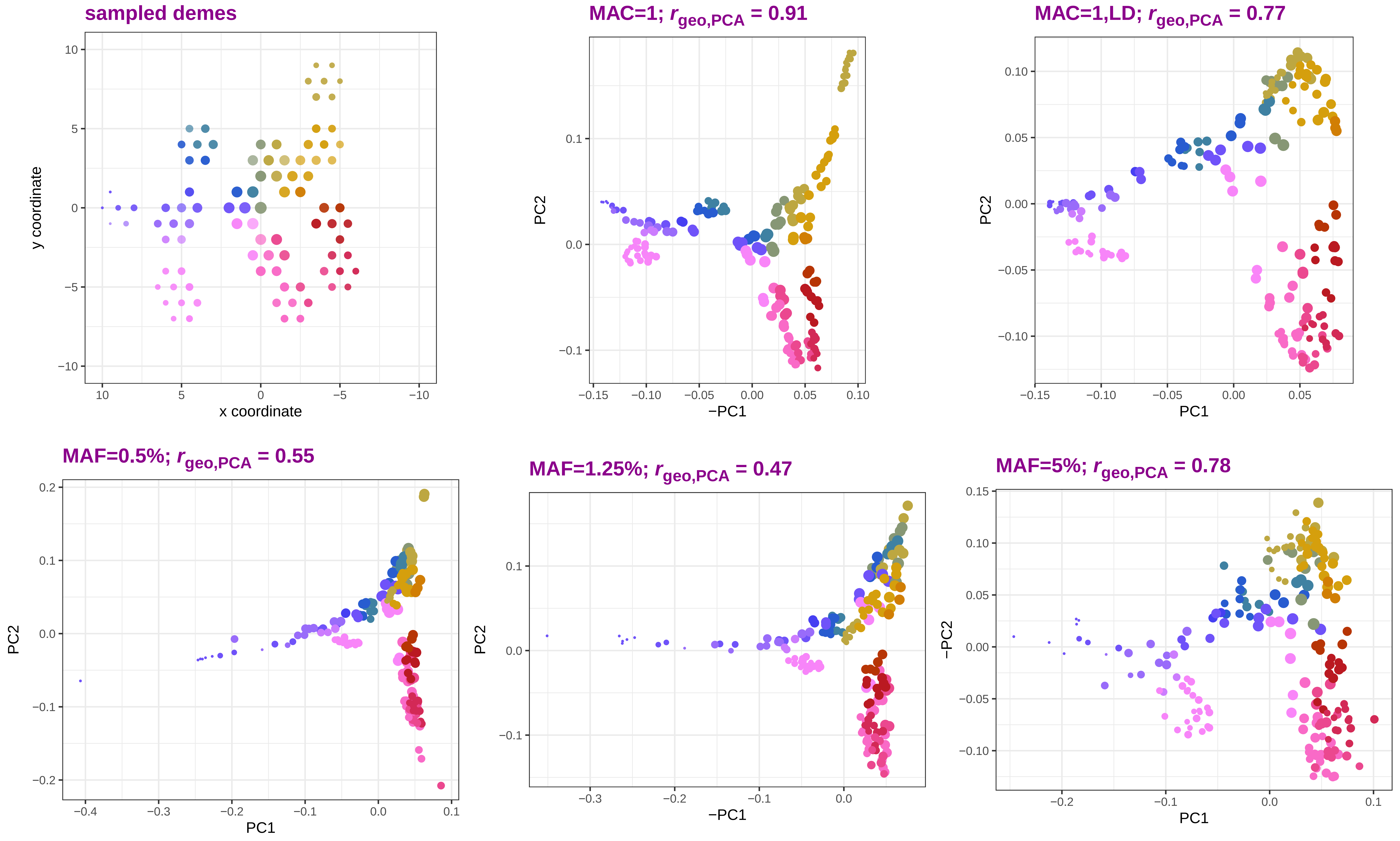

I

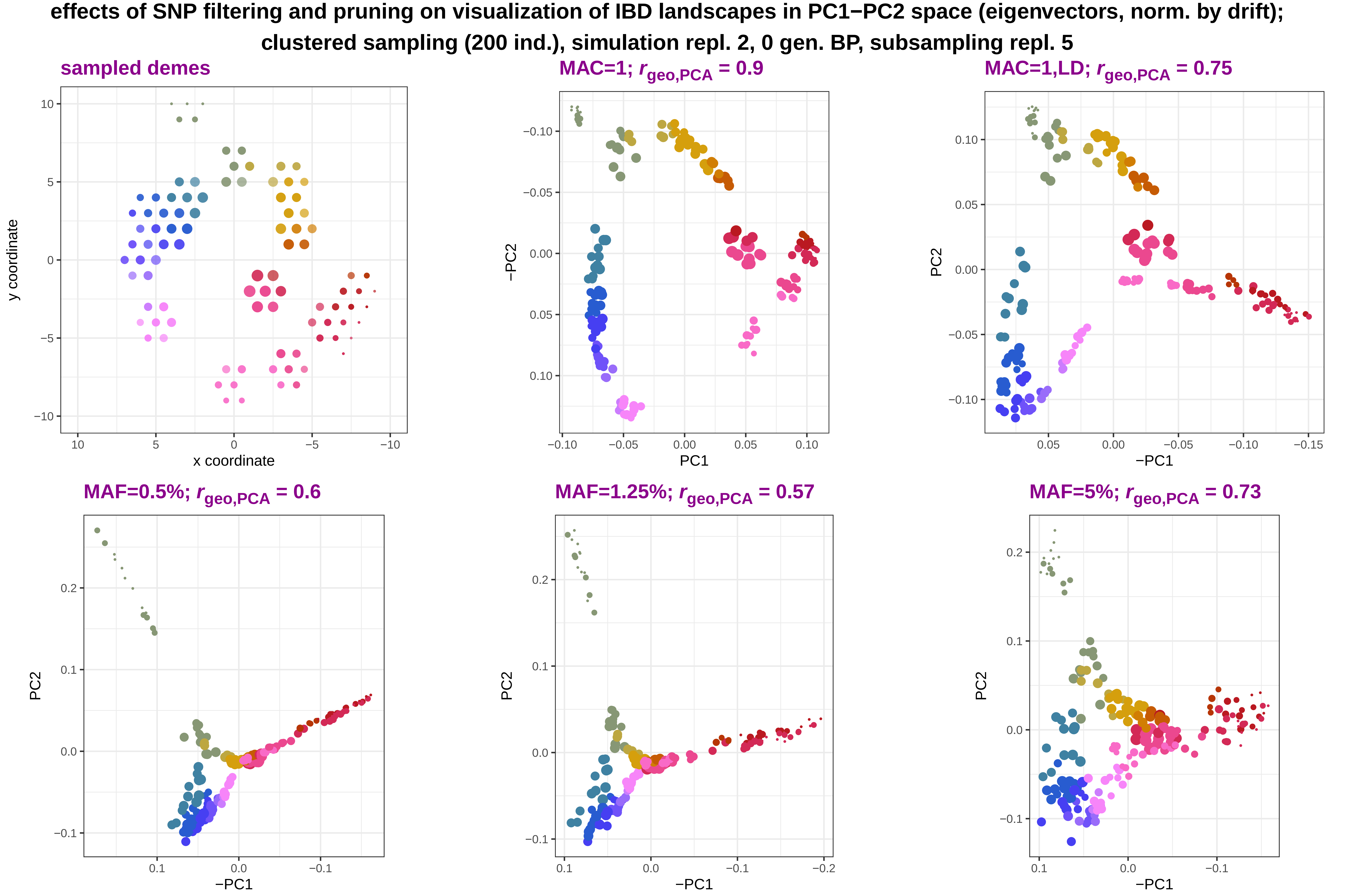

effects of SNP filtering and pruning on visualization of IBD landscapes in PC1−PC2 space (eigenvectors, norm. by drift);
clustered sampling (200 ind.), simulation repl. 2, 0 gen. BP, subsampling repl. 5
sampled demes
MAC=1; $r_{geo,PCA}$ = 0.9
MAC=1,LD; $r_{geo,PCA}$ = 0.75
MAF=0.5%; $r_{geo,PCA}$ = 0.6
MAF=1.25%; $r_{geo,PCA}$ = 0.57
MAF=5%; $r_{geo,PCA}$ = 0.73
y coordinate
x coordinate
−PC2
PC1
PC2
−PC1

**m** **effects of SNP filtering and pruning on visualization of IBD landscapes in PC1−PC2 space (eigenvectors, norm. by drift); clustered sampling (200 ind.), simulation repl. 3, 0 gen. BP, subsampling repl. 1**

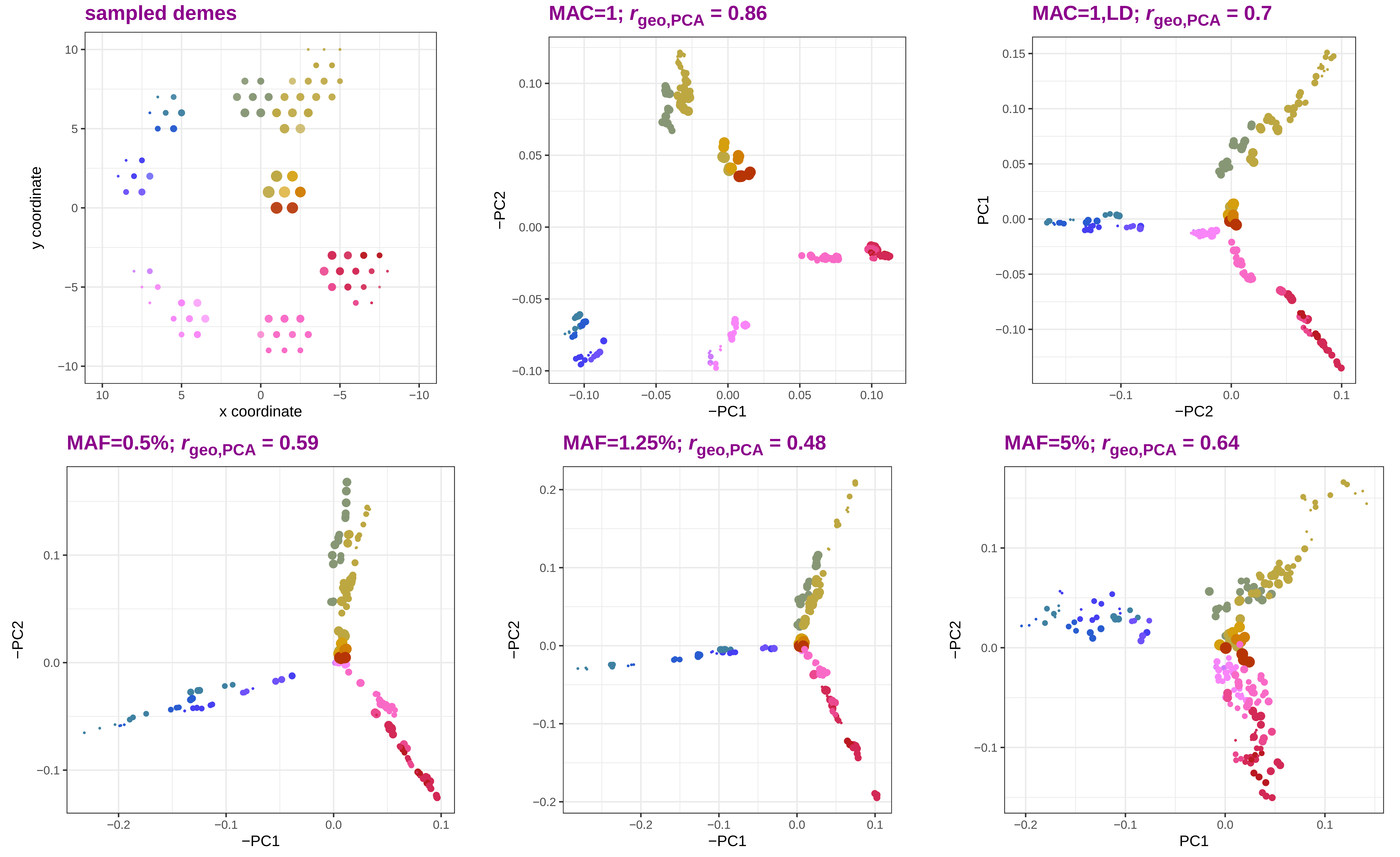

**n**

**effects of SNP filtering and pruning on visualization of IBD landscapes in PC1−PC2 space (eigenvectors, norm. by drift); clustered sampling (200 ind.), simulation repl. 3, 0 gen. BP, subsampling repl. 5**

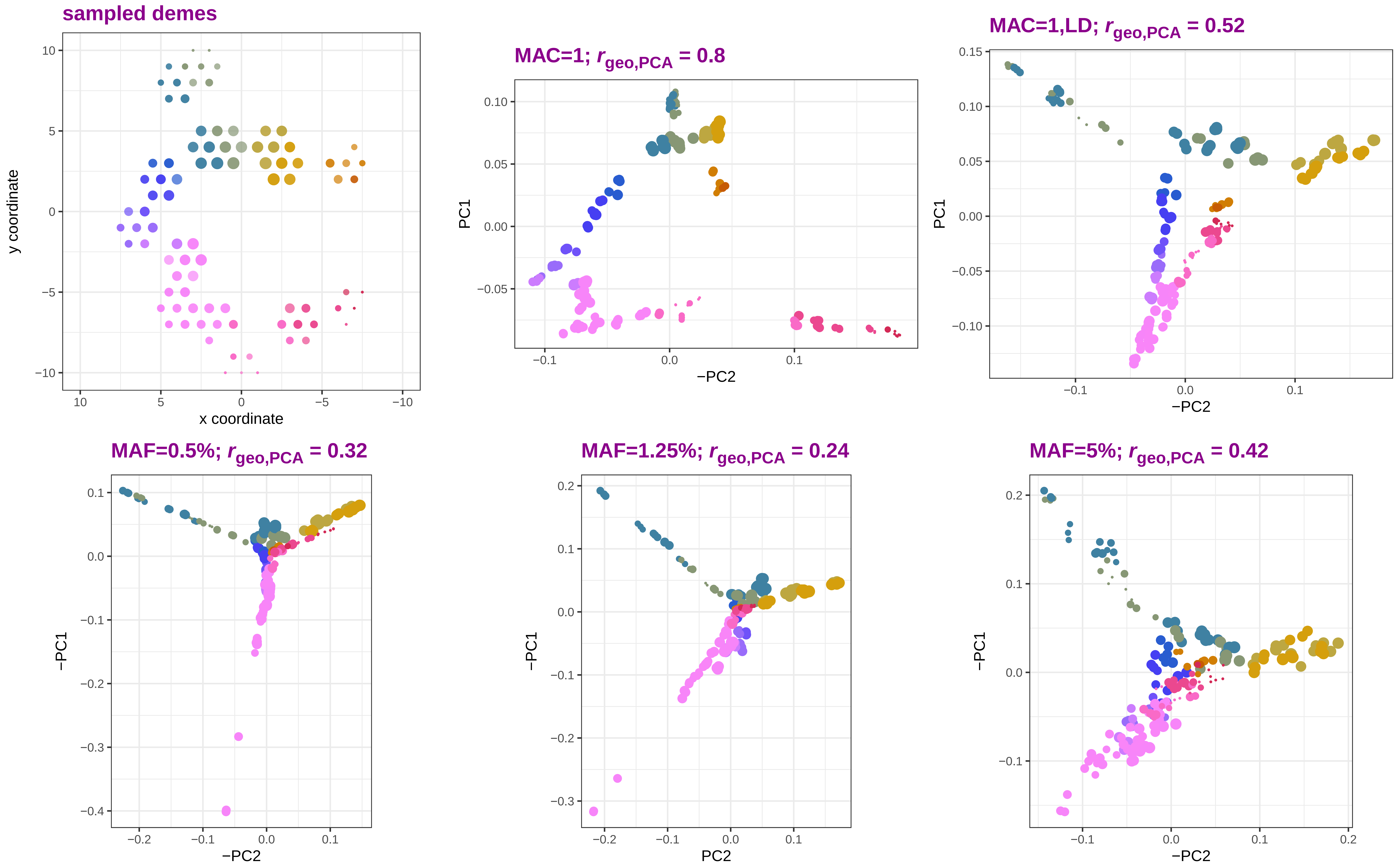

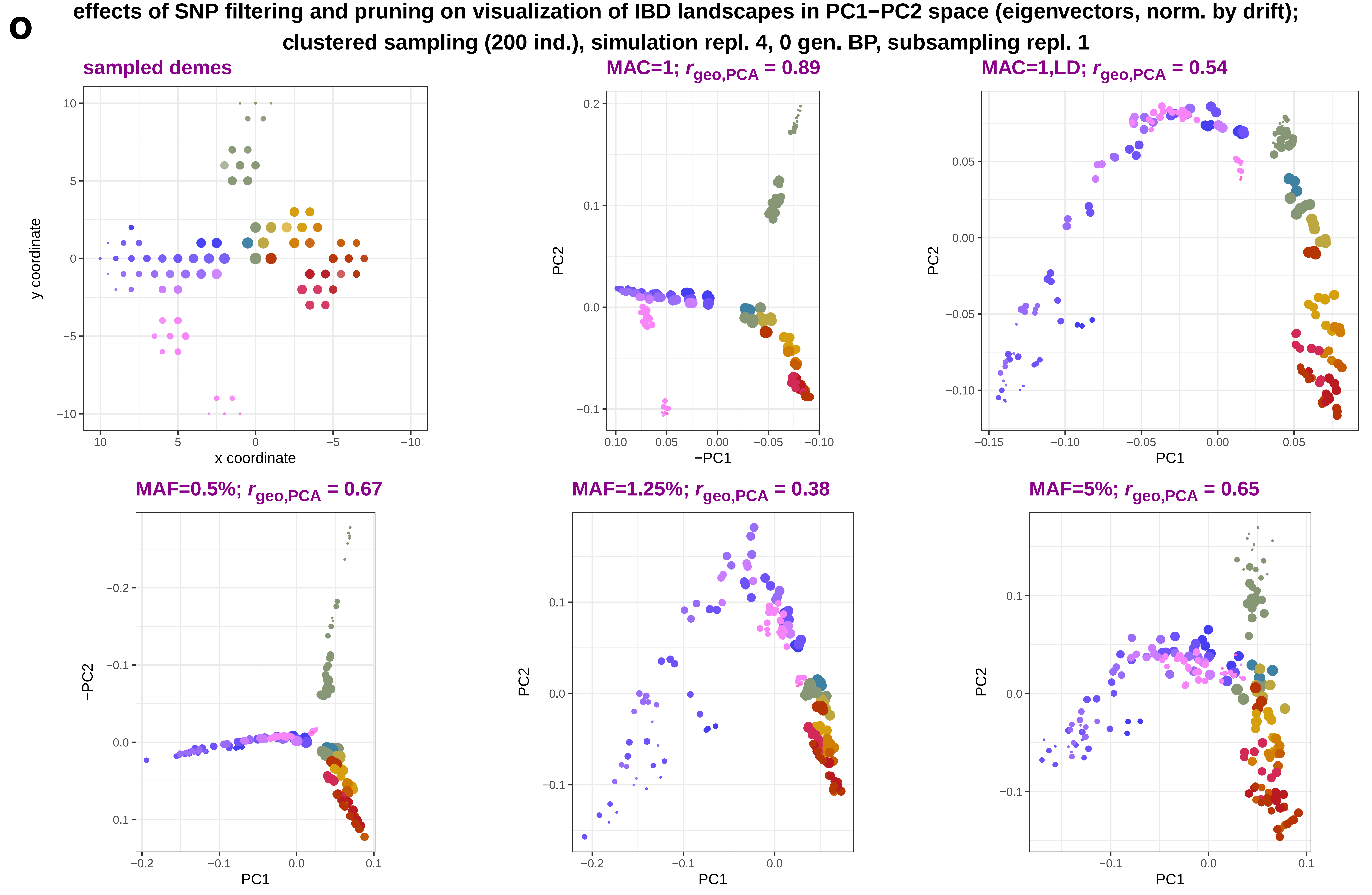
o
effects of SNP filtering and pruning on visualization of IBD landscapes in PC1−PC2 space (eigenvectors, norm. by drift);
clustered sampling (200 ind.), simulation repl. 4, 0 gen. BP, subsampling repl. 1
sampled demes
y coordinate
x coordinate
MAC=1; r geo,PCA = 0.89
PC2
−PC1
MAC=1,LD; r geo,PCA = 0.54
PC2
PC1
MAF=0.5%; r geo,PCA = 0.67
−PC2
PC1
MAF=1.25%; r geo,PCA = 0.38
PC2
PC1
MAF=5%; r geo,PCA = 0.65
PC2
PC1

**p** effects of SNP filtering and pruning on visualization of IBD landscapes in PC1−PC2 space (eigenvectors, norm. by drift); clustered sampling (200 ind.), simulation repl. 5, 300 gen. BP, subsampling repl. 5

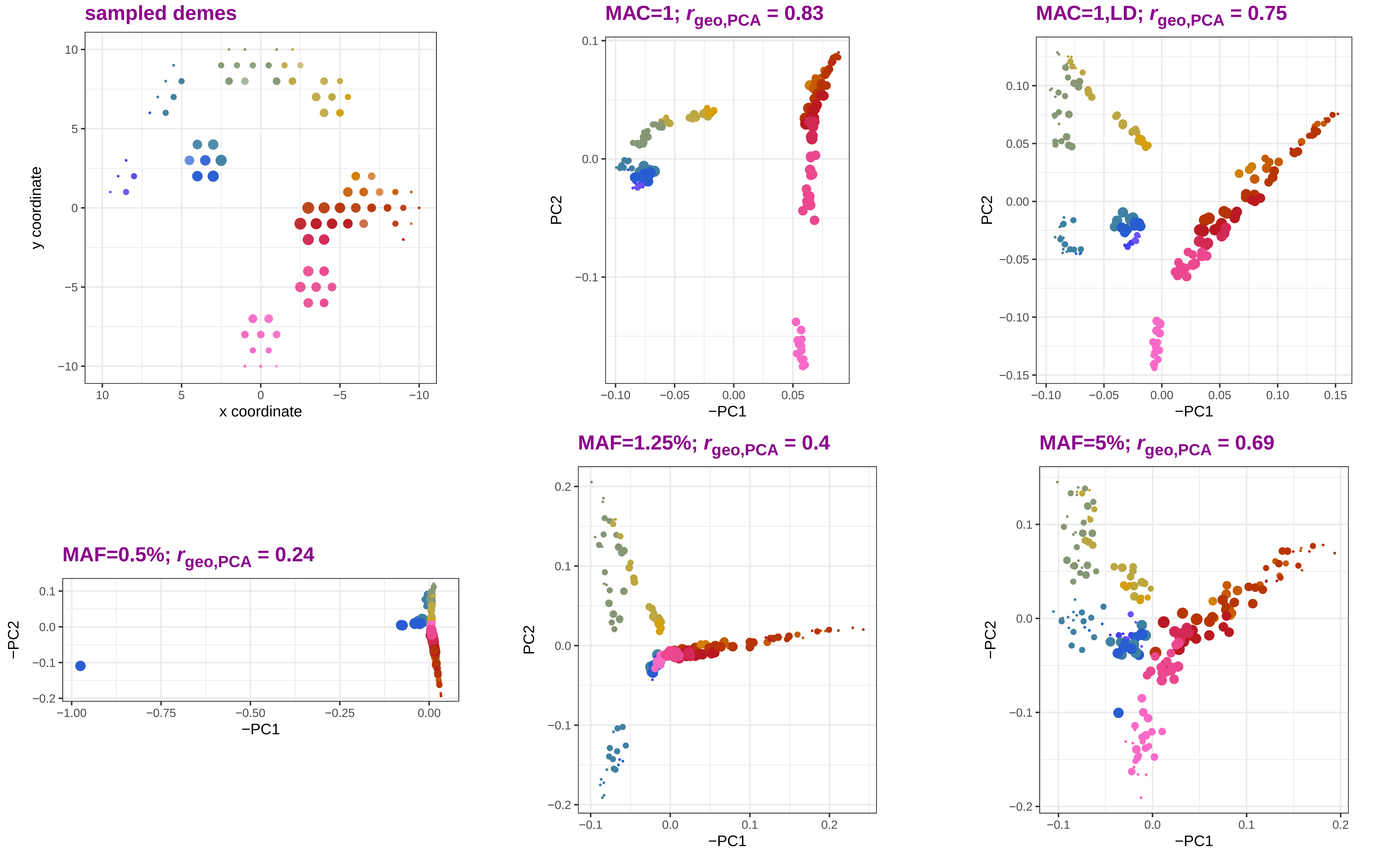
