## Supplementary Figure 4 for "Sparse sampling and rare-variant depletion distort PCA visualizations of population structure: recovery with objective-guided manifold learning"

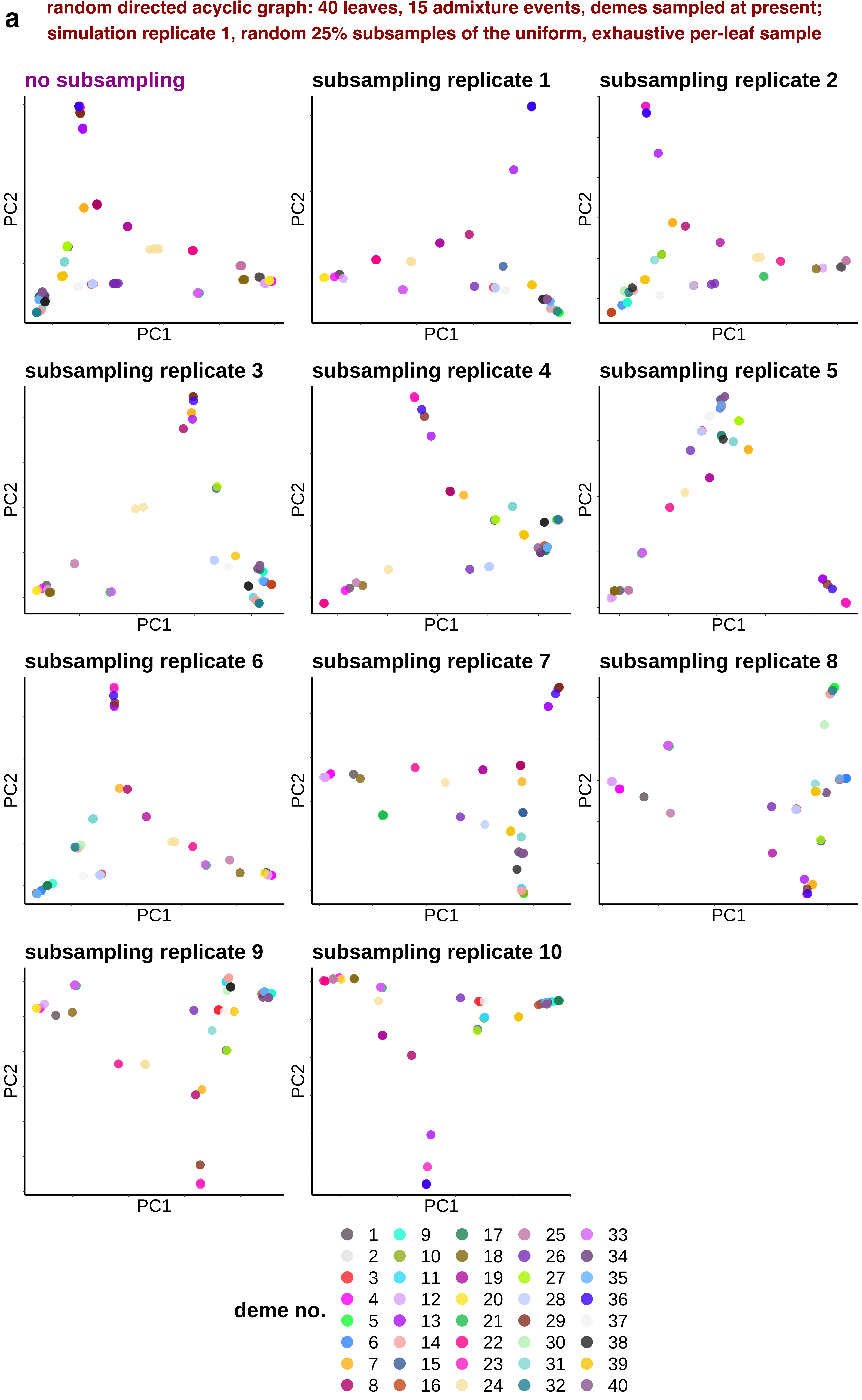
a
random directed acyclic graph: 40 leaves, 15 admixture events, demes sampled at present;
simulation replicate 1, random 25% subsamples of the uniform, exhaustive per-leaf sample
no subsampling
subsampling replicate 1
subsampling replicate 2
subsampling replicate 3
subsampling replicate 4
subsampling replicate 5
subsampling replicate 6
subsampling replicate 7
subsampling replicate 8
subsampling replicate 9
subsampling replicate 10
PC1
PC2
deme no.
1 2 3 4 5 6 7 8 9 10 11 12 13 14 15 16 17 18 19 20 21 22 23 24 25 26 27 28 29 30 31 32 33 34 35 36 37 38 39 40

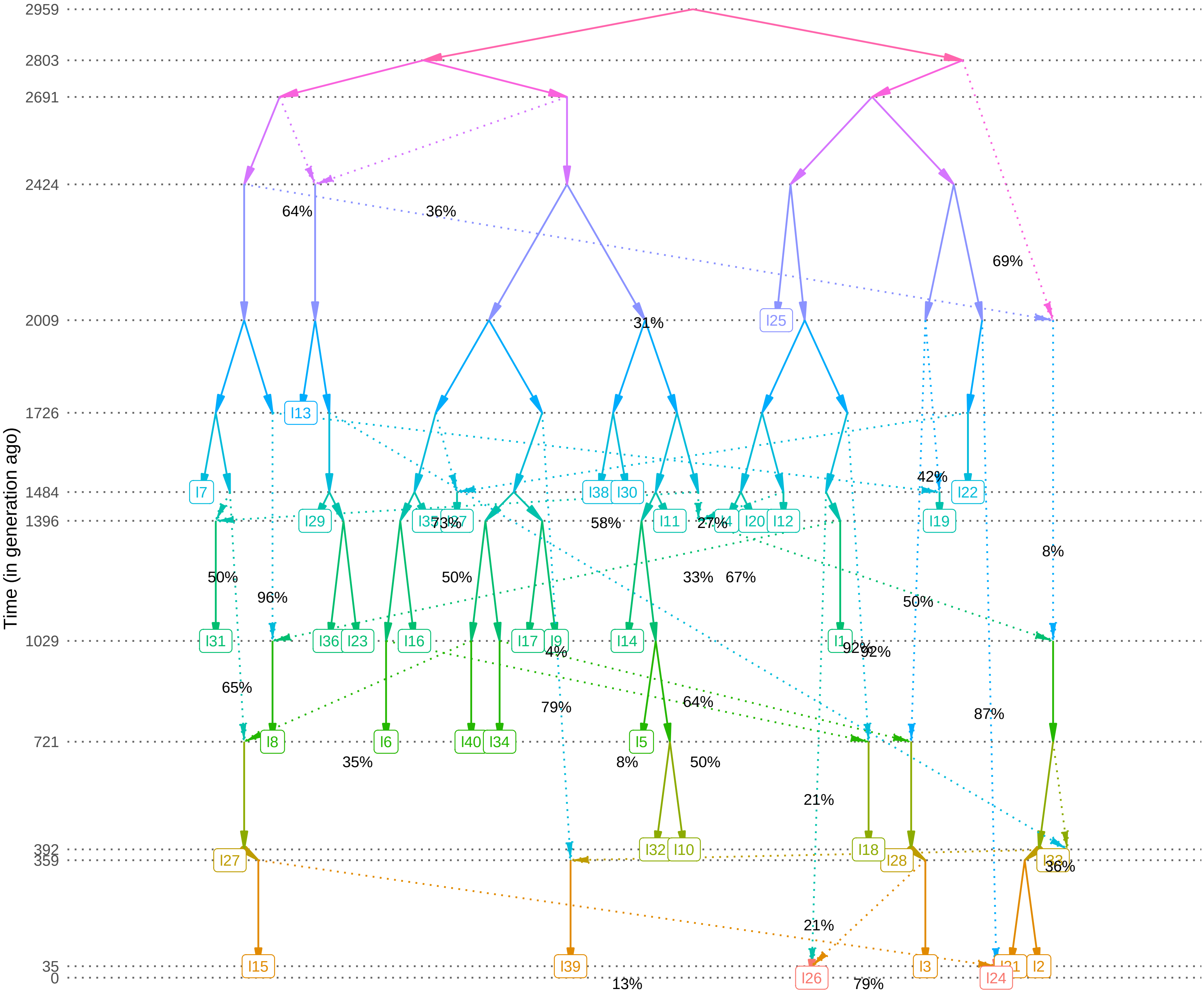
Time (in generation ago)
2959
2803
2691
2424
2009
1726
1484
1396
1029
721
392
359
35
0
64%
36%
69%
31%
I25
I13
I7
I38
I30
I22
42%
I29
73%
58%
I11
27%
I20
I12
I19
8%
50%
50%
96%
33%
67%
50%
I31
I36
I23
I16
I17
I9
I14
I1
92%
4%
65%
79%
64%
87%
I8
I6
I40
I34
I5
35%
8%
50%
21%
I32
I10
I18
I27
I28
36%
21%
I15
I39
I26
I3
I24
I2
13%
79%

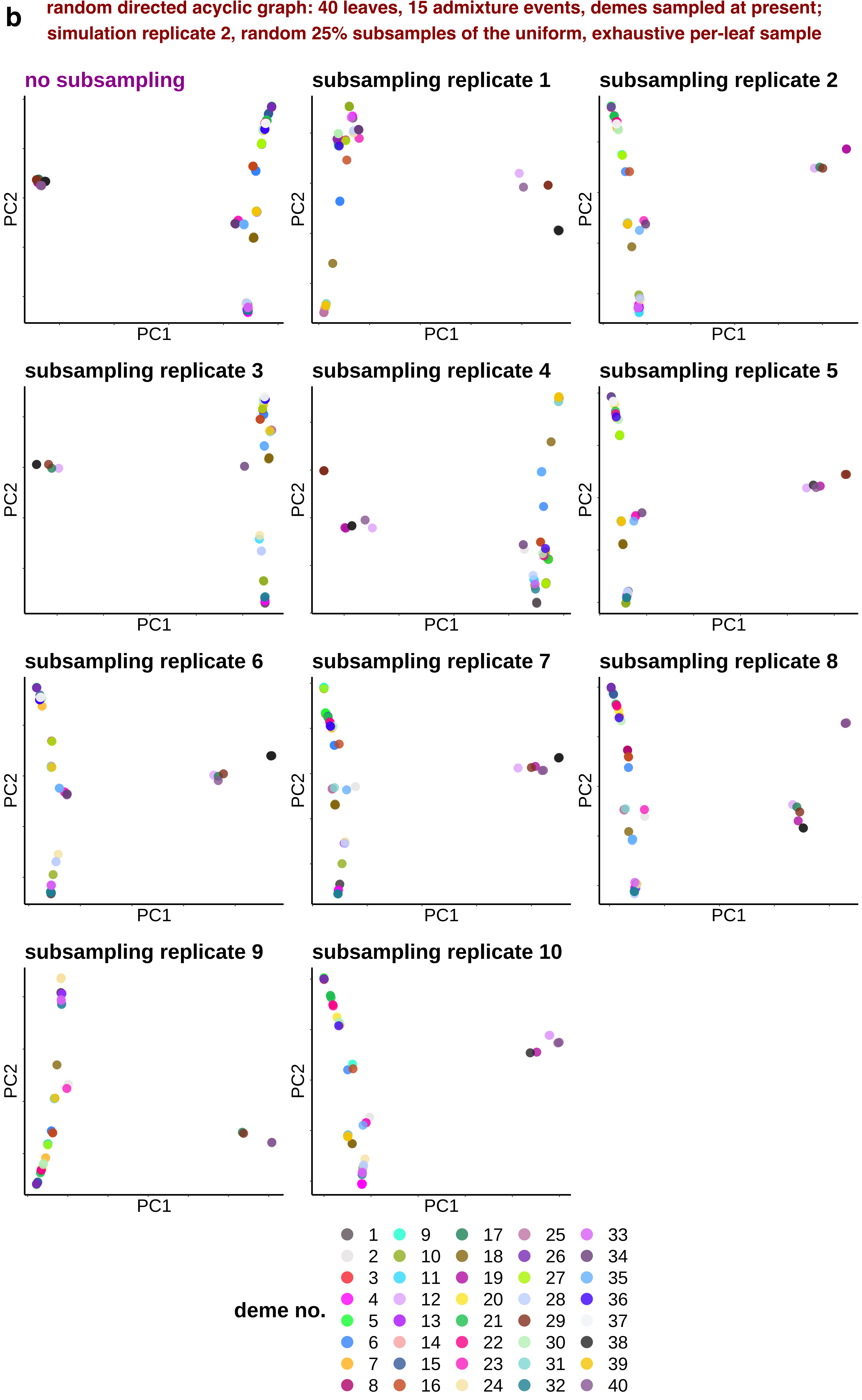
b
random directed acyclic graph: 40 leaves, 15 admixture events, demes sampled at present;
simulation replicate 2, random 25% subsamples of the uniform, exhaustive per-leaf sample
no subsampling
subsampling replicate 1
subsampling replicate 2
subsampling replicate 3
subsampling replicate 4
subsampling replicate 5
subsampling replicate 6
subsampling replicate 7
subsampling replicate 8
subsampling replicate 9
subsampling replicate 10
PC1
PC2
deme no.
1 2 3 4 5 6 7 8 9 10 11 12 13 14 15 16 17 18 19 20 21 22 23 24 25 26 27 28 29 30 31 32 33 34 35 36 37 38 39 40

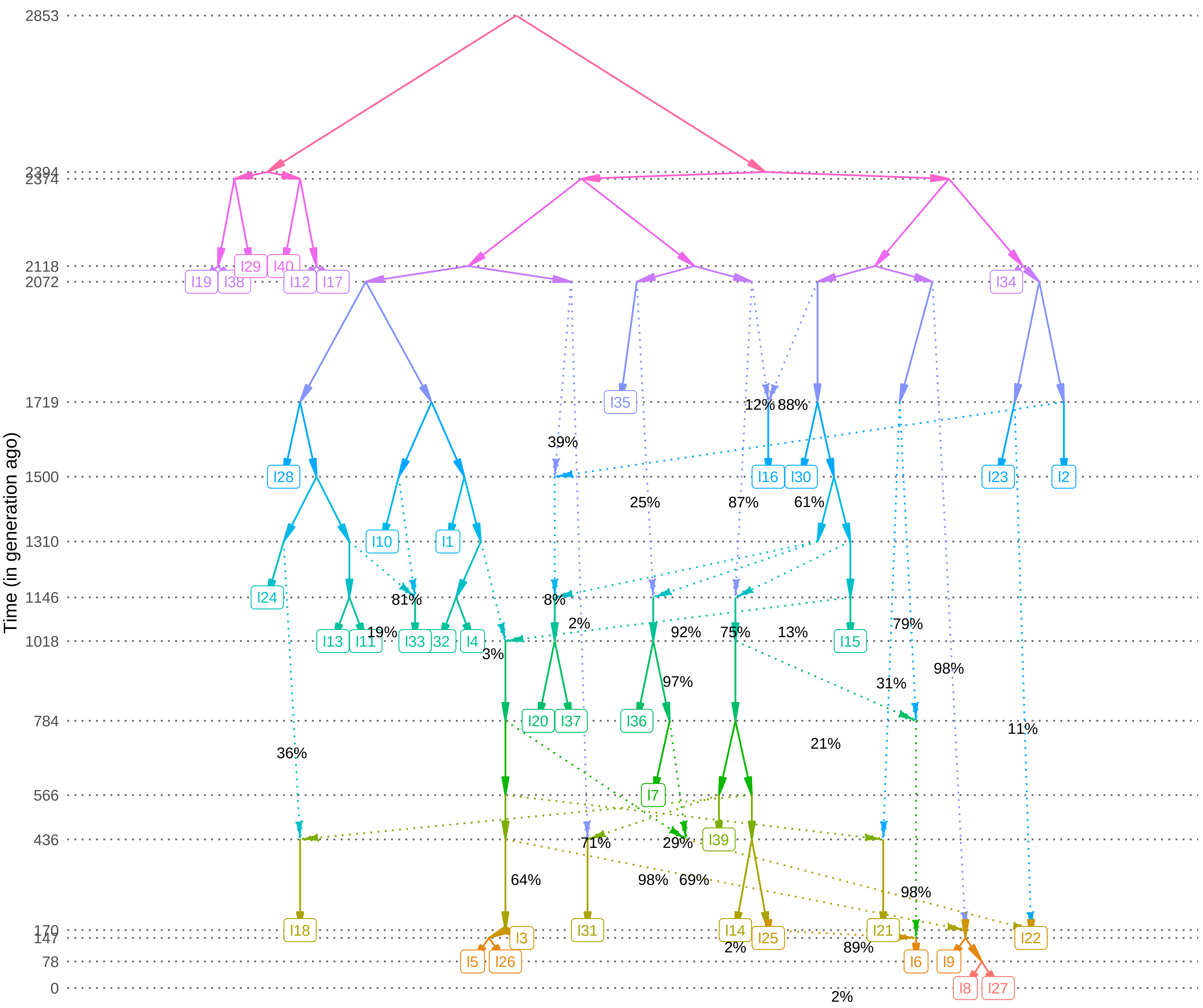
Time (in generation ago)
2853
2394
2374
2118
2072
1719
1500
1310
1146
1018
784
566
436
170
147
78
0
I19
I38
I29
I40
I12
I17
I34
I35
I28
I16
I30
I23
I2
I10
I1
I24
I13
I11
I33
I32
I4
I15
I20
I37
I36
I7
I39
I18
I31
I14
I25
I21
I22
I3
I5
I26
I6
I9
I8
I27
39%
12%
88%
25%
87%
61%
81%
19%
8%
2%
3%
92%
75%
13%
79%
97%
98%
31%
11%
21%
36%
71%
29%
64%
98%
69%
98%
2%
89%
2%

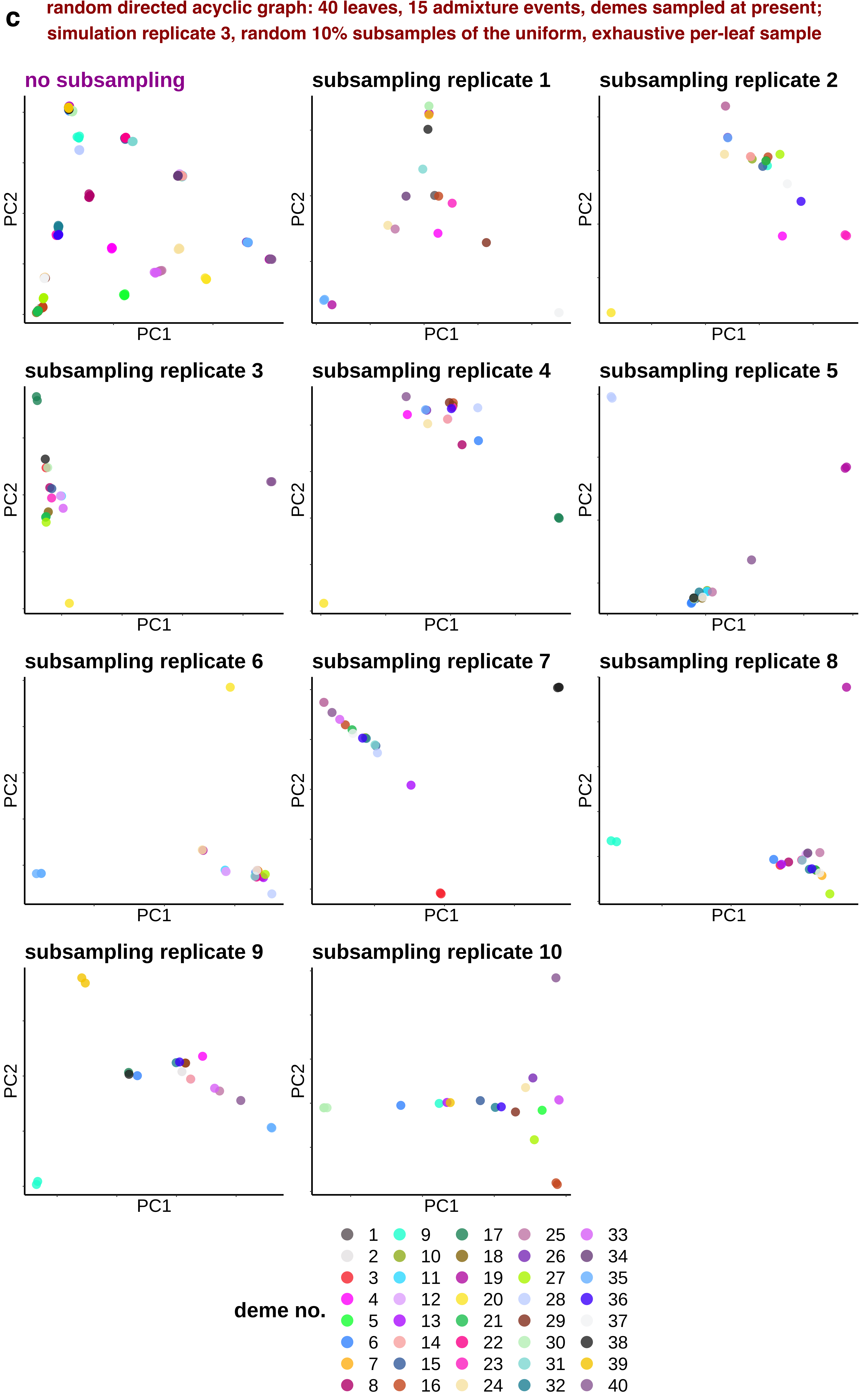
c
random directed acyclic graph: 40 leaves, 15 admixture events, demes sampled at present;
simulation replicate 3, random 10% subsamples of the uniform, exhaustive per-leaf sample
no subsampling
subsampling replicate 1
subsampling replicate 2
subsampling replicate 3
subsampling replicate 4
subsampling replicate 5
subsampling replicate 6
subsampling replicate 7
subsampling replicate 8
subsampling replicate 9
subsampling replicate 10
PC1
PC2
deme no.
1 2 3 4 5 6 7 8 9 10 11 12 13 14 15 16 17 18 19 20 21 22 23 24 25 26 27 28 29 30 31 32 33 34 35 36 37 38 39 40

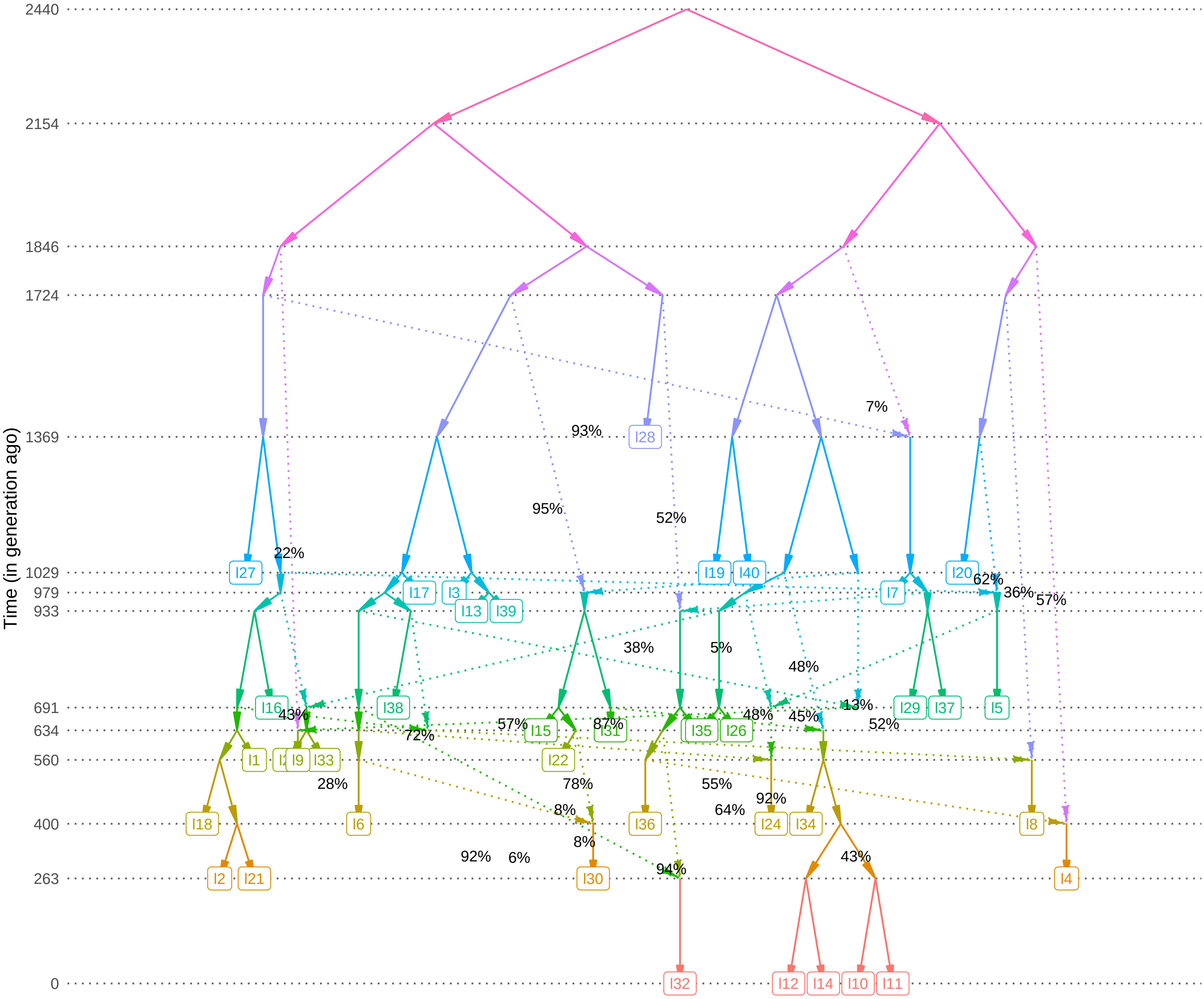

Time (in generation ago)
2440
2154
1846
1724
1369
1029
979
933
691
634
560
400
263
0
I27
I28
I19
I40
I20
I17
I3
I7
I13
I39
I16
I38
I29
I37
I5
I15
I31
I35
I26
I1
I9
I33
I22
I18
I6
I36
I24
I34
I8
I2
I21
I30
I4
I32
I12
I14
I10
I11
7%
93%
95%
52%
22%
62%
36%
57%
38%
5%
48%
13%
43%
48%
45%
52%
57%
87%
72%
28%
78%
55%
92%
8%
64%
8%
92%
6%
94%
43%

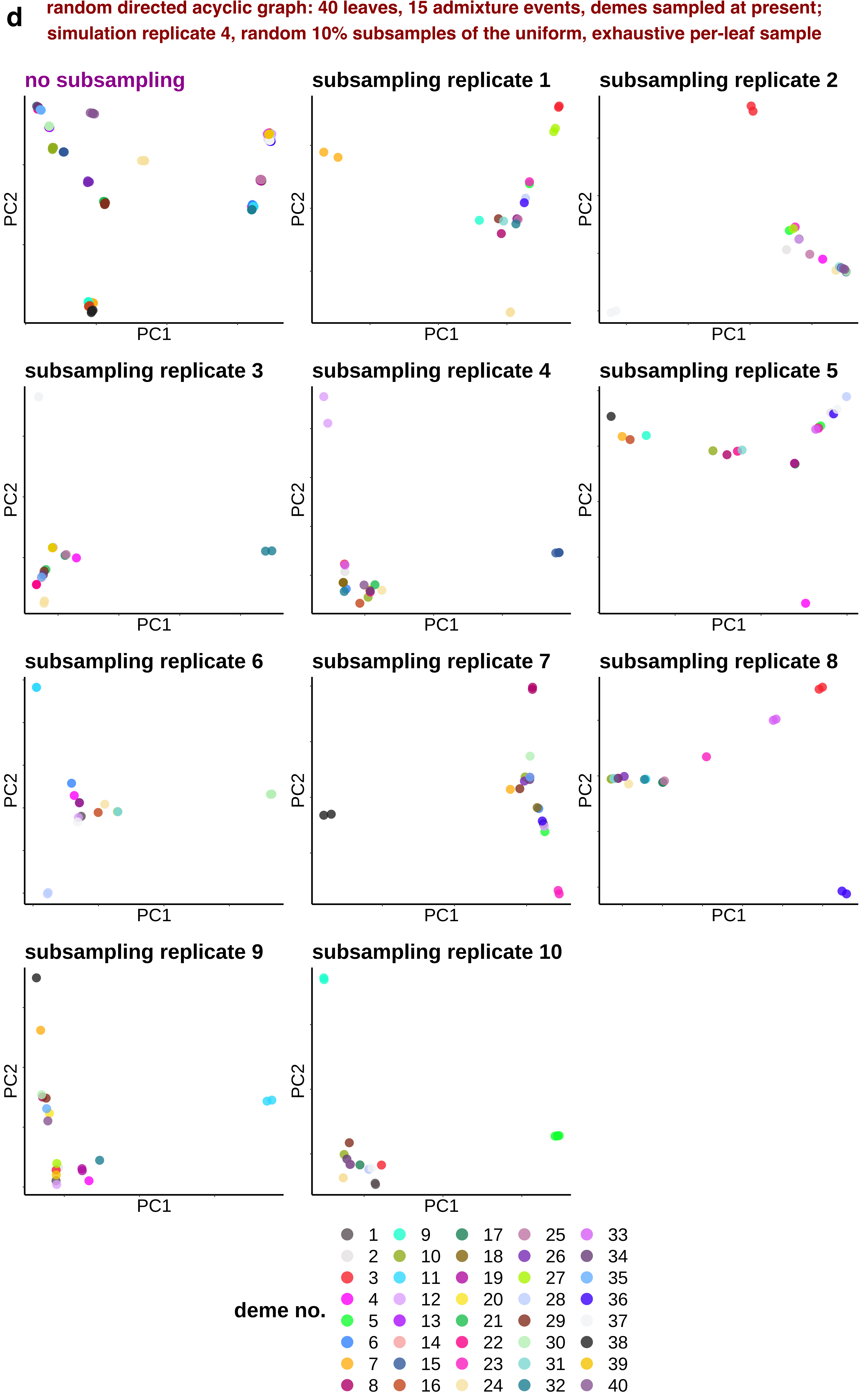

d
random directed acyclic graph: 40 leaves, 15 admixture events, demes sampled at present;
simulation replicate 4, random 10% subsamples of the uniform, exhaustive per-leaf sample
no subsampling
subsampling replicate 1
subsampling replicate 2
subsampling replicate 3
subsampling replicate 4
subsampling replicate 5
subsampling replicate 6
subsampling replicate 7
subsampling replicate 8
subsampling replicate 9
subsampling replicate 10
PC1
PC2
deme no.
1 2 3 4 5 6 7 8 9 10 11 12 13 14 15 16 17 18 19 20 21 22 23 24 25 26 27 28 29 30 31 32 33 34 35 36 37 38 39 40

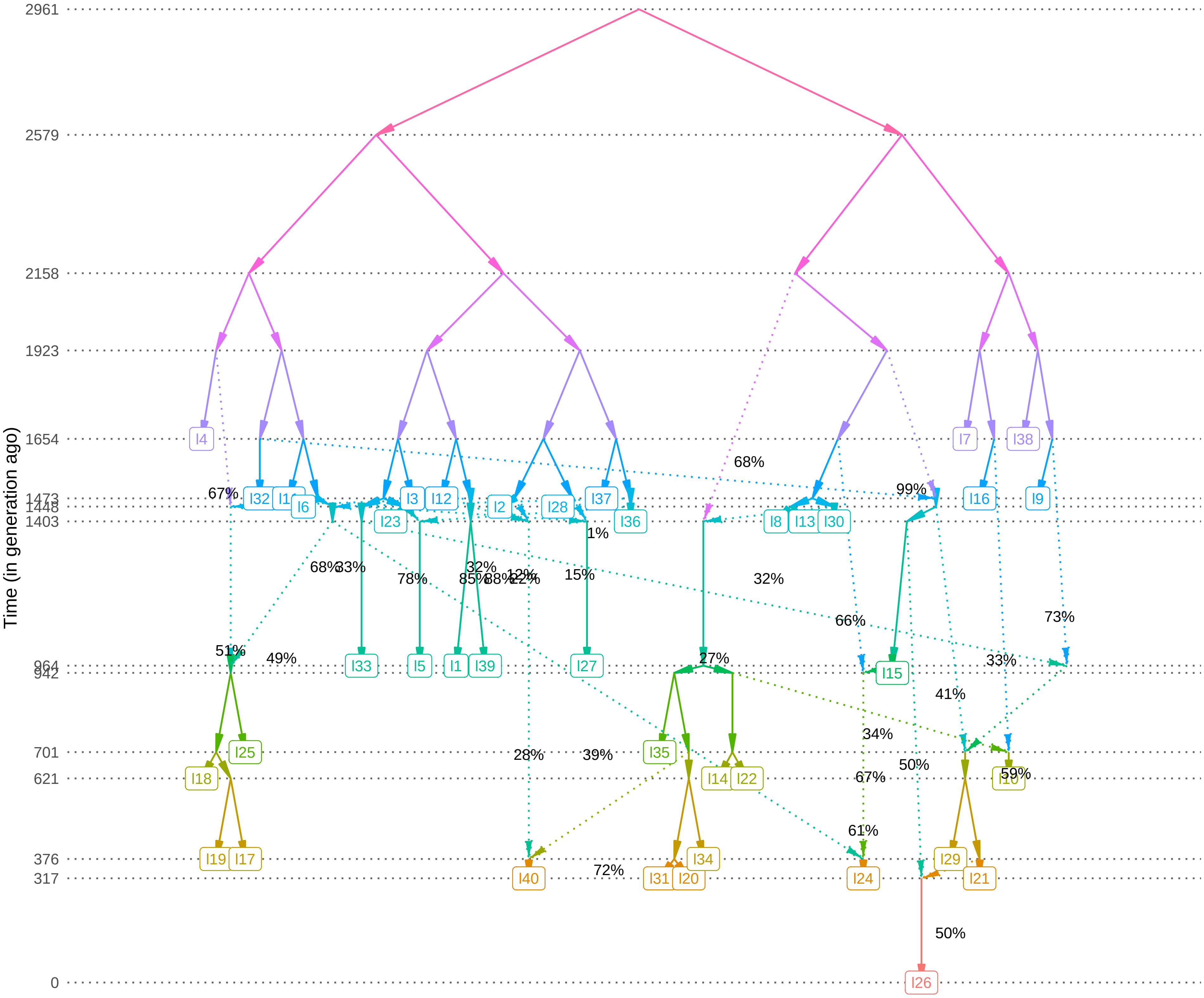

Time (in generation ago)
2961
2579
2158
1923
1654
1473
1448
1403
964
942
701
621
376
317
0
I4
I32
I6
I23
I3
I12
I2
I28
I37
I36
I8
I13
I30
I7
I38
I16
I9
I33
I5
I1
I39
I27
I15
I25
I18
I35
I14
I22
I10
I19
I17
I34
I29
I40
I31
I20
I24
I21
I26
67%
68%
33%
78%
32%
85%
88%
12%
15%
1%
68%
99%
32%
66%
73%
51%
49%
27%
33%
41%
34%
28%
39%
50%
67%
59%
61%
72%
50%

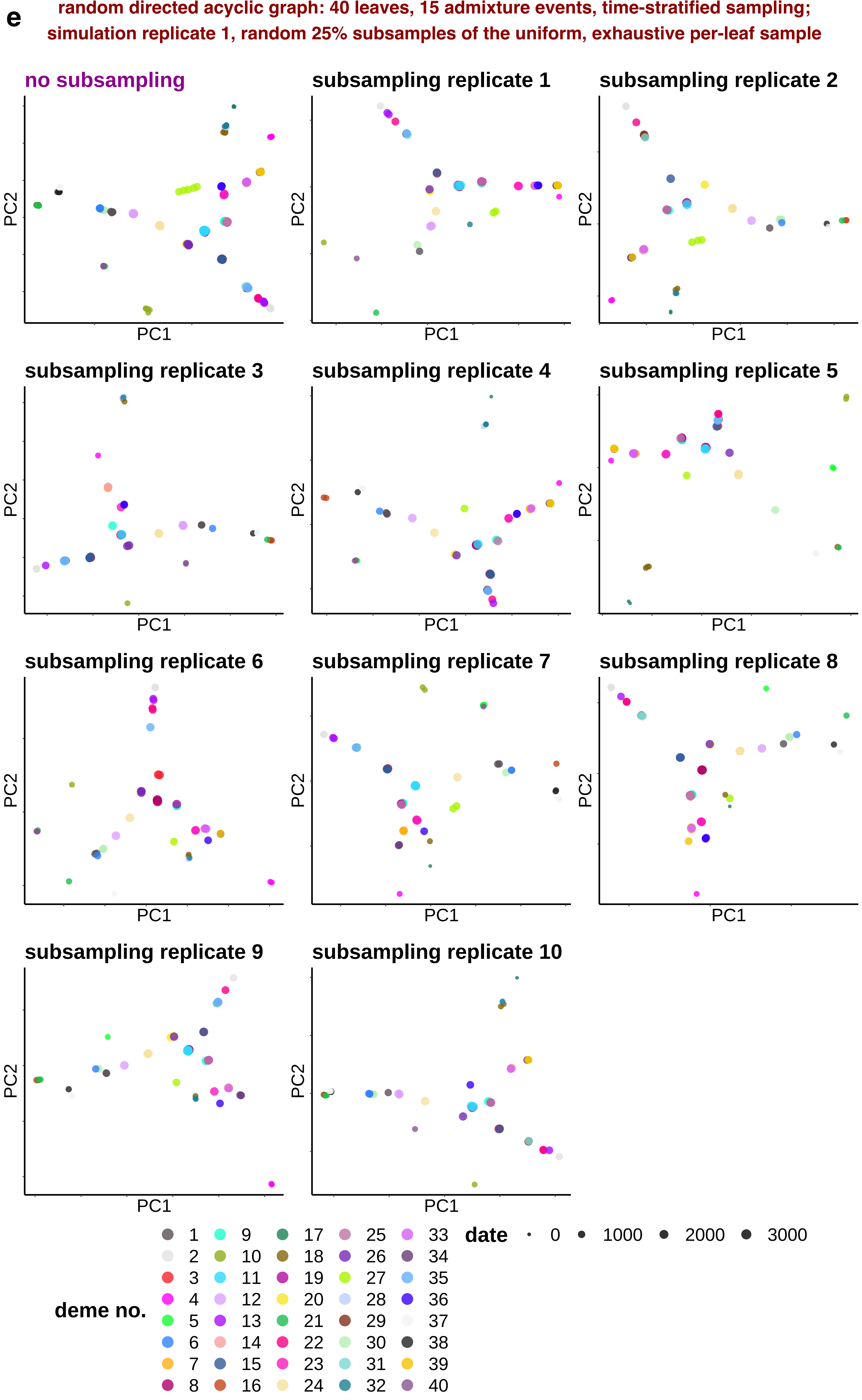
e
random directed acyclic graph: 40 leaves, 15 admixture events, time-stratified sampling;
simulation replicate 1, random 25% subsamples of the uniform, exhaustive per-leaf sample
no subsampling
subsampling replicate 1
subsampling replicate 2
subsampling replicate 3
subsampling replicate 4
subsampling replicate 5
subsampling replicate 6
subsampling replicate 7
subsampling replicate 8
subsampling replicate 9
subsampling replicate 10
PC1
PC2
deme no.
1 2 3 4 5 6 7 8 9 10 11 12 13 14 15 16 17 18 19 20 21 22 23 24 25 26 27 28 29 30 31 32 33 34 35 36 37 38 39 40
date
0
1000
2000
3000

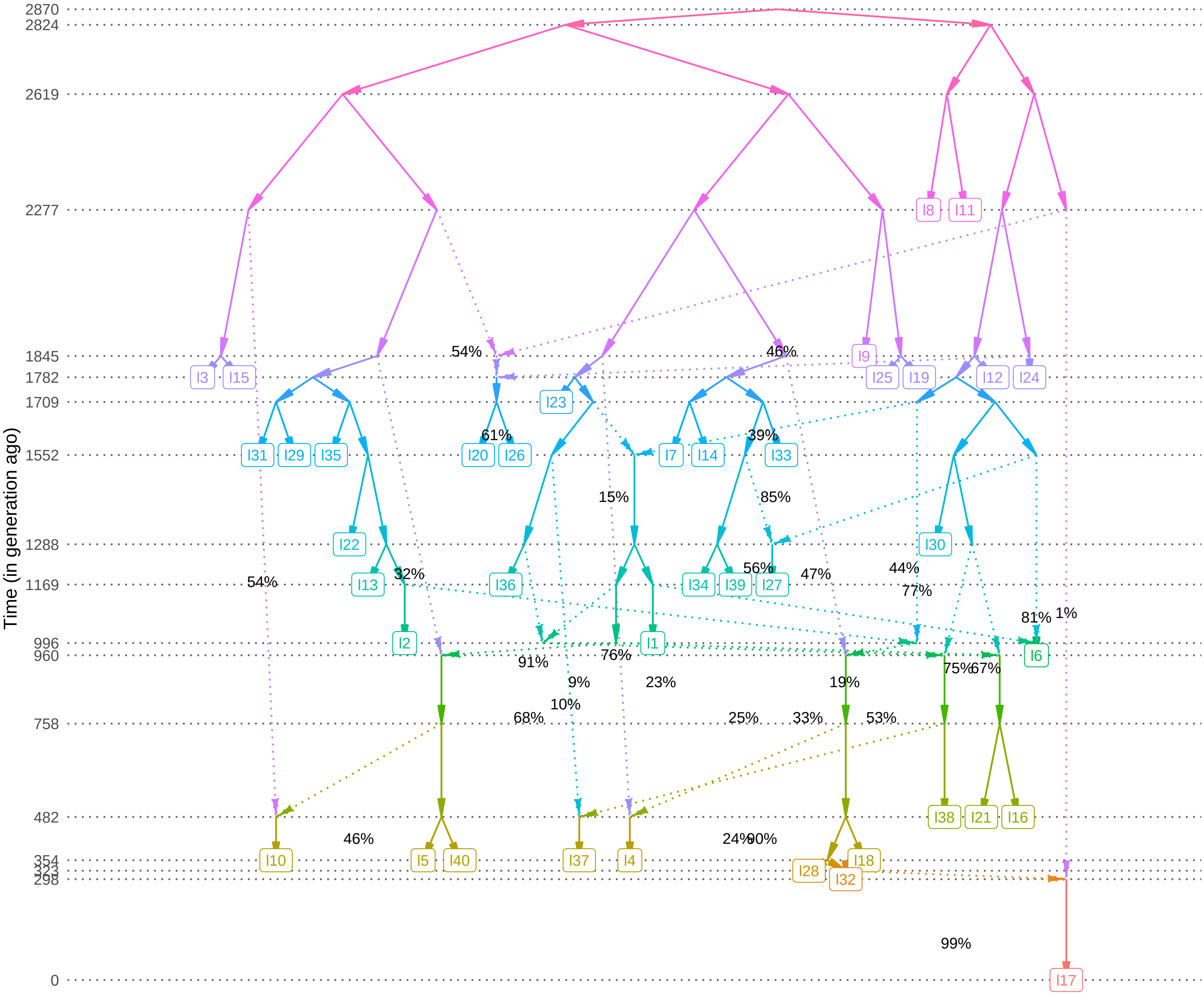

Time (in generation ago)
2870
2824
2619
2277
1845
1782
1709
1552
1288
1169
996
960
758
482
354
298
0
I8
I11
I9
I3
I15
I25
I19
I12
I24
I23
I31
I29
I35
I20
I26
I7
I14
I33
I22
I30
I13
I36
I34
I39
I27
I2
I1
I6
I38
I21
I16
I10
I5
I40
I37
I4
I28
I18
I32
I17
54%
46%
61%
39%
15%
85%
54%
32%
56%
47%
44%
77%
81%
1%
91%
76%
9%
23%
19%
75%
67%
10%
68%
25%
33%
53%
46%
24%
90%
99%

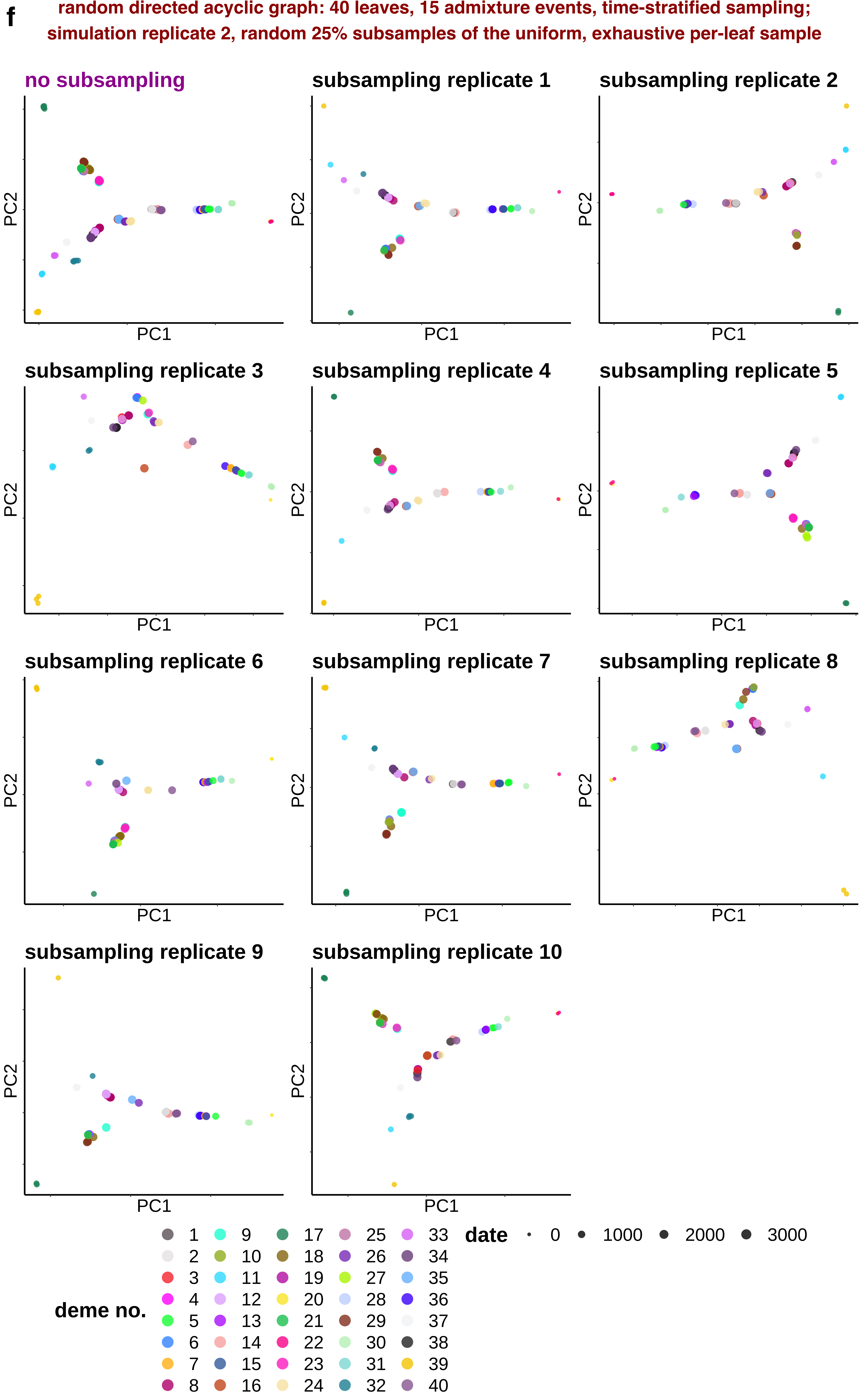
f
random directed acyclic graph: 40 leaves, 15 admixture events, time-stratified sampling;
simulation replicate 2, random 25% subsamples of the uniform, exhaustive per-leaf sample
no subsampling
subsampling replicate 1
subsampling replicate 2
subsampling replicate 3
subsampling replicate 4
subsampling replicate 5
subsampling replicate 6
subsampling replicate 7
subsampling replicate 8
subsampling replicate 9
subsampling replicate 10
PC1
PC2
deme no.
1 2 3 4 5 6 7 8 9 10 11 12 13 14 15 16 17 18 19 20 21 22 23 24 25 26 27 28 29 30 31 32 33 34 35 36 37 38 39 40
date
0 1000 2000 3000

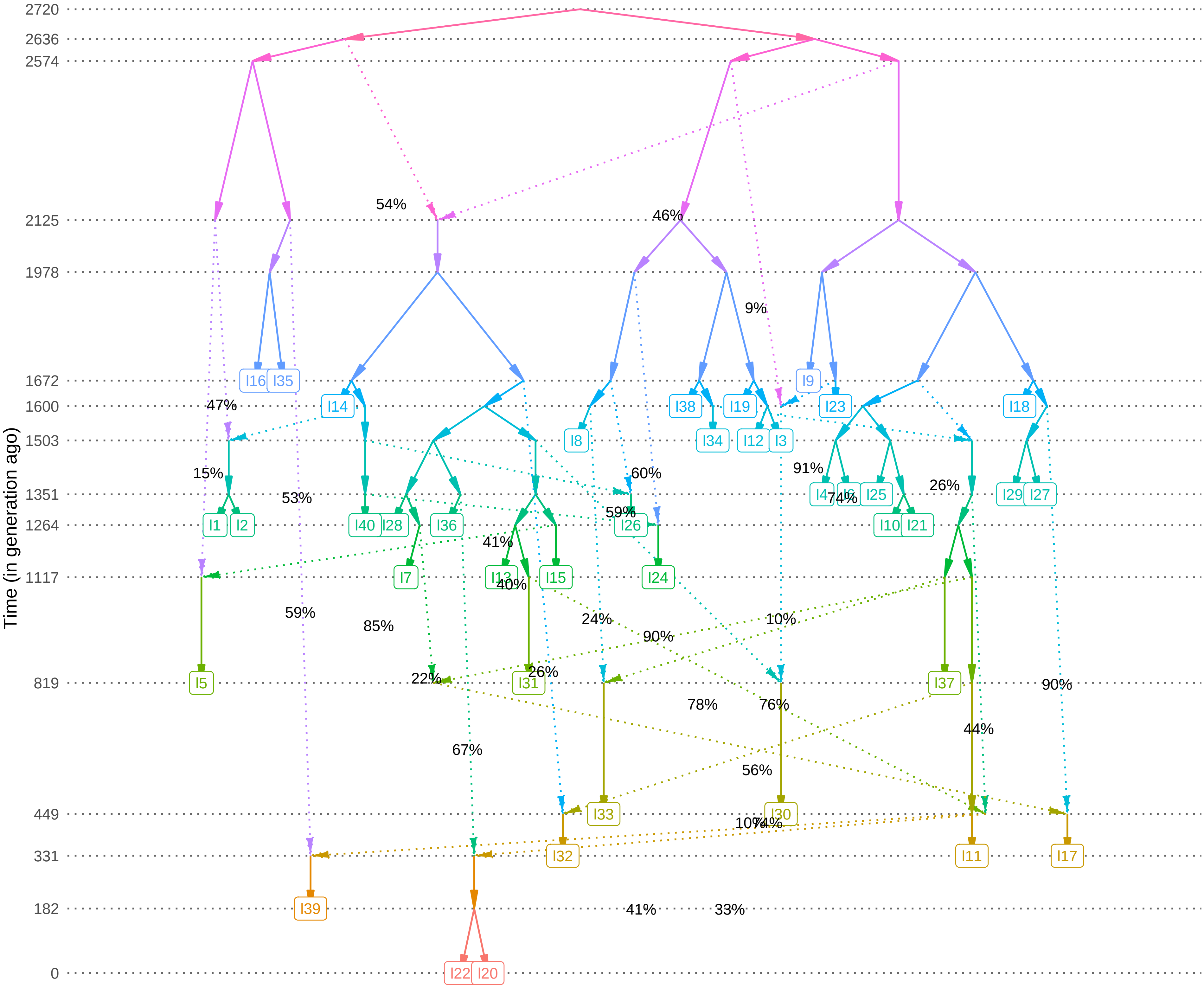
Time (in generation ago)
2720
2636
2574
2125
1978
1672
1600
1503
1351
1264
1117
819
449
331
182
0
I16
I35
I14
I9
I23
I38
I19
I18
I8
I34
I12
I3
I1
I2
I40
I28
I36
I26
I4
I6
I25
I29
I27
I10
I21
I7
I13
I15
I24
I5
I31
I37
I33
I30
I32
I11
I17
I39
I22
I20
54%
46%
9%
47%
15%
53%
60%
59%
91%
74%
26%
41%
40%
59%
85%
24%
90%
10%
22%
26%
78%
76%
90%
44%
67%
56%
10%
74%
41%
33%

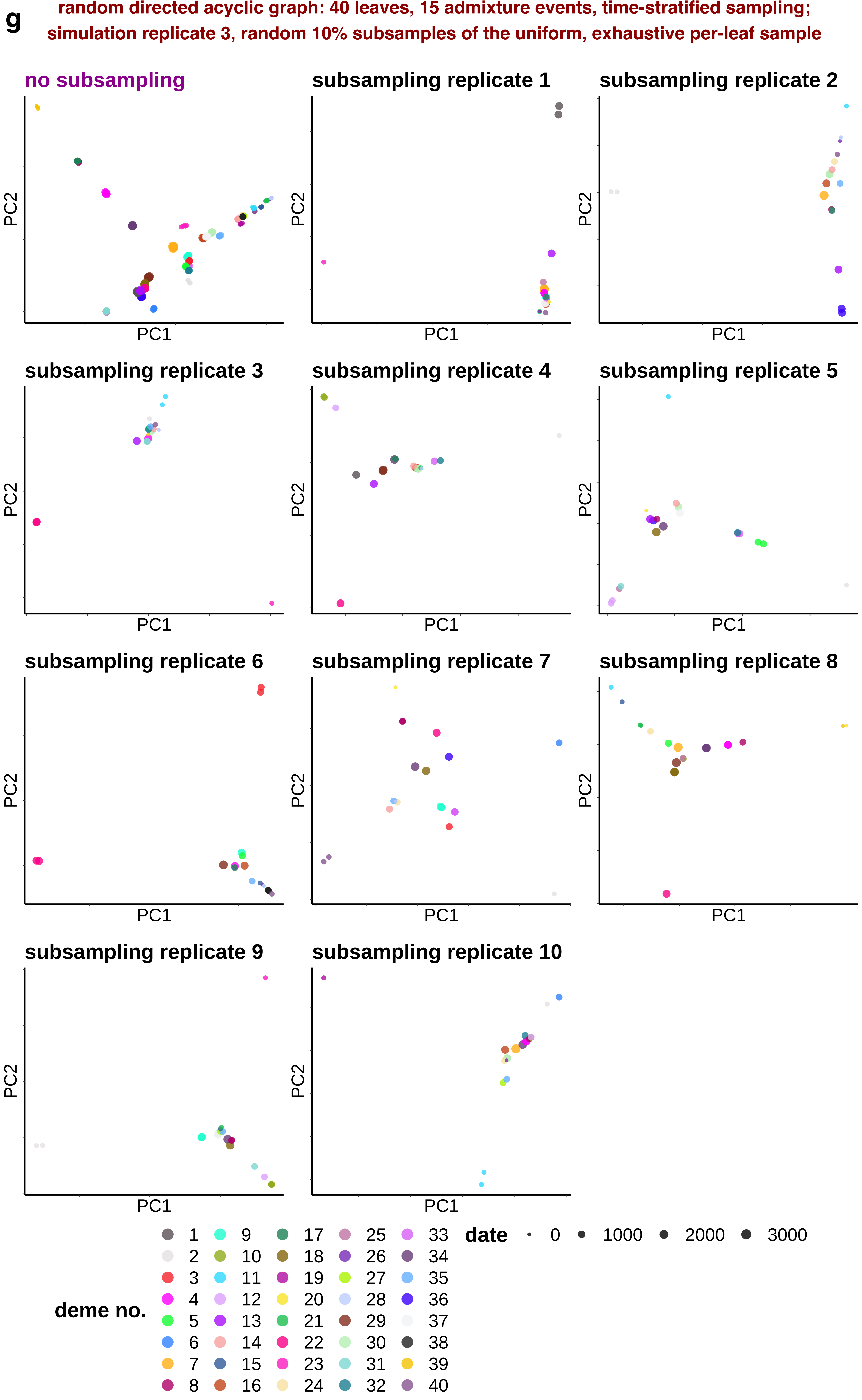
g
random directed acyclic graph: 40 leaves, 15 admixture events, time-stratified sampling;
simulation replicate 3, random 10% subsamples of the uniform, exhaustive per-leaf sample
no subsampling
subsampling replicate 1
subsampling replicate 2
subsampling replicate 3
subsampling replicate 4
subsampling replicate 5
subsampling replicate 6
subsampling replicate 7
subsampling replicate 8
subsampling replicate 9
subsampling replicate 10
PC1
PC2
deme no.
1 2 3 4 5 6 7 8 9 10 11 12 13 14 15 16 17 18 19 20 21 22 23 24 25 26 27 28 29 30 31 32 33 34 35 36 37 38 39 40
date
0 1000 2000 3000

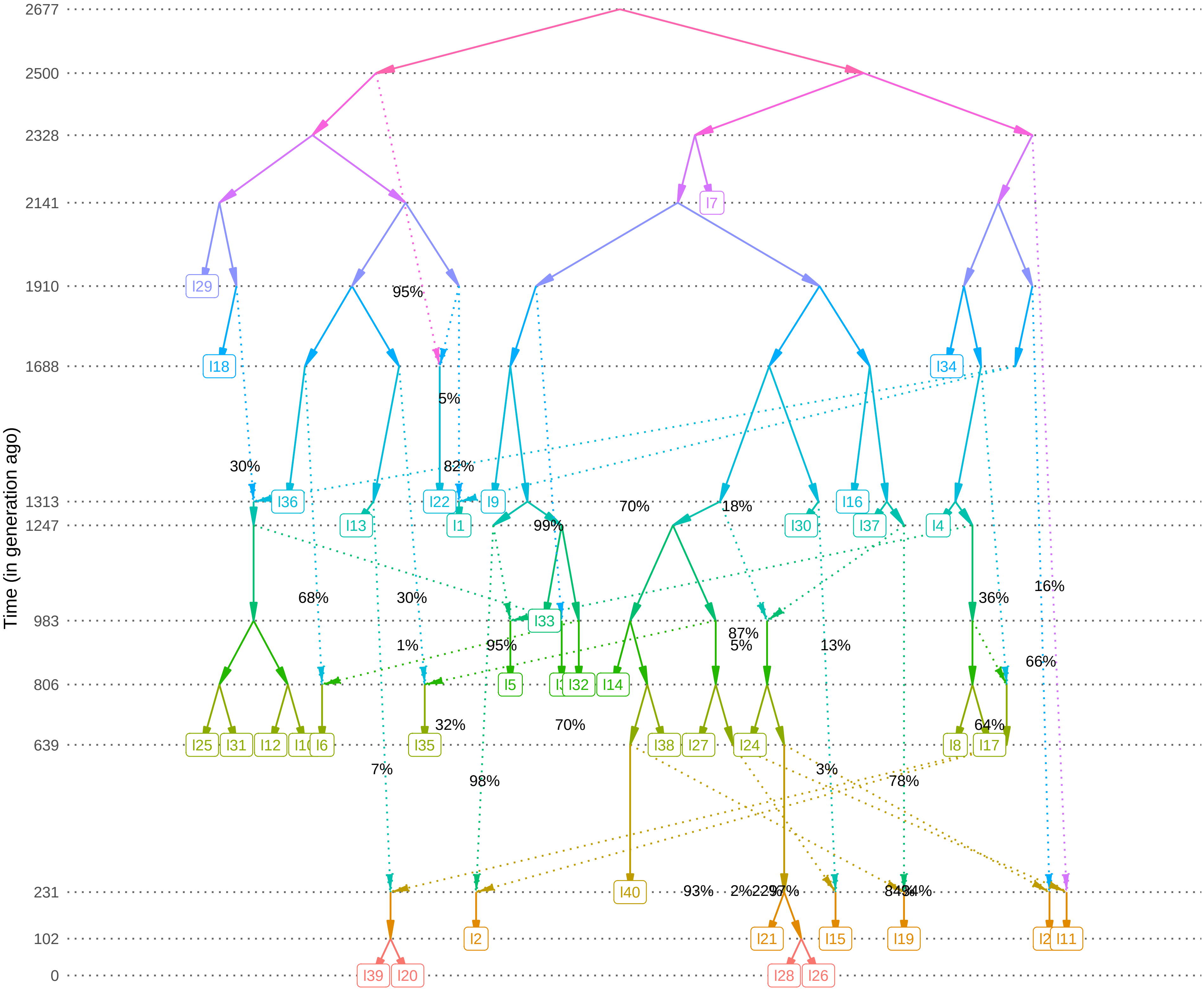

Time (in generation ago)
2677
2500
2328
2141
1910
1688
1313
1247
983
806
639
231
102
0
I7
I29
I18
I34
I36
I22
I9
I16
I13
I1
I30
I37
I4
I33
I5
I32
I14
I25
I31
I12
I6
I35
I38
I27
I24
I8
I17
I40
I2
I21
I15
I19
I11
I39
I20
I28
I26
95%
5%
30%
82%
70%
18%
99%
16%
68%
30%
36%
87%
1%
95%
5%
13%
66%
32%
70%
64%
7%
98%
3%
78%
93%

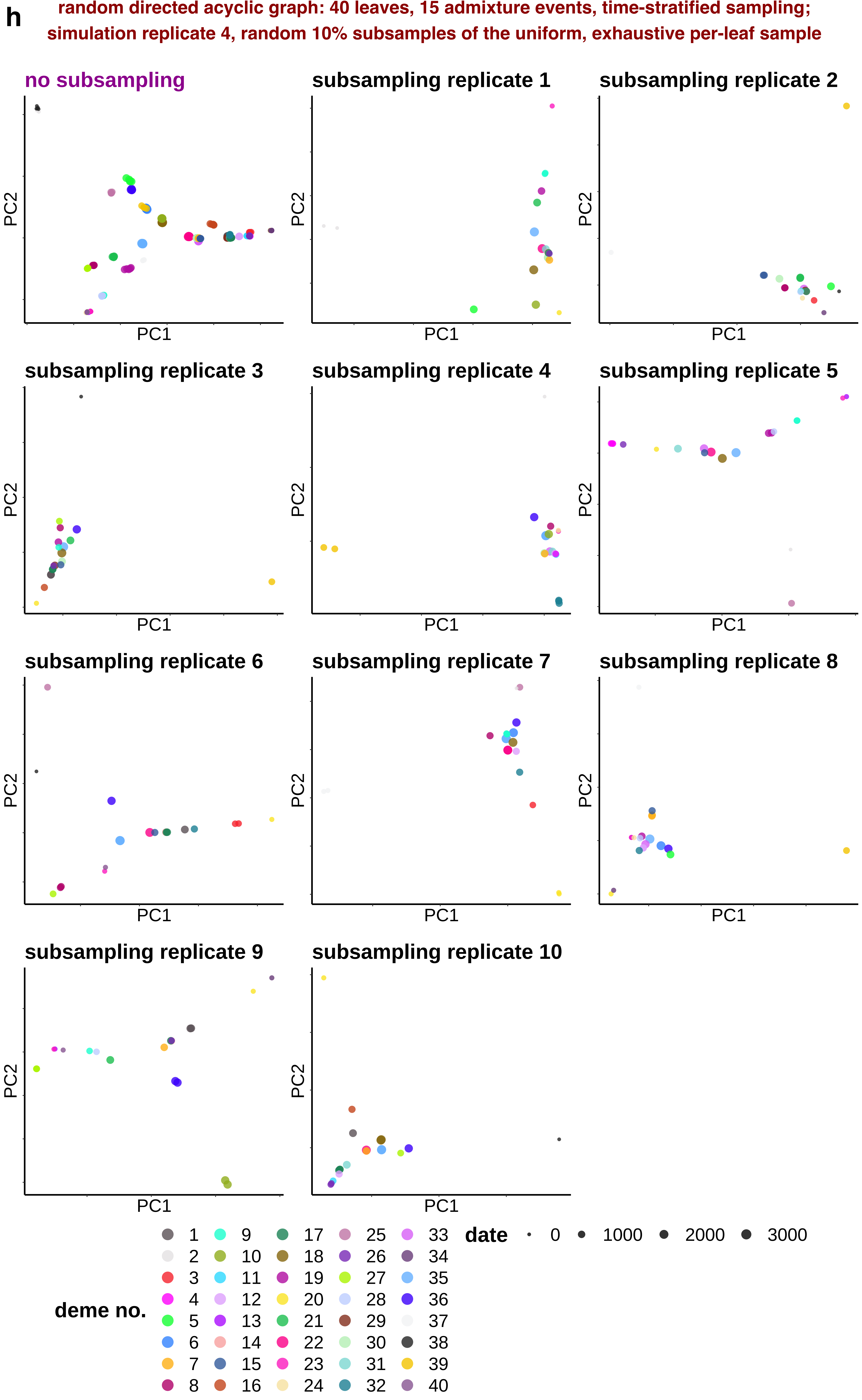
h
random directed acyclic graph: 40 leaves, 15 admixture events, time-stratified sampling;
simulation replicate 4, random 10% subsamples of the uniform, exhaustive per-leaf sample
no subsampling
subsampling replicate 1
subsampling replicate 2
subsampling replicate 3
subsampling replicate 4
subsampling replicate 5
subsampling replicate 6
subsampling replicate 7
subsampling replicate 8
subsampling replicate 9
subsampling replicate 10
PC1
PC2
deme no.
1 2 3 4 5 6 7 8 9 10 11 12 13 14 15 16 17 18 19 20 21 22 23 24 25 26 27 28 29 30 31 32 33 34 35 36 37 38 39 40
date
0 1000 2000 3000

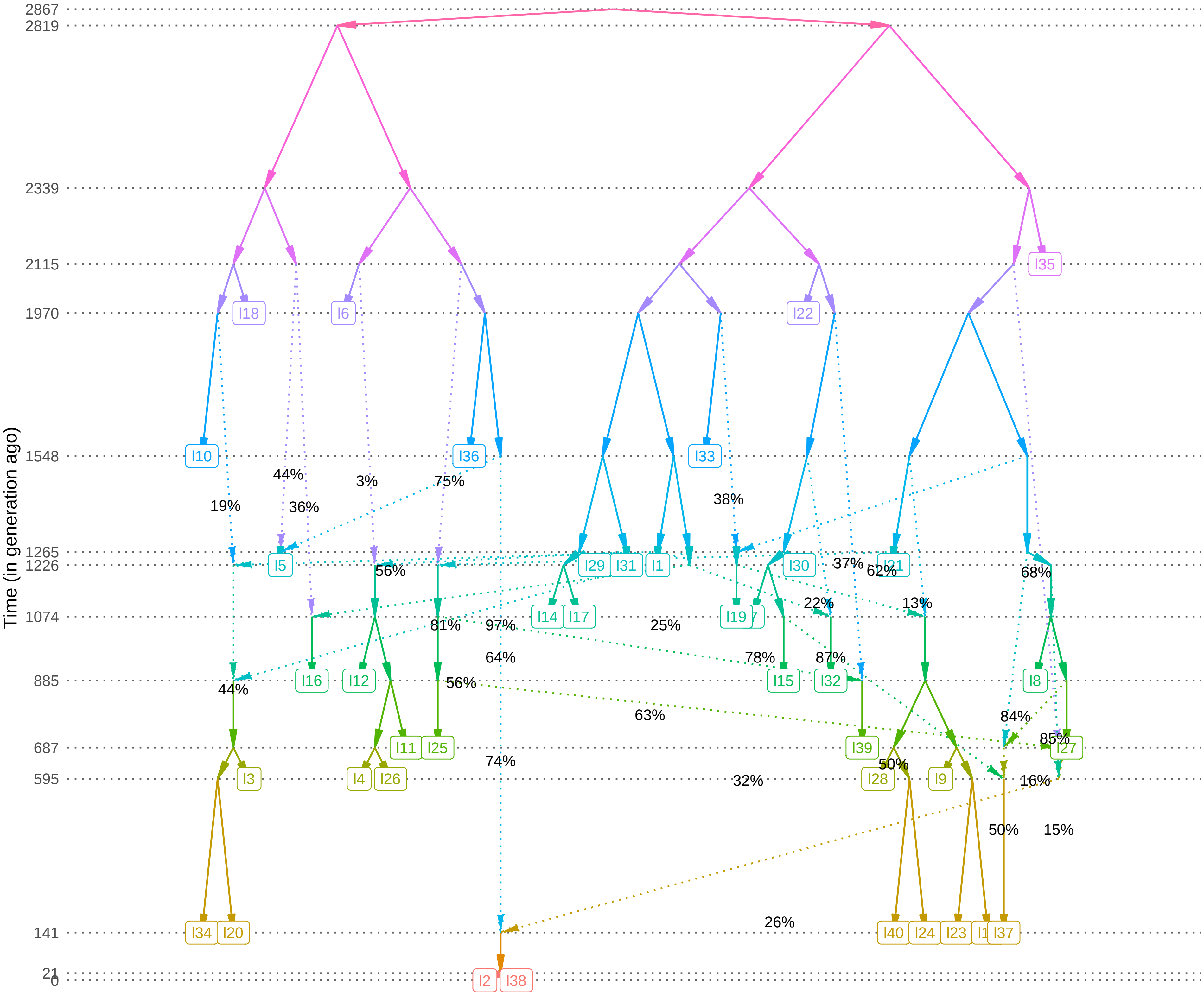

Time (in generation ago)
2867
2819
2339
2115
1970
1548
1265
1226
1074
885
687
595
141
21
0
I18
I6
I22
I35
I10
I36
I33
I5
I29
I31
I1
I30
I21
I14
I17
I19
I16
I12
I15
I32
I8
I11
I25
I39
I27
I4
I26
I3
I28
I9
I34
I20
I40
I24
I23
I37
I2
I38
19%
44%
36%
3%
75%
38%
56%
37%
62%
68%
81%
97%
25%
22%
13%
64%
78%
87%
44%
56%
63%
84%
85%
74%
50%
32%
16%
50%
15%
26%

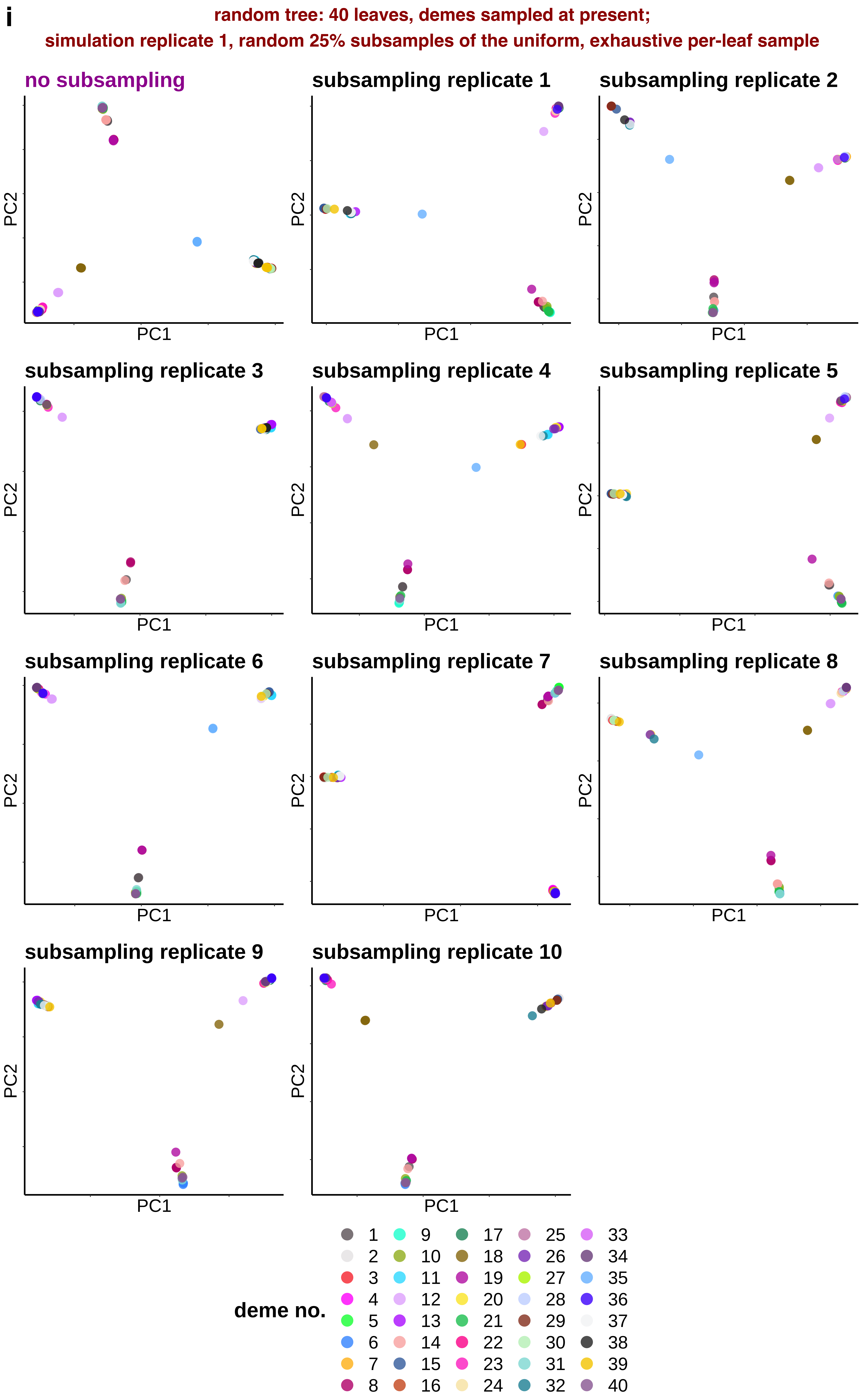
i
random tree: 40 leaves, demes sampled at present;
simulation replicate 1, random 25% subsamples of the uniform, exhaustive per-leaf sample
no subsampling
subsampling replicate 1
subsampling replicate 2
subsampling replicate 3
subsampling replicate 4
subsampling replicate 5
subsampling replicate 6
subsampling replicate 7
subsampling replicate 8
subsampling replicate 9
subsampling replicate 10
PC1
PC2
deme no.
1 2 3 4 5 6 7 8 9 10 11 12 13 14 15 16 17 18 19 20 21 22 23 24 25 26 27 28 29 30 31 32 33 34 35 36 37 38 39 40

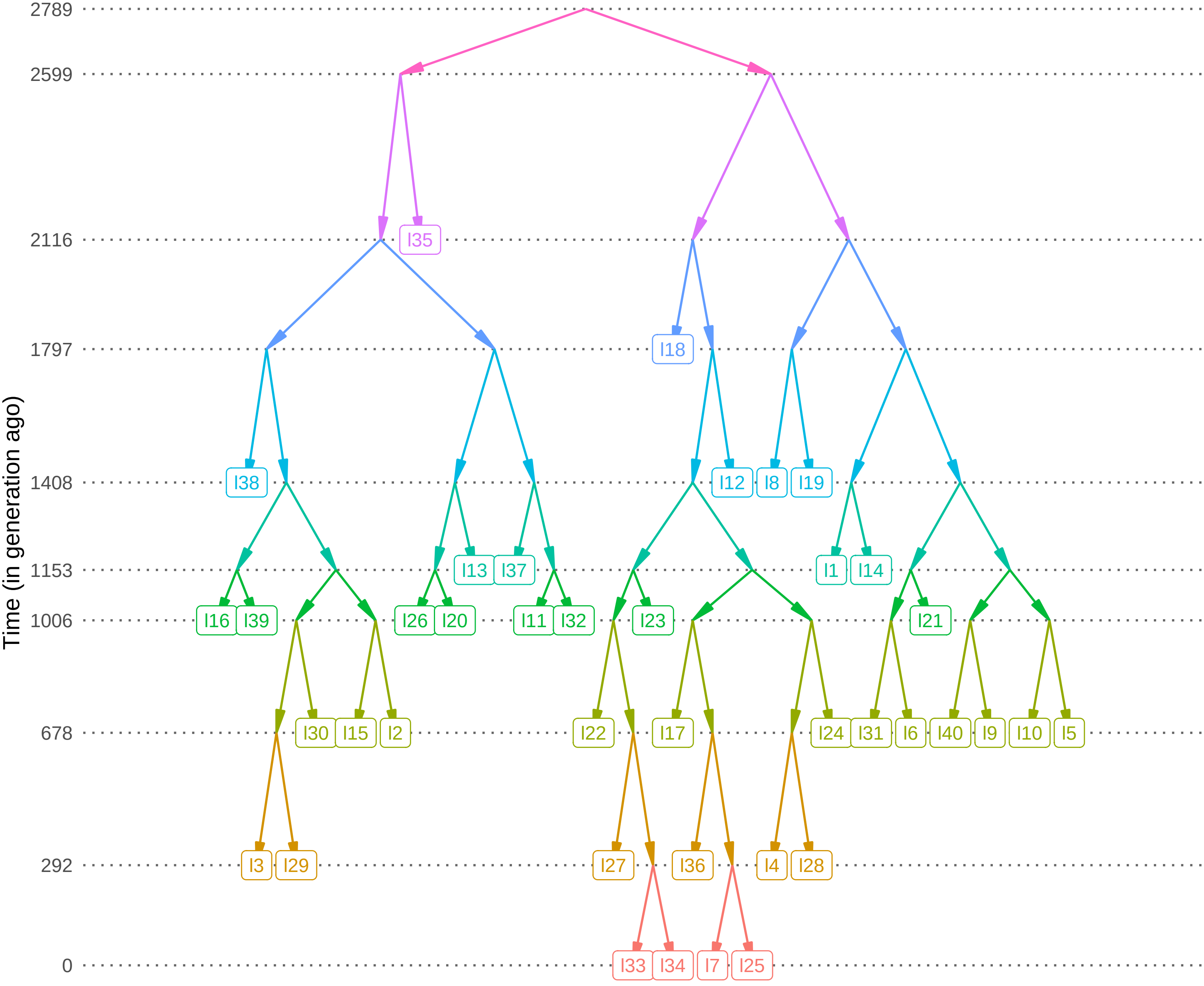

Time (in generation ago)
2789
2599
2116
1797
1408
1153
1006
678
292
0
I35
I18
I38
I12
I8
I19
I13
I37
I1
I14
I16
I39
I26
I20
I11
I32
I23
I21
I30
I15
I2
I22
I17
I24
I31
I6
I40
I9
I10
I5
I3
I29
I27
I36
I4
I28
I33
I34
I7
I25

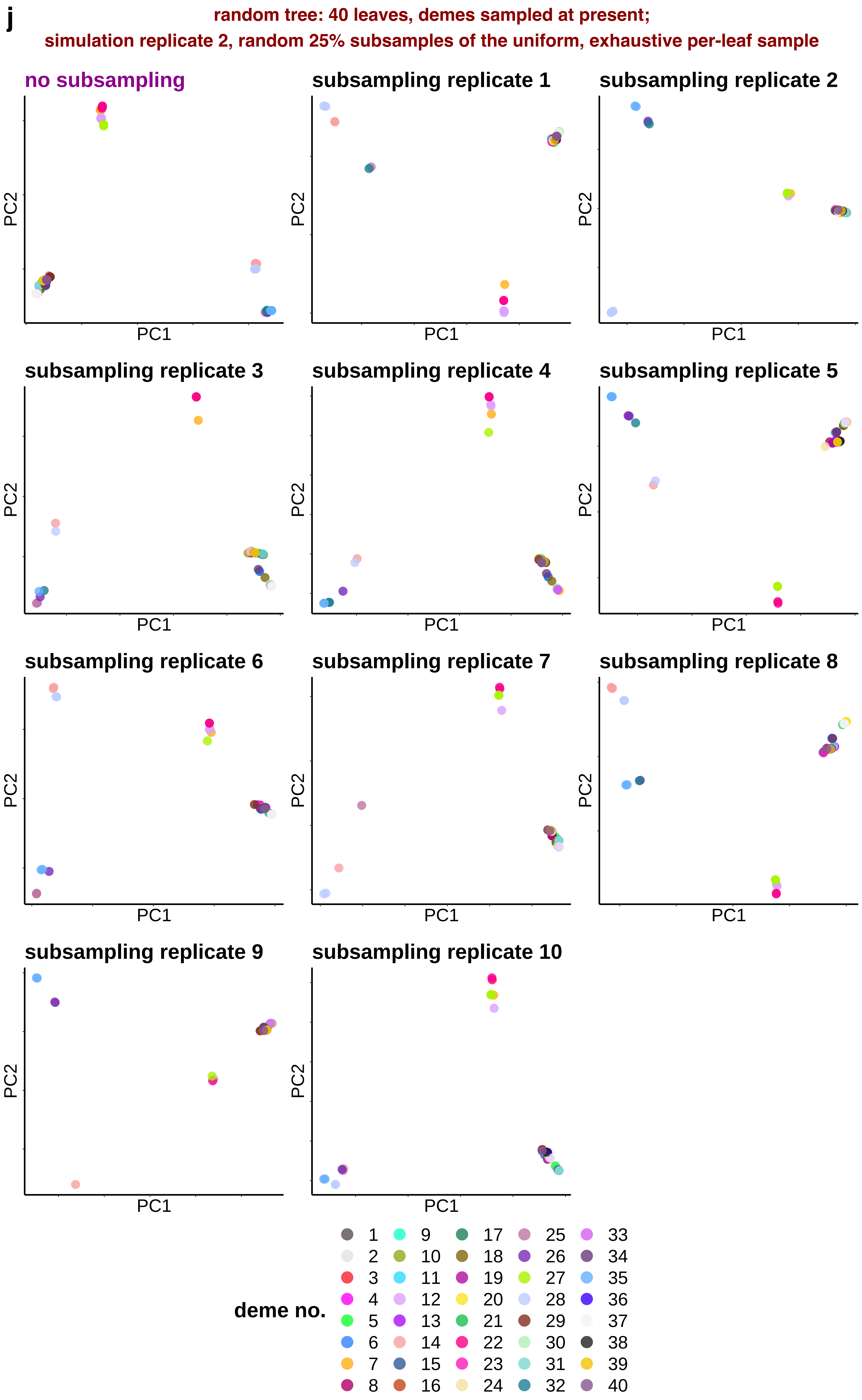
j
random tree: 40 leaves, demes sampled at present;
simulation replicate 2, random 25% subsamples of the uniform, exhaustive per-leaf sample
no subsampling
subsampling replicate 1
subsampling replicate 2
subsampling replicate 3
subsampling replicate 4
subsampling replicate 5
subsampling replicate 6
subsampling replicate 7
subsampling replicate 8
subsampling replicate 9
subsampling replicate 10
PC1
PC2
deme no.
1 2 3 4 5 6 7 8 9 10 11 12 13 14 15 16 17 18 19 20 21 22 23 24 25 26 27 28 29 30 31 32 33 34 35 36 37 38 39 40

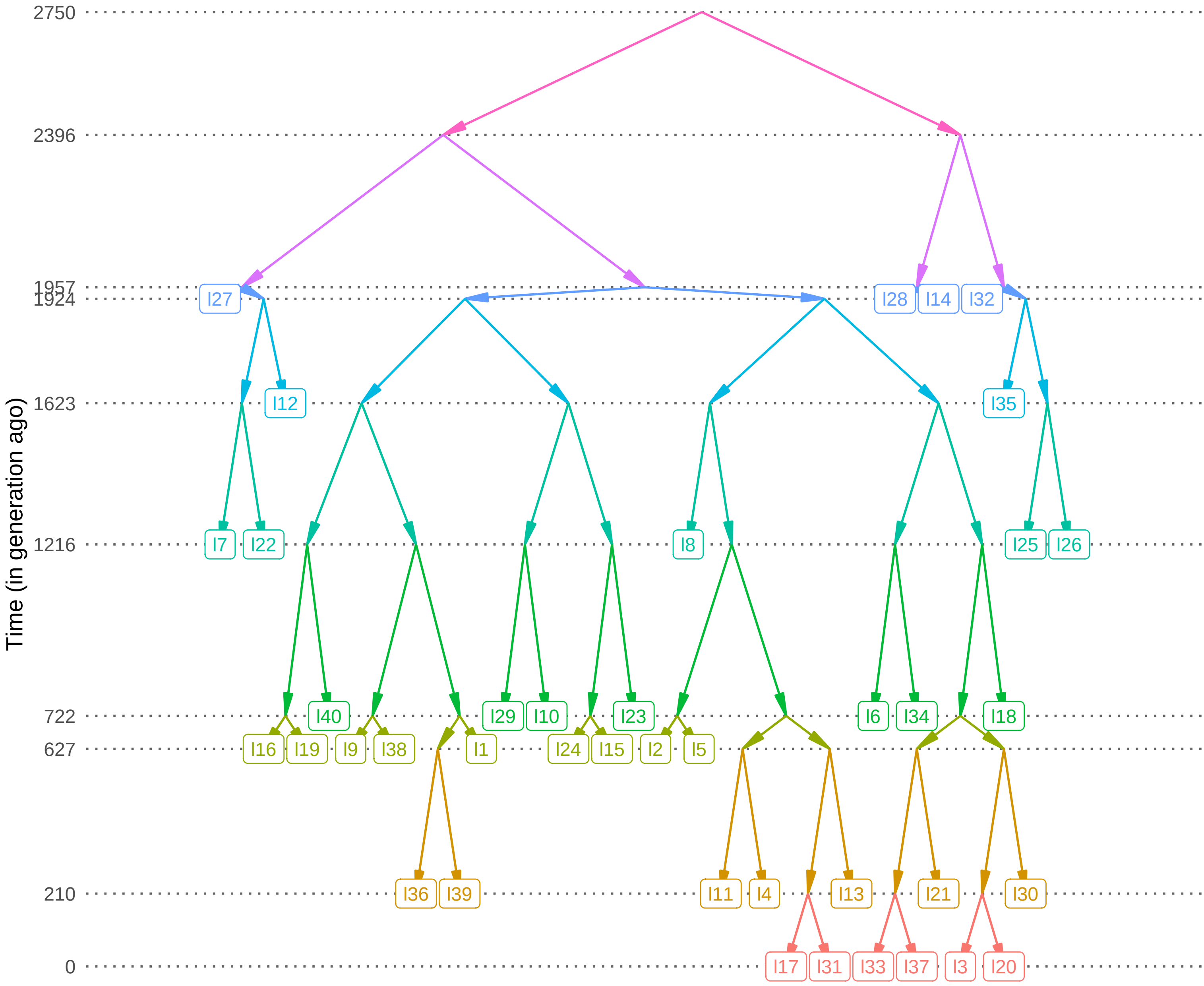
2750
2396
1957
1924
1623
1216
722
627
210
0
Time (in generation ago)
I27
I28
I14
I32
I12
I35
I7
I22
I8
I25
I26
I40
I29
I10
I23
I6
I34
I18
I16
I19
I9
I38
I1
I24
I15
I2
I5
I36
I39
I11
I4
I13
I21
I30
I17
I31
I33
I37
I3
I20

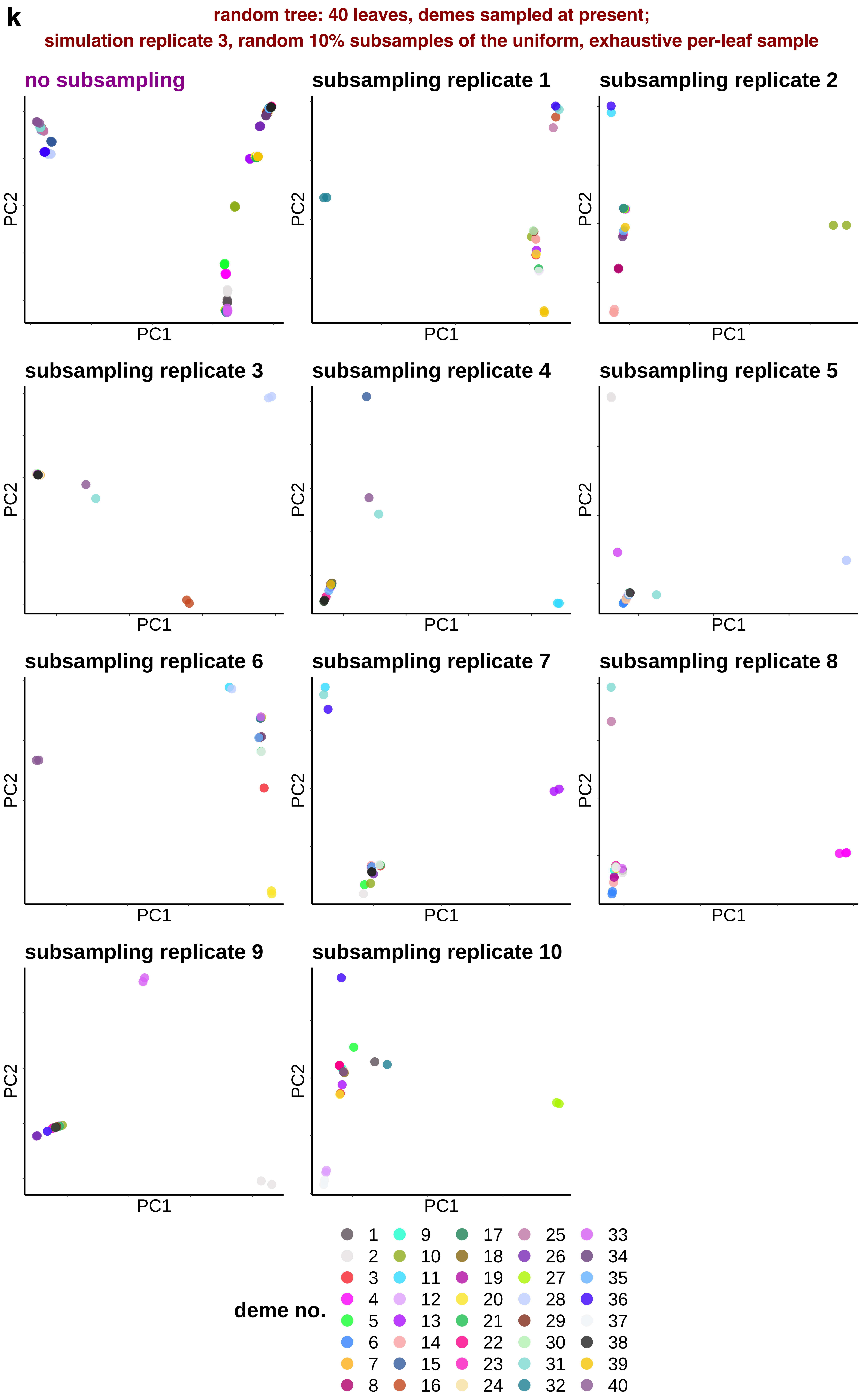

k
random tree: 40 leaves, demes sampled at present;
simulation replicate 3, random 10% subsamples of the uniform, exhaustive per-leaf sample
no subsampling
subsampling replicate 1
subsampling replicate 2
subsampling replicate 3
subsampling replicate 4
subsampling replicate 5
subsampling replicate 6
subsampling replicate 7
subsampling replicate 8
subsampling replicate 9
subsampling replicate 10
PC1
PC2
deme no.
1 2 3 4 5 6 7 8 9 10 11 12 13 14 15 16 17 18 19 20 21 22 23 24 25 26 27 28 29 30 31 32 33 34 35 36 37 38 39 40

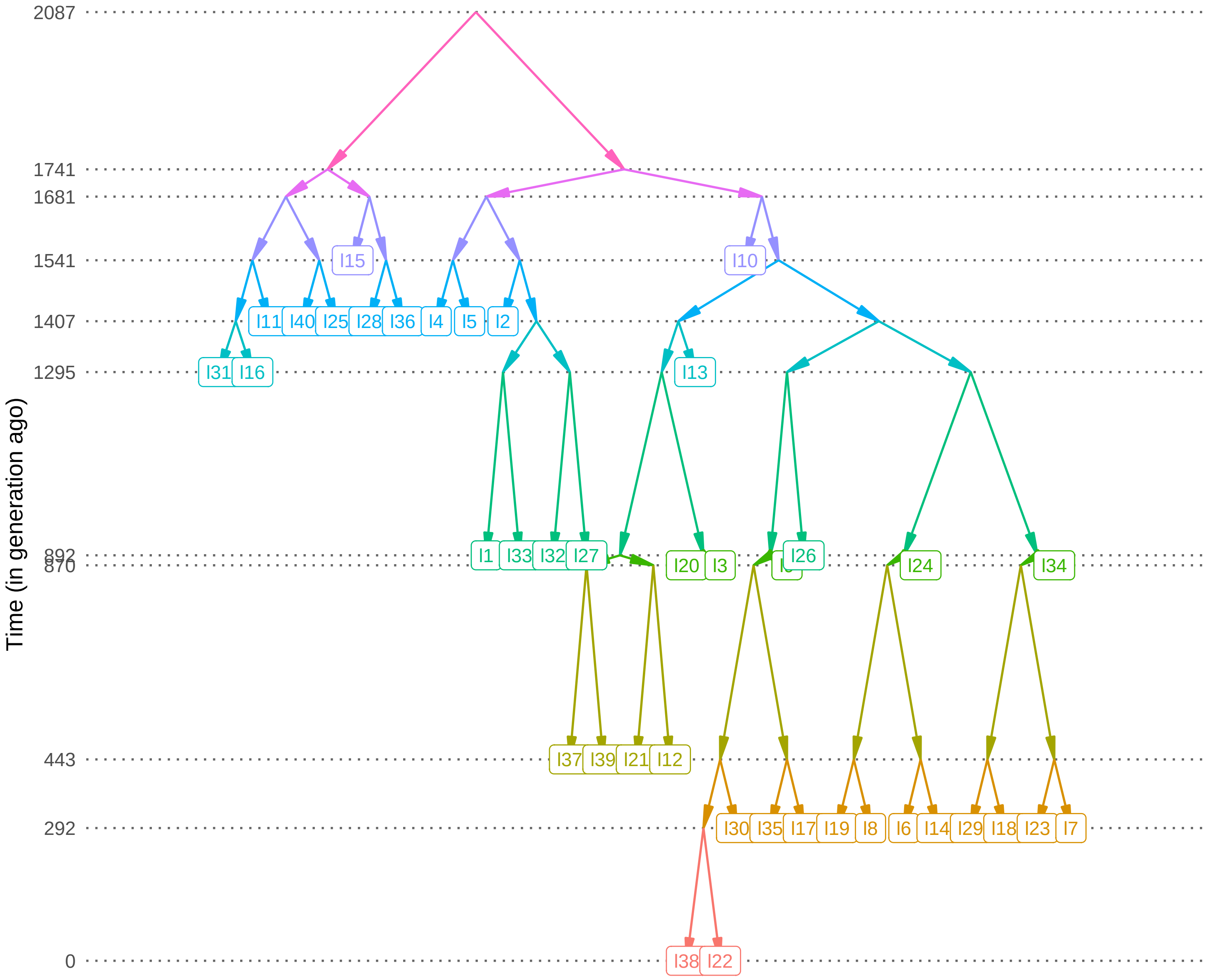

Time (in generation ago)
2087
1741
1681
1541
1407
1295
892
870
443
292
0
I15
I10
I11
I40
I25
I28
I36
I4
I5
I2
I31
I16
I13
I1
I33
I32
I27
I20
I3
I26
I24
I34
I37
I39
I21
I12
I30
I35
I17
I19
I8
I6
I14
I29
I18
I23
I7
I38
I22

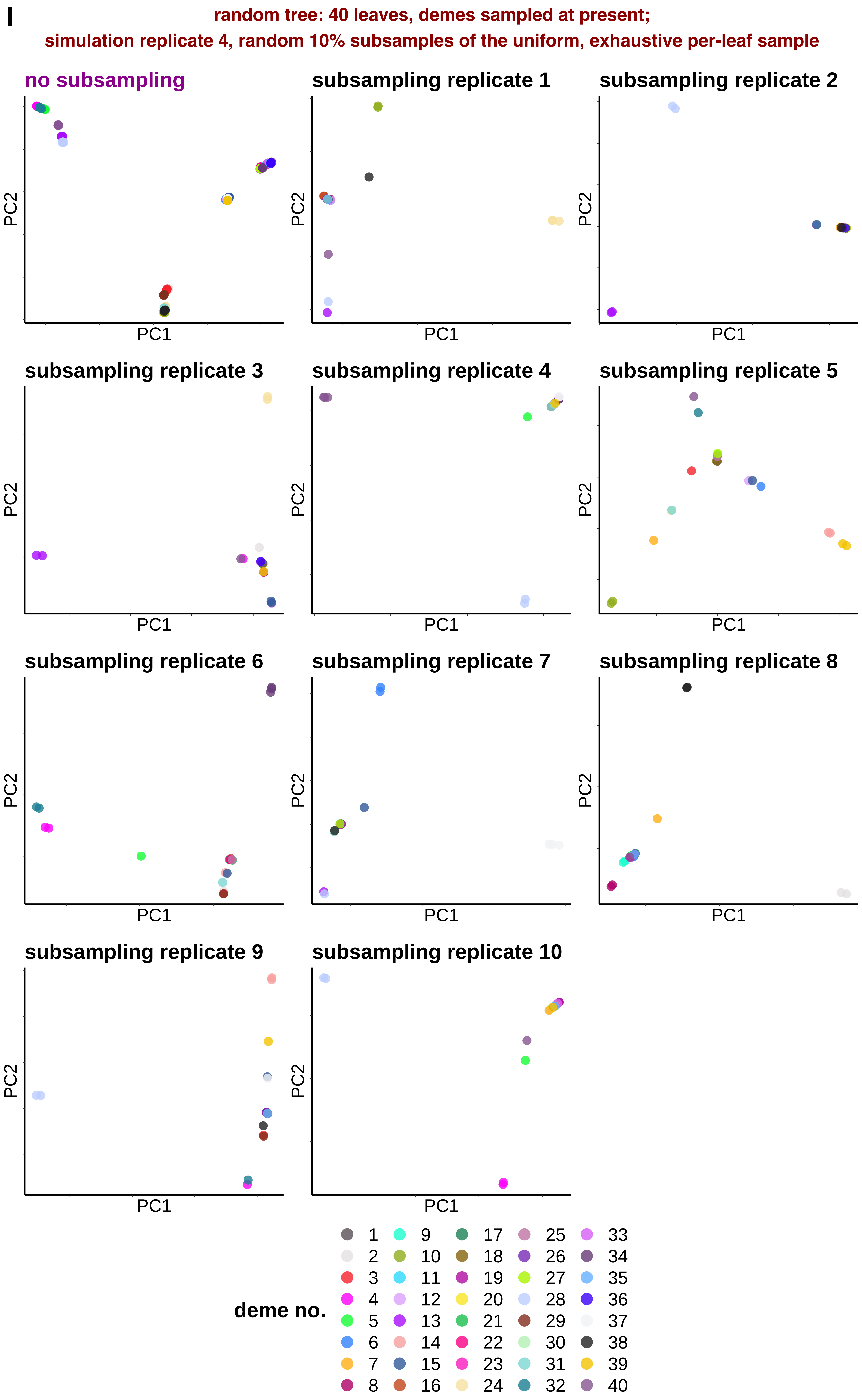
I
random tree: 40 leaves, demes sampled at present;
simulation replicate 4, random 10% subsamples of the uniform, exhaustive per-leaf sample
no subsampling
subsampling replicate 1
subsampling replicate 2
subsampling replicate 3
subsampling replicate 4
subsampling replicate 5
subsampling replicate 6
subsampling replicate 7
subsampling replicate 8
subsampling replicate 9
subsampling replicate 10
PC1
PC2
deme no.
1 2 3 4 5 6 7 8 9 10 11 12 13 14 15 16 17 18 19 20 21 22 23 24 25 26 27 28 29 30 31 32 33 34 35 36 37 38 39 40

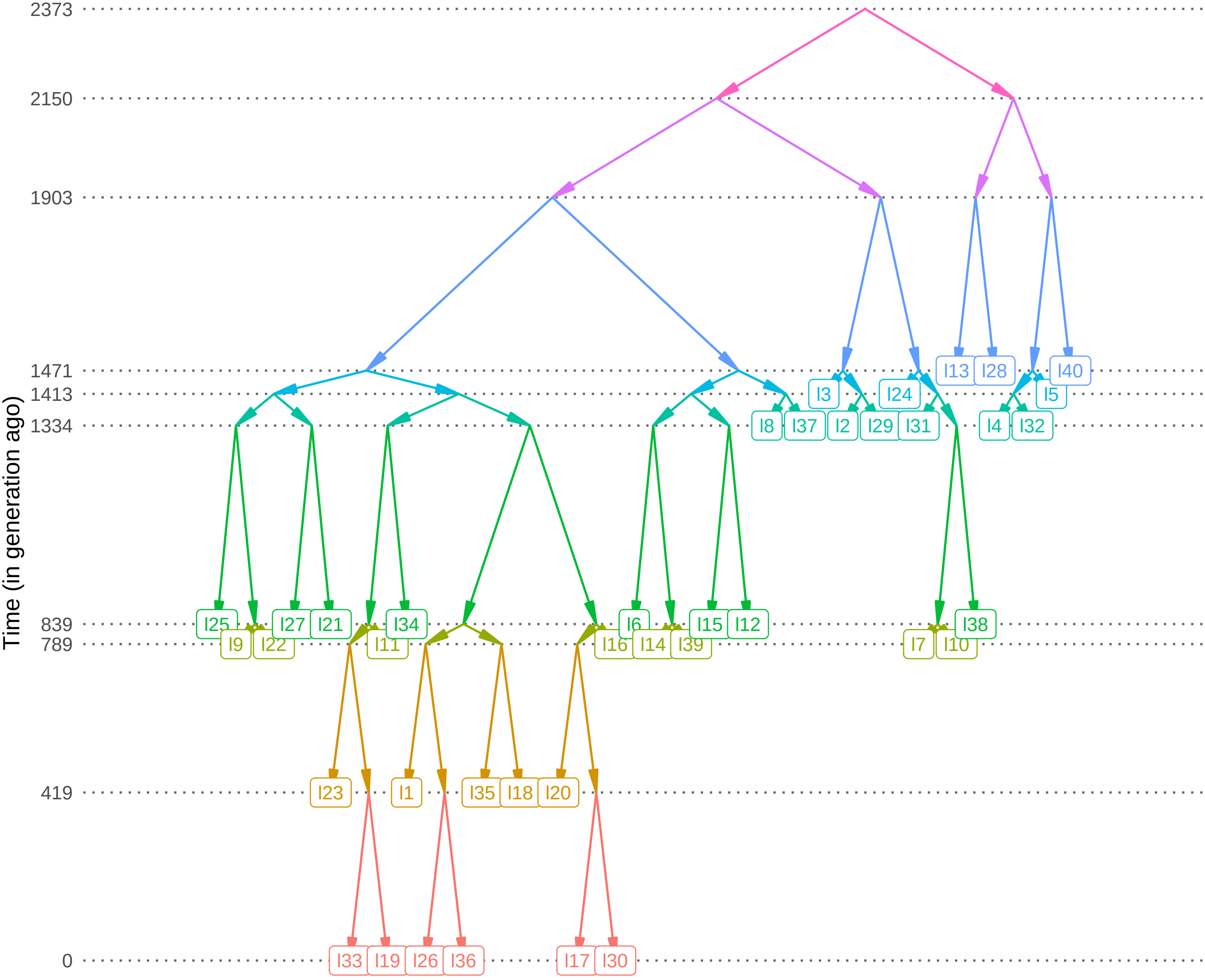

Time (in generation ago)
2373
2150
1903
1471
1413
1334
839
789
419
0
I13
I28
I40
I3
I24
I5
I8
I37
I2
I29
I31
I4
I32
I25
I27
I21
I34
I6
I15
I12
I38
I9
I22
I11
I16
I14
I39
I7
I10
I23
I1
I35
I18
I20
I33
I19
I26
I36
I17
I30

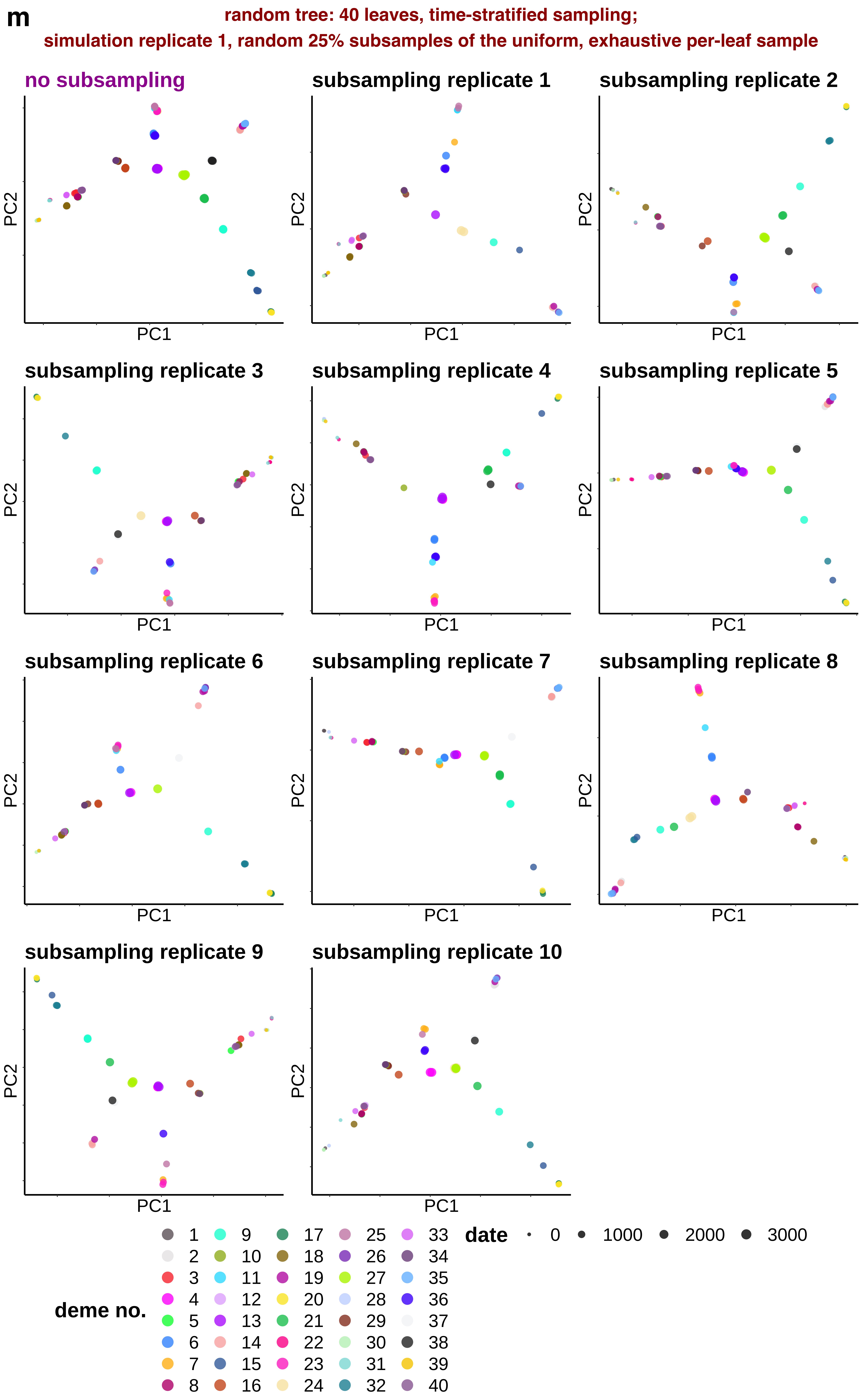
m
random tree: 40 leaves, time-stratified sampling;
simulation replicate 1, random 25% subsamples of the uniform, exhaustive per-leaf sample
no subsampling
subsampling replicate 1
subsampling replicate 2
subsampling replicate 3
subsampling replicate 4
subsampling replicate 5
subsampling replicate 6
subsampling replicate 7
subsampling replicate 8
subsampling replicate 9
subsampling replicate 10
PC1
PC2
deme no.
1 2 3 4 5 6 7 8 9 10 11 12 13 14 15 16 17 18 19 20 21 22 23 24 25 26 27 28 29 30 31 32 33 34 35 36 37 38 39 40
date
0 1000 2000 3000

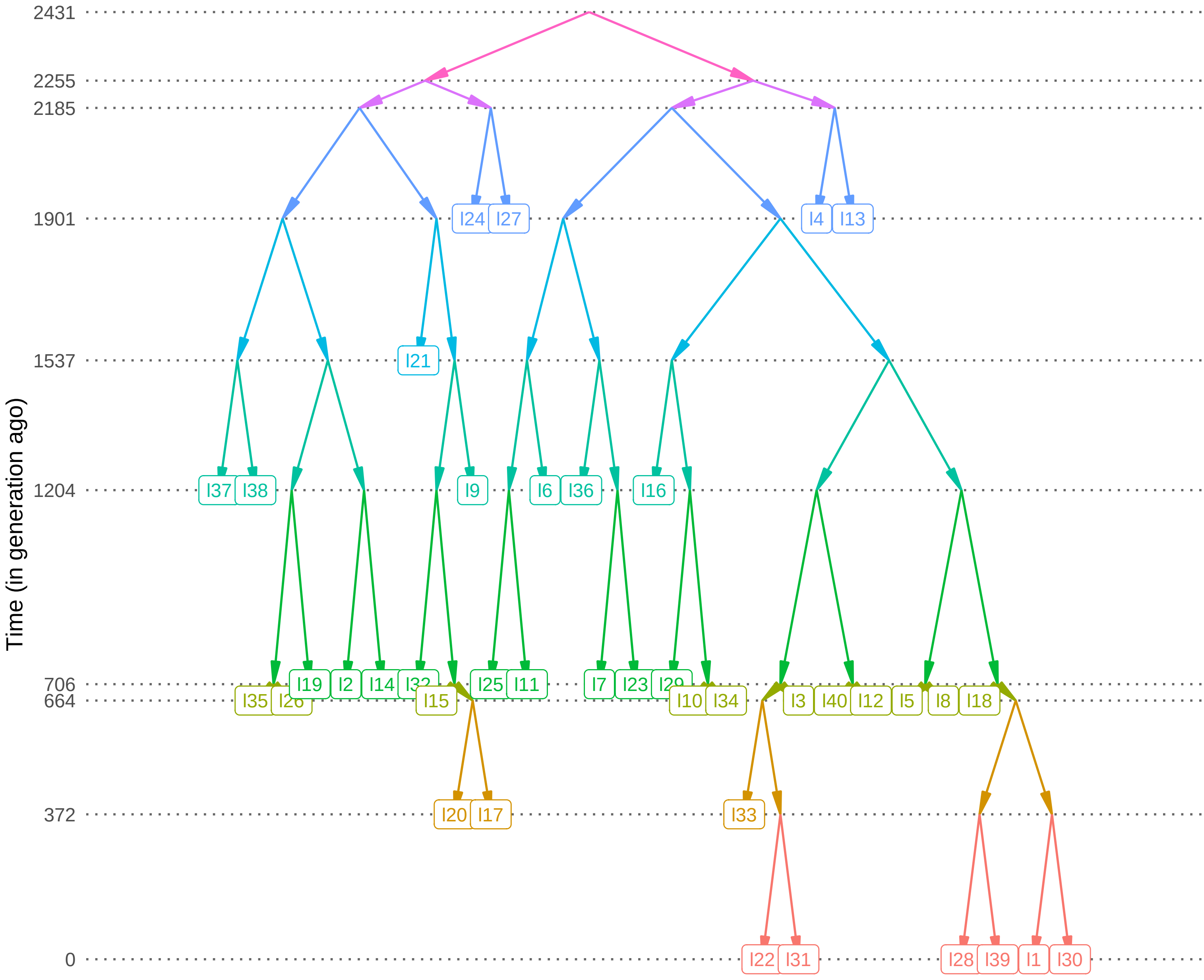
Time (in generation ago)
2431
2255
2185
1901
1537
1204
706
664
372
0
I24
I27
I4
I13
I21
I37
I38
I9
I6
I36
I16
I19
I2
I14
I25
I11
I7
I23
I35
I15
I10
I34
I3
I40
I12
I5
I8
I18
I20
I17
I33
I22
I31
I28
I39
I1
I30

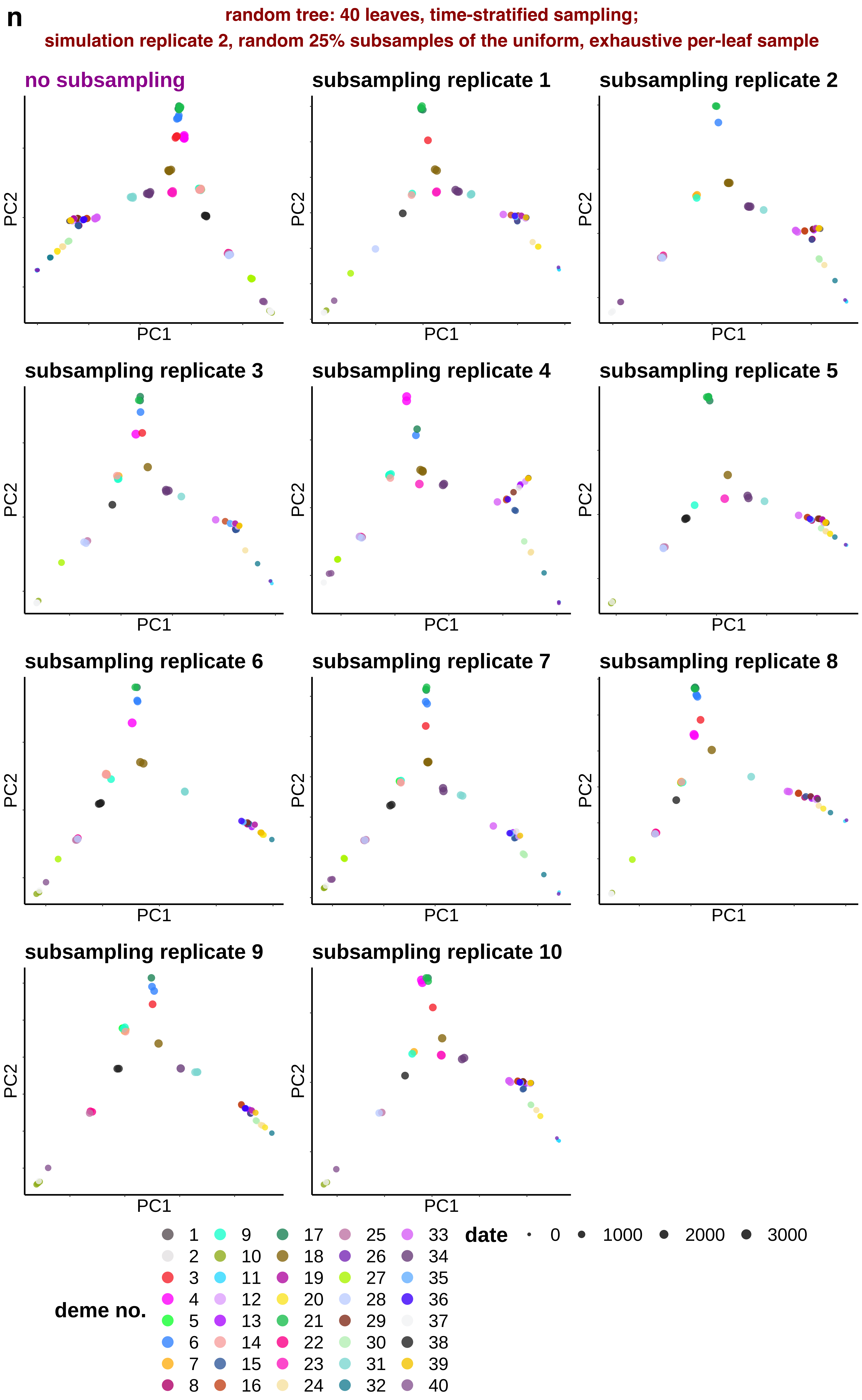
n
random tree: 40 leaves, time-stratified sampling;
simulation replicate 2, random 25% subsamples of the uniform, exhaustive per-leaf sample
no subsampling
subsampling replicate 1
subsampling replicate 2
subsampling replicate 3
subsampling replicate 4
subsampling replicate 5
subsampling replicate 6
subsampling replicate 7
subsampling replicate 8
subsampling replicate 9
subsampling replicate 10
PC1
PC2
deme no.
1 2 3 4 5 6 7 8 9 10 11 12 13 14 15 16 17 18 19 20 21 22 23 24 25 26 27 28 29 30 31 32 33 34 35 36 37 38 39 40
date
0
1000
2000
3000

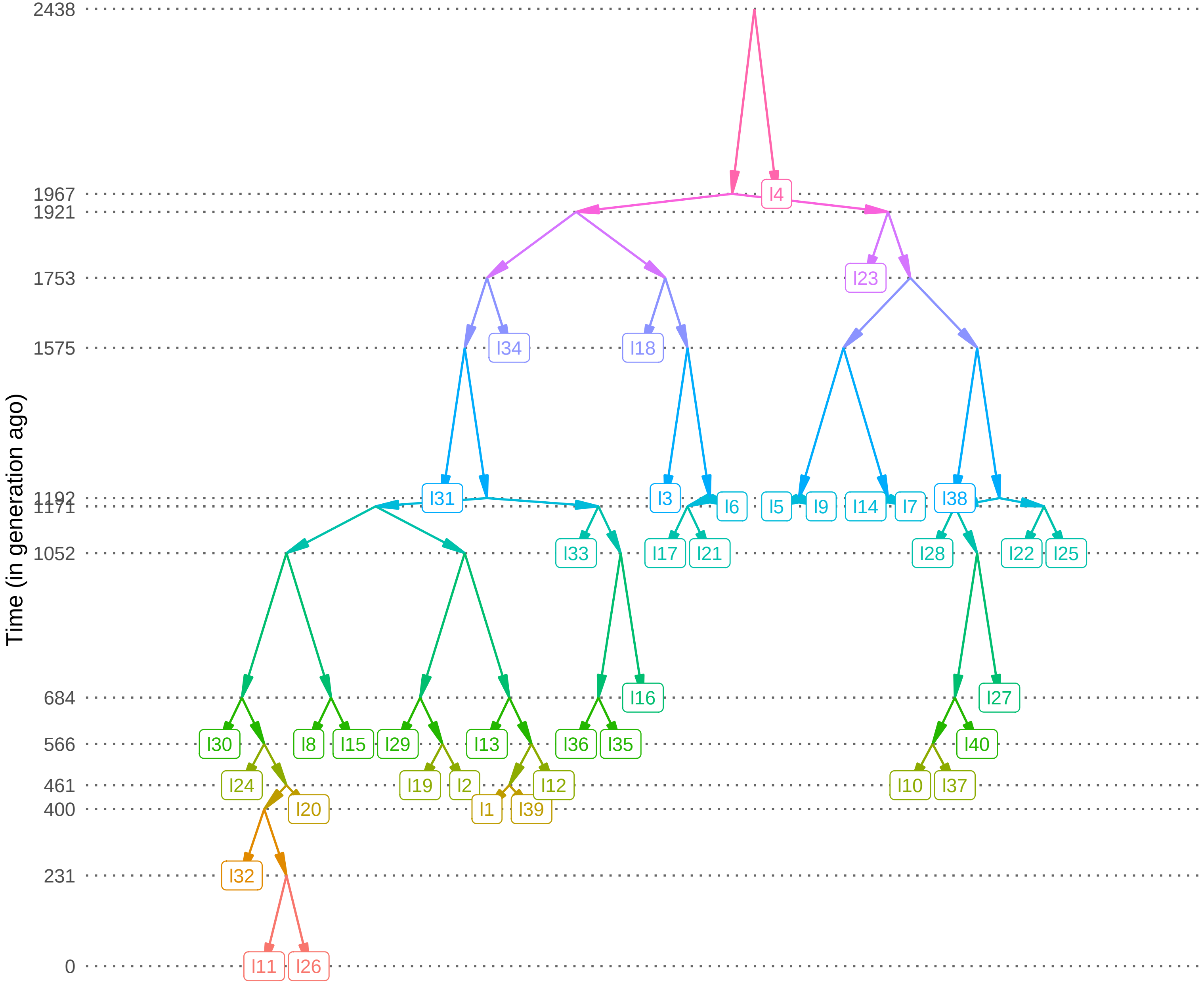
Time (in generation ago)
2438
1967
1921
1753
1575
1192
1171
1052
684
566
461
400
231
0
I4
I23
I34
I18
I31
I3
I6
I5
I9
I14
I7
I38
I33
I17
I21
I28
I22
I25
I16
I27
I30
I8
I15
I29
I13
I36
I35
I40
I24
I19
I2
I12
I10
I37
I20
I1
I39
I32
I11
I26

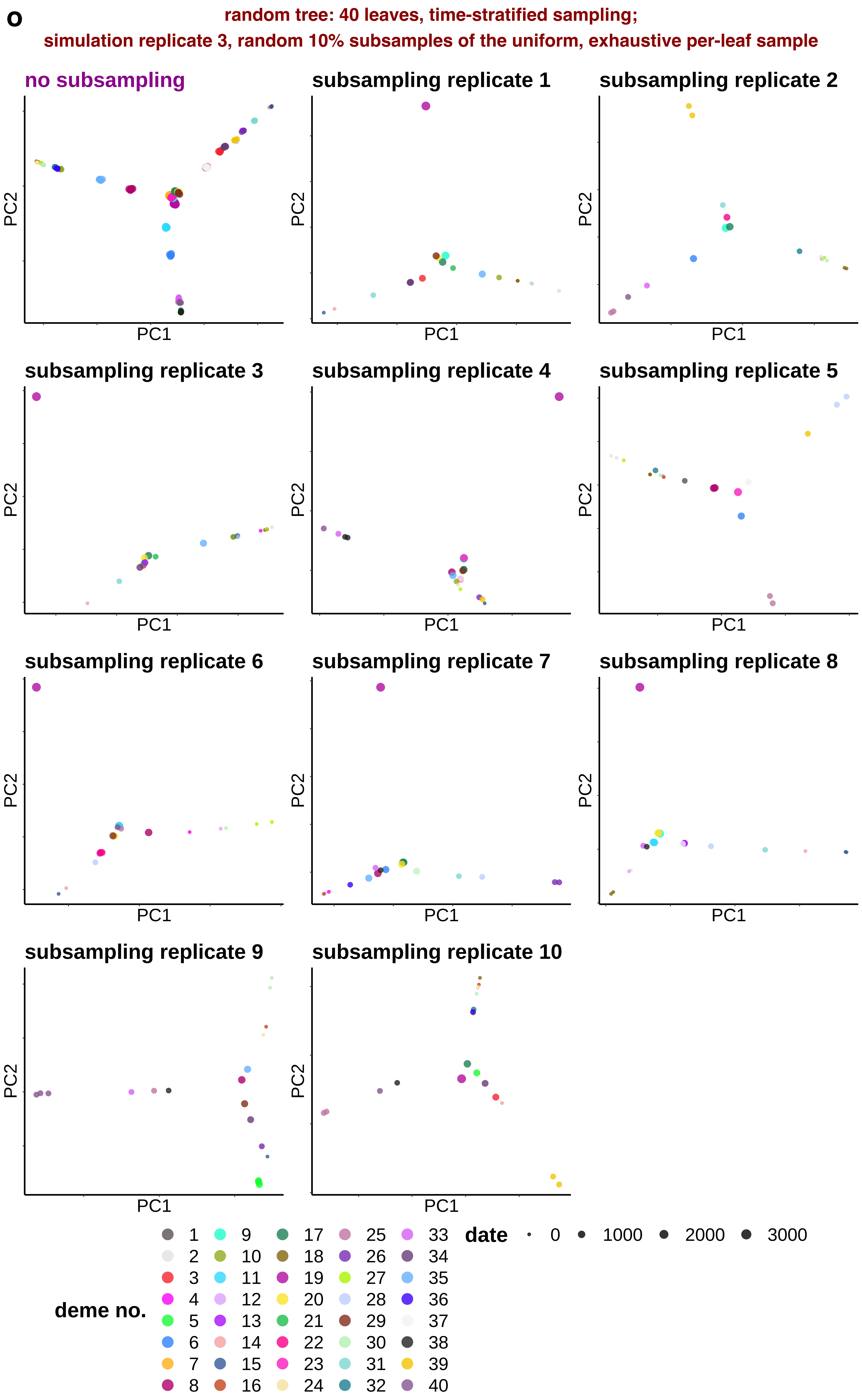
o
random tree: 40 leaves, time-stratified sampling;
simulation replicate 3, random 10% subsamples of the uniform, exhaustive per-leaf sample
no subsampling
subsampling replicate 1
subsampling replicate 2
subsampling replicate 3
subsampling replicate 4
subsampling replicate 5
subsampling replicate 6
subsampling replicate 7
subsampling replicate 8
subsampling replicate 9
subsampling replicate 10
PC1
PC2
deme no.
1 2 3 4 5 6 7 8 9 10 11 12 13 14 15 16 17 18 19 20 21 22 23 24 25 26 27 28 29 30 31 32 33 34 35 36 37 38 39 40
date
0
1000
2000
3000

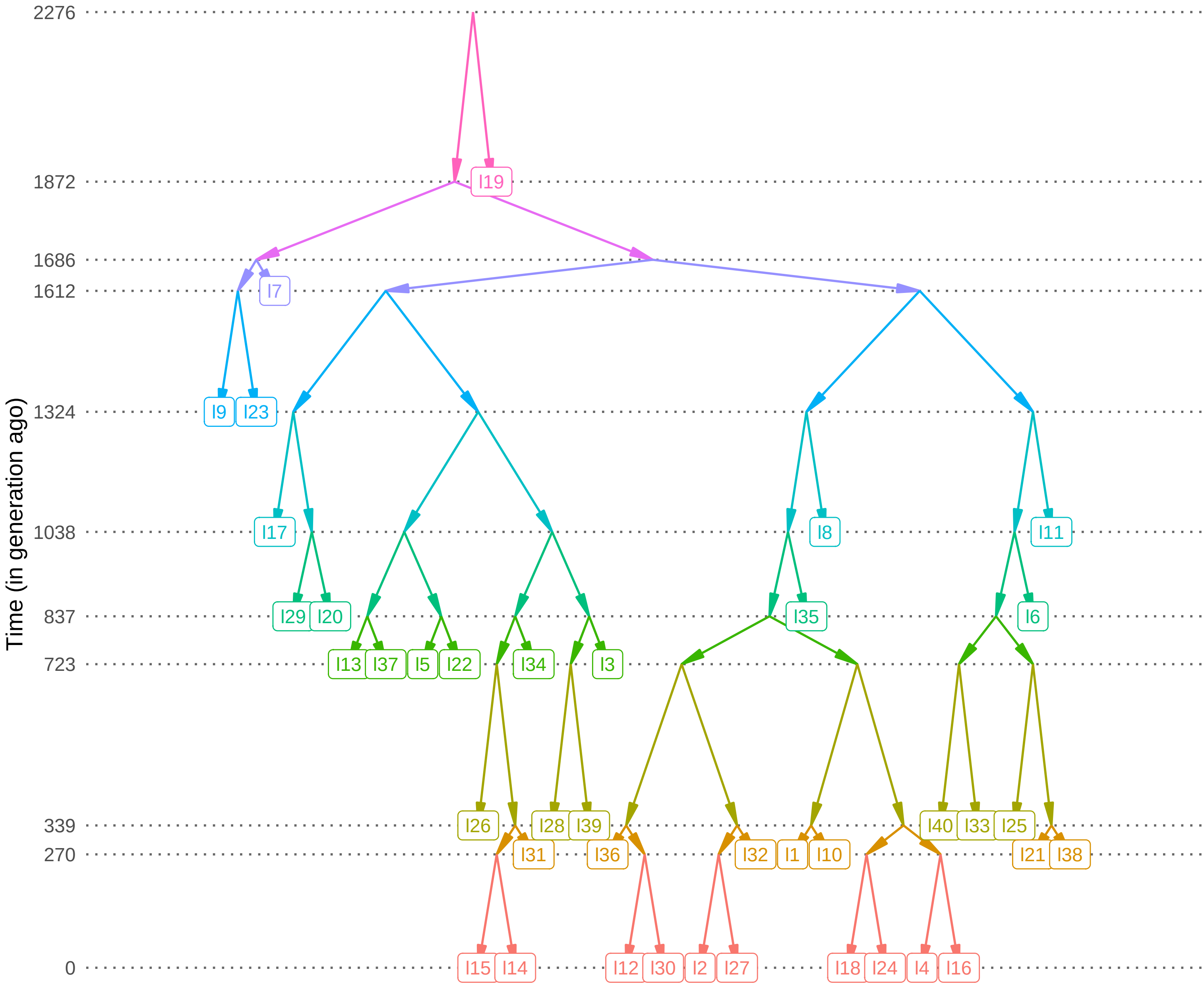

Time (in generation ago)
2276
1872
1686
1612
1324
1038
837
723
339
270
0
I19
I7
I9
I23
I17
I8
I11
I29
I20
I35
I6
I13
I37
I5
I22
I34
I3
I26
I28
I39
I40
I33
I25
I31
I36
I32
I1
I10
I21
I38
I15
I14
I12
I30
I2
I27
I18
I24
I4
I16

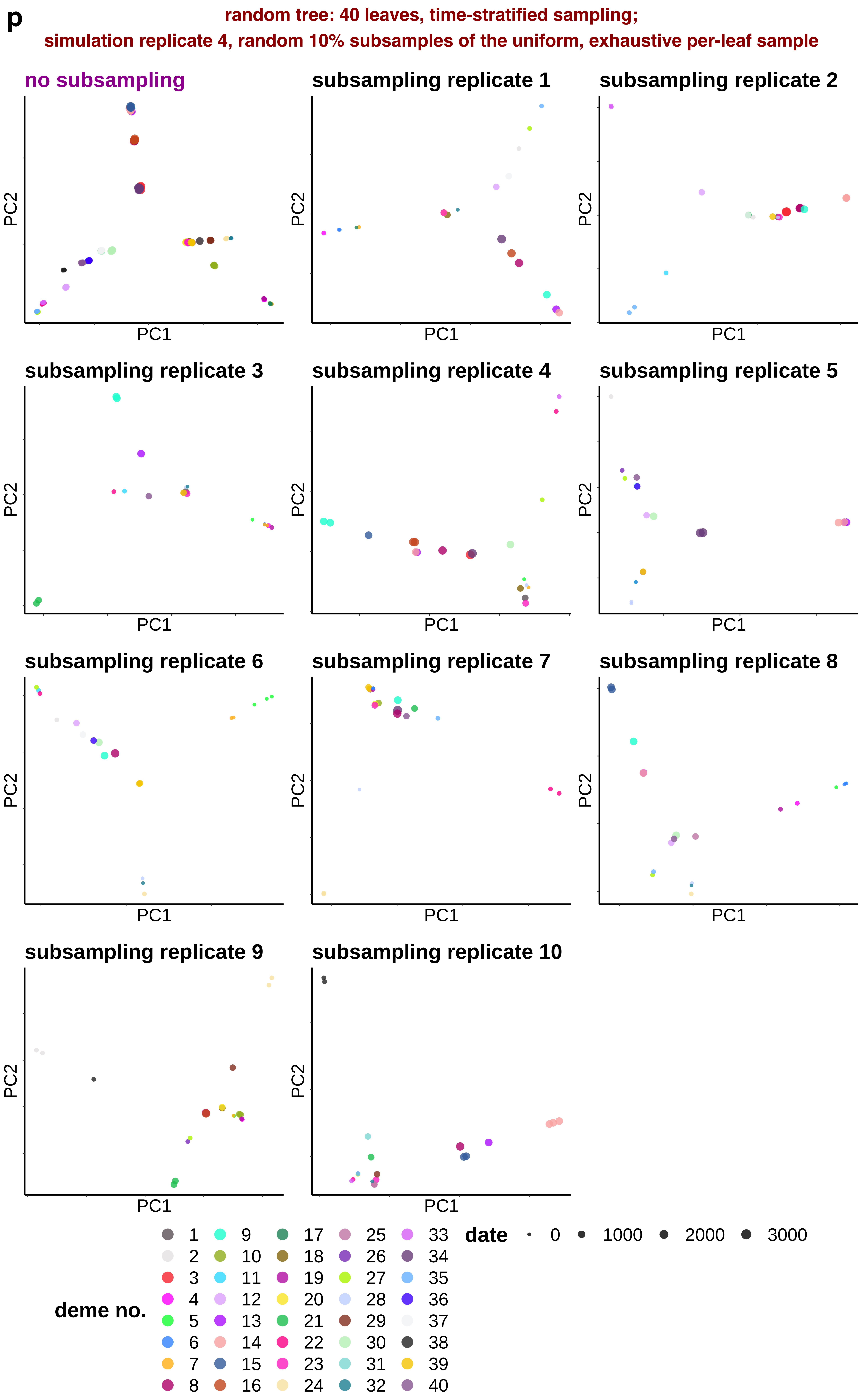
p
random tree: 40 leaves, time-stratified sampling;
simulation replicate 4, random 10% subsamples of the uniform, exhaustive per-leaf sample
no subsampling
subsampling replicate 1
subsampling replicate 2
subsampling replicate 3
subsampling replicate 4
subsampling replicate 5
subsampling replicate 6
subsampling replicate 7
subsampling replicate 8
subsampling replicate 9
subsampling replicate 10
PC1
PC2
deme no.
1 2 3 4 5 6 7 8 9 10 11 12 13 14 15 16 17 18 19 20 21 22 23 24 25 26 27 28 29 30 31 32 33 34 35 36 37 38 39 40
date
0 1000 2000 3000

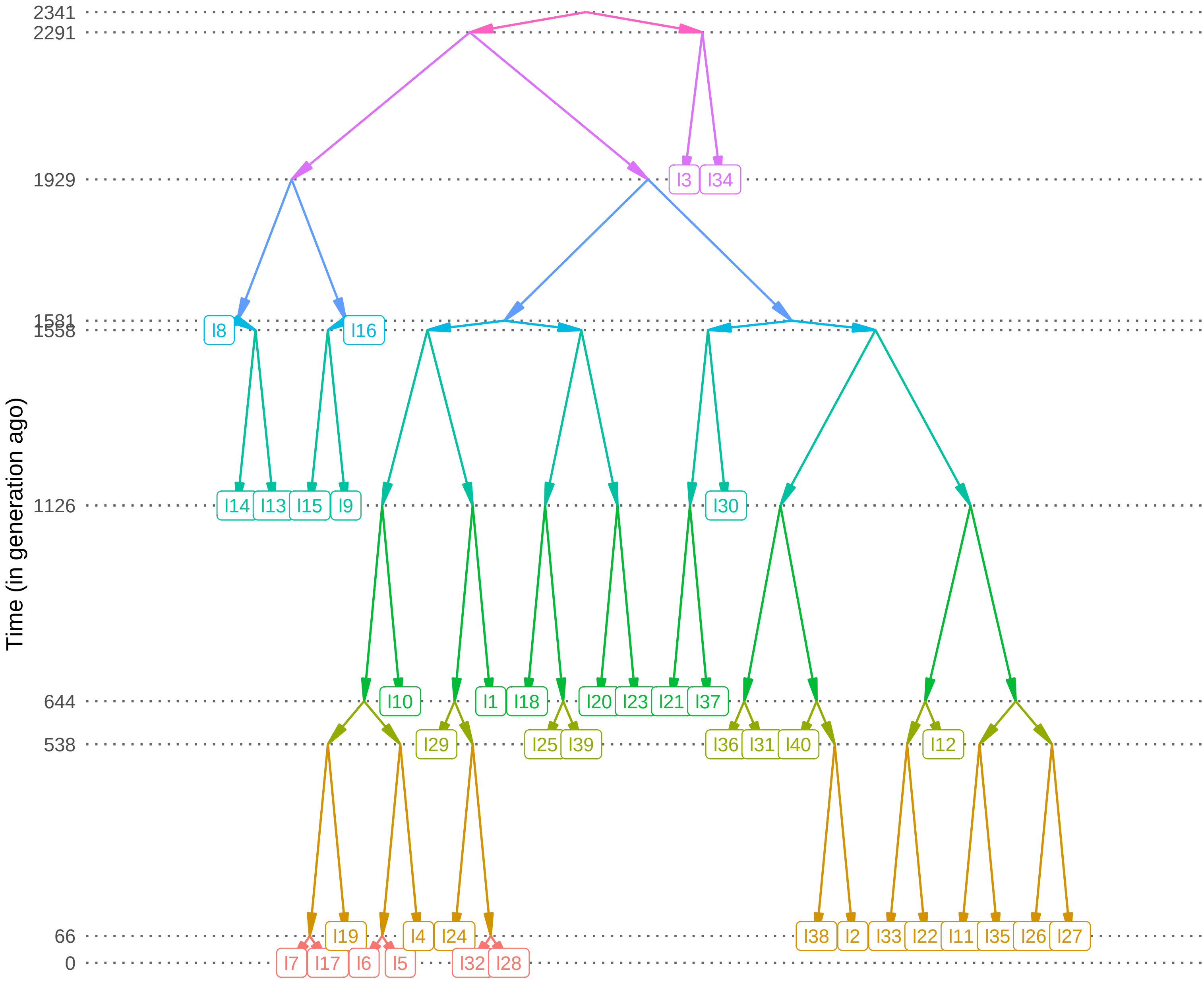
Time (in generation ago)
2341
2291
1929
1581
1558
1126
644
538
66
0
I3
I34
I8
I16
I14
I13
I15
I9
I30
I10
I1
I18
I20
I23
I21
I37
I29
I25
I39
I36
I31
I40
I12
I19
I4
I24
I38
I2
I33
I22
I11
I35
I26
I27
I7
I17
I6
I5
I32
I28
