## Supplementary Figure 5 for "Sparse sampling and rare-variant depletion distort PCA visualizations of population structure: recovery with objective-guided manifold learning"

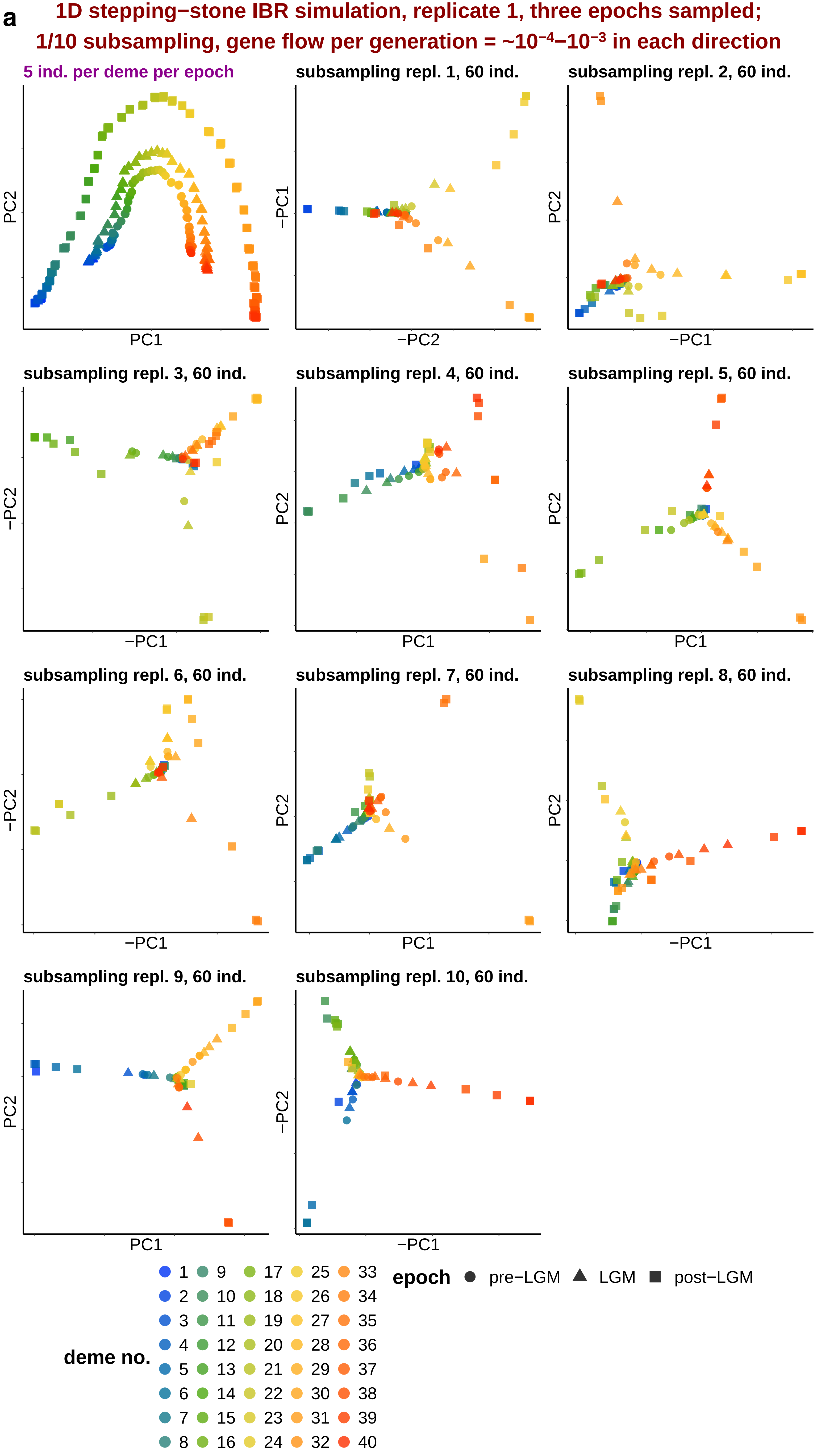
a
1D stepping−stone IBR simulation, replicate 1, three epochs sampled;
1/10 subsampling, gene flow per generation = ~10−4–10−3 in each direction
5 ind. per deme per epoch
PC1
PC2
subsampling repl. 1, 60 ind.
−PC2
−PC1
subsampling repl. 2, 60 ind.
−PC1
PC2
subsampling repl. 3, 60 ind.
−PC1
−PC2
subsampling repl. 4, 60 ind.
PC1
PC2
subsampling repl. 5, 60 ind.
PC1
PC2
subsampling repl. 6, 60 ind.
−PC1
−PC2
subsampling repl. 7, 60 ind.
PC1
PC2
subsampling repl. 8, 60 ind.
−PC1
PC2
subsampling repl. 9, 60 ind.
PC1
PC2
subsampling repl. 10, 60 ind.
−PC1
−PC2
deme no.
1 9 17 25 33
2 10 18 26 34
3 11 19 27 35
4 12 20 28 36
5 13 21 29 37
6 14 22 30 38
7 15 23 31 39
8 16 24 32 40
epoch
pre−LGM
LGM
post−LGM

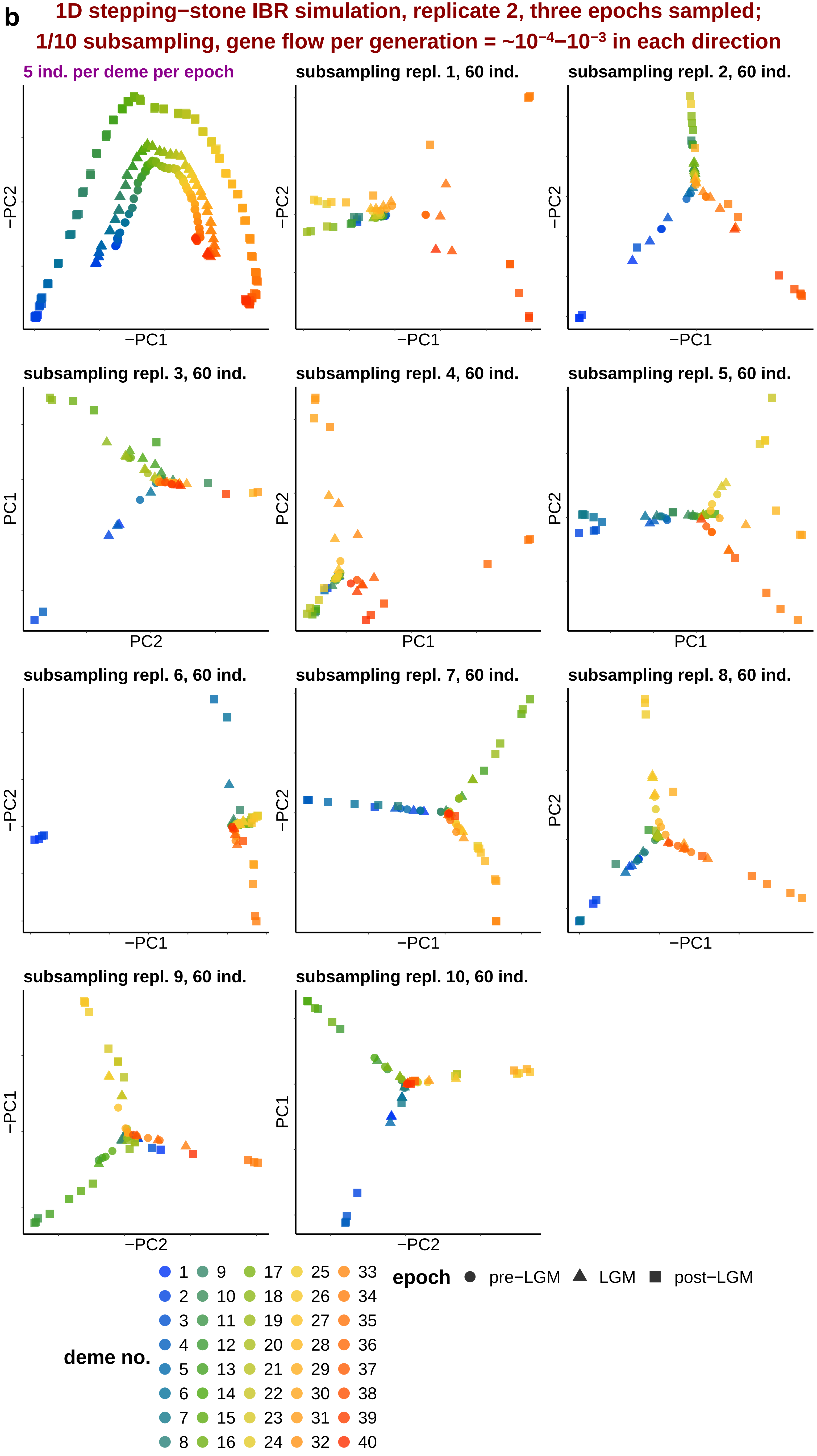
b
1D stepping−stone IBR simulation, replicate 2, three epochs sampled;
1/10 subsampling, gene flow per generation = ~10−4–10−3 in each direction
5 ind. per deme per epoch
subsampling repl. 1, 60 ind.
subsampling repl. 2, 60 ind.
subsampling repl. 3, 60 ind.
subsampling repl. 4, 60 ind.
subsampling repl. 5, 60 ind.
subsampling repl. 6, 60 ind.
subsampling repl. 7, 60 ind.
subsampling repl. 8, 60 ind.
subsampling repl. 9, 60 ind.
subsampling repl. 10, 60 ind.
−PC1
−PC2
PC1
PC2
deme no.
1 2 3 4 5 6 7 8 9 10 11 12 13 14 15 16 17 18 19 20 21 22 23 24 25 26 27 28 29 30 31 32 33 34 35 36 37 38 39 40
epoch
pre−LGM
LGM
post−LGM

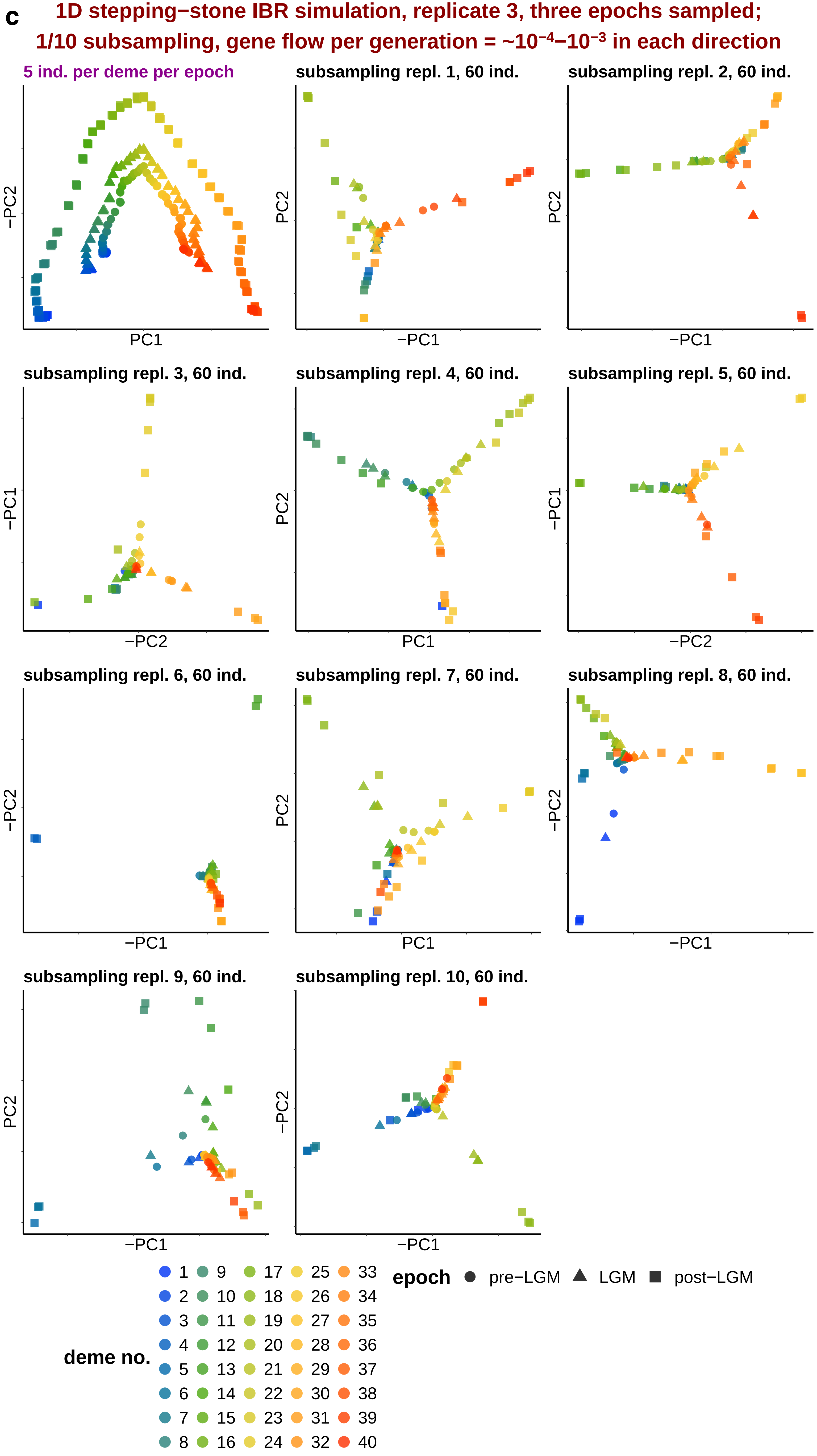
c
1D stepping−stone IBR simulation, replicate 3, three epochs sampled;
1/10 subsampling, gene flow per generation = ~10⁻⁴–10⁻³ in each direction
5 ind. per deme per epoch
subsampling repl. 1, 60 ind.
subsampling repl. 2, 60 ind.
subsampling repl. 3, 60 ind.
subsampling repl. 4, 60 ind.
subsampling repl. 5, 60 ind.
subsampling repl. 6, 60 ind.
subsampling repl. 7, 60 ind.
subsampling repl. 8, 60 ind.
subsampling repl. 9, 60 ind.
subsampling repl. 10, 60 ind.
PC1
−PC1
PC2
−PC2
deme no.
1 2 3 4 5 6 7 8 9 10 11 12 13 14 15 16 17 18 19 20 21 22 23 24 25 26 27 28 29 30 31 32 33 34 35 36 37 38 39 40
epoch
pre−LGM
LGM
post−LGM

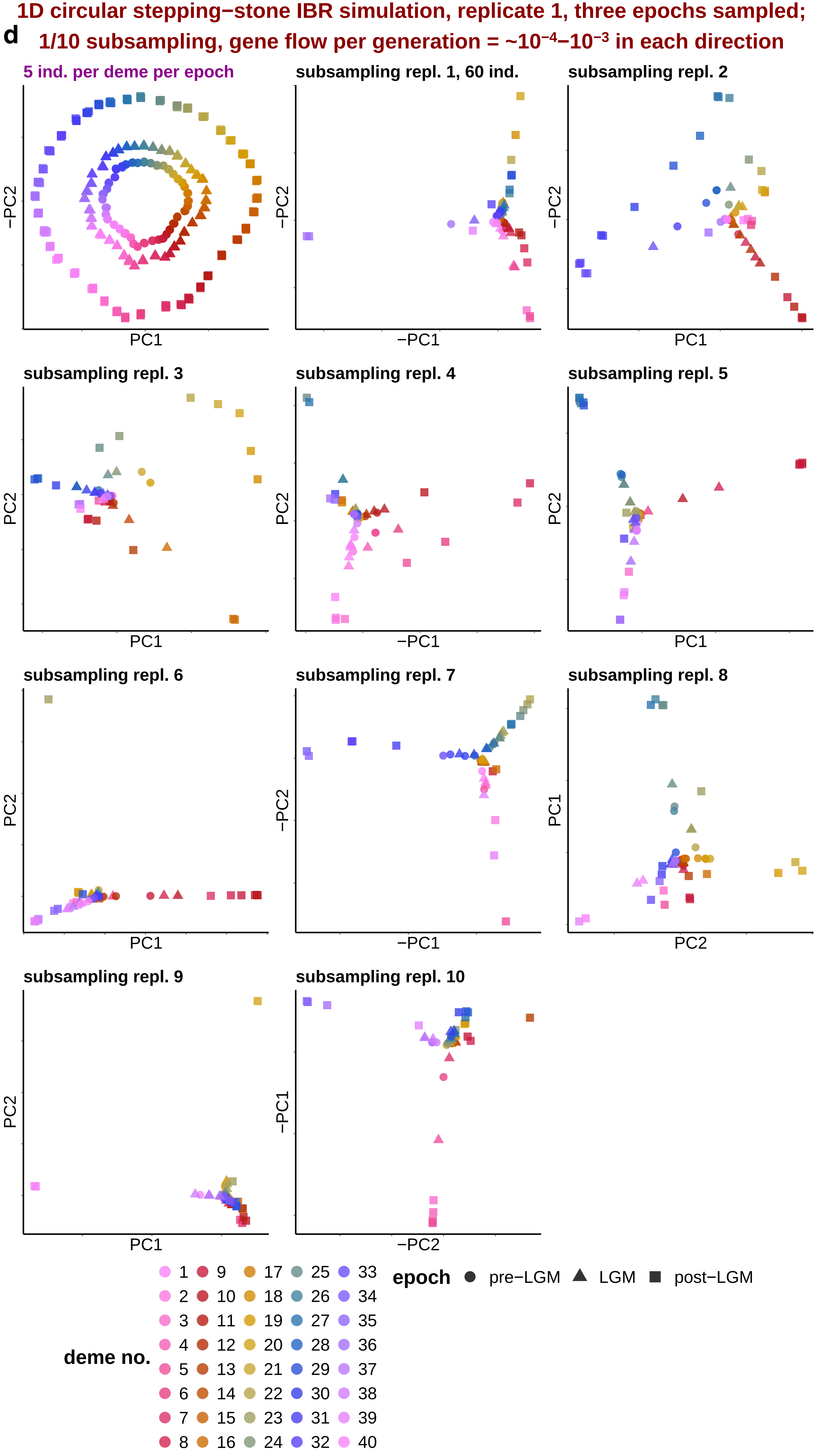
1D circular stepping−stone IBR simulation, replicate 1, three epochs sampled;
d
1/10 subsampling, gene flow per generation = ~10−4−10−3 in each direction
5 ind. per deme per epoch
subsampling repl. 1, 60 ind.
subsampling repl. 2
subsampling repl. 3
subsampling repl. 4
subsampling repl. 5
subsampling repl. 6
subsampling repl. 7
subsampling repl. 8
subsampling repl. 9
subsampling repl. 10
PC1
−PC1
PC2
−PC2
PC1
−PC1
deme no.
1 2 3 4 5 6 7 8 9 10 11 12 13 14 15 16 17 18 19 20 21 22 23 24 25 26 27 28 29 30 31 32 33 34 35 36 37 38 39 40
epoch
pre−LGM
LGM
post−LGM

**e** **1D circular stepping−stone IBR simulation, replicate 2, three epochs sampled; 1/10 subsampling, gene flow per generation = ~$10^{-4}$−$10^{-3}$ in each direction**

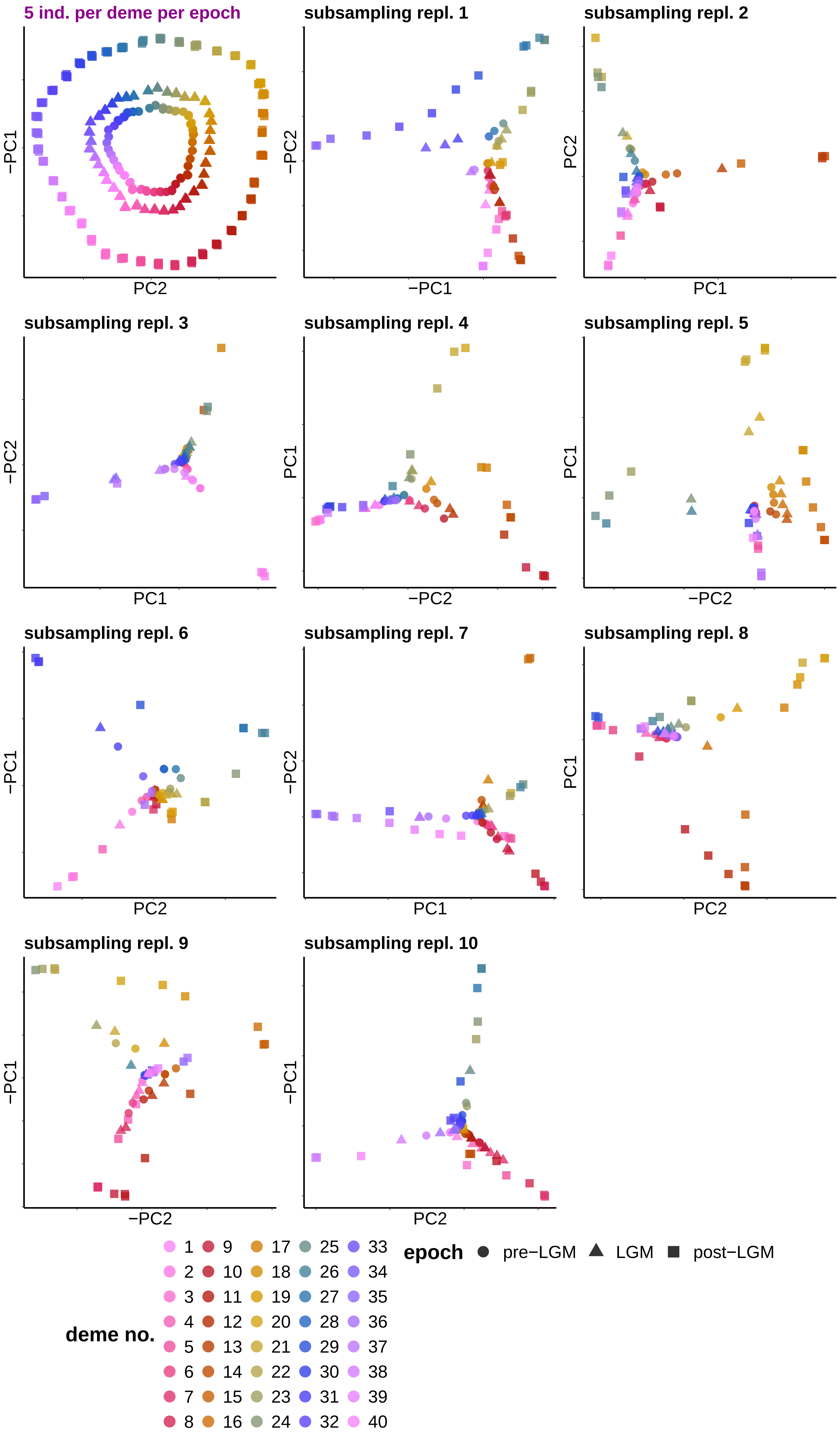

**f** 1D circular stepping−stone IBR simulation, replicate 3, three epochs sampled; 1/10 subsampling, gene flow per generation = ~$10^{-4}$−$10^{-3}$ in each direction

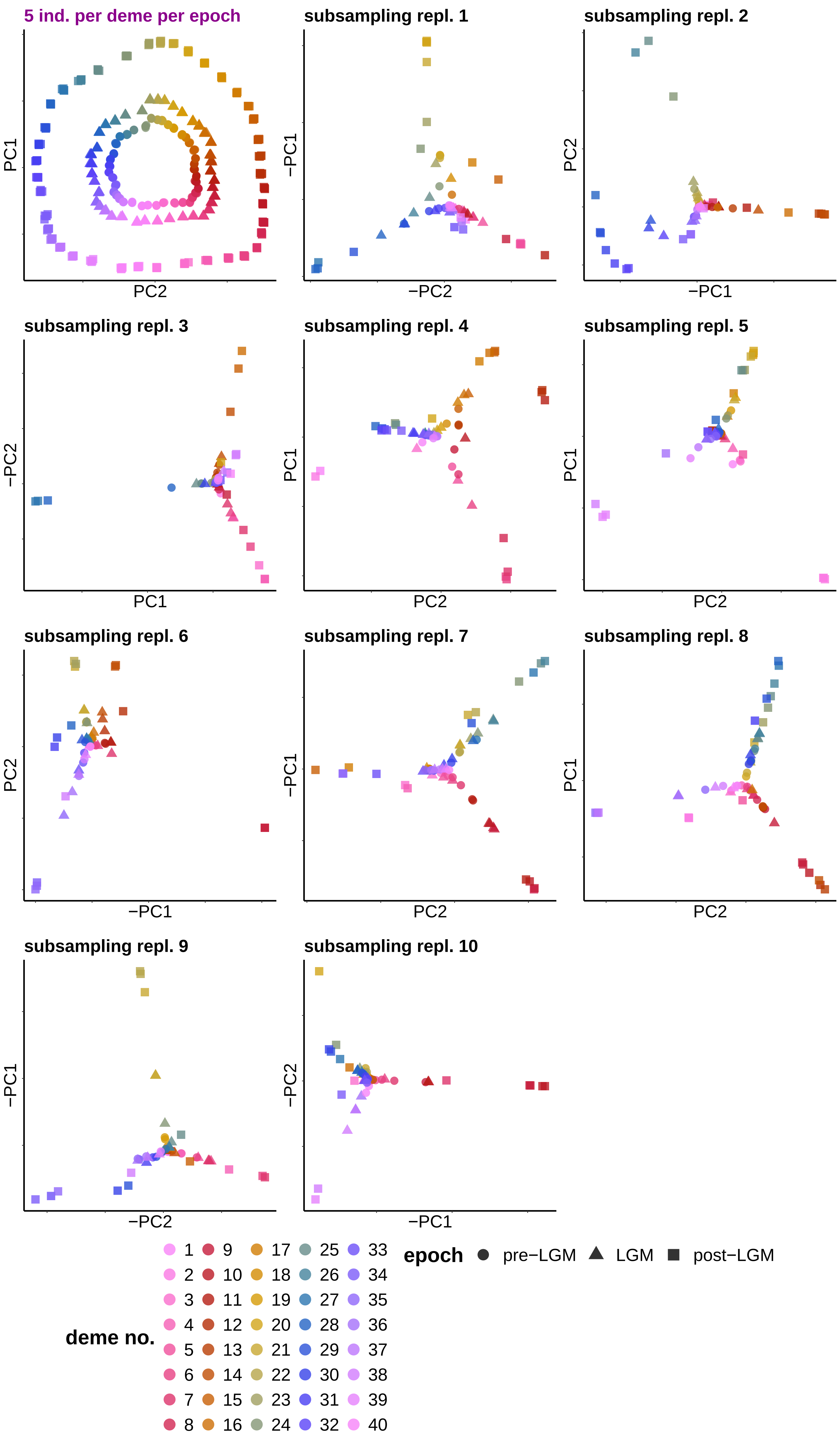
