## Supplementary Figure 6 for "Sparse sampling and rare-variant depletion distort PCA visualizations of population structure: recovery with objective-guided manifold learning"

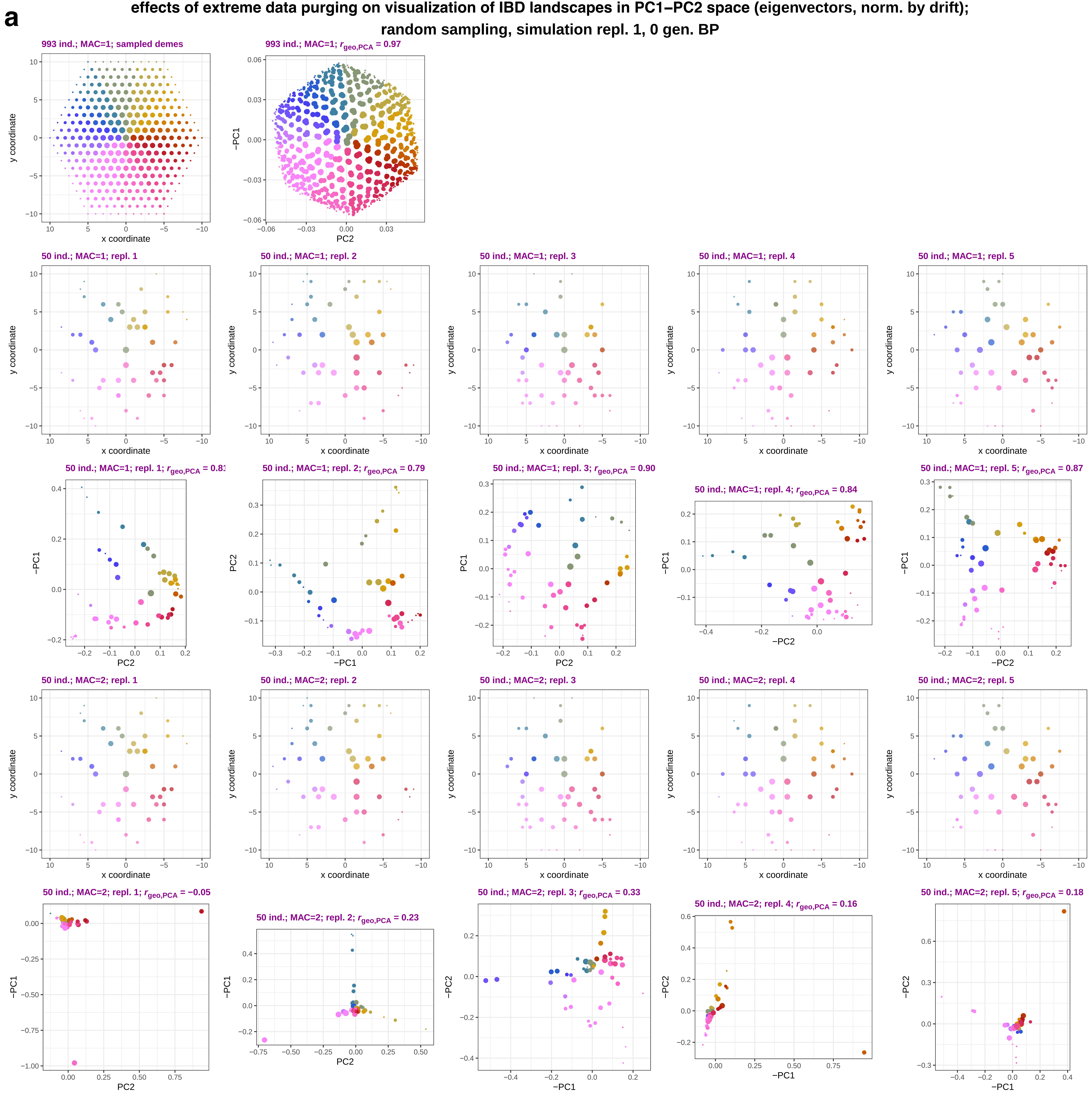

a
effects of extreme data purging on visualization of IBD landscapes in PC1–PC2 space (eigenvectors, norm. by drift);
random sampling, simulation repl. 1, 0 gen. BP
993 ind.; MAC=1; sampled demes
993 ind.; MAC=1; r_geo,PCA = 0.97
50 ind.; MAC=1; repl. 1
50 ind.; MAC=1; repl. 2
50 ind.; MAC=1; repl. 3
50 ind.; MAC=1; repl. 4
50 ind.; MAC=1; repl. 5
50 ind.; MAC=1; repl. 1; r_geo,PCA = 0.81
50 ind.; MAC=1; repl. 2; r_geo,PCA = 0.79
50 ind.; MAC=1; repl. 3; r_geo,PCA = 0.90
50 ind.; MAC=1; repl. 4; r_geo,PCA = 0.84
50 ind.; MAC=1; repl. 5; r_geo,PCA = 0.87
50 ind.; MAC=2; repl. 1
50 ind.; MAC=2; repl. 2
50 ind.; MAC=2; repl. 3
50 ind.; MAC=2; repl. 4
50 ind.; MAC=2; repl. 5
50 ind.; MAC=2; repl. 1; r_geo,PCA = −0.05
50 ind.; MAC=2; repl. 2; r_geo,PCA = 0.23
50 ind.; MAC=2; repl. 3; r_geo,PCA = 0.33
50 ind.; MAC=2; repl. 4; r_geo,PCA = 0.16
50 ind.; MAC=2; repl. 5; r_geo,PCA = 0.18
x coordinate
y coordinate
PC1
−PC1
PC2
−PC2

**b** **effects of extreme data purging on visualization of IBD landscapes in PC1–PC2 space (eigenvectors, norm. by drift); random sampling, simulation repl. 2, 0 gen. BP**

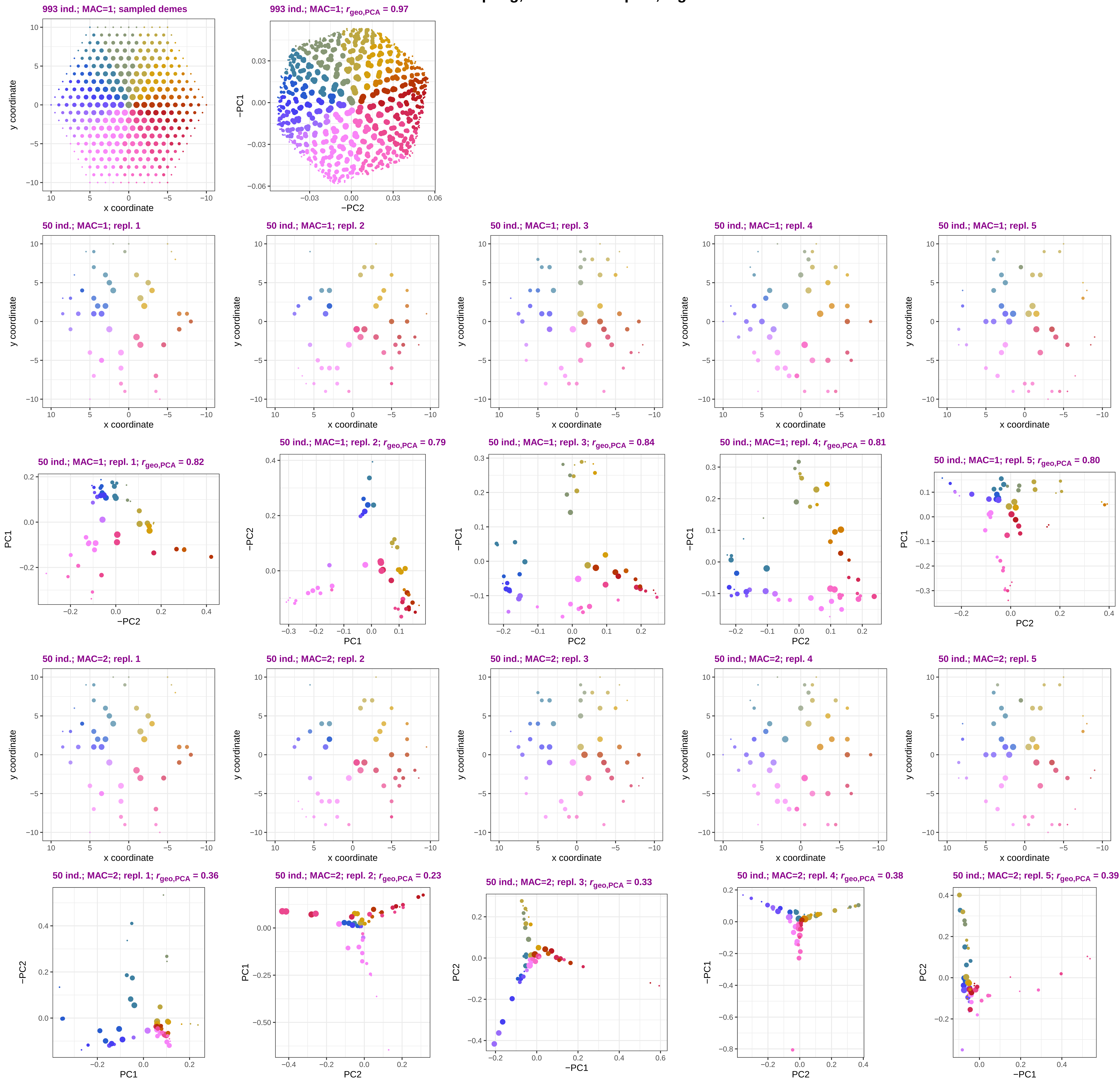

**c** effects of extreme data purging on visualization of IBD landscapes in PC1–PC2 space (eigenvectors, norm. by drift); random sampling, simulation repl. 3, 0 gen. BP

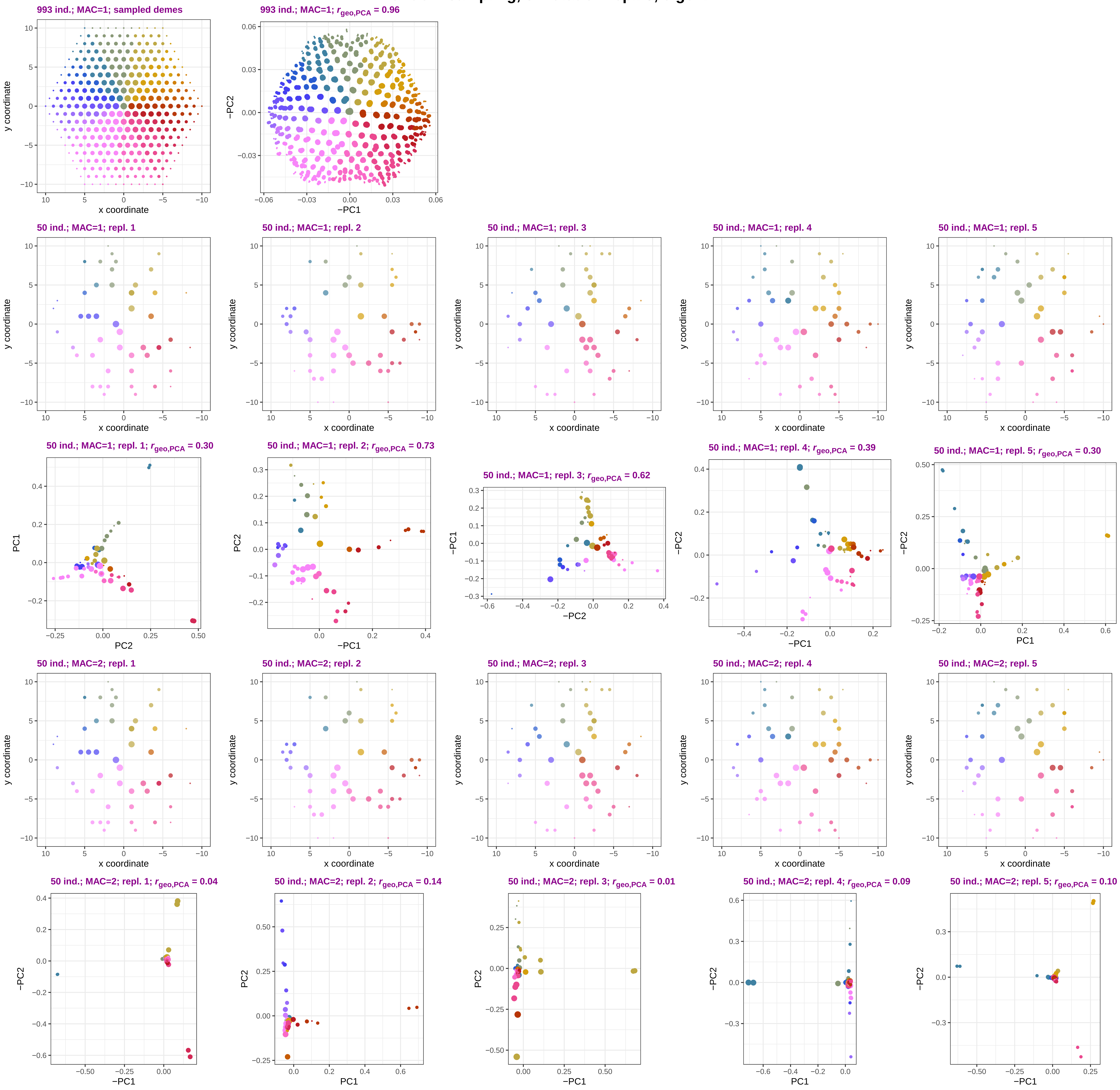

**d**

**effects of extreme data purging on visualization of IBD landscapes in PC1–PC2 space (eigenvectors, norm. by drift); clustered sampling, simulation repl. 1, 0 gen. BP**

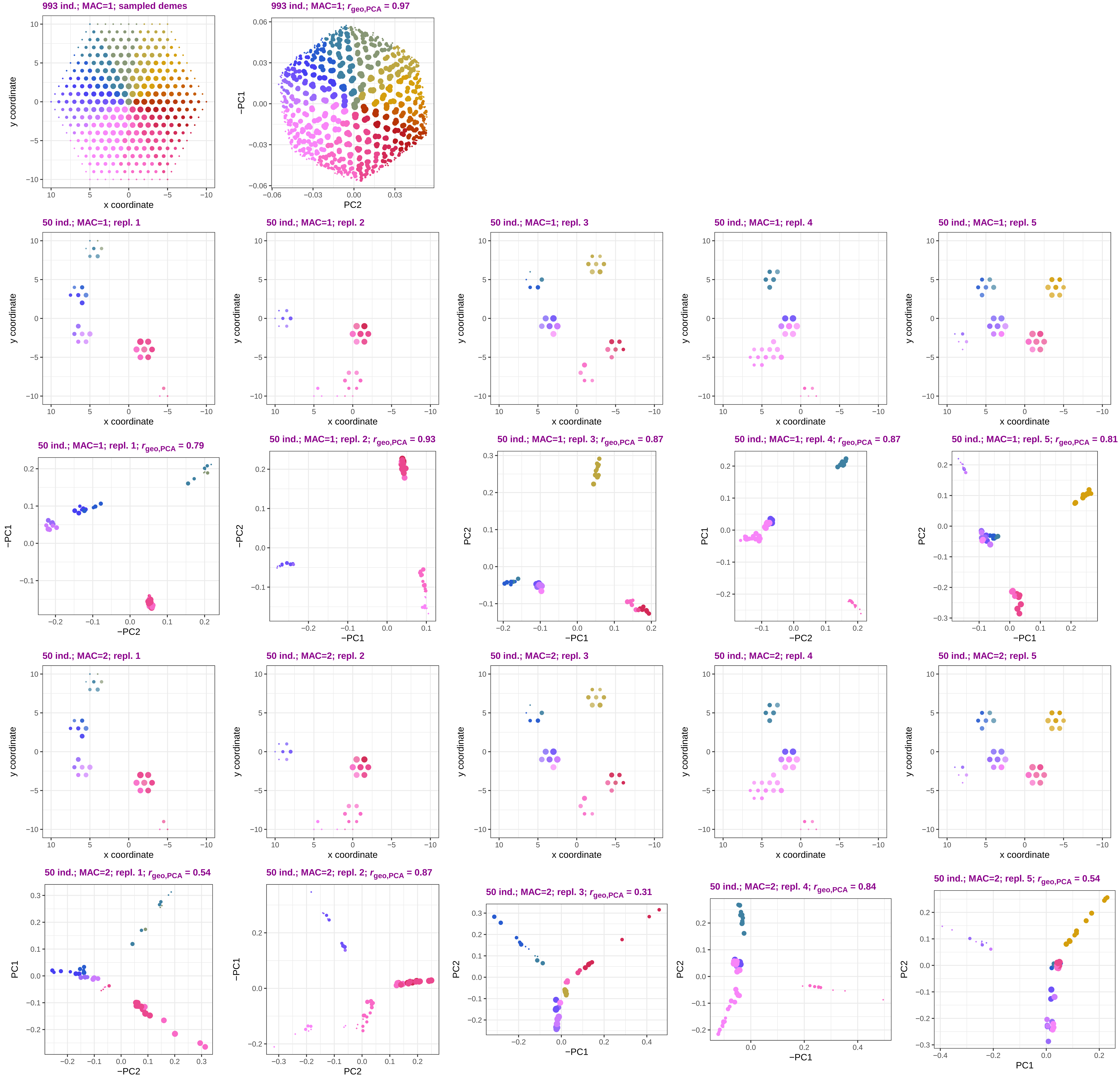

**e** effects of extreme data purging on visualization of IBD landscapes in PC1–PC2 space (eigenvectors, norm. by drift); clustered sampling, simulation repl. 2, 0 gen. BP

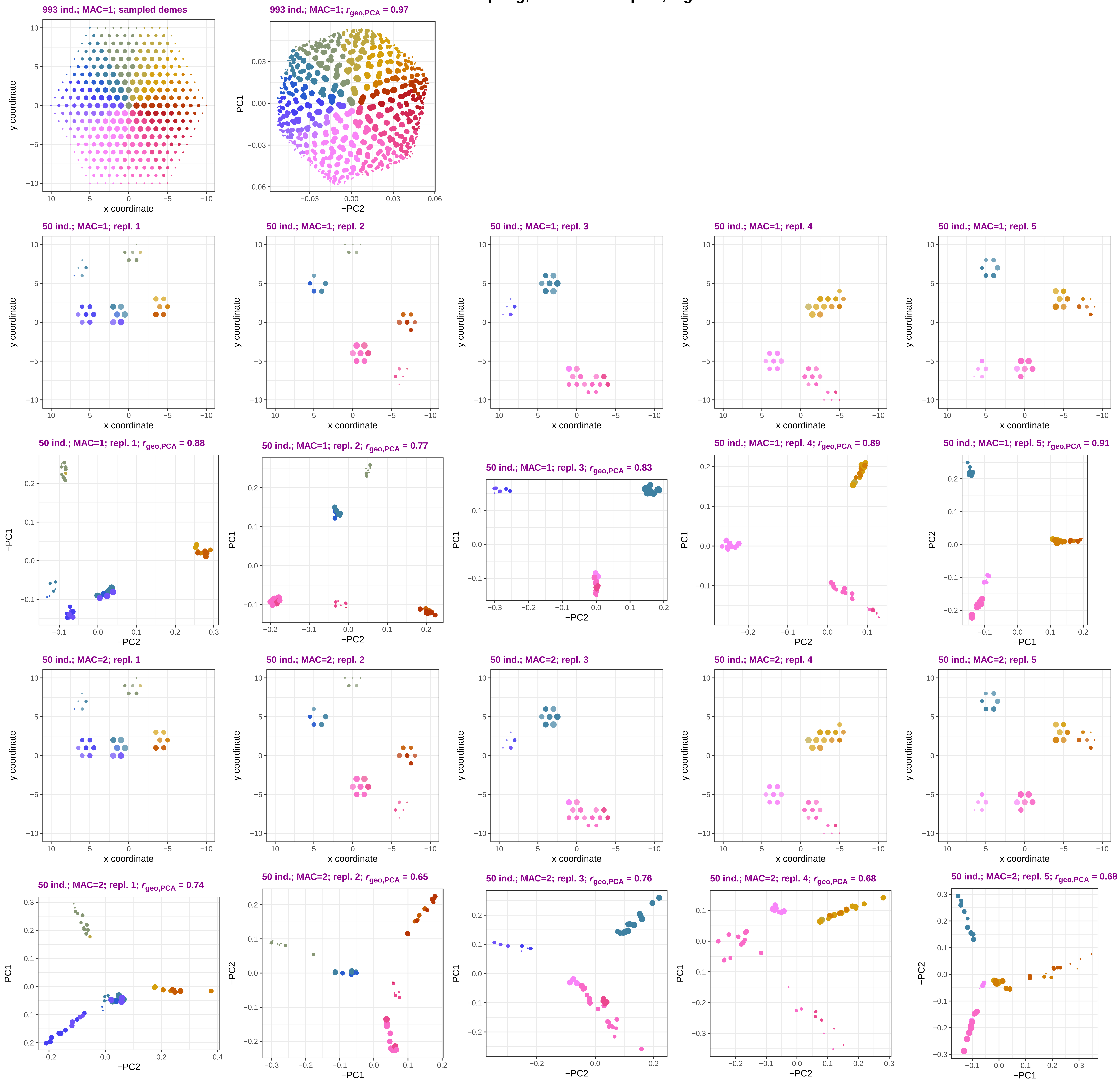

**f**

**effects of extreme data purging on visualization of IBD landscapes in PC1–PC2 space (eigenvectors, norm. by drift); clustered sampling, simulation repl. 3, 0 gen. BP**

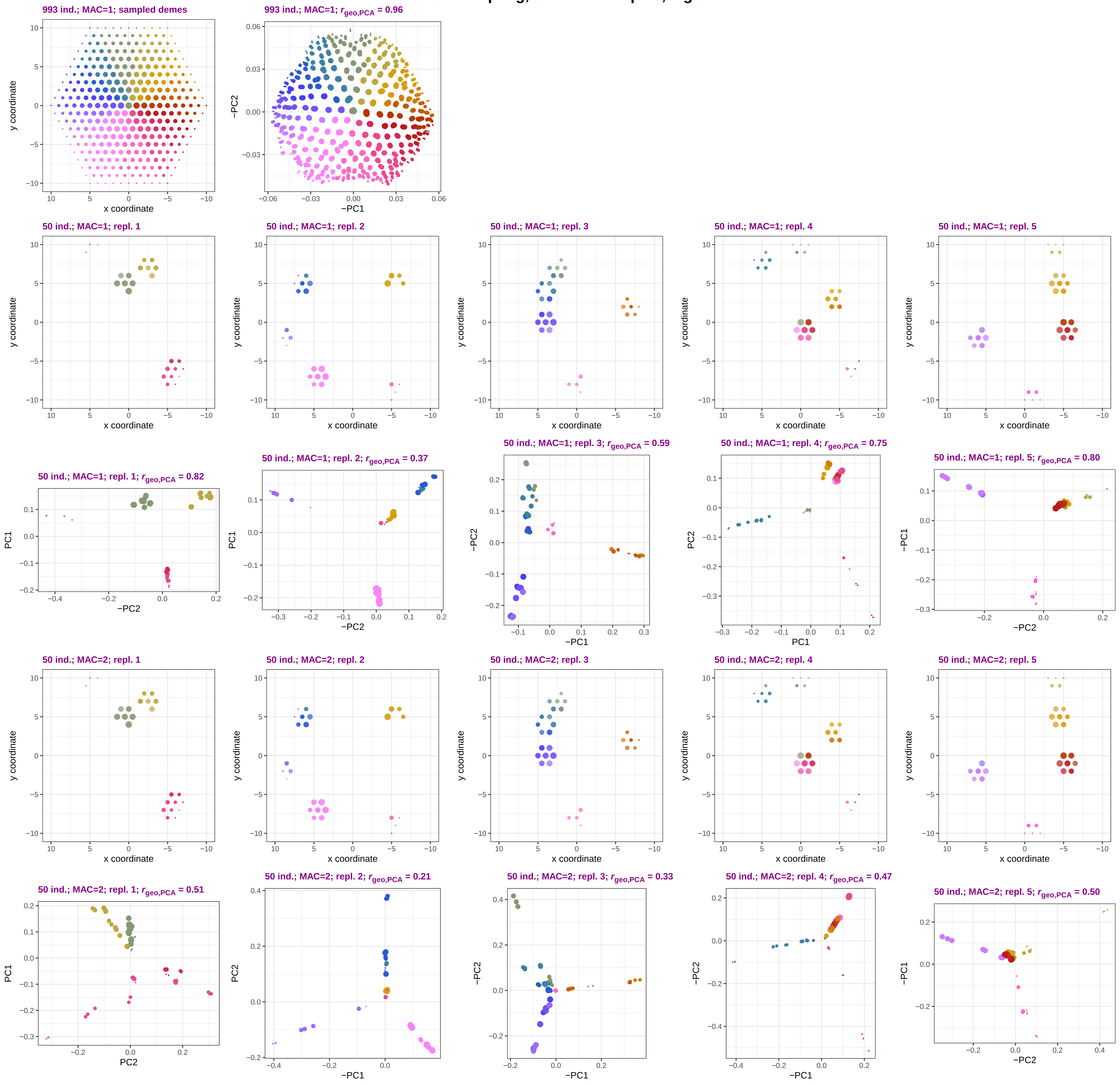
