## Supplementary Figure 8a for "Sparse sampling and rare-variant depletion distort PCA visualizations of population structure: recovery with objective-guided manifold learning"

**Mathematical properties of eigenvectors/PC scores on selected simulated data: eigenvalue and sum of squares; each dot stands for one PC and one simulation/sampling epoch/subsampling replicate**

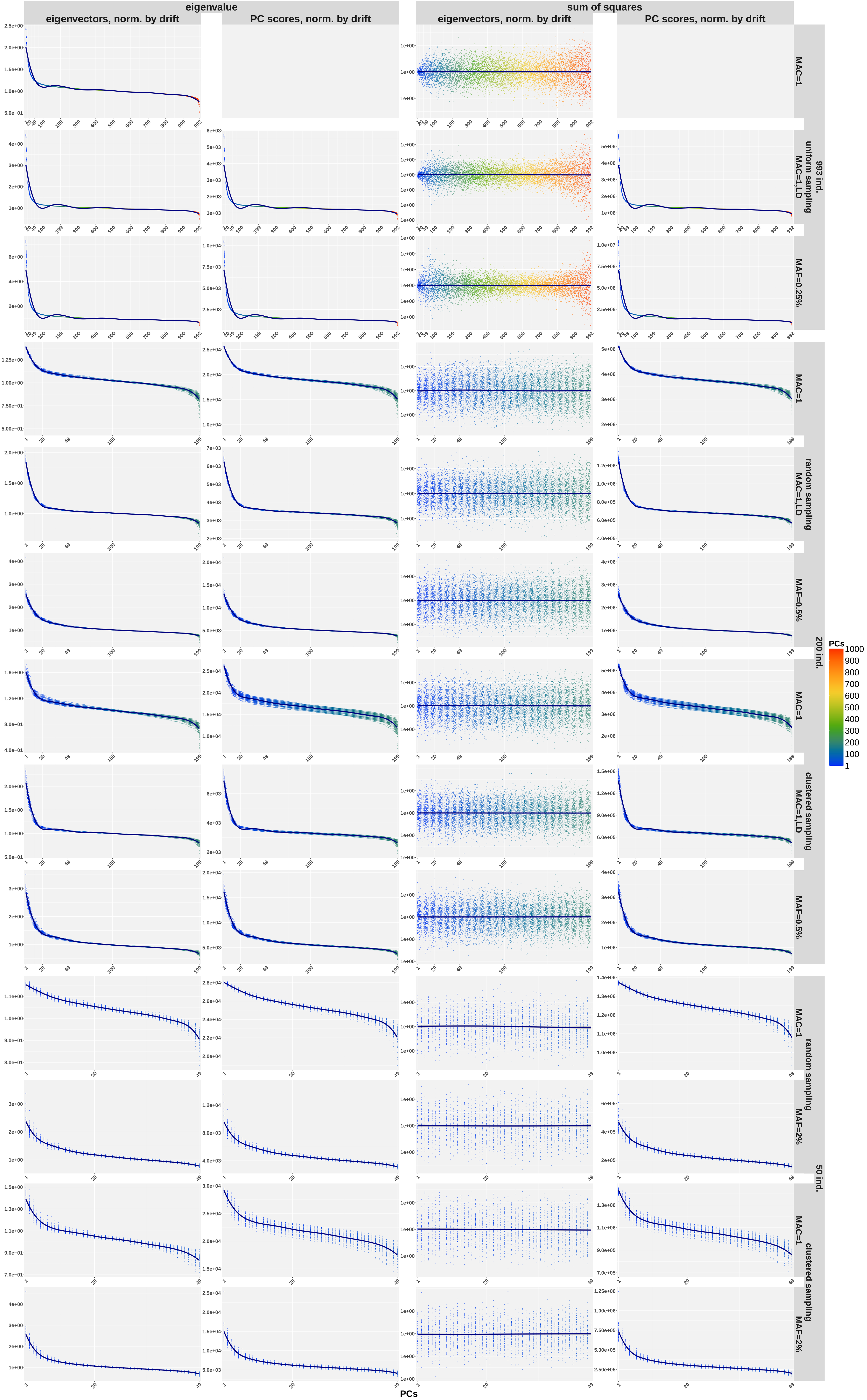
