## Supplementary Figure 12 for "Sparse sampling and rare-variant depletion distort PCA visualizations of population structure: recovery with objective-guided manifold learning"

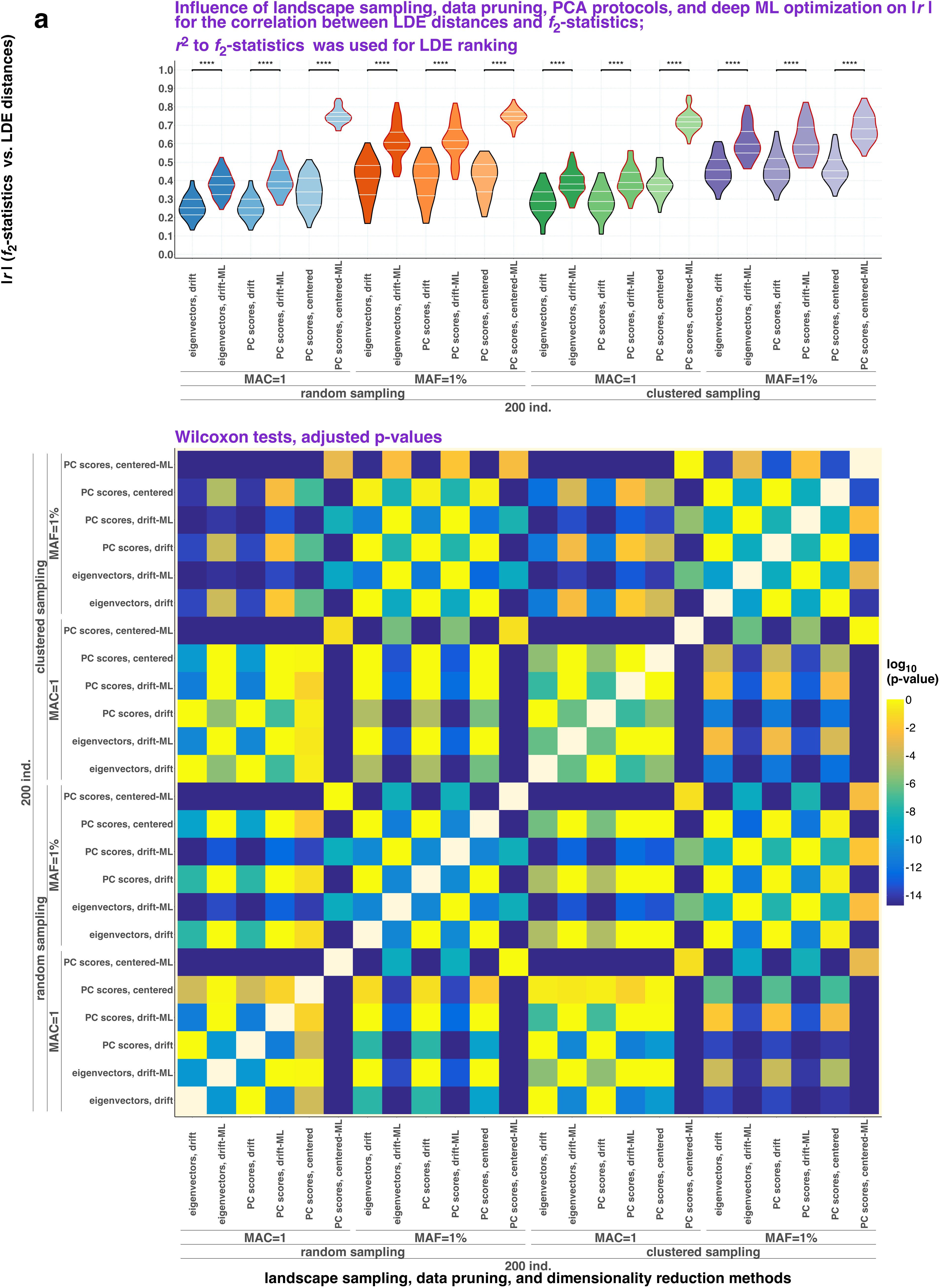
a
Influence of landscape sampling, data pruning, PCA protocols, and deep ML optimization on |r| for the correlation between LDE distances and $f_2$-statistics;
$r^2$ to $f_2$-statistics was used for LDE ranking
|r| ($f_2$-statistics vs. LDE distances)
eigenvectors, drift
eigenvectors, drift-ML
PC scores, drift
PC scores, drift-ML
PC scores, centered
PC scores, centered-ML
MAC=1
MAF=1%
random sampling
clustered sampling
200 ind.
Wilcoxon tests, adjusted p-values
log10 (p-value)
landscape sampling, data pruning, and dimensionality reduction methods

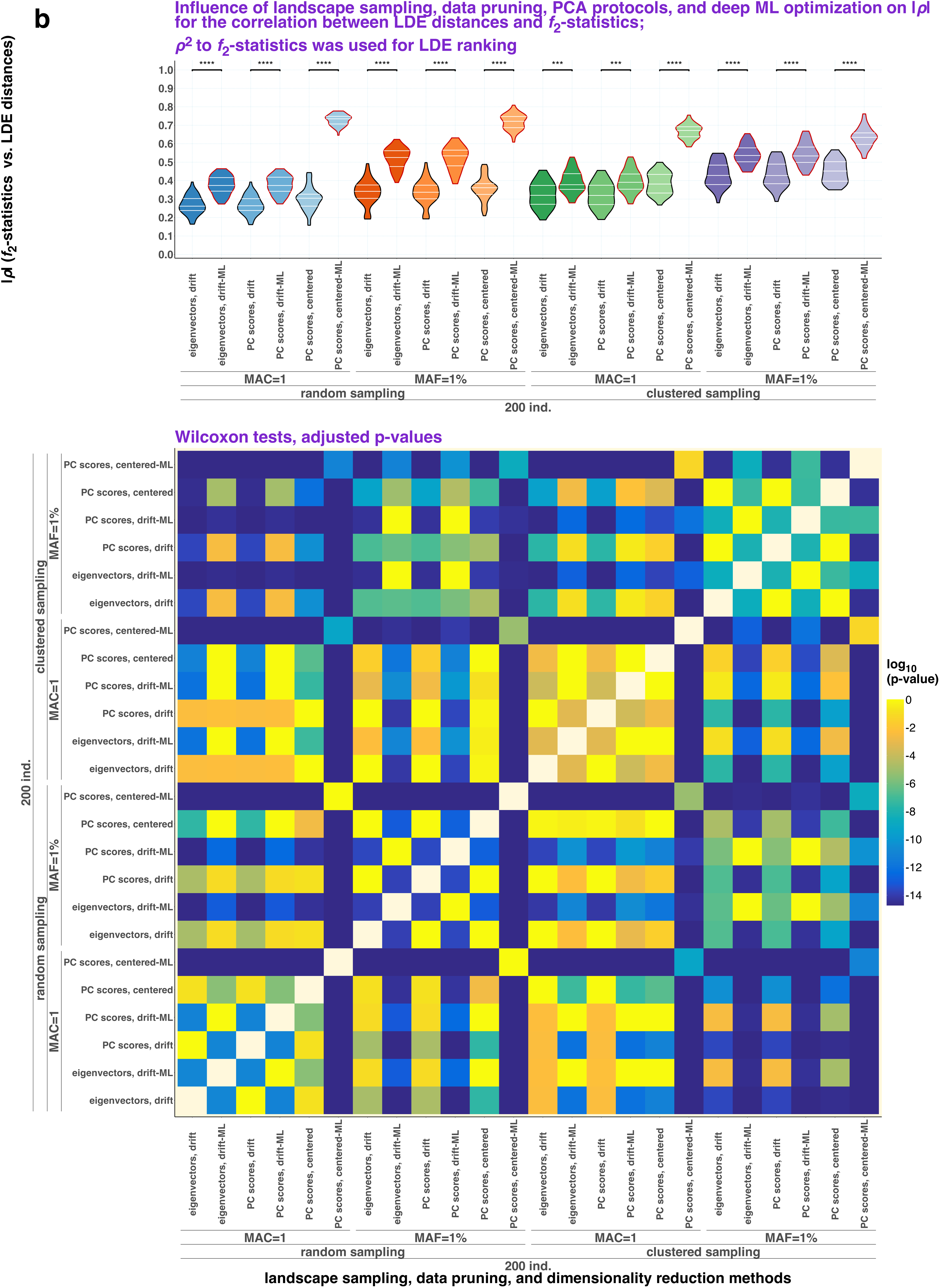
b
Influence of landscape sampling, data pruning, PCA protocols, and deep ML optimization on |ρ| for the correlation between LDE distances and f2-statistics;
ρ2 to f2-statistics was used for LDE ranking
|ρ| (f2-statistics vs. LDE distances)
****
***
1.0
0.9
0.8
0.7
0.6
0.5
0.4
0.3
0.2
0.1
0.0
eigenvectors, drift
eigenvectors, drift-ML
PC scores, drift
PC scores, drift-ML
PC scores, centered
PC scores, centered-ML
MAC=1
MAF=1%
random sampling
clustered sampling
200 ind.
Wilcoxon tests, adjusted p-values
log10 (p-value)
0
-2
-4
-6
-8
-10
-12
-14
landscape sampling, data pruning, and dimensionality reduction methods

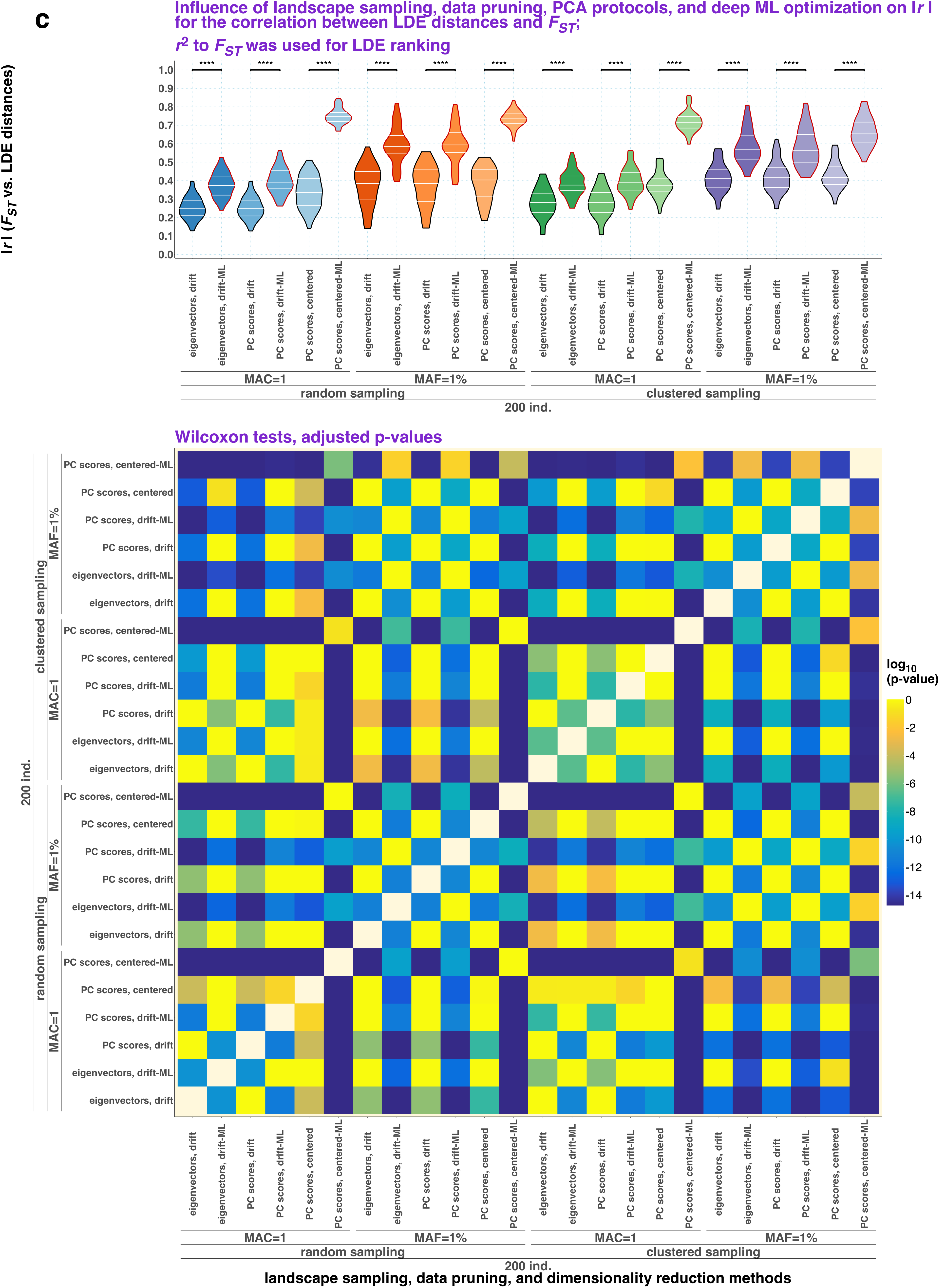
c
Influence of landscape sampling, data pruning, PCA protocols, and deep ML optimization on |r| for the correlation between LDE distances and $F_{ST}$;
$r^2$ to $F_{ST}$ was used for LDE ranking
|r| ($F_{ST}$ vs. LDE distances)
****
eigenvectors, drift
eigenvectors, drift-ML
PC scores, drift
PC scores, drift-ML
PC scores, centered
PC scores, centered-ML
MAC=1
MAF=1%
random sampling
clustered sampling
200 ind.
Wilcoxon tests, adjusted p-values
log10 (p-value)
landscape sampling, data pruning, and dimensionality reduction methods

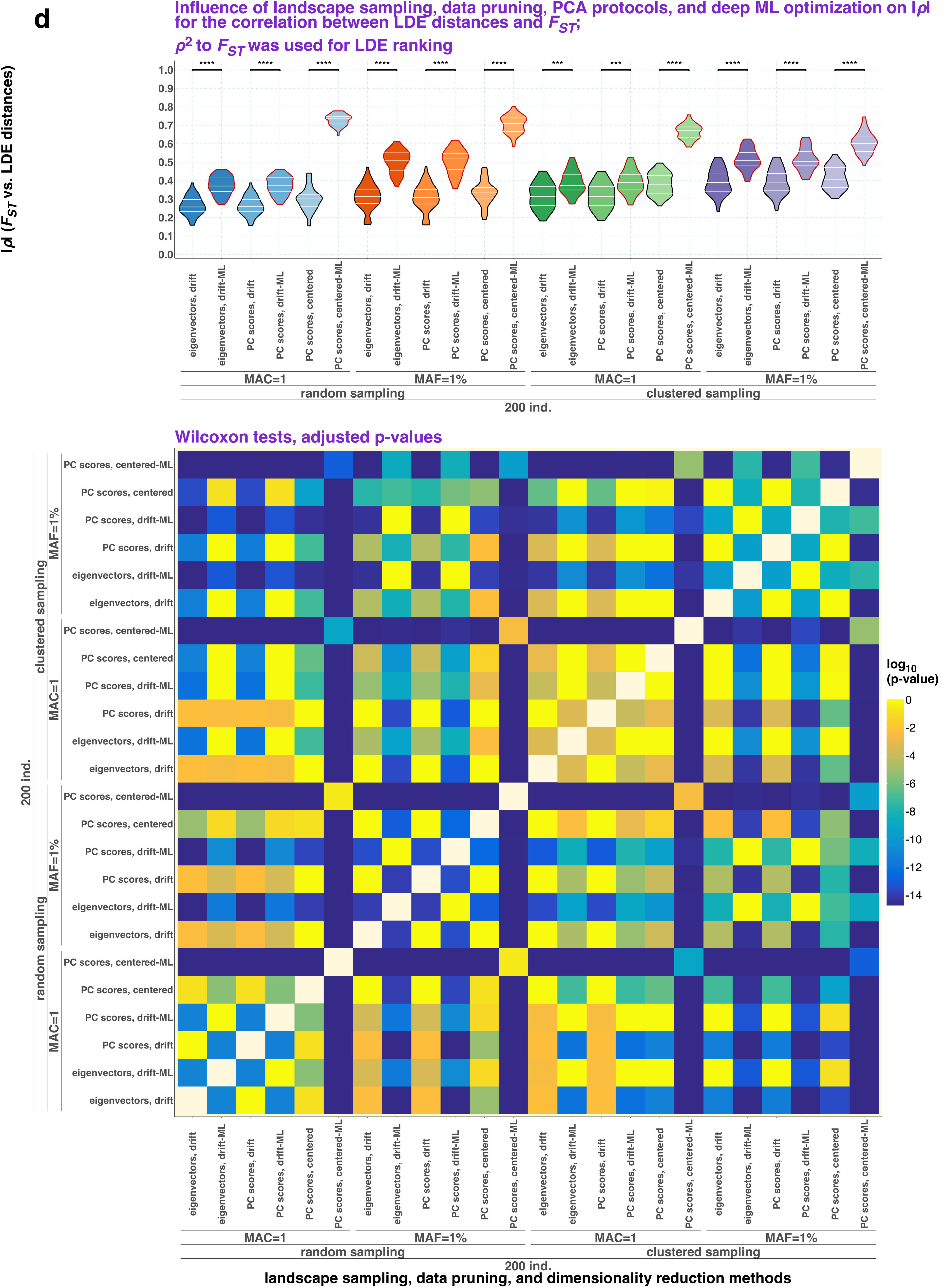
d
Influence of landscape sampling, data pruning, PCA protocols, and deep ML optimization on |ρ| for the correlation between LDE distances and $F_{ST}$;
$\rho^2$ to $F_{ST}$ was used for LDE ranking
|ρ| ($F_{ST}$ vs. LDE distances)
eigenvectors, drift
eigenvectors, drift-ML
PC scores, drift
PC scores, drift-ML
PC scores, centered
PC scores, centered-ML
MAC=1
MAF=1%
random sampling
clustered sampling
200 ind.
Wilcoxon tests, adjusted p-values
log10 (p-value)
landscape sampling, data pruning, and dimensionality reduction methods

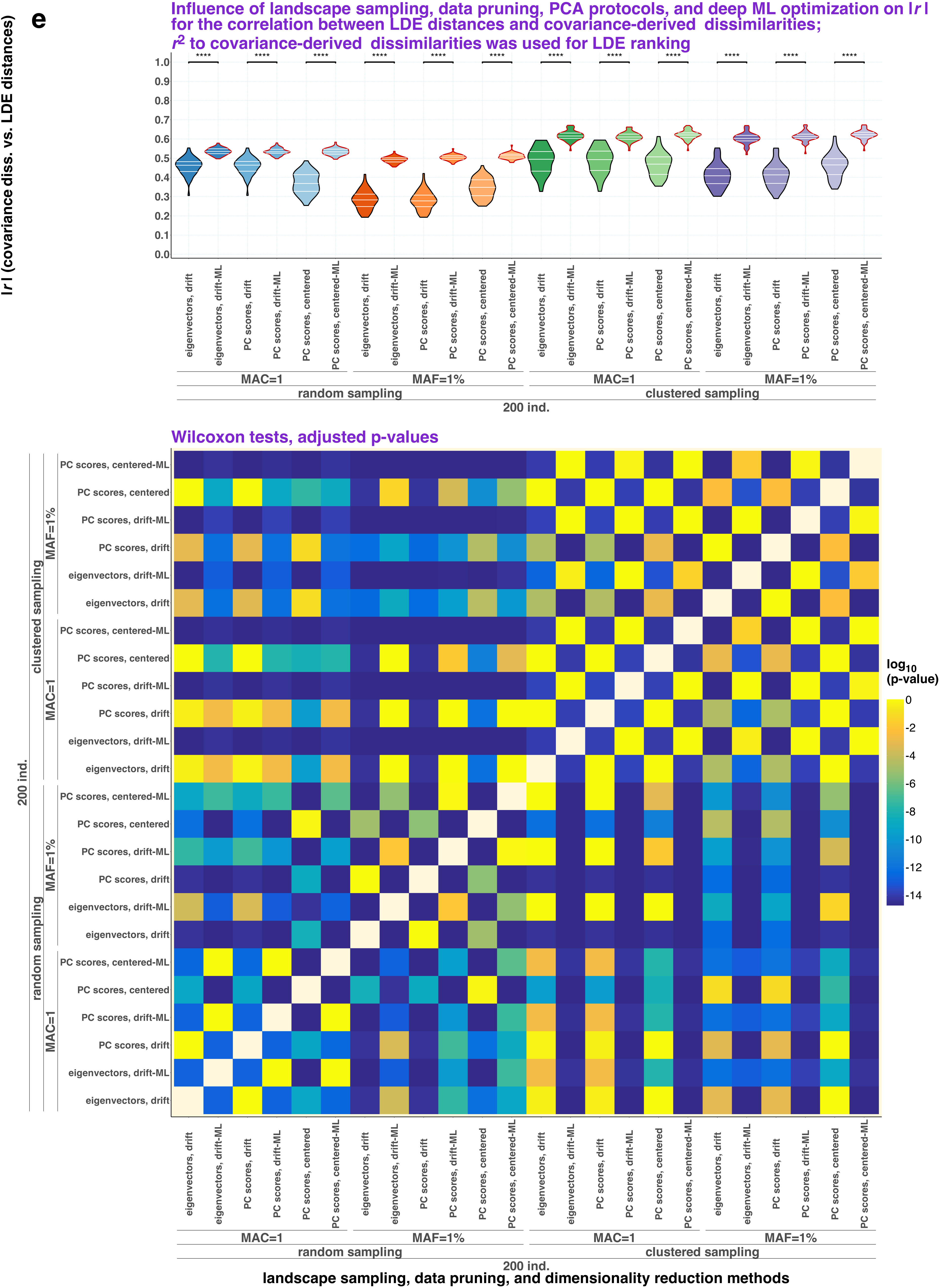

e
Influence of landscape sampling, data pruning, PCA protocols, and deep ML optimization on |r| for the correlation between LDE distances and covariance-derived dissimilarities; $r^2$ to covariance-derived dissimilarities was used for LDE ranking
|r| (covariance diss. vs. LDE distances)
****
eigenvectors, drift
eigenvectors, drift-ML
PC scores, drift
PC scores, drift-ML
PC scores, centered
PC scores, centered-ML
MAC=1
MAF=1%
random sampling
clustered sampling
200 ind.
Wilcoxon tests, adjusted p-values
$\log_{10}$ (p-value)
landscape sampling, data pruning, and dimensionality reduction methods

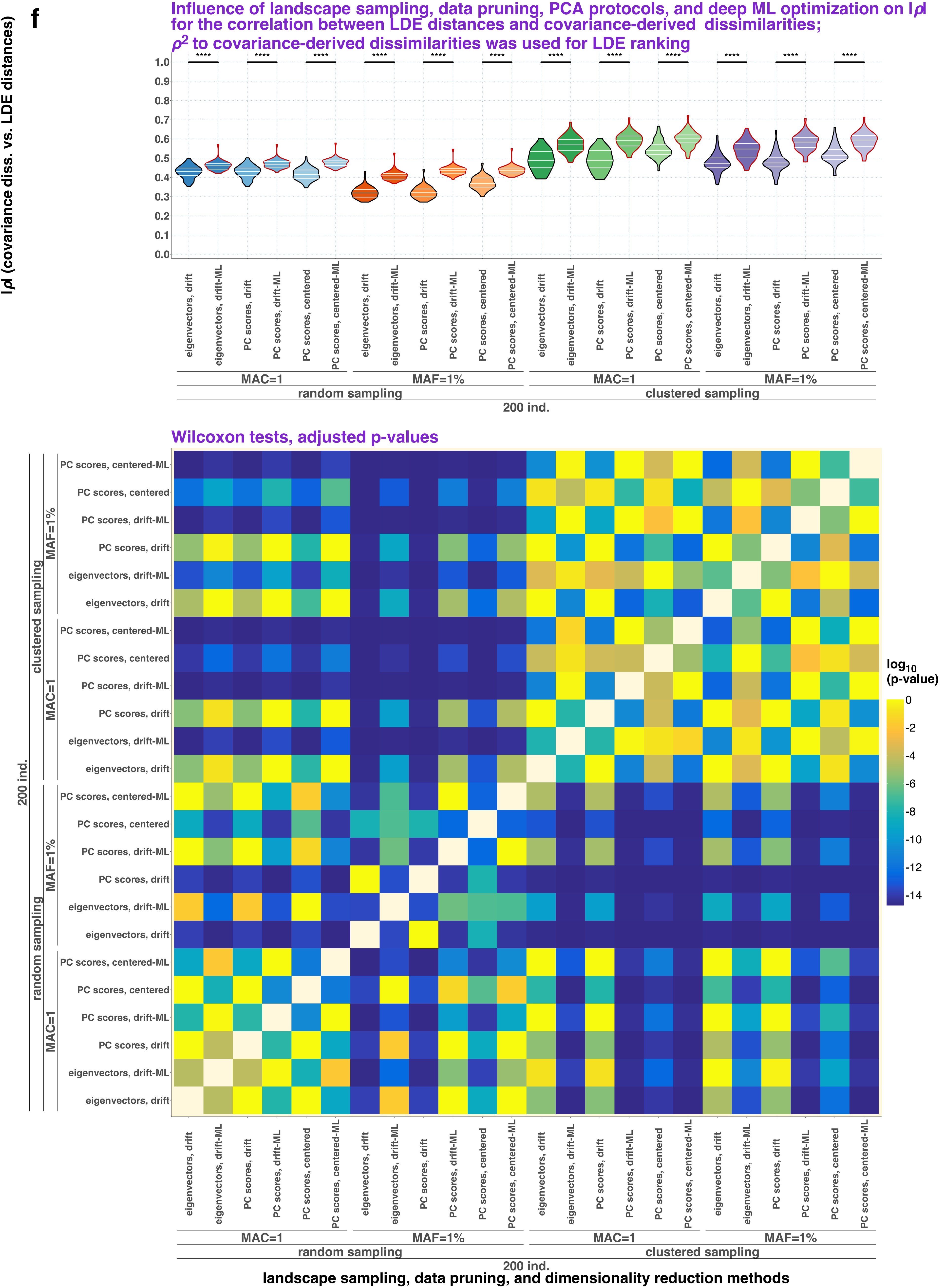

f
Influence of landscape sampling, data pruning, PCA protocols, and deep ML optimization on |ρ| for the correlation between LDE distances and covariance-derived dissimilarities;
ρ2 to covariance-derived dissimilarities was used for LDE ranking
|ρ| (covariance diss. vs. LDE distances)
eigenvectors, drift
eigenvectors, drift-ML
PC scores, drift
PC scores, drift-ML
PC scores, centered
PC scores, centered-ML
MAC=1
MAF=1%
random sampling
clustered sampling
200 ind.
Wilcoxon tests, adjusted p-values
log10 (p-value)
landscape sampling, data pruning, and dimensionality reduction methods

g

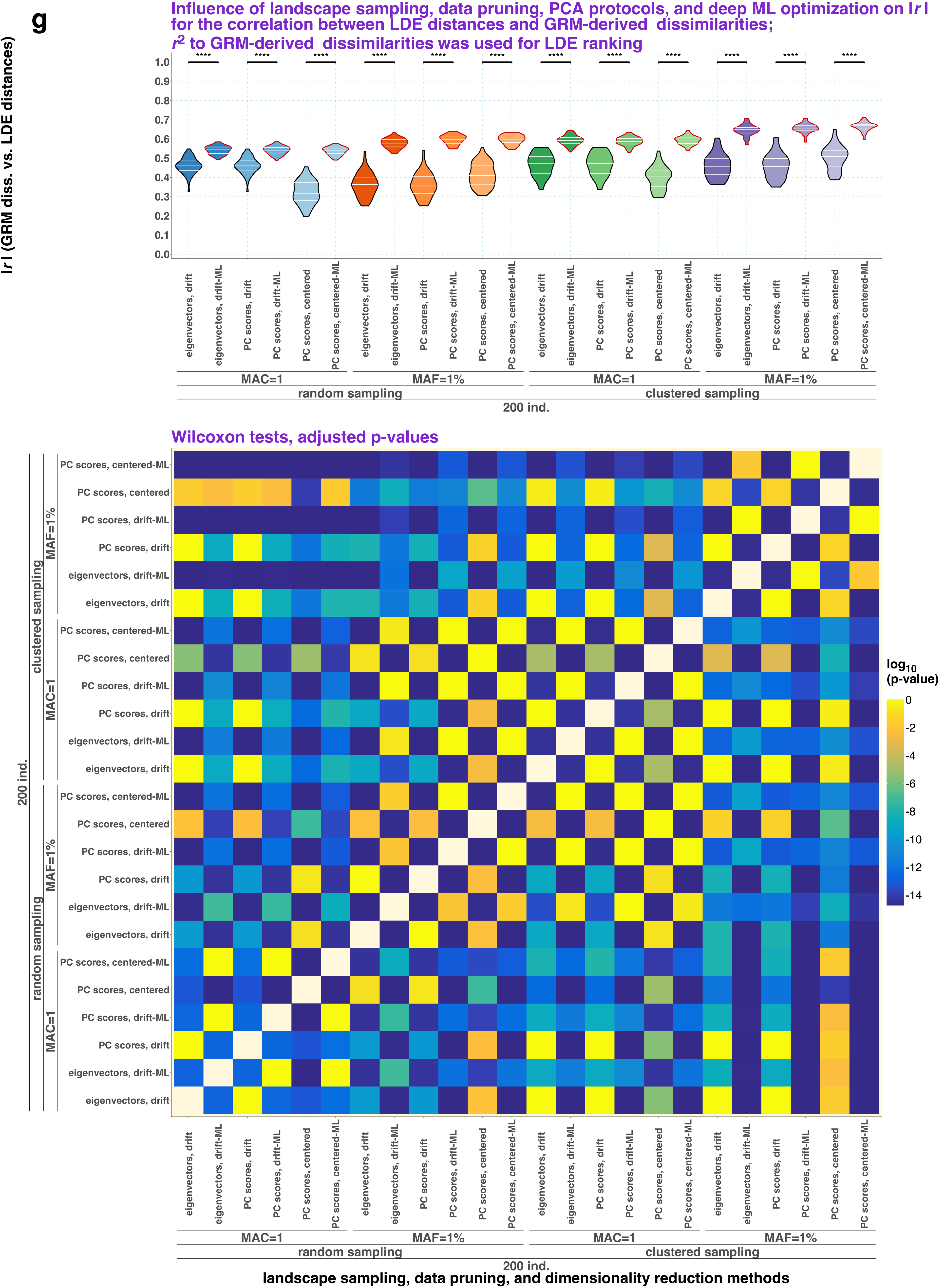

Influence of landscape sampling, data pruning, PCA protocols, and deep ML optimization on |r| for the correlation between LDE distances and GRM-derived dissimilarities; $r^2$ to GRM-derived dissimilarities was used for LDE ranking
|r| (GRM diss. vs. LDE distances)
****
eigenvectors, drift
eigenvectors, drift-ML
PC scores, drift
PC scores, drift-ML
PC scores, centered
PC scores, centered-ML
MAC=1
MAF=1%
random sampling
clustered sampling
200 ind.
Wilcoxon tests, adjusted p-values
log10 (p-value)
landscape sampling, data pruning, and dimensionality reduction methods

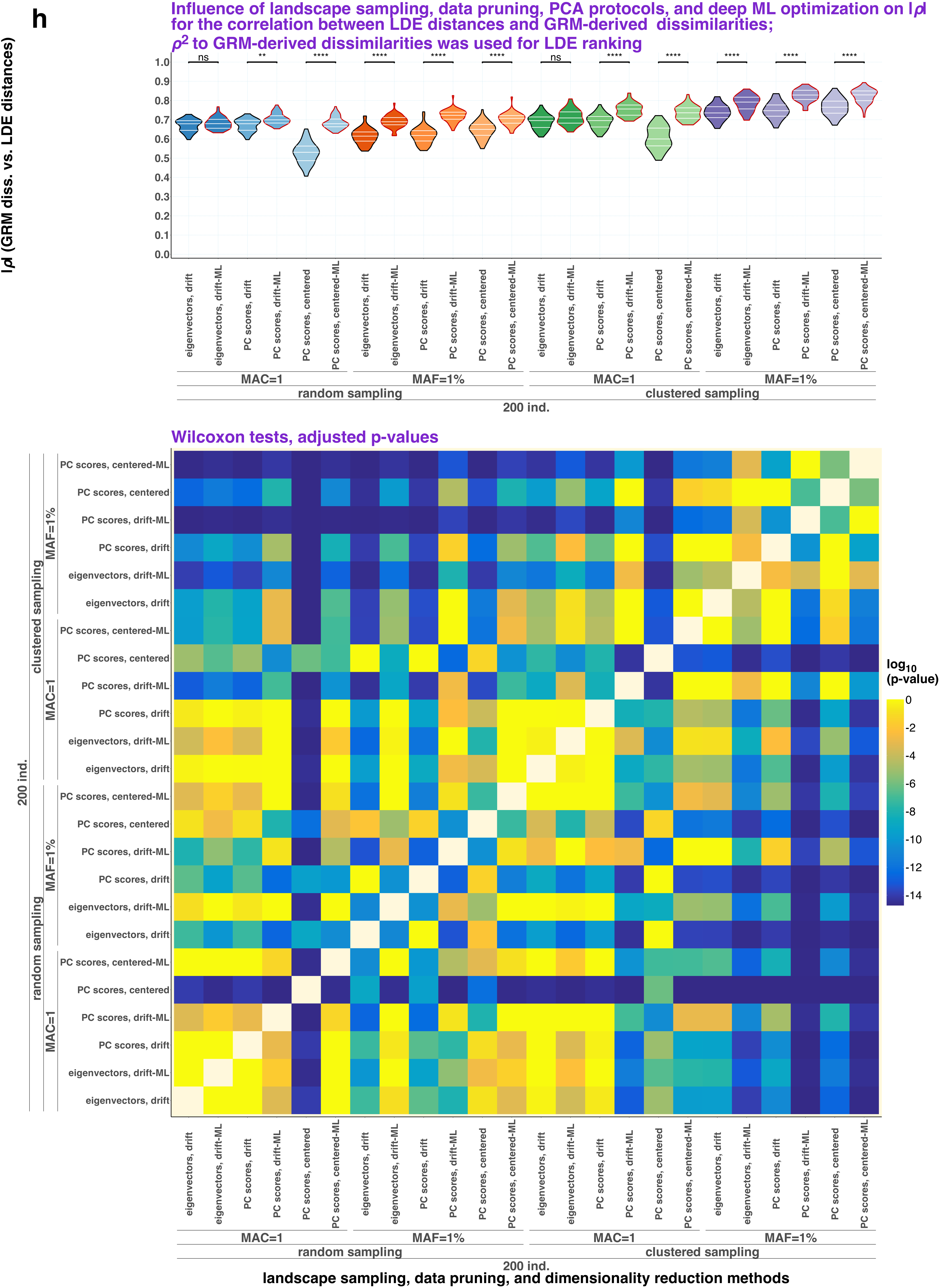
h
Influence of landscape sampling, data pruning, PCA protocols, and deep ML optimization on |ρ| for the correlation between LDE distances and GRM-derived dissimilarities;
ρ² to GRM-derived dissimilarities was used for LDE ranking
|ρ| (GRM diss. vs. LDE distances)
ns
**
****
****
****
****
ns
****
****
****
****
****
eigenvectors, drift
eigenvectors, drift-ML
PC scores, drift
PC scores, drift-ML
PC scores, centered
PC scores, centered-ML
MAC=1
MAF=1%
random sampling
clustered sampling
200 ind.
Wilcoxon tests, adjusted p-values
log10 (p-value)
landscape sampling, data pruning, and dimensionality reduction methods

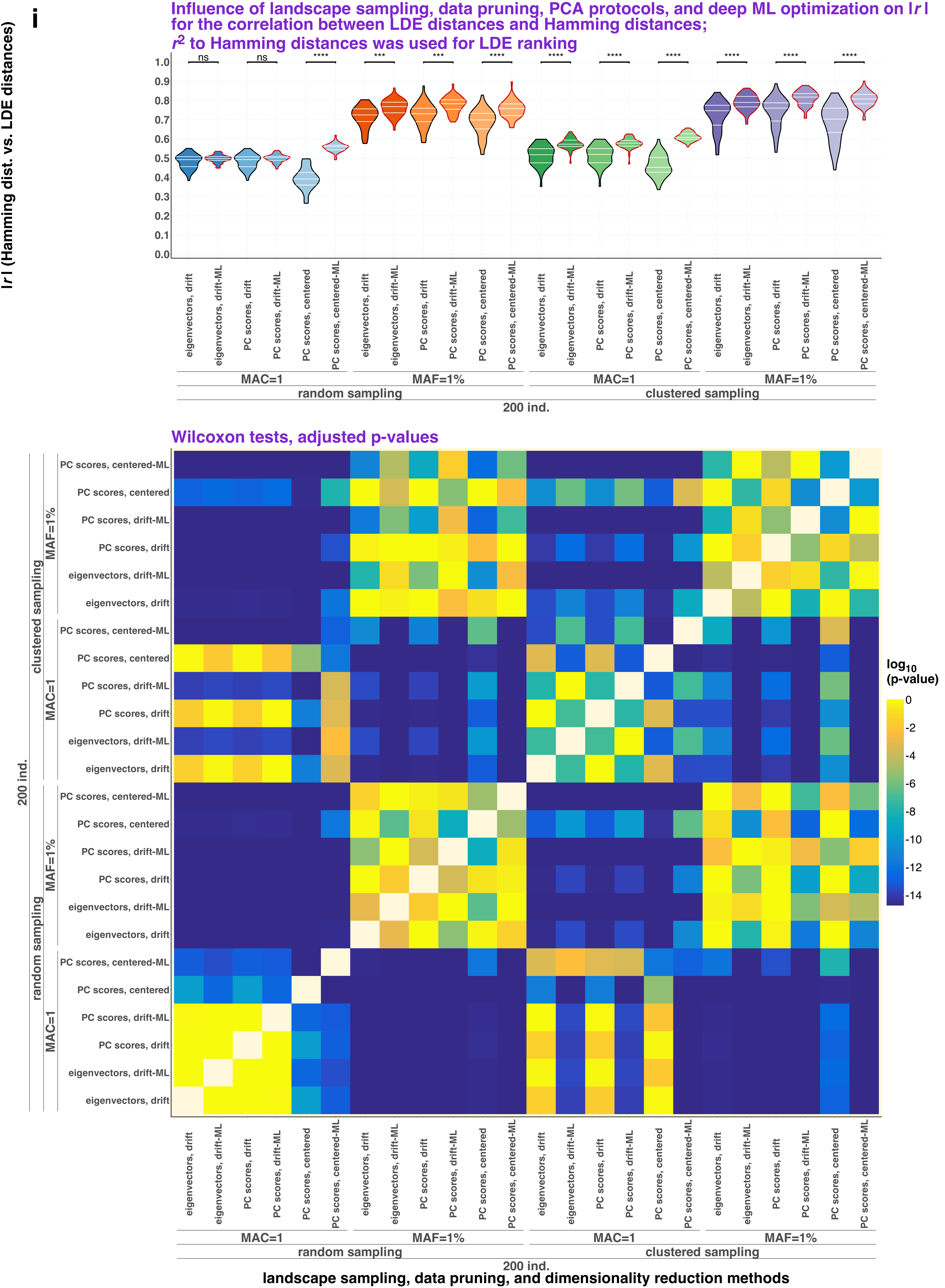
i
Influence of landscape sampling, data pruning, PCA protocols, and deep ML optimization on |r| for the correlation between LDE distances and Hamming distances;
r2 to Hamming distances was used for LDE ranking
|r| (Hamming dist. vs. LDE distances)
ns
****
***
eigenvectors, drift
eigenvectors, drift-ML
PC scores, drift
PC scores, drift-ML
PC scores, centered
PC scores, centered-ML
MAC=1
MAF=1%
random sampling
clustered sampling
200 ind.
Wilcoxon tests, adjusted p-values
log10 (p-value)
landscape sampling, data pruning, and dimensionality reduction methods

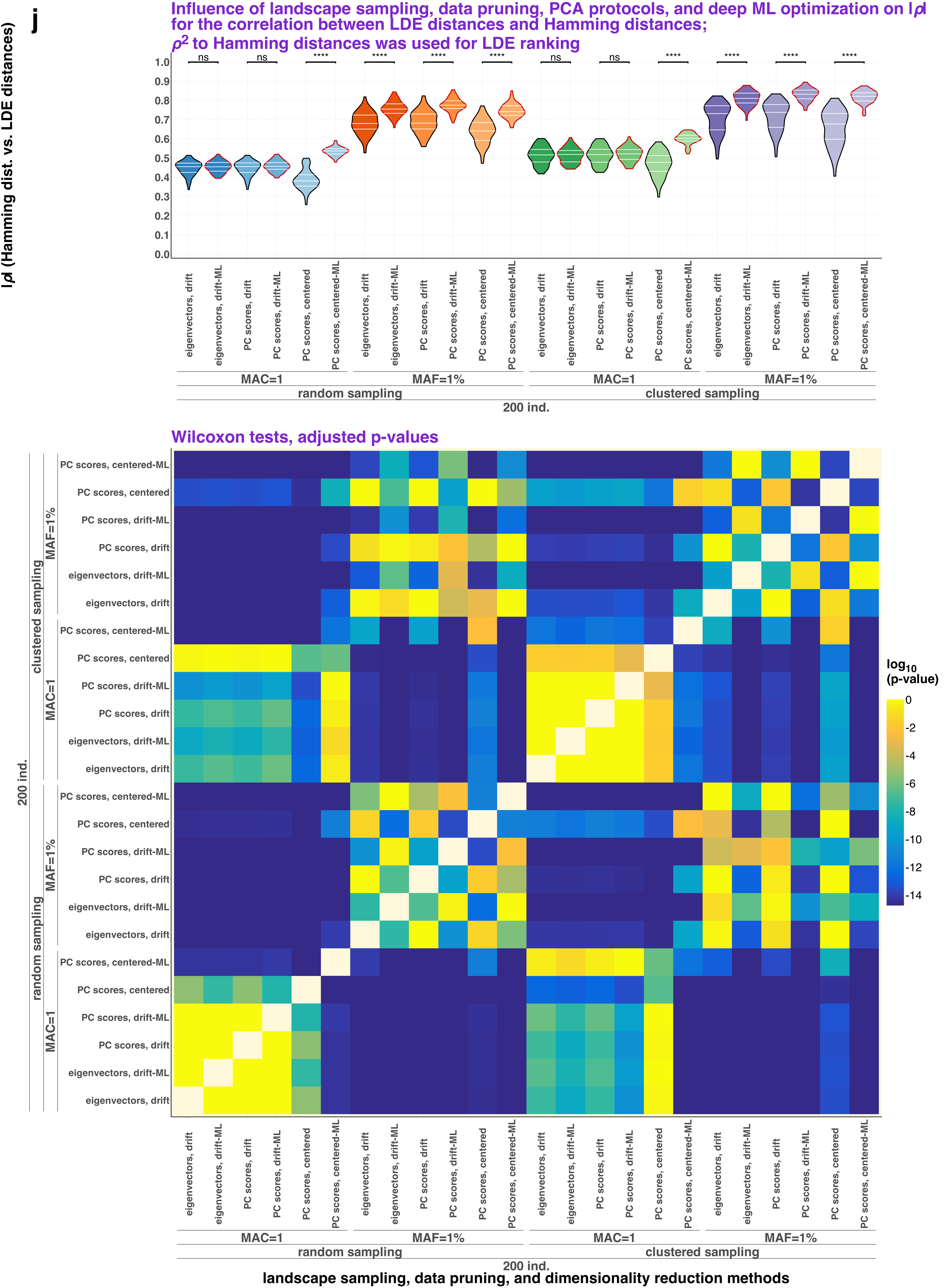

j
Influence of landscape sampling, data pruning, PCA protocols, and deep ML optimization on |ρ| for the correlation between LDE distances and Hamming distances;
ρ² to Hamming distances was used for LDE ranking
|ρ| (Hamming dist. vs. LDE distances)
ns
ns
****
****
****
****
ns
ns
****
****
****
****
1.0
0.9
0.8
0.7
0.6
0.5
0.4
0.3
0.2
0.1
0.0
eigenvectors, drift
eigenvectors, drift-ML
PC scores, drift
PC scores, drift-ML
PC scores, centered
PC scores, centered-ML
MAC=1
MAF=1%
random sampling
clustered sampling
200 ind.
Wilcoxon tests, adjusted p-values
log10 (p-value)
0
-2
-4
-6
-8
-10
-12
-14
landscape sampling, data pruning, and dimensionality reduction methods

**k**

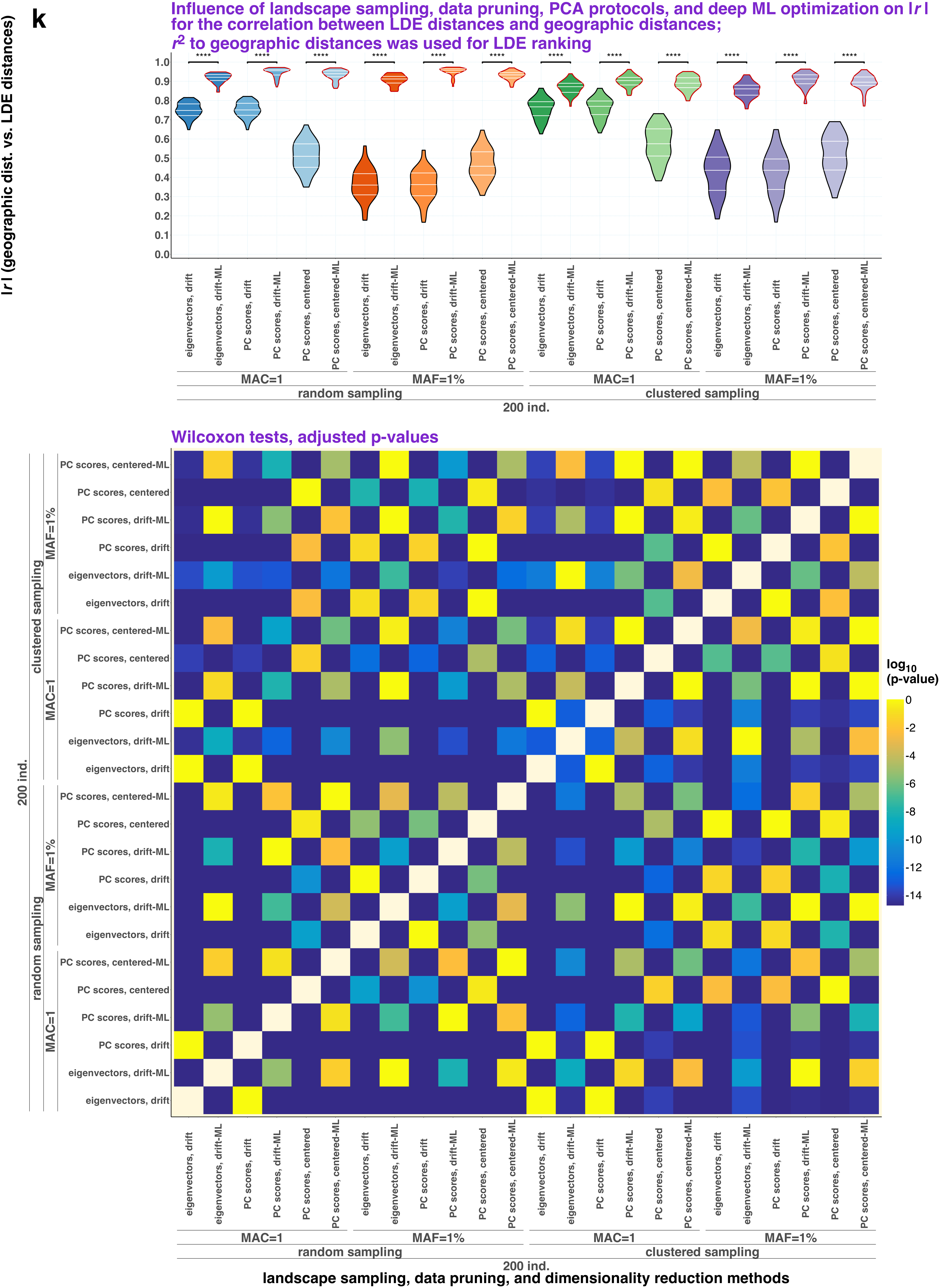
Influence of landscape sampling, data pruning, PCA protocols, and deep ML optimization on |r| for the correlation between LDE distances and geographic distances;
r2 to geographic distances was used for LDE ranking
|r| (geographic dist. vs. LDE distances)
1.0
0.9
0.8
0.7
0.6
0.5
0.4
0.3
0.2
0.1
0.0
****
eigenvectors, drift
eigenvectors, drift-ML
PC scores, drift
PC scores, drift-ML
PC scores, centered
PC scores, centered-ML
MAC=1
MAF=1%
random sampling
clustered sampling
200 ind.
Wilcoxon tests, adjusted p-values
log10 (p-value)
0
-2
-4
-6
-8
-10
-12
-14
landscape sampling, data pruning, and dimensionality reduction methods

I

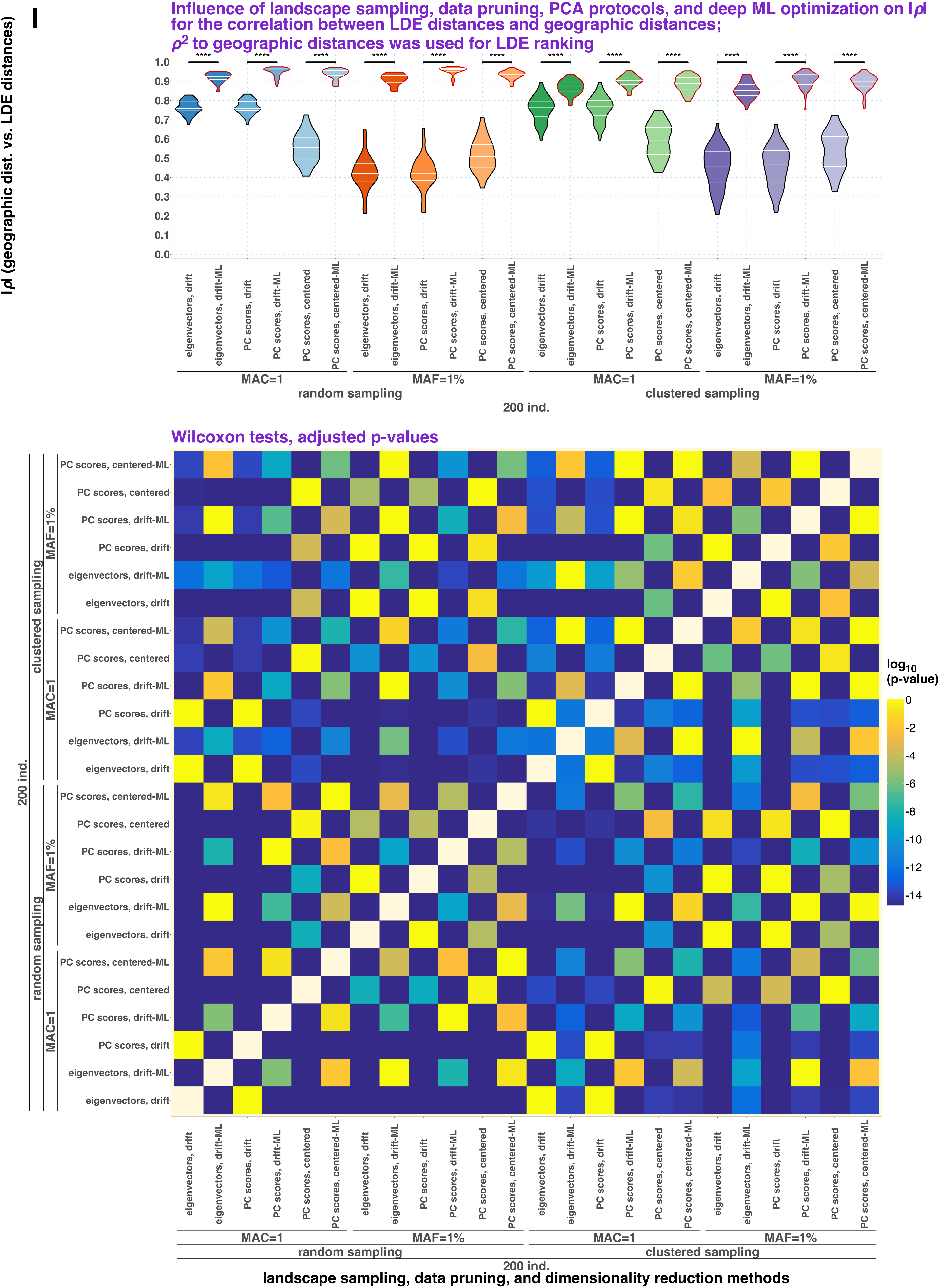
Influence of landscape sampling, data pruning, PCA protocols, and deep ML optimization on I$\rho$I for the correlation between LDE distances and geographic distances; $\rho^2$ to geographic distances was used for LDE ranking
I$\rho$I (geographic dist. vs. LDE distances)
eigenvectors, drift
eigenvectors, drift-ML
PC scores, drift
PC scores, drift-ML
PC scores, centered
PC scores, centered-ML
MAC=1
MAF=1%
random sampling
clustered sampling
200 ind.
Wilcoxon tests, adjusted p-values
log10 (p-value)
landscape sampling, data pruning, and dimensionality reduction methods
