## Supplementary Figure 13 for "Sparse sampling and rare-variant depletion distort PCA visualizations of population structure: recovery with objective-guided manifold learning"

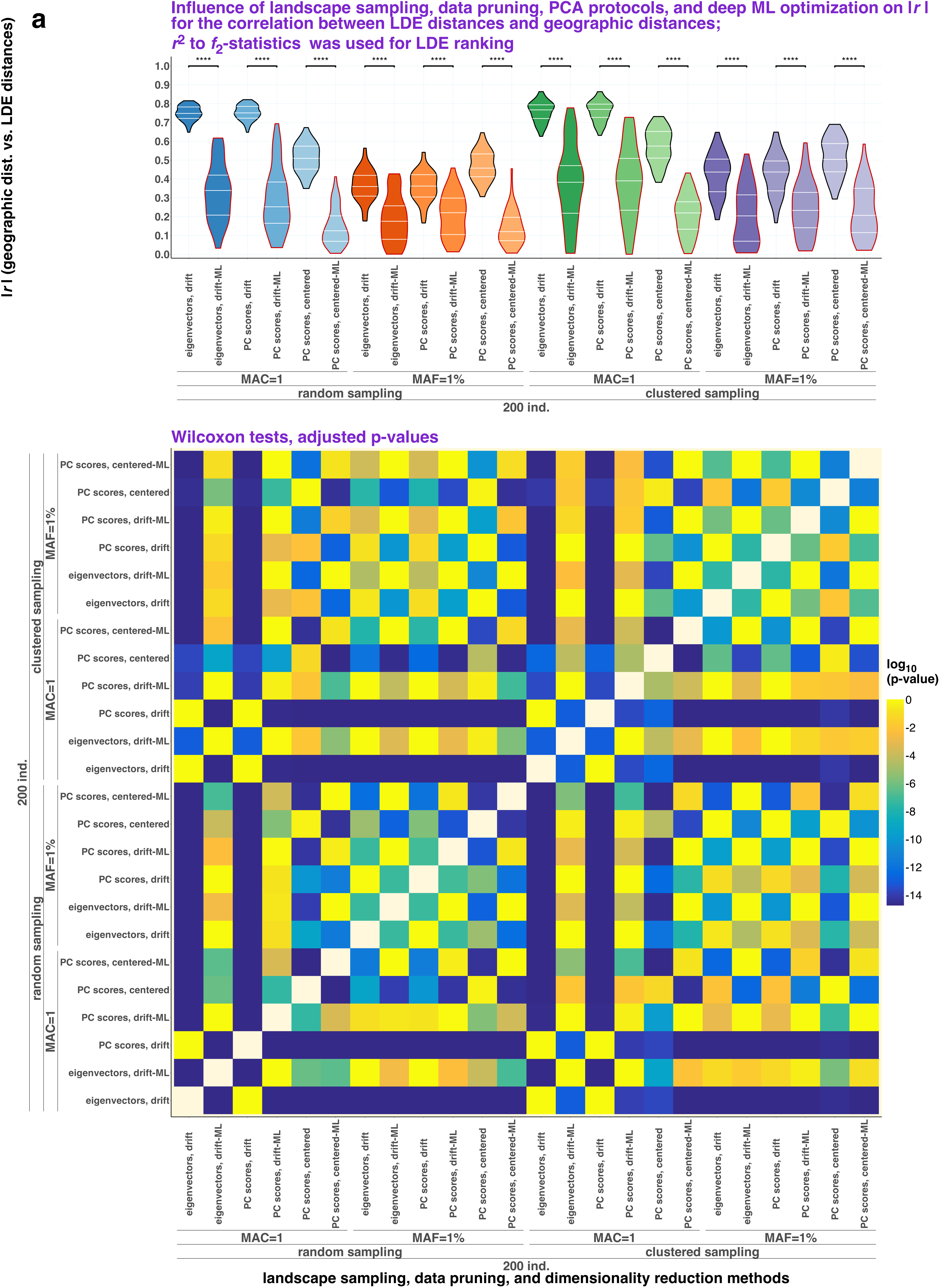
a
Influence of landscape sampling, data pruning, PCA protocols, and deep ML optimization on |r| for the correlation between LDE distances and geographic distances;
r2 to f2-statistics was used for LDE ranking
|r| (geographic dist. vs. LDE distances)
****
1.0
0.9
0.8
0.7
0.6
0.5
0.4
0.3
0.2
0.1
0.0
eigenvectors, drift
eigenvectors, drift-ML
PC scores, drift
PC scores, drift-ML
PC scores, centered
PC scores, centered-ML
MAC=1
MAF=1%
random sampling
clustered sampling
200 ind.
Wilcoxon tests, adjusted p-values
log10 (p-value)
0
-2
-4
-6
-8
-10
-12
-14
landscape sampling, data pruning, and dimensionality reduction methods

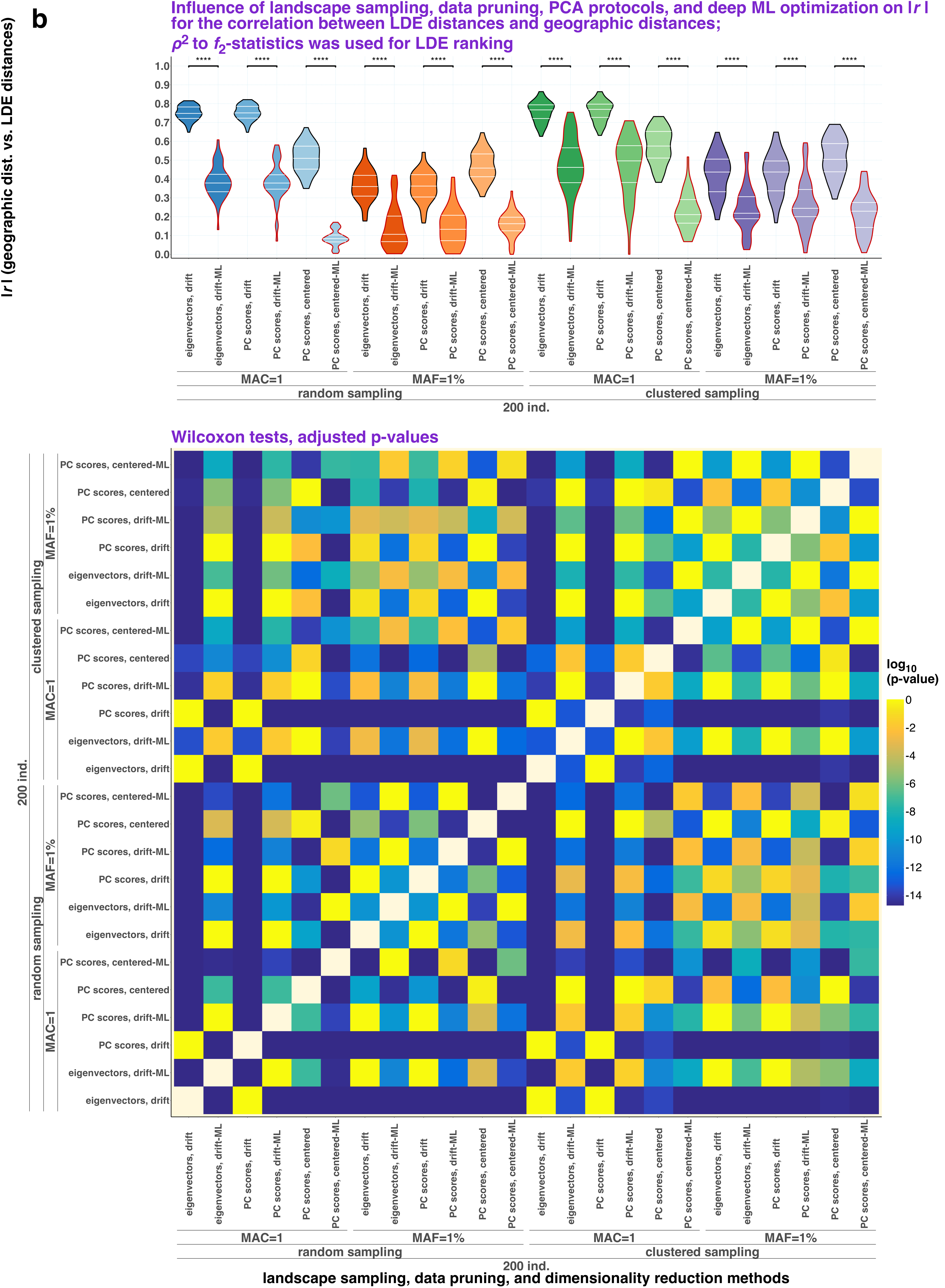
b
Influence of landscape sampling, data pruning, PCA protocols, and deep ML optimization on |r| for the correlation between LDE distances and geographic distances;
$\rho^2$ to $f_2$-statistics was used for LDE ranking
|r| (geographic dist. vs. LDE distances)
****
1.0
0.9
0.8
0.7
0.6
0.5
0.4
0.3
0.2
0.1
0.0
eigenvectors, drift
eigenvectors, drift-ML
PC scores, drift
PC scores, drift-ML
PC scores, centered
PC scores, centered-ML
MAC=1
MAF=1%
random sampling
clustered sampling
200 ind.
Wilcoxon tests, adjusted p-values
log10 (p-value)
0
-2
-4
-6
-8
-10
-12
-14
landscape sampling, data pruning, and dimensionality reduction methods

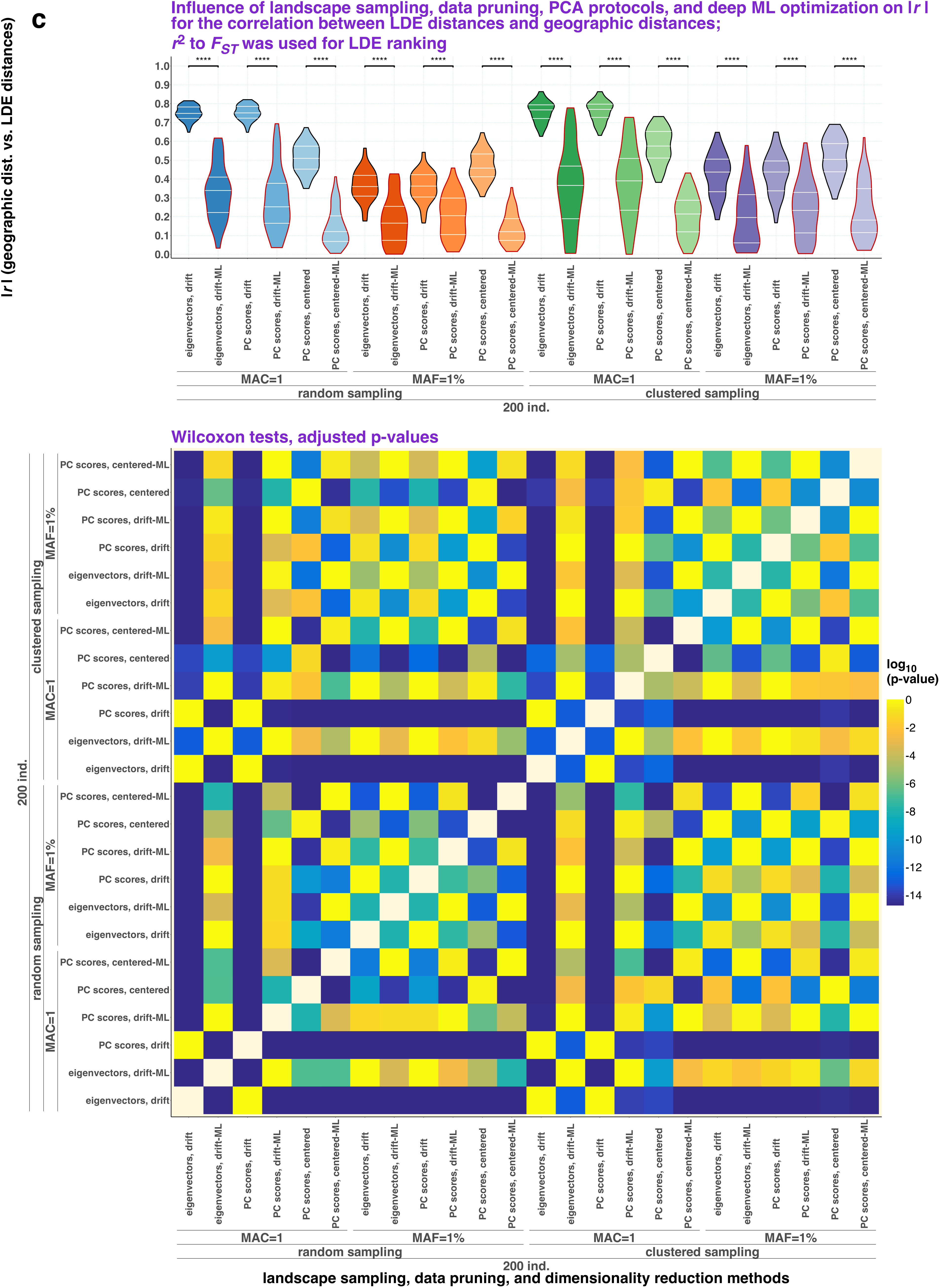

c
Influence of landscape sampling, data pruning, PCA protocols, and deep ML optimization on |r| for the correlation between LDE distances and geographic distances; $r^2$ to $F_{ST}$ was used for LDE ranking
|r| (geographic dist. vs. LDE distances)
eigenvectors, drift
eigenvectors, drift-ML
PC scores, drift
PC scores, drift-ML
PC scores, centered
PC scores, centered-ML
MAC=1
MAF=1%
random sampling
clustered sampling
200 ind.
Wilcoxon tests, adjusted p-values
log10 (p-value)
landscape sampling, data pruning, and dimensionality reduction methods

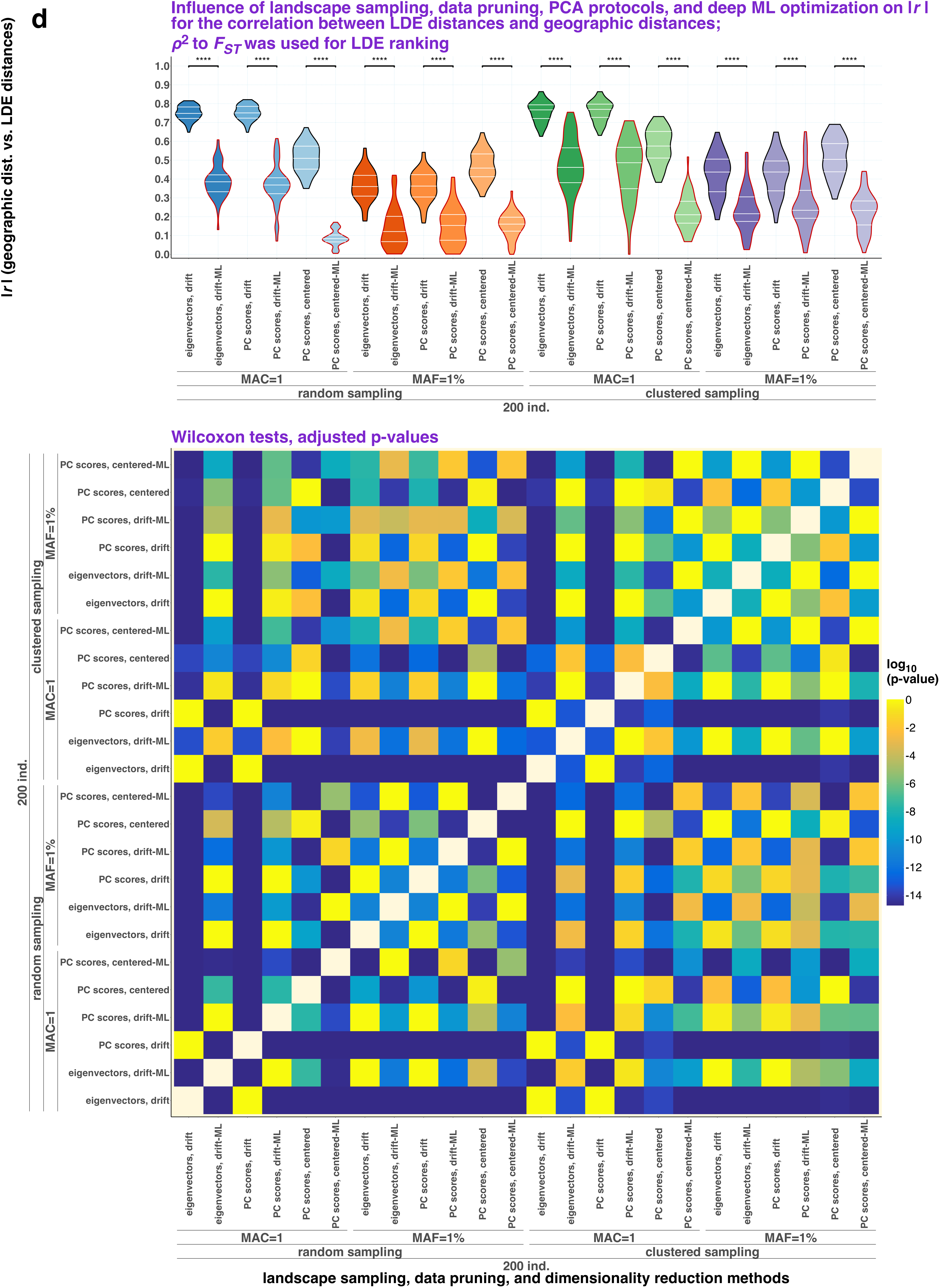

d
Influence of landscape sampling, data pruning, PCA protocols, and deep ML optimization on |r| for the correlation between LDE distances and geographic distances; ρ2 to FST was used for LDE ranking
|r| (geographic dist. vs. LDE distances)
eigenvectors, drift
eigenvectors, drift-ML
PC scores, drift
PC scores, drift-ML
PC scores, centered
PC scores, centered-ML
MAC=1
MAF=1%
random sampling
clustered sampling
200 ind.
Wilcoxon tests, adjusted p-values
log10 (p-value)
landscape sampling, data pruning, and dimensionality reduction methods

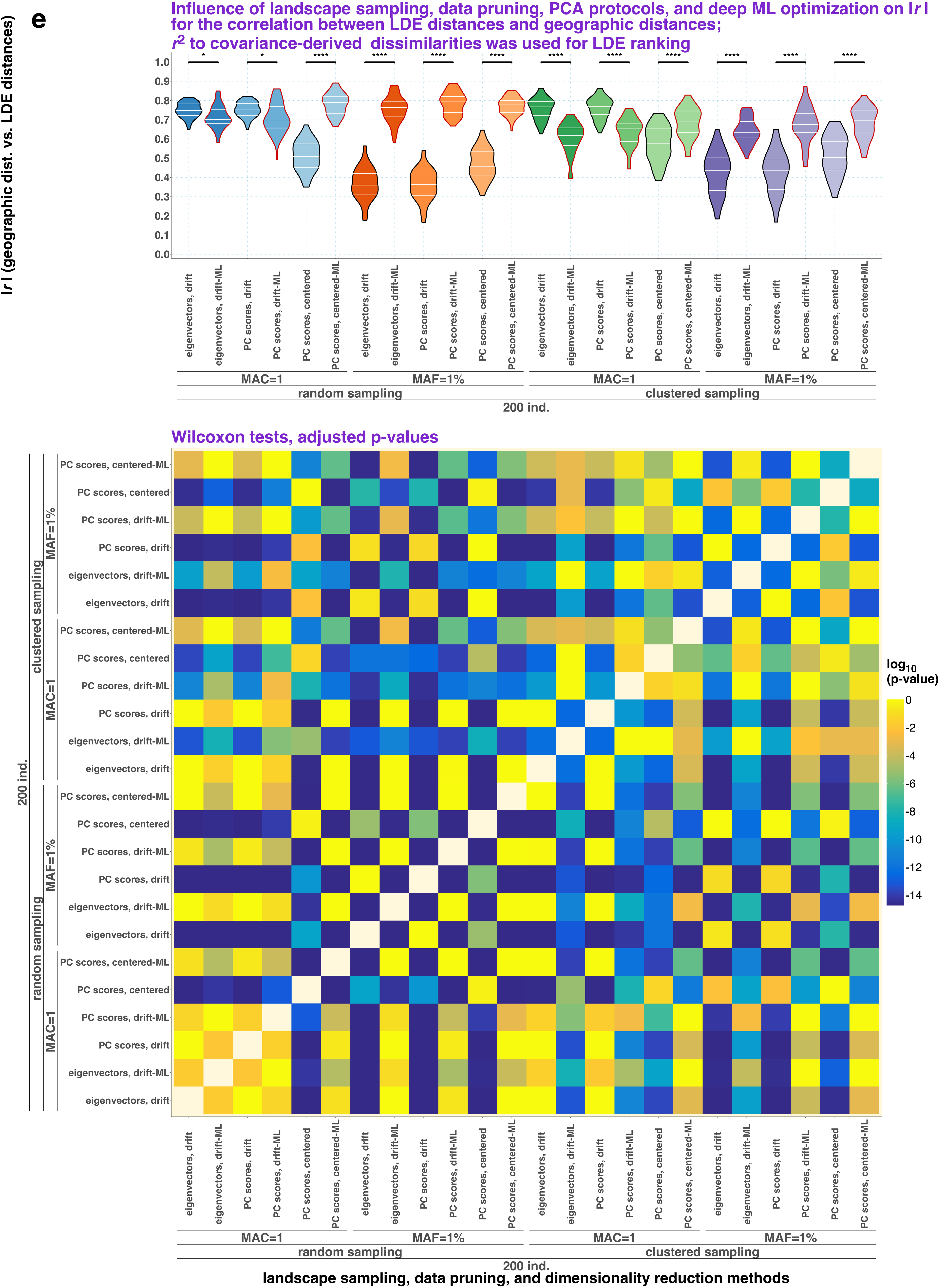
e
Influence of landscape sampling, data pruning, PCA protocols, and deep ML optimization on |r|
for the correlation between LDE distances and geographic distances;
r2 to covariance-derived dissimilarities was used for LDE ranking
|r| (geographic dist. vs. LDE distances)
*
*
****
****
****
****
****
****
****
****
****
****
eigenvectors, drift
eigenvectors, drift-ML
PC scores, drift
PC scores, drift-ML
PC scores, centered
PC scores, centered-ML
MAC=1
MAF=1%
random sampling
clustered sampling
200 ind.
Wilcoxon tests, adjusted p-values
log10 (p-value)
landscape sampling, data pruning, and dimensionality reduction methods

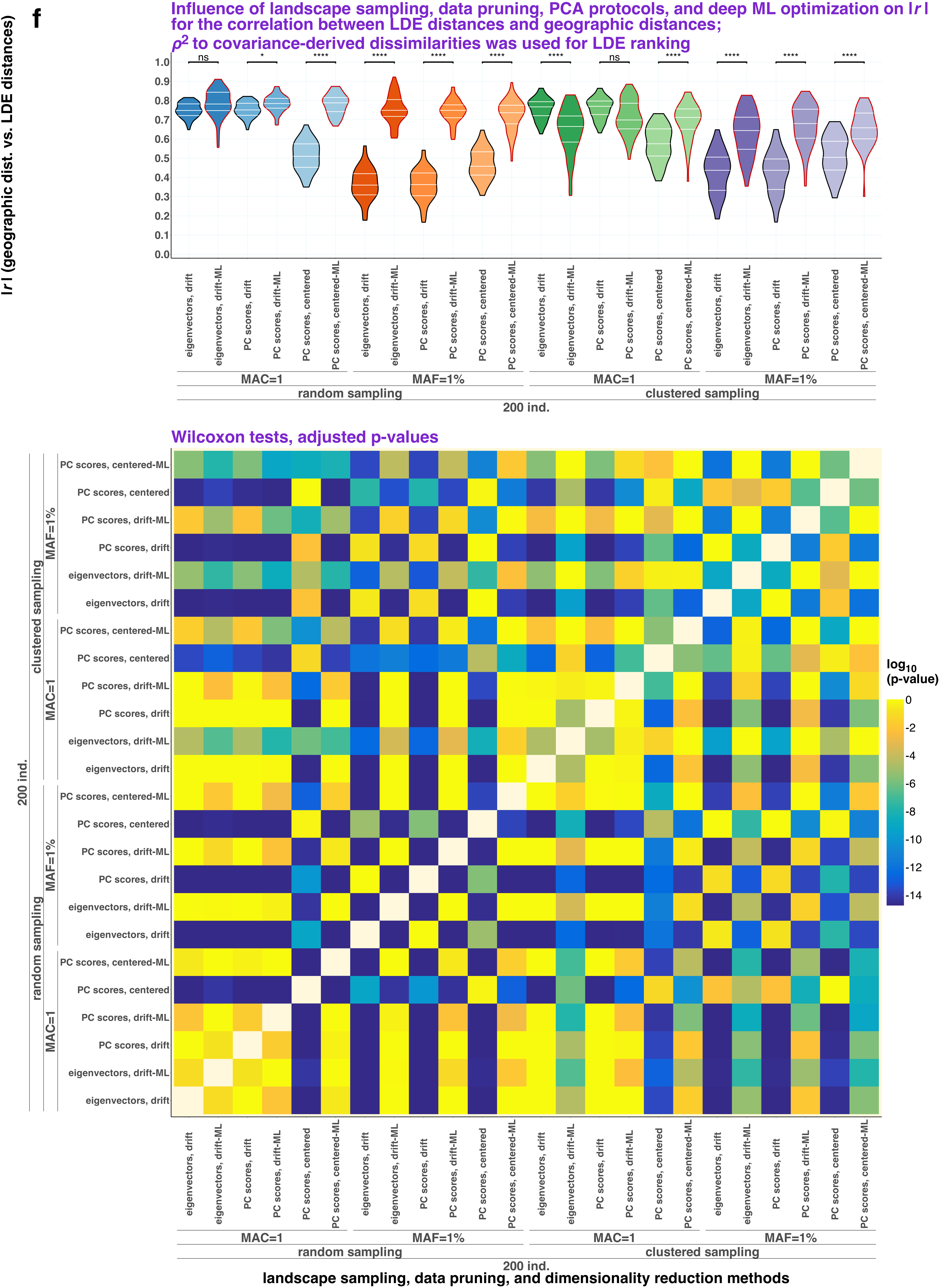
f
Influence of landscape sampling, data pruning, PCA protocols, and deep ML optimization on |r| for the correlation between LDE distances and geographic distances;
ρ² to covariance-derived dissimilarities was used for LDE ranking
|r| (geographic dist. vs. LDE distances)
ns
*
****
ns
1.0
0.9
0.8
0.7
0.6
0.5
0.4
0.3
0.2
0.1
0.0
eigenvectors, drift
eigenvectors, drift-ML
PC scores, drift
PC scores, drift-ML
PC scores, centered
PC scores, centered-ML
MAC=1
MAF=1%
random sampling
clustered sampling
200 ind.
Wilcoxon tests, adjusted p-values
log10 (p-value)
0
-2
-4
-6
-8
-10
-12
-14
landscape sampling, data pruning, and dimensionality reduction methods

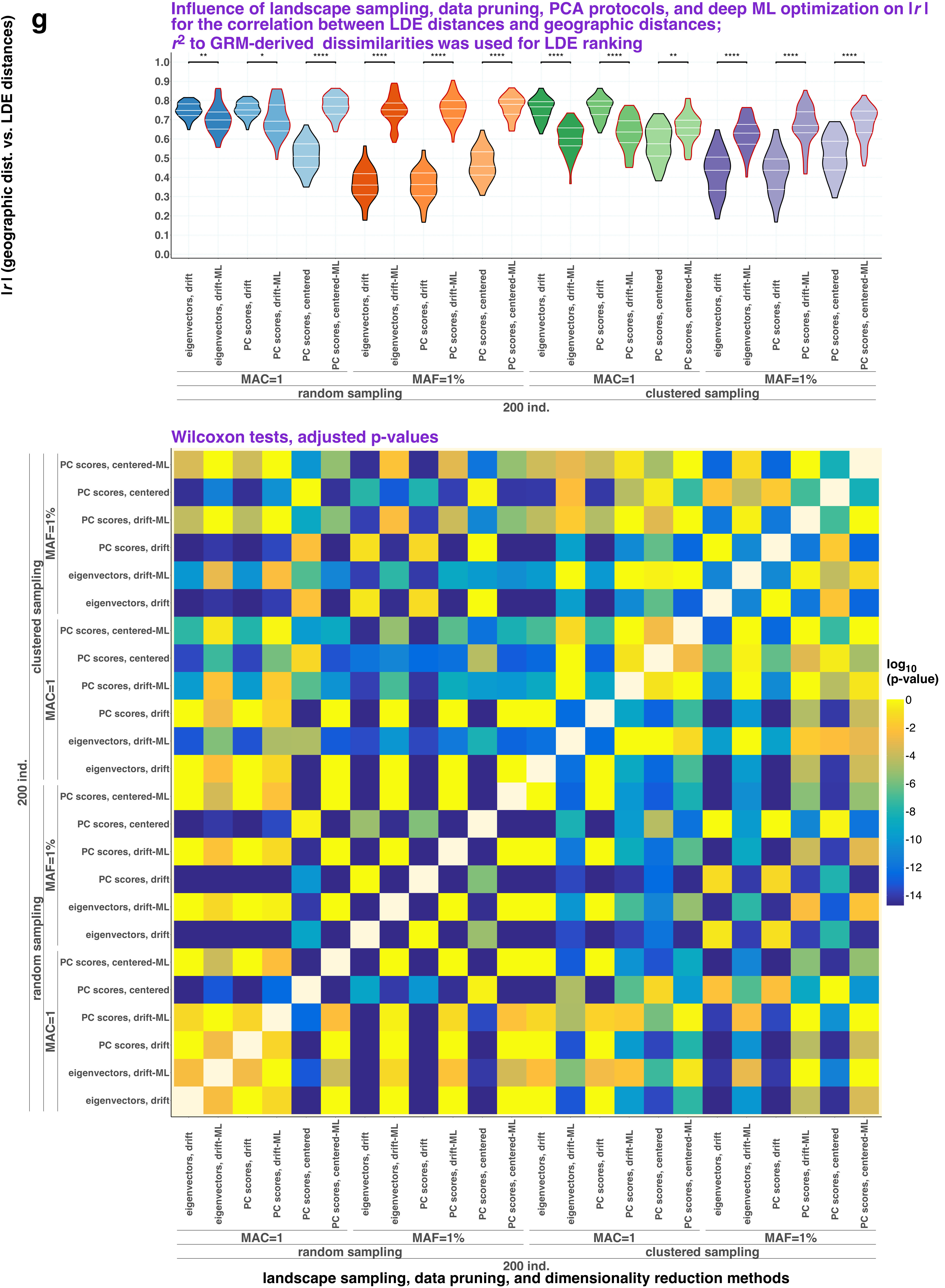

g
Influence of landscape sampling, data pruning, PCA protocols, and deep ML optimization on |r| for the correlation between LDE distances and geographic distances;
r² to GRM-derived dissimilarities was used for LDE ranking
|r| (geographic dist. vs. LDE distances)
eigenvectors, drift
eigenvectors, drift-ML
PC scores, drift
PC scores, drift-ML
PC scores, centered
PC scores, centered-ML
MAC=1
MAF=1%
random sampling
clustered sampling
200 ind.
Wilcoxon tests, adjusted p-values
log10 (p-value)
landscape sampling, data pruning, and dimensionality reduction methods

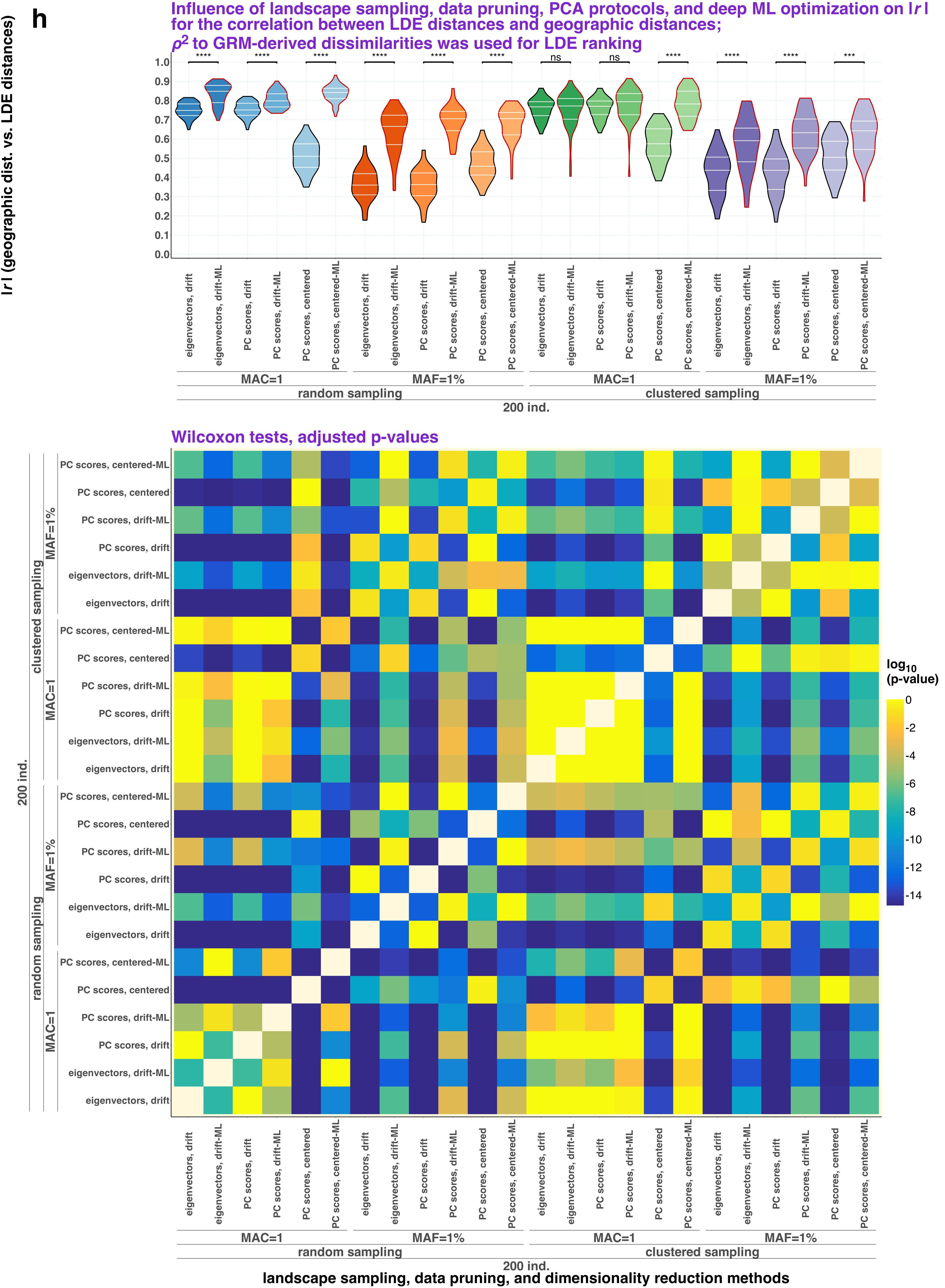
h
Influence of landscape sampling, data pruning, PCA protocols, and deep ML optimization on |r| for the correlation between LDE distances and geographic distances;
ρ2 to GRM-derived dissimilarities was used for LDE ranking
|r| (geographic dist. vs. LDE distances)
eigenvectors, drift
eigenvectors, drift-ML
PC scores, drift
PC scores, drift-ML
PC scores, centered
PC scores, centered-ML
MAC=1
MAF=1%
random sampling
clustered sampling
200 ind.
Wilcoxon tests, adjusted p-values
log10 (p-value)
landscape sampling, data pruning, and dimensionality reduction methods

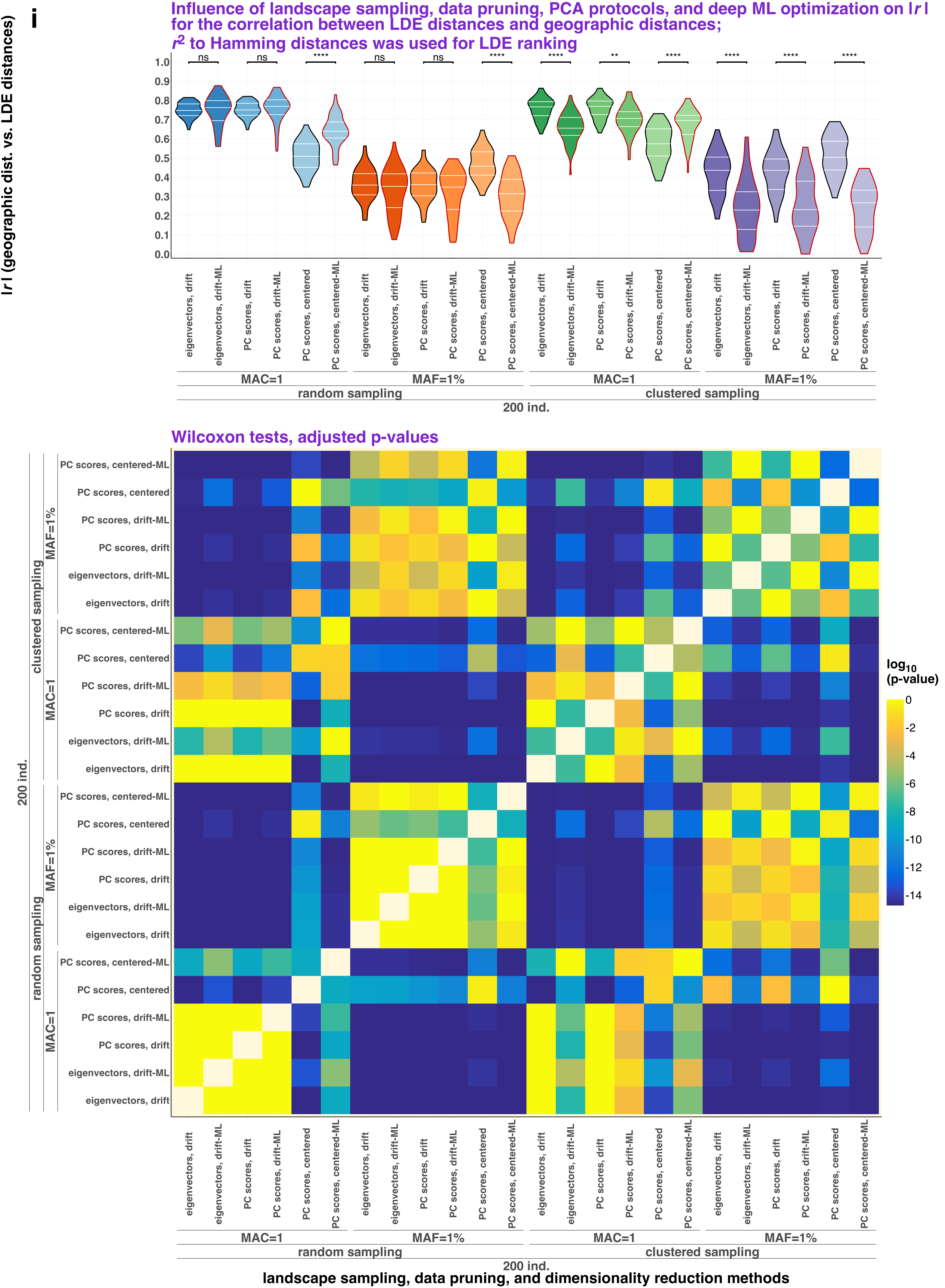

i
Influence of landscape sampling, data pruning, PCA protocols, and deep ML optimization on |r| for the correlation between LDE distances and geographic distances; $r^2$ to Hamming distances was used for LDE ranking
|r| (geographic dist. vs. LDE distances)
ns ns **** ns ns **** **** ** **** **** **** ****
eigenvectors, drift
eigenvectors, drift-ML
PC scores, drift
PC scores, drift-ML
PC scores, centered
PC scores, centered-ML
MAC=1
MAF=1%
random sampling
clustered sampling
200 ind.
Wilcoxon tests, adjusted p-values
log10 (p-value)
0 -2 -4 -6 -8 -10 -12 -14
landscape sampling, data pruning, and dimensionality reduction methods

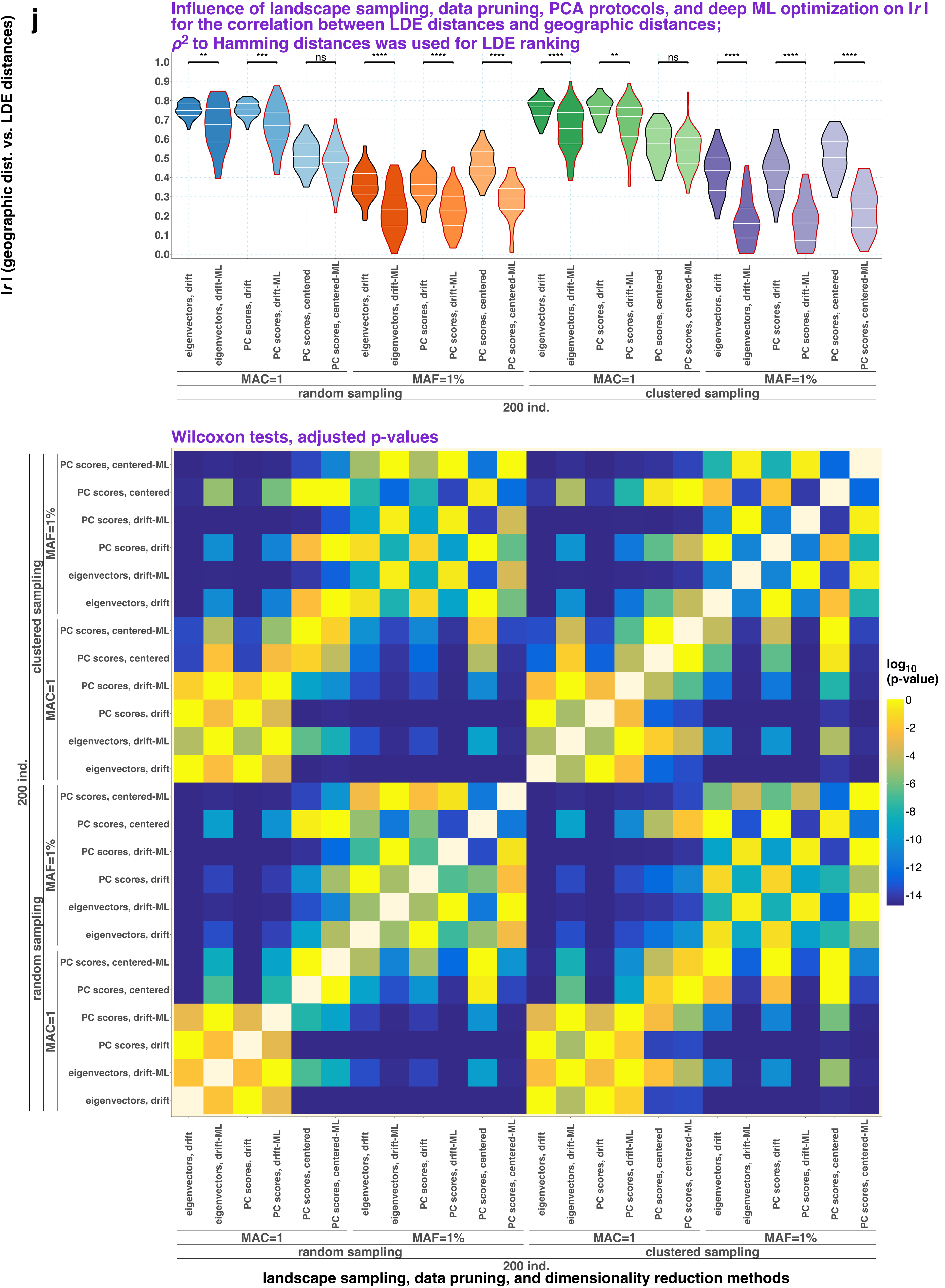
j
Influence of landscape sampling, data pruning, PCA protocols, and deep ML optimization on |r|
for the correlation between LDE distances and geographic distances;
ρ2 to Hamming distances was used for LDE ranking
|r| (geographic dist. vs. LDE distances)
** *** ns **** **** **** **** ** ns **** **** ****
1.0 0.9 0.8 0.7 0.6 0.5 0.4 0.3 0.2 0.1 0.0
eigenvectors, drift
eigenvectors, drift-ML
PC scores, drift
PC scores, drift-ML
PC scores, centered
PC scores, centered-ML
MAC=1
MAF=1%
random sampling
clustered sampling
200 ind.
Wilcoxon tests, adjusted p-values
log10 (p-value)
0 -2 -4 -6 -8 -10 -12 -14
landscape sampling, data pruning, and dimensionality reduction methods

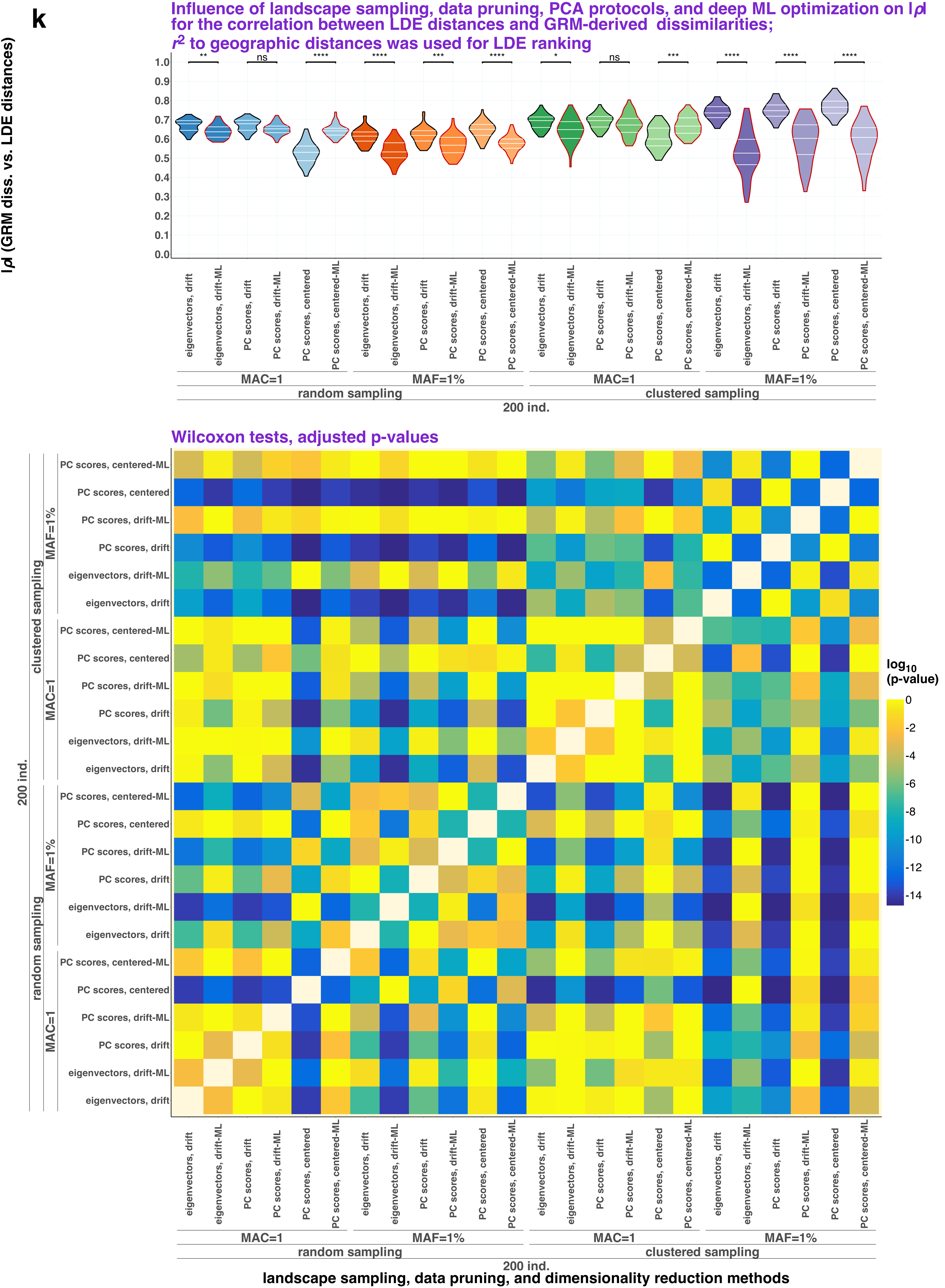
k
Influence of landscape sampling, data pruning, PCA protocols, and deep ML optimization on |ρ| for the correlation between LDE distances and GRM-derived dissimilarities; r2 to geographic distances was used for LDE ranking
|ρ| (GRM diss. vs. LDE distances)
eigenvectors, drift
eigenvectors, drift-ML
PC scores, drift
PC scores, drift-ML
PC scores, centered
PC scores, centered-ML
MAC=1
MAF=1%
random sampling
clustered sampling
200 ind.
Wilcoxon tests, adjusted p-values
log10 (p-value)
landscape sampling, data pruning, and dimensionality reduction methods

I

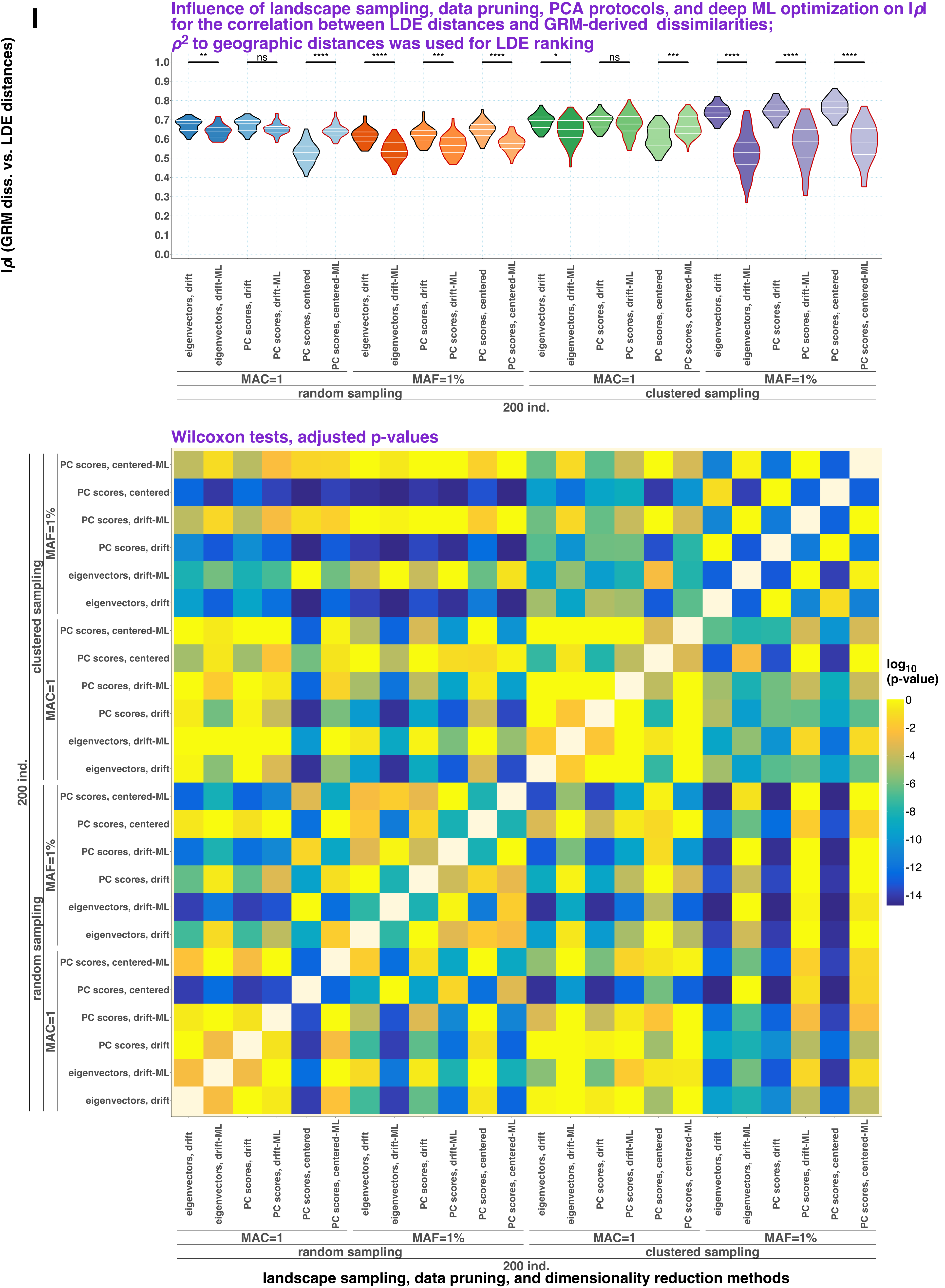
Influence of landscape sampling, data pruning, PCA protocols, and deep ML optimization on |ρ| for the correlation between LDE distances and GRM-derived dissimilarities; ρ2 to geographic distances was used for LDE ranking
|ρ| (GRM diss. vs. LDE distances)
1.0
0.9
0.8
0.7
0.6
0.5
0.4
0.3
0.2
0.1
0.0
**
ns
****
****
***
****
*
ns
***
****
****
****
eigenvectors, drift
eigenvectors, drift-ML
PC scores, drift
PC scores, drift-ML
PC scores, centered
PC scores, centered-ML
MAC=1
MAF=1%
random sampling
clustered sampling
200 ind.
Wilcoxon tests, adjusted p-values
log10 (p-value)
0
-2
-4
-6
-8
-10
-12
-14
landscape sampling, data pruning, and dimensionality reduction methods
