## Supplementary Figure 15 for "Sparse sampling and rare-variant depletion distort PCA visualizations of population structure: recovery with objective-guided manifold learning"

**a**

**Top 100 LDEs per dataset/MAF level selected according to $r^2$ of LDE dist. vs. $f_2$-statistics and stratified by: sampling protocol, PC space dimensionality, MAF filtering, PCA protocol, distance metric in PC space, & ML algorithm**

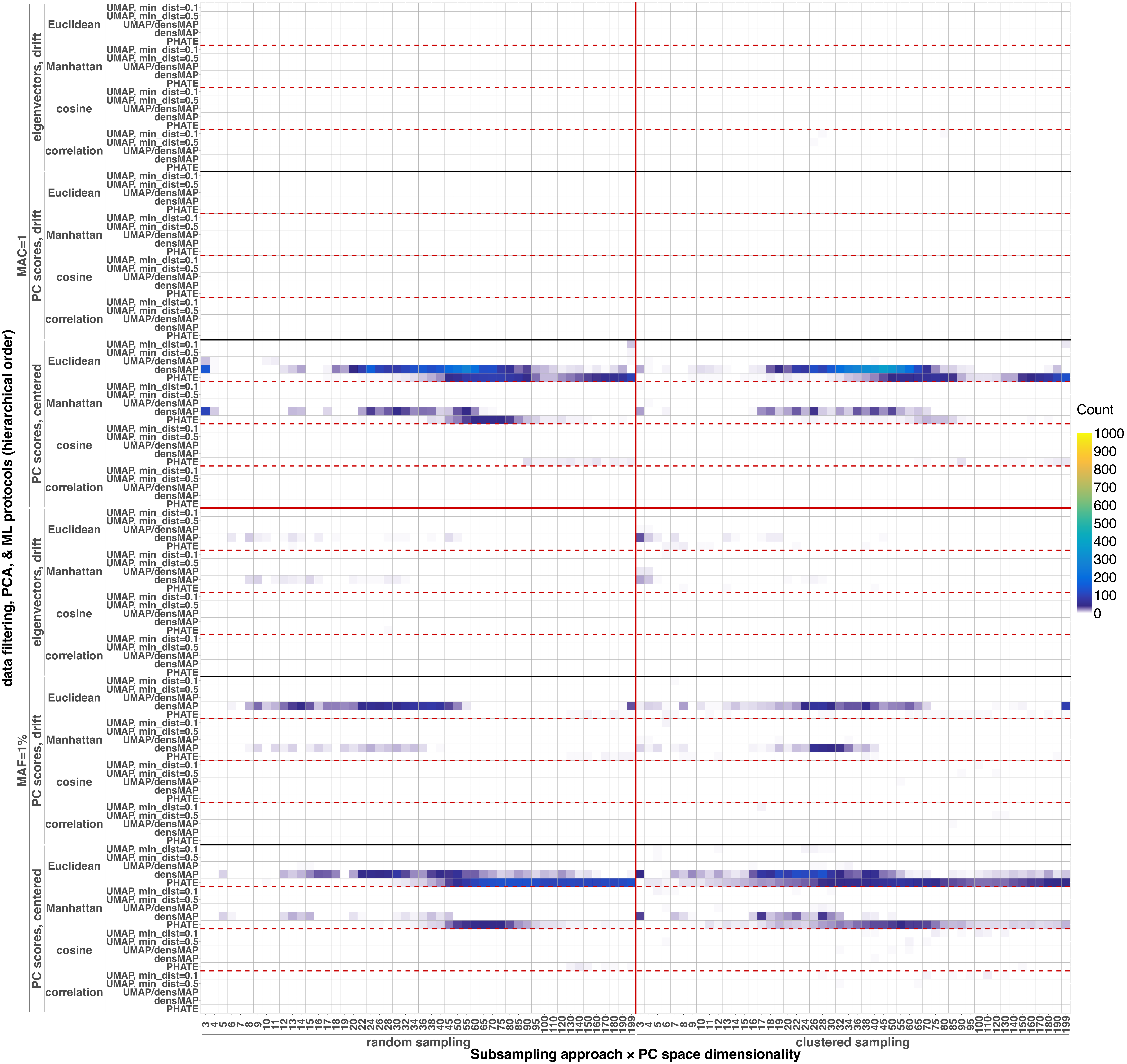

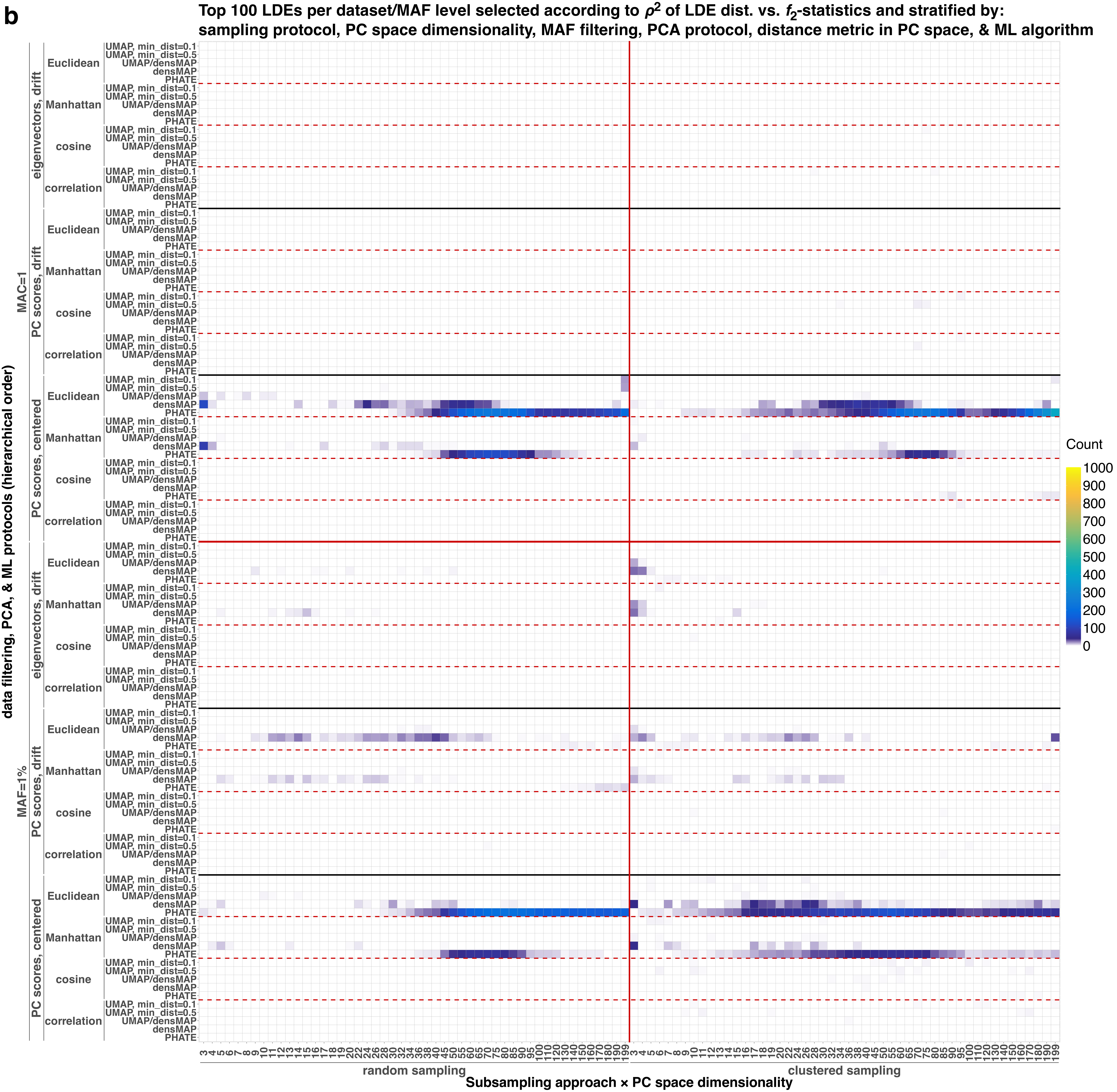

b
Top 100 LDEs per dataset/MAF level selected according to $\rho^2$ of LDE dist. vs. $f_2$-statistics and stratified by:
sampling protocol, PC space dimensionality, MAF filtering, PCA protocol, distance metric in PC space, & ML algorithm
data filtering, PCA, & ML protocols (hierarchical order)
MAC=1
MAF=1%
eigenvectors, drift
PC scores, drift
PC scores, centered
Euclidean
Manhattan
cosine
correlation
UMAP, min_dist=0.1
UMAP, min_dist=0.5
UMAP/densMAP
densMAP
PHATE
Count
1000
900
800
700
600
500
400
300
200
100
0
3 4 5 6 7 8 9 10 11 12 13 14 15 16 17 18 19 20 22 24 26 28 30 32 34 36 38 40 45 50 55 60 65 70 75 80 85 90 95 100 110 120 130 140 150 160 170 180 190 199
random sampling
clustered sampling
Subsampling approach × PC space dimensionality

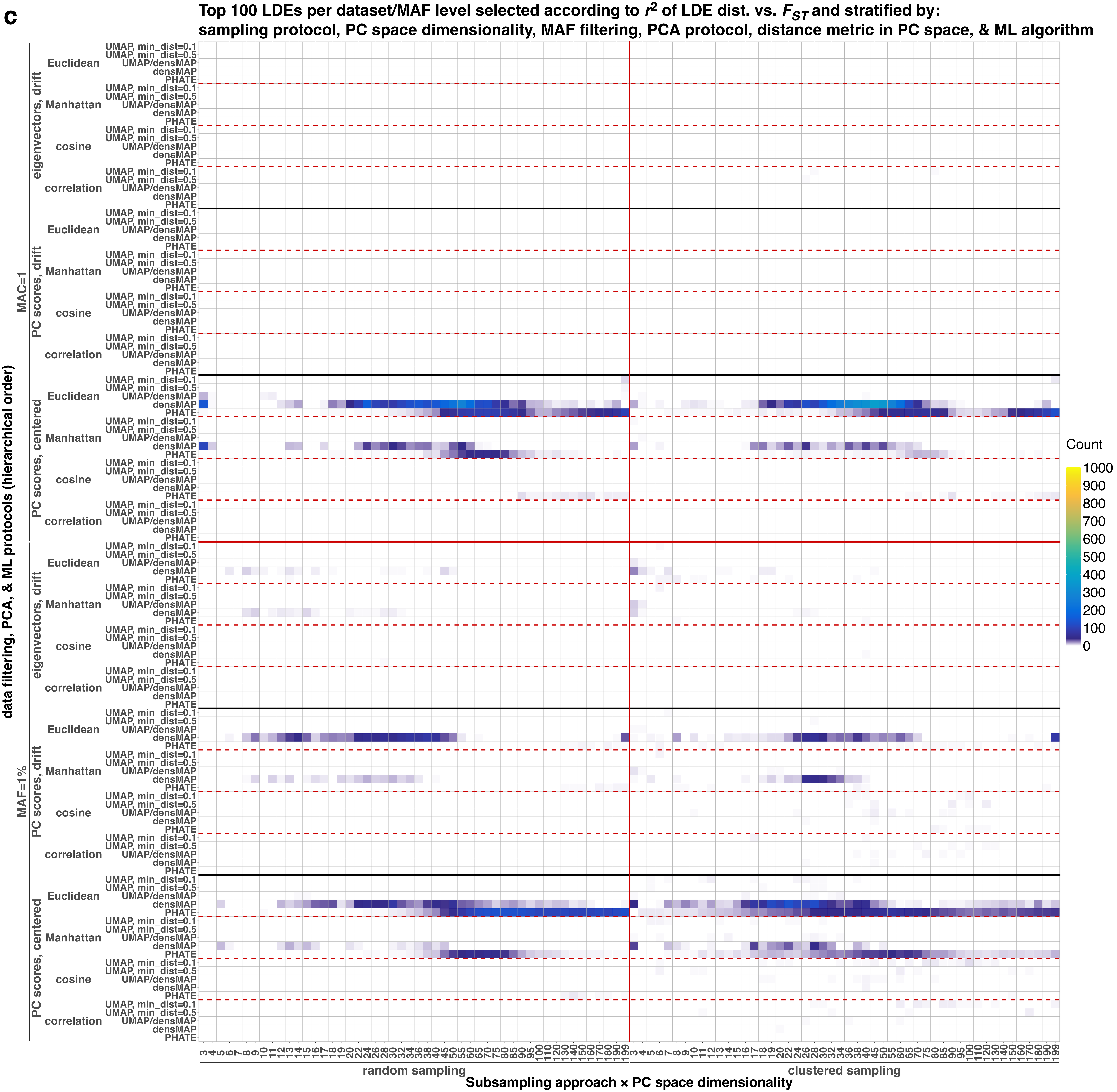
c
Top 100 LDEs per dataset/MAF level selected according to $r^2$ of LDE dist. vs. $F_{ST}$ and stratified by:
sampling protocol, PC space dimensionality, MAF filtering, PCA protocol, distance metric in PC space, & ML algorithm
data filtering, PCA, & ML protocols (hierarchical order)
MAC=1
MAF=1%
eigenvectors, drift
PC scores, drift
PC scores, centered
Euclidean
Manhattan
cosine
correlation
UMAP, min_dist=0.1
UMAP, min_dist=0.5
UMAP/densMAP
densMAP
PHATE
Count
1000
900
800
700
600
500
400
300
200
100
0
random sampling
clustered sampling
Subsampling approach × PC space dimensionality

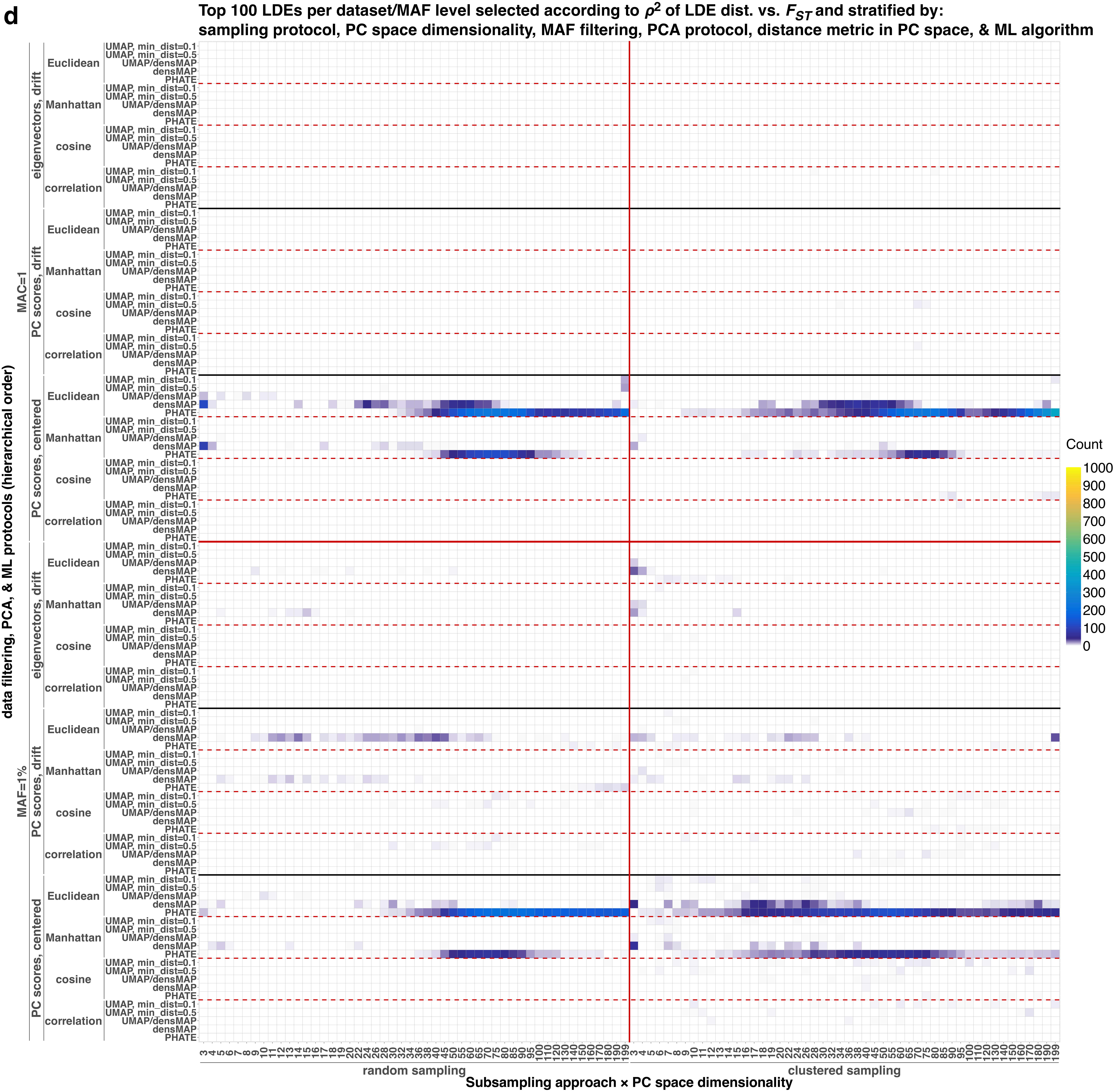

d
Top 100 LDEs per dataset/MAF level selected according to ρ2 of LDE dist. vs. FST and stratified by:
sampling protocol, PC space dimensionality, MAF filtering, PCA protocol, distance metric in PC space, & ML algorithm
data filtering, PCA, & ML protocols (hierarchical order)
MAC=1
MAF=1%
eigenvectors, drift
PC scores, drift
PC scores, centered
Euclidean
Manhattan
cosine
correlation
UMAP, min_dist=0.1
UMAP, min_dist=0.5
UMAP/densMAP
densMAP
PHATE
Count
1000
900
800
700
600
500
400
300
200
100
0
random sampling
clustered sampling
Subsampling approach × PC space dimensionality

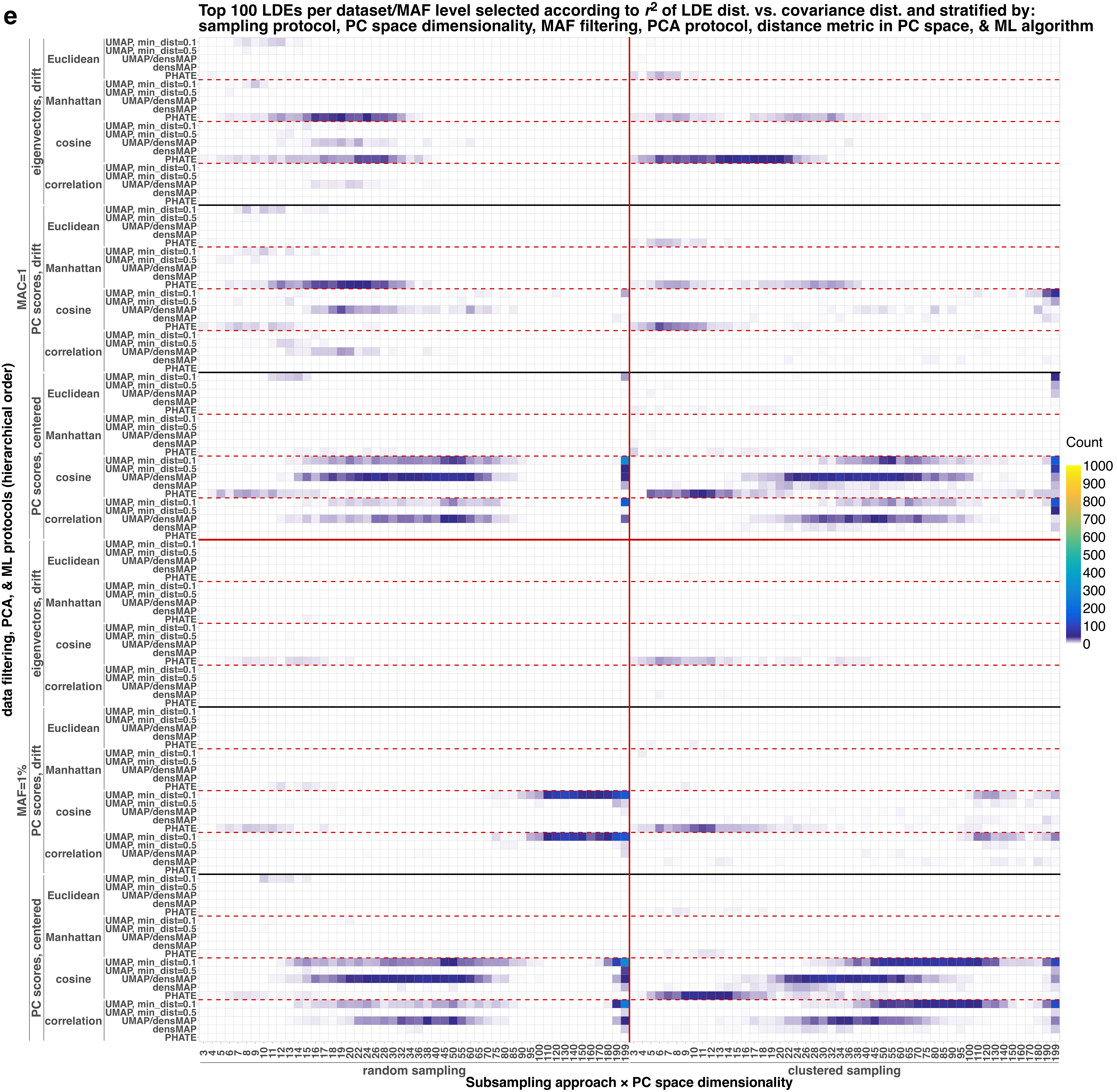
e
Top 100 LDEs per dataset/MAF level selected according to $r^2$ of LDE dist. vs. covariance dist. and stratified by: sampling protocol, PC space dimensionality, MAF filtering, PCA protocol, distance metric in PC space, & ML algorithm
data filtering, PCA, & ML protocols (hierarchical order)
MAC=1
MAF=1%
eigenvectors, drift
PC scores, drift
PC scores, centered
Euclidean
Manhattan
cosine
correlation
UMAP, min_dist=0.1
UMAP, min_dist=0.5
UMAP/densMAP
densMAP
PHATE
Count
1000
900
800
700
600
500
400
300
200
100
0
3 4 5 6 7 8 9 10 11 12 13 14 15 16 17 18 19 20 22 24 26 28 30 32 34 36 38 40 45 50 55 60 65 70 75 80 85 90 95 100 110 120 130 140 150 160 170 180 190 199
random sampling
clustered sampling
Subsampling approach × PC space dimensionality

f

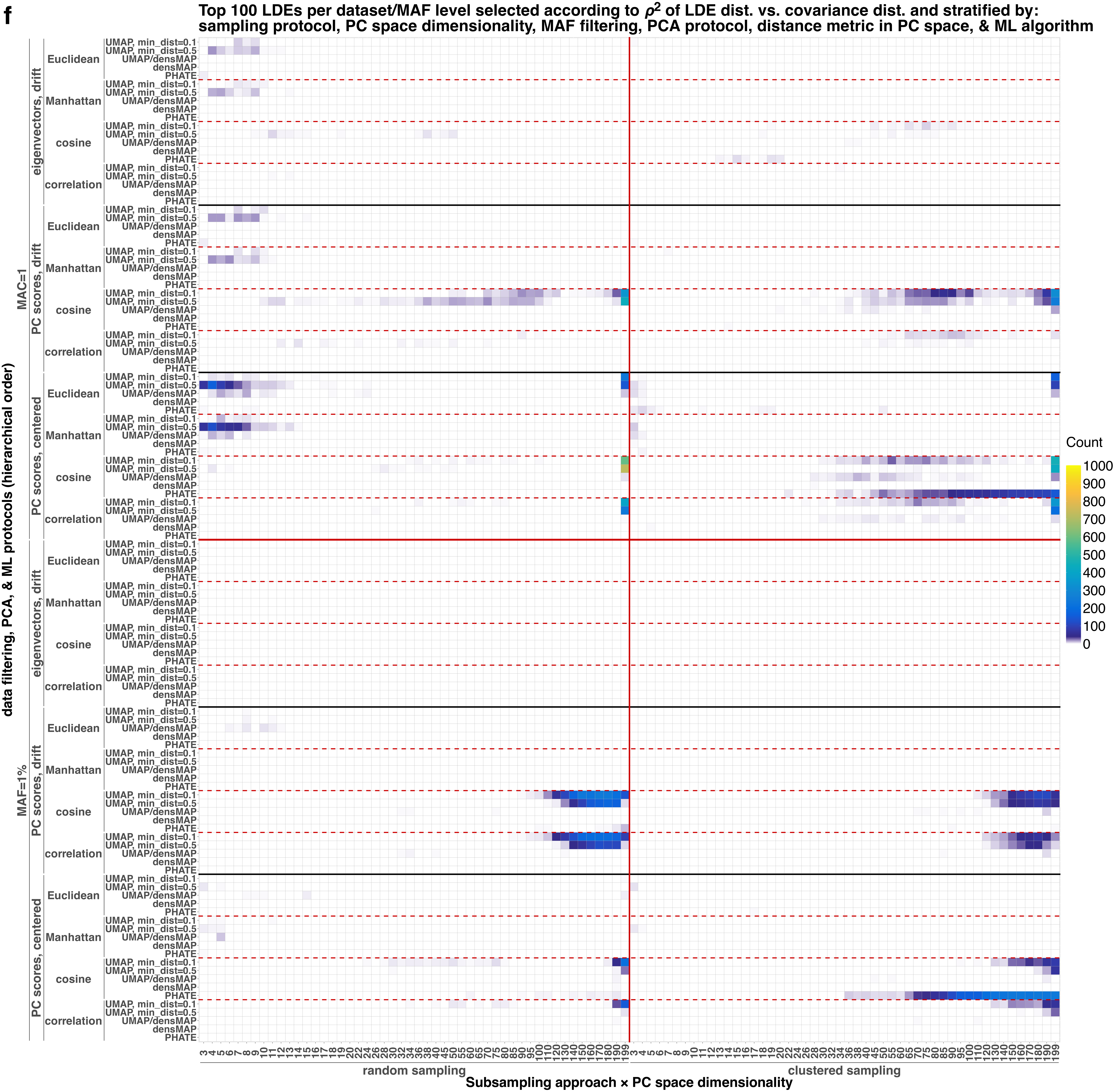

Top 100 LDEs per dataset/MAF level selected according to ρ2 of LDE dist. vs. covariance dist. and stratified by:
sampling protocol, PC space dimensionality, MAF filtering, PCA protocol, distance metric in PC space, & ML algorithm
data filtering, PCA, & ML protocols (hierarchical order)
MAC=1
MAF=1%
eigenvectors, drift
PC scores, drift
PC scores, centered
Euclidean
Manhattan
cosine
correlation
UMAP, min_dist=0.1
UMAP, min_dist=0.5
UMAP/densMAP
densMAP
PHATE
Count
1000
900
800
700
600
500
400
300
200
100
0
random sampling
clustered sampling
Subsampling approach × PC space dimensionality

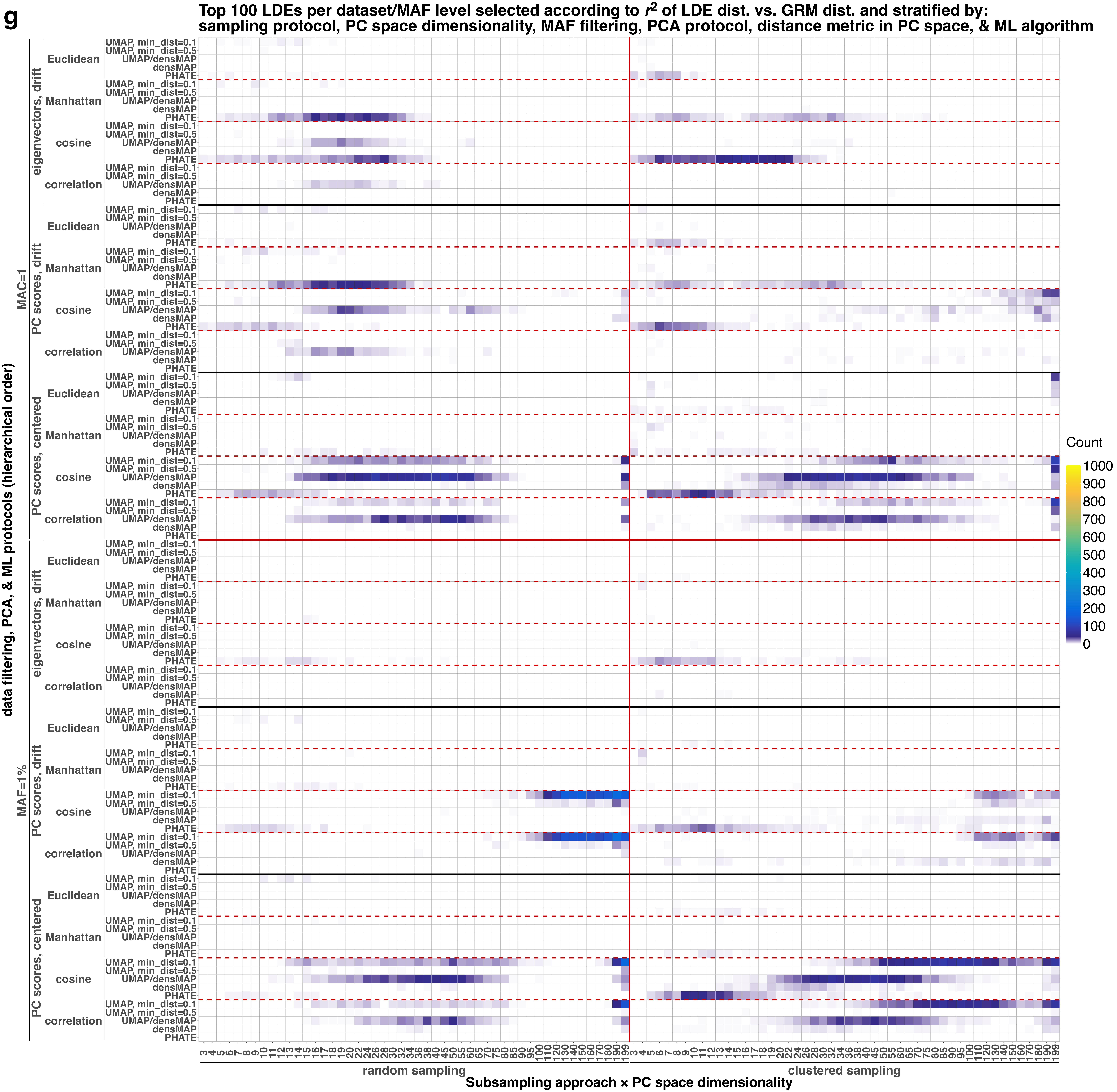
g
Top 100 LDEs per dataset/MAF level selected according to $r^2$ of LDE dist. vs. GRM dist. and stratified by:
sampling protocol, PC space dimensionality, MAF filtering, PCA protocol, distance metric in PC space, & ML algorithm
data filtering, PCA, & ML protocols (hierarchical order)
MAC=1
MAF=1%
eigenvectors, drift
PC scores, drift
PC scores, centered
Euclidean
Manhattan
cosine
correlation
UMAP, min_dist=0.1
UMAP, min_dist=0.5
UMAP/densMAP
densMAP
PHATE
Count
1000
900
800
700
600
500
400
300
200
100
0
3 4 5 6 7 8 9 10 11 12 13 14 15 16 17 18 19 20 22 24 26 28 30 32 34 36 38 40 45 50 55 60 65 70 75 80 85 90 95 100 110 120 130 140 150 160 170 180 190 199
random sampling
clustered sampling
Subsampling approach × PC space dimensionality

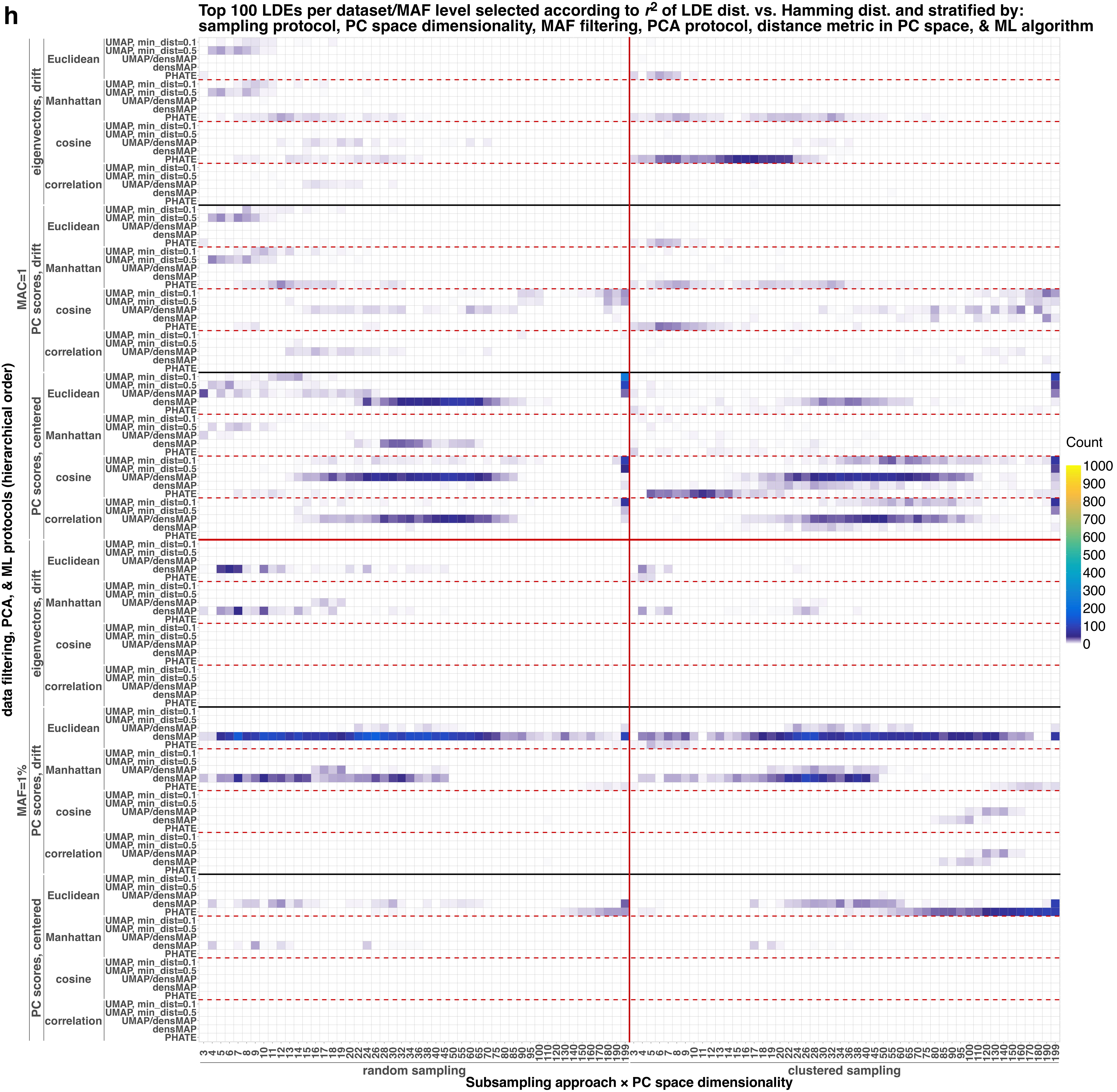
h
Top 100 LDEs per dataset/MAF level selected according to $r^2$ of LDE dist. vs. Hamming dist. and stratified by: sampling protocol, PC space dimensionality, MAF filtering, PCA protocol, distance metric in PC space, & ML algorithm
data filtering, PCA, & ML protocols (hierarchical order)
MAC=1
MAF=1%
eigenvectors, drift
PC scores, drift
PC scores, centered
Euclidean
Manhattan
cosine
correlation
UMAP, min_dist=0.1
UMAP, min_dist=0.5
UMAP/densMAP
densMAP
PHATE
Count
1000
900
800
700
600
500
400
300
200
100
0
3 4 5 6 7 8 9 10 11 12 13 14 15 16 17 18 19 20 22 24 26 28 30 32 34 36 38 40 45 50 55 60 65 70 75 80 85 90 95 100 110 120 130 140 150 160 170 180 190 199
random sampling
clustered sampling
Subsampling approach × PC space dimensionality

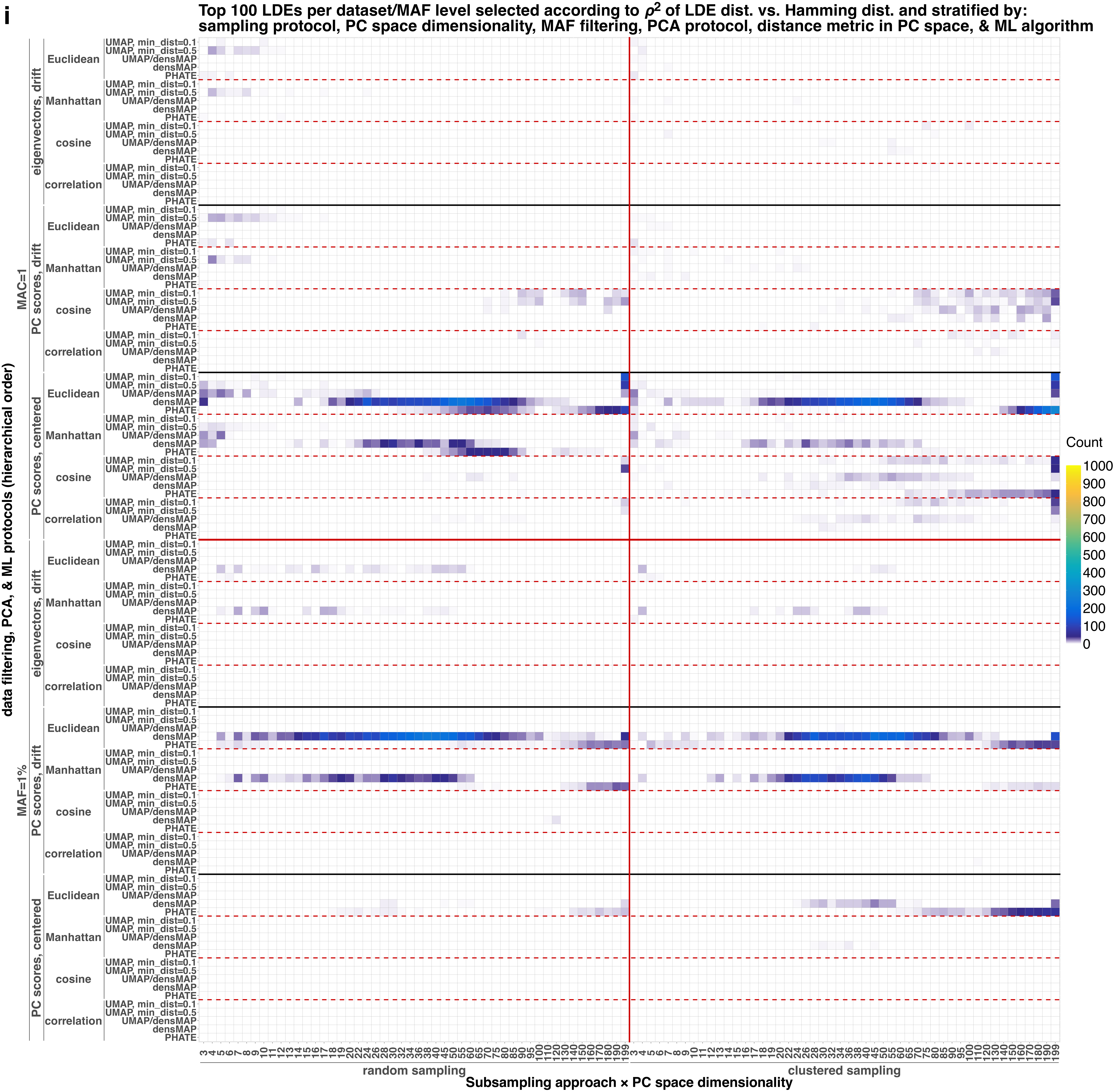
i
Top 100 LDEs per dataset/MAF level selected according to ρ2 of LDE dist. vs. Hamming dist. and stratified by:
sampling protocol, PC space dimensionality, MAF filtering, PCA protocol, distance metric in PC space, & ML algorithm
data filtering, PCA, & ML protocols (hierarchical order)
MAC=1
MAF=1%
eigenvectors, drift
PC scores, drift
PC scores, centered
Euclidean
Manhattan
cosine
correlation
UMAP, min_dist=0.1
UMAP, min_dist=0.5
UMAP/densMAP
densMAP
PHATE
Count
1000
900
800
700
600
500
400
300
200
100
0
random sampling
clustered sampling
Subsampling approach × PC space dimensionality

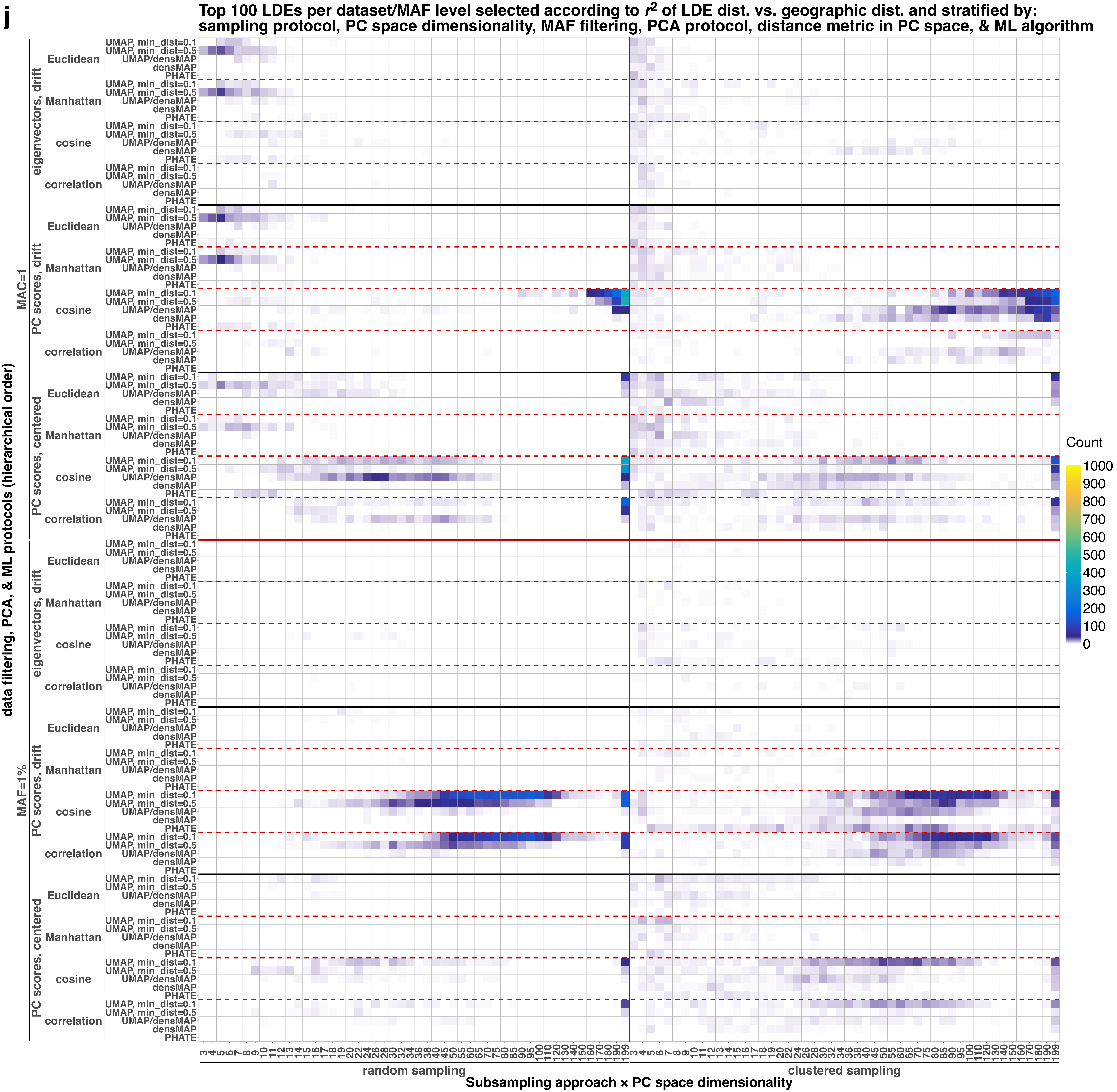
j
Top 100 LDEs per dataset/MAF level selected according to r2 of LDE dist. vs. geographic dist. and stratified by:
sampling protocol, PC space dimensionality, MAF filtering, PCA protocol, distance metric in PC space, & ML algorithm
data filtering, PCA, & ML protocols (hierarchical order)
MAC=1
MAF=1%
eigenvectors, drift
PC scores, drift
PC scores, centered
Euclidean
Manhattan
cosine
correlation
UMAP, min_dist=0.1
UMAP, min_dist=0.5
UMAP/densMAP
densMAP
PHATE
Count
1000
900
800
700
600
500
400
300
200
100
0
3 4 5 6 7 8 9 10 11 12 13 14 15 16 17 18 19 20 22 24 26 28 30 32 34 36 38 40 45 50 55 60 65 70 75 80 85 90 95 100 110 120 130 140 150 160 170 180 190 199
random sampling
clustered sampling
Subsampling approach × PC space dimensionality

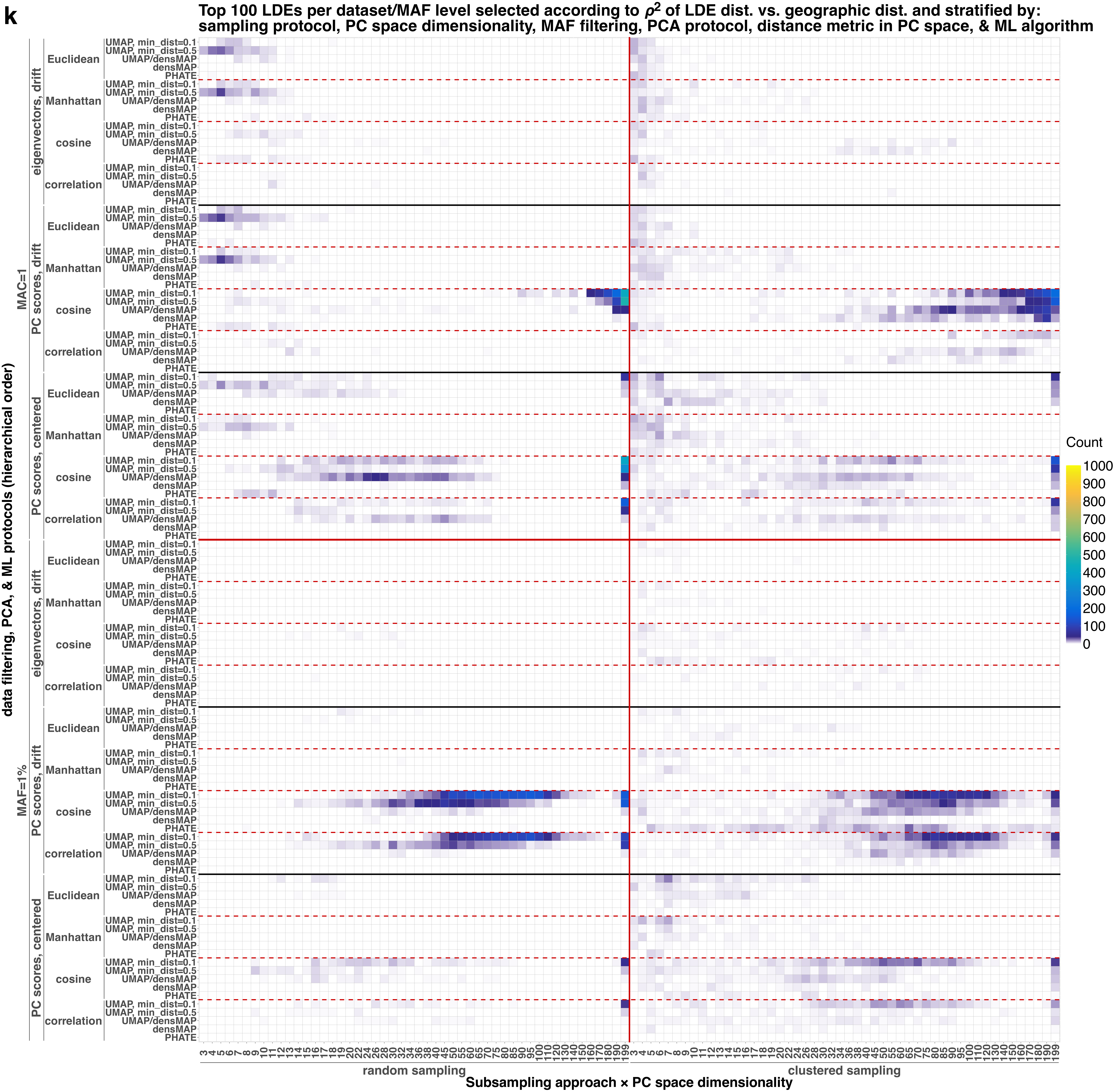
k
Top 100 LDEs per dataset/MAF level selected according to ρ2 of LDE dist. vs. geographic dist. and stratified by:
sampling protocol, PC space dimensionality, MAF filtering, PCA protocol, distance metric in PC space, & ML algorithm
data filtering, PCA, & ML protocols (hierarchical order)
MAC=1
MAF=1%
eigenvectors, drift
PC scores, drift
PC scores, centered
Euclidean
Manhattan
cosine
correlation
UMAP, min_dist=0.1
UMAP, min_dist=0.5
UMAP/densMAP
densMAP
PHATE
Count
1000
900
800
700
600
500
400
300
200
100
0
3 4 5 6 7 8 9 10 11 12 13 14 15 16 17 18 19 20 22 24 26 28 30 32 34 36 38 40 45 50 55 60 65 70 75 80 85 90 95 100 110 120 130 140 150 160 170 180 190 199
random sampling
clustered sampling
Subsampling approach × PC space dimensionality
