## Supplementary Figure 17 for "Sparse sampling and rare-variant depletion distort PCA visualizations of population structure: recovery with objective-guided manifold learning"

**(a) Correlation between $F_{ST}$ and Euclidean/Manhattan/cosine/correlation distances in PC spaces of varying dimensionality; each dot represents a given number of PCs used to calculate distances; *Cervus elaphus* case study: results are stratified by PCA protocol, distance metric in PC space, correlation coefficient, missing-data treatment, and SNP-filtering approach**

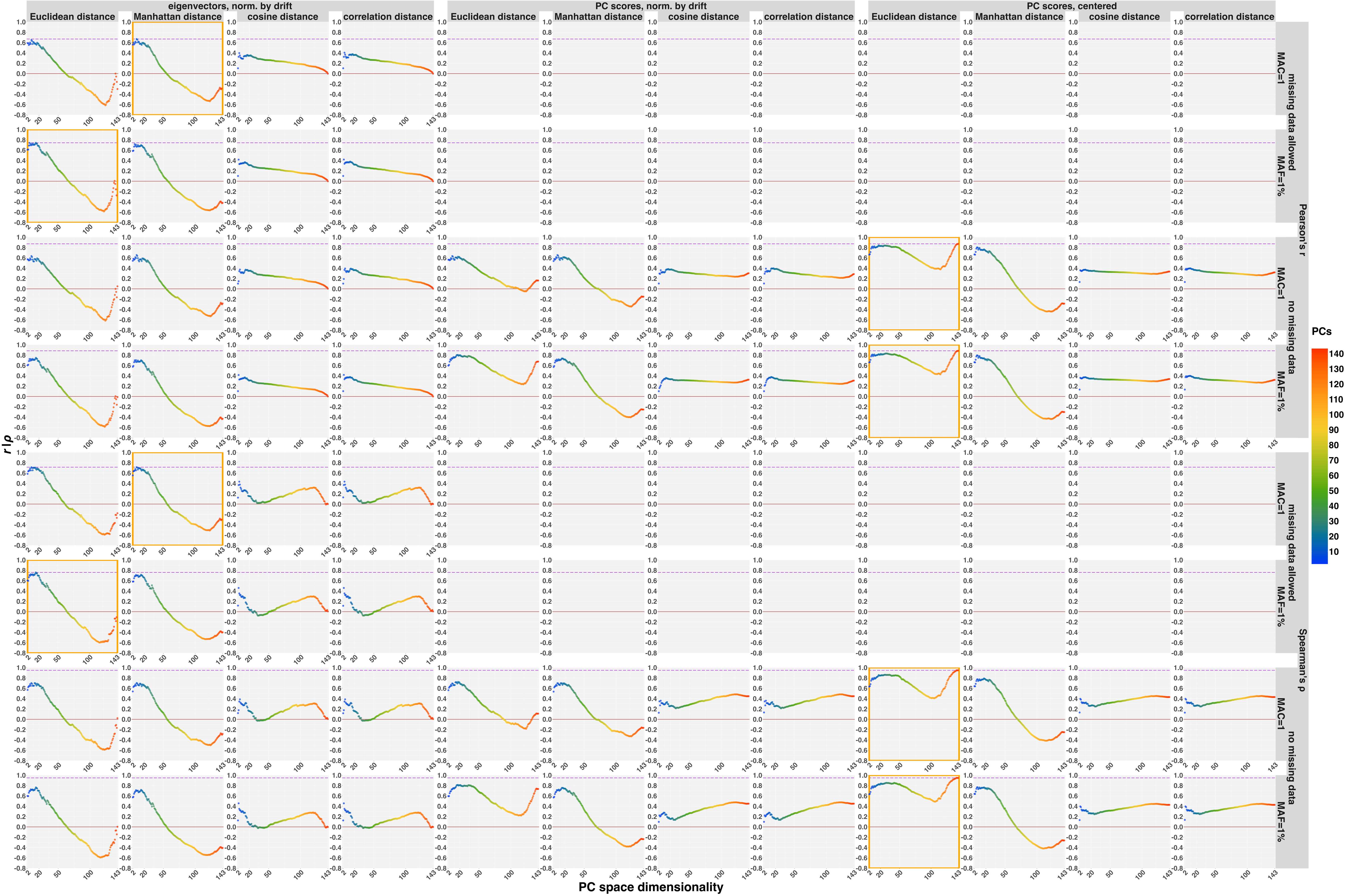

**(b) Correlation between Hamming genetic distances and Euclidean/Manhattan/cosine/correlation distances in PC spaces of varying dimensionality; each dot represents a given number of PCs used to calculate distances; *Cervus elaphus* case study: results are stratified by PCA protocol, distance metric in PC space, correlation coefficient, missing-data treatment, and SNP-filtering approach**

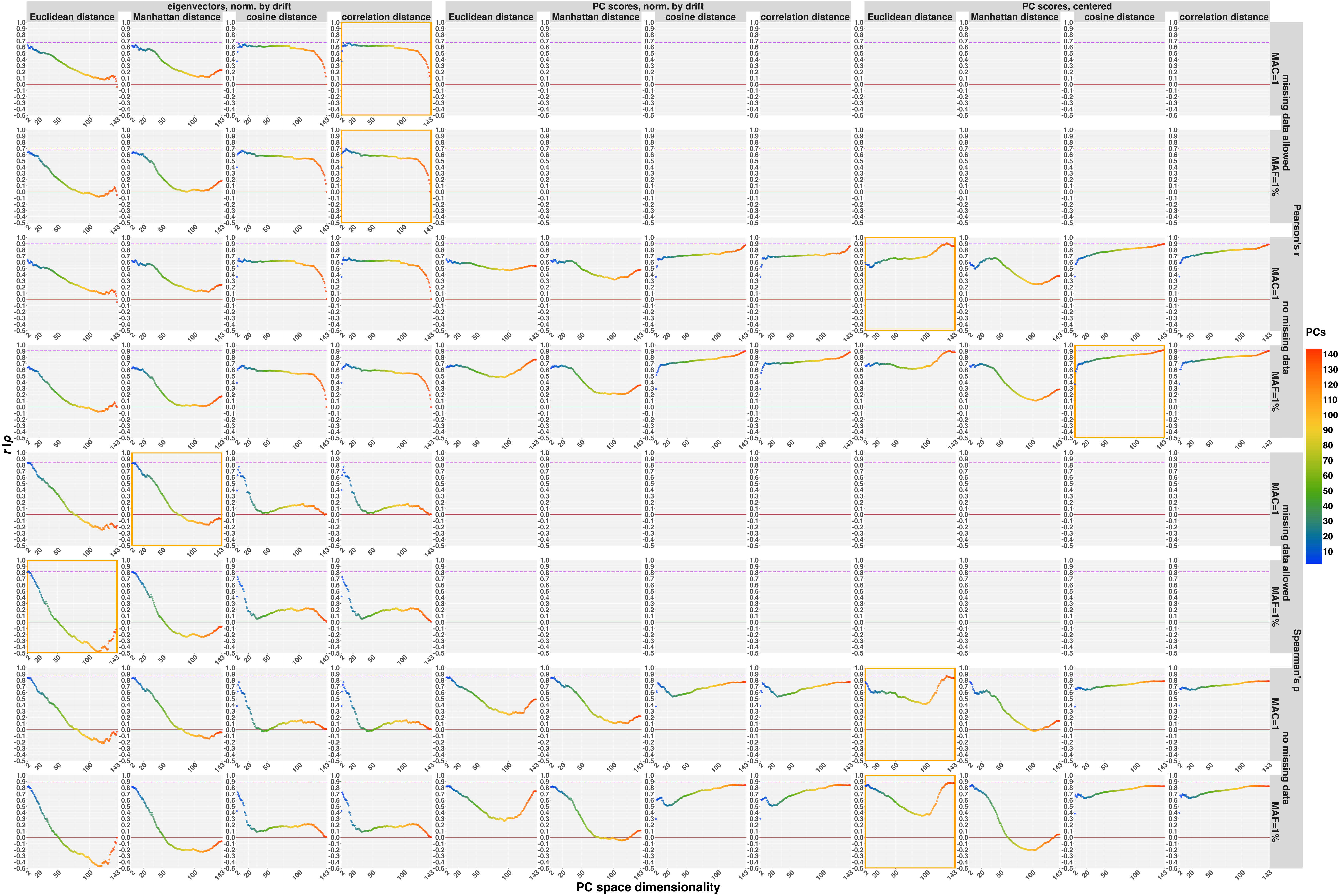

(c) Correlation between GRM-derived dissimilarities and Euclidean/Manhattan/cosine/correlation distances in PC spaces of varying dimensionality; each dot represents a given number of PCs used to calculate distances; *Cervus elaphus* case study: results are stratified by PCA protocol, distance metric in PC space, correlation coefficient, missing-data treatment, and SNP-filtering approach

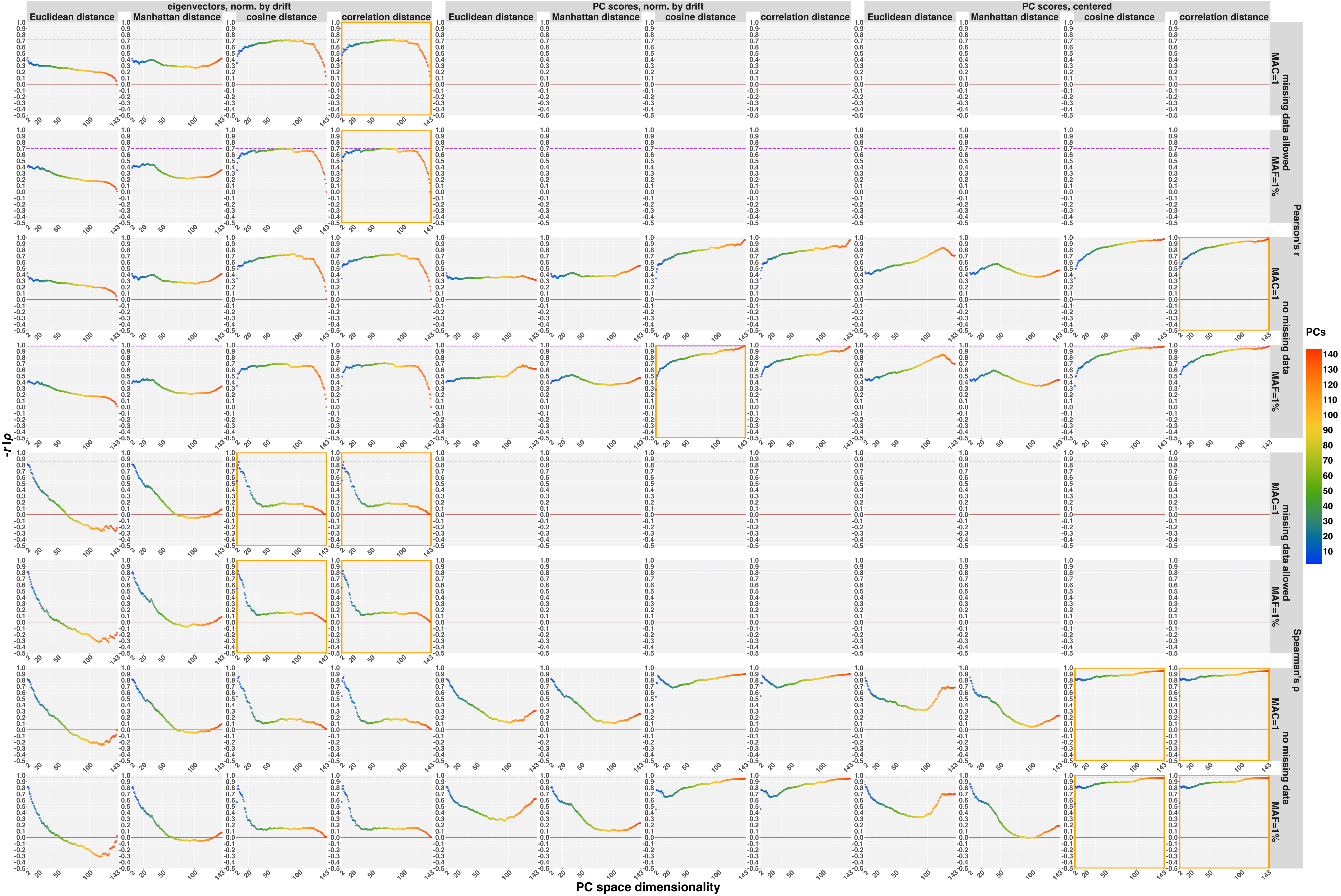

**(d) Correlation between outgroup $f_3$-statistics and Euclidean/Manhattan/cosine/correlation distances in PC spaces of varying dimensionality; each dot represents a given number of PCs used to calculate distances; *Cervus elaphus* case study: results are stratified by PCA protocol, distance metric in PC space, correlation coefficient, missing-data treatment, and SNP-filtering approach**

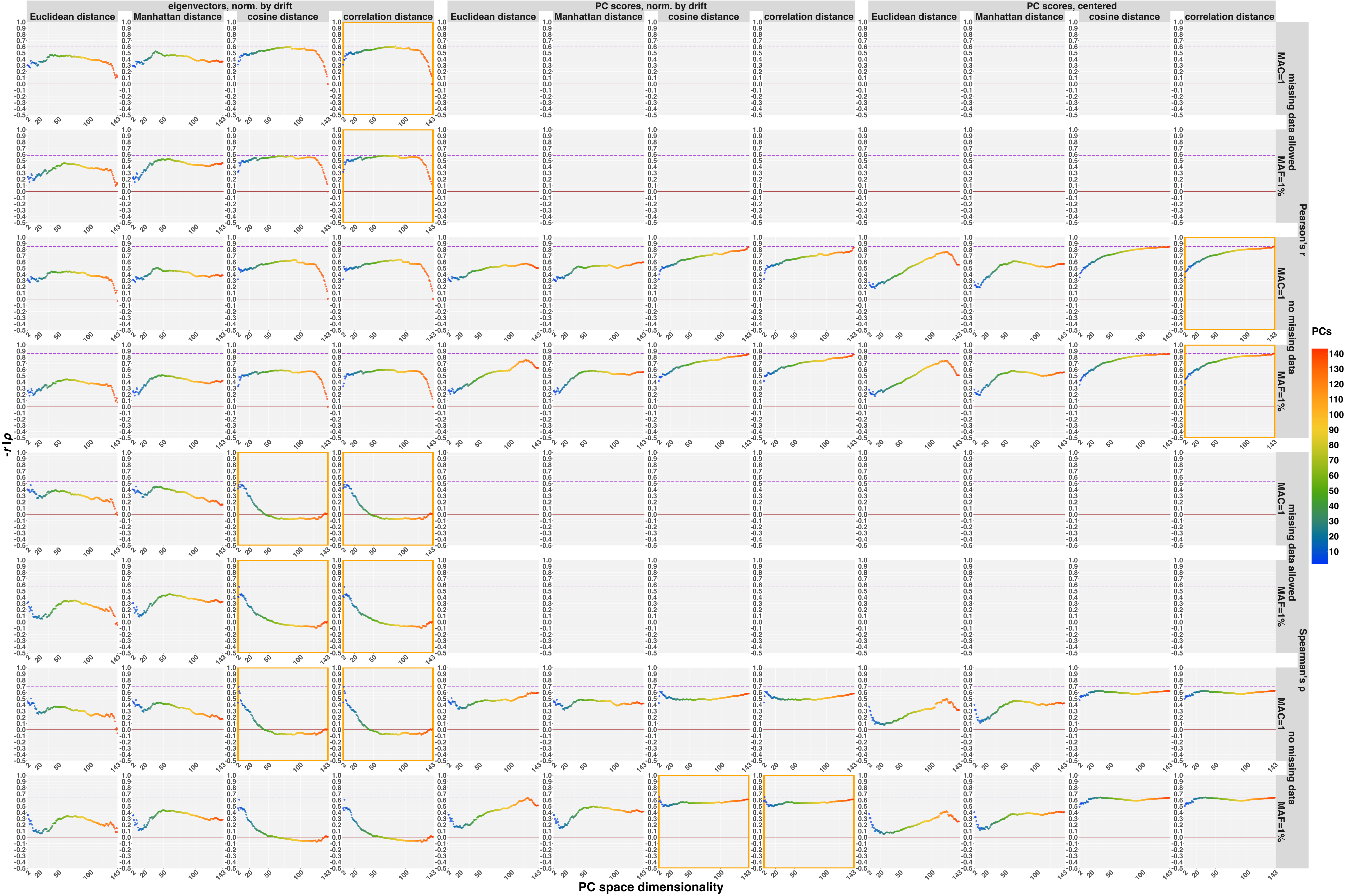

**(e) Correlation between great-circle geographic distances and Euclidean/Manhattan/cosine/correlation distances in PC spaces of varying dimensionality; each dot represents a given number of PCs used to calculate distances; *Cervus elaphus* case study: results are stratified by PCA protocol, distance metric in PC space, correlation coefficient, missing-data treatment, and SNP-filtering approach**

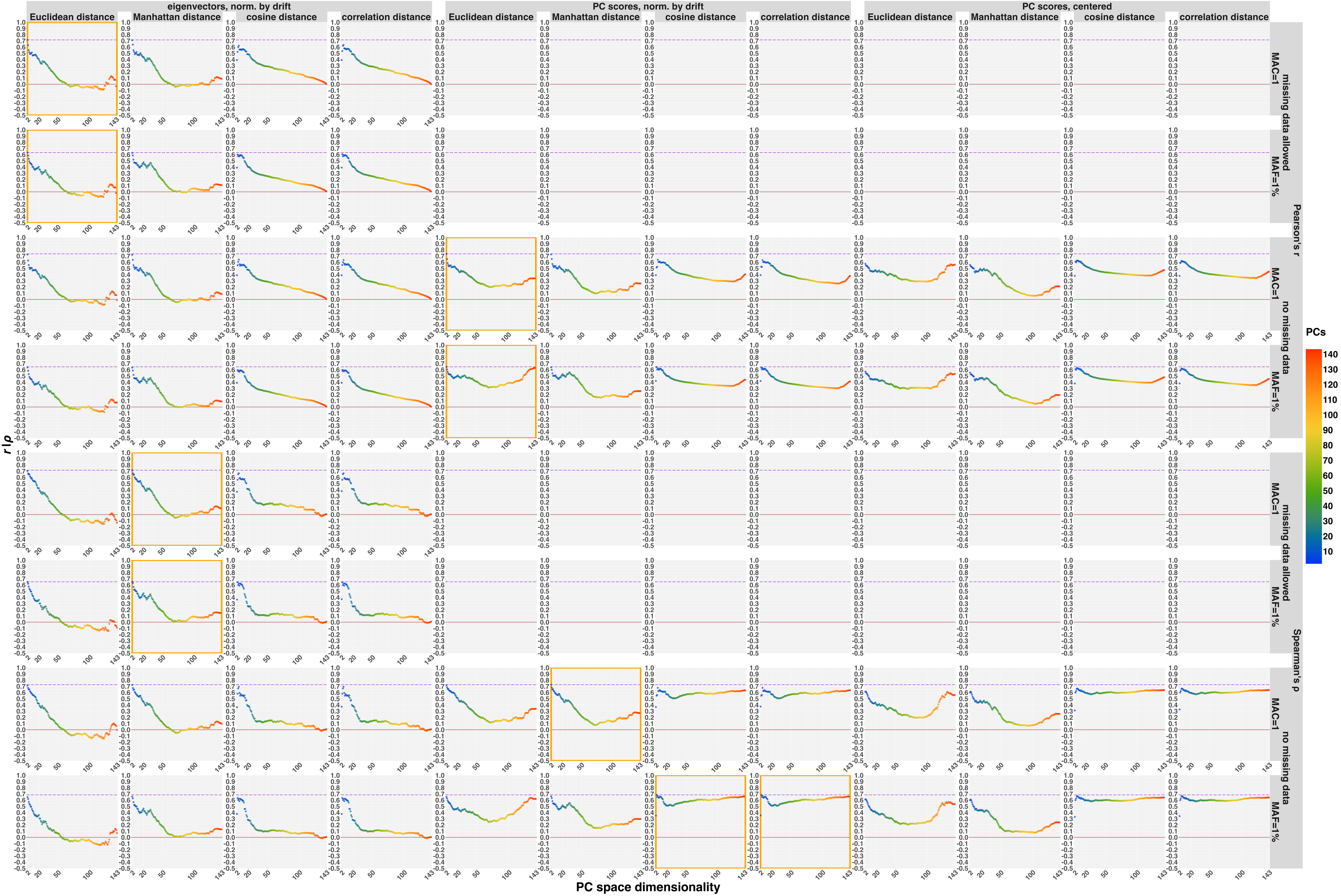

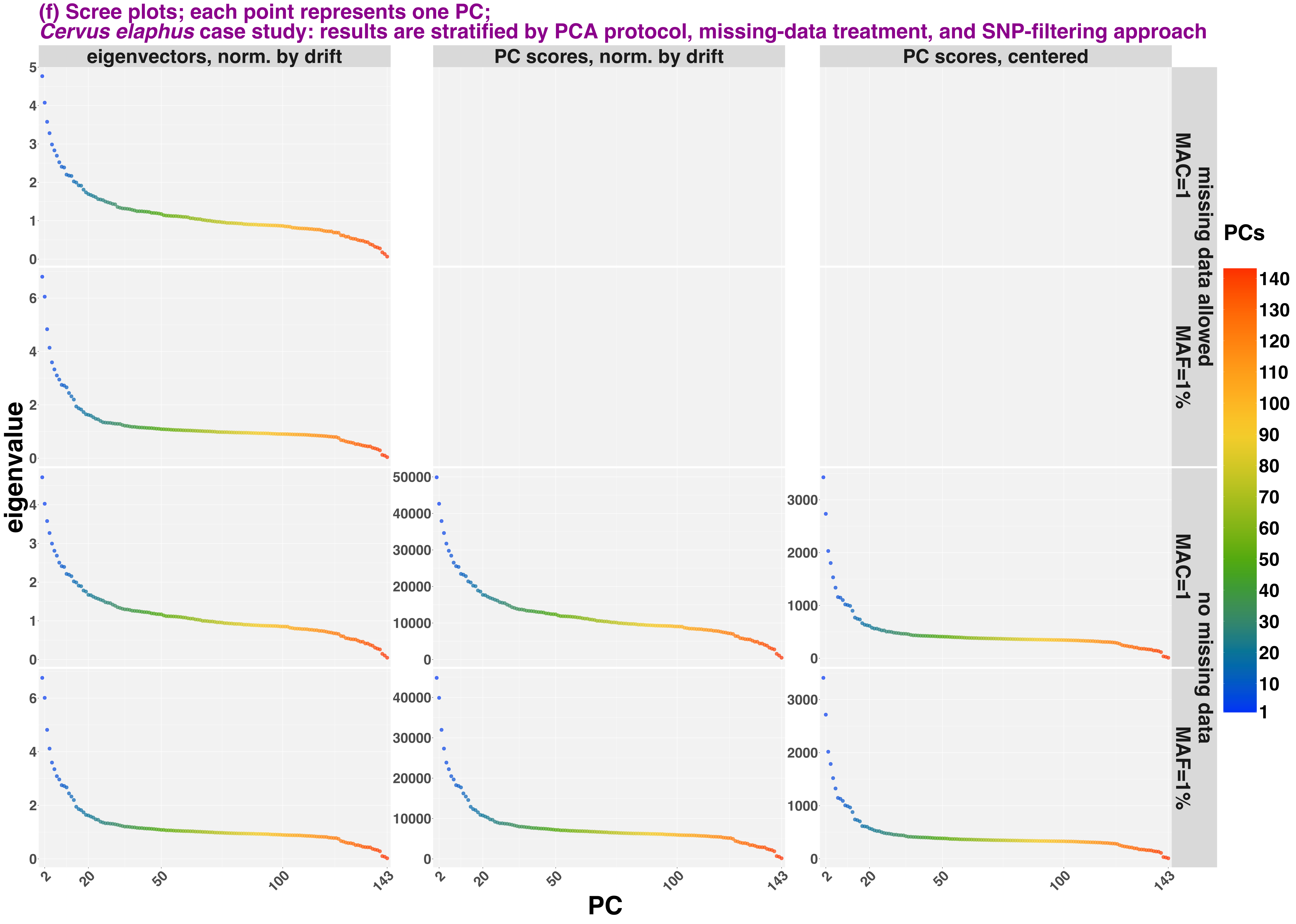

(f) Scree plots; each point represents one PC;
*Cervus elaphus* case study: results are stratified by PCA protocol, missing-data treatment, and SNP-filtering approach
eigenvectors, norm. by drift
PC scores, norm. by drift
PC scores, centered
missing data allowed
no missing data
MAC=1
MAF=1%
eigenvalue
PC
PCs
2
20
50
100
143
1
10
20
30
40
50
60
70
80
90
100
110
120
130
140
