## Supplementary Figure 19 for "Sparse sampling and rare-variant depletion distort PCA visualizations of population structure: recovery with objective-guided manifold learning"

**(a) Correlation between $F_{ST}$ and Euclidean/Manhattan/cosine/correlation distances in PC spaces of varying dimensionality; each dot represents a given number of PCs used to calculate distances; SEA case study: results are stratified by PCA protocol, distance metric in PC space, correlation coefficient, missing-data treatment, and SNP-filtering approach**

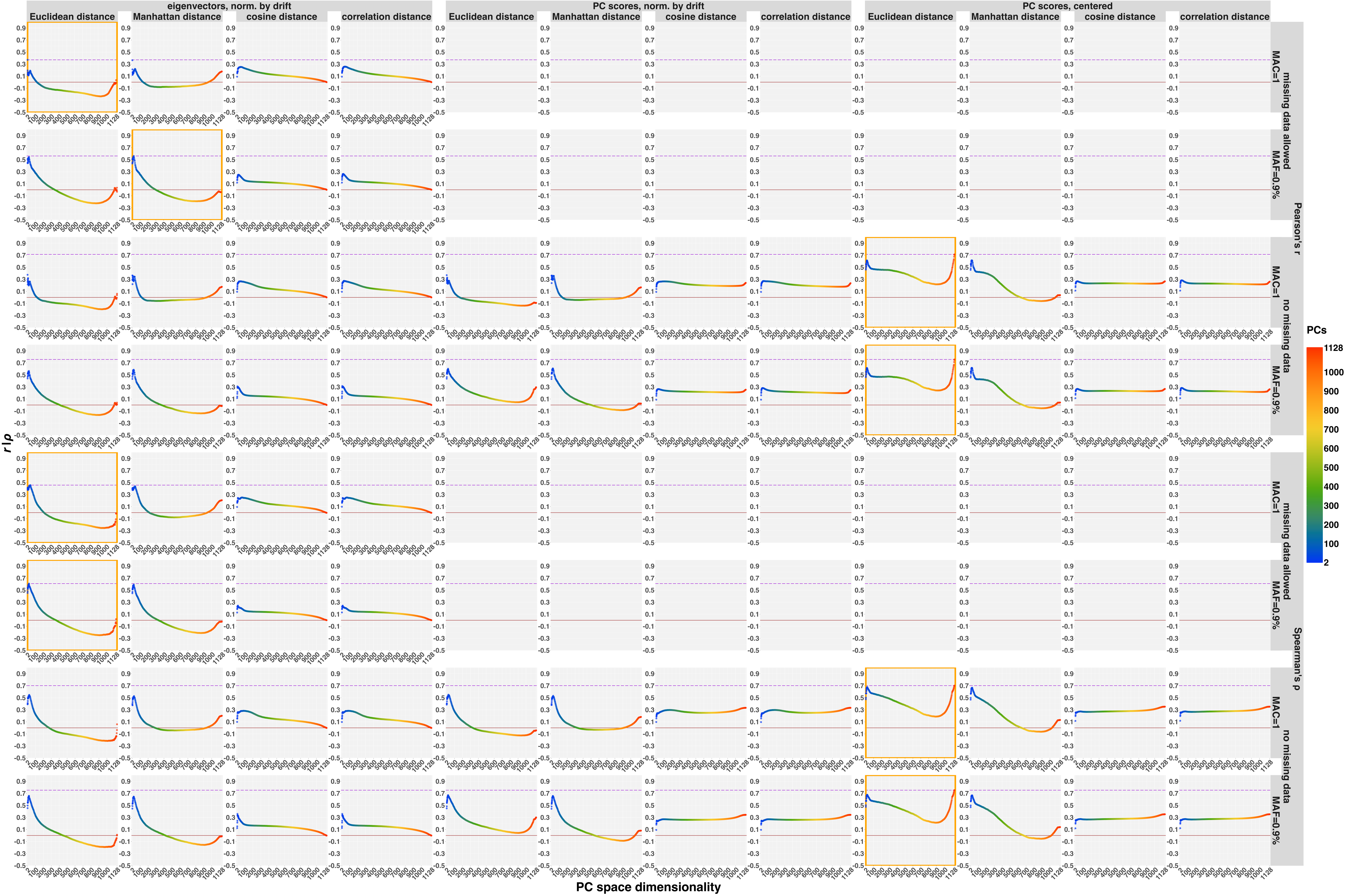

(b) Correlation between Hamming genetic distances and Euclidean/Manhattan/cosine/correlation distances in PC spaces of varying dimensionality; each dot represents a given number of PCs used to calculate distances; SEA case study: results are stratified by PCA protocol, distance metric in PC space, correlation coefficient, missing-data treatment, and SNP-filtering approach

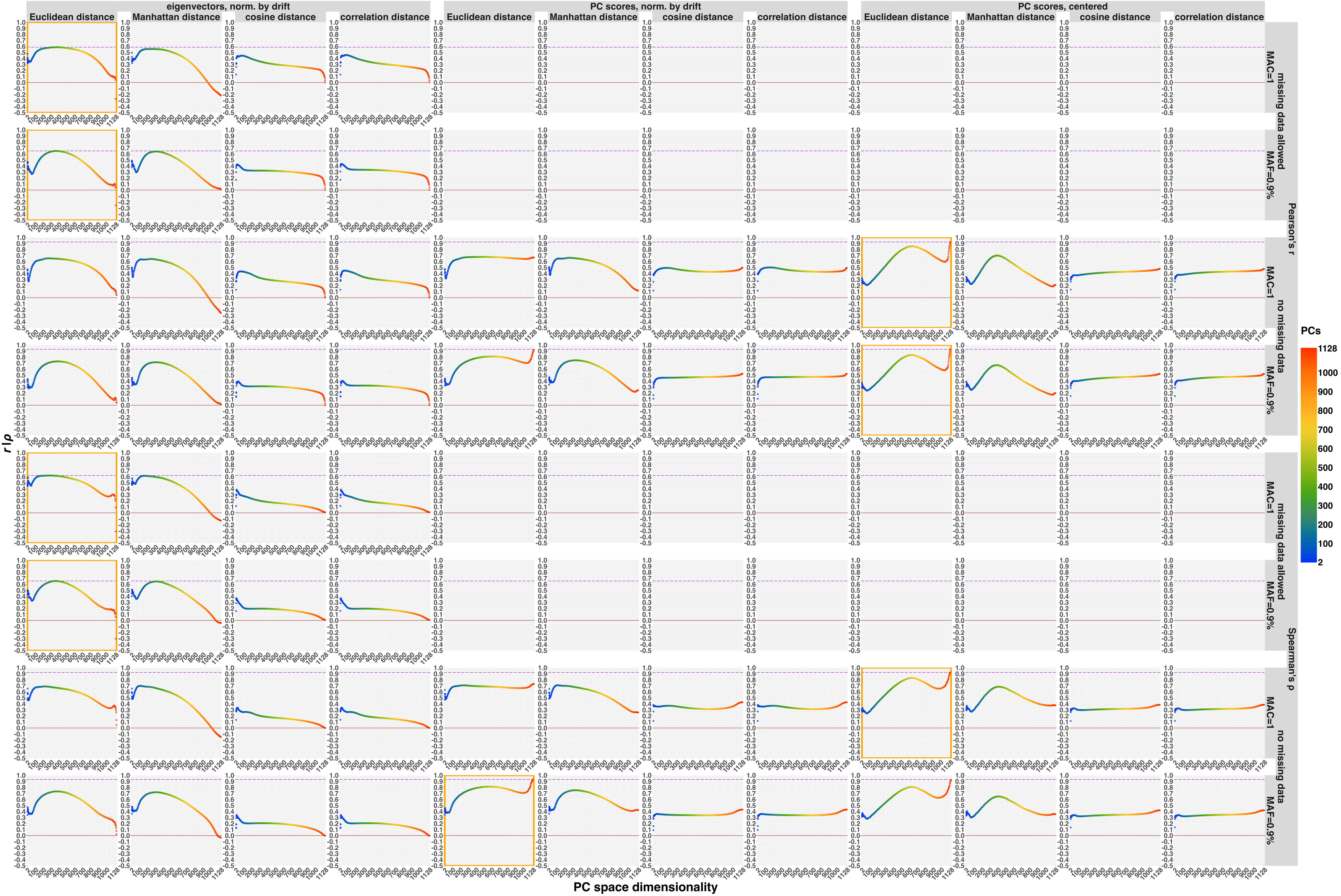

**(c) Correlation between GRM-derived dissimilarities and Euclidean/Manhattan/cosine/correlation distances in PC spaces of varying dimensionality; each dot represents a given number of PCs used to calculate distances; SEA case study: results are stratified by PCA protocol, distance metric in PC space, correlation coefficient, missing-data treatment, and SNP-filtering approach**

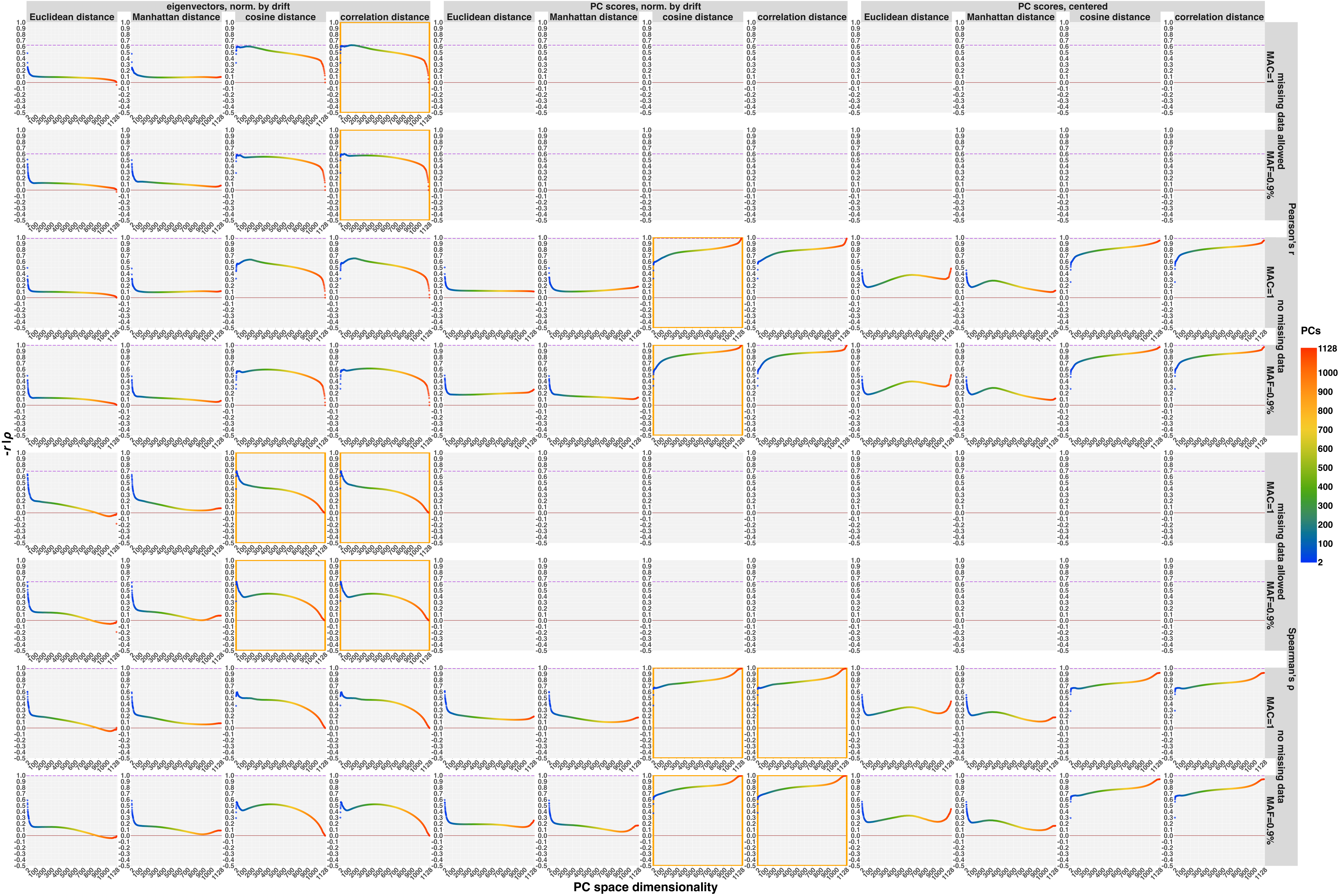

**(d) Correlation between outgroup $f_3$-statistics and Euclidean/Manhattan/cosine/correlation distances in PC spaces of varying dimensionality; each dot represents a given number of PCs used to calculate distances; SEA case study: results are stratified by PCA protocol, distance metric in PC space, correlation coefficient, missing-data treatment, and SNP-filtering approach**

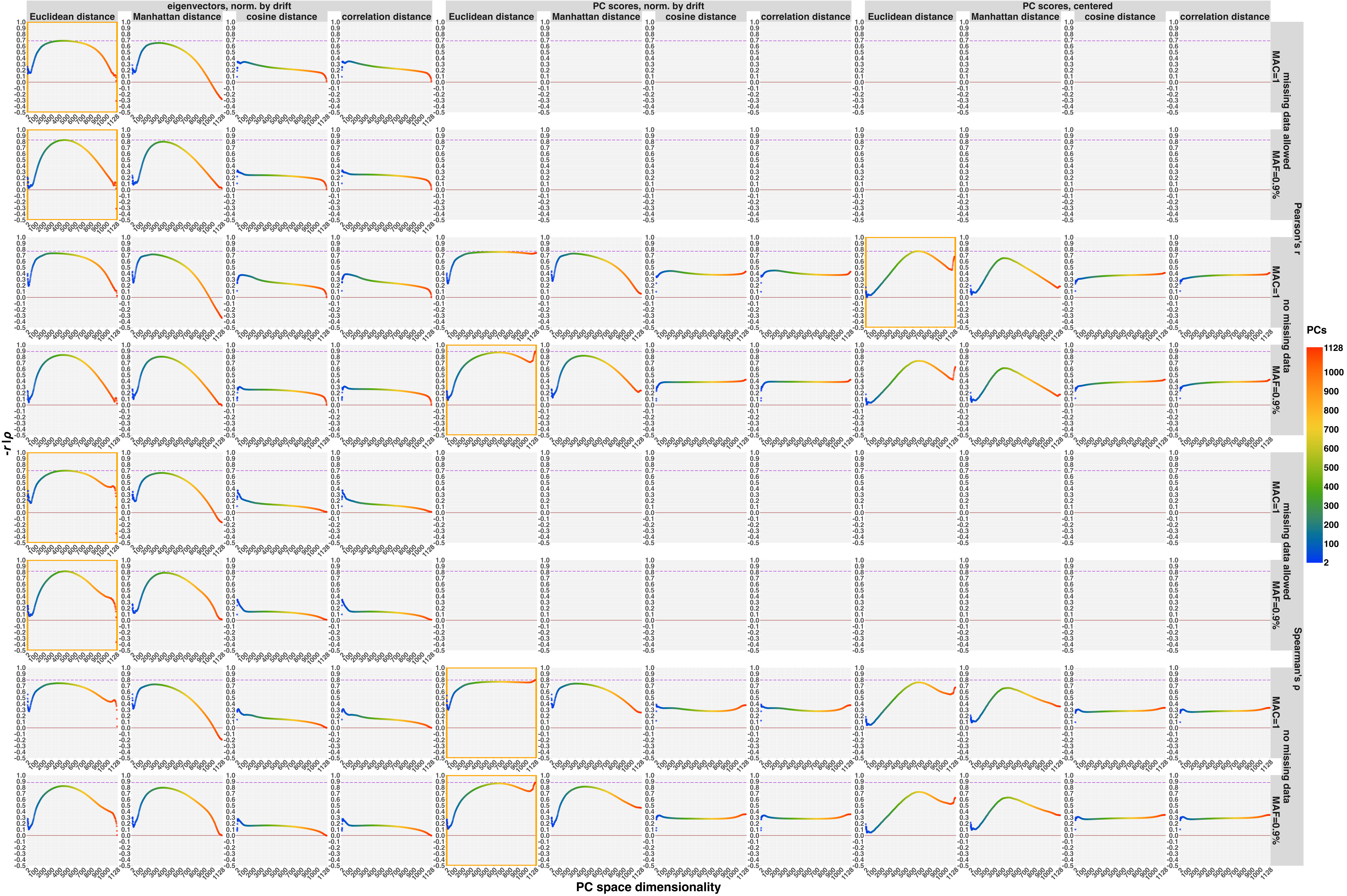

**(e) Correlation between great-circle geographic distances and Euclidean/Manhattan/cosine/correlation distances in PC spaces of varying dimensionality; each dot represents a given number of PCs used to calculate distances; SEA case study: results are stratified by PCA protocol, distance metric in PC space, correlation coefficient, missing-data treatment, and SNP-filtering approach**

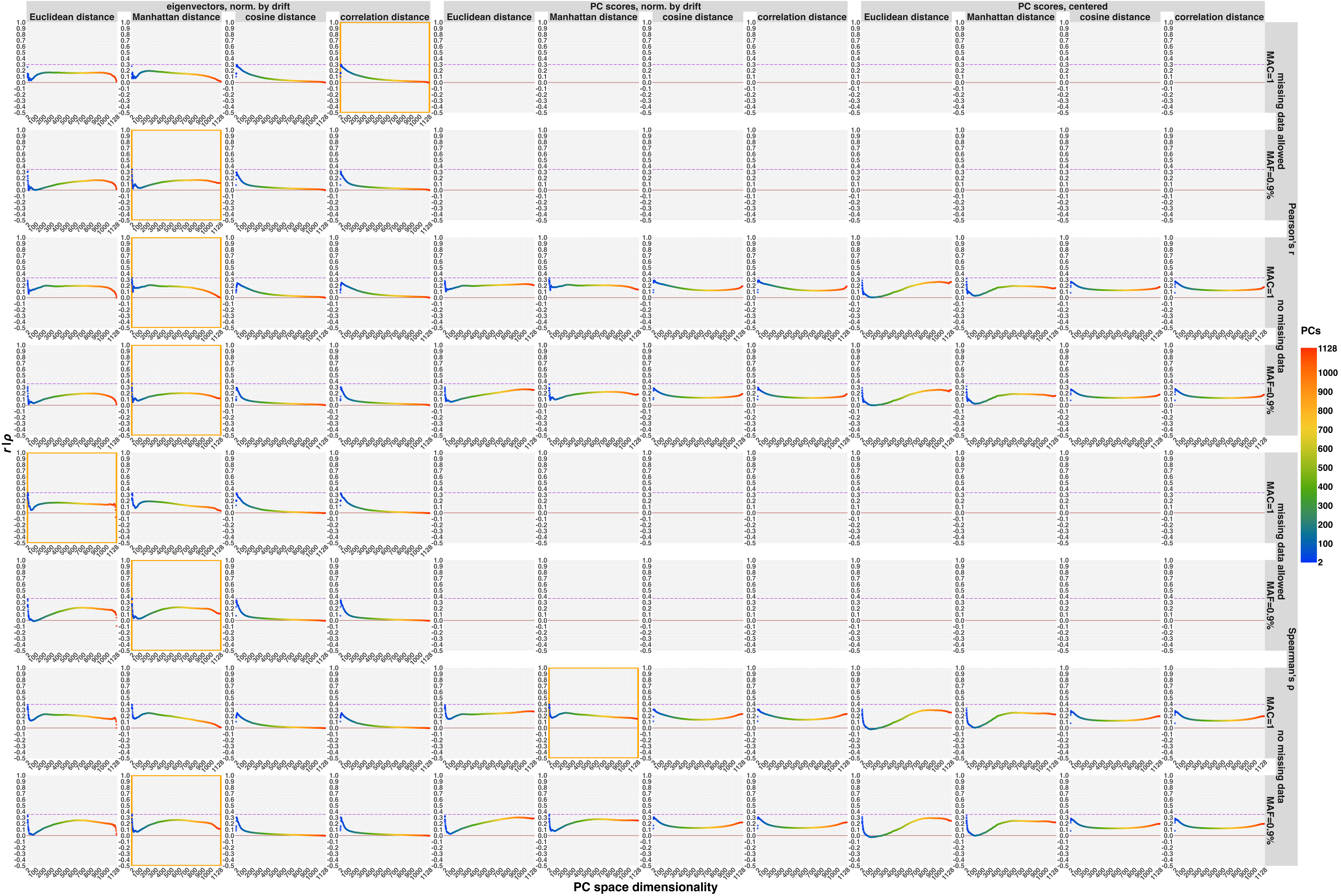

**(f) Scree plots; each point represents one PC;**
**SEA case study: results are stratified by PCA protocol, missing-data treatment, and SNP-filtering approach**

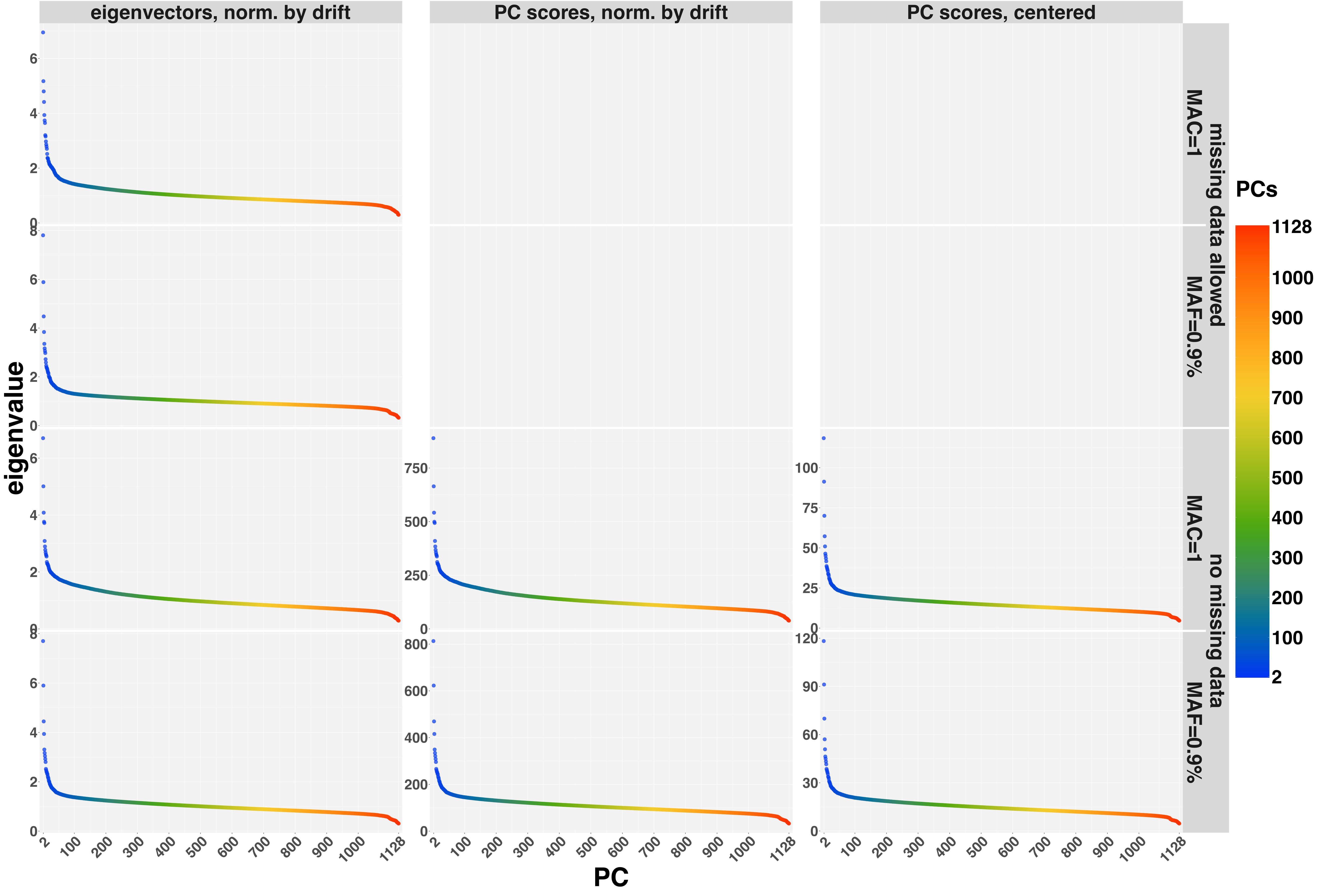

(g) Correlation between GRM-derived dissimilarities and Euclidean/Manhattan/cosine/correlation distances in PC spaces of varying dimensionality; each dot represents a given number of PCs used to calculate distances; SEA case study: results are stratified by PCA protocol, distance metric in PC space, correlation coefficient, missing-data treatment, and SNP-filtering approach

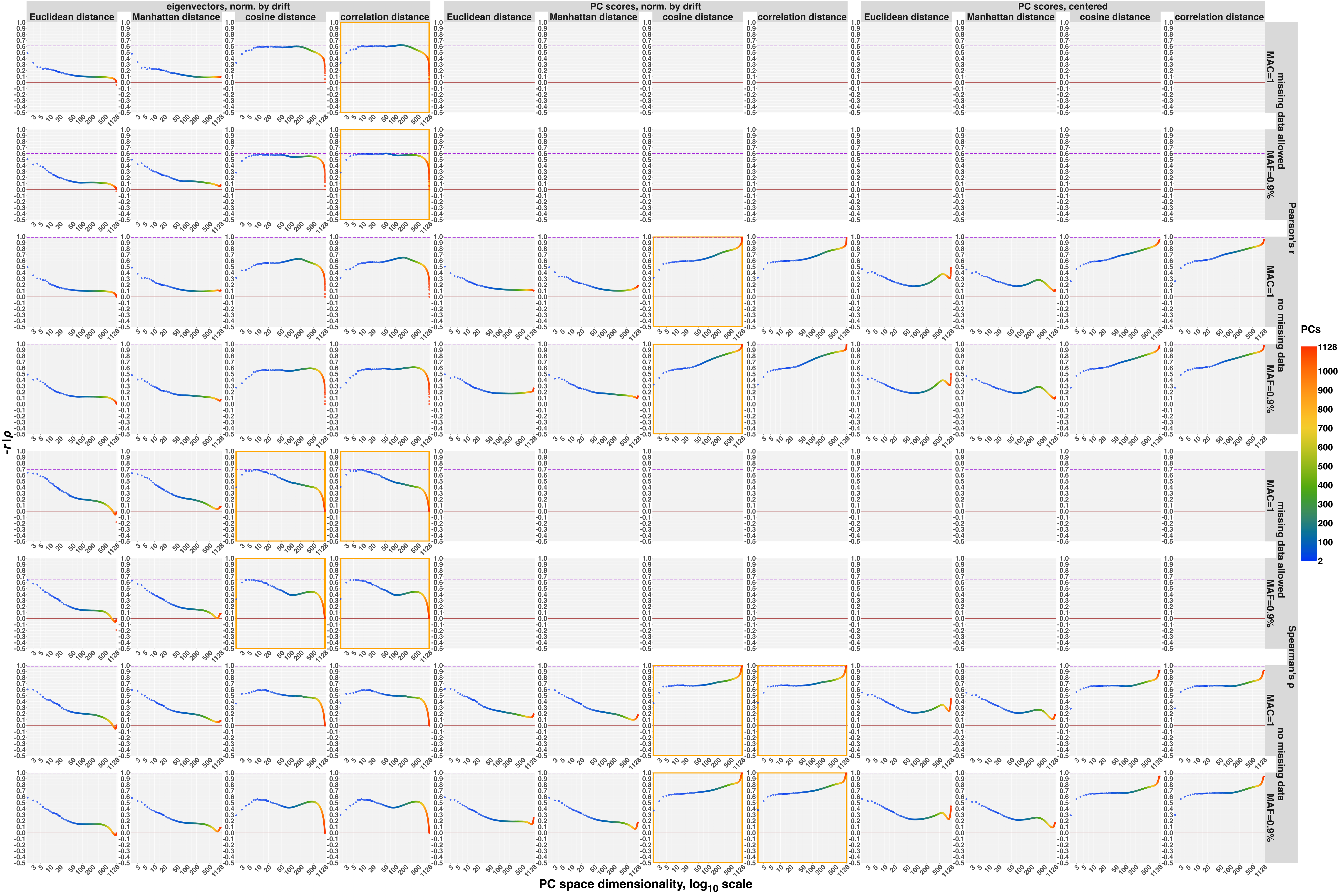
