## Supplementary Figure 21 for "Sparse sampling and rare-variant depletion distort PCA visualizations of population structure: recovery with objective-guided manifold learning"

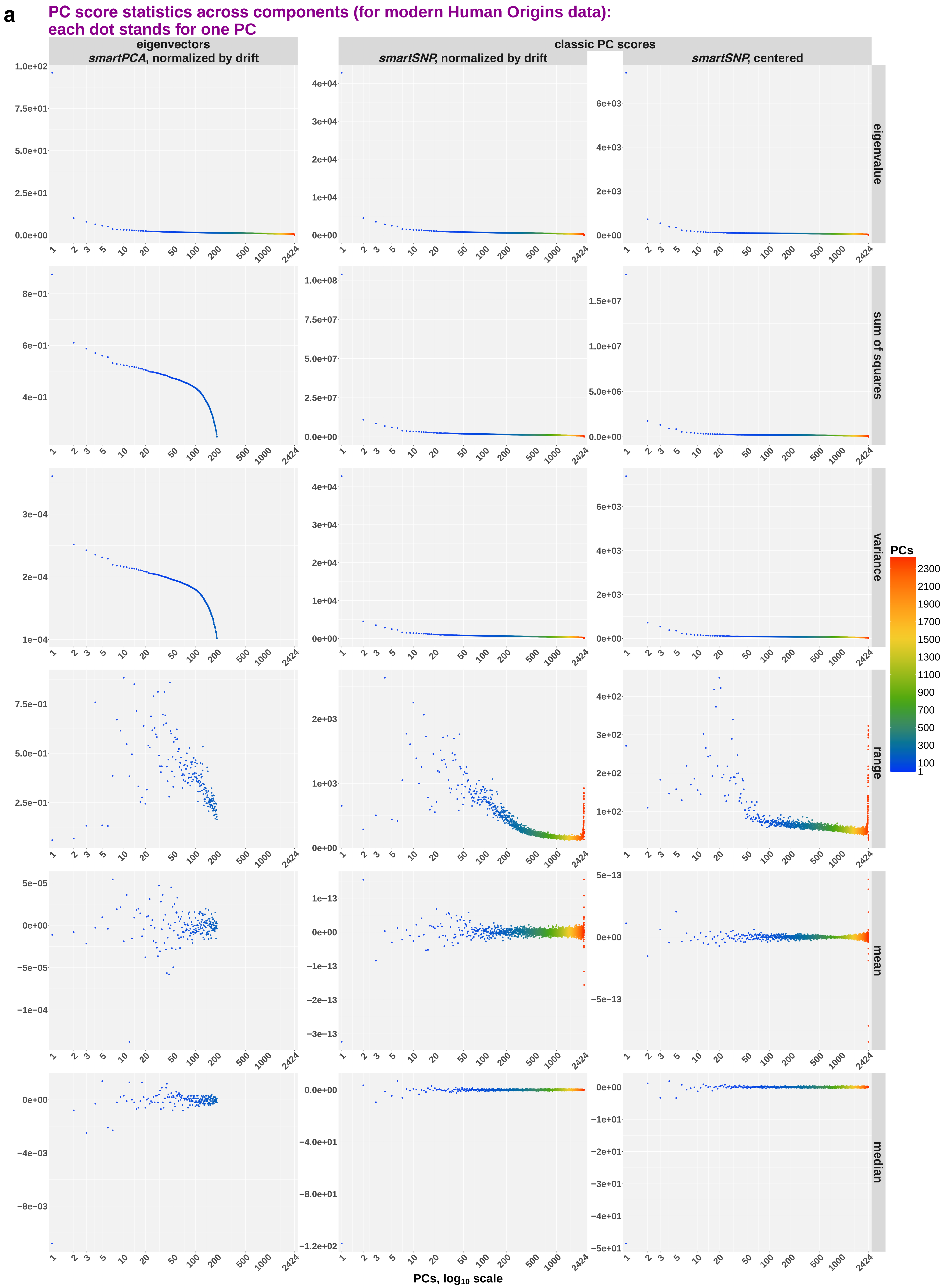
a
PC score statistics across components (for modern Human Origins data):
each dot stands for one PC
eigenvectors
smartPCA, normalized by drift
classic PC scores
smartSNP, normalized by drift
smartSNP, centered
eigenvalue
sum of squares
variance
range
mean
median
PCs
2300
2100
1900
1700
1500
1300
1100
900
700
500
300
100
1
PCs, log10 scale

**b** PC score statistics across components (for ancient individuals projected on modern Human Origins data): each dot stands for one PC

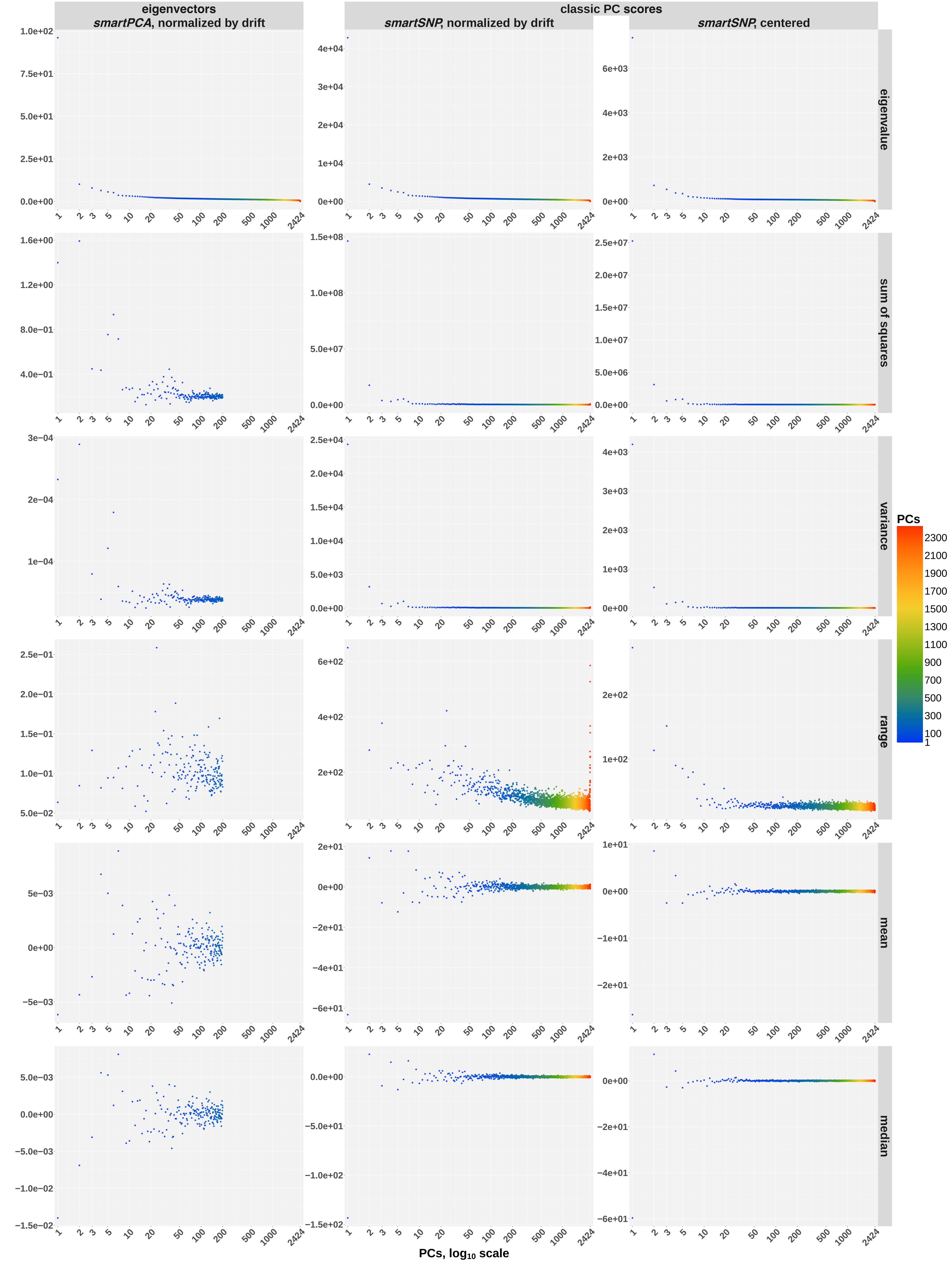
